# Granger-Causal Testing for Irregularly Sampled Time Series with Application to Nitrogen Signaling in Arabidopsis

**DOI:** 10.1101/2020.06.15.152819

**Authors:** Sachin Heerah, Roberto Molinari, Stéphane Guerrier, Amy Marshall-Colon

## Abstract

**Motivation:** Identification of system-wide causal relationships can contribute to our understanding of long-distance, intercellular signaling in biological organisms. Dynamic transcriptome analysis holds great potential to uncover coordinated biological processes between organs. However, many existing dynamic transcriptome studies are characterized by sparse and often unevenly spaced time points that make the identification of causal relationships across organs analytically challenging. Application of existing statistical models, designed for regular time series with abundant time points, to sparse data may fail to reveal biologically significant, causal relationships. With increasing research interest in biological time series data, there is a need for new statistical methods that are able to determine causality within and between time series data sets. Here, a statistical framework was developed to identify (Granger) causal gene-gene relationships of unevenly spaced, multivariate time series data from two different tissues of *Arabidopsis thaliana* in response to a nitrogen signal.

**Results:** This work delivers a statistical approach for modelling irregularly sampled bivariate signals which embeds functions from the domain of engineering that allow to adapt the model’s dependence structure to the specific sampling time. Using Maximum-Likelihood to estimate the parameters of this model for each bivariate time series, it is then possible to use bootstrap procedures for small samples (or asymptotics for large samples) in order to test for Granger-Causality. When applied to the *Arabidopsis thaliana* data, the proposed approach produced 3,078 significant interactions, in which 2,012 interactions have root causal genes and 1,066 interactions have shoot causal genes. Many of the predicted causal and target genes are known players in local and long-distance nitrogen signaling, including genes encoding transcription factors, hormones, and signaling peptides. Of the 1,007 total causal genes (either organ), 384 are either known or predicted mobile transcripts, suggesting that the identified causal genes may be directly involved in long-distance nitrogen signaling through intercellular interactions. The model predictions and subsequent network analysis identified nitrogen-responsive genes that can be further tested for their specific roles in long-distance nitrogen signaling.

**Availability:** The method was developed with the R statistical software and is made available thorugh the R package “irg” hosted on the GitHub repository https://github.com/SMAC-Group/irg. A sample data set is made available as an example to apply the method and the complete *Arabidopsis thaliana* data can be found at: https://www.ncbi.nlm.nih.gov/geo/query/acc.cgi?acc=GSE97500.

## 1 Introduction

Time series data are important for understanding the biological processes that are activated at different times and for inferring causality (Bar-Joseph *et al*., 2012). Many time series studies are designed to capture both dynamic and stationary phases in response to perturbations, which result in unevenly spaced time points, with dense sampling early and sparse sampling at later time points (Spellman *et al*., 1998; Colón *et al*., 2010; Krouk *et al*., 2010; Gargouri *et al*., 2015; Zhu *et al*., 2000). In biology, this is a commonly used sampling scheme to efficiently capture transient transcriptional and metabolic responses for example. However, the analysis of this irregular data is challenging, among others, since traditional time-lagged or cross-correlation analyses, designed for regularly spaced intervals, cannot be used. To date, it can be argued that no statistical approach has been able to comprehensively account for these unique features common to many biological time series (see e.g. Rehfeld *et al*., 2011).

Among the current approaches, methods designed for time-independent or regularly-spaced processes have been used to analyze unevenly-spaced time series data. For example, “static-based” clustering methods like hierarchical clustering and K-means have been used to organize and identify genes differentially expressed over developmental time in *Zea mays* (Chen *et al*., 2014), or in response to drought stress in *Arabidopsis thaliana* (Bechtold *et al*., 2016). However, clustering methods are not suitable to predict causal relationships between genes. Hence, other employed approaches include, among others, the transformation of irregularly sampled data into evenly spaced time series (Hamilton, 1994), in which the irregularity of the time interval can be approximated by forced regular intervals (Maller *et al*., 2008), or (resampling) strategies that estimate missing data points to fill in lags between observations (Broersen and Bos, 2006; Thiebaut and Roques, 2005; Remondini *et al*., 2005). Other methods directly address the irregular nature of the processes but do not consider the multivariate dependence and, consequently, the causal relation between signals (see e.g. Erdogan *et al*., 2005; Eyheramendy *et al*., 2018). These approaches have different drawbacks (Eckner, 2014) including: i) an inability to capture the variable nature of multivariate dynamic transcriptome experiments; and ii) resampling strategies often change the (Granger) causal relationship of the multivariate time series (Bahadori and Liu, 2012). All of these approximations can lead to incorrect correlations and predictions, and are unable to determine causal relationships within or between time series. Another commonly used approach in the analysis of (biological) time series is to perform a correlation analysis which however often does not account for non-stationary features of the data (Gargouri *et al*., 2015; Zhao *et al*., 2006). Indeed, the latter form of analysis can be highly misleading if, for example, the mean and/or variance of the series change over time which can often be the case for many experimental settings.

In response to the above limitations, this work puts forward a statistical approach that provides a general framework to determine Granger-Causality (Granger, 1969) for (short) irregularly sampled bivariate signals. We use this approach to describe causal gene-gene relationships from above- (shoot) and below- (root) ground organs of *Arabidopsis thaliana* in response to a nitrogen signal. Through identification and bioinformatic exploration of the detected causal relationships, we achieve a greater understanding of the underlying molecular and biochemical pathways involved in the nitrogen-signal response. This increase in understanding of nitrogen-responsive biochemical pathways in different plant tissues may help to predict emergent plant properties under nitrogen sufficiency and deficiency. Further testing of model-predicted causal relationships may uncover new molecules, pathways, and processes involved in the root-to-shoot-to-root nitrogen-signal relay, providing biological insight into complex, whole-plant nitrogen response.

## 2 Granger-Causal Analysis for Irregular Data

An irregularly spaced time series is a sequence of observations that are observed in time in a strictly increasing manner but where the spacing of observation times is not necessarily constant. More formally, let

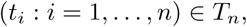

denote a strictly increasing time sequence of length *n* where:

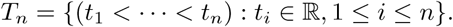

In addition, let (*X*_*i*_: *i* = 1, …, *n*) ∈ ℝ^*n*^ and (*Y*_*i*_ : *i* = 1, …, *n*) ℝ^*n*^ denote two sequences of real-valued random variables such that we can denote a bivariate irregularly spaced time series with *n* time points, as (*t*_*i*_, *X*_*i*_, *Y*_*i*_ : *i* = 1, …, *n*), where *t*_*i*_ denotes the time at which *X*_*i*_ and *Y*_*i*_ are to be observed. In the context of this paper, we focus on those random sequences that are observed at the same points in time (i.e. the sequences (*t*_*i*_ : *i* = 1, …, *n*) correspond for both series). However, this condition can also be relaxed as a result of the research developed in this work.

As highlighted previously, the literature on irregularly spaced time series is not abundant and methods available to practitioners for estimation and inference in these cases are lacking as well. In this section we therefore put forward a pertinent statistical model that we will denote as *F* = {*F*_***θ***_ : ***θ*** ∈ **Θ** ⊂ ℝ^*p*^}, with ***θ*** being the vector containing the parameters of this model. The latter model needs to deal with irregularly spaced bivariate time series and should allow to test for Granger causal links between the series themselves. In order to achieve this goal, we firstly define 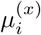 and 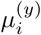 as the expected values of 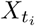 and 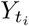, respectively. These quantities represent, in the case of dynamic transcriptome and metabolome data, the natural (deterministic) variation in gene expressions due, for example, to changes in environmental conditions or natural cycles. If we were considering evenly spaced observations, it would appear reasonable to consider the class of AutoRegressive Moving Average (ARMA) models to describe the variations of 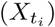 around its mean (see e.g. Box *et al*., 2015, for details). A commonly used model within this class, especially when dealing with small sample sizes, is the first-order autoregressive model, i.e. an AR(1), which is defined as

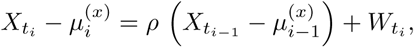

where *ρ* represents the parameter which explains the dependence between consecutive observations and 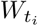 is an independent sequence of random variables with a certain (finite) variance *σ*^2^. This model allows to approximate many covariance structures delivering a behavior that is often reasonable for many biological and natural phenomena. In order to determine whether another time series (signal) has an impact on the time series under consideration, the above model can be extended to the following:

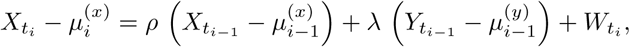

where *λ* therefore represents the impact that the time series 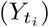 has on the time series 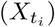. In general terms, we can say that 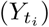 *Granger-causes* 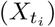 if the latter model explains the behavior of 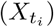 better than the previously defined AR(1) model that only depends on the sequence 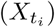. The concept of Granger-Causality was introduced in Granger (1969) and the goal of the biological study considered in this work would therefore be to perform a statistical test to confirm the stronger explanatory power of the second model over the first.

However, these models are not well-adapted to irregularly spaced time series that are the focus of this work. For example the parameter *ρ*, that measures the relation between consecutive observations, remains constant regardless of the distance in time between 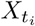 and 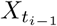 (as well as the parameters *λ* and *σ*^2^). For this reason, the next sections put forward a new framework for these settings.

### 2.1 The Proposed Model

The first step required to address the problem of modelling irregularly spaced time series consists in integrating the distance in time between observations within the model specification. Assuming an appropriate technique is used to estimate 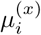 (e.g. splines or other semi- or non-parametric approaches), we denote the centred observations as 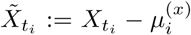 and the distance in time as 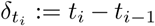, with 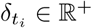 by definition. Based on this, the AR(1) model for irregularly spaced data can be represented as follows:

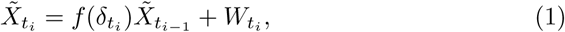

where *f* (·) is a deterministic function, possibly known up to some parameter values, that plays the same role as the constant *ρ* but takes into account the distance between observations. For different reasons, among which estimation feasibility, the independent sequence 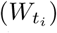 is usually considered as being Gaussian (although other distributions can be considered). Without loss of generality, we will make this assumption and therefore state that 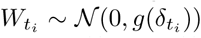 with *g*(·) being another deterministic function. Both the functions *f* (·) and *g*(·) need to respect certain properties which will be discussed further on. The model defined in (1) could be extended in several ways, for example, by considering a dependence between 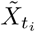and 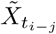 with *j >* 1 or between 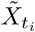 and 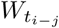 as in general ARMA models (as well as considering non-Gaussian distributions for 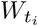 as mentioned earlier). However, given the small sample sizes usually encountered in dynamic transcriptome and metabolome studies, it is rather unlikely that more complex models can be appropriately estimated and the model in (1) is a very reasonable approximation for more general dependence structures.

Considering the extension of AR(1) processes to irregularly spaced settings, we can consider the same extension when modelling the joint behavior of two time series. For this purpose we define the following bivariate model, which is a natural extension of a vector AR(1) model for irregularly spaced data:

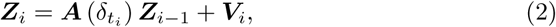

where

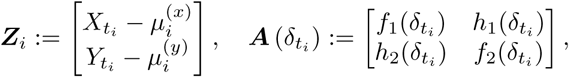

and where *h*(·) is another deterministic function (which may depend on unknown parameters). In addition, we have 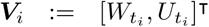 with 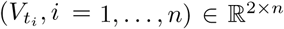 denoting a bivariate independent sequence with distribution 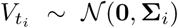, with **0** being a two-dimensional zero vector and

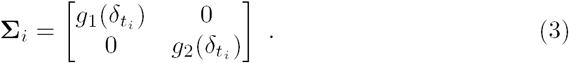

It can be observed how the matrix 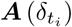 plays the main role in describing the dependence “within” and “between” the two time series. Indeed, on one hand the functions 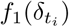 and 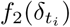 determine to what extent the time series depend on themselves to describe the behavior of their future observations while the functions 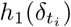 and 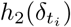, on the other hand, determine the degree of dependence between the two time series. Also within this setting it is possible to recognize the idea of Granger causality where one is interested in assessing whether past values of a certain time series can significantly increase the explanation of the behavior of another time series. In general, this assessment is based on statistical tests which are typically related to characteristics of the matrix 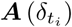. In fact, if this matrix is diagonal, this implies that the two time series are independent from each other (under the Gaussian assumption) while if it is full this entails that the two time series are also inter-dependent. Moreover, if the matrix is upper or lower triangular, this would imply that only one of the series depends on itself *and* on the other series (the latter therefore only depending on itself).

Considering the above modelling framework, there is a need to estimate the unknown parameters in the model and test whether the estimated models appear to explain the data sufficiently well to draw reliable conclusions. Firstly, to estimate these kind of models we propose a likelihood approach based on the assumption of a jointly normal distribution of the observations which, for the bivariate series, gives the following conditional distribution:

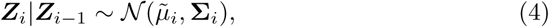

where **Σ**_*i*_ is defined in (3), and

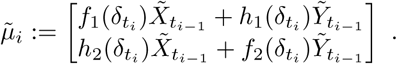

If we denote the unconditional distribution of ***Z***_*i*_ as *l*(***Z***_*i*_), then the likelihood function is given by

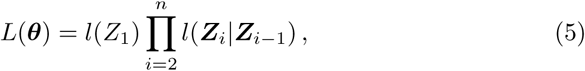

where, using (4), we have

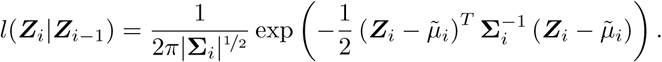

Applying the log(·) function to *L*(***θ***) and fixing *l*(***Z***_1_) as constant (neglecting constant terms) we obtain the following estimating equation which defines the Maximum Likelihood Estimator (MLE):

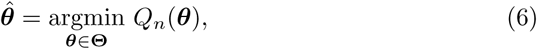

where

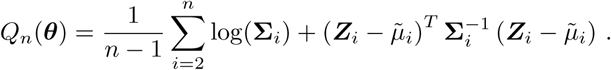

Under a set of conditions (see Section 1 in the supplementary material), the estimator defined in (6) has appropriate statistical properties. Among these conditions there are constraints on the deterministic functions that characterize the dependence structure of the model defined in (2). For this reason, we define these functions accordingly taking from the domain of (navigation) engineering (see e.g. Titterton *et al*., 2004). In the latter field, a model that is often used is the discrete-time first-order Gauss-Markov model that can be defined as:

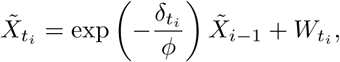

where *ϕ ∈* ℝ^+^ is a parameter that determines the “range” of dependence in the data and

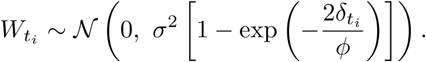

Having been mainly proposed to deal with time series measured at different frequencies, the idea behind this model is very close to the structure of an exponential model for spatial data (see e.g. Ripley, 2005). Indeed, the latter explains the dependence in space through an exponential structure and roughly corresponds to the above-mentioned Gauss-Markov process when considering 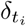 as a measure of Euclidean distance. The above model therefore gives an explicit form to the functions *f*_*·*_(·) and *g*_*·*_(·) mentioned earlier but of course other explicit forms can be envisaged.

While the above defined functions characterize the dependence of a time series on itself, it is still necessary to give an adequate form to the function *h*_*·*_(·) that describes the behaviour of a signal based on another. Given the short time series available, we decide to impose a reasonable structure to this behaviour which allows the dependence of a signal on the other signal to grow exponentially over time (reaching its maximum) and then decay exponentially. Indeed, while we consider the impact of past values of a time series on its future values as a function only of their distance in time, we postulate that the impact of another time series is not constant but increases and then decreases as a function of the distance in time over the chosen experimental time-frame. This behaviour can be justified from a biological point of view since genes have been shown to influence the expression of other genes in a “hit and run” manner (Doidy *et al*., 2016). The causal gene physically interacts with the target gene then dissociates, but the transient target gene’s expression continues to be affected after the dissociation. For this reason we propose to use the following function:

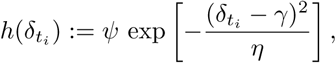

where *ψ* ∈ (− 1, 1) is a parameter that describes the “intensity” and “direction” of the dependence of a time series on the other while *γ* ∈ ℝ^+^ denotes the distance in time at which the dependence of a time series on another is maximal. Finally, *η ∈* ℝ^+^ plays a similar role to *ϕ* in the previously defined function *h*_*·*_(*·*).

As stated earlier, other explicit (more complex) forms can be defined for these functions. However other forms would probably require more parameters to characterize them and would be complicated (if not impossible) to estimate in practice given the small sample sizes collected in many experimental settings such as the one considered in this work. Hence, in order to respond to the need to balance model complexity with practical feasibility, we will consider the above functions to understand the relationship between different root and shoot signals since they can be considered as appropriate approximations to the underlying dependence structure.

### 2.2 Testing Procedure

Once the model is defined, the goal of this work is therefore to understand which structure of the matrix 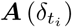 in (2) best describes the observed data (e.g. diagonal, lower/upper triangular). In this perspective, we are interested in making a decision on the following set of hypotheses:

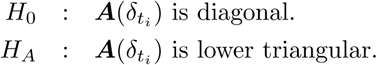

Hence, the null hypothesis *H*_0_ states that neither signal has an impact on the other (i.e. no Granger causality in the bivariate time series) while the alternative *H*_*A*_ states that the first signal Granger-causes the second. This alternative can of course be changed to “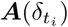 is upper triangular” therefore reversing the direction of dependence.

The MLE defined in (6) allows to estimate the parameters of the proposed model using the likelihood function in (5). Based on the latter, a commonly used test to determine the performance of a more “simple” model (such as the one considered in the null hypothesis stated above) with respect to a more “complex” model (such as the one in the alternative hypothesis) is the likelihood-ratio test whose statistic is given by

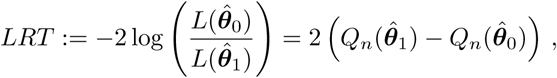

where 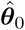 and 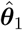 represent the estimated parameters of the models under the null and alternative hypothesis respectively. In order to perform this test one needs to derive the distribution of the *LRT* statistic under the null hypothesis which is asymptotically chi-squared with *p*^⋆^ degress of freedom, where *p*^⋆^ represents the number of extra parameters contained in 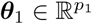 with respect to 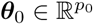 (i.e. *p*^⋆^ := *p*_1_ − *p*_0_). Using this distribution and the observed *LRT* statistic one can then reject or not the null hypothesis thereby concluding whether or not a signal Granger-causes the other.

### 2.3 Implementation

As highlighted before, the sample sizes coming from target biological applications are typically small (i.e. 5 *< n <* 20 time points) and it therefore seems unreasonable to make use of asymptotic properties in these cases. For this reason, Monte-Carlo-based techniques appear to be a natural alternative that are able to consider the small sample distribution of the test statistics of interest. More specifically, we propose to use parametric bootstrap to derive the small sample distribution of the *LRT* statistic under the null hypothesis as described in Algorithm 1.

#### Algorithm 1: Parametric Bootstrap for *LRT* Statistic

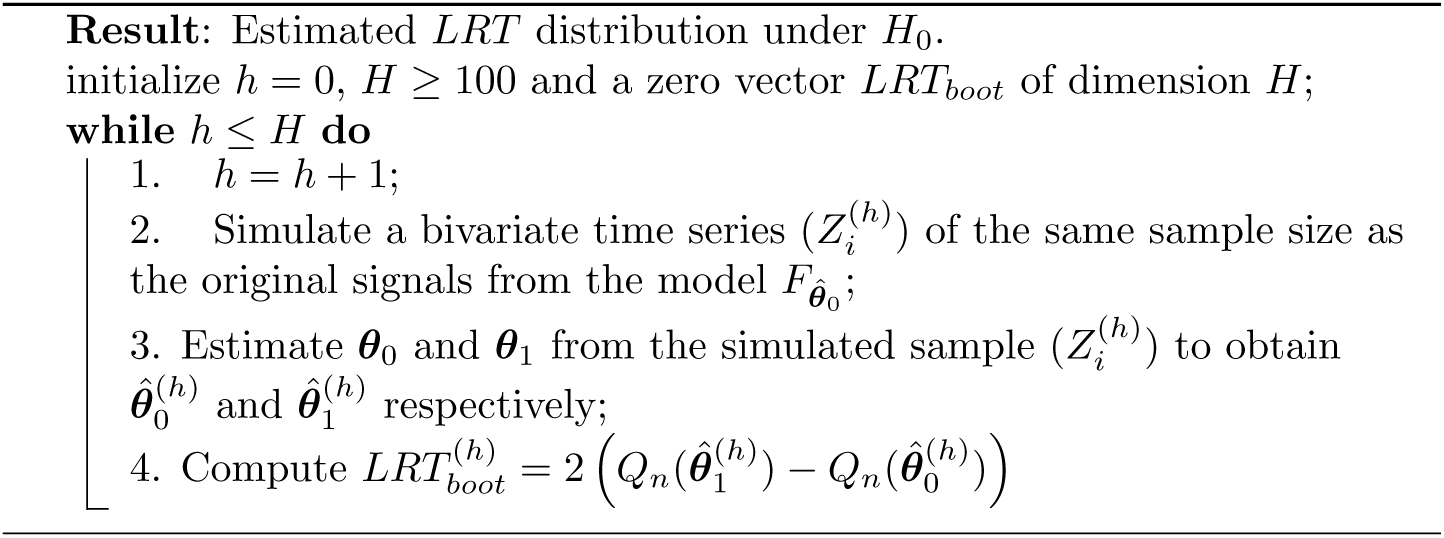

The parametric bootstrap approach allows for a good approximation (for large *H*) of the *LRT* statistic distribution under the null hypothesis by using the empirical distribution of the 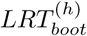 values. Given this distribution, it is possible to obtain an approximate *p*-value (see Davison and Hinkley, 1997) as follows

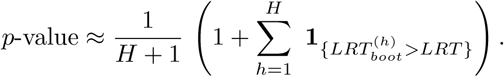

If this *p*-value is smaller than a chosen level of significance *α*, then we can reject the null hypothesis *H*_0_ that there is no Granger causality in favor of the specific alternative hypothesis *H*_*A*_ being tested.

Given this testing framework, there are a couple of issues that need to be considered, the first of which is the computational burden of Algorithm 1. In fact, the above defined *p*-value needs to be computed for all possible bivariate signal combinations and alternative hypotheses resulting in 2 *× N*_*X*_ *× N*_*Y*_ tests, where *N*_*X*_ and *N*_*Y*_ are the number of measured expressions in the two considered signals. Considering that the computational complexity to obtain the above *p*-value, given our assumptions, is approximately of order 𝒪 (*nH*), the final algorithmic complexity of the entire procedure would be of order 𝒪 (*nHM*) with *M* = *N*_*X*_ × *N*_*Y*_ and *N*_*X*_, *N*_*Y*_ ≫ 10^3^. This implies that the time required to obtain the results can be considerable. Another issue consists in the multiple testing framework this procedure entails, which therefore has consequences in terms of False Discovery Rate (FDR). Indeed, each 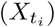 signal is tested 2*N*_*Y*_ times (and vice-versa for the 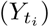 signals) which would require to compare the *p*-value to the level *α/*(2*N*_*X*_ *N*_*Y*_) if applying, for example, a Bonferroni correction. If the sizes *N*_*X*_ and *N*_*Y*_ are considerable, this would require to increase the number of simulations *H* in a proportional manner consequently increasing the computational burden. Unless one uses the asymptotic approximation to obtain a *p*-value (which would be highly unreliable for the small sample sizes used in these settings), there is currently no way of avoiding such a computational bottleneck.

## 3 Results and Discussion

The described approach was applied to the time-evolved transcriptome of Arabidopsis roots and shoots (the 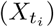 and 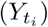 signals respectively) whose measurements were made through an experimental setup described more in detail in the supplemental material along with the chosen pre-processing (Section 2). These signals, each of length *n* = 10 and collected at higher frequencies in the initial experimental phase, generate 1,234,264 possible gene pairs from significantly differentially expressed root and shoot genes. Using *H* = 10^3^, we apply the procedure described in Section 2 which produces a final list of 3,078 gene pair interactions whose details are listed in Table 1, Section 3 of supplementary material (only *p*-values equal to zero were considered to reduce FDR as much as possible given computational constraints). Out of these interactions, 2,012 had a predicted root-to-shoot direction of influence meaning that the root gene was identified as being the (Granger) causal gene, or the influencer on the expression of the shoot gene. The remaining 1,066 interactions had a predicted shoot-to-root direction of influence. In addition, the approach predicted 1,616 positive interactions (i.e. *ψ >* 0) and 1,462 negative interactions (i.e. *ψ <* 0). Due to the limited and irregular number of samples across time, we choose to classify the time of influence at which the maximum influence between two genes occurred (measured by the *γ* parameter) into three general groups: Early (0-15 min), Middle (20-45 min), and Late (60-120 min). Based on this, among the 3,078 interactions, 2,502 occur Early, 548 occur during the Middle time frame, and 28 occur Late. In the following paragraphs we analyse only some of the model-predicted interactions in terms of their known properties and/or based on how they have a coherent biological interpretation. To do so, we will use the term “causal” to indicate genes that impact another gene, the latter being referred to as “target”.

**Table 1:**
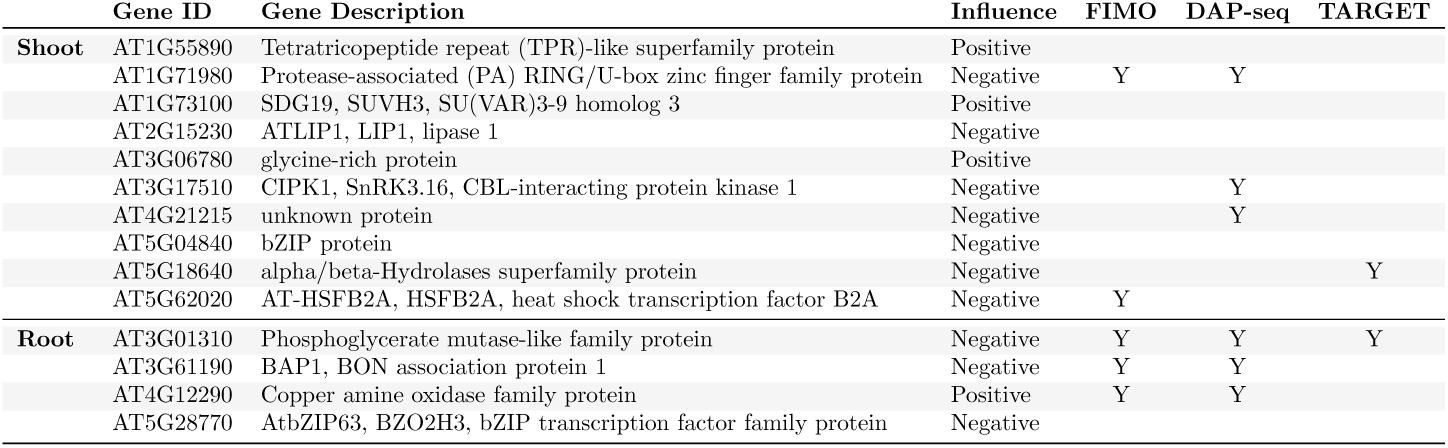
TGA1 target genes in root and shoot with which genes have a TGA1 motif occurrence of p *<* 0.0001 from the FIMO promoter analysis, and genes with which TGA1 has been shown to physically bind to based on DAP-Seq and TARGET experiments. “Y” indicates existing evidence for a predicted interaction from a specific experiment, whereas empty cells indicate possible avenues of future investigation.

### 3.1 Global analysis of model-predicted interactions reveal links between biological processes and pathways

Gene Ontology (GO) term analysis was performed to understand what pathways and processes are influenced across tissues over time (see Section 2 in supplemental material for more details). As highlighted also in Fig. 1, at early time points (0 - 15 minutes), causal root-genes reflect the early nitrogen response, while target-shoot genes reflect post-transcriptional and translational processes (see Tables 2 and 11, Section 3 of supplemental material). At later time points there is a shift in metabolism in which causal root-genes are involved in degradation and catabolic processes (45 - 120 minutes) (GO enrichment p-value *<* 0.01), while the predicted shoot target genes are involved in peptide biosynthesis (15 - 45 minutes) (GO enrichment p-value *<* 0.01) and sugar/carbohydrate response and signaling (45 - 120 minutes) (GO enrichment p-value *<* 0.01) (see Tables 3,4,12 and 13, Section 3 of supplemental material). GO analysis of the causal shoot-genes reflect the synthesis of shoot-derived signals, such as peptides and hormones, while the identified target root-genes are involved in phosphorus metabolic processes (0 - 15 minutes), lateral root development (15 - 45 minutes), and response to cytokinin (45 - 120 minutes) (see Tables 5-10, Section 3 of supplemental material). This analysis reflects much of the current knowledge about long distance nitrogen signaling between roots and shoots (Ruffel *et al*., 2011; Ko and Helariutta, 2017; Poitout *et al*., 2018).

**Figure 1:**
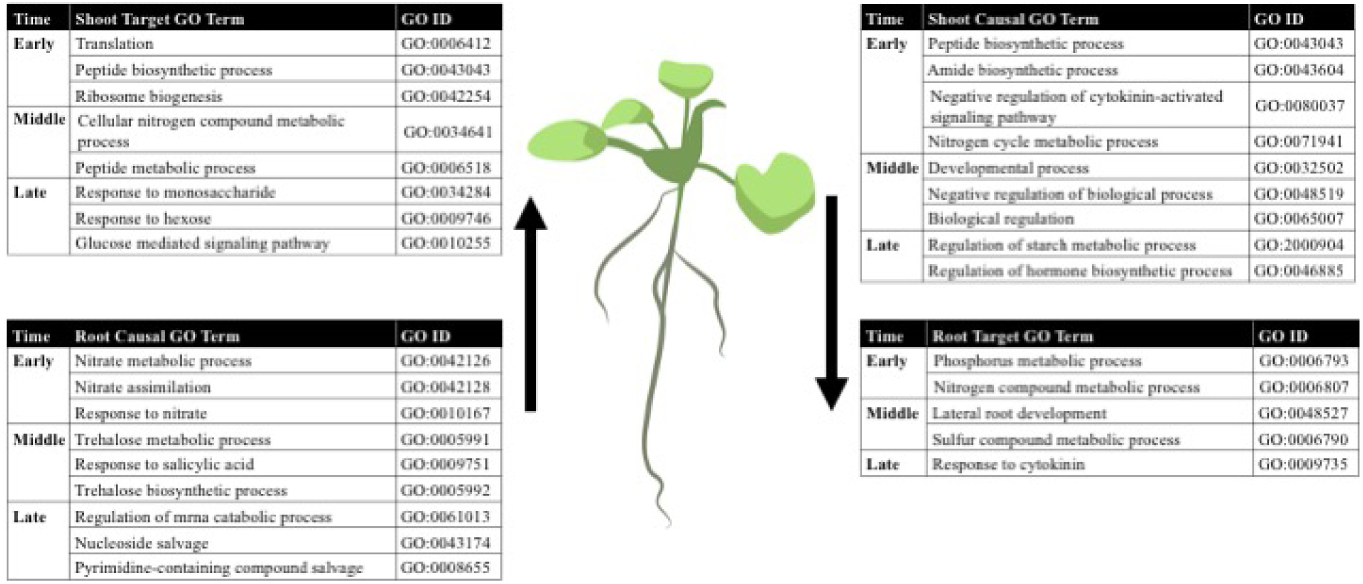
Selected enriched GO terms for root causal, shoot causal, root target and shoot target genes.

### 3.2 Model predictions are supported by in planta observations

A gene network was constructed where nodes (1,322 nodes) represent genes and edges (3,078 edges) constitute the model-predicted interactions described above (see Section 2 in supplemental material). Network analysis revealed that the gene interaction network with model-defined edges closely follows a power law distribution (*R*^2^ = 0.92), indicative of a scale-free biological network (Barabási, 2003; Albert, 2005). The validity of this finding was supported by a simulation of 10^3^ randomly generated networks using the same number of nodes and edges whose *R*^2^ values for the power law distribution were all between 0 and 0.35 (see Fig. 2). Network analysis for out-degree identified causal hub genes that are predicted to be highly influential in the temporal root-shoot transcriptomes in response to nitrogen treatment. Taking into consideration directionality, the top ten hubbiest genes in the network, based on out-degree, include three transcription factors previously implicated in the Arabidopsis nitrogen response: AFB3 (AT1G12820) (Vidal *et al*., 2013b, 2014; Xu and Cai, 2019), BT1 (AT5G63160) (Vidal *et al*., 2013a; Araus *et al*., 2016; Sato *et al*., 2017), and WRKY38 (AT5G22570) (Scheible *et al*., 2004; Gaudinier *et al*., 2018) (see Table 4, Section 3 of supplementary material). Other network hubs include the TF RD21A (AT1G47128) that is involved in autophagy and senescence which are key nitrogen turnover processes; and the RNA binding protein CID10 (AT3G49390), which is a poly(A) binding protein (PABP) potentially involved in mRNA stability or degradation (see “Supplemental Network File”). Further investigation of the interaction network revealed a number of previously identified genes and gene-gene relationships involved in local and long-distance nitrogen signaling, namely those involved in transcriptional regulation and in long-distance signaling by hormones and peptides, which are described in detail in the following sections.

**Figure 2:**
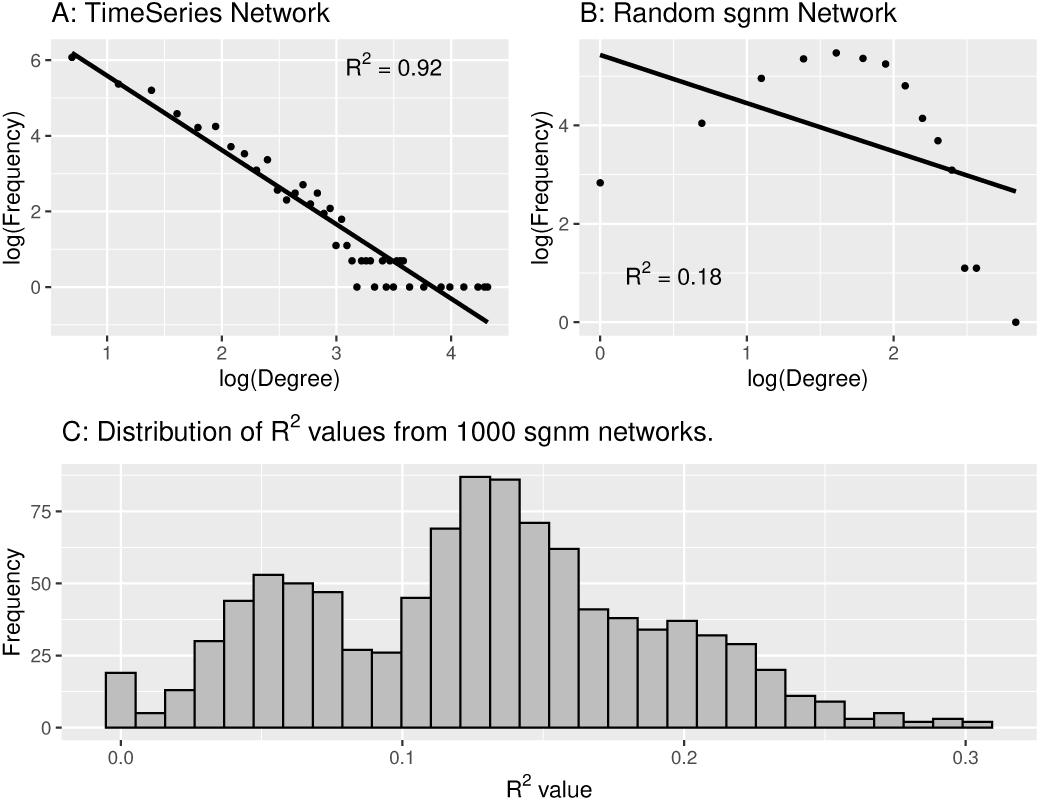
A: Node degree distribution for the network generated from model predictions (*R*^2^ = 0.92). B: Node degree distribution for a randomly generated network (*R*^2^ = 0.18). C: Histogram of the *R*^2^ values for the node degree distribution of 1,000 randomly generated networks (0 *≤ R*^2^ *<* 0.35).

### 3.3 Regulators of nitrogen processes

The transcription factors TGA1 and TGA4 were shown to be involved in mediating the primary nitrate response in roots by regulating the expression of the nitrate transporters NRT1.1 and NRT2.2, and also by coordinating the root developmental response to nitrate (Alvarez *et al*., 2014). From our analysis, root-expressed TGA1 is predicted to influence the expression of ten shoot genes, while shoot-expressed TGA1 is predicted to influence the expression of four root genes (see Tab. 1). To further investigate these predicted relationships, promoter analysis using FIMO from MEME Suite (Bailey and Machanick, 2012) was performed (as outlined in Section 2 of supplemental material). At least one TGA1 binding motif had a significant occurrence (FIMO p-value *<* 0.0001) in the putative promoters of two of the targeted shoot genes: a protease-associated RING/U-Box zinc finger family protein (AT1G71980) and HSFB2A heatshock transcription factor B2A (AT5G62020). The TGA1 motif also had a significant occurrence (FIMO p-value *<* 0.0001) in the three root target genes: a phospho-glycerate mutase-like family protein (AT3G01310), BAP1 BON association protein 1 (AT3G61190) and a copper amine oxidase family protein (AT4G12290). DAP-seq (DNA affinity purification sequencing) is an experimental technique allowing for the discovery of transcription factor binding sites on genomic DNA in vitro. A recent DAP-seq experiment showed that TGA1 actively binds to three shoot genes, AT1G71980, CIPK1 (AT3G17510) and an unknown protein (AT4G21215), as well as the three root target genes from the promoter analysis (O’Malley *et al*., 2016) (see Tab. 1). Furthermore, the model-predicted targets of TGA1, Phosphoglycerate mutase-like family protein (AT3G01310), and alpha/beta-Hydrolases superfamily protein (AT5G18640) were predicted to be direct targets of TGA1 in a TARGET (Transient Assay Reporting Genome-wide Effects of Transcription factors) assay experiment in root protoplasts by Brooks *et al*. (2019). A TARGET assay can identify candidate transcription factor targets based on TF-induced changes in gene expression (Brooks *et al*., 2019). These in-planta results provide support for the predicted interactions between TGA1 and its target genes within the same tissue, but additional studies will be needed to test if these interactions occur directly or indirectly between tissues.

### 3.4 Long-distance signaling by hormones and peptides Cytoknin Response Factors (CRFs)

Transcription factors (TF) with previously described regulatory roles in nitrogen uptake and assimilation include members of the ERF, bZIP, and NLP TF families (Konishi and Yanagisawa, 2013; Krapp *et al*., 2014; Vidal *et al*., 2015; Varala *et al*., 2018; Brooks *et al*., 2019). Of particular interest are the ERF TFs CRF 1-5. These CRFs were previously implicated in nitrogen signaling, targeting genes involved in nitrogen uptake and assimilation (Varala *et al*., 2018; Brooks *et al*., 2019). In our analysis, CRF5 expressed in the shoot was predicted to positively influence the expression of a heavy metal transport/detoxification protein (AT5G03380) expressed in the root. Using Elefinder (Hudson, 2005), CRF5 has been shown to bind to the GCC-box motif (GCCGCC) (Fujimoto *et al*., 2000; Sakuma *et al*., 2002; Liang *et al*., 2010) which is over-represented in the 2kb promoter region of AT5G03380 (E-value = 5.85 · 10^*-*4^, see Section 2 of supplemental material), indicating potential for a physical protein-DNA binding interaction. Shoot-expressed CRF3 is a predicted target of the causal root-expressed gene AT4G34419 (an unknown protein) in which AT4G34419 positively influences the expression of CRF3. Root-expressed CRF4 is predicted to influence the expression of the shoot genes SAUR-like auxin responsive protein family (AT4G34750), and Late embryogenesis abundant (LEA) hydroxyproline-rich glycoprotein family (AT3G44380). CRF4 is predicted to positively influence both of these genes during the early time interval. Like CRF5, CRF4 binds to the GCC-box motif, and this motif is overepresented in the 2kb upstream region of AT4G34750 (E-value = 1.43 · 10^*-*2^, see Section 2 of supplemental material). Root CRF4 is also predicted to negatively influence the shoot gene Homeobox Protein 6 (HB6, AT2G22430) during the middle time interval. CRF4 was shown to bind to HB6 via DAP-Seq (O’Malley *et al*., 2016). HB6 is a known negative regulator of the abscisic acid (ABA) signaling pathway (Himmelbach *et al*., 2003; Fujita *et al*., 2011). The ABA pathway is a phytohormone signalling pathway that was previously implicated in coordinating the long-distance nitrogen response (Kiba *et al*., 2011; Guan, 2017). A recent study by Varala *et al*. (2018) showed that CRF4 targets the TFs SNZ1 and CDF1, which in turn target HB6. The overexpression of CRF4 decreased the rate of nitrate uptake and altered root architecture in response to nitrogen treatment compared to WT plants (Varala *et al*., 2018). In CRF4 overexpressors, there was a decrease in primary root length and lateral root number under low nitrate conditions. Lateral root development has been shown to be inhibited under low nitrate conditions, which trigger ABA accumulation (Signora *et al*., 2001; Vidal *et al*., 2010; Léran *et al*., 2015; Sun *et al*., 2017). Thus, the results of our analysis suggest a coherent type 4 feed-forward loop (Mangan and Alon, 2003) in which root CRF4 represses shoot HB6 which represses whole plant ABA signaling (see Fig. 3), and may have physiological consequences for the observed changes in lateral root formation (Varala *et al*., 2018).

**Figure 3:**
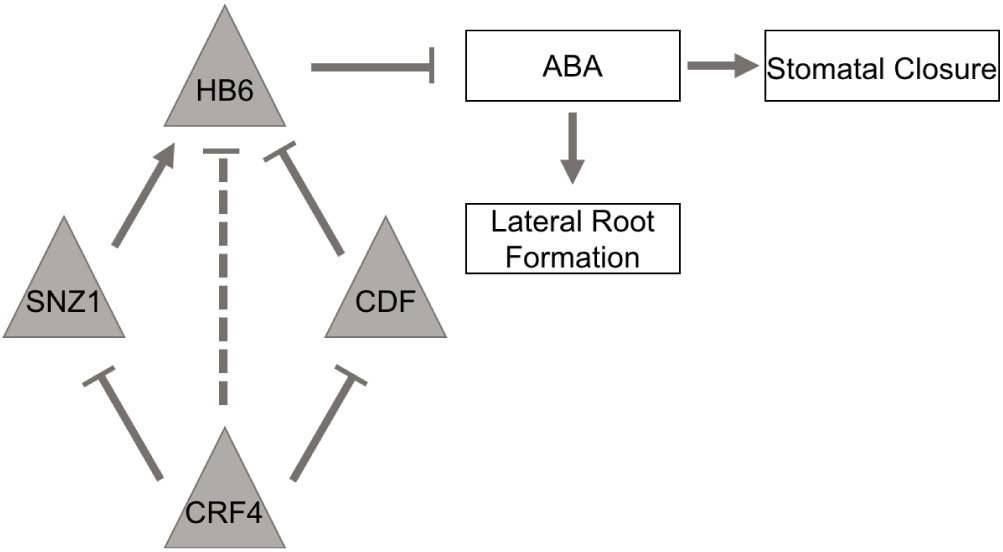
CRF4 interaction pathway, with flat-head arrows indicating negative interactions, pointed-head arrows indicating positive interactions, dashed arrows representing predicted interactions from the model, and solid arrows representing known interactions.

### Arabidopsis Response Regulators (ARRs)

The cytokinin signaling pathway is triggered by nitrogen and has been shown to be involved in the coordination of both root-to-shoot and shoot-to-root nitrogen-responses. In the shoots, cytokinins stimulate cell division and differentiation, whereas in the roots cytokinins reduce the activity of nitrogen uptake (Sakakibara *et al*., 2006). Cytokinins have also been shown to induce the expression of ARRs, which then regulate cytokinin signaling through feedback (To *et al*., 2007, 2004). For example, ARR4 (AT1G10470) is a Type-A response regulator that negatively regulates the cytokinin response (To *et al*., 2007). In our study, root ARR4 is predicted to influence the expression of three shoot genes (see Table 1, Section 3 of supplementary material), including a transmembrane amino acid transporter family protein (ATAVT1B; AT3G54830). During the middle time interval, root-expressed ARR4 is predicted to negatively influence the expression of AVT1 in shoots. Yeast AVT1 homologues have been implicated in the vacuolar uptake of large neutral amino acids including glutamine, asparagine, isoleucine, and tyrosine (Russnak *et al*., 2001; Tone *et al*., 2015) where they are stored in the vacuole under high nitrogen conditions (Sekito *et al*., 2008). When nitrogen starvation occurs, several AVT genes are upregulated to facilitate the export of the stored amino acids from the vacuole to the cytoplasm for protein synthesis (Fujiki *et al*., 2017). The analysis detected a relationship between ARR4 and AVT1B suggesting a potential mechanism by which cytokinin-induced ARR4 in the root may provide a long-distance signal to regulate shoot vacuolar amino acid import under high nitrogen conditions, like those used in this study.

### Peptides

Signal peptides have been implicated in the whole plant response to nitrogen (Tabata *et al*., 2014; Ohkubo *et al*., 2017; Oh *et al*., 2018). In the present study, seven peptides were uncovered as causal genes involved in 20 interactions (see Table 16, Section 3 of supplementary material). ATPSK4 is a Phytosulfokine 3 precursor and was shown to influence plant growth and cellular longevity, in particular root growth (Matsubayashi *et al*., 2006). CLE (Clavata3/ESR-related) peptides have long been known to be involved in long-distance nitrogen-signaling in legumes and have also been shown to be involved in nitrogen-signaling in Arabidopsis (Bidadi *et al*., 2014; Okamoto *et al*., 2016). In the present study, three CLE peptides are present in the predicted long-distance signaling network; CLE3 (AT1G06225), CLE4 (AT2G31081), and CLE27 (AT3G25905). CLE3 is a predicted causal gene expressed in the shoot that influences the root-expressed gene AT5G52530 (dentin sialophosphoprotein-related), while CLE4 is a causal root-gene predicted to influence the expression of four shoot-expressed genes either negatively (AT5G67510 Translation protein SH3-like family protein; AT1G55890 Tetratricopeptide repeat (TPR)-like superfamily protein) or positively (AT3G61620 RRP41, 3’-5’-exoribonuclease family protein; AT5G18640 alpha/beta-Hydrolases superfamily protein). Lastly, CLE27 is a Clavata family gene that was previously shown to be repressed by auxin (Wang *et al*., 2016). In the our study, CLE27 is a shoot expressed causal gene predicted to positively influence the expression of AT5G03380 (Heavy metal transport/detoxification superfamily protein) in the root. Devil/Rotundifolia Like (DVL) peptides are non-secretory peptides, conserved in plants, that can act as small signaling molecules and influence development in Arabidopsis (Wen *et al*., 2004). MTDVL1 was previously shown to be involved in symbiosis in *Medicago truncatula*, in which it has a negative regulatory role in nodulation (Combier *et al*., 2008). Two Devil peptides were identified in our analysis: DVL4 and DVL11. Of the four interactions involving DVL11, root DVL11 is predicted to be the causal gene influencing three shoot genes. Of these, DVL11 is predicted to positively influence the expression of ICK1, a cyclin-dependent kinase inhibitor family protein (AT2G23430). ICK1 is a known key regulator in development, and can inhibit entry into mitosis (Weinl *et al*., 2005). Root DVL4 is also predicted by the analysis to influence three shoot genes. Specifically, root DVL4 is predicted to positively influence shoot TCP-1/cpn60 chaperonin family protein (AT3G13470) at a middle time point. A previous study explored the transcriptional landscape of a DVL4 overexpressor line and showed that over-expression of DVL4 resulted in the up-regualtion of a number of genes encoding transcription factors (Larue *et al*., 2010). Our re-analysis of the microarray data from this study (see Section 2 of supplemental material) revealed that TCP1 was downregulated in DVL4 overexpressor plants compared to wild type Arabidopsis plants, providing support for a gene-gene interaction between DVL4 and TCP1 (see Fig. 4); however, this needs further exploration in the context of a nitrogen-signal.

**Figure 4:**
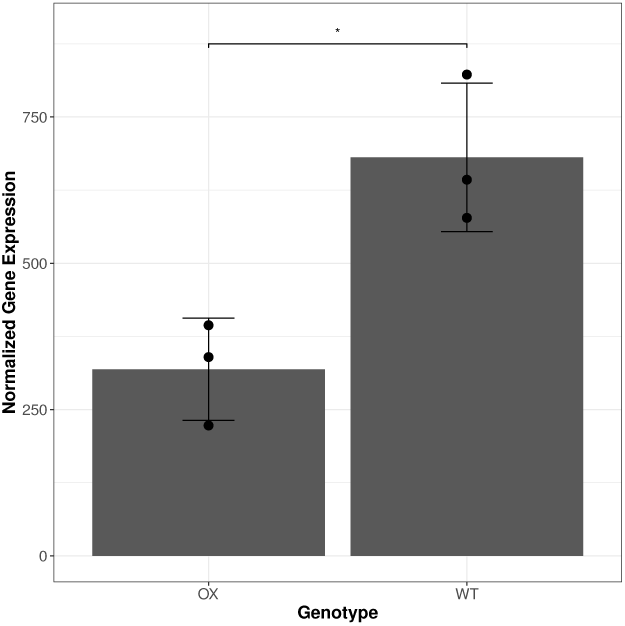
Bar chart of the normalized gene expression obtained from GSE8975 via GEO2R for TCP-1 in DVL4 overexpressor (OX) and wild-type (WT) Arabidopsis plants (t-test: p-value *<* 0.05).

### 3.5 Model predictions contain an over-representation of mobile causal gene products

The proposed approach, as stated previously, aims at understanding if the expression of one gene influences the expression of its target gene through the notion of Granger-Causality. Biologically, this influence may be direct or indirect.

It has previously been shown that mobile mRNAs that originate from one cell-type or organ can translocate to another cell-type or organ and have functional activity there (Lough and Lucas, 2006; Banerjee *et al*., 2009; Luo *et al*., 2018). To identify potential direct, long-distance interactions, we took advantage of two recent publications (Thieme *et al*., 2015; Guan *et al*., 2016) with extensive lists of experimentally determined mobile mRNAs that travel from root-to-shoot and from shoot-to-root. The lists of directional, causal genes from our model were intersected with the mobile transcripts identified by these studies. This analysis provided support for 204 causal genes involved in 340 predicted root-to-shoot, and 241 predicted shoot-to-root relationships; meaning that the direction of influence of the causal gene was the same in our analysis as that experimentally determined by these studies. An over-representation analysis (see Section 2 in supplemental material) was performed with the following hypotheses: “*H*_0_: the proposed approach (model) is equivalent to detecting known mobile transcripts randomly” and (alternative) “*H*_*A*_: the proposed approach (model) detects more known mobile transcripts than random selection”. In this case the *p*-value is 0 allowing us to reject the null hypothesis and hence the model is able to detect mobile transcripts which are potentially able to interact directly with their target genes. At least 36 of the total causal genes are known RNA-binding proteins (Marondedze *et al*., 2016), and 21 of these are mobile (see Table 17, Section 3 of supplemental material). In general, RNA-binding proteins can form ribonucleoprotein complexes (RNPs) that facilitate phloem transport and long distance trafficking of RNA molecules (Ham *et al*., 2009; Kehr and Kragler, 2018). An additional 79 causal genes involved in 203 relationships (121 root-to-shoot and 82 shoot-to-root) have not been experimentally shown to be mobile, but are predicted to produce an mRNA molecule that possesses a t-RNA like motif. Guan *et al*. (2016) also have hypothesised that some mRNA have a tRNA-like structure in their sequence. This allows the mRNA to fold into a tRNA-like shape that confers some stability to the mRNA strand. This stability allows the mRNA to move long distances in the plant. These results suggest that a large proportion of the model-predicted causal genes have the potential to influence the expression of its target gene (directly or indirectly) via long-distance vascular trafficking. One example of a model-predicted gene interaction that may function through interaction of a mobile causal gene with its target is the relationship between root derived aconitase 2 (ACO2), predicted to have a negative influence on the expression of malate dehydrogenase (MDH2) in the shoot. ACO2 is the only isoform of aconitase that is specifically induced by nitrogen treatment. Root ACO2 is involved in the TCA cycle, while shoot MDH2 is localized in the mitochondria and involved in gluconeogenesis. One possibility is that a direct or downstream gene product of root ACO2 represses shoot MDH2, resulting in possible down-regulation of shoot gluconeogenesis in response to a large, transient nitrogen signal. Although the specific mechanism of this relationship needs experimental exploration, it is partially supported by existing data describing the tight relationship between carbon and nitrogen metabolism to maintain whole plant C:N balance (Palenchar *et al*., 2004; Zheng, 2009; Goel *et al*., 2016). Alternatively, aconitase, an iron-sulphur protein, has been shown to be a bifunctional enzyme/RNA-binding protein that binds to iron-responsive elements in target RNA to stabilize the transcript and function in iron homeostasis (Hentze and Argos, 1991). Our analysis predicted a positive relationship between ACO2 (causal root) and Ironman 1 (target shoot), an Fe-uptake inducing peptide 3 that is involved in the regulation of iron deficiency response genes (Grillet *et al*., 2018). It was previously shown that nitrogen treatment induces the expression of genes involved in iron uptake, transport, and homeostasis in plants (Wang *et al*., 2000, 2003), and that the form of nitrogen taken up by roots influences the amount of iron accumulation in leaves (Zou *et al*., 2001). There is also a well-established relationship between nitrogen and Fe pathways since Fe is a component of many enzymes involved in nitrate assimilation (Wang *et al*., 2003).

## 4 Conclusions

This work puts forward an approach to perform Granger-Causal analysis for (small-sample) irregularly-spaced bivariate signals which overcomes existing limitations in the analysis of biological time series data following this common sampling scheme. Based on this new framework, (Granger) causal relationships were detected and whole-organism molecular response to a nitrogen signal were predicted. The survey of genes with predicted temporal cause-and-effect relationships enabled discovery of coordinated biological processes and chemical pathways that communicate the nitrogen-signal between roots and shoots of plants. These coordinated processes can now be further investigated to identify potential regulatory bottlenecks that influence whole plant nitrogen uptake/utilization efficiency. The abundance of genes involved in the known transcriptional nitrogen-response (nitrogen-transport and assimilation) as both causal and target genes indicate that the proposed approach was able to capture whole-plant response to a transient nitrogen-treatment across tissues. The predicted cross-organ dependencies provide insights and hypotheses about potential signaling cascades that are triggered sequentially as the nitrogen-signal propagates from roots-to-shoots-to-roots. Importantly, regulatory factors that have not previously been implicated in whole plant nitrogen-response were highlighted by the proposed approach. These novel factors can be targets for engineering to enhance plant nitrogen uptake/utilization efficiency. The findings from this research will have implications for predicting causal molecular relationships that influence intercellular, long-distance nitrogen-signaling, and the methodological framework proposed in this work is applicable to researchers struggling with meaningful integration of dynamic, system-wide transcriptome data.

## Supporting information

Supplemental Network

