## Supplemental Network for "Granger-Causal Testing for Irregularly Sampled Time Series with Application to Nitrogen Signaling in Arabidopsis": networks.js

```
var networks = {"TSnetwork.csv": {
  "format_version" : "1.0",
  "generated_by" : "cytoscape-3.7.2",
  "target_cytoscapejs_version" : "~2.1",
  "data" : {
    "shared_name" : "TSnetwork.csv",
    "name" : "TSnetwork.csv",
    "SUID" : 62,
    "__Annotations" : [ ],
    "selected" : true
  },
  "elements" : {
    "nodes" : [ {
      "data" : {
        "id" : "1393",
        "symbol" : "AT5G14480",
        "degree_layout" : 8,
        "Shoot_Descriptions" : "",
        "PLAMOM_Match_Shoot" : "",
        "Causal_TF" : "4-N-acetylglucosaminyltransferase family protein\"",
        "Root_Descriptions" : "beta-1,4-N-acetylglucosaminyltransferase family protein",
        "Eccentricity" : 10,
        "Outdegree" : 5,
        "Target_PLAMOM" : "no",
        "PLAMOM_Match_Root" : "NO",
        "shared_name" : "AT5G14480",
        "Indegree" : 6,
        "Target_tRNA" : "no",
        "t_RNA_motif_shoot" : "",
        "SUID" : 1393,
        "SelfLoops" : 0,
        "Target_TF" : "4-N-acetylglucosaminyltransferase family protein\"",
        "MolecularAttributes" : "protein_coding",
        "AverageShortestPathLength" : 5.79916318,
        "selected" : false,
        "Name2" : "AT5G14480",
        "ClosenessCentrality" : 0.17243867,
        "Casual_Name" : "\"beta-1",
        "PartnerOfMultiEdgedNodePairs" : 0,
        "ClusteringCoefficient" : 0.0,
        "Target_Name" : "\"beta-1",
        "Stress" : 6429,
        "BetweennessCentrality" : 0.00118153,
        "t_RNA_motif_Root" : "NO",
        "EdgeCount" : 11,
        "Causal_PLAMOM" : "no",
        "name" : "AT5G14480",
        "Causal_tRNA" : "no",
        "IsSingleNode" : false,
        "briefDescription" : "beta-1",
        "NeighborhoodConnectivity" : 8.545454545454545
      },
      "position" : {
        "x" : -6655.68359375,
        "y" : 2761.765625
      },
      "selected" : false
    }, {
      "data" : {
        "id" : "1392",
        "symbol" : "AT1G01180",
        "degree_layout" : 3,
        "Shoot_Descriptions" : "S-adenosyl-L-methionine-dependent methyltransferases superfamily protein",
        "PLAMOM_Match_Shoot" : "NO",
        "Causal_TF" : "no",
        "Root_Descriptions" : "",
        "Eccentricity" : 11,
        "Outdegree" : 2,
        "Target_PLAMOM" : "no",
        "PLAMOM_Match_Root" : "",
        "shared_name" : "AT1G01180",
        "Indegree" : 1,
        "Target_tRNA" : "no",
        "t_RNA_motif_shoot" : "NO",
        "SUID" : 1392,
        "SelfLoops" : 0,
        "Target_TF" : "S-adenosyl-L-methionine-dependent methyltransferases superfamily protein",
        "MolecularAttributes" : "protein_coding",
        "AverageShortestPathLength" : 6.76150628,
        "selected" : false,
        "Name2" : "AT1G01180",
        "ClosenessCentrality" : 0.14789604,
        "Casual_Name" : "S-adenosyl-L-methionine-dependent methyltransferases superfamily protein",
        "PartnerOfMultiEdgedNodePairs" : 0,
        "ClusteringCoefficient" : 0.0,
        "Target_Name" : "no",
        "Stress" : 235,
        "BetweennessCentrality" : 5.394E-5,
        "t_RNA_motif_Root" : "",
        "EdgeCount" : 3,
        "Causal_PLAMOM" : "no",
        "name" : "AT1G01180",
        "Causal_tRNA" : "no",
        "IsSingleNode" : false,
        "briefDescription" : "S-adenosyl-L-methionine-dependent methyltransferases superfamily protein",
        "NeighborhoodConnectivity" : 11.333333333333334
      },
      "position" : {
        "x" : 3983.552685189421,
        "y" : 2940.468017578125
      },
      "selected" : false
    }, {
      "data" : {
        "id" : "1391",
        "symbol" : "AT2G31990",
        "Shoot_Descriptions" : "",
        "PLAMOM_Match_Shoot" : "",
        "Causal_TF" : "no",
        "Root_Descriptions" : "Exostosin family protein",
        "Eccentricity" : 9,
        "Outdegree" : 14,
        "Target_PLAMOM" : "no",
        "PLAMOM_Match_Root" : "NO",
        "shared_name" : "AT2G31990",
        "Indegree" : 2,
        "Target_tRNA" : "no",
        "t_RNA_motif_shoot" : "",
        "SUID" : 1391,
        "SelfLoops" : 0,
        "Target_TF" : "Exostosin family protein",
        "MolecularAttributes" : "protein_coding",
        "AverageShortestPathLength" : 4.00627615,
        "selected" : false,
        "Name2" : "AT2G31990",
        "ClosenessCentrality" : 0.24960836,
        "Casual_Name" : "Exostosin family protein",
        "PartnerOfMultiEdgedNodePairs" : 0,
        "ClusteringCoefficient" : 0.00416667,
        "Target_Name" : "no",
        "Stress" : 11041,
        "BetweennessCentrality" : 0.00265464,
        "t_RNA_motif_Root" : "NO",
        "EdgeCount" : 16,
        "Causal_PLAMOM" : "no",
        "name" : "AT2G31990",
        "Causal_tRNA" : "no",
        "IsSingleNode" : false,
        "briefDescription" : "Exostosin family protein",
        "NeighborhoodConnectivity" : 10.0
      },
      "position" : {
        "x" : -6794.6064453125,
        "y" : 3290.95849609375
      },
      "selected" : false
    }, {
      "data" : {
        "id" : "1390",
        "symbol" : "FKGP",
        "Shoot_Descriptions" : "AtFKGP, FKGP, L-fucokinase/GDP-L-fucose pyrophosphorylase",
        "PLAMOM_Match_Shoot" : "NO",
        "Causal_TF" : "FKGP",
        "Root_Descriptions" : "AtFKGP, FKGP, L-fucokinase/GDP-L-fucose pyrophosphorylase",
        "Eccentricity" : 9,
        "Outdegree" : 8,
        "Target_PLAMOM" : "AT5G41410",
        "PLAMOM_Match_Root" : "NO",
        "shared_name" : "AT1G01220",
        "Indegree" : 11,
        "Target_tRNA" : "no",
        "t_RNA_motif_shoot" : "NO",
        "SUID" : 1390,
        "SelfLoops" : 0,
        "Target_TF" : "\"AtFKGP",
        "MolecularAttributes" : "METABOLIC",
        "AverageShortestPathLength" : 4.78451883,
        "selected" : false,
        "Name2" : "FKGP",
        "ClosenessCentrality" : 0.20900743,
        "Casual_Name" : "\"AtFKGP",
        "PartnerOfMultiEdgedNodePairs" : 0,
        "ClusteringCoefficient" : 0.0,
        "Target_Name" : "no",
        "Stress" : 87208,
        "BetweennessCentrality" : 0.01903462,
        "t_RNA_motif_Root" : "NO",
        "EdgeCount" : 19,
        "Causal_PLAMOM" : "no",
        "name" : "AT1G01220",
        "Causal_tRNA" : "no",
        "IsSingleNode" : false,
        "briefDescription" : "L-fucokinase/GDP-L-fucose pyrophosphorylase",
        "NeighborhoodConnectivity" : 9.0
      },
      "position" : {
        "x" : -1025.881152701204,
        "y" : 3551.968017578125
      },
      "selected" : false
    }, {
      "data" : {
        "id" : "1389",
        "symbol" : "UGT84A2",
        "Shoot_Descriptions" : "",
        "PLAMOM_Match_Shoot" : "",
        "Causal_TF" : "UDP-Glycosyltransferase superfamily protein\"",
        "Root_Descriptions" : "UGT84A2, UDP-Glycosyltransferase superfamily protein",
        "Eccentricity" : 9,
        "Outdegree" : 14,
        "Target_PLAMOM" : "PPS",
        "PLAMOM_Match_Root" : "NO",
        "shared_name" : "AT3G21560",
        "Indegree" : 1,
        "Target_tRNA" : "RNI-like superfamily protein\"",
        "t_RNA_motif_shoot" : "",
        "SUID" : 1389,
        "SelfLoops" : 0,
        "Target_TF" : "no",
        "MolecularAttributes" : "METABOLIC",
        "AverageShortestPathLength" : 4.91493776,
        "selected" : false,
        "Name2" : "UGT84A2",
        "ClosenessCentrality" : 0.20346138,
        "Casual_Name" : "\"UGT84A2",
        "PartnerOfMultiEdgedNodePairs" : 0,
        "ClusteringCoefficient" : 0.0,
        "Target_Name" : "no",
        "Stress" : 522,
        "BetweennessCentrality" : 1.3011E-4,
        "t_RNA_motif_Root" : "NO",
        "EdgeCount" : 15,
        "Causal_PLAMOM" : "no",
        "name" : "AT3G21560",
        "Causal_tRNA" : "no",
        "IsSingleNode" : false,
        "briefDescription" : "UDP-Glycosyltransferase superfamily protein",
        "NeighborhoodConnectivity" : 7.133333333333334
      },
      "position" : {
        "x" : -6738.8076171875,
        "y" : 3436.499755859375
      },
      "selected" : false
    }, {
      "data" : {
        "id" : "1388",
        "symbol" : "FUT4",
        "degree_layout" : 1,
        "Shoot_Descriptions" : "",
        "PLAMOM_Match_Shoot" : "",
        "Causal_TF" : "FUT4",
        "Root_Descriptions" : "atfut4, FUT4, fucosyltransferase 4",
        "Eccentricity" : 1,
        "Outdegree" : 1,
        "Target_PLAMOM" : "no",
        "PLAMOM_Match_Root" : "NO",
        "shared_name" : "AT2G15390",
        "Indegree" : 2,
        "Target_tRNA" : "no",
        "t_RNA_motif_shoot" : "",
        "SUID" : 1388,
        "SelfLoops" : 0,
        "Target_TF" : "FUT4",
        "MolecularAttributes" : "protein_coding",
        "AverageShortestPathLength" : 1.0,
        "selected" : false,
        "Name2" : "FUT4",
        "ClosenessCentrality" : 1.0,
        "Casual_Name" : "\"atfut4",
        "PartnerOfMultiEdgedNodePairs" : 0,
        "ClusteringCoefficient" : 0.0,
        "Target_Name" : "\"atfut4",
        "Stress" : 697,
        "BetweennessCentrality" : 6.253E-5,
        "t_RNA_motif_Root" : "NO",
        "EdgeCount" : 3,
        "Causal_PLAMOM" : "no",
        "name" : "AT2G15390",
        "Causal_tRNA" : "no",
        "IsSingleNode" : false,
        "briefDescription" : "fucosyltransferase 4",
        "NeighborhoodConnectivity" : 7.0
      },
      "position" : {
        "x" : -6393.400390625,
        "y" : 2481.475830078125
      },
      "selected" : false
    }, {
      "data" : {
        "id" : "1387",
        "symbol" : "RCAR1",
        "Shoot_Descriptions" : "PYL9, RCAR1, regulatory component of ABA receptor 1",
        "PLAMOM_Match_Shoot" : "YES",
        "Causal_TF" : "",
        "Root_Descriptions" : "",
        "Eccentricity" : 0,
        "Outdegree" : 0,
        "Target_PLAMOM" : "AT1G01360",
        "PLAMOM_Match_Root" : "",
        "shared_name" : "AT1G01360",
        "Indegree" : 3,
        "Target_tRNA" : "no",
        "t_RNA_motif_shoot" : "NO",
        "SUID" : 1387,
        "SelfLoops" : 0,
        "Target_TF" : "RCAR1",
        "MolecularAttributes" : "protein_coding",
        "AverageShortestPathLength" : 0.0,
        "selected" : false,
        "Name2" : "RCAR1",
        "ClosenessCentrality" : 0.0,
        "Casual_Name" : "",
        "PartnerOfMultiEdgedNodePairs" : 0,
        "ClusteringCoefficient" : 0.0,
        "Target_Name" : "\"PYL9",
        "Stress" : 0,
        "BetweennessCentrality" : 0.0,
        "t_RNA_motif_Root" : "",
        "EdgeCount" : 3,
        "Causal_PLAMOM" : "",
        "name" : "AT1G01360",
        "Causal_tRNA" : "",
        "IsSingleNode" : false,
        "briefDescription" : "regulatory component of ABA receptor 1",
        "NeighborhoodConnectivity" : 12.333333333333334
      },
      "position" : {
        "x" : 4058.9096783965824,
        "y" : -498.49700927734375
      },
      "selected" : false
    }, {
      "data" : {
        "id" : "1386",
        "symbol" : "AT5G39590",
        "Shoot_Descriptions" : "",
        "PLAMOM_Match_Shoot" : "",
        "Causal_TF" : "no",
        "Root_Descriptions" : "TLD-domain containing nucleolar protein",
        "Eccentricity" : 14,
        "Outdegree" : 2,
        "Target_PLAMOM" : "",
        "PLAMOM_Match_Root" : "NO",
        "shared_name" : "AT5G39590",
        "Indegree" : 0,
        "Target_tRNA" : "",
        "t_RNA_motif_shoot" : "",
        "SUID" : 1386,
        "SelfLoops" : 0,
        "Target_TF" : "",
        "MolecularAttributes" : "protein_coding",
        "AverageShortestPathLength" : 8.93347193,
        "selected" : false,
        "Name2" : "AT5G39590",
        "ClosenessCentrality" : 0.11193856,
        "Casual_Name" : "TLD-domain containing nucleolar protein",
        "PartnerOfMultiEdgedNodePairs" : 0,
        "ClusteringCoefficient" : 0.0,
        "Target_Name" : "",
        "Stress" : 0,
        "BetweennessCentrality" : 0.0,
        "t_RNA_motif_Root" : "NO",
        "EdgeCount" : 2,
        "Causal_PLAMOM" : "no",
        "name" : "AT5G39590",
        "Causal_tRNA" : "no",
        "IsSingleNode" : false,
        "briefDescription" : "TLD-domain containing nucleolar protein",
        "NeighborhoodConnectivity" : 4.0
      },
      "position" : {
        "x" : -5591.45361328125,
        "y" : -356.202880859375
      },
      "selected" : false
    }, {
      "data" : {
        "id" : "1385",
        "symbol" : "BRG1",
        "degree_layout" : 6,
        "Shoot_Descriptions" : "",
        "PLAMOM_Match_Shoot" : "",
        "Causal_TF" : "no",
        "Root_Descriptions" : "SBP (S-ribonuclease binding protein) family protein",
        "Eccentricity" : 11,
        "Outdegree" : 6,
        "Target_PLAMOM" : "no",
        "PLAMOM_Match_Root" : "NO",
        "shared_name" : "AT5G45100",
        "Indegree" : 26,
        "Target_tRNA" : "no",
        "t_RNA_motif_shoot" : "",
        "SUID" : 1385,
        "SelfLoops" : 0,
        "Target_TF" : "no",
        "MolecularAttributes" : "protein_coding",
        "AverageShortestPathLength" : 6.10251046,
        "selected" : false,
        "Name2" : "BRG1",
        "ClosenessCentrality" : 0.16386699,
        "Casual_Name" : "SBP (S-ribonuclease binding protein) family protein",
        "PartnerOfMultiEdgedNodePairs" : 0,
        "ClusteringCoefficient" : 0.0,
        "Target_Name" : "SBP (S-ribonuclease binding protein) family protein",
        "Stress" : 107551,
        "BetweennessCentrality" : 0.02217023,
        "t_RNA_motif_Root" : "NO",
        "EdgeCount" : 32,
        "Causal_PLAMOM" : "no",
        "name" : "AT5G45100",
        "Causal_tRNA" : "no",
        "IsSingleNode" : false,
        "briefDescription" : "SBP (S-ribonuclease binding protein) family protein",
        "NeighborhoodConnectivity" : 5.5625
      },
      "position" : {
        "x" : -6629.3525390625,
        "y" : 3256.275390625
      },
      "selected" : false
    }, {
      "data" : {
        "id" : "1384",
        "symbol" : "AFB3",
        "degree_layout" : 18,
        "Shoot_Descriptions" : "AFB3, auxin signaling F-box 3",
        "PLAMOM_Match_Shoot" : "NO",
        "Causal_TF" : "auxin signaling F-box 3\"",
        "Root_Descriptions" : "AFB3, auxin signaling F-box 3",
        "Eccentricity" : 9,
        "Outdegree" : 24,
        "Target_PLAMOM" : "no",
        "PLAMOM_Match_Root" : "NO",
        "shared_name" : "AT1G12820",
        "Indegree" : 10,
        "Target_tRNA" : "no",
        "t_RNA_motif_shoot" : "NO",
        "SUID" : 1384,
        "SelfLoops" : 0,
        "Target_TF" : "auxin signaling F-box 3\"",
        "MolecularAttributes" : "protein_coding",
        "AverageShortestPathLength" : 4.59414226,
        "selected" : false,
        "Name2" : "AFB3",
        "ClosenessCentrality" : 0.21766849,
        "Casual_Name" : "\"AFB3",
        "PartnerOfMultiEdgedNodePairs" : 0,
        "ClusteringCoefficient" : 0.0,
        "Target_Name" : "\"AFB3",
        "Stress" : 54247,
        "BetweennessCentrality" : 0.00801656,
        "t_RNA_motif_Root" : "NO",
        "EdgeCount" : 34,
        "Causal_PLAMOM" : "no",
        "name" : "AT1G12820",
        "Causal_tRNA" : "no",
        "IsSingleNode" : false,
        "briefDescription" : "auxin signaling F-box 3",
        "NeighborhoodConnectivity" : 7.0
      },
      "position" : {
        "x" : -1153.811572623079,
        "y" : 3059.88525390625
      },
      "selected" : false
    }, {
      "data" : {
        "id" : "1383",
        "symbol" : "AT1G02370",
        "Shoot_Descriptions" : "Tetratricopeptide repeat (TPR)-like superfamily protein",
        "PLAMOM_Match_Shoot" : "NO",
        "Causal_TF" : "",
        "Root_Descriptions" : "",
        "Eccentricity" : 0,
        "Outdegree" : 0,
        "Target_PLAMOM" : "no",
        "PLAMOM_Match_Root" : "",
        "shared_name" : "AT1G02370",
        "Indegree" : 4,
        "Target_tRNA" : "no",
        "t_RNA_motif_shoot" : "NO",
        "SUID" : 1383,
        "SelfLoops" : 0,
        "Target_TF" : "Tetratricopeptide repeat (TPR)-like superfamily protein",
        "MolecularAttributes" : "protein_coding",
        "AverageShortestPathLength" : 0.0,
        "selected" : false,
        "Name2" : "AT1G02370",
        "ClosenessCentrality" : 0.0,
        "Casual_Name" : "",
        "PartnerOfMultiEdgedNodePairs" : 0,
        "ClusteringCoefficient" : 0.0,
        "Target_Name" : "no",
        "Stress" : 0,
        "BetweennessCentrality" : 0.0,
        "t_RNA_motif_Root" : "",
        "EdgeCount" : 4,
        "Causal_PLAMOM" : "",
        "name" : "AT1G02370",
        "Causal_tRNA" : "",
        "IsSingleNode" : false,
        "briefDescription" : "Tetratricopeptide repeat (TPR)-like superfamily protein",
        "NeighborhoodConnectivity" : 19.25
      },
      "position" : {
        "x" : 3720.944775033171,
        "y" : 309.9804382324219
      },
      "selected" : false
    }, {
      "data" : {
        "id" : "1382",
        "symbol" : "AT4G32480",
        "Shoot_Descriptions" : "",
        "PLAMOM_Match_Shoot" : "",
        "Causal_TF" : "no",
        "Root_Descriptions" : "Protein of unknown function (DUF506)",
        "Eccentricity" : 11,
        "Outdegree" : 14,
        "Target_PLAMOM" : "",
        "PLAMOM_Match_Root" : "YES",
        "shared_name" : "AT4G32480",
        "Indegree" : 0,
        "Target_tRNA" : "",
        "t_RNA_motif_shoot" : "",
        "SUID" : 1382,
        "SelfLoops" : 0,
        "Target_TF" : "",
        "MolecularAttributes" : "protein_coding",
        "AverageShortestPathLength" : 6.24123711,
        "selected" : false,
        "Name2" : "AT4G32480",
        "ClosenessCentrality" : 0.16022464,
        "Casual_Name" : "Protein of unknown function (DUF506)",
        "PartnerOfMultiEdgedNodePairs" : 0,
        "ClusteringCoefficient" : 0.0,
        "Target_Name" : "",
        "Stress" : 0,
        "BetweennessCentrality" : 0.0,
        "t_RNA_motif_Root" : "NO",
        "EdgeCount" : 14,
        "Causal_PLAMOM" : "AT4G32480",
        "name" : "AT4G32480",
        "Causal_tRNA" : "no",
        "IsSingleNode" : false,
        "briefDescription" : "sugar phosphate exchanger",
        "NeighborhoodConnectivity" : 8.214285714285714
      },
      "position" : {
        "x" : -6306.08984375,
        "y" : 364.8254089355469
      },
      "selected" : false
    }, {
      "data" : {
        "id" : "1381",
        "symbol" : "GSR2",
        "Shoot_Descriptions" : "",
        "PLAMOM_Match_Shoot" : "",
        "Causal_TF" : "GSR2",
        "Root_Descriptions" : "ATGSR2, GSR2, glutamine synthase clone F11",
        "Eccentricity" : 1,
        "Outdegree" : 2,
        "Target_PLAMOM" : "",
        "PLAMOM_Match_Root" : "YES",
        "shared_name" : "AT1G66200",
        "Indegree" : 0,
        "Target_tRNA" : "",
        "t_RNA_motif_shoot" : "",
        "SUID" : 1381,
        "SelfLoops" : 0,
        "Target_TF" : "",
        "MolecularAttributes" : "METABOLIC",
        "AverageShortestPathLength" : 1.0,
        "selected" : false,
        "Name2" : "GSR2",
        "ClosenessCentrality" : 1.0,
        "Casual_Name" : "\"ATGSR2",
        "PartnerOfMultiEdgedNodePairs" : 0,
        "ClusteringCoefficient" : 0.0,
        "Target_Name" : "",
        "Stress" : 0,
        "BetweennessCentrality" : 0.0,
        "t_RNA_motif_Root" : "NO",
        "EdgeCount" : 2,
        "Causal_PLAMOM" : "AT1G66200",
        "name" : "AT1G66200",
        "Causal_tRNA" : "no",
        "IsSingleNode" : false,
        "briefDescription" : "hypothetical protein",
        "NeighborhoodConnectivity" : 5.5
      },
      "position" : {
        "x" : -6629.42724609375,
        "y" : 481.2294006347656
      },
      "selected" : false
    }, {
      "data" : {
        "id" : "1380",
        "symbol" : "AOC3",
        "degree_layout" : 2,
        "Shoot_Descriptions" : "",
        "PLAMOM_Match_Shoot" : "",
        "Causal_TF" : "allene oxide cyclase 3\"",
        "Root_Descriptions" : "AOC3, allene oxide cyclase 3",
        "Eccentricity" : 12,
        "Outdegree" : 3,
        "Target_PLAMOM" : "AT3G25780",
        "PLAMOM_Match_Root" : "YES",
        "shared_name" : "AT3G25780",
        "Indegree" : 24,
        "Target_tRNA" : "no",
        "t_RNA_motif_shoot" : "",
        "SUID" : 1380,
        "SelfLoops" : 0,
        "Target_TF" : "allene oxide cyclase 3\"",
        "MolecularAttributes" : "METABOLIC",
        "AverageShortestPathLength" : 6.9874477,
        "selected" : false,
        "Name2" : "AOC3",
        "ClosenessCentrality" : 0.14311377,
        "Casual_Name" : "\"AOC3",
        "PartnerOfMultiEdgedNodePairs" : 0,
        "ClusteringCoefficient" : 0.0,
        "Target_Name" : "\"AOC3",
        "Stress" : 24803,
        "BetweennessCentrality" : 0.00542753,
        "t_RNA_motif_Root" : "NO",
        "EdgeCount" : 27,
        "Causal_PLAMOM" : "AT3G25780",
        "name" : "AT3G25780",
        "Causal_tRNA" : "no",
        "IsSingleNode" : false,
        "briefDescription" : "allene oxide cyclase 3",
        "NeighborhoodConnectivity" : 4.814814814814815
      },
      "position" : {
        "x" : -6894.26171875,
        "y" : 3124.17919921875
      },
      "selected" : false
    }, {
      "data" : {
        "id" : "1379",
        "symbol" : "AT2G27830",
        "Shoot_Descriptions" : "",
        "PLAMOM_Match_Shoot" : "",
        "Causal_TF" : "no",
        "Root_Descriptions" : "unknown protein; BEST Arabidopsis thaliana protein match is: unknown protein (TAIR:AT4G22758.1); Has 131 Blast hits to 131 proteins in 17 species: Archae - 0; Bacteria - 0; Metazoa - 0; Fungi - 0; Plants - 131; Viruses - 0; Other Eukaryotes - 0 (source: NCBI BLink).",
        "Eccentricity" : 10,
        "Outdegree" : 6,
        "Target_PLAMOM" : "",
        "PLAMOM_Match_Root" : "YES",
        "shared_name" : "AT2G27830",
        "Indegree" : 0,
        "Target_tRNA" : "",
        "t_RNA_motif_shoot" : "",
        "SUID" : 1379,
        "SelfLoops" : 0,
        "Target_TF" : "",
        "MolecularAttributes" : "protein_coding",
        "AverageShortestPathLength" : 5.37160752,
        "selected" : false,
        "Name2" : "AT2G27830",
        "ClosenessCentrality" : 0.18616401,
        "Casual_Name" : "unknown protein; BEST Arabidopsis thaliana protein match is: unknown protein (TAIR:AT4G22758.1); Has 131 Blast hits to 131 proteins in 17 species: Archae - 0; Bacteria - 0; Metazoa - 0; Fungi - 0; Plants - 131; Viruses - 0; Other Eukaryotes - 0 (source: NCBI BLink).",
        "PartnerOfMultiEdgedNodePairs" : 0,
        "ClusteringCoefficient" : 0.0,
        "Target_Name" : "",
        "Stress" : 0,
        "BetweennessCentrality" : 0.0,
        "t_RNA_motif_Root" : "NO",
        "EdgeCount" : 6,
        "Causal_PLAMOM" : "AT2G27830",
        "name" : "AT2G27830",
        "Causal_tRNA" : "no",
        "IsSingleNode" : false,
        "briefDescription" : "hypothetical protein",
        "NeighborhoodConnectivity" : 8.833333333333334
      },
      "position" : {
        "x" : -6117.1142578125,
        "y" : -97.63868713378906
      },
      "selected" : false
    }, {
      "data" : {
        "id" : "1378",
        "symbol" : "AT1G02870",
        "degree_layout" : 1,
        "Shoot_Descriptions" : "FUNCTIONS IN: molecular_function unknown; INVOLVED IN: biological_process unknown; EXPRESSED IN: 23 plant structures; EXPRESSED DURING: 13 growth stages; CONTAINS InterPro DOMAIN/s: Ribosome biogenesis protein Nop16 (InterPro:IPR019002); Has 104 Blast hits to 104 proteins in 57 species: Archae - 0; Bacteria - 0; Metazoa - 0; Fungi - 63; Plants - 35; Viruses - 0; Other Eukaryotes - 6 (source: NCBI BLink).",
        "PLAMOM_Match_Shoot" : "NO",
        "Causal_TF" : "",
        "Root_Descriptions" : "",
        "Eccentricity" : 0,
        "Outdegree" : 0,
        "Target_PLAMOM" : "no",
        "PLAMOM_Match_Root" : "",
        "shared_name" : "AT1G02870",
        "Indegree" : 2,
        "Target_tRNA" : "no",
        "t_RNA_motif_shoot" : "NO",
        "SUID" : 1378,
        "SelfLoops" : 0,
        "Target_TF" : "FUNCTIONS IN: molecular_function unknown; INVOLVED IN: biological_process unknown; EXPRESSED IN: 23 plant structures; EXPRESSED DURING: 13 growth stages; CONTAINS InterPro DOMAIN/s: Ribosome biogenesis protein Nop16 (InterPro:IPR019002); Has 104 Blast hits to 104 proteins in 57 species: Archae - 0; Bacteria - 0; Metazoa - 0; Fungi - 63; Plants - 35; Viruses - 0; Other Eukaryotes - 6 (source: NCBI BLink).",
        "MolecularAttributes" : "protein_coding",
        "AverageShortestPathLength" : 0.0,
        "selected" : false,
        "Name2" : "AT1G02870",
        "ClosenessCentrality" : 0.0,
        "Casual_Name" : "",
        "PartnerOfMultiEdgedNodePairs" : 0,
        "ClusteringCoefficient" : 0.0,
        "Target_Name" : "no",
        "Stress" : 0,
        "BetweennessCentrality" : 0.0,
        "t_RNA_motif_Root" : "",
        "EdgeCount" : 2,
        "Causal_PLAMOM" : "",
        "name" : "AT1G02870",
        "Causal_tRNA" : "",
        "IsSingleNode" : false,
        "briefDescription" : "nucleolar-like protein",
        "NeighborhoodConnectivity" : 24.0
      },
      "position" : {
        "x" : 3904.009960580046,
        "y" : 28.708532333374023
      },
      "selected" : false
    }, {
      "data" : {
        "id" : "1377",
        "symbol" : "BT1",
        "degree_layout" : 29,
        "Shoot_Descriptions" : "BT1, BTB and TAZ domain protein 1",
        "PLAMOM_Match_Shoot" : "YES",
        "Causal_TF" : "BTB and TAZ domain protein 1\"",
        "Root_Descriptions" : "BT1, BTB and TAZ domain protein 1",
        "Eccentricity" : 8,
        "Outdegree" : 27,
        "Target_PLAMOM" : "no",
        "PLAMOM_Match_Root" : "YES",
        "shared_name" : "AT5G63160",
        "Indegree" : 15,
        "Target_tRNA" : "AT5G63160",
        "t_RNA_motif_shoot" : "NO",
        "SUID" : 1377,
        "SelfLoops" : 0,
        "Target_TF" : "\"BT1",
        "MolecularAttributes" : "protein_coding",
        "AverageShortestPathLength" : 3.74476987,
        "selected" : false,
        "Name2" : "BT1",
        "ClosenessCentrality" : 0.26703911,
        "Casual_Name" : "\"BT1",
        "PartnerOfMultiEdgedNodePairs" : 0,
        "ClusteringCoefficient" : 0.00813008,
        "Target_Name" : "no",
        "Stress" : 156779,
        "BetweennessCentrality" : 0.02979059,
        "t_RNA_motif_Root" : "NO",
        "EdgeCount" : 42,
        "Causal_PLAMOM" : "AT5G63160",
        "name" : "AT5G63160",
        "Causal_tRNA" : "no",
        "IsSingleNode" : false,
        "briefDescription" : "BTB and TAZ domain protein 1",
        "NeighborhoodConnectivity" : 15.976190476190476
      },
      "position" : {
        "x" : -1126.878711294954,
        "y" : 3095.7841796875
      },
      "selected" : false
    }, {
      "data" : {
        "id" : "1376",
        "symbol" : "AT3G07340",
        "Shoot_Descriptions" : "",
        "PLAMOM_Match_Shoot" : "",
        "Causal_TF" : "AT3G07340",
        "Root_Descriptions" : "basic helix-loop-helix (bHLH) DNA-binding superfamily protein",
        "Eccentricity" : 8,
        "Outdegree" : 19,
        "Target_PLAMOM" : "no",
        "PLAMOM_Match_Root" : "NO",
        "shared_name" : "AT3G07340",
        "Indegree" : 2,
        "Target_tRNA" : "no",
        "t_RNA_motif_shoot" : "",
        "SUID" : 1376,
        "SelfLoops" : 0,
        "Target_TF" : "AT3G07340",
        "MolecularAttributes" : "TXNFACTOR",
        "AverageShortestPathLength" : 3.958159,
        "selected" : false,
        "Name2" : "AT3G07340",
        "ClosenessCentrality" : 0.25264271,
        "Casual_Name" : "basic helix-loop-helix (bHLH) DNA-binding superfamily protein",
        "PartnerOfMultiEdgedNodePairs" : 0,
        "ClusteringCoefficient" : 0.0,
        "Target_Name" : "basic helix-loop-helix (bHLH) DNA-binding superfamily protein",
        "Stress" : 21163,
        "BetweennessCentrality" : 0.00397224,
        "t_RNA_motif_Root" : "NO",
        "EdgeCount" : 21,
        "Causal_PLAMOM" : "no",
        "name" : "AT3G07340",
        "Causal_tRNA" : "no",
        "IsSingleNode" : false,
        "briefDescription" : "basic helix-loop-helix (bHLH) DNA-binding superfamily protein",
        "NeighborhoodConnectivity" : 13.952380952380953
      },
      "position" : {
        "x" : -6561.47265625,
        "y" : 2912.824951171875
      },
      "selected" : false
    }, {
      "data" : {
        "id" : "1375",
        "symbol" : "AT1G03220",
        "degree_layout" : 2,
        "Shoot_Descriptions" : "Eukaryotic aspartyl protease family protein",
        "PLAMOM_Match_Shoot" : "YES",
        "Causal_TF" : "no",
        "Root_Descriptions" : "",
        "Eccentricity" : 11,
        "Outdegree" : 1,
        "Target_PLAMOM" : "AT1G03220",
        "PLAMOM_Match_Root" : "",
        "shared_name" : "AT1G03220",
        "Indegree" : 6,
        "Target_tRNA" : "no",
        "t_RNA_motif_shoot" : "NO",
        "SUID" : 1375,
        "SelfLoops" : 0,
        "Target_TF" : "no",
        "MolecularAttributes" : "protein_coding",
        "AverageShortestPathLength" : 6.70920502,
        "selected" : false,
        "Name2" : "AT1G03220",
        "ClosenessCentrality" : 0.14904896,
        "Casual_Name" : "Eukaryotic aspartyl protease family protein",
        "PartnerOfMultiEdgedNodePairs" : 0,
        "ClusteringCoefficient" : 0.0,
        "Target_Name" : "Eukaryotic aspartyl protease family protein",
        "Stress" : 2338,
        "BetweennessCentrality" : 4.0181E-4,
        "t_RNA_motif_Root" : "",
        "EdgeCount" : 7,
        "Causal_PLAMOM" : "AT1G03220",
        "name" : "AT1G03220",
        "Causal_tRNA" : "no",
        "IsSingleNode" : false,
        "briefDescription" : "Eukaryotic aspartyl protease family protein",
        "NeighborhoodConnectivity" : 19.285714285714285
      },
      "position" : {
        "x" : 3345.680126595671,
        "y" : 2817.074462890625
      },
      "selected" : false
    }, {
      "data" : {
        "id" : "1374",
        "symbol" : "UCP5",
        "Shoot_Descriptions" : "",
        "PLAMOM_Match_Shoot" : "",
        "Causal_TF" : "DIC1",
        "Root_Descriptions" : "ATPUMP5, DIC1, UCP5, uncoupling protein 5",
        "Eccentricity" : 10,
        "Outdegree" : 9,
        "Target_PLAMOM" : "AT2G22500",
        "PLAMOM_Match_Root" : "YES",
        "shared_name" : "AT2G22500",
        "Indegree" : 1,
        "Target_tRNA" : "no",
        "t_RNA_motif_shoot" : "",
        "SUID" : 1374,
        "SelfLoops" : 0,
        "Target_TF" : "DIC1",
        "MolecularAttributes" : "protein_coding",
        "AverageShortestPathLength" : 5.20876827,
        "selected" : false,
        "Name2" : "UCP5",
        "ClosenessCentrality" : 0.19198397,
        "Casual_Name" : "\"ATPUMP5",
        "PartnerOfMultiEdgedNodePairs" : 0,
        "ClusteringCoefficient" : 0.0,
        "Target_Name" : "\"ATPUMP5",
        "Stress" : 264,
        "BetweennessCentrality" : 2.261E-5,
        "t_RNA_motif_Root" : "NO",
        "EdgeCount" : 10,
        "Causal_PLAMOM" : "AT2G22500",
        "name" : "AT2G22500",
        "Causal_tRNA" : "no",
        "IsSingleNode" : false,
        "briefDescription" : "uncoupling protein 5",
        "NeighborhoodConnectivity" : 10.2
      },
      "position" : {
        "x" : -6581.728515625,
        "y" : 2684.008056640625
      },
      "selected" : false
    }, {
      "data" : {
        "id" : "1373",
        "symbol" : "AT5G57655",
        "Shoot_Descriptions" : "",
        "PLAMOM_Match_Shoot" : "",
        "Causal_TF" : "no",
        "Root_Descriptions" : "xylose isomerase family protein",
        "Eccentricity" : 10,
        "Outdegree" : 10,
        "Target_PLAMOM" : "AT5G57655",
        "PLAMOM_Match_Root" : "YES",
        "shared_name" : "AT5G57655",
        "Indegree" : 7,
        "Target_tRNA" : "no",
        "t_RNA_motif_shoot" : "",
        "SUID" : 1373,
        "SelfLoops" : 0,
        "Target_TF" : "no",
        "MolecularAttributes" : "METABOLIC",
        "AverageShortestPathLength" : 5.4790795,
        "selected" : false,
        "Name2" : "AT5G57655",
        "ClosenessCentrality" : 0.18251241,
        "Casual_Name" : "xylose isomerase family protein",
        "PartnerOfMultiEdgedNodePairs" : 0,
        "ClusteringCoefficient" : 0.00367647,
        "Target_Name" : "xylose isomerase family protein",
        "Stress" : 13739,
        "BetweennessCentrality" : 0.00240115,
        "t_RNA_motif_Root" : "NO",
        "EdgeCount" : 17,
        "Causal_PLAMOM" : "AT5G57655",
        "name" : "AT5G57655",
        "Causal_tRNA" : "no",
        "IsSingleNode" : false,
        "briefDescription" : "xylose isomerase family protein",
        "NeighborhoodConnectivity" : 12.176470588235293
      },
      "position" : {
        "x" : -6675.962890625,
        "y" : 2817.814697265625
      },
      "selected" : false
    }, {
      "data" : {
        "id" : "1372",
        "symbol" : "NET4A",
        "Shoot_Descriptions" : "Kinase interacting (KIP1-like) family protein",
        "PLAMOM_Match_Shoot" : "NO",
        "Causal_TF" : "no",
        "Root_Descriptions" : "Kinase interacting (KIP1-like) family protein",
        "Eccentricity" : 10,
        "Outdegree" : 9,
        "Target_PLAMOM" : "no",
        "PLAMOM_Match_Root" : "NO",
        "shared_name" : "AT5G58320",
        "Indegree" : 2,
        "Target_tRNA" : "no",
        "t_RNA_motif_shoot" : "NO",
        "SUID" : 1372,
        "SelfLoops" : 0,
        "Target_TF" : "Kinase interacting (KIP1-like) family protein",
        "MolecularAttributes" : "protein_coding",
        "AverageShortestPathLength" : 4.9539749,
        "selected" : false,
        "Name2" : "NET4A",
        "ClosenessCentrality" : 0.20185811,
        "Casual_Name" : "Kinase interacting (KIP1-like) family protein",
        "PartnerOfMultiEdgedNodePairs" : 0,
        "ClusteringCoefficient" : 0.00909091,
        "Target_Name" : "no",
        "Stress" : 8825,
        "BetweennessCentrality" : 0.00184708,
        "t_RNA_motif_Root" : "NO",
        "EdgeCount" : 11,
        "Causal_PLAMOM" : "no",
        "name" : "AT5G58320",
        "Causal_tRNA" : "no",
        "IsSingleNode" : false,
        "briefDescription" : "Kinase interacting (KIP1-like) family protein",
        "NeighborhoodConnectivity" : 15.090909090909092
      },
      "position" : {
        "x" : -1177.71240234375,
        "y" : 2797.2209368377926
      },
      "selected" : false
    }, {
      "data" : {
        "id" : "1371",
        "symbol" : "AT5G66520",
        "Shoot_Descriptions" : "",
        "PLAMOM_Match_Shoot" : "",
        "Causal_TF" : "no",
        "Root_Descriptions" : "Tetratricopeptide repeat (TPR)-like superfamily protein",
        "Eccentricity" : 12,
        "Outdegree" : 1,
        "Target_PLAMOM" : "",
        "PLAMOM_Match_Root" : "NO",
        "shared_name" : "AT5G66520",
        "Indegree" : 0,
        "Target_tRNA" : "",
        "t_RNA_motif_shoot" : "",
        "SUID" : 1371,
        "SelfLoops" : 0,
        "Target_TF" : "",
        "MolecularAttributes" : "protein_coding",
        "AverageShortestPathLength" : 7.69519833,
        "selected" : false,
        "Name2" : "AT5G66520",
        "ClosenessCentrality" : 0.12995117,
        "Casual_Name" : "Tetratricopeptide repeat (TPR)-like superfamily protein",
        "PartnerOfMultiEdgedNodePairs" : 0,
        "ClusteringCoefficient" : 0.0,
        "Target_Name" : "",
        "Stress" : 0,
        "BetweennessCentrality" : 0.0,
        "t_RNA_motif_Root" : "NO",
        "EdgeCount" : 1,
        "Causal_PLAMOM" : "no",
        "name" : "AT5G66520",
        "Causal_tRNA" : "no",
        "IsSingleNode" : false,
        "briefDescription" : "Tetratricopeptide repeat (TPR)-like superfamily protein",
        "NeighborhoodConnectivity" : 7.0
      },
      "position" : {
        "x" : -6368.60546875,
        "y" : -547.7074584960938
      },
      "selected" : false
    }, {
      "data" : {
        "id" : "1370",
        "symbol" : "GATA17",
        "Shoot_Descriptions" : "",
        "PLAMOM_Match_Shoot" : "",
        "Causal_TF" : "GATA transcription factor 17\"",
        "Root_Descriptions" : "GATA17, GATA transcription factor 17",
        "Eccentricity" : 10,
        "Outdegree" : 10,
        "Target_PLAMOM" : "no",
        "PLAMOM_Match_Root" : "NO",
        "shared_name" : "AT3G16870",
        "Indegree" : 1,
        "Target_tRNA" : "no",
        "t_RNA_motif_shoot" : "",
        "SUID" : 1370,
        "SelfLoops" : 0,
        "Target_TF" : "GATA transcription factor 17\"",
        "MolecularAttributes" : "TXNFACTOR",
        "AverageShortestPathLength" : 5.06004141,
        "selected" : false,
        "Name2" : "GATA17",
        "ClosenessCentrality" : 0.19762684,
        "Casual_Name" : "\"GATA17",
        "PartnerOfMultiEdgedNodePairs" : 0,
        "ClusteringCoefficient" : 0.0,
        "Target_Name" : "\"GATA17",
        "Stress" : 767,
        "BetweennessCentrality" : 2.096E-4,
        "t_RNA_motif_Root" : "NO",
        "EdgeCount" : 11,
        "Causal_PLAMOM" : "no",
        "name" : "AT3G16870",
        "Causal_tRNA" : "no",
        "IsSingleNode" : false,
        "briefDescription" : "GATA transcription factor 17",
        "NeighborhoodConnectivity" : 9.636363636363637
      },
      "position" : {
        "x" : -6645.5390625,
        "y" : 2959.201904296875
      },
      "selected" : false
    }, {
      "data" : {
        "id" : "1369",
        "symbol" : "AT1G03230",
        "degree_layout" : 2,
        "Shoot_Descriptions" : "Eukaryotic aspartyl protease family protein",
        "PLAMOM_Match_Shoot" : "YES",
        "Causal_TF" : "",
        "Root_Descriptions" : "",
        "Eccentricity" : 0,
        "Outdegree" : 0,
        "Target_PLAMOM" : "family 707",
        "PLAMOM_Match_Root" : "",
        "shared_name" : "AT1G03230",
        "Indegree" : 7,
        "Target_tRNA" : "subfamily A",
        "t_RNA_motif_shoot" : "YES",
        "SUID" : 1369,
        "SelfLoops" : 0,
        "Target_TF" : "no",
        "MolecularAttributes" : "protein_coding",
        "AverageShortestPathLength" : 0.0,
        "selected" : false,
        "Name2" : "AT1G03230",
        "ClosenessCentrality" : 0.0,
        "Casual_Name" : "",
        "PartnerOfMultiEdgedNodePairs" : 0,
        "ClusteringCoefficient" : 0.0,
        "Target_Name" : "polypeptide 3\"",
        "Stress" : 0,
        "BetweennessCentrality" : 0.0,
        "t_RNA_motif_Root" : "",
        "EdgeCount" : 7,
        "Causal_PLAMOM" : "",
        "name" : "AT1G03230",
        "Causal_tRNA" : "",
        "IsSingleNode" : false,
        "briefDescription" : "Eukaryotic aspartyl protease family protein",
        "NeighborhoodConnectivity" : 16.857142857142858
      },
      "position" : {
        "x" : 4018.353954720671,
        "y" : 99.90956115722656
      },
      "selected" : false
    }, {
      "data" : {
        "id" : "1368",
        "symbol" : "AT1G08400",
        "degree_layout" : 18,
        "Shoot_Descriptions" : "",
        "PLAMOM_Match_Shoot" : "",
        "Causal_TF" : "no",
        "Root_Descriptions" : "RINT-1 / TIP-1 family",
        "Eccentricity" : 9,
        "Outdegree" : 16,
        "Target_PLAMOM" : "no",
        "PLAMOM_Match_Root" : "NO",
        "shared_name" : "AT1G08400",
        "Indegree" : 19,
        "Target_tRNA" : "AT1G08400",
        "t_RNA_motif_shoot" : "",
        "SUID" : 1368,
        "SelfLoops" : 0,
        "Target_TF" : "no",
        "MolecularAttributes" : "protein_coding",
        "AverageShortestPathLength" : 4.39121339,
        "selected" : false,
        "Name2" : "AT1G08400",
        "ClosenessCentrality" : 0.22772749,
        "Casual_Name" : "RINT-1 / TIP-1 family",
        "PartnerOfMultiEdgedNodePairs" : 0,
        "ClusteringCoefficient" : 0.0,
        "Target_Name" : "RINT-1 / TIP-1 family",
        "Stress" : 179652,
        "BetweennessCentrality" : 0.02962492,
        "t_RNA_motif_Root" : "YES",
        "EdgeCount" : 35,
        "Causal_PLAMOM" : "no",
        "name" : "AT1G08400",
        "Causal_tRNA" : "AT1G08400",
        "IsSingleNode" : false,
        "briefDescription" : "RINT-1 / TIP-1 family",
        "NeighborhoodConnectivity" : 12.17142857142857
      },
      "position" : {
        "x" : -6515.3505859375,
        "y" : 2946.300048828125
      },
      "selected" : false
    }, {
      "data" : {
        "id" : "1367",
        "symbol" : "LBD37",
        "Shoot_Descriptions" : "",
        "PLAMOM_Match_Shoot" : "",
        "Causal_TF" : "LBD37",
        "Root_Descriptions" : "ASL39, LBD37, LOB domain-containing protein 37",
        "Eccentricity" : 10,
        "Outdegree" : 4,
        "Target_PLAMOM" : "no",
        "PLAMOM_Match_Root" : "NO",
        "shared_name" : "AT5G67420",
        "Indegree" : 2,
        "Target_tRNA" : "no",
        "t_RNA_motif_shoot" : "",
        "SUID" : 1367,
        "SelfLoops" : 0,
        "Target_TF" : "LBD37",
        "MolecularAttributes" : "protein_coding",
        "AverageShortestPathLength" : 5.36610879,
        "selected" : false,
        "Name2" : "LBD37",
        "ClosenessCentrality" : 0.18635478,
        "Casual_Name" : "\"ASL39",
        "PartnerOfMultiEdgedNodePairs" : 0,
        "ClusteringCoefficient" : 0.0,
        "Target_Name" : "\"ASL39",
        "Stress" : 3676,
        "BetweennessCentrality" : 6.74E-4,
        "t_RNA_motif_Root" : "NO",
        "EdgeCount" : 6,
        "Causal_PLAMOM" : "no",
        "name" : "AT5G67420",
        "Causal_tRNA" : "no",
        "IsSingleNode" : false,
        "briefDescription" : "LOB domain-containing protein 37",
        "NeighborhoodConnectivity" : 16.333333333333332
      },
      "position" : {
        "x" : -6239.56591796875,
        "y" : 2994.921630859375
      },
      "selected" : false
    }, {
      "data" : {
        "id" : "1366",
        "symbol" : "AT2G30600",
        "Shoot_Descriptions" : "",
        "PLAMOM_Match_Shoot" : "",
        "Causal_TF" : "no",
        "Root_Descriptions" : "BTB/POZ domain-containing protein",
        "Eccentricity" : 10,
        "Outdegree" : 6,
        "Target_PLAMOM" : "",
        "PLAMOM_Match_Root" : "YES",
        "shared_name" : "AT2G30600",
        "Indegree" : 0,
        "Target_tRNA" : "",
        "t_RNA_motif_shoot" : "",
        "SUID" : 1366,
        "SelfLoops" : 0,
        "Target_TF" : "",
        "MolecularAttributes" : "protein_coding",
        "AverageShortestPathLength" : 6.08768267,
        "selected" : false,
        "Name2" : "AT2G30600",
        "ClosenessCentrality" : 0.16426612,
        "Casual_Name" : "BTB/POZ domain-containing protein",
        "PartnerOfMultiEdgedNodePairs" : 0,
        "ClusteringCoefficient" : 0.0,
        "Target_Name" : "",
        "Stress" : 0,
        "BetweennessCentrality" : 0.0,
        "t_RNA_motif_Root" : "NO",
        "EdgeCount" : 6,
        "Causal_PLAMOM" : "AT2G30600",
        "name" : "AT2G30600",
        "Causal_tRNA" : "no",
        "IsSingleNode" : false,
        "briefDescription" : "BTB/POZ domain-containing protein",
        "NeighborhoodConnectivity" : 10.666666666666666
      },
      "position" : {
        "x" : -6157.41845703125,
        "y" : 15.033241271972656
      },
      "selected" : false
    }, {
      "data" : {
        "id" : "1365",
        "symbol" : "AT5G05520",
        "Shoot_Descriptions" : "Outer membrane OMP85 family protein",
        "PLAMOM_Match_Shoot" : "NO",
        "Causal_TF" : "no",
        "Root_Descriptions" : "Outer membrane OMP85 family protein",
        "Eccentricity" : 12,
        "Outdegree" : 11,
        "Target_PLAMOM" : "",
        "PLAMOM_Match_Root" : "NO",
        "shared_name" : "AT5G05520",
        "Indegree" : 0,
        "Target_tRNA" : "",
        "t_RNA_motif_shoot" : "NO",
        "SUID" : 1365,
        "SelfLoops" : 0,
        "Target_TF" : "",
        "MolecularAttributes" : "protein_coding",
        "AverageShortestPathLength" : 6.76763485,
        "selected" : false,
        "Name2" : "AT5G05520",
        "ClosenessCentrality" : 0.14776211,
        "Casual_Name" : "Outer membrane OMP85 family protein",
        "PartnerOfMultiEdgedNodePairs" : 0,
        "ClusteringCoefficient" : 0.0,
        "Target_Name" : "",
        "Stress" : 0,
        "BetweennessCentrality" : 0.0,
        "t_RNA_motif_Root" : "NO",
        "EdgeCount" : 11,
        "Causal_PLAMOM" : "no",
        "name" : "AT5G05520",
        "Causal_tRNA" : "no",
        "IsSingleNode" : false,
        "briefDescription" : "Outer membrane OMP85 family protein",
        "NeighborhoodConnectivity" : 9.545454545454545
      },
      "position" : {
        "x" : -1435.83251953125,
        "y" : 2893.8193255096676
      },
      "selected" : false
    }, {
      "data" : {
        "id" : "1364",
        "symbol" : "CYP707A3",
        "Shoot_Descriptions" : "",
        "PLAMOM_Match_Shoot" : "",
        "Causal_TF" : "cytochrome P450",
        "Root_Descriptions" : "CYP707A3, cytochrome P450, family 707, subfamily A, polypeptide 3",
        "Eccentricity" : 9,
        "Outdegree" : 7,
        "Target_PLAMOM" : "",
        "PLAMOM_Match_Root" : "NO",
        "shared_name" : "AT5G45340",
        "Indegree" : 0,
        "Target_tRNA" : "",
        "t_RNA_motif_shoot" : "",
        "SUID" : 1364,
        "SelfLoops" : 0,
        "Target_TF" : "",
        "MolecularAttributes" : "METABOLIC",
        "AverageShortestPathLength" : 4.83924843,
        "selected" : false,
        "Name2" : "CYP707A3",
        "ClosenessCentrality" : 0.20664366,
        "Casual_Name" : "\"CYP707A3",
        "PartnerOfMultiEdgedNodePairs" : 0,
        "ClusteringCoefficient" : 0.0,
        "Target_Name" : "",
        "Stress" : 0,
        "BetweennessCentrality" : 0.0,
        "t_RNA_motif_Root" : "NO",
        "EdgeCount" : 7,
        "Causal_PLAMOM" : "no",
        "name" : "AT5G45340",
        "Causal_tRNA" : "no",
        "IsSingleNode" : false,
        "briefDescription" : "cytochrome P450",
        "NeighborhoodConnectivity" : 12.142857142857142
      },
      "position" : {
        "x" : -6034.4248046875,
        "y" : 415.1164245605469
      },
      "selected" : false
    }, {
      "data" : {
        "id" : "1363",
        "symbol" : "AT3G45210",
        "Shoot_Descriptions" : "Protein of unknown function, DUF584",
        "PLAMOM_Match_Shoot" : "NO",
        "Causal_TF" : "DUF584\"",
        "Root_Descriptions" : "Protein of unknown function, DUF584",
        "Eccentricity" : 9,
        "Outdegree" : 6,
        "Target_PLAMOM" : "no",
        "PLAMOM_Match_Root" : "NO",
        "shared_name" : "AT3G45210",
        "Indegree" : 4,
        "Target_tRNA" : "no",
        "t_RNA_motif_shoot" : "NO",
        "SUID" : 1363,
        "SelfLoops" : 0,
        "Target_TF" : "DUF584\"",
        "MolecularAttributes" : "protein_coding",
        "AverageShortestPathLength" : 4.89958159,
        "selected" : false,
        "Name2" : "AT3G45210",
        "ClosenessCentrality" : 0.20409906,
        "Casual_Name" : "\"Protein of unknown function",
        "PartnerOfMultiEdgedNodePairs" : 0,
        "ClusteringCoefficient" : 0.04444444,
        "Target_Name" : "\"Protein of unknown function",
        "Stress" : 11489,
        "BetweennessCentrality" : 0.00178629,
        "t_RNA_motif_Root" : "NO",
        "EdgeCount" : 10,
        "Causal_PLAMOM" : "no",
        "name" : "AT3G45210",
        "Causal_tRNA" : "no",
        "IsSingleNode" : false,
        "briefDescription" : "transcription initiation factor TFIID subunit (Protein of unknown function",
        "NeighborhoodConnectivity" : 18.6
      },
      "position" : {
        "x" : -830.3681522129227,
        "y" : 3424.896728515625
      },
      "selected" : false
    }, {
      "data" : {
        "id" : "1362",
        "symbol" : "AT1G03560",
        "Shoot_Descriptions" : "Pentatricopeptide repeat (PPR-like) superfamily protein",
        "PLAMOM_Match_Shoot" : "NO",
        "Causal_TF" : "no",
        "Root_Descriptions" : "",
        "Eccentricity" : 9,
        "Outdegree" : 3,
        "Target_PLAMOM" : "family 707",
        "PLAMOM_Match_Root" : "",
        "shared_name" : "AT1G03560",
        "Indegree" : 3,
        "Target_tRNA" : "subfamily A",
        "t_RNA_motif_shoot" : "YES",
        "SUID" : 1362,
        "SelfLoops" : 0,
        "Target_TF" : "no",
        "MolecularAttributes" : "protein_coding",
        "AverageShortestPathLength" : 5.41004184,
        "selected" : false,
        "Name2" : "AT1G03560",
        "ClosenessCentrality" : 0.18484145,
        "Casual_Name" : "Pentatricopeptide repeat (PPR-like) superfamily protein",
        "PartnerOfMultiEdgedNodePairs" : 0,
        "ClusteringCoefficient" : 0.06666667,
        "Target_Name" : "polypeptide 3\"",
        "Stress" : 33321,
        "BetweennessCentrality" : 0.00622528,
        "t_RNA_motif_Root" : "",
        "EdgeCount" : 6,
        "Causal_PLAMOM" : "no",
        "name" : "AT1G03560",
        "Causal_tRNA" : "AT1G03560",
        "IsSingleNode" : false,
        "briefDescription" : "Pentatricopeptide repeat (PPR-like) superfamily protein",
        "NeighborhoodConnectivity" : 12.5
      },
      "position" : {
        "x" : 3755.607616830046,
        "y" : 3509.501220703125
      },
      "selected" : false
    }, {
      "data" : {
        "id" : "1361",
        "symbol" : "AT5G10210",
        "Shoot_Descriptions" : "CONTAINS InterPro DOMAIN/s: C2 calcium-dependent membrane targeting (InterPro:IPR000008); BEST Arabidopsis thaliana protein match is: unknown protein (TAIR:AT5G65030.1); Has 1807 Blast hits to 1807 proteins in 277 species: Archae - 0; Bacteria - 0; Metazoa - 736; Fungi - 347; Plants - 385; Viruses - 0; Other Eukaryotes - 339 (source: NCBI BLink).",
        "PLAMOM_Match_Shoot" : "NO",
        "Causal_TF" : "no",
        "Root_Descriptions" : "CONTAINS InterPro DOMAIN/s: C2 calcium-dependent membrane targeting (InterPro:IPR000008); BEST Arabidopsis thaliana protein match is: unknown protein (TAIR:AT5G65030.1); Has 1807 Blast hits to 1807 proteins in 277 species: Archae - 0; Bacteria - 0; Metazoa - 736; Fungi - 347; Plants - 385; Viruses - 0; Other Eukaryotes - 339 (source: NCBI BLink).",
        "Eccentricity" : 9,
        "Outdegree" : 12,
        "Target_PLAMOM" : "no",
        "PLAMOM_Match_Root" : "NO",
        "shared_name" : "AT5G10210",
        "Indegree" : 8,
        "Target_tRNA" : "no",
        "t_RNA_motif_shoot" : "NO",
        "SUID" : 1361,
        "SelfLoops" : 0,
        "Target_TF" : "no",
        "MolecularAttributes" : "protein_coding",
        "AverageShortestPathLength" : 4.79079498,
        "selected" : false,
        "Name2" : "AT5G10210",
        "ClosenessCentrality" : 0.20873362,
        "Casual_Name" : "CONTAINS InterPro DOMAIN/s: C2 calcium-dependent membrane targeting (InterPro:IPR000008); BEST Arabidopsis thaliana protein match is: unknown protein (TAIR:AT5G65030.1); Has 1807 Blast hits to 1807 proteins in 277 species: Archae - 0; Bacteria - 0; Metazoa - 736; Fungi - 347; Plants - 385; Viruses - 0; Other Eukaryotes - 339 (source: NCBI BLink).",
        "PartnerOfMultiEdgedNodePairs" : 0,
        "ClusteringCoefficient" : 0.0,
        "Target_Name" : "CONTAINS InterPro DOMAIN/s: C2 calcium-dependent membrane targeting (InterPro:IPR000008); BEST Arabidopsis thaliana protein match is: unknown protein (TAIR:AT5G65030.1); Has 1807 Blast hits to 1807 proteins in 277 species: Archae - 0; Bacteria - 0; Metazoa - 736; Fungi - 347; Plants - 385; Viruses - 0; Other Eukaryotes - 339 (source: NCBI BLink).",
        "Stress" : 138319,
        "BetweennessCentrality" : 0.02903694,
        "t_RNA_motif_Root" : "NO",
        "EdgeCount" : 20,
        "Causal_PLAMOM" : "no",
        "name" : "AT5G10210",
        "Causal_tRNA" : "no",
        "IsSingleNode" : false,
        "briefDescription" : "nitric oxide synthase-interacting protein",
        "NeighborhoodConnectivity" : 9.15
      },
      "position" : {
        "x" : -1105.938281607454,
        "y" : 3352.41455078125
      },
      "selected" : false
    }, {
      "data" : {
        "id" : "1360",
        "symbol" : "AT2G38820",
        "Shoot_Descriptions" : "",
        "PLAMOM_Match_Shoot" : "",
        "Causal_TF" : "no",
        "Root_Descriptions" : "Protein of unknown function (DUF506)",
        "Eccentricity" : 8,
        "Outdegree" : 10,
        "Target_PLAMOM" : "NAC domain containing protein 96\"",
        "PLAMOM_Match_Root" : "NO",
        "shared_name" : "AT2G38820",
        "Indegree" : 4,
        "Target_tRNA" : "AT5G46590",
        "t_RNA_motif_shoot" : "",
        "SUID" : 1360,
        "SelfLoops" : 0,
        "Target_TF" : "no",
        "MolecularAttributes" : "protein_coding",
        "AverageShortestPathLength" : 4.76778243,
        "selected" : false,
        "Name2" : "AT2G38820",
        "ClosenessCentrality" : 0.20974111,
        "Casual_Name" : "Protein of unknown function (DUF506)",
        "PartnerOfMultiEdgedNodePairs" : 0,
        "ClusteringCoefficient" : 0.0,
        "Target_Name" : "no",
        "Stress" : 6709,
        "BetweennessCentrality" : 0.00127531,
        "t_RNA_motif_Root" : "NO",
        "EdgeCount" : 14,
        "Causal_PLAMOM" : "no",
        "name" : "AT2G38820",
        "Causal_tRNA" : "no",
        "IsSingleNode" : false,
        "briefDescription" : "DNA-directed RNA polymerase subunit beta-beta protein",
        "NeighborhoodConnectivity" : 10.0
      },
      "position" : {
        "x" : -6476.36181640625,
        "y" : 3197.174072265625
      },
      "selected" : false
    }, {
      "data" : {
        "id" : "1359",
        "symbol" : "AT1G05410",
        "Shoot_Descriptions" : "Protein of unknown function (DUF1423)",
        "PLAMOM_Match_Shoot" : "NO",
        "Causal_TF" : "",
        "Root_Descriptions" : "",
        "Eccentricity" : 0,
        "Outdegree" : 0,
        "Target_PLAMOM" : "no",
        "PLAMOM_Match_Root" : "",
        "shared_name" : "AT1G05410",
        "Indegree" : 1,
        "Target_tRNA" : "no",
        "t_RNA_motif_shoot" : "NO",
        "SUID" : 1359,
        "SelfLoops" : 0,
        "Target_TF" : "no",
        "MolecularAttributes" : "protein_coding",
        "AverageShortestPathLength" : 0.0,
        "selected" : false,
        "Name2" : "AT1G05410",
        "ClosenessCentrality" : 0.0,
        "Casual_Name" : "",
        "PartnerOfMultiEdgedNodePairs" : 0,
        "ClusteringCoefficient" : 0.0,
        "Target_Name" : "Protein of unknown function (DUF1423)",
        "Stress" : 0,
        "BetweennessCentrality" : 0.0,
        "t_RNA_motif_Root" : "",
        "EdgeCount" : 1,
        "Causal_PLAMOM" : "",
        "name" : "AT1G05410",
        "Causal_tRNA" : "",
        "IsSingleNode" : false,
        "briefDescription" : "CDPK adapter",
        "NeighborhoodConnectivity" : 14.0
      },
      "position" : {
        "x" : 4469.873974251921,
        "y" : 274.1626281738281
      },
      "selected" : false
    }, {
      "data" : {
        "id" : "1358",
        "symbol" : "AT2G36320",
        "Shoot_Descriptions" : "",
        "PLAMOM_Match_Shoot" : "",
        "Causal_TF" : "no",
        "Root_Descriptions" : "A20/AN1-like zinc finger family protein",
        "Eccentricity" : 10,
        "Outdegree" : 5,
        "Target_PLAMOM" : "AT2G36320",
        "PLAMOM_Match_Root" : "YES",
        "shared_name" : "AT2G36320",
        "Indegree" : 1,
        "Target_tRNA" : "no",
        "t_RNA_motif_shoot" : "",
        "SUID" : 1358,
        "SelfLoops" : 0,
        "Target_TF" : "no",
        "MolecularAttributes" : "protein_coding",
        "AverageShortestPathLength" : 6.32572614,
        "selected" : false,
        "Name2" : "AT2G36320",
        "ClosenessCentrality" : 0.15808462,
        "Casual_Name" : "A20/AN1-like zinc finger family protein",
        "PartnerOfMultiEdgedNodePairs" : 0,
        "ClusteringCoefficient" : 0.0,
        "Target_Name" : "A20/AN1-like zinc finger family protein",
        "Stress" : 40,
        "BetweennessCentrality" : 1.91E-5,
        "t_RNA_motif_Root" : "NO",
        "EdgeCount" : 6,
        "Causal_PLAMOM" : "AT2G36320",
        "name" : "AT2G36320",
        "Causal_tRNA" : "no",
        "IsSingleNode" : false,
        "briefDescription" : "A20/AN1-like zinc finger family protein",
        "NeighborhoodConnectivity" : 3.6666666666666665
      },
      "position" : {
        "x" : -7134.0234375,
        "y" : 3019.769287109375
      },
      "selected" : false
    }, {
      "data" : {
        "id" : "1357",
        "symbol" : "MEE5",
        "Shoot_Descriptions" : "CLO, GFA1, MEE5, Ribosomal protein S5/Elongation factor G/III/V family protein",
        "PLAMOM_Match_Shoot" : "YES",
        "Causal_TF" : "",
        "Root_Descriptions" : "",
        "Eccentricity" : 0,
        "Outdegree" : 0,
        "Target_PLAMOM" : "AT1G06220",
        "PLAMOM_Match_Root" : "",
        "shared_name" : "AT1G06220",
        "Indegree" : 2,
        "Target_tRNA" : "AT1G06220",
        "t_RNA_motif_shoot" : "YES",
        "SUID" : 1357,
        "SelfLoops" : 0,
        "Target_TF" : "GFA1",
        "MolecularAttributes" : "protein_coding",
        "AverageShortestPathLength" : 0.0,
        "selected" : false,
        "Name2" : "MEE5",
        "ClosenessCentrality" : 0.0,
        "Casual_Name" : "",
        "PartnerOfMultiEdgedNodePairs" : 0,
        "ClusteringCoefficient" : 0.0,
        "Target_Name" : "\"CLO",
        "Stress" : 0,
        "BetweennessCentrality" : 0.0,
        "t_RNA_motif_Root" : "",
        "EdgeCount" : 2,
        "Causal_PLAMOM" : "",
        "name" : "AT1G06220",
        "Causal_tRNA" : "",
        "IsSingleNode" : false,
        "briefDescription" : "Ribosomal protein S5/Elongation factor G/III/V family protein",
        "NeighborhoodConnectivity" : 6.0
      },
      "position" : {
        "x" : 3408.919384408171,
        "y" : -52.5946159362793
      },
      "selected" : false
    }, {
      "data" : {
        "id" : "1356",
        "symbol" : "AT3G11280",
        "Shoot_Descriptions" : "",
        "PLAMOM_Match_Shoot" : "",
        "Causal_TF" : "AT3G11280",
        "Root_Descriptions" : "Duplicated homeodomain-like superfamily protein",
        "Eccentricity" : 10,
        "Outdegree" : 5,
        "Target_PLAMOM" : "no",
        "PLAMOM_Match_Root" : "YES",
        "shared_name" : "AT3G11280",
        "Indegree" : 1,
        "Target_tRNA" : "AT3G11280",
        "t_RNA_motif_shoot" : "",
        "SUID" : 1356,
        "SelfLoops" : 0,
        "Target_TF" : "Duplicated homeodomain-like superfamily protein",
        "MolecularAttributes" : "TXNFACTOR",
        "AverageShortestPathLength" : 6.16666667,
        "selected" : false,
        "Name2" : "AT3G11280",
        "ClosenessCentrality" : 0.16216216,
        "Casual_Name" : "Duplicated homeodomain-like superfamily protein",
        "PartnerOfMultiEdgedNodePairs" : 0,
        "ClusteringCoefficient" : 0.0,
        "Target_Name" : "no",
        "Stress" : 735,
        "BetweennessCentrality" : 1.3318E-4,
        "t_RNA_motif_Root" : "NO",
        "EdgeCount" : 6,
        "Causal_PLAMOM" : "AT3G11280",
        "name" : "AT3G11280",
        "Causal_tRNA" : "no",
        "IsSingleNode" : false,
        "briefDescription" : "Duplicated homeodomain-like superfamily protein",
        "NeighborhoodConnectivity" : 4.833333333333333
      },
      "position" : {
        "x" : -6988.873046875,
        "y" : 3085.48828125
      },
      "selected" : false
    }, {
      "data" : {
        "id" : "1355",
        "symbol" : "ZFP4",
        "Shoot_Descriptions" : "",
        "PLAMOM_Match_Shoot" : "",
        "Causal_TF" : "zinc finger protein 4\"",
        "Root_Descriptions" : "ZFP4, zinc finger protein 4",
        "Eccentricity" : 9,
        "Outdegree" : 13,
        "Target_PLAMOM" : "no",
        "PLAMOM_Match_Root" : "NO",
        "shared_name" : "AT1G66140",
        "Indegree" : 1,
        "Target_tRNA" : "no",
        "t_RNA_motif_shoot" : "",
        "SUID" : 1355,
        "SelfLoops" : 0,
        "Target_TF" : "zinc finger protein 4\"",
        "MolecularAttributes" : "TXNFACTOR",
        "AverageShortestPathLength" : 4.41631799,
        "selected" : false,
        "Name2" : "ZFP4",
        "ClosenessCentrality" : 0.22643297,
        "Casual_Name" : "\"ZFP4",
        "PartnerOfMultiEdgedNodePairs" : 0,
        "ClusteringCoefficient" : 0.0,
        "Target_Name" : "\"ZFP4",
        "Stress" : 24257,
        "BetweennessCentrality" : 0.00465532,
        "t_RNA_motif_Root" : "NO",
        "EdgeCount" : 14,
        "Causal_PLAMOM" : "no",
        "name" : "AT1G66140",
        "Causal_tRNA" : "no",
        "IsSingleNode" : false,
        "briefDescription" : "zinc finger protein 4",
        "NeighborhoodConnectivity" : 18.357142857142858
      },
      "position" : {
        "x" : -6630.1552734375,
        "y" : 2996.445068359375
      },
      "selected" : false
    }, {
      "data" : {
        "id" : "1354",
        "symbol" : "AT1G07300",
        "degree_layout" : 1,
        "Shoot_Descriptions" : "josephin protein-related",
        "PLAMOM_Match_Shoot" : "YES",
        "Causal_TF" : "",
        "Root_Descriptions" : "",
        "Eccentricity" : 0,
        "Outdegree" : 0,
        "Target_PLAMOM" : "no",
        "PLAMOM_Match_Root" : "",
        "shared_name" : "AT1G07300",
        "Indegree" : 7,
        "Target_tRNA" : "AT1G07300",
        "t_RNA_motif_shoot" : "NO",
        "SUID" : 1354,
        "SelfLoops" : 0,
        "Target_TF" : "josephin protein-related",
        "MolecularAttributes" : "protein_coding",
        "AverageShortestPathLength" : 0.0,
        "selected" : false,
        "Name2" : "AT1G07300",
        "ClosenessCentrality" : 0.0,
        "Casual_Name" : "",
        "PartnerOfMultiEdgedNodePairs" : 0,
        "ClusteringCoefficient" : 0.0,
        "Target_Name" : "no",
        "Stress" : 0,
        "BetweennessCentrality" : 0.0,
        "t_RNA_motif_Root" : "",
        "EdgeCount" : 7,
        "Causal_PLAMOM" : "",
        "name" : "AT1G07300",
        "Causal_tRNA" : "",
        "IsSingleNode" : false,
        "briefDescription" : "josephin protein-like protein",
        "NeighborhoodConnectivity" : 19.428571428571427
      },
      "position" : {
        "x" : 4119.327099251921,
        "y" : 24.376256942749023
      },
      "selected" : false
    }, {
      "data" : {
        "id" : "1353",
        "symbol" : "AT5G04840",
        "Shoot_Descriptions" : "bZIP protein",
        "PLAMOM_Match_Shoot" : "NO",
        "Causal_TF" : "no",
        "Root_Descriptions" : "bZIP protein",
        "Eccentricity" : 8,
        "Outdegree" : 28,
        "Target_PLAMOM" : "phloem protein 2-A13\"",
        "PLAMOM_Match_Root" : "NO",
        "shared_name" : "AT5G04840",
        "Indegree" : 9,
        "Target_tRNA" : "no",
        "t_RNA_motif_shoot" : "NO",
        "SUID" : 1353,
        "SelfLoops" : 0,
        "Target_TF" : "no",
        "MolecularAttributes" : "protein_coding",
        "AverageShortestPathLength" : 3.75313808,
        "selected" : false,
        "Name2" : "AT5G04840",
        "ClosenessCentrality" : 0.2664437,
        "Casual_Name" : "bZIP protein",
        "PartnerOfMultiEdgedNodePairs" : 0,
        "ClusteringCoefficient" : 0.00375375,
        "Target_Name" : "no",
        "Stress" : 146681,
        "BetweennessCentrality" : 0.03402828,
        "t_RNA_motif_Root" : "NO",
        "EdgeCount" : 37,
        "Causal_PLAMOM" : "no",
        "name" : "AT5G04840",
        "Causal_tRNA" : "no",
        "IsSingleNode" : false,
        "briefDescription" : "bZIP protein",
        "NeighborhoodConnectivity" : 14.486486486486486
      },
      "position" : {
        "x" : -948.9677615879227,
        "y" : 3091.97216796875
      },
      "selected" : false
    }, {
      "data" : {
        "id" : "1352",
        "symbol" : "GDH1",
        "Shoot_Descriptions" : "",
        "PLAMOM_Match_Shoot" : "",
        "Causal_TF" : "glutamate dehydrogenase 1\"",
        "Root_Descriptions" : "GDH1, glutamate dehydrogenase 1",
        "Eccentricity" : 9,
        "Outdegree" : 8,
        "Target_PLAMOM" : "",
        "PLAMOM_Match_Root" : "YES",
        "shared_name" : "AT5G18170",
        "Indegree" : 0,
        "Target_tRNA" : "",
        "t_RNA_motif_shoot" : "",
        "SUID" : 1352,
        "SelfLoops" : 0,
        "Target_TF" : "",
        "MolecularAttributes" : "METABOLIC",
        "AverageShortestPathLength" : 4.71189979,
        "selected" : false,
        "Name2" : "GDH1",
        "ClosenessCentrality" : 0.21222862,
        "Casual_Name" : "\"GDH1",
        "PartnerOfMultiEdgedNodePairs" : 0,
        "ClusteringCoefficient" : 0.0,
        "Target_Name" : "",
        "Stress" : 0,
        "BetweennessCentrality" : 0.0,
        "t_RNA_motif_Root" : "NO",
        "EdgeCount" : 8,
        "Causal_PLAMOM" : "AT5G18170",
        "name" : "AT5G18170",
        "Causal_tRNA" : "no",
        "IsSingleNode" : false,
        "briefDescription" : "glutamate dehydrogenase 1",
        "NeighborhoodConnectivity" : 18.0
      },
      "position" : {
        "x" : -6245.49560546875,
        "y" : -89.09169006347656
      },
      "selected" : false
    }, {
      "data" : {
        "id" : "1351",
        "symbol" : "SNZ",
        "Shoot_Descriptions" : "",
        "PLAMOM_Match_Shoot" : "",
        "Causal_TF" : "Integrase-type DNA-binding superfamily protein\"",
        "Root_Descriptions" : "SNZ, Integrase-type DNA-binding superfamily protein",
        "Eccentricity" : 12,
        "Outdegree" : 5,
        "Target_PLAMOM" : "",
        "PLAMOM_Match_Root" : "NO",
        "shared_name" : "AT2G39250",
        "Indegree" : 0,
        "Target_tRNA" : "",
        "t_RNA_motif_shoot" : "",
        "SUID" : 1351,
        "SelfLoops" : 0,
        "Target_TF" : "",
        "MolecularAttributes" : "TXNFACTOR",
        "AverageShortestPathLength" : 7.1954262,
        "selected" : false,
        "Name2" : "SNZ",
        "ClosenessCentrality" : 0.13897717,
        "Casual_Name" : "\"SNZ",
        "PartnerOfMultiEdgedNodePairs" : 0,
        "ClusteringCoefficient" : 0.0,
        "Target_Name" : "",
        "Stress" : 0,
        "BetweennessCentrality" : 0.0,
        "t_RNA_motif_Root" : "NO",
        "EdgeCount" : 5,
        "Causal_PLAMOM" : "no",
        "name" : "AT2G39250",
        "Causal_tRNA" : "no",
        "IsSingleNode" : false,
        "briefDescription" : "Integrase-type DNA-binding superfamily protein",
        "NeighborhoodConnectivity" : 5.0
      },
      "position" : {
        "x" : -5831.8271484375,
        "y" : 555.9466552734375
      },
      "selected" : false
    }, {
      "data" : {
        "id" : "1350",
        "symbol" : "WRKY38",
        "Shoot_Descriptions" : "",
        "PLAMOM_Match_Shoot" : "",
        "Causal_TF" : "WRKY38",
        "Root_Descriptions" : "ATWRKY38, WRKY38, WRKY DNA-binding protein 38",
        "Eccentricity" : 8,
        "Outdegree" : 19,
        "Target_PLAMOM" : "",
        "PLAMOM_Match_Root" : "NO",
        "shared_name" : "AT5G22570",
        "Indegree" : 0,
        "Target_tRNA" : "",
        "t_RNA_motif_shoot" : "",
        "SUID" : 1350,
        "SelfLoops" : 0,
        "Target_TF" : "",
        "MolecularAttributes" : "TXNFACTOR",
        "AverageShortestPathLength" : 3.9125,
        "selected" : false,
        "Name2" : "WRKY38",
        "ClosenessCentrality" : 0.25559105,
        "Casual_Name" : "\"ATWRKY38",
        "PartnerOfMultiEdgedNodePairs" : 0,
        "ClusteringCoefficient" : 0.00292398,
        "Target_Name" : "",
        "Stress" : 0,
        "BetweennessCentrality" : 0.0,
        "t_RNA_motif_Root" : "NO",
        "EdgeCount" : 19,
        "Causal_PLAMOM" : "no",
        "name" : "AT5G22570",
        "Causal_tRNA" : "no",
        "IsSingleNode" : false,
        "briefDescription" : "WRKY DNA-binding protein 38",
        "NeighborhoodConnectivity" : 9.789473684210526
      },
      "position" : {
        "x" : -6042.14697265625,
        "y" : -84.26258850097656
      },
      "selected" : false
    }, {
      "data" : {
        "id" : "1349",
        "symbol" : "TLP9",
        "Shoot_Descriptions" : "",
        "PLAMOM_Match_Shoot" : "",
        "Causal_TF" : "TLP9",
        "Root_Descriptions" : "ATTLP9, TLP9, tubby-like protein 9",
        "Eccentricity" : 1,
        "Outdegree" : 3,
        "Target_PLAMOM" : "no",
        "PLAMOM_Match_Root" : "NO",
        "shared_name" : "AT3G06380",
        "Indegree" : 1,
        "Target_tRNA" : "no",
        "t_RNA_motif_shoot" : "",
        "SUID" : 1349,
        "SelfLoops" : 0,
        "Target_TF" : "TLP9",
        "MolecularAttributes" : "TXNFACTOR",
        "AverageShortestPathLength" : 1.0,
        "selected" : false,
        "Name2" : "TLP9",
        "ClosenessCentrality" : 1.0,
        "Casual_Name" : "\"ATTLP9",
        "PartnerOfMultiEdgedNodePairs" : 0,
        "ClusteringCoefficient" : 0.0,
        "Target_Name" : "\"ATTLP9",
        "Stress" : 3,
        "BetweennessCentrality" : 1.49E-6,
        "t_RNA_motif_Root" : "NO",
        "EdgeCount" : 4,
        "Causal_PLAMOM" : "no",
        "name" : "AT3G06380",
        "Causal_tRNA" : "no",
        "IsSingleNode" : false,
        "briefDescription" : "tubby-like protein 9",
        "NeighborhoodConnectivity" : 5.5
      },
      "position" : {
        "x" : -6764.3837890625,
        "y" : 2778.443359375
      },
      "selected" : false
    }, {
      "data" : {
        "id" : "1348",
        "symbol" : "OBP2",
        "Shoot_Descriptions" : "OBP2, Dof-type zinc finger DNA-binding family protein",
        "PLAMOM_Match_Shoot" : "NO",
        "Causal_TF" : "",
        "Root_Descriptions" : "",
        "Eccentricity" : 0,
        "Outdegree" : 0,
        "Target_PLAMOM" : "AT3G61850",
        "PLAMOM_Match_Root" : "",
        "shared_name" : "AT1G07640",
        "Indegree" : 3,
        "Target_tRNA" : "no",
        "t_RNA_motif_shoot" : "NO",
        "SUID" : 1348,
        "SelfLoops" : 0,
        "Target_TF" : "\"OBP2",
        "MolecularAttributes" : "TXNFACTOR",
        "AverageShortestPathLength" : 0.0,
        "selected" : false,
        "Name2" : "OBP2",
        "ClosenessCentrality" : 0.0,
        "Casual_Name" : "",
        "PartnerOfMultiEdgedNodePairs" : 0,
        "ClusteringCoefficient" : 0.0,
        "Target_Name" : "no",
        "Stress" : 0,
        "BetweennessCentrality" : 0.0,
        "t_RNA_motif_Root" : "",
        "EdgeCount" : 3,
        "Causal_PLAMOM" : "",
        "name" : "AT1G07640",
        "Causal_tRNA" : "",
        "IsSingleNode" : false,
        "briefDescription" : "Dof-type zinc finger DNA-binding family protein",
        "NeighborhoodConnectivity" : 4.666666666666667
      },
      "position" : {
        "x" : 4138.422314095671,
        "y" : -402.34332275390625
      },
      "selected" : false
    }, {
      "data" : {
        "id" : "1347",
        "symbol" : "AT1G20640",
        "Shoot_Descriptions" : "",
        "PLAMOM_Match_Shoot" : "",
        "Causal_TF" : "AT1G20640",
        "Root_Descriptions" : "Plant regulator RWP-RK family protein",
        "Eccentricity" : 1,
        "Outdegree" : 2,
        "Target_PLAMOM" : "",
        "PLAMOM_Match_Root" : "NO",
        "shared_name" : "AT1G20640",
        "Indegree" : 0,
        "Target_tRNA" : "",
        "t_RNA_motif_shoot" : "",
        "SUID" : 1347,
        "SelfLoops" : 0,
        "Target_TF" : "",
        "MolecularAttributes" : "TXNFACTOR",
        "AverageShortestPathLength" : 1.0,
        "selected" : false,
        "Name2" : "AT1G20640",
        "ClosenessCentrality" : 1.0,
        "Casual_Name" : "Plant regulator RWP-RK family protein",
        "PartnerOfMultiEdgedNodePairs" : 0,
        "ClusteringCoefficient" : 0.0,
        "Target_Name" : "",
        "Stress" : 0,
        "BetweennessCentrality" : 0.0,
        "t_RNA_motif_Root" : "NO",
        "EdgeCount" : 2,
        "Causal_PLAMOM" : "no",
        "name" : "AT1G20640",
        "Causal_tRNA" : "no",
        "IsSingleNode" : false,
        "briefDescription" : "Plant regulator RWP-RK family protein",
        "NeighborhoodConnectivity" : 11.5
      },
      "position" : {
        "x" : -5938.52734375,
        "y" : -539.1010131835938
      },
      "selected" : false
    }, {
      "data" : {
        "id" : "1346",
        "symbol" : "DAG1",
        "Shoot_Descriptions" : "",
        "PLAMOM_Match_Shoot" : "",
        "Causal_TF" : "Dof-type zinc finger DNA-binding family protein\"",
        "Root_Descriptions" : "DAG1, Dof-type zinc finger DNA-binding family protein",
        "Eccentricity" : 9,
        "Outdegree" : 7,
        "Target_PLAMOM" : "no",
        "PLAMOM_Match_Root" : "NO",
        "shared_name" : "AT3G61850",
        "Indegree" : 1,
        "Target_tRNA" : "no",
        "t_RNA_motif_shoot" : "",
        "SUID" : 1346,
        "SelfLoops" : 0,
        "Target_TF" : "Dof-type zinc finger DNA-binding family protein\"",
        "MolecularAttributes" : "TXNFACTOR",
        "AverageShortestPathLength" : 4.65481172,
        "selected" : false,
        "Name2" : "DAG1",
        "ClosenessCentrality" : 0.21483146,
        "Casual_Name" : "\"DAG1",
        "PartnerOfMultiEdgedNodePairs" : 0,
        "ClusteringCoefficient" : 0.0,
        "Target_Name" : "\"DAG1",
        "Stress" : 7204,
        "BetweennessCentrality" : 0.00161089,
        "t_RNA_motif_Root" : "NO",
        "EdgeCount" : 8,
        "Causal_PLAMOM" : "no",
        "name" : "AT3G61850",
        "Causal_tRNA" : "no",
        "IsSingleNode" : false,
        "briefDescription" : "Dof-type zinc finger DNA-binding family protein",
        "NeighborhoodConnectivity" : 10.625
      },
      "position" : {
        "x" : -6175.95556640625,
        "y" : 2824.094970703125
      },
      "selected" : false
    }, {
      "data" : {
        "id" : "1345",
        "symbol" : "CP1",
        "Shoot_Descriptions" : "",
        "PLAMOM_Match_Shoot" : "",
        "Causal_TF" : "CP1",
        "Root_Descriptions" : "ATCP1, CP1, Ca2+-binding protein 1",
        "Eccentricity" : 12,
        "Outdegree" : 3,
        "Target_PLAMOM" : "",
        "PLAMOM_Match_Root" : "YES",
        "shared_name" : "AT5G49480",
        "Indegree" : 0,
        "Target_tRNA" : "",
        "t_RNA_motif_shoot" : "",
        "SUID" : 1345,
        "SelfLoops" : 0,
        "Target_TF" : "",
        "MolecularAttributes" : "protein_coding",
        "AverageShortestPathLength" : 7.65970772,
        "selected" : false,
        "Name2" : "CP1",
        "ClosenessCentrality" : 0.13055328,
        "Casual_Name" : "\"ATCP1",
        "PartnerOfMultiEdgedNodePairs" : 0,
        "ClusteringCoefficient" : 0.0,
        "Target_Name" : "",
        "Stress" : 0,
        "BetweennessCentrality" : 0.0,
        "t_RNA_motif_Root" : "NO",
        "EdgeCount" : 3,
        "Causal_PLAMOM" : "AT5G49480",
        "name" : "AT5G49480",
        "Causal_tRNA" : "no",
        "IsSingleNode" : false,
        "briefDescription" : "Ca2 -binding protein 1",
        "NeighborhoodConnectivity" : 7.0
      },
      "position" : {
        "x" : -6019.63671875,
        "y" : -6.320960998535156
      },
      "selected" : false
    }, {
      "data" : {
        "id" : "1344",
        "symbol" : "NRT2:1",
        "Shoot_Descriptions" : "ACH1, ATNRT2.1, ATNRT2:1, LIN1, NRT2, NRT2.1, NRT2:1, NRT2;1AT, nitrate transporter 2:1",
        "PLAMOM_Match_Shoot" : "NO",
        "Causal_TF" : "ATNRT2.1",
        "Root_Descriptions" : "ACH1, ATNRT2.1, ATNRT2:1, LIN1, NRT2, NRT2.1, NRT2:1, NRT2;1AT, nitrate transporter 2:1",
        "Eccentricity" : 1,
        "Outdegree" : 1,
        "Target_PLAMOM" : "ATGLB2",
        "PLAMOM_Match_Root" : "NO",
        "shared_name" : "AT1G08090",
        "Indegree" : 3,
        "Target_tRNA" : "GLB2",
        "t_RNA_motif_shoot" : "NO",
        "SUID" : 1344,
        "SelfLoops" : 0,
        "Target_TF" : "NSHB2",
        "MolecularAttributes" : "protein_coding",
        "AverageShortestPathLength" : 1.0,
        "selected" : false,
        "Name2" : "NRT2:1",
        "ClosenessCentrality" : 1.0,
        "Casual_Name" : "\"ACH1",
        "PartnerOfMultiEdgedNodePairs" : 0,
        "ClusteringCoefficient" : 0.0,
        "Target_Name" : "HB2",
        "Stress" : 3,
        "BetweennessCentrality" : 1.79E-6,
        "t_RNA_motif_Root" : "NO",
        "EdgeCount" : 4,
        "Causal_PLAMOM" : "no",
        "name" : "AT1G08090",
        "Causal_tRNA" : "no",
        "IsSingleNode" : false,
        "briefDescription" : "nitrate transporter 2:1",
        "NeighborhoodConnectivity" : 7.25
      },
      "position" : {
        "x" : -560.1822391269852,
        "y" : -282.0693054199219
      },
      "selected" : false
    }, {
      "data" : {
        "id" : "1343",
        "symbol" : "SULTR1;3",
        "Shoot_Descriptions" : "",
        "PLAMOM_Match_Shoot" : "",
        "Causal_TF" : "sulfate transporter 1;3\"",
        "Root_Descriptions" : "SULTR1;3, sulfate transporter 1;3",
        "Eccentricity" : 13,
        "Outdegree" : 4,
        "Target_PLAMOM" : "",
        "PLAMOM_Match_Root" : "NO",
        "shared_name" : "AT1G22150",
        "Indegree" : 0,
        "Target_tRNA" : "",
        "t_RNA_motif_shoot" : "",
        "SUID" : 1343,
        "SelfLoops" : 0,
        "Target_TF" : "",
        "MolecularAttributes" : "protein_coding",
        "AverageShortestPathLength" : 8.42588727,
        "selected" : false,
        "Name2" : "SULTR1;3",
        "ClosenessCentrality" : 0.11868186,
        "Casual_Name" : "\"SULTR1;3",
        "PartnerOfMultiEdgedNodePairs" : 0,
        "ClusteringCoefficient" : 0.0,
        "Target_Name" : "",
        "Stress" : 0,
        "BetweennessCentrality" : 0.0,
        "t_RNA_motif_Root" : "NO",
        "EdgeCount" : 4,
        "Causal_PLAMOM" : "no",
        "name" : "AT1G22150",
        "Causal_tRNA" : "no",
        "IsSingleNode" : false,
        "briefDescription" : "sulfate transporter 1;3",
        "NeighborhoodConnectivity" : 19.25
      },
      "position" : {
        "x" : -5993.6806640625,
        "y" : -487.29632568359375
      },
      "selected" : false
    }, {
      "data" : {
        "id" : "1342",
        "symbol" : "HB2",
        "Shoot_Descriptions" : "AHB2, ARATH GLB2, ATGLB2, GLB2, HB2, NSHB2, haemoglobin 2",
        "PLAMOM_Match_Shoot" : "YES",
        "Causal_TF" : "ARATH GLB2",
        "Root_Descriptions" : "AHB2, ARATH GLB2, ATGLB2, GLB2, HB2, NSHB2, haemoglobin 2",
        "Eccentricity" : 9,
        "Outdegree" : 7,
        "Target_PLAMOM" : "no",
        "PLAMOM_Match_Root" : "YES",
        "shared_name" : "AT3G10520",
        "Indegree" : 2,
        "Target_tRNA" : "AT3G10520",
        "t_RNA_motif_shoot" : "NO",
        "SUID" : 1342,
        "SelfLoops" : 0,
        "Target_TF" : "\"AHB2",
        "MolecularAttributes" : "protein_coding",
        "AverageShortestPathLength" : 4.78661088,
        "selected" : false,
        "Name2" : "HB2",
        "ClosenessCentrality" : 0.20891608,
        "Casual_Name" : "\"AHB2",
        "PartnerOfMultiEdgedNodePairs" : 0,
        "ClusteringCoefficient" : 0.0,
        "Target_Name" : "no",
        "Stress" : 7568,
        "BetweennessCentrality" : 0.0016199,
        "t_RNA_motif_Root" : "NO",
        "EdgeCount" : 9,
        "Causal_PLAMOM" : "AT3G10520",
        "name" : "AT3G10520",
        "Causal_tRNA" : "no",
        "IsSingleNode" : false,
        "briefDescription" : "hemoglobin 2",
        "NeighborhoodConnectivity" : 24.0
      },
      "position" : {
        "x" : -1329.7685546875,
        "y" : 2789.7897844940426
      },
      "selected" : false
    }, {
      "data" : {
        "id" : "1341",
        "symbol" : "AT1G80400",
        "Shoot_Descriptions" : "",
        "PLAMOM_Match_Shoot" : "",
        "Causal_TF" : "no",
        "Root_Descriptions" : "RING/U-box superfamily protein",
        "Eccentricity" : 13,
        "Outdegree" : 3,
        "Target_PLAMOM" : "",
        "PLAMOM_Match_Root" : "NO",
        "shared_name" : "AT1G80400",
        "Indegree" : 0,
        "Target_tRNA" : "",
        "t_RNA_motif_shoot" : "",
        "SUID" : 1341,
        "SelfLoops" : 0,
        "Target_TF" : "",
        "MolecularAttributes" : "TXNFACTOR",
        "AverageShortestPathLength" : 8.60416667,
        "selected" : false,
        "Name2" : "AT1G80400",
        "ClosenessCentrality" : 0.11622276,
        "Casual_Name" : "RING/U-box superfamily protein",
        "PartnerOfMultiEdgedNodePairs" : 0,
        "ClusteringCoefficient" : 0.0,
        "Target_Name" : "",
        "Stress" : 0,
        "BetweennessCentrality" : 0.0,
        "t_RNA_motif_Root" : "NO",
        "EdgeCount" : 3,
        "Causal_PLAMOM" : "no",
        "name" : "AT1G80400",
        "Causal_tRNA" : "no",
        "IsSingleNode" : false,
        "briefDescription" : "RING/U-box superfamily protein",
        "NeighborhoodConnectivity" : 3.6666666666666665
      },
      "position" : {
        "x" : -5702.4814453125,
        "y" : 7.594202041625977
      },
      "selected" : false
    }, {
      "data" : {
        "id" : "1340",
        "symbol" : "AT1G09640",
        "Shoot_Descriptions" : "Translation elongation factor EF1B, gamma chain",
        "PLAMOM_Match_Shoot" : "YES",
        "Causal_TF" : "gamma chain\"",
        "Root_Descriptions" : "",
        "Eccentricity" : 13,
        "Outdegree" : 1,
        "Target_PLAMOM" : "AT1G09640",
        "PLAMOM_Match_Root" : "",
        "shared_name" : "AT1G09640",
        "Indegree" : 2,
        "Target_tRNA" : "no",
        "t_RNA_motif_shoot" : "NO",
        "SUID" : 1340,
        "SelfLoops" : 0,
        "Target_TF" : "gamma chain\"",
        "MolecularAttributes" : "protein_coding",
        "AverageShortestPathLength" : 8.0125261,
        "selected" : false,
        "Name2" : "AT1G09640",
        "ClosenessCentrality" : 0.12480459,
        "Casual_Name" : "\"Translation elongation factor EF1B",
        "PartnerOfMultiEdgedNodePairs" : 0,
        "ClusteringCoefficient" : 0.0,
        "Target_Name" : "\"Translation elongation factor EF1B",
        "Stress" : 596,
        "BetweennessCentrality" : 1.9285E-4,
        "t_RNA_motif_Root" : "",
        "EdgeCount" : 3,
        "Causal_PLAMOM" : "AT1G09640",
        "name" : "AT1G09640",
        "Causal_tRNA" : "no",
        "IsSingleNode" : false,
        "briefDescription" : "Translation elongation factor EF1B",
        "NeighborhoodConnectivity" : 27.333333333333332
      },
      "position" : {
        "x" : 3528.693798470671,
        "y" : 3395.303955078125
      },
      "selected" : false
    }, {
      "data" : {
        "id" : "1339",
        "symbol" : "AT5G15790",
        "Shoot_Descriptions" : "",
        "PLAMOM_Match_Shoot" : "",
        "Causal_TF" : "no",
        "Root_Descriptions" : "RING/U-box superfamily protein",
        "Eccentricity" : 11,
        "Outdegree" : 7,
        "Target_PLAMOM" : "",
        "PLAMOM_Match_Root" : "NO",
        "shared_name" : "AT5G15790",
        "Indegree" : 0,
        "Target_tRNA" : "",
        "t_RNA_motif_shoot" : "",
        "SUID" : 1339,
        "SelfLoops" : 0,
        "Target_TF" : "",
        "MolecularAttributes" : "TXNFACTOR",
        "AverageShortestPathLength" : 5.77366255,
        "selected" : false,
        "Name2" : "AT5G15790",
        "ClosenessCentrality" : 0.17320029,
        "Casual_Name" : "RING/U-box superfamily protein",
        "PartnerOfMultiEdgedNodePairs" : 0,
        "ClusteringCoefficient" : 0.0,
        "Target_Name" : "",
        "Stress" : 0,
        "BetweennessCentrality" : 0.0,
        "t_RNA_motif_Root" : "NO",
        "EdgeCount" : 7,
        "Causal_PLAMOM" : "no",
        "name" : "AT5G15790",
        "Causal_tRNA" : "no",
        "IsSingleNode" : false,
        "briefDescription" : "RING/U-box superfamily protein",
        "NeighborhoodConnectivity" : 5.571428571428571
      },
      "position" : {
        "x" : -6322.2587890625,
        "y" : 391.7510986328125
      },
      "selected" : false
    }, {
      "data" : {
        "id" : "1338",
        "symbol" : "MDAR6",
        "Shoot_Descriptions" : "",
        "PLAMOM_Match_Shoot" : "",
        "Causal_TF" : "monodehydroascorbate reductase 6\"",
        "Root_Descriptions" : "MDAR6, monodehydroascorbate reductase 6",
        "Eccentricity" : 12,
        "Outdegree" : 11,
        "Target_PLAMOM" : "no",
        "PLAMOM_Match_Root" : "YES",
        "shared_name" : "AT1G63940",
        "Indegree" : 5,
        "Target_tRNA" : "AT1G63940",
        "t_RNA_motif_shoot" : "",
        "SUID" : 1338,
        "SelfLoops" : 0,
        "Target_TF" : "\"MDAR6",
        "MolecularAttributes" : "protein_coding",
        "AverageShortestPathLength" : 6.79916318,
        "selected" : false,
        "Name2" : "MDAR6",
        "ClosenessCentrality" : 0.14707692,
        "Casual_Name" : "\"MDAR6",
        "PartnerOfMultiEdgedNodePairs" : 0,
        "ClusteringCoefficient" : 0.0,
        "Target_Name" : "no",
        "Stress" : 22988,
        "BetweennessCentrality" : 0.00487363,
        "t_RNA_motif_Root" : "NO",
        "EdgeCount" : 16,
        "Causal_PLAMOM" : "AT1G63940",
        "name" : "AT1G63940",
        "Causal_tRNA" : "no",
        "IsSingleNode" : false,
        "briefDescription" : "monodehydroascorbate reductase 6",
        "NeighborhoodConnectivity" : 2.5
      },
      "position" : {
        "x" : -6389.92431640625,
        "y" : 3809.177978515625
      },
      "selected" : false
    }, {
      "data" : {
        "id" : "1337",
        "symbol" : "AT1G09900",
        "Shoot_Descriptions" : "Pentatricopeptide repeat (PPR-like) superfamily protein",
        "PLAMOM_Match_Shoot" : "NO",
        "Causal_TF" : "",
        "Root_Descriptions" : "",
        "Eccentricity" : 0,
        "Outdegree" : 0,
        "Target_PLAMOM" : "no",
        "PLAMOM_Match_Root" : "",
        "shared_name" : "AT1G09900",
        "Indegree" : 1,
        "Target_tRNA" : "no",
        "t_RNA_motif_shoot" : "NO",
        "SUID" : 1337,
        "SelfLoops" : 0,
        "Target_TF" : "Pentatricopeptide repeat (PPR-like) superfamily protein",
        "MolecularAttributes" : "protein_coding",
        "AverageShortestPathLength" : 0.0,
        "selected" : false,
        "Name2" : "AT1G09900",
        "ClosenessCentrality" : 0.0,
        "Casual_Name" : "",
        "PartnerOfMultiEdgedNodePairs" : 0,
        "ClusteringCoefficient" : 0.0,
        "Target_Name" : "no",
        "Stress" : 0,
        "BetweennessCentrality" : 0.0,
        "t_RNA_motif_Root" : "",
        "EdgeCount" : 1,
        "Causal_PLAMOM" : "",
        "name" : "AT1G09900",
        "Causal_tRNA" : "",
        "IsSingleNode" : false,
        "briefDescription" : "Pentatricopeptide repeat (PPR-like) superfamily protein",
        "NeighborhoodConnectivity" : 16.0
      },
      "position" : {
        "x" : 4115.941904959082,
        "y" : 979.1416015625
      },
      "selected" : false
    }, {
      "data" : {
        "id" : "1336",
        "symbol" : "AT1G23840",
        "Shoot_Descriptions" : "",
        "PLAMOM_Match_Shoot" : "",
        "Causal_TF" : "no",
        "Root_Descriptions" : "unknown protein; LOCATED IN: endomembrane system; EXPRESSED IN: 18 plant structures; EXPRESSED DURING: 12 growth stages; BEST Arabidopsis thaliana protein match is: unknown protein (TAIR:AT1G23850.1); Has 53 Blast hits to 48 proteins in 5 species: Archae - 0; Bacteria - 0; Metazoa - 0; Fungi - 0; Plants - 53; Viruses - 0; Other Eukaryotes - 0 (source: NCBI BLink).",
        "Eccentricity" : 15,
        "Outdegree" : 4,
        "Target_PLAMOM" : "",
        "PLAMOM_Match_Root" : "NO",
        "shared_name" : "AT1G23840",
        "Indegree" : 0,
        "Target_tRNA" : "",
        "t_RNA_motif_shoot" : "",
        "SUID" : 1336,
        "SelfLoops" : 0,
        "Target_TF" : "",
        "MolecularAttributes" : "protein_coding",
        "AverageShortestPathLength" : 9.93195876,
        "selected" : false,
        "Name2" : "AT1G23840",
        "ClosenessCentrality" : 0.10068507,
        "Casual_Name" : "unknown protein; LOCATED IN: endomembrane system; EXPRESSED IN: 18 plant structures; EXPRESSED DURING: 12 growth stages; BEST Arabidopsis thaliana protein match is: unknown protein (TAIR:AT1G23850.1); Has 53 Blast hits to 48 proteins in 5 species: Archae - 0; Bacteria - 0; Metazoa - 0; Fungi - 0; Plants - 53; Viruses - 0; Other Eukaryotes - 0 (source: NCBI BLink).",
        "PartnerOfMultiEdgedNodePairs" : 0,
        "ClusteringCoefficient" : 0.0,
        "Target_Name" : "",
        "Stress" : 0,
        "BetweennessCentrality" : 0.0,
        "t_RNA_motif_Root" : "NO",
        "EdgeCount" : 4,
        "Causal_PLAMOM" : "no",
        "name" : "AT1G23840",
        "Causal_tRNA" : "no",
        "IsSingleNode" : false,
        "briefDescription" : "transmembrane protein",
        "NeighborhoodConnectivity" : 10.5
      },
      "position" : {
        "x" : -5696.54052734375,
        "y" : -84.35731506347656
      },
      "selected" : false
    }, {
      "data" : {
        "id" : "1335",
        "symbol" : "NFXL1",
        "Shoot_Descriptions" : "ATNFXL1, NFXL1, NF-X-like 1",
        "PLAMOM_Match_Shoot" : "NO",
        "Causal_TF" : "NFXL1",
        "Root_Descriptions" : "",
        "Eccentricity" : 3,
        "Outdegree" : 1,
        "Target_PLAMOM" : "no",
        "PLAMOM_Match_Root" : "",
        "shared_name" : "AT1G10170",
        "Indegree" : 1,
        "Target_tRNA" : "no",
        "t_RNA_motif_shoot" : "NO",
        "SUID" : 1335,
        "SelfLoops" : 0,
        "Target_TF" : "NFXL1",
        "MolecularAttributes" : "protein_coding",
        "AverageShortestPathLength" : 2.0,
        "selected" : false,
        "Name2" : "NFXL1",
        "ClosenessCentrality" : 0.5,
        "Casual_Name" : "\"ATNFXL1",
        "PartnerOfMultiEdgedNodePairs" : 0,
        "ClusteringCoefficient" : 0.0,
        "Target_Name" : "\"ATNFXL1",
        "Stress" : 4,
        "BetweennessCentrality" : 2.39E-6,
        "t_RNA_motif_Root" : "",
        "EdgeCount" : 2,
        "Causal_PLAMOM" : "no",
        "name" : "AT1G10170",
        "Causal_tRNA" : "no",
        "IsSingleNode" : false,
        "briefDescription" : "NF-X-like 1",
        "NeighborhoodConnectivity" : 4.0
      },
      "position" : {
        "x" : 4049.490234375,
        "y" : 2438.333984375
      },
      "selected" : false
    }, {
      "data" : {
        "id" : "1334",
        "symbol" : "AT2G36470",
        "Shoot_Descriptions" : "",
        "PLAMOM_Match_Shoot" : "",
        "Causal_TF" : "no",
        "Root_Descriptions" : "Plant protein of unknown function (DUF868)",
        "Eccentricity" : 10,
        "Outdegree" : 11,
        "Target_PLAMOM" : "receptor like protein 4\"",
        "PLAMOM_Match_Root" : "NO",
        "shared_name" : "AT2G36470",
        "Indegree" : 1,
        "Target_tRNA" : "no",
        "t_RNA_motif_shoot" : "",
        "SUID" : 1334,
        "SelfLoops" : 0,
        "Target_TF" : "no",
        "MolecularAttributes" : "protein_coding",
        "AverageShortestPathLength" : 5.60460251,
        "selected" : false,
        "Name2" : "AT2G36470",
        "ClosenessCentrality" : 0.17842479,
        "Casual_Name" : "Plant protein of unknown function (DUF868)",
        "PartnerOfMultiEdgedNodePairs" : 0,
        "ClusteringCoefficient" : 0.0,
        "Target_Name" : "no",
        "Stress" : 2363,
        "BetweennessCentrality" : 5.2956E-4,
        "t_RNA_motif_Root" : "NO",
        "EdgeCount" : 12,
        "Causal_PLAMOM" : "no",
        "name" : "AT2G36470",
        "Causal_tRNA" : "no",
        "IsSingleNode" : false,
        "briefDescription" : "DUF868 family protein",
        "NeighborhoodConnectivity" : 7.583333333333333
      },
      "position" : {
        "x" : -6758.9375,
        "y" : 3003.739013671875
      },
      "selected" : false
    }, {
      "data" : {
        "id" : "1333",
        "symbol" : "AT1G11710",
        "Shoot_Descriptions" : "Pentatricopeptide repeat (PPR) superfamily protein",
        "PLAMOM_Match_Shoot" : "NO",
        "Causal_TF" : "no",
        "Root_Descriptions" : "",
        "Eccentricity" : 13,
        "Outdegree" : 1,
        "Target_PLAMOM" : "no",
        "PLAMOM_Match_Root" : "",
        "shared_name" : "AT1G11710",
        "Indegree" : 1,
        "Target_tRNA" : "no",
        "t_RNA_motif_shoot" : "NO",
        "SUID" : 1333,
        "SelfLoops" : 0,
        "Target_TF" : "no",
        "MolecularAttributes" : "protein_coding",
        "AverageShortestPathLength" : 7.9623431,
        "selected" : false,
        "Name2" : "AT1G11710",
        "ClosenessCentrality" : 0.12559117,
        "Casual_Name" : "Pentatricopeptide repeat (PPR) superfamily protein",
        "PartnerOfMultiEdgedNodePairs" : 0,
        "ClusteringCoefficient" : 0.0,
        "Target_Name" : "Pentatricopeptide repeat (PPR) superfamily protein",
        "Stress" : 34,
        "BetweennessCentrality" : 3.81E-6,
        "t_RNA_motif_Root" : "",
        "EdgeCount" : 2,
        "Causal_PLAMOM" : "no",
        "name" : "AT1G11710",
        "Causal_tRNA" : "no",
        "IsSingleNode" : false,
        "briefDescription" : "Pentatricopeptide repeat (PPR) superfamily protein",
        "NeighborhoodConnectivity" : 19.5
      },
      "position" : {
        "x" : 2952.867431640625,
        "y" : 2678.280029296875
      },
      "selected" : false
    }, {
      "data" : {
        "id" : "1332",
        "symbol" : "AT1G12020",
        "Shoot_Descriptions" : "unknown protein; BEST Arabidopsis thaliana protein match is: unknown protein (TAIR:AT1G62422.1); Has 89 Blast hits to 88 proteins in 16 species: Archae - 0; Bacteria - 0; Metazoa - 0; Fungi - 0; Plants - 87; Viruses - 0; Other Eukaryotes - 2 (source: NCBI BLink).",
        "PLAMOM_Match_Shoot" : "NO",
        "Causal_TF" : "",
        "Root_Descriptions" : "",
        "Eccentricity" : 0,
        "Outdegree" : 0,
        "Target_PLAMOM" : "no",
        "PLAMOM_Match_Root" : "",
        "shared_name" : "AT1G12020",
        "Indegree" : 1,
        "Target_tRNA" : "no",
        "t_RNA_motif_shoot" : "YES",
        "SUID" : 1332,
        "SelfLoops" : 0,
        "Target_TF" : "unknown protein; BEST Arabidopsis thaliana protein match is: unknown protein (TAIR:AT1G62422.1); Has 89 Blast hits to 88 proteins in 16 species: Archae - 0; Bacteria - 0; Metazoa - 0; Fungi - 0; Plants - 87; Viruses - 0; Other Eukaryotes - 2 (source: NCBI BLink).",
        "MolecularAttributes" : "protein_coding",
        "AverageShortestPathLength" : 0.0,
        "selected" : false,
        "Name2" : "AT1G12020",
        "ClosenessCentrality" : 0.0,
        "Casual_Name" : "",
        "PartnerOfMultiEdgedNodePairs" : 0,
        "ClusteringCoefficient" : 0.0,
        "Target_Name" : "AT1G12020",
        "Stress" : 0,
        "BetweennessCentrality" : 0.0,
        "t_RNA_motif_Root" : "",
        "EdgeCount" : 1,
        "Causal_PLAMOM" : "",
        "name" : "AT1G12020",
        "Causal_tRNA" : "",
        "IsSingleNode" : false,
        "briefDescription" : "hypothetical protein",
        "NeighborhoodConnectivity" : 16.0
      },
      "position" : {
        "x" : 4135.278759408171,
        "y" : 1101.3218994140625
      },
      "selected" : false
    }, {
      "data" : {
        "id" : "1331",
        "symbol" : "MC8",
        "degree_layout" : 4,
        "Shoot_Descriptions" : "",
        "PLAMOM_Match_Shoot" : "",
        "Causal_TF" : "MC8",
        "Root_Descriptions" : "ATMC8, MC8, metacaspase 8",
        "Eccentricity" : 9,
        "Outdegree" : 10,
        "Target_PLAMOM" : "NAC domain containing protein 96\"",
        "PLAMOM_Match_Root" : "NO",
        "shared_name" : "AT1G16420",
        "Indegree" : 5,
        "Target_tRNA" : "AT5G46590",
        "t_RNA_motif_shoot" : "",
        "SUID" : 1331,
        "SelfLoops" : 0,
        "Target_TF" : "AT1G16420",
        "MolecularAttributes" : "protein_coding",
        "AverageShortestPathLength" : 4.77615063,
        "selected" : false,
        "Name2" : "MC8",
        "ClosenessCentrality" : 0.20937363,
        "Casual_Name" : "\"ATMC8",
        "PartnerOfMultiEdgedNodePairs" : 0,
        "ClusteringCoefficient" : 0.0,
        "Target_Name" : "no",
        "Stress" : 26271,
        "BetweennessCentrality" : 0.00599422,
        "t_RNA_motif_Root" : "YES",
        "EdgeCount" : 15,
        "Causal_PLAMOM" : "no",
        "name" : "AT1G16420",
        "Causal_tRNA" : "AT1G16420",
        "IsSingleNode" : false,
        "briefDescription" : "metacaspase 8",
        "NeighborhoodConnectivity" : 10.4
      },
      "position" : {
        "x" : -6160.23828125,
        "y" : 3006.800537109375
      },
      "selected" : false
    }, {
      "data" : {
        "id" : "1330",
        "symbol" : "AT1G12650",
        "Shoot_Descriptions" : "unknown protein; CONTAINS InterPro DOMAIN/s: Protein of unknown function DUF947 (InterPro:IPR009292); Has 700 Blast hits to 631 proteins in 192 species: Archae - 0; Bacteria - 32; Metazoa - 138; Fungi - 168; Plants - 60; Viruses - 0; Other Eukaryotes - 302 (source: NCBI BLink).",
        "PLAMOM_Match_Shoot" : "NO",
        "Causal_TF" : "no",
        "Root_Descriptions" : "",
        "Eccentricity" : 2,
        "Outdegree" : 1,
        "Target_PLAMOM" : "metacaspase 8\"",
        "PLAMOM_Match_Root" : "",
        "shared_name" : "AT1G12650",
        "Indegree" : 1,
        "Target_tRNA" : "no",
        "t_RNA_motif_shoot" : "NO",
        "SUID" : 1330,
        "SelfLoops" : 0,
        "Target_TF" : "no",
        "MolecularAttributes" : "protein_coding",
        "AverageShortestPathLength" : 1.5,
        "selected" : false,
        "Name2" : "AT1G12650",
        "ClosenessCentrality" : 0.66666667,
        "Casual_Name" : "unknown protein; CONTAINS InterPro DOMAIN/s: Protein of unknown function DUF947 (InterPro:IPR009292); Has 700 Blast hits to 631 proteins in 192 species: Archae - 0; Bacteria - 32; Metazoa - 138; Fungi - 168; Plants - 60; Viruses - 0; Other Eukaryotes - 302 (source: NCBI BLink).",
        "PartnerOfMultiEdgedNodePairs" : 0,
        "ClusteringCoefficient" : 0.0,
        "Target_Name" : "no",
        "Stress" : 243,
        "BetweennessCentrality" : 4.224E-5,
        "t_RNA_motif_Root" : "",
        "EdgeCount" : 2,
        "Causal_PLAMOM" : "no",
        "name" : "AT1G12650",
        "Causal_tRNA" : "no",
        "IsSingleNode" : false,
        "briefDescription" : "rRNA biogenesis RRP36-like protein",
        "NeighborhoodConnectivity" : 14.0
      },
      "position" : {
        "x" : 4238.895946908171,
        "y" : 2889.342041015625
      },
      "selected" : false
    }, {
      "data" : {
        "id" : "1329",
        "symbol" : "AAE17",
        "Shoot_Descriptions" : "",
        "PLAMOM_Match_Shoot" : "",
        "Causal_TF" : "acyl-activating enzyme 17\"",
        "Root_Descriptions" : "AAE17, acyl-activating enzyme 17",
        "Eccentricity" : 11,
        "Outdegree" : 4,
        "Target_PLAMOM" : "no",
        "PLAMOM_Match_Root" : "NO",
        "shared_name" : "AT5G23050",
        "Indegree" : 1,
        "Target_tRNA" : "no",
        "t_RNA_motif_shoot" : "",
        "SUID" : 1329,
        "SelfLoops" : 0,
        "Target_TF" : "\"AAE17",
        "MolecularAttributes" : "METABOLIC",
        "AverageShortestPathLength" : 5.97708333,
        "selected" : false,
        "Name2" : "AAE17",
        "ClosenessCentrality" : 0.16730568,
        "Casual_Name" : "\"AAE17",
        "PartnerOfMultiEdgedNodePairs" : 0,
        "ClusteringCoefficient" : 0.0,
        "Target_Name" : "no",
        "Stress" : 376,
        "BetweennessCentrality" : 9.825E-5,
        "t_RNA_motif_Root" : "NO",
        "EdgeCount" : 5,
        "Causal_PLAMOM" : "no",
        "name" : "AT5G23050",
        "Causal_tRNA" : "no",
        "IsSingleNode" : false,
        "briefDescription" : "acyl-activating enzyme 17",
        "NeighborhoodConnectivity" : 8.2
      },
      "position" : {
        "x" : -6246.0771484375,
        "y" : 3137.00146484375
      },
      "selected" : false
    }, {
      "data" : {
        "id" : "1328",
        "symbol" : "ESR1",
        "Shoot_Descriptions" : "DRN, ESR1, Integrase-type DNA-binding superfamily protein",
        "PLAMOM_Match_Shoot" : "NO",
        "Causal_TF" : "ESR1",
        "Root_Descriptions" : "",
        "Eccentricity" : 11,
        "Outdegree" : 2,
        "Target_PLAMOM" : "no",
        "PLAMOM_Match_Root" : "",
        "shared_name" : "AT1G12980",
        "Indegree" : 1,
        "Target_tRNA" : "no",
        "t_RNA_motif_shoot" : "NO",
        "SUID" : 1328,
        "SelfLoops" : 0,
        "Target_TF" : "\"DRN",
        "MolecularAttributes" : "TXNFACTOR",
        "AverageShortestPathLength" : 5.98121086,
        "selected" : false,
        "Name2" : "ESR1",
        "ClosenessCentrality" : 0.16719023,
        "Casual_Name" : "\"DRN",
        "PartnerOfMultiEdgedNodePairs" : 0,
        "ClusteringCoefficient" : 0.0,
        "Target_Name" : "no",
        "Stress" : 119,
        "BetweennessCentrality" : 2.359E-5,
        "t_RNA_motif_Root" : "",
        "EdgeCount" : 3,
        "Causal_PLAMOM" : "no",
        "name" : "AT1G12980",
        "Causal_tRNA" : "no",
        "IsSingleNode" : false,
        "briefDescription" : "Integrase-type DNA-binding superfamily protein",
        "NeighborhoodConnectivity" : 48.333333333333336
      },
      "position" : {
        "x" : 3700.859081673796,
        "y" : 3377.21533203125
      },
      "selected" : false
    }, {
      "data" : {
        "id" : "1327",
        "symbol" : "AT1G15630",
        "Shoot_Descriptions" : "",
        "PLAMOM_Match_Shoot" : "",
        "Causal_TF" : "no",
        "Root_Descriptions" : "unknown protein; BEST Arabidopsis thaliana protein match is: unknown protein (TAIR:AT1G15610.1); Has 122 Blast hits to 122 proteins in 14 species: Archae - 0; Bacteria - 0; Metazoa - 0; Fungi - 0; Plants - 122; Viruses - 0; Other Eukaryotes - 0 (source: NCBI BLink).",
        "Eccentricity" : 10,
        "Outdegree" : 12,
        "Target_PLAMOM" : "ferric reduction oxidase 8\"",
        "PLAMOM_Match_Root" : "NO",
        "shared_name" : "AT1G15630",
        "Indegree" : 3,
        "Target_tRNA" : "no",
        "t_RNA_motif_shoot" : "",
        "SUID" : 1327,
        "SelfLoops" : 0,
        "Target_TF" : "no",
        "MolecularAttributes" : "protein_coding",
        "AverageShortestPathLength" : 5.84518828,
        "selected" : false,
        "Name2" : "AT1G15630",
        "ClosenessCentrality" : 0.17108089,
        "Casual_Name" : "unknown protein; BEST Arabidopsis thaliana protein match is: unknown protein (TAIR:AT1G15610.1); Has 122 Blast hits to 122 proteins in 14 species: Archae - 0; Bacteria - 0; Metazoa - 0; Fungi - 0; Plants - 122; Viruses - 0; Other Eukaryotes - 0 (source: NCBI BLink).",
        "PartnerOfMultiEdgedNodePairs" : 0,
        "ClusteringCoefficient" : 0.0,
        "Target_Name" : "no",
        "Stress" : 37749,
        "BetweennessCentrality" : 0.00749409,
        "t_RNA_motif_Root" : "NO",
        "EdgeCount" : 15,
        "Causal_PLAMOM" : "no",
        "name" : "AT1G15630",
        "Causal_tRNA" : "no",
        "IsSingleNode" : false,
        "briefDescription" : "transmembrane protein",
        "NeighborhoodConnectivity" : 6.933333333333334
      },
      "position" : {
        "x" : -6987.7568359375,
        "y" : 3362.45263671875
      },
      "selected" : false
    }, {
      "data" : {
        "id" : "1326",
        "symbol" : "AT1G12990",
        "Shoot_Descriptions" : "beta-1,4-N-acetylglucosaminyltransferase family protein",
        "PLAMOM_Match_Shoot" : "NO",
        "Causal_TF" : "4-N-acetylglucosaminyltransferase family protein\"",
        "Root_Descriptions" : "",
        "Eccentricity" : 11,
        "Outdegree" : 1,
        "Target_PLAMOM" : "no",
        "PLAMOM_Match_Root" : "",
        "shared_name" : "AT1G12990",
        "Indegree" : 3,
        "Target_tRNA" : "no",
        "t_RNA_motif_shoot" : "NO",
        "SUID" : 1326,
        "SelfLoops" : 0,
        "Target_TF" : "4-N-acetylglucosaminyltransferase family protein\"",
        "MolecularAttributes" : "protein_coding",
        "AverageShortestPathLength" : 6.60878661,
        "selected" : false,
        "Name2" : "AT1G12990",
        "ClosenessCentrality" : 0.15131371,
        "Casual_Name" : "\"beta-1",
        "PartnerOfMultiEdgedNodePairs" : 0,
        "ClusteringCoefficient" : 0.0,
        "Target_Name" : "\"beta-1",
        "Stress" : 4778,
        "BetweennessCentrality" : 0.00135879,
        "t_RNA_motif_Root" : "",
        "EdgeCount" : 4,
        "Causal_PLAMOM" : "no",
        "name" : "AT1G12990",
        "Causal_tRNA" : "no",
        "IsSingleNode" : false,
        "briefDescription" : "beta-1",
        "NeighborhoodConnectivity" : 16.0
      },
      "position" : {
        "x" : 2981.763866830046,
        "y" : 3222.810546875
      },
      "selected" : false
    }, {
      "data" : {
        "id" : "1325",
        "symbol" : "AT3G57070",
        "Shoot_Descriptions" : "Glutaredoxin family protein",
        "PLAMOM_Match_Shoot" : "NO",
        "Causal_TF" : "no",
        "Root_Descriptions" : "Glutaredoxin family protein",
        "Eccentricity" : 11,
        "Outdegree" : 8,
        "Target_PLAMOM" : "no",
        "PLAMOM_Match_Root" : "NO",
        "shared_name" : "AT3G57070",
        "Indegree" : 10,
        "Target_tRNA" : "no",
        "t_RNA_motif_shoot" : "NO",
        "SUID" : 1325,
        "SelfLoops" : 0,
        "Target_TF" : "no",
        "MolecularAttributes" : "protein_coding",
        "AverageShortestPathLength" : 6.91422594,
        "selected" : false,
        "Name2" : "AT3G57070",
        "ClosenessCentrality" : 0.14462935,
        "Casual_Name" : "Glutaredoxin family protein",
        "PartnerOfMultiEdgedNodePairs" : 0,
        "ClusteringCoefficient" : 0.00980392,
        "Target_Name" : "Glutaredoxin family protein",
        "Stress" : 11688,
        "BetweennessCentrality" : 0.00291079,
        "t_RNA_motif_Root" : "NO",
        "EdgeCount" : 18,
        "Causal_PLAMOM" : "no",
        "name" : "AT3G57070",
        "Causal_tRNA" : "no",
        "IsSingleNode" : false,
        "briefDescription" : "Glutaredoxin family protein",
        "NeighborhoodConnectivity" : 8.222222222222221
      },
      "position" : {
        "x" : -1629.041064810579,
        "y" : 3494.675048828125
      },
      "selected" : false
    }, {
      "data" : {
        "id" : "1324",
        "symbol" : "CKX4",
        "Shoot_Descriptions" : "",
        "PLAMOM_Match_Shoot" : "",
        "Causal_TF" : "CKX4",
        "Root_Descriptions" : "ATCKX4, CKX4, cytokinin oxidase 4",
        "Eccentricity" : 10,
        "Outdegree" : 6,
        "Target_PLAMOM" : "no",
        "PLAMOM_Match_Root" : "NO",
        "shared_name" : "AT4G29740",
        "Indegree" : 1,
        "Target_tRNA" : "no",
        "t_RNA_motif_shoot" : "",
        "SUID" : 1324,
        "SelfLoops" : 0,
        "Target_TF" : "CKX4",
        "MolecularAttributes" : "METABOLIC",
        "AverageShortestPathLength" : 5.54037267,
        "selected" : false,
        "Name2" : "CKX4",
        "ClosenessCentrality" : 0.18049327,
        "Casual_Name" : "\"ATCKX4",
        "PartnerOfMultiEdgedNodePairs" : 0,
        "ClusteringCoefficient" : 0.0,
        "Target_Name" : "\"ATCKX4",
        "Stress" : 1250,
        "BetweennessCentrality" : 1.5672E-4,
        "t_RNA_motif_Root" : "NO",
        "EdgeCount" : 7,
        "Causal_PLAMOM" : "no",
        "name" : "AT4G29740",
        "Causal_tRNA" : "no",
        "IsSingleNode" : false,
        "briefDescription" : "cytokinin oxidase 4",
        "NeighborhoodConnectivity" : 2.7142857142857144
      },
      "position" : {
        "x" : -6177.07373046875,
        "y" : 3767.876708984375
      },
      "selected" : false
    }, {
      "data" : {
        "id" : "1323",
        "symbol" : "HRS1",
        "Shoot_Descriptions" : "HRS1, myb-like transcription factor family protein",
        "PLAMOM_Match_Shoot" : "YES",
        "Causal_TF" : "myb-like transcription factor family protein\"",
        "Root_Descriptions" : "HRS1, myb-like transcription factor family protein",
        "Eccentricity" : 1,
        "Outdegree" : 1,
        "Target_PLAMOM" : "no",
        "PLAMOM_Match_Root" : "YES",
        "shared_name" : "AT1G13300",
        "Indegree" : 2,
        "Target_tRNA" : "AT1G13300",
        "t_RNA_motif_shoot" : "NO",
        "SUID" : 1323,
        "SelfLoops" : 0,
        "Target_TF" : "\"HRS1",
        "MolecularAttributes" : "TXNFACTOR",
        "AverageShortestPathLength" : 1.0,
        "selected" : false,
        "Name2" : "HRS1",
        "ClosenessCentrality" : 1.0,
        "Casual_Name" : "\"HRS1",
        "PartnerOfMultiEdgedNodePairs" : 0,
        "ClusteringCoefficient" : 0.0,
        "Target_Name" : "no",
        "Stress" : 8,
        "BetweennessCentrality" : 4.48E-6,
        "t_RNA_motif_Root" : "NO",
        "EdgeCount" : 3,
        "Causal_PLAMOM" : "AT1G13300",
        "name" : "AT1G13300",
        "Causal_tRNA" : "no",
        "IsSingleNode" : false,
        "briefDescription" : "myb-like transcription factor family protein",
        "NeighborhoodConnectivity" : 11.666666666666666
      },
      "position" : {
        "x" : -457.8033328769852,
        "y" : 665.509521484375
      },
      "selected" : false
    }, {
      "data" : {
        "id" : "1322",
        "symbol" : "AT3G57490",
        "Shoot_Descriptions" : "",
        "PLAMOM_Match_Shoot" : "",
        "Causal_TF" : "no",
        "Root_Descriptions" : "Ribosomal protein S5 family protein",
        "Eccentricity" : 11,
        "Outdegree" : 10,
        "Target_PLAMOM" : "",
        "PLAMOM_Match_Root" : "NO",
        "shared_name" : "AT3G57490",
        "Indegree" : 0,
        "Target_tRNA" : "",
        "t_RNA_motif_shoot" : "",
        "SUID" : 1322,
        "SelfLoops" : 0,
        "Target_TF" : "",
        "MolecularAttributes" : "protein_coding",
        "AverageShortestPathLength" : 5.6371134,
        "selected" : false,
        "Name2" : "AT3G57490",
        "ClosenessCentrality" : 0.17739576,
        "Casual_Name" : "Ribosomal protein S5 family protein",
        "PartnerOfMultiEdgedNodePairs" : 0,
        "ClusteringCoefficient" : 0.0,
        "Target_Name" : "",
        "Stress" : 0,
        "BetweennessCentrality" : 0.0,
        "t_RNA_motif_Root" : "NO",
        "EdgeCount" : 10,
        "Causal_PLAMOM" : "no",
        "name" : "AT3G57490",
        "Causal_tRNA" : "no",
        "IsSingleNode" : false,
        "briefDescription" : "Ribosomal protein S5 family protein",
        "NeighborhoodConnectivity" : 5.5
      },
      "position" : {
        "x" : -6310.17724609375,
        "y" : 299.05224609375
      },
      "selected" : false
    }, {
      "data" : {
        "id" : "1321",
        "symbol" : "GAPC2",
        "Shoot_Descriptions" : "GAPC-2, GAPC2, glyceraldehyde-3-phosphate dehydrogenase C2",
        "PLAMOM_Match_Shoot" : "YES",
        "Causal_TF" : "GAPC2",
        "Root_Descriptions" : "",
        "Eccentricity" : 11,
        "Outdegree" : 1,
        "Target_PLAMOM" : "AT1G13440",
        "PLAMOM_Match_Root" : "",
        "shared_name" : "AT1G13440",
        "Indegree" : 1,
        "Target_tRNA" : "no",
        "t_RNA_motif_shoot" : "NO",
        "SUID" : 1321,
        "SelfLoops" : 0,
        "Target_TF" : "GAPC2",
        "MolecularAttributes" : "METABOLIC",
        "AverageShortestPathLength" : 7.33402923,
        "selected" : false,
        "Name2" : "GAPC2",
        "ClosenessCentrality" : 0.1363507,
        "Casual_Name" : "\"GAPC-2",
        "PartnerOfMultiEdgedNodePairs" : 0,
        "ClusteringCoefficient" : 0.0,
        "Target_Name" : "\"GAPC-2",
        "Stress" : 43,
        "BetweennessCentrality" : 1.17E-5,
        "t_RNA_motif_Root" : "",
        "EdgeCount" : 2,
        "Causal_PLAMOM" : "AT1G13440",
        "name" : "AT1G13440",
        "Causal_tRNA" : "no",
        "IsSingleNode" : false,
        "briefDescription" : "glyceraldehyde-3-phosphate dehydrogenase C2",
        "NeighborhoodConnectivity" : 8.0
      },
      "position" : {
        "x" : 3191.288525033171,
        "y" : 3670.172607421875
      },
      "selected" : false
    }, {
      "data" : {
        "id" : "1320",
        "symbol" : "MYB305",
        "Shoot_Descriptions" : "",
        "PLAMOM_Match_Shoot" : "",
        "Causal_TF" : "MYB305",
        "Root_Descriptions" : "ATMYB71, MYB305, myb domain protein 305",
        "Eccentricity" : 14,
        "Outdegree" : 4,
        "Target_PLAMOM" : "indeterminate(ID)-domain 14\"",
        "PLAMOM_Match_Root" : "NO",
        "shared_name" : "AT3G24310",
        "Indegree" : 1,
        "Target_tRNA" : "AT1G68130",
        "t_RNA_motif_shoot" : "",
        "SUID" : 1320,
        "SelfLoops" : 0,
        "Target_TF" : "AT3G24310",
        "MolecularAttributes" : "TXNFACTOR",
        "AverageShortestPathLength" : 8.8946281,
        "selected" : false,
        "Name2" : "MYB305",
        "ClosenessCentrality" : 0.11242741,
        "Casual_Name" : "\"ATMYB71",
        "PartnerOfMultiEdgedNodePairs" : 0,
        "ClusteringCoefficient" : 0.0,
        "Target_Name" : "no",
        "Stress" : 1320,
        "BetweennessCentrality" : 2.8644E-4,
        "t_RNA_motif_Root" : "YES",
        "EdgeCount" : 5,
        "Causal_PLAMOM" : "no",
        "name" : "AT3G24310",
        "Causal_tRNA" : "AT3G24310",
        "IsSingleNode" : false,
        "briefDescription" : "myb domain protein 305",
        "NeighborhoodConnectivity" : 2.0
      },
      "position" : {
        "x" : -5516.5126953125,
        "y" : 2825.508056640625
      },
      "selected" : false
    }, {
      "data" : {
        "id" : "1319",
        "symbol" : "AT1G14130",
        "Shoot_Descriptions" : "2-oxoglutarate (2OG) and Fe(II)-dependent oxygenase superfamily protein",
        "PLAMOM_Match_Shoot" : "NO",
        "Causal_TF" : "",
        "Root_Descriptions" : "",
        "Eccentricity" : 0,
        "Outdegree" : 0,
        "Target_PLAMOM" : "myb domain protein 305\"",
        "PLAMOM_Match_Root" : "",
        "shared_name" : "AT1G14130",
        "Indegree" : 1,
        "Target_tRNA" : "AT3G24310",
        "t_RNA_motif_shoot" : "NO",
        "SUID" : 1319,
        "SelfLoops" : 0,
        "Target_TF" : "no",
        "MolecularAttributes" : "METABOLIC",
        "AverageShortestPathLength" : 0.0,
        "selected" : false,
        "Name2" : "AT1G14130",
        "ClosenessCentrality" : 0.0,
        "Casual_Name" : "",
        "PartnerOfMultiEdgedNodePairs" : 0,
        "ClusteringCoefficient" : 0.0,
        "Target_Name" : "no",
        "Stress" : 0,
        "BetweennessCentrality" : 0.0,
        "t_RNA_motif_Root" : "",
        "EdgeCount" : 1,
        "Causal_PLAMOM" : "",
        "name" : "AT1G14130",
        "Causal_tRNA" : "",
        "IsSingleNode" : false,
        "briefDescription" : "2-oxoglutarate (2OG) and Fe(II)-dependent oxygenase superfamily protein",
        "NeighborhoodConnectivity" : 5.0
      },
      "position" : {
        "x" : 5080.472666677832,
        "y" : -415.04296875
      },
      "selected" : false
    }, {
      "data" : {
        "id" : "1318",
        "symbol" : "AT2G39400",
        "Shoot_Descriptions" : "",
        "PLAMOM_Match_Shoot" : "",
        "Causal_TF" : "no",
        "Root_Descriptions" : "alpha/beta-Hydrolases superfamily protein",
        "Eccentricity" : 11,
        "Outdegree" : 3,
        "Target_PLAMOM" : "",
        "PLAMOM_Match_Root" : "YES",
        "shared_name" : "AT2G39400",
        "Indegree" : 0,
        "Target_tRNA" : "",
        "t_RNA_motif_shoot" : "",
        "SUID" : 1318,
        "SelfLoops" : 0,
        "Target_TF" : "",
        "MolecularAttributes" : "METABOLIC",
        "AverageShortestPathLength" : 5.55532359,
        "selected" : false,
        "Name2" : "AT2G39400",
        "ClosenessCentrality" : 0.18000752,
        "Casual_Name" : "alpha/beta-Hydrolases superfamily protein",
        "PartnerOfMultiEdgedNodePairs" : 0,
        "ClusteringCoefficient" : 0.0,
        "Target_Name" : "",
        "Stress" : 0,
        "BetweennessCentrality" : 0.0,
        "t_RNA_motif_Root" : "NO",
        "EdgeCount" : 3,
        "Causal_PLAMOM" : "AT2G39400",
        "name" : "AT2G39400",
        "Causal_tRNA" : "no",
        "IsSingleNode" : false,
        "briefDescription" : "alpha/beta-Hydrolases superfamily protein",
        "NeighborhoodConnectivity" : 11.333333333333334
      },
      "position" : {
        "x" : -5933.79638671875,
        "y" : 7.475549697875977
      },
      "selected" : false
    }, {
      "data" : {
        "id" : "1317",
        "symbol" : "AT1G14260",
        "Shoot_Descriptions" : "RING/FYVE/PHD zinc finger superfamily protein",
        "PLAMOM_Match_Shoot" : "NO",
        "Causal_TF" : "no",
        "Root_Descriptions" : "RING/FYVE/PHD zinc finger superfamily protein",
        "Eccentricity" : 10,
        "Outdegree" : 14,
        "Target_PLAMOM" : "phloem protein 2-A13\"",
        "PLAMOM_Match_Root" : "NO",
        "shared_name" : "AT1G14260",
        "Indegree" : 5,
        "Target_tRNA" : "no",
        "t_RNA_motif_shoot" : "NO",
        "SUID" : 1317,
        "SelfLoops" : 0,
        "Target_TF" : "no",
        "MolecularAttributes" : "protein_coding",
        "AverageShortestPathLength" : 4.58158996,
        "selected" : false,
        "Name2" : "AT1G14260",
        "ClosenessCentrality" : 0.21826484,
        "Casual_Name" : "RING/FYVE/PHD zinc finger superfamily protein",
        "PartnerOfMultiEdgedNodePairs" : 0,
        "ClusteringCoefficient" : 0.0,
        "Target_Name" : "no",
        "Stress" : 44480,
        "BetweennessCentrality" : 0.00992848,
        "t_RNA_motif_Root" : "NO",
        "EdgeCount" : 19,
        "Causal_PLAMOM" : "no",
        "name" : "AT1G14260",
        "Causal_tRNA" : "no",
        "IsSingleNode" : false,
        "briefDescription" : "RING/FYVE/PHD zinc finger superfamily protein",
        "NeighborhoodConnectivity" : 8.31578947368421
      },
      "position" : {
        "x" : -1327.581592154329,
        "y" : 3297.13330078125
      },
      "selected" : false
    }, {
      "data" : {
        "id" : "1316",
        "symbol" : "AAE5",
        "Shoot_Descriptions" : "",
        "PLAMOM_Match_Shoot" : "",
        "Causal_TF" : "acyl activating enzyme 5\"",
        "Root_Descriptions" : "AAE5, acyl activating enzyme 5",
        "Eccentricity" : 11,
        "Outdegree" : 2,
        "Target_PLAMOM" : "",
        "PLAMOM_Match_Root" : "NO",
        "shared_name" : "AT5G16370",
        "Indegree" : 0,
        "Target_tRNA" : "",
        "t_RNA_motif_shoot" : "",
        "SUID" : 1316,
        "SelfLoops" : 0,
        "Target_TF" : "",
        "MolecularAttributes" : "protein_coding",
        "AverageShortestPathLength" : 5.565762,
        "selected" : false,
        "Name2" : "AAE5",
        "ClosenessCentrality" : 0.17966992,
        "Casual_Name" : "\"AAE5",
        "PartnerOfMultiEdgedNodePairs" : 0,
        "ClusteringCoefficient" : 0.0,
        "Target_Name" : "",
        "Stress" : 0,
        "BetweennessCentrality" : 0.0,
        "t_RNA_motif_Root" : "NO",
        "EdgeCount" : 2,
        "Causal_PLAMOM" : "no",
        "name" : "AT5G16370",
        "Causal_tRNA" : "no",
        "IsSingleNode" : false,
        "briefDescription" : "acyl activating enzyme 5",
        "NeighborhoodConnectivity" : 15.5
      },
      "position" : {
        "x" : -6572.90771484375,
        "y" : 41.75220489501953
      },
      "selected" : false
    }, {
      "data" : {
        "id" : "1315",
        "symbol" : "ATRPAC42",
        "Shoot_Descriptions" : "AAC42, ATRPAC42, DNA-directed RNA polymerase family protein",
        "PLAMOM_Match_Shoot" : "NO",
        "Causal_TF" : "ATRPAC42",
        "Root_Descriptions" : "AAC42, ATRPAC42, DNA-directed RNA polymerase family protein",
        "Eccentricity" : 10,
        "Outdegree" : 5,
        "Target_PLAMOM" : "no",
        "PLAMOM_Match_Root" : "NO",
        "shared_name" : "AT1G60850",
        "Indegree" : 9,
        "Target_tRNA" : "no",
        "t_RNA_motif_shoot" : "NO",
        "SUID" : 1315,
        "SelfLoops" : 0,
        "Target_TF" : "ATRPAC42",
        "MolecularAttributes" : "protein_coding",
        "AverageShortestPathLength" : 4.99372385,
        "selected" : false,
        "Name2" : "ATRPAC42",
        "ClosenessCentrality" : 0.20025136,
        "Casual_Name" : "\"AAC42",
        "PartnerOfMultiEdgedNodePairs" : 0,
        "ClusteringCoefficient" : 0.0,
        "Target_Name" : "\"AAC42",
        "Stress" : 27688,
        "BetweennessCentrality" : 0.00751993,
        "t_RNA_motif_Root" : "NO",
        "EdgeCount" : 14,
        "Causal_PLAMOM" : "no",
        "name" : "AT1G60850",
        "Causal_tRNA" : "no",
        "IsSingleNode" : false,
        "briefDescription" : "DNA-directed RNA polymerase family protein",
        "NeighborhoodConnectivity" : 5.285714285714286
      },
      "position" : {
        "x" : -1594.191455435579,
        "y" : 3312.24365234375
      },
      "selected" : false
    }, {
      "data" : {
        "id" : "1314",
        "symbol" : "ROXY2",
        "Shoot_Descriptions" : "",
        "PLAMOM_Match_Shoot" : "",
        "Causal_TF" : "Thioredoxin superfamily protein\"",
        "Root_Descriptions" : "ROXY2, Thioredoxin superfamily protein",
        "Eccentricity" : 12,
        "Outdegree" : 3,
        "Target_PLAMOM" : "",
        "PLAMOM_Match_Root" : "NO",
        "shared_name" : "AT5G14070",
        "Indegree" : 0,
        "Target_tRNA" : "",
        "t_RNA_motif_shoot" : "",
        "SUID" : 1314,
        "SelfLoops" : 0,
        "Target_TF" : "",
        "MolecularAttributes" : "protein_coding",
        "AverageShortestPathLength" : 7.62217659,
        "selected" : false,
        "Name2" : "ROXY2",
        "ClosenessCentrality" : 0.13119612,
        "Casual_Name" : "\"ROXY2",
        "PartnerOfMultiEdgedNodePairs" : 0,
        "ClusteringCoefficient" : 0.0,
        "Target_Name" : "",
        "Stress" : 0,
        "BetweennessCentrality" : 0.0,
        "t_RNA_motif_Root" : "NO",
        "EdgeCount" : 3,
        "Causal_PLAMOM" : "no",
        "name" : "AT5G14070",
        "Causal_tRNA" : "no",
        "IsSingleNode" : false,
        "briefDescription" : "Thioredoxin superfamily protein",
        "NeighborhoodConnectivity" : 4.0
      },
      "position" : {
        "x" : -5448.2421875,
        "y" : 622.921630859375
      },
      "selected" : false
    }, {
      "data" : {
        "id" : "1313",
        "symbol" : "SAC52",
        "degree_layout" : 1,
        "Shoot_Descriptions" : "RPL10, RPL10A, SAC52, Ribosomal protein L16p/L10e family protein",
        "PLAMOM_Match_Shoot" : "YES",
        "Causal_TF" : "",
        "Root_Descriptions" : "",
        "Eccentricity" : 0,
        "Outdegree" : 0,
        "Target_PLAMOM" : "polyamine oxidase 5\"",
        "PLAMOM_Match_Root" : "",
        "shared_name" : "AT1G14320",
        "Indegree" : 3,
        "Target_tRNA" : "no",
        "t_RNA_motif_shoot" : "NO",
        "SUID" : 1313,
        "SelfLoops" : 0,
        "Target_TF" : "no",
        "MolecularAttributes" : "protein_coding",
        "AverageShortestPathLength" : 0.0,
        "selected" : false,
        "Name2" : "SAC52",
        "ClosenessCentrality" : 0.0,
        "Casual_Name" : "",
        "PartnerOfMultiEdgedNodePairs" : 0,
        "ClusteringCoefficient" : 0.0,
        "Target_Name" : "AT1G14320",
        "Stress" : 0,
        "BetweennessCentrality" : 0.0,
        "t_RNA_motif_Root" : "",
        "EdgeCount" : 3,
        "Causal_PLAMOM" : "",
        "name" : "AT1G14320",
        "Causal_tRNA" : "",
        "IsSingleNode" : false,
        "briefDescription" : "Ribosomal protein L16p/L10e family protein",
        "NeighborhoodConnectivity" : 8.333333333333334
      },
      "position" : {
        "x" : 4457.131786751921,
        "y" : 308.1650085449219
      },
      "selected" : false
    }, {
      "data" : {
        "id" : "1312",
        "symbol" : "AT3G01310",
        "degree_layout" : 13,
        "Shoot_Descriptions" : "",
        "PLAMOM_Match_Shoot" : "",
        "Causal_TF" : "no",
        "Root_Descriptions" : "Phosphoglycerate mutase-like family protein",
        "Eccentricity" : 9,
        "Outdegree" : 11,
        "Target_PLAMOM" : "AT5G65210",
        "PLAMOM_Match_Root" : "YES",
        "shared_name" : "AT3G01310",
        "Indegree" : 5,
        "Target_tRNA" : "AT3G01310",
        "t_RNA_motif_shoot" : "",
        "SUID" : 1312,
        "SelfLoops" : 0,
        "Target_TF" : "Phosphoglycerate mutase-like family protein",
        "MolecularAttributes" : "protein_coding",
        "AverageShortestPathLength" : 4.64853556,
        "selected" : false,
        "Name2" : "AT3G01310",
        "ClosenessCentrality" : 0.21512151,
        "Casual_Name" : "Phosphoglycerate mutase-like family protein",
        "PartnerOfMultiEdgedNodePairs" : 0,
        "ClusteringCoefficient" : 0.05,
        "Target_Name" : "no",
        "Stress" : 13430,
        "BetweennessCentrality" : 0.00300771,
        "t_RNA_motif_Root" : "NO",
        "EdgeCount" : 16,
        "Causal_PLAMOM" : "AT3G01310",
        "name" : "AT3G01310",
        "Causal_tRNA" : "no",
        "IsSingleNode" : false,
        "briefDescription" : "Phosphoglycerate mutase-like family protein",
        "NeighborhoodConnectivity" : 13.625
      },
      "position" : {
        "x" : -4798.48291015625,
        "y" : 3046.336669921875
      },
      "selected" : false
    }, {
      "data" : {
        "id" : "1311",
        "symbol" : "PAO5",
        "Shoot_Descriptions" : "",
        "PLAMOM_Match_Shoot" : "",
        "Causal_TF" : "PAO5",
        "Root_Descriptions" : "ATPAO5, PAO5, polyamine oxidase 5",
        "Eccentricity" : 14,
        "Outdegree" : 5,
        "Target_PLAMOM" : "no",
        "PLAMOM_Match_Root" : "NO",
        "shared_name" : "AT4G29720",
        "Indegree" : 1,
        "Target_tRNA" : "no",
        "t_RNA_motif_shoot" : "",
        "SUID" : 1311,
        "SelfLoops" : 0,
        "Target_TF" : "PAO5",
        "MolecularAttributes" : "METABOLIC",
        "AverageShortestPathLength" : 8.94375,
        "selected" : false,
        "Name2" : "PAO5",
        "ClosenessCentrality" : 0.11180992,
        "Casual_Name" : "\"ATPAO5",
        "PartnerOfMultiEdgedNodePairs" : 0,
        "ClusteringCoefficient" : 0.0,
        "Target_Name" : "\"ATPAO5",
        "Stress" : 11,
        "BetweennessCentrality" : 3.9E-6,
        "t_RNA_motif_Root" : "NO",
        "EdgeCount" : 6,
        "Causal_PLAMOM" : "no",
        "name" : "AT4G29720",
        "Causal_tRNA" : "no",
        "IsSingleNode" : false,
        "briefDescription" : "polyamine oxidase 5",
        "NeighborhoodConnectivity" : 5.833333333333333
      },
      "position" : {
        "x" : -6211.48486328125,
        "y" : 3253.31640625
      },
      "selected" : false
    }, {
      "data" : {
        "id" : "1310",
        "symbol" : "AT1G14650",
        "Shoot_Descriptions" : "SWAP (Suppressor-of-White-APricot)/surp domain-containing protein / ubiquitin family protein",
        "PLAMOM_Match_Shoot" : "NO",
        "Causal_TF" : "",
        "Root_Descriptions" : "",
        "Eccentricity" : 0,
        "Outdegree" : 0,
        "Target_PLAMOM" : "no",
        "PLAMOM_Match_Root" : "",
        "shared_name" : "AT1G14650",
        "Indegree" : 1,
        "Target_tRNA" : "no",
        "t_RNA_motif_shoot" : "NO",
        "SUID" : 1310,
        "SelfLoops" : 0,
        "Target_TF" : "no",
        "MolecularAttributes" : "protein_coding",
        "AverageShortestPathLength" : 0.0,
        "selected" : false,
        "Name2" : "AT1G14650",
        "ClosenessCentrality" : 0.0,
        "Casual_Name" : "",
        "PartnerOfMultiEdgedNodePairs" : 0,
        "ClusteringCoefficient" : 0.0,
        "Target_Name" : "SWAP (Suppressor-of-White-APricot)/surp domain-containing protein / ubiquitin family protein",
        "Stress" : 0,
        "BetweennessCentrality" : 0.0,
        "t_RNA_motif_Root" : "",
        "EdgeCount" : 1,
        "Causal_PLAMOM" : "",
        "name" : "AT1G14650",
        "Causal_tRNA" : "",
        "IsSingleNode" : false,
        "briefDescription" : "SWAP (Suppressor-of-White-APricot)/surp domain-containing protein / ubiquitin family protein",
        "NeighborhoodConnectivity" : 16.0
      },
      "position" : {
        "x" : 3422.725048470671,
        "y" : 148.30760192871094
      },
      "selected" : false
    }, {
      "data" : {
        "id" : "1309",
        "symbol" : "SULTR1;2",
        "Shoot_Descriptions" : "",
        "PLAMOM_Match_Shoot" : "",
        "Causal_TF" : "SULTR1;2",
        "Root_Descriptions" : "SEL1, SULTR1;2, sulfate transporter 1;2",
        "Eccentricity" : 1,
        "Outdegree" : 1,
        "Target_PLAMOM" : "AT1G78000",
        "PLAMOM_Match_Root" : "YES",
        "shared_name" : "AT1G78000",
        "Indegree" : 5,
        "Target_tRNA" : "no",
        "t_RNA_motif_shoot" : "",
        "SUID" : 1309,
        "SelfLoops" : 0,
        "Target_TF" : "SULTR1;2",
        "MolecularAttributes" : "protein_coding",
        "AverageShortestPathLength" : 1.0,
        "selected" : false,
        "Name2" : "SULTR1;2",
        "ClosenessCentrality" : 1.0,
        "Casual_Name" : "\"SEL1",
        "PartnerOfMultiEdgedNodePairs" : 0,
        "ClusteringCoefficient" : 0.0,
        "Target_Name" : "\"SEL1",
        "Stress" : 10,
        "BetweennessCentrality" : 5.97E-6,
        "t_RNA_motif_Root" : "NO",
        "EdgeCount" : 6,
        "Causal_PLAMOM" : "AT1G78000",
        "name" : "AT1G78000",
        "Causal_tRNA" : "no",
        "IsSingleNode" : false,
        "briefDescription" : "sulfate transporter 1;2",
        "NeighborhoodConnectivity" : 2.1666666666666665
      },
      "position" : {
        "x" : -6111.4658203125,
        "y" : 3911.932373046875
      },
      "selected" : false
    }, {
      "data" : {
        "id" : "1308",
        "symbol" : "AT1G15060",
        "Shoot_Descriptions" : "Uncharacterised conserved protein UCP031088, alpha/beta hydrolase",
        "PLAMOM_Match_Shoot" : "NO",
        "Causal_TF" : "",
        "Root_Descriptions" : "",
        "Eccentricity" : 0,
        "Outdegree" : 0,
        "Target_PLAMOM" : "sulfate transporter 1;2\"",
        "PLAMOM_Match_Root" : "",
        "shared_name" : "AT1G15060",
        "Indegree" : 1,
        "Target_tRNA" : "no",
        "t_RNA_motif_shoot" : "NO",
        "SUID" : 1308,
        "SelfLoops" : 0,
        "Target_TF" : "no",
        "MolecularAttributes" : "protein_coding",
        "AverageShortestPathLength" : 0.0,
        "selected" : false,
        "Name2" : "AT1G15060",
        "ClosenessCentrality" : 0.0,
        "Casual_Name" : "",
        "PartnerOfMultiEdgedNodePairs" : 0,
        "ClusteringCoefficient" : 0.0,
        "Target_Name" : "no",
        "Stress" : 0,
        "BetweennessCentrality" : 0.0,
        "t_RNA_motif_Root" : "",
        "EdgeCount" : 1,
        "Causal_PLAMOM" : "",
        "name" : "AT1G15060",
        "Causal_tRNA" : "",
        "IsSingleNode" : false,
        "briefDescription" : "alpha/beta hydrolase family protein",
        "NeighborhoodConnectivity" : 6.0
      },
      "position" : {
        "x" : 4529.367686209082,
        "y" : 917.39990234375
      },
      "selected" : false
    }, {
      "data" : {
        "id" : "1307",
        "symbol" : "AT1G15410",
        "Shoot_Descriptions" : "aspartate-glutamate racemase family",
        "PLAMOM_Match_Shoot" : "NO",
        "Causal_TF" : "",
        "Root_Descriptions" : "",
        "Eccentricity" : 0,
        "Outdegree" : 0,
        "Target_PLAMOM" : "no",
        "PLAMOM_Match_Root" : "",
        "shared_name" : "AT1G15410",
        "Indegree" : 2,
        "Target_tRNA" : "no",
        "t_RNA_motif_shoot" : "NO",
        "SUID" : 1307,
        "SelfLoops" : 0,
        "Target_TF" : "no",
        "MolecularAttributes" : "protein_coding",
        "AverageShortestPathLength" : 0.0,
        "selected" : false,
        "Name2" : "AT1G15410",
        "ClosenessCentrality" : 0.0,
        "Casual_Name" : "",
        "PartnerOfMultiEdgedNodePairs" : 0,
        "ClusteringCoefficient" : 0.0,
        "Target_Name" : "aspartate-glutamate racemase family",
        "Stress" : 0,
        "BetweennessCentrality" : 0.0,
        "t_RNA_motif_Root" : "",
        "EdgeCount" : 2,
        "Causal_PLAMOM" : "",
        "name" : "AT1G15410",
        "Causal_tRNA" : "",
        "IsSingleNode" : false,
        "briefDescription" : "aspartate-glutamate racemase family",
        "NeighborhoodConnectivity" : 20.0
      },
      "position" : {
        "x" : 4132.935069021582,
        "y" : -46.541404724121094
      },
      "selected" : false
    }, {
      "data" : {
        "id" : "1306",
        "symbol" : "SQD2",
        "Shoot_Descriptions" : "SQD2, sulfoquinovosyldiacylglycerol 2",
        "PLAMOM_Match_Shoot" : "NO",
        "Causal_TF" : "sulfoquinovosyldiacylglycerol 2\"",
        "Root_Descriptions" : "SQD2, sulfoquinovosyldiacylglycerol 2",
        "Eccentricity" : 10,
        "Outdegree" : 9,
        "Target_PLAMOM" : "acyl-CoA binding protein 1\"",
        "PLAMOM_Match_Root" : "NO",
        "shared_name" : "AT5G01220",
        "Indegree" : 6,
        "Target_tRNA" : "no",
        "t_RNA_motif_shoot" : "NO",
        "SUID" : 1306,
        "SelfLoops" : 0,
        "Target_TF" : "no",
        "MolecularAttributes" : "METABOLIC",
        "AverageShortestPathLength" : 5.62133891,
        "selected" : false,
        "Name2" : "SQD2",
        "ClosenessCentrality" : 0.17789356,
        "Casual_Name" : "\"SQD2",
        "PartnerOfMultiEdgedNodePairs" : 0,
        "ClusteringCoefficient" : 0.0,
        "Target_Name" : "no",
        "Stress" : 26749,
        "BetweennessCentrality" : 0.00587888,
        "t_RNA_motif_Root" : "NO",
        "EdgeCount" : 15,
        "Causal_PLAMOM" : "no",
        "name" : "AT5G01220",
        "Causal_tRNA" : "no",
        "IsSingleNode" : false,
        "briefDescription" : "sulfoquinovosyldiacylglycerol 2",
        "NeighborhoodConnectivity" : 6.133333333333334
      },
      "position" : {
        "x" : -1575.83740234375,
        "y" : 2793.2424212127926
      },
      "selected" : false
    }, {
      "data" : {
        "id" : "1305",
        "symbol" : "AT1G15480",
        "Shoot_Descriptions" : "Tetratricopeptide repeat (TPR)-like superfamily protein",
        "PLAMOM_Match_Shoot" : "NO",
        "Causal_TF" : "",
        "Root_Descriptions" : "",
        "Eccentricity" : 0,
        "Outdegree" : 0,
        "Target_PLAMOM" : "no",
        "PLAMOM_Match_Root" : "",
        "shared_name" : "AT1G15480",
        "Indegree" : 1,
        "Target_tRNA" : "no",
        "t_RNA_motif_shoot" : "NO",
        "SUID" : 1305,
        "SelfLoops" : 0,
        "Target_TF" : "Tetratricopeptide repeat (TPR)-like superfamily protein",
        "MolecularAttributes" : "protein_coding",
        "AverageShortestPathLength" : 0.0,
        "selected" : false,
        "Name2" : "AT1G15480",
        "ClosenessCentrality" : 0.0,
        "Casual_Name" : "",
        "PartnerOfMultiEdgedNodePairs" : 0,
        "ClusteringCoefficient" : 0.0,
        "Target_Name" : "no",
        "Stress" : 0,
        "BetweennessCentrality" : 0.0,
        "t_RNA_motif_Root" : "",
        "EdgeCount" : 1,
        "Causal_PLAMOM" : "",
        "name" : "AT1G15480",
        "Causal_tRNA" : "",
        "IsSingleNode" : false,
        "briefDescription" : "Tetratricopeptide repeat (TPR)-like superfamily protein",
        "NeighborhoodConnectivity" : 15.0
      },
      "position" : {
        "x" : 3242.9509381622074,
        "y" : -476.31439208984375
      },
      "selected" : false
    }, {
      "data" : {
        "id" : "1304",
        "symbol" : "AT5G56840",
        "Shoot_Descriptions" : "",
        "PLAMOM_Match_Shoot" : "",
        "Causal_TF" : "AT5G56840",
        "Root_Descriptions" : "myb-like transcription factor family protein",
        "Eccentricity" : 9,
        "Outdegree" : 8,
        "Target_PLAMOM" : "",
        "PLAMOM_Match_Root" : "NO",
        "shared_name" : "AT5G56840",
        "Indegree" : 0,
        "Target_tRNA" : "",
        "t_RNA_motif_shoot" : "",
        "SUID" : 1304,
        "SelfLoops" : 0,
        "Target_TF" : "",
        "MolecularAttributes" : "TXNFACTOR",
        "AverageShortestPathLength" : 4.53638254,
        "selected" : false,
        "Name2" : "AT5G56840",
        "ClosenessCentrality" : 0.22043996,
        "Casual_Name" : "myb-like transcription factor family protein",
        "PartnerOfMultiEdgedNodePairs" : 0,
        "ClusteringCoefficient" : 0.01785714,
        "Target_Name" : "",
        "Stress" : 0,
        "BetweennessCentrality" : 0.0,
        "t_RNA_motif_Root" : "NO",
        "EdgeCount" : 8,
        "Causal_PLAMOM" : "no",
        "name" : "AT5G56840",
        "Causal_tRNA" : "no",
        "IsSingleNode" : false,
        "briefDescription" : "myb-like transcription factor family protein",
        "NeighborhoodConnectivity" : 16.625
      },
      "position" : {
        "x" : -6160.38525390625,
        "y" : -33.653175354003906
      },
      "selected" : false
    }, {
      "data" : {
        "id" : "1303",
        "symbol" : "GA3OX1",
        "degree_layout" : 3,
        "Shoot_Descriptions" : "ATGA3OX1, GA3OX1, GA4, gibberellin 3-oxidase 1",
        "PLAMOM_Match_Shoot" : "NO",
        "Causal_TF" : "GA3OX1",
        "Root_Descriptions" : "ATGA3OX1, GA3OX1, GA4, gibberellin 3-oxidase 1",
        "Eccentricity" : 10,
        "Outdegree" : 4,
        "Target_PLAMOM" : "no",
        "PLAMOM_Match_Root" : "NO",
        "shared_name" : "AT1G15550",
        "Indegree" : 5,
        "Target_tRNA" : "no",
        "t_RNA_motif_shoot" : "NO",
        "SUID" : 1303,
        "SelfLoops" : 0,
        "Target_TF" : "GA3OX1",
        "MolecularAttributes" : "METABOLIC",
        "AverageShortestPathLength" : 5.42050209,
        "selected" : false,
        "Name2" : "GA3OX1",
        "ClosenessCentrality" : 0.18448475,
        "Casual_Name" : "\"ATGA3OX1",
        "PartnerOfMultiEdgedNodePairs" : 0,
        "ClusteringCoefficient" : 0.01388889,
        "Target_Name" : "\"ATGA3OX1",
        "Stress" : 26291,
        "BetweennessCentrality" : 0.00357057,
        "t_RNA_motif_Root" : "NO",
        "EdgeCount" : 9,
        "Causal_PLAMOM" : "no",
        "name" : "AT1G15550",
        "Causal_tRNA" : "no",
        "IsSingleNode" : false,
        "briefDescription" : "gibberellin 3-oxidase 1",
        "NeighborhoodConnectivity" : 22.666666666666668
      },
      "position" : {
        "x" : -1337.629443716829,
        "y" : 2995.945068359375
      },
      "selected" : false
    }, {
      "data" : {
        "id" : "1302",
        "symbol" : "GAT1_2.1",
        "Shoot_Descriptions" : "",
        "PLAMOM_Match_Shoot" : "",
        "Causal_TF" : "no",
        "Root_Descriptions" : "Class I glutamine amidotransferase-like superfamily protein",
        "Eccentricity" : 9,
        "Outdegree" : 15,
        "Target_PLAMOM" : "4-trisphosphate 5/6-kinase family protein\"",
        "PLAMOM_Match_Root" : "NO",
        "shared_name" : "AT1G15040",
        "Indegree" : 5,
        "Target_tRNA" : "no",
        "t_RNA_motif_shoot" : "",
        "SUID" : 1302,
        "SelfLoops" : 0,
        "Target_TF" : "no",
        "MolecularAttributes" : "protein_coding",
        "AverageShortestPathLength" : 4.4707113,
        "selected" : false,
        "Name2" : "GAT1_2.1",
        "ClosenessCentrality" : 0.22367805,
        "Casual_Name" : "Class I glutamine amidotransferase-like superfamily protein",
        "PartnerOfMultiEdgedNodePairs" : 0,
        "ClusteringCoefficient" : 0.00789474,
        "Target_Name" : "no",
        "Stress" : 33647,
        "BetweennessCentrality" : 0.00423487,
        "t_RNA_motif_Root" : "NO",
        "EdgeCount" : 20,
        "Causal_PLAMOM" : "no",
        "name" : "AT1G15040",
        "Causal_tRNA" : "no",
        "IsSingleNode" : false,
        "briefDescription" : "Class I glutamine amidotransferase-like superfamily protein",
        "NeighborhoodConnectivity" : 13.75
      },
      "position" : {
        "x" : -6636.4384765625,
        "y" : 2889.262939453125
      },
      "selected" : false
    }, {
      "data" : {
        "id" : "1301",
        "symbol" : "AT3G14260",
        "Shoot_Descriptions" : "",
        "PLAMOM_Match_Shoot" : "",
        "Causal_TF" : "no",
        "Root_Descriptions" : "Protein of unknown function (DUF567)",
        "Eccentricity" : 9,
        "Outdegree" : 8,
        "Target_PLAMOM" : "",
        "PLAMOM_Match_Root" : "NO",
        "shared_name" : "AT3G14260",
        "Indegree" : 0,
        "Target_tRNA" : "",
        "t_RNA_motif_shoot" : "",
        "SUID" : 1301,
        "SelfLoops" : 0,
        "Target_TF" : "",
        "MolecularAttributes" : "protein_coding",
        "AverageShortestPathLength" : 5.13900415,
        "selected" : false,
        "Name2" : "AT3G14260",
        "ClosenessCentrality" : 0.19459023,
        "Casual_Name" : "Protein of unknown function (DUF567)",
        "PartnerOfMultiEdgedNodePairs" : 0,
        "ClusteringCoefficient" : 0.0,
        "Target_Name" : "",
        "Stress" : 0,
        "BetweennessCentrality" : 0.0,
        "t_RNA_motif_Root" : "NO",
        "EdgeCount" : 8,
        "Causal_PLAMOM" : "no",
        "name" : "AT3G14260",
        "Causal_tRNA" : "no",
        "IsSingleNode" : false,
        "briefDescription" : "LURP-one-like protein (DUF567)",
        "NeighborhoodConnectivity" : 13.625
      },
      "position" : {
        "x" : -6256.9375,
        "y" : 100.65809631347656
      },
      "selected" : false
    }, {
      "data" : {
        "id" : "1300",
        "symbol" : "LSU2",
        "Shoot_Descriptions" : "",
        "PLAMOM_Match_Shoot" : "",
        "Causal_TF" : "response to low sulfur 2\"",
        "Root_Descriptions" : "LSU2, response to low sulfur 2",
        "Eccentricity" : 11,
        "Outdegree" : 3,
        "Target_PLAMOM" : "",
        "PLAMOM_Match_Root" : "NO",
        "shared_name" : "AT5G24660",
        "Indegree" : 0,
        "Target_tRNA" : "",
        "t_RNA_motif_shoot" : "",
        "SUID" : 1300,
        "SelfLoops" : 0,
        "Target_TF" : "",
        "MolecularAttributes" : "protein_coding",
        "AverageShortestPathLength" : 6.34719335,
        "selected" : false,
        "Name2" : "LSU2",
        "ClosenessCentrality" : 0.15754995,
        "Casual_Name" : "\"LSU2",
        "PartnerOfMultiEdgedNodePairs" : 0,
        "ClusteringCoefficient" : 0.0,
        "Target_Name" : "",
        "Stress" : 0,
        "BetweennessCentrality" : 0.0,
        "t_RNA_motif_Root" : "NO",
        "EdgeCount" : 3,
        "Causal_PLAMOM" : "no",
        "name" : "AT5G24660",
        "Causal_tRNA" : "no",
        "IsSingleNode" : false,
        "briefDescription" : "response to low sulfur 2",
        "NeighborhoodConnectivity" : 5.333333333333333
      },
      "position" : {
        "x" : -5673.10205078125,
        "y" : 535.4794921875
      },
      "selected" : false
    }, {
      "data" : {
        "id" : "1299",
        "symbol" : "AT1G16220",
        "Shoot_Descriptions" : "Protein phosphatase 2C family protein",
        "PLAMOM_Match_Shoot" : "NO",
        "Causal_TF" : "",
        "Root_Descriptions" : "",
        "Eccentricity" : 0,
        "Outdegree" : 0,
        "Target_PLAMOM" : "no",
        "PLAMOM_Match_Root" : "",
        "shared_name" : "AT1G16220",
        "Indegree" : 1,
        "Target_tRNA" : "no",
        "t_RNA_motif_shoot" : "NO",
        "SUID" : 1299,
        "SelfLoops" : 0,
        "Target_TF" : "Protein phosphatase 2C family protein",
        "MolecularAttributes" : "protein_coding",
        "AverageShortestPathLength" : 0.0,
        "selected" : false,
        "Name2" : "AT1G16220",
        "ClosenessCentrality" : 0.0,
        "Casual_Name" : "",
        "PartnerOfMultiEdgedNodePairs" : 0,
        "ClusteringCoefficient" : 0.0,
        "Target_Name" : "no",
        "Stress" : 0,
        "BetweennessCentrality" : 0.0,
        "t_RNA_motif_Root" : "",
        "EdgeCount" : 1,
        "Causal_PLAMOM" : "",
        "name" : "AT1G16220",
        "Causal_tRNA" : "",
        "IsSingleNode" : false,
        "briefDescription" : "Protein phosphatase 2C family protein",
        "NeighborhoodConnectivity" : 3.0
      },
      "position" : {
        "x" : 4545.888622689421,
        "y" : 687.053955078125
      },
      "selected" : false
    }, {
      "data" : {
        "id" : "1298",
        "symbol" : "PUB9",
        "degree_layout" : 5,
        "Shoot_Descriptions" : "",
        "PLAMOM_Match_Shoot" : "",
        "Causal_TF" : "PUB9",
        "Root_Descriptions" : "ATPUB9, PUB9, plant U-box 9",
        "Eccentricity" : 11,
        "Outdegree" : 11,
        "Target_PLAMOM" : "no",
        "PLAMOM_Match_Root" : "NO",
        "shared_name" : "AT3G07360",
        "Indegree" : 3,
        "Target_tRNA" : "no",
        "t_RNA_motif_shoot" : "",
        "SUID" : 1298,
        "SelfLoops" : 0,
        "Target_TF" : "PUB9",
        "MolecularAttributes" : "protein_coding",
        "AverageShortestPathLength" : 5.89958159,
        "selected" : false,
        "Name2" : "PUB9",
        "ClosenessCentrality" : 0.16950355,
        "Casual_Name" : "\"ATPUB9",
        "PartnerOfMultiEdgedNodePairs" : 0,
        "ClusteringCoefficient" : 0.0,
        "Target_Name" : "\"ATPUB9",
        "Stress" : 16637,
        "BetweennessCentrality" : 0.00390233,
        "t_RNA_motif_Root" : "NO",
        "EdgeCount" : 14,
        "Causal_PLAMOM" : "no",
        "name" : "AT3G07360",
        "Causal_tRNA" : "no",
        "IsSingleNode" : false,
        "briefDescription" : "plant U-box 9",
        "NeighborhoodConnectivity" : 6.071428571428571
      },
      "position" : {
        "x" : -6758.5791015625,
        "y" : 3325.880859375
      },
      "selected" : false
    }, {
      "data" : {
        "id" : "1297",
        "symbol" : "CYP72C1",
        "Shoot_Descriptions" : "CYP72C1, SOB7, cytochrome p450 72c1",
        "PLAMOM_Match_Shoot" : "NO",
        "Causal_TF" : "",
        "Root_Descriptions" : "",
        "Eccentricity" : 0,
        "Outdegree" : 0,
        "Target_PLAMOM" : "plant U-box 9\"",
        "PLAMOM_Match_Root" : "",
        "shared_name" : "AT1G17060",
        "Indegree" : 1,
        "Target_tRNA" : "no",
        "t_RNA_motif_shoot" : "NO",
        "SUID" : 1297,
        "SelfLoops" : 0,
        "Target_TF" : "no",
        "MolecularAttributes" : "protein_coding",
        "AverageShortestPathLength" : 0.0,
        "selected" : false,
        "Name2" : "CYP72C1",
        "ClosenessCentrality" : 0.0,
        "Casual_Name" : "",
        "PartnerOfMultiEdgedNodePairs" : 0,
        "ClusteringCoefficient" : 0.0,
        "Target_Name" : "no",
        "Stress" : 0,
        "BetweennessCentrality" : 0.0,
        "t_RNA_motif_Root" : "",
        "EdgeCount" : 1,
        "Causal_PLAMOM" : "",
        "name" : "AT1G17060",
        "Causal_tRNA" : "",
        "IsSingleNode" : false,
        "briefDescription" : "cytochrome p450 72c1",
        "NeighborhoodConnectivity" : 14.0
      },
      "position" : {
        "x" : 3825.609325814421,
        "y" : 761.4168701171875
      },
      "selected" : false
    }, {
      "data" : {
        "id" : "1296",
        "symbol" : "AT1G28260",
        "degree_layout" : 39,
        "Shoot_Descriptions" : "Telomerase activating protein Est1",
        "PLAMOM_Match_Shoot" : "NO",
        "Causal_TF" : "no",
        "Root_Descriptions" : "Telomerase activating protein Est1",
        "Eccentricity" : 8,
        "Outdegree" : 34,
        "Target_PLAMOM" : "no",
        "PLAMOM_Match_Root" : "NO",
        "shared_name" : "AT1G28260",
        "Indegree" : 19,
        "Target_tRNA" : "no",
        "t_RNA_motif_shoot" : "NO",
        "SUID" : 1296,
        "SelfLoops" : 0,
        "Target_TF" : "no",
        "MolecularAttributes" : "protein_coding",
        "AverageShortestPathLength" : 3.44769874,
        "selected" : false,
        "Name2" : "AT1G28260",
        "ClosenessCentrality" : 0.29004854,
        "Casual_Name" : "Telomerase activating protein Est1",
        "PartnerOfMultiEdgedNodePairs" : 1,
        "ClusteringCoefficient" : 0.01508296,
        "Target_Name" : "Telomerase activating protein Est1",
        "Stress" : 203669,
        "BetweennessCentrality" : 0.04341402,
        "t_RNA_motif_Root" : "NO",
        "EdgeCount" : 53,
        "Causal_PLAMOM" : "no",
        "name" : "AT1G28260",
        "Causal_tRNA" : "no",
        "IsSingleNode" : false,
        "briefDescription" : "Telomerase activating protein Est1",
        "NeighborhoodConnectivity" : 9.788461538461538
      },
      "position" : {
        "x" : -996.8552127597977,
        "y" : 3180.69873046875
      },
      "selected" : false
    }, {
      "data" : {
        "id" : "1295",
        "symbol" : "GSTU24",
        "degree_layout" : 3,
        "Shoot_Descriptions" : "ATGSTU24, GST, GSTU24, glutathione S-transferase TAU 24",
        "PLAMOM_Match_Shoot" : "NO",
        "Causal_TF" : "GST",
        "Root_Descriptions" : "",
        "Eccentricity" : 9,
        "Outdegree" : 2,
        "Target_PLAMOM" : "no",
        "PLAMOM_Match_Root" : "",
        "shared_name" : "AT1G17170",
        "Indegree" : 1,
        "Target_tRNA" : "no",
        "t_RNA_motif_shoot" : "NO",
        "SUID" : 1295,
        "SelfLoops" : 0,
        "Target_TF" : "GST",
        "MolecularAttributes" : "protein_coding",
        "AverageShortestPathLength" : 5.28242678,
        "selected" : false,
        "Name2" : "GSTU24",
        "ClosenessCentrality" : 0.18930693,
        "Casual_Name" : "\"ATGSTU24",
        "PartnerOfMultiEdgedNodePairs" : 0,
        "ClusteringCoefficient" : 0.0,
        "Target_Name" : "\"ATGSTU24",
        "Stress" : 988,
        "BetweennessCentrality" : 1.0343E-4,
        "t_RNA_motif_Root" : "",
        "EdgeCount" : 3,
        "Causal_PLAMOM" : "no",
        "name" : "AT1G17170",
        "Causal_tRNA" : "no",
        "IsSingleNode" : false,
        "briefDescription" : "glutathione S-transferase TAU 24",
        "NeighborhoodConnectivity" : 47.333333333333336
      },
      "position" : {
        "x" : 3960.141552376921,
        "y" : 3098.279296875
      },
      "selected" : false
    }, {
      "data" : {
        "id" : "1294",
        "symbol" : "AT1G71970",
        "Shoot_Descriptions" : "",
        "PLAMOM_Match_Shoot" : "",
        "Causal_TF" : "no",
        "Root_Descriptions" : "unknown protein; FUNCTIONS IN: molecular_function unknown; INVOLVED IN: biological_process unknown; LOCATED IN: chloroplast; EXPRESSED IN: 22 plant structures; EXPRESSED DURING: 13 growth stages; BEST Arabidopsis thaliana protein match is: unknown protein (TAIR:AT1G22680.1); Has 58 Blast hits to 58 proteins in 13 species: Archae - 0; Bacteria - 0; Metazoa - 4; Fungi - 0; Plants - 49; Viruses - 0; Other Eukaryotes - 5 (source: NCBI BLink).",
        "Eccentricity" : 9,
        "Outdegree" : 4,
        "Target_PLAMOM" : "",
        "PLAMOM_Match_Root" : "NO",
        "shared_name" : "AT1G71970",
        "Indegree" : 0,
        "Target_tRNA" : "",
        "t_RNA_motif_shoot" : "",
        "SUID" : 1294,
        "SelfLoops" : 0,
        "Target_TF" : "",
        "MolecularAttributes" : "protein_coding",
        "AverageShortestPathLength" : 5.34303534,
        "selected" : false,
        "Name2" : "AT1G71970",
        "ClosenessCentrality" : 0.18715953,
        "Casual_Name" : "unknown protein; FUNCTIONS IN: molecular_function unknown; INVOLVED IN: biological_process unknown; LOCATED IN: chloroplast; EXPRESSED IN: 22 plant structures; EXPRESSED DURING: 13 growth stages; BEST Arabidopsis thaliana protein match is: unknown protein (TAIR:AT1G22680.1); Has 58 Blast hits to 58 proteins in 13 species: Archae - 0; Bacteria - 0; Metazoa - 4; Fungi - 0; Plants - 49; Viruses - 0; Other Eukaryotes - 5 (source: NCBI BLink).",
        "PartnerOfMultiEdgedNodePairs" : 0,
        "ClusteringCoefficient" : 0.0,
        "Target_Name" : "",
        "Stress" : 0,
        "BetweennessCentrality" : 0.0,
        "t_RNA_motif_Root" : "NO",
        "EdgeCount" : 4,
        "Causal_PLAMOM" : "no",
        "name" : "AT1G71970",
        "Causal_tRNA" : "no",
        "IsSingleNode" : false,
        "briefDescription" : "hypothetical protein",
        "NeighborhoodConnectivity" : 6.5
      },
      "position" : {
        "x" : -6196.22021484375,
        "y" : 723.6434936523438
      },
      "selected" : false
    }, {
      "data" : {
        "id" : "1293",
        "symbol" : "ATARCA",
        "Shoot_Descriptions" : "ATARCA, RACK1A, RACK1A_AT, Transducin/WD40 repeat-like superfamily protein",
        "PLAMOM_Match_Shoot" : "YES",
        "Causal_TF" : "",
        "Root_Descriptions" : "",
        "Eccentricity" : 0,
        "Outdegree" : 0,
        "Target_PLAMOM" : "AT1G18080",
        "PLAMOM_Match_Root" : "",
        "shared_name" : "AT1G18080",
        "Indegree" : 1,
        "Target_tRNA" : "no",
        "t_RNA_motif_shoot" : "NO",
        "SUID" : 1293,
        "SelfLoops" : 0,
        "Target_TF" : "RACK1A",
        "MolecularAttributes" : "protein_coding",
        "AverageShortestPathLength" : 0.0,
        "selected" : false,
        "Name2" : "ATARCA",
        "ClosenessCentrality" : 0.0,
        "Casual_Name" : "",
        "PartnerOfMultiEdgedNodePairs" : 0,
        "ClusteringCoefficient" : 0.0,
        "Target_Name" : "\"ATARCA",
        "Stress" : 0,
        "BetweennessCentrality" : 0.0,
        "t_RNA_motif_Root" : "",
        "EdgeCount" : 1,
        "Causal_PLAMOM" : "",
        "name" : "AT1G18080",
        "Causal_tRNA" : "",
        "IsSingleNode" : false,
        "briefDescription" : "Transducin/WD40 repeat-like superfamily protein",
        "NeighborhoodConnectivity" : 4.0
      },
      "position" : {
        "x" : 3761.721142220671,
        "y" : 841.7733154296875
      },
      "selected" : false
    }, {
      "data" : {
        "id" : "1292",
        "symbol" : "AT1G30200",
        "Shoot_Descriptions" : "",
        "PLAMOM_Match_Shoot" : "",
        "Causal_TF" : "no",
        "Root_Descriptions" : "F-box family protein",
        "Eccentricity" : 13,
        "Outdegree" : 8,
        "Target_PLAMOM" : "",
        "PLAMOM_Match_Root" : "NO",
        "shared_name" : "AT1G30200",
        "Indegree" : 0,
        "Target_tRNA" : "",
        "t_RNA_motif_shoot" : "",
        "SUID" : 1292,
        "SelfLoops" : 0,
        "Target_TF" : "",
        "MolecularAttributes" : "protein_coding",
        "AverageShortestPathLength" : 7.79296066,
        "selected" : false,
        "Name2" : "AT1G30200",
        "ClosenessCentrality" : 0.12832094,
        "Casual_Name" : "F-box family protein",
        "PartnerOfMultiEdgedNodePairs" : 0,
        "ClusteringCoefficient" : 0.0,
        "Target_Name" : "",
        "Stress" : 0,
        "BetweennessCentrality" : 0.0,
        "t_RNA_motif_Root" : "NO",
        "EdgeCount" : 8,
        "Causal_PLAMOM" : "no",
        "name" : "AT1G30200",
        "Causal_tRNA" : "no",
        "IsSingleNode" : false,
        "briefDescription" : "F-box family protein",
        "NeighborhoodConnectivity" : 4.75
      },
      "position" : {
        "x" : -6692.41162109375,
        "y" : 484.85443115234375
      },
      "selected" : false
    }, {
      "data" : {
        "id" : "1291",
        "symbol" : "AT1G18540",
        "Shoot_Descriptions" : "Ribosomal protein L6 family protein",
        "PLAMOM_Match_Shoot" : "YES",
        "Causal_TF" : "",
        "Root_Descriptions" : "",
        "Eccentricity" : 0,
        "Outdegree" : 0,
        "Target_PLAMOM" : "AT1G18540",
        "PLAMOM_Match_Root" : "",
        "shared_name" : "AT1G18540",
        "Indegree" : 1,
        "Target_tRNA" : "no",
        "t_RNA_motif_shoot" : "NO",
        "SUID" : 1291,
        "SelfLoops" : 0,
        "Target_TF" : "no",
        "MolecularAttributes" : "protein_coding",
        "AverageShortestPathLength" : 0.0,
        "selected" : false,
        "Name2" : "AT1G18540",
        "ClosenessCentrality" : 0.0,
        "Casual_Name" : "",
        "PartnerOfMultiEdgedNodePairs" : 0,
        "ClusteringCoefficient" : 0.0,
        "Target_Name" : "Ribosomal protein L6 family protein",
        "Stress" : 0,
        "BetweennessCentrality" : 0.0,
        "t_RNA_motif_Root" : "",
        "EdgeCount" : 1,
        "Causal_PLAMOM" : "",
        "name" : "AT1G18540",
        "Causal_tRNA" : "",
        "IsSingleNode" : false,
        "briefDescription" : "Ribosomal protein L6 family protein",
        "NeighborhoodConnectivity" : 8.0
      },
      "position" : {
        "x" : 3011.6479596465824,
        "y" : 466.9003601074219
      },
      "selected" : false
    }, {
      "data" : {
        "id" : "1290",
        "symbol" : "AT2G44000",
        "degree_layout" : 2,
        "Shoot_Descriptions" : "",
        "PLAMOM_Match_Shoot" : "",
        "Causal_TF" : "no",
        "Root_Descriptions" : "Late embryogenesis abundant (LEA) hydroxyproline-rich glycoprotein family",
        "Eccentricity" : 11,
        "Outdegree" : 5,
        "Target_PLAMOM" : "no",
        "PLAMOM_Match_Root" : "NO",
        "shared_name" : "AT2G44000",
        "Indegree" : 7,
        "Target_tRNA" : "no",
        "t_RNA_motif_shoot" : "",
        "SUID" : 1290,
        "SelfLoops" : 0,
        "Target_TF" : "no",
        "MolecularAttributes" : "protein_coding",
        "AverageShortestPathLength" : 6.20502092,
        "selected" : false,
        "Name2" : "AT2G44000",
        "ClosenessCentrality" : 0.16115981,
        "Casual_Name" : "Late embryogenesis abundant (LEA) hydroxyproline-rich glycoprotein family",
        "PartnerOfMultiEdgedNodePairs" : 0,
        "ClusteringCoefficient" : 0.0,
        "Target_Name" : "Late embryogenesis abundant (LEA) hydroxyproline-rich glycoprotein family",
        "Stress" : 12282,
        "BetweennessCentrality" : 0.00252336,
        "t_RNA_motif_Root" : "NO",
        "EdgeCount" : 12,
        "Causal_PLAMOM" : "no",
        "name" : "AT2G44000",
        "Causal_tRNA" : "no",
        "IsSingleNode" : false,
        "briefDescription" : "Late embryogenesis abundant (LEA) hydroxyproline-rich glycoprotein family",
        "NeighborhoodConnectivity" : 7.25
      },
      "position" : {
        "x" : -6236.6025390625,
        "y" : 2948.864501953125
      },
      "selected" : false
    }, {
      "data" : {
        "id" : "1289",
        "symbol" : "AT1G18850",
        "Shoot_Descriptions" : "unknown protein; FUNCTIONS IN: molecular_function unknown; INVOLVED IN: biological_process unknown; LOCATED IN: nucleolus; EXPRESSED IN: 22 plant structures; EXPRESSED DURING: 13 growth stages; Has 40 Blast hits to 40 proteins in 13 species: Archae - 0; Bacteria - 0; Metazoa - 0; Fungi - 0; Plants - 40; Viruses - 0; Other Eukaryotes - 0 (source: NCBI BLink).",
        "PLAMOM_Match_Shoot" : "NO",
        "Causal_TF" : "",
        "Root_Descriptions" : "",
        "Eccentricity" : 0,
        "Outdegree" : 0,
        "Target_PLAMOM" : "no",
        "PLAMOM_Match_Root" : "",
        "shared_name" : "AT1G18850",
        "Indegree" : 3,
        "Target_tRNA" : "no",
        "t_RNA_motif_shoot" : "NO",
        "SUID" : 1289,
        "SelfLoops" : 0,
        "Target_TF" : "no",
        "MolecularAttributes" : "protein_coding",
        "AverageShortestPathLength" : 0.0,
        "selected" : false,
        "Name2" : "AT1G18850",
        "ClosenessCentrality" : 0.0,
        "Casual_Name" : "",
        "PartnerOfMultiEdgedNodePairs" : 0,
        "ClusteringCoefficient" : 0.0,
        "Target_Name" : "unknown protein; FUNCTIONS IN: molecular_function unknown; INVOLVED IN: biological_process unknown; LOCATED IN: nucleolus; EXPRESSED IN: 22 plant structures; EXPRESSED DURING: 13 growth stages; Has 40 Blast hits to 40 proteins in 13 species: Archae - 0; Bacteria - 0; Metazoa - 0; Fungi - 0; Plants - 40; Viruses - 0; Other Eukaryotes - 0 (source: NCBI BLink).",
        "Stress" : 0,
        "BetweennessCentrality" : 0.0,
        "t_RNA_motif_Root" : "",
        "EdgeCount" : 3,
        "Causal_PLAMOM" : "",
        "name" : "AT1G18850",
        "Causal_tRNA" : "",
        "IsSingleNode" : false,
        "briefDescription" : "hypothetical protein",
        "NeighborhoodConnectivity" : 10.666666666666666
      },
      "position" : {
        "x" : 4405.424755501921,
        "y" : -12.063819885253906
      },
      "selected" : false
    }, {
      "data" : {
        "id" : "1288",
        "symbol" : "AT1G66930",
        "degree_layout" : 1,
        "Shoot_Descriptions" : "",
        "PLAMOM_Match_Shoot" : "",
        "Causal_TF" : "no",
        "Root_Descriptions" : "Protein kinase superfamily protein",
        "Eccentricity" : 9,
        "Outdegree" : 6,
        "Target_PLAMOM" : "no",
        "PLAMOM_Match_Root" : "NO",
        "shared_name" : "AT1G66930",
        "Indegree" : 9,
        "Target_tRNA" : "no",
        "t_RNA_motif_shoot" : "",
        "SUID" : 1288,
        "SelfLoops" : 0,
        "Target_TF" : "no",
        "MolecularAttributes" : "protein_coding",
        "AverageShortestPathLength" : 4.80962343,
        "selected" : false,
        "Name2" : "AT1G66930",
        "ClosenessCentrality" : 0.20791649,
        "Casual_Name" : "Protein kinase superfamily protein",
        "PartnerOfMultiEdgedNodePairs" : 0,
        "ClusteringCoefficient" : 0.0047619,
        "Target_Name" : "Protein kinase superfamily protein",
        "Stress" : 24628,
        "BetweennessCentrality" : 0.00388561,
        "t_RNA_motif_Root" : "NO",
        "EdgeCount" : 15,
        "Causal_PLAMOM" : "no",
        "name" : "AT1G66930",
        "Causal_tRNA" : "no",
        "IsSingleNode" : false,
        "briefDescription" : "Protein kinase superfamily protein",
        "NeighborhoodConnectivity" : 7.666666666666667
      },
      "position" : {
        "x" : -6303.96630859375,
        "y" : 3205.742431640625
      },
      "selected" : false
    }, {
      "data" : {
        "id" : "1287",
        "symbol" : "AT3G46650",
        "Shoot_Descriptions" : "",
        "PLAMOM_Match_Shoot" : "",
        "Causal_TF" : "no",
        "Root_Descriptions" : "UDP-Glycosyltransferase superfamily protein",
        "Eccentricity" : 11,
        "Outdegree" : 5,
        "Target_PLAMOM" : "",
        "PLAMOM_Match_Root" : "NO",
        "shared_name" : "AT3G46650",
        "Indegree" : 0,
        "Target_tRNA" : "",
        "t_RNA_motif_shoot" : "",
        "SUID" : 1287,
        "SelfLoops" : 0,
        "Target_TF" : "",
        "MolecularAttributes" : "protein_coding",
        "AverageShortestPathLength" : 5.94572025,
        "selected" : false,
        "Name2" : "AT3G46650",
        "ClosenessCentrality" : 0.1681882,
        "Casual_Name" : "UDP-Glycosyltransferase superfamily protein",
        "PartnerOfMultiEdgedNodePairs" : 0,
        "ClusteringCoefficient" : 0.0,
        "Target_Name" : "",
        "Stress" : 0,
        "BetweennessCentrality" : 0.0,
        "t_RNA_motif_Root" : "NO",
        "EdgeCount" : 5,
        "Causal_PLAMOM" : "no",
        "name" : "AT3G46650",
        "Causal_tRNA" : "no",
        "IsSingleNode" : false,
        "briefDescription" : "UDP-Glycosyltransferase superfamily protein",
        "NeighborhoodConnectivity" : 8.6
      },
      "position" : {
        "x" : -5901.06982421875,
        "y" : -144.2696075439453
      },
      "selected" : false
    }, {
      "data" : {
        "id" : "1286",
        "symbol" : "MYB68",
        "Shoot_Descriptions" : "",
        "PLAMOM_Match_Shoot" : "",
        "Causal_TF" : "MYB68",
        "Root_Descriptions" : "ATMYB68, MYB68, myb domain protein 68",
        "Eccentricity" : 8,
        "Outdegree" : 24,
        "Target_PLAMOM" : "no",
        "PLAMOM_Match_Root" : "NO",
        "shared_name" : "AT5G65790",
        "Indegree" : 5,
        "Target_tRNA" : "no",
        "t_RNA_motif_shoot" : "",
        "SUID" : 1286,
        "SelfLoops" : 0,
        "Target_TF" : "MYB68",
        "MolecularAttributes" : "TXNFACTOR",
        "AverageShortestPathLength" : 3.85355649,
        "selected" : false,
        "Name2" : "MYB68",
        "ClosenessCentrality" : 0.25950054,
        "Casual_Name" : "\"ATMYB68",
        "PartnerOfMultiEdgedNodePairs" : 0,
        "ClusteringCoefficient" : 0.00246305,
        "Target_Name" : "\"ATMYB68",
        "Stress" : 99645,
        "BetweennessCentrality" : 0.01959893,
        "t_RNA_motif_Root" : "NO",
        "EdgeCount" : 29,
        "Causal_PLAMOM" : "no",
        "name" : "AT5G65790",
        "Causal_tRNA" : "no",
        "IsSingleNode" : false,
        "briefDescription" : "myb domain protein 68",
        "NeighborhoodConnectivity" : 8.655172413793103
      },
      "position" : {
        "x" : -6641.037109375,
        "y" : 3357.56884765625
      },
      "selected" : false
    }, {
      "data" : {
        "id" : "1285",
        "symbol" : "NFD5",
        "Shoot_Descriptions" : "NFD5, pentatricopeptide (PPR) repeat-containing protein",
        "PLAMOM_Match_Shoot" : "NO",
        "Causal_TF" : "pentatricopeptide (PPR) repeat-containing protein\"",
        "Root_Descriptions" : "",
        "Eccentricity" : 11,
        "Outdegree" : 1,
        "Target_PLAMOM" : "SHM5",
        "PLAMOM_Match_Root" : "",
        "shared_name" : "AT1G19520",
        "Indegree" : 2,
        "Target_tRNA" : "Pyridoxal phosphate (PLP)-dependent transferases superfamily protein\"",
        "t_RNA_motif_shoot" : "NO",
        "SUID" : 1285,
        "SelfLoops" : 0,
        "Target_TF" : "no",
        "MolecularAttributes" : "protein_coding",
        "AverageShortestPathLength" : 6.0251046,
        "selected" : false,
        "Name2" : "NFD5",
        "ClosenessCentrality" : 0.16597222,
        "Casual_Name" : "\"NFD5",
        "PartnerOfMultiEdgedNodePairs" : 0,
        "ClusteringCoefficient" : 0.0,
        "Target_Name" : "no",
        "Stress" : 823,
        "BetweennessCentrality" : 1.0009E-4,
        "t_RNA_motif_Root" : "",
        "EdgeCount" : 3,
        "Causal_PLAMOM" : "no",
        "name" : "AT1G19520",
        "Causal_tRNA" : "no",
        "IsSingleNode" : false,
        "briefDescription" : "pentatricopeptide (PPR) repeat-containing protein",
        "NeighborhoodConnectivity" : 37.0
      },
      "position" : {
        "x" : 3569.218212533171,
        "y" : 3480.873779296875
      },
      "selected" : false
    }, {
      "data" : {
        "id" : "1284",
        "symbol" : "EDA36",
        "Shoot_Descriptions" : "",
        "PLAMOM_Match_Shoot" : "",
        "Causal_TF" : "EDA37",
        "Root_Descriptions" : "EDA36, EDA37, SHM5, Pyridoxal phosphate (PLP)-dependent transferases superfamily protein",
        "Eccentricity" : 9,
        "Outdegree" : 14,
        "Target_PLAMOM" : "",
        "PLAMOM_Match_Root" : "NO",
        "shared_name" : "AT4G13890",
        "Indegree" : 0,
        "Target_tRNA" : "",
        "t_RNA_motif_shoot" : "",
        "SUID" : 1284,
        "SelfLoops" : 0,
        "Target_TF" : "",
        "MolecularAttributes" : "METABOLIC",
        "AverageShortestPathLength" : 4.32016632,
        "selected" : false,
        "Name2" : "EDA36",
        "ClosenessCentrality" : 0.23147257,
        "Casual_Name" : "\"EDA36",
        "PartnerOfMultiEdgedNodePairs" : 0,
        "ClusteringCoefficient" : 0.00549451,
        "Target_Name" : "",
        "Stress" : 0,
        "BetweennessCentrality" : 0.0,
        "t_RNA_motif_Root" : "NO",
        "EdgeCount" : 14,
        "Causal_PLAMOM" : "no",
        "name" : "AT4G13890",
        "Causal_tRNA" : "no",
        "IsSingleNode" : false,
        "briefDescription" : "Pyridoxal phosphate (PLP)-dependent transferases superfamily protein",
        "NeighborhoodConnectivity" : 10.642857142857142
      },
      "position" : {
        "x" : -6250.0595703125,
        "y" : 204.27561950683594
      },
      "selected" : false
    }, {
      "data" : {
        "id" : "1283",
        "symbol" : "MDH",
        "Shoot_Descriptions" : "MDH, malate dehydrogenase",
        "PLAMOM_Match_Shoot" : "YES",
        "Causal_TF" : "malate dehydrogenase\"",
        "Root_Descriptions" : "MDH, malate dehydrogenase",
        "Eccentricity" : 11,
        "Outdegree" : 3,
        "Target_PLAMOM" : "no",
        "PLAMOM_Match_Root" : "YES",
        "shared_name" : "AT3G47520",
        "Indegree" : 1,
        "Target_tRNA" : "AT3G47520",
        "t_RNA_motif_shoot" : "NO",
        "SUID" : 1283,
        "SelfLoops" : 0,
        "Target_TF" : "\"MDH",
        "MolecularAttributes" : "METABOLIC",
        "AverageShortestPathLength" : 6.71757322,
        "selected" : false,
        "Name2" : "MDH",
        "ClosenessCentrality" : 0.14886328,
        "Casual_Name" : "\"MDH",
        "PartnerOfMultiEdgedNodePairs" : 0,
        "ClusteringCoefficient" : 0.0,
        "Target_Name" : "no",
        "Stress" : 2596,
        "BetweennessCentrality" : 4.895E-4,
        "t_RNA_motif_Root" : "NO",
        "EdgeCount" : 4,
        "Causal_PLAMOM" : "AT3G47520",
        "name" : "AT3G47520",
        "Causal_tRNA" : "no",
        "IsSingleNode" : false,
        "briefDescription" : "malate dehydrogenase",
        "NeighborhoodConnectivity" : 22.5
      },
      "position" : {
        "x" : -399.3862186191727,
        "y" : -49.923709869384766
      },
      "selected" : false
    }, {
      "data" : {
        "id" : "1282",
        "symbol" : "AT1G19660",
        "Shoot_Descriptions" : "Wound-responsive family protein",
        "PLAMOM_Match_Shoot" : "YES",
        "Causal_TF" : "",
        "Root_Descriptions" : "",
        "Eccentricity" : 0,
        "Outdegree" : 0,
        "Target_PLAMOM" : "TDT",
        "PLAMOM_Match_Root" : "",
        "shared_name" : "AT1G19660",
        "Indegree" : 3,
        "Target_tRNA" : "tonoplast dicarboxylate transporter\"",
        "t_RNA_motif_shoot" : "NO",
        "SUID" : 1282,
        "SelfLoops" : 0,
        "Target_TF" : "AT1G19660",
        "MolecularAttributes" : "protein_coding",
        "AverageShortestPathLength" : 0.0,
        "selected" : false,
        "Name2" : "AT1G19660",
        "ClosenessCentrality" : 0.0,
        "Casual_Name" : "",
        "PartnerOfMultiEdgedNodePairs" : 0,
        "ClusteringCoefficient" : 0.0,
        "Target_Name" : "no",
        "Stress" : 0,
        "BetweennessCentrality" : 0.0,
        "t_RNA_motif_Root" : "",
        "EdgeCount" : 3,
        "Causal_PLAMOM" : "",
        "name" : "AT1G19660",
        "Causal_tRNA" : "",
        "IsSingleNode" : false,
        "briefDescription" : "Wound-responsive family protein",
        "NeighborhoodConnectivity" : 5.666666666666667
      },
      "position" : {
        "x" : 4139.260752615332,
        "y" : -399.71954345703125
      },
      "selected" : false
    }, {
      "data" : {
        "id" : "1281",
        "symbol" : "AT5G04250",
        "Shoot_Descriptions" : "Cysteine proteinases superfamily protein",
        "PLAMOM_Match_Shoot" : "NO",
        "Causal_TF" : "no",
        "Root_Descriptions" : "Cysteine proteinases superfamily protein",
        "Eccentricity" : 10,
        "Outdegree" : 6,
        "Target_PLAMOM" : "no",
        "PLAMOM_Match_Root" : "NO",
        "shared_name" : "AT5G04250",
        "Indegree" : 1,
        "Target_tRNA" : "no",
        "t_RNA_motif_shoot" : "NO",
        "SUID" : 1281,
        "SelfLoops" : 0,
        "Target_TF" : "Cysteine proteinases superfamily protein",
        "MolecularAttributes" : "protein_coding",
        "AverageShortestPathLength" : 5.217119,
        "selected" : false,
        "Name2" : "AT5G04250",
        "ClosenessCentrality" : 0.19167667,
        "Casual_Name" : "Cysteine proteinases superfamily protein",
        "PartnerOfMultiEdgedNodePairs" : 0,
        "ClusteringCoefficient" : 0.02380952,
        "Target_Name" : "no",
        "Stress" : 3262,
        "BetweennessCentrality" : 4.9877E-4,
        "t_RNA_motif_Root" : "NO",
        "EdgeCount" : 7,
        "Causal_PLAMOM" : "no",
        "name" : "AT5G04250",
        "Causal_tRNA" : "no",
        "IsSingleNode" : false,
        "briefDescription" : "Cysteine proteinases superfamily protein",
        "NeighborhoodConnectivity" : 13.285714285714286
      },
      "position" : {
        "x" : -1174.672168326204,
        "y" : 2858.645751953125
      },
      "selected" : false
    }, {
      "data" : {
        "id" : "1280",
        "symbol" : "TDT",
        "Shoot_Descriptions" : "",
        "PLAMOM_Match_Shoot" : "",
        "Causal_TF" : "ATTDT",
        "Root_Descriptions" : "ATSDAT, ATTDT, TDT, tonoplast dicarboxylate transporter",
        "Eccentricity" : 10,
        "Outdegree" : 6,
        "Target_PLAMOM" : "",
        "PLAMOM_Match_Root" : "YES",
        "shared_name" : "AT5G47560",
        "Indegree" : 0,
        "Target_tRNA" : "",
        "t_RNA_motif_shoot" : "",
        "SUID" : 1280,
        "SelfLoops" : 0,
        "Target_TF" : "",
        "MolecularAttributes" : "protein_coding",
        "AverageShortestPathLength" : 5.36625514,
        "selected" : false,
        "Name2" : "TDT",
        "ClosenessCentrality" : 0.18634969,
        "Casual_Name" : "\"ATSDAT",
        "PartnerOfMultiEdgedNodePairs" : 0,
        "ClusteringCoefficient" : 0.0,
        "Target_Name" : "",
        "Stress" : 0,
        "BetweennessCentrality" : 0.0,
        "t_RNA_motif_Root" : "NO",
        "EdgeCount" : 6,
        "Causal_PLAMOM" : "AT5G47560",
        "name" : "AT5G47560",
        "Causal_tRNA" : "no",
        "IsSingleNode" : false,
        "briefDescription" : "tonoplast dicarboxylate transporter",
        "NeighborhoodConnectivity" : 18.0
      },
      "position" : {
        "x" : -5756.90771484375,
        "y" : 224.1102752685547
      },
      "selected" : false
    }, {
      "data" : {
        "id" : "1279",
        "symbol" : "AT1G19880",
        "Shoot_Descriptions" : "Regulator of chromosome condensation (RCC1) family protein",
        "PLAMOM_Match_Shoot" : "NO",
        "Causal_TF" : "no",
        "Root_Descriptions" : "",
        "Eccentricity" : 9,
        "Outdegree" : 3,
        "Target_PLAMOM" : "no",
        "PLAMOM_Match_Root" : "",
        "shared_name" : "AT1G19880",
        "Indegree" : 4,
        "Target_tRNA" : "no",
        "t_RNA_motif_shoot" : "NO",
        "SUID" : 1279,
        "SelfLoops" : 0,
        "Target_TF" : "no",
        "MolecularAttributes" : "protein_coding",
        "AverageShortestPathLength" : 4.71129707,
        "selected" : false,
        "Name2" : "AT1G19880",
        "ClosenessCentrality" : 0.21225577,
        "Casual_Name" : "Regulator of chromosome condensation (RCC1) family protein",
        "PartnerOfMultiEdgedNodePairs" : 0,
        "ClusteringCoefficient" : 0.0,
        "Target_Name" : "Regulator of chromosome condensation (RCC1) family protein",
        "Stress" : 21433,
        "BetweennessCentrality" : 0.00479492,
        "t_RNA_motif_Root" : "",
        "EdgeCount" : 7,
        "Causal_PLAMOM" : "no",
        "name" : "AT1G19880",
        "Causal_tRNA" : "no",
        "IsSingleNode" : false,
        "briefDescription" : "Regulator of chromosome condensation (RCC1) family protein",
        "NeighborhoodConnectivity" : 22.285714285714285
      },
      "position" : {
        "x" : 3607.014599251921,
        "y" : 2812.469970703125
      },
      "selected" : false
    }, {
      "data" : {
        "id" : "1278",
        "symbol" : "AT2G32560",
        "Shoot_Descriptions" : "",
        "PLAMOM_Match_Shoot" : "",
        "Causal_TF" : "no",
        "Root_Descriptions" : "F-box family protein",
        "Eccentricity" : 10,
        "Outdegree" : 9,
        "Target_PLAMOM" : "no",
        "PLAMOM_Match_Root" : "NO",
        "shared_name" : "AT2G32560",
        "Indegree" : 4,
        "Target_tRNA" : "no",
        "t_RNA_motif_shoot" : "",
        "SUID" : 1278,
        "SelfLoops" : 0,
        "Target_TF" : "no",
        "MolecularAttributes" : "protein_coding",
        "AverageShortestPathLength" : 5.16945607,
        "selected" : false,
        "Name2" : "AT2G32560",
        "ClosenessCentrality" : 0.19344395,
        "Casual_Name" : "F-box family protein",
        "PartnerOfMultiEdgedNodePairs" : 0,
        "ClusteringCoefficient" : 0.0,
        "Target_Name" : "F-box family protein",
        "Stress" : 11534,
        "BetweennessCentrality" : 0.00305656,
        "t_RNA_motif_Root" : "NO",
        "EdgeCount" : 13,
        "Causal_PLAMOM" : "no",
        "name" : "AT2G32560",
        "Causal_tRNA" : "no",
        "IsSingleNode" : false,
        "briefDescription" : "F-box family protein",
        "NeighborhoodConnectivity" : 11.615384615384615
      },
      "position" : {
        "x" : -6611.5126953125,
        "y" : 2774.9814453125
      },
      "selected" : false
    }, {
      "data" : {
        "id" : "1277",
        "symbol" : "AT1G76580",
        "Shoot_Descriptions" : "",
        "PLAMOM_Match_Shoot" : "",
        "Causal_TF" : "AT1G76580",
        "Root_Descriptions" : "Squamosa promoter-binding protein-like (SBP domain) transcription factor family protein",
        "Eccentricity" : 10,
        "Outdegree" : 4,
        "Target_PLAMOM" : "",
        "PLAMOM_Match_Root" : "NO",
        "shared_name" : "AT1G76580",
        "Indegree" : 0,
        "Target_tRNA" : "",
        "t_RNA_motif_shoot" : "",
        "SUID" : 1277,
        "SelfLoops" : 0,
        "Target_TF" : "",
        "MolecularAttributes" : "TXNFACTOR",
        "AverageShortestPathLength" : 5.2776618,
        "selected" : false,
        "Name2" : "AT1G76580",
        "ClosenessCentrality" : 0.18947785,
        "Casual_Name" : "Squamosa promoter-binding protein-like (SBP domain) transcription factor family protein",
        "PartnerOfMultiEdgedNodePairs" : 0,
        "ClusteringCoefficient" : 0.0,
        "Target_Name" : "",
        "Stress" : 0,
        "BetweennessCentrality" : 0.0,
        "t_RNA_motif_Root" : "NO",
        "EdgeCount" : 4,
        "Causal_PLAMOM" : "no",
        "name" : "AT1G76580",
        "Causal_tRNA" : "no",
        "IsSingleNode" : false,
        "briefDescription" : "Squamosa promoter-binding protein-like (SBP domain) transcription factor family protein",
        "NeighborhoodConnectivity" : 28.25
      },
      "position" : {
        "x" : -5919.60595703125,
        "y" : -141.2344512939453
      },
      "selected" : false
    }, {
      "data" : {
        "id" : "1276",
        "symbol" : "CAT3",
        "Shoot_Descriptions" : "ATCAT3, CAT3, SEN2, catalase 3",
        "PLAMOM_Match_Shoot" : "YES",
        "Causal_TF" : "",
        "Root_Descriptions" : "",
        "Eccentricity" : 0,
        "Outdegree" : 0,
        "Target_PLAMOM" : "L-fucokinase/GDP-L-fucose pyrophosphorylase\"",
        "PLAMOM_Match_Root" : "",
        "shared_name" : "AT1G20620",
        "Indegree" : 1,
        "Target_tRNA" : "no",
        "t_RNA_motif_shoot" : "NO",
        "SUID" : 1276,
        "SelfLoops" : 0,
        "Target_TF" : "no",
        "MolecularAttributes" : "METABOLIC",
        "AverageShortestPathLength" : 0.0,
        "selected" : false,
        "Name2" : "CAT3",
        "ClosenessCentrality" : 0.0,
        "Casual_Name" : "",
        "PartnerOfMultiEdgedNodePairs" : 0,
        "ClusteringCoefficient" : 0.0,
        "Target_Name" : "AT1G20620",
        "Stress" : 0,
        "BetweennessCentrality" : 0.0,
        "t_RNA_motif_Root" : "",
        "EdgeCount" : 1,
        "Causal_PLAMOM" : "",
        "name" : "AT1G20620",
        "Causal_tRNA" : "",
        "IsSingleNode" : false,
        "briefDescription" : "catalase 3",
        "NeighborhoodConnectivity" : 19.0
      },
      "position" : {
        "x" : 3880.6350201934574,
        "y" : 554.603515625
      },
      "selected" : false
    }, {
      "data" : {
        "id" : "1275",
        "symbol" : "AT1G21780",
        "degree_layout" : 5,
        "Shoot_Descriptions" : "BTB/POZ domain-containing protein",
        "PLAMOM_Match_Shoot" : "NO",
        "Causal_TF" : "no",
        "Root_Descriptions" : "",
        "Eccentricity" : 9,
        "Outdegree" : 1,
        "Target_PLAMOM" : "no",
        "PLAMOM_Match_Root" : "",
        "shared_name" : "AT1G21780",
        "Indegree" : 8,
        "Target_tRNA" : "AT1G21780",
        "t_RNA_motif_shoot" : "YES",
        "SUID" : 1275,
        "SelfLoops" : 0,
        "Target_TF" : "no",
        "MolecularAttributes" : "protein_coding",
        "AverageShortestPathLength" : 5.10460251,
        "selected" : false,
        "Name2" : "AT1G21780",
        "ClosenessCentrality" : 0.19590164,
        "Casual_Name" : "BTB/POZ domain-containing protein",
        "PartnerOfMultiEdgedNodePairs" : 0,
        "ClusteringCoefficient" : 0.01388889,
        "Target_Name" : "BTB/POZ domain-containing protein",
        "Stress" : 17963,
        "BetweennessCentrality" : 0.00368505,
        "t_RNA_motif_Root" : "",
        "EdgeCount" : 9,
        "Causal_PLAMOM" : "no",
        "name" : "AT1G21780",
        "Causal_tRNA" : "AT1G21780",
        "IsSingleNode" : false,
        "briefDescription" : "BTB/POZ domain-containing protein",
        "NeighborhoodConnectivity" : 25.555555555555557
      },
      "position" : {
        "x" : 3448.022900033171,
        "y" : 2889.835693359375
      },
      "selected" : false
    }, {
      "data" : {
        "id" : "1274",
        "symbol" : "CID10",
        "degree_layout" : 22,
        "Shoot_Descriptions" : "",
        "PLAMOM_Match_Shoot" : "",
        "Causal_TF" : "CTC-interacting domain 10\"",
        "Root_Descriptions" : "CID10, CTC-interacting domain 10",
        "Eccentricity" : 9,
        "Outdegree" : 20,
        "Target_PLAMOM" : "no",
        "PLAMOM_Match_Root" : "NO",
        "shared_name" : "AT3G49390",
        "Indegree" : 6,
        "Target_tRNA" : "no",
        "t_RNA_motif_shoot" : "",
        "SUID" : 1274,
        "SelfLoops" : 0,
        "Target_TF" : "\"CID10",
        "MolecularAttributes" : "protein_coding",
        "AverageShortestPathLength" : 3.86192469,
        "selected" : false,
        "Name2" : "CID10",
        "ClosenessCentrality" : 0.25893824,
        "Casual_Name" : "\"CID10",
        "PartnerOfMultiEdgedNodePairs" : 0,
        "ClusteringCoefficient" : 0.02307692,
        "Target_Name" : "no",
        "Stress" : 74634,
        "BetweennessCentrality" : 0.01479753,
        "t_RNA_motif_Root" : "NO",
        "EdgeCount" : 26,
        "Causal_PLAMOM" : "no",
        "name" : "AT3G49390",
        "Causal_tRNA" : "no",
        "IsSingleNode" : false,
        "briefDescription" : "CTC-interacting domain 10",
        "NeighborhoodConnectivity" : 11.461538461538462
      },
      "position" : {
        "x" : -6705.0654296875,
        "y" : 3142.9697265625
      },
      "selected" : false
    }, {
      "data" : {
        "id" : "1273",
        "symbol" : "AT3G57630",
        "Shoot_Descriptions" : "",
        "PLAMOM_Match_Shoot" : "",
        "Causal_TF" : "no",
        "Root_Descriptions" : "exostosin family protein",
        "Eccentricity" : 9,
        "Outdegree" : 9,
        "Target_PLAMOM" : "4-trisphosphate 5/6-kinase family protein\"",
        "PLAMOM_Match_Root" : "NO",
        "shared_name" : "AT3G57630",
        "Indegree" : 2,
        "Target_tRNA" : "no",
        "t_RNA_motif_shoot" : "",
        "SUID" : 1273,
        "SelfLoops" : 0,
        "Target_TF" : "no",
        "MolecularAttributes" : "protein_coding",
        "AverageShortestPathLength" : 4.62552301,
        "selected" : false,
        "Name2" : "AT3G57630",
        "ClosenessCentrality" : 0.21619177,
        "Casual_Name" : "exostosin family protein",
        "PartnerOfMultiEdgedNodePairs" : 0,
        "ClusteringCoefficient" : 0.00909091,
        "Target_Name" : "no",
        "Stress" : 8696,
        "BetweennessCentrality" : 0.00161356,
        "t_RNA_motif_Root" : "NO",
        "EdgeCount" : 11,
        "Causal_PLAMOM" : "no",
        "name" : "AT3G57630",
        "Causal_tRNA" : "no",
        "IsSingleNode" : false,
        "briefDescription" : "exostosin family protein",
        "NeighborhoodConnectivity" : 13.636363636363637
      },
      "position" : {
        "x" : -6532.12939453125,
        "y" : 2913.342041015625
      },
      "selected" : false
    }, {
      "data" : {
        "id" : "1272",
        "symbol" : "AT4G03500",
        "Shoot_Descriptions" : "",
        "PLAMOM_Match_Shoot" : "",
        "Causal_TF" : "no",
        "Root_Descriptions" : "Ankyrin repeat family protein",
        "Eccentricity" : 13,
        "Outdegree" : 6,
        "Target_PLAMOM" : "",
        "PLAMOM_Match_Root" : "NO",
        "shared_name" : "AT4G03500",
        "Indegree" : 0,
        "Target_tRNA" : "",
        "t_RNA_motif_shoot" : "",
        "SUID" : 1272,
        "SelfLoops" : 0,
        "Target_TF" : "",
        "MolecularAttributes" : "protein_coding",
        "AverageShortestPathLength" : 8.02277433,
        "selected" : false,
        "Name2" : "AT4G03500",
        "ClosenessCentrality" : 0.12464516,
        "Casual_Name" : "Ankyrin repeat family protein",
        "PartnerOfMultiEdgedNodePairs" : 0,
        "ClusteringCoefficient" : 0.0,
        "Target_Name" : "",
        "Stress" : 0,
        "BetweennessCentrality" : 0.0,
        "t_RNA_motif_Root" : "NO",
        "EdgeCount" : 6,
        "Causal_PLAMOM" : "no",
        "name" : "AT4G03500",
        "Causal_tRNA" : "no",
        "IsSingleNode" : false,
        "briefDescription" : "Ankyrin repeat family protein",
        "NeighborhoodConnectivity" : 11.166666666666666
      },
      "position" : {
        "x" : -5723.435546875,
        "y" : 253.99368286132812
      },
      "selected" : false
    }, {
      "data" : {
        "id" : "1271",
        "symbol" : "AT1G22270",
        "Shoot_Descriptions" : "Trm112p-like protein",
        "PLAMOM_Match_Shoot" : "YES",
        "Causal_TF" : "",
        "Root_Descriptions" : "",
        "Eccentricity" : 0,
        "Outdegree" : 0,
        "Target_PLAMOM" : "AT1G22270",
        "PLAMOM_Match_Root" : "",
        "shared_name" : "AT1G22270",
        "Indegree" : 2,
        "Target_tRNA" : "no",
        "t_RNA_motif_shoot" : "NO",
        "SUID" : 1271,
        "SelfLoops" : 0,
        "Target_TF" : "no",
        "MolecularAttributes" : "protein_coding",
        "AverageShortestPathLength" : 0.0,
        "selected" : false,
        "Name2" : "AT1G22270",
        "ClosenessCentrality" : 0.0,
        "Casual_Name" : "",
        "PartnerOfMultiEdgedNodePairs" : 0,
        "ClusteringCoefficient" : 0.0,
        "Target_Name" : "Trm112p-like protein",
        "Stress" : 0,
        "BetweennessCentrality" : 0.0,
        "t_RNA_motif_Root" : "",
        "EdgeCount" : 2,
        "Causal_PLAMOM" : "",
        "name" : "AT1G22270",
        "Causal_tRNA" : "",
        "IsSingleNode" : false,
        "briefDescription" : "Trm112p-like protein",
        "NeighborhoodConnectivity" : 3.5
      },
      "position" : {
        "x" : 4886.811045584082,
        "y" : 204.53689575195312
      },
      "selected" : false
    }, {
      "data" : {
        "id" : "1270",
        "symbol" : "AT5G48970",
        "Shoot_Descriptions" : "",
        "PLAMOM_Match_Shoot" : "",
        "Causal_TF" : "no",
        "Root_Descriptions" : "Mitochondrial substrate carrier family protein",
        "Eccentricity" : 1,
        "Outdegree" : 1,
        "Target_PLAMOM" : "",
        "PLAMOM_Match_Root" : "NO",
        "shared_name" : "AT5G48970",
        "Indegree" : 0,
        "Target_tRNA" : "",
        "t_RNA_motif_shoot" : "",
        "SUID" : 1270,
        "SelfLoops" : 0,
        "Target_TF" : "",
        "MolecularAttributes" : "protein_coding",
        "AverageShortestPathLength" : 1.0,
        "selected" : false,
        "Name2" : "AT5G48970",
        "ClosenessCentrality" : 1.0,
        "Casual_Name" : "Mitochondrial substrate carrier family protein",
        "PartnerOfMultiEdgedNodePairs" : 0,
        "ClusteringCoefficient" : 0.0,
        "Target_Name" : "",
        "Stress" : 0,
        "BetweennessCentrality" : 0.0,
        "t_RNA_motif_Root" : "NO",
        "EdgeCount" : 1,
        "Causal_PLAMOM" : "no",
        "name" : "AT5G48970",
        "Causal_tRNA" : "no",
        "IsSingleNode" : false,
        "briefDescription" : "Mitochondrial substrate carrier family protein",
        "NeighborhoodConnectivity" : 2.0
      },
      "position" : {
        "x" : -4823.66650390625,
        "y" : 277.1971740722656
      },
      "selected" : false
    }, {
      "data" : {
        "id" : "1269",
        "symbol" : "NAC011",
        "Shoot_Descriptions" : "",
        "PLAMOM_Match_Shoot" : "",
        "Causal_TF" : "NAC011",
        "Root_Descriptions" : "ANAC011, NAC011, NAC domain containing protein 11",
        "Eccentricity" : 1,
        "Outdegree" : 1,
        "Target_PLAMOM" : "",
        "PLAMOM_Match_Root" : "NO",
        "shared_name" : "AT1G32510",
        "Indegree" : 0,
        "Target_tRNA" : "",
        "t_RNA_motif_shoot" : "",
        "SUID" : 1269,
        "SelfLoops" : 0,
        "Target_TF" : "",
        "MolecularAttributes" : "TXNFACTOR",
        "AverageShortestPathLength" : 1.0,
        "selected" : false,
        "Name2" : "NAC011",
        "ClosenessCentrality" : 1.0,
        "Casual_Name" : "\"ANAC011",
        "PartnerOfMultiEdgedNodePairs" : 0,
        "ClusteringCoefficient" : 0.0,
        "Target_Name" : "",
        "Stress" : 0,
        "BetweennessCentrality" : 0.0,
        "t_RNA_motif_Root" : "NO",
        "EdgeCount" : 1,
        "Causal_PLAMOM" : "no",
        "name" : "AT1G32510",
        "Causal_tRNA" : "no",
        "IsSingleNode" : false,
        "briefDescription" : "NAC domain containing protein 11",
        "NeighborhoodConnectivity" : 10.0
      },
      "position" : {
        "x" : -6026.73193359375,
        "y" : -671.6905517578125
      },
      "selected" : false
    }, {
      "data" : {
        "id" : "1268",
        "symbol" : "PFL",
        "Shoot_Descriptions" : "PFL, PFL1, RPS18A, Ribosomal protein S13/S18 family",
        "PLAMOM_Match_Shoot" : "YES",
        "Causal_TF" : "",
        "Root_Descriptions" : "",
        "Eccentricity" : 0,
        "Outdegree" : 0,
        "Target_PLAMOM" : "AT1G22780",
        "PLAMOM_Match_Root" : "",
        "shared_name" : "AT1G22780",
        "Indegree" : 10,
        "Target_tRNA" : "no",
        "t_RNA_motif_shoot" : "NO",
        "SUID" : 1268,
        "SelfLoops" : 0,
        "Target_TF" : "PFL1",
        "MolecularAttributes" : "protein_coding",
        "AverageShortestPathLength" : 0.0,
        "selected" : false,
        "Name2" : "PFL",
        "ClosenessCentrality" : 0.0,
        "Casual_Name" : "",
        "PartnerOfMultiEdgedNodePairs" : 0,
        "ClusteringCoefficient" : 0.0,
        "Target_Name" : "\"PFL",
        "Stress" : 0,
        "BetweennessCentrality" : 0.0,
        "t_RNA_motif_Root" : "",
        "EdgeCount" : 10,
        "Causal_PLAMOM" : "",
        "name" : "AT1G22780",
        "Causal_tRNA" : "",
        "IsSingleNode" : false,
        "briefDescription" : "Ribosomal protein S13/S18 family",
        "NeighborhoodConnectivity" : 3.9
      },
      "position" : {
        "x" : 4112.564892220671,
        "y" : -347.591064453125
      },
      "selected" : false
    }, {
      "data" : {
        "id" : "1267",
        "symbol" : "AT1G63860",
        "Shoot_Descriptions" : "Disease resistance protein (TIR-NBS-LRR class) family",
        "PLAMOM_Match_Shoot" : "NO",
        "Causal_TF" : "no",
        "Root_Descriptions" : "Disease resistance protein (TIR-NBS-LRR class) family",
        "Eccentricity" : 11,
        "Outdegree" : 8,
        "Target_PLAMOM" : "no",
        "PLAMOM_Match_Root" : "NO",
        "shared_name" : "AT1G63860",
        "Indegree" : 1,
        "Target_tRNA" : "no",
        "t_RNA_motif_shoot" : "NO",
        "SUID" : 1267,
        "SelfLoops" : 0,
        "Target_TF" : "no",
        "MolecularAttributes" : "protein_coding",
        "AverageShortestPathLength" : 5.92916667,
        "selected" : false,
        "Name2" : "AT1G63860",
        "ClosenessCentrality" : 0.16865777,
        "Casual_Name" : "Disease resistance protein (TIR-NBS-LRR class) family",
        "PartnerOfMultiEdgedNodePairs" : 0,
        "ClusteringCoefficient" : 0.0,
        "Target_Name" : "Disease resistance protein (TIR-NBS-LRR class) family",
        "Stress" : 1151,
        "BetweennessCentrality" : 2.7417E-4,
        "t_RNA_motif_Root" : "NO",
        "EdgeCount" : 9,
        "Causal_PLAMOM" : "no",
        "name" : "AT1G63860",
        "Causal_tRNA" : "no",
        "IsSingleNode" : false,
        "briefDescription" : "Disease resistance protein (TIR-NBS-LRR class) family",
        "NeighborhoodConnectivity" : 29.22222222222222
      },
      "position" : {
        "x" : -1268.905810904329,
        "y" : 3162.4140625
      },
      "selected" : false
    }, {
      "data" : {
        "id" : "1266",
        "symbol" : "BUBR1",
        "Shoot_Descriptions" : "",
        "PLAMOM_Match_Shoot" : "",
        "Causal_TF" : "BUB1-related (BUB1: budding uninhibited by benzymidazol 1)\"",
        "Root_Descriptions" : "BUBR1, BUB1-related (BUB1: budding uninhibited by benzymidazol 1)",
        "Eccentricity" : 1,
        "Outdegree" : 1,
        "Target_PLAMOM" : "",
        "PLAMOM_Match_Root" : "NO",
        "shared_name" : "AT2G33560",
        "Indegree" : 0,
        "Target_tRNA" : "",
        "t_RNA_motif_shoot" : "",
        "SUID" : 1266,
        "SelfLoops" : 0,
        "Target_TF" : "",
        "MolecularAttributes" : "protein_coding",
        "AverageShortestPathLength" : 1.0,
        "selected" : false,
        "Name2" : "BUBR1",
        "ClosenessCentrality" : 1.0,
        "Casual_Name" : "\"BUBR1",
        "PartnerOfMultiEdgedNodePairs" : 0,
        "ClusteringCoefficient" : 0.0,
        "Target_Name" : "",
        "Stress" : 0,
        "BetweennessCentrality" : 0.0,
        "t_RNA_motif_Root" : "YES",
        "EdgeCount" : 1,
        "Causal_PLAMOM" : "no",
        "name" : "AT2G33560",
        "Causal_tRNA" : "AT2G33560",
        "IsSingleNode" : false,
        "briefDescription" : "BUB1-related (BUB1: budding uninhibited by benzymidazol 1)",
        "NeighborhoodConnectivity" : 10.0
      },
      "position" : {
        "x" : -5987.837890625,
        "y" : -721.3017578125
      },
      "selected" : false
    }, {
      "data" : {
        "id" : "1265",
        "symbol" : "AT2G34170",
        "Shoot_Descriptions" : "",
        "PLAMOM_Match_Shoot" : "",
        "Causal_TF" : "no",
        "Root_Descriptions" : "Protein of unknown function (DUF688)",
        "Eccentricity" : 13,
        "Outdegree" : 2,
        "Target_PLAMOM" : "no",
        "PLAMOM_Match_Root" : "NO",
        "shared_name" : "AT2G34170",
        "Indegree" : 3,
        "Target_tRNA" : "no",
        "t_RNA_motif_shoot" : "",
        "SUID" : 1265,
        "SelfLoops" : 0,
        "Target_TF" : "no",
        "MolecularAttributes" : "protein_coding",
        "AverageShortestPathLength" : 8.05230126,
        "selected" : false,
        "Name2" : "AT2G34170",
        "ClosenessCentrality" : 0.1241881,
        "Casual_Name" : "Protein of unknown function (DUF688)",
        "PartnerOfMultiEdgedNodePairs" : 0,
        "ClusteringCoefficient" : 0.0,
        "Target_Name" : "Protein of unknown function (DUF688)",
        "Stress" : 5046,
        "BetweennessCentrality" : 0.00144245,
        "t_RNA_motif_Root" : "NO",
        "EdgeCount" : 5,
        "Causal_PLAMOM" : "no",
        "name" : "AT2G34170",
        "Causal_tRNA" : "no",
        "IsSingleNode" : false,
        "briefDescription" : "hypothetical protein (DUF688)",
        "NeighborhoodConnectivity" : 7.4
      },
      "position" : {
        "x" : -6216.07470703125,
        "y" : 2730.10888671875
      },
      "selected" : false
    }, {
      "data" : {
        "id" : "1264",
        "symbol" : "GDU6",
        "Shoot_Descriptions" : "",
        "PLAMOM_Match_Shoot" : "",
        "Causal_TF" : "GDU6",
        "Root_Descriptions" : "AtGDU6, GDU6, glutamine dumper 6",
        "Eccentricity" : 1,
        "Outdegree" : 1,
        "Target_PLAMOM" : "",
        "PLAMOM_Match_Root" : "NO",
        "shared_name" : "AT3G30725",
        "Indegree" : 0,
        "Target_tRNA" : "",
        "t_RNA_motif_shoot" : "",
        "SUID" : 1264,
        "SelfLoops" : 0,
        "Target_TF" : "",
        "MolecularAttributes" : "protein_coding",
        "AverageShortestPathLength" : 1.0,
        "selected" : false,
        "Name2" : "GDU6",
        "ClosenessCentrality" : 1.0,
        "Casual_Name" : "\"AtGDU6",
        "PartnerOfMultiEdgedNodePairs" : 0,
        "ClusteringCoefficient" : 0.0,
        "Target_Name" : "",
        "Stress" : 0,
        "BetweennessCentrality" : 0.0,
        "t_RNA_motif_Root" : "NO",
        "EdgeCount" : 1,
        "Causal_PLAMOM" : "no",
        "name" : "AT3G30725",
        "Causal_tRNA" : "no",
        "IsSingleNode" : false,
        "briefDescription" : "glutamine dumper 6",
        "NeighborhoodConnectivity" : 10.0
      },
      "position" : {
        "x" : -6143.744140625,
        "y" : -616.0703125
      },
      "selected" : false
    }, {
      "data" : {
        "id" : "1263",
        "symbol" : "AT3G55690",
        "Shoot_Descriptions" : "",
        "PLAMOM_Match_Shoot" : "",
        "Causal_TF" : "no",
        "Root_Descriptions" : "unknown protein; BEST Arabidopsis thaliana protein match is: unknown protein (TAIR:AT2G39870.1); Has 76 Blast hits to 69 proteins in 15 species: Archae - 0; Bacteria - 0; Metazoa - 4; Fungi - 3; Plants - 69; Viruses - 0; Other Eukaryotes - 0 (source: NCBI BLink).",
        "Eccentricity" : 12,
        "Outdegree" : 2,
        "Target_PLAMOM" : "",
        "PLAMOM_Match_Root" : "NO",
        "shared_name" : "AT3G55690",
        "Indegree" : 0,
        "Target_tRNA" : "",
        "t_RNA_motif_shoot" : "",
        "SUID" : 1263,
        "SelfLoops" : 0,
        "Target_TF" : "",
        "MolecularAttributes" : "protein_coding",
        "AverageShortestPathLength" : 7.05010438,
        "selected" : false,
        "Name2" : "AT3G55690",
        "ClosenessCentrality" : 0.14184187,
        "Casual_Name" : "unknown protein; BEST Arabidopsis thaliana protein match is: unknown protein (TAIR:AT2G39870.1); Has 76 Blast hits to 69 proteins in 15 species: Archae - 0; Bacteria - 0; Metazoa - 4; Fungi - 3; Plants - 69; Viruses - 0; Other Eukaryotes - 0 (source: NCBI BLink).",
        "PartnerOfMultiEdgedNodePairs" : 0,
        "ClusteringCoefficient" : 0.0,
        "Target_Name" : "",
        "Stress" : 0,
        "BetweennessCentrality" : 0.0,
        "t_RNA_motif_Root" : "NO",
        "EdgeCount" : 2,
        "Causal_PLAMOM" : "no",
        "name" : "AT3G55690",
        "Causal_tRNA" : "no",
        "IsSingleNode" : false,
        "briefDescription" : "hypothetical protein",
        "NeighborhoodConnectivity" : 7.0
      },
      "position" : {
        "x" : -6354.7109375,
        "y" : -220.30946350097656
      },
      "selected" : false
    }, {
      "data" : {
        "id" : "1262",
        "symbol" : "AT5G60520",
        "Shoot_Descriptions" : "",
        "PLAMOM_Match_Shoot" : "",
        "Causal_TF" : "no",
        "Root_Descriptions" : "Late embryogenesis abundant (LEA) protein-related",
        "Eccentricity" : 14,
        "Outdegree" : 6,
        "Target_PLAMOM" : "",
        "PLAMOM_Match_Root" : "NO",
        "shared_name" : "AT5G60520",
        "Indegree" : 0,
        "Target_tRNA" : "",
        "t_RNA_motif_shoot" : "",
        "SUID" : 1262,
        "SelfLoops" : 0,
        "Target_TF" : "",
        "MolecularAttributes" : "protein_coding",
        "AverageShortestPathLength" : 9.84854772,
        "selected" : false,
        "Name2" : "AT5G60520",
        "ClosenessCentrality" : 0.10153781,
        "Casual_Name" : "Late embryogenesis abundant (LEA) protein-related",
        "PartnerOfMultiEdgedNodePairs" : 0,
        "ClusteringCoefficient" : 0.0,
        "Target_Name" : "",
        "Stress" : 0,
        "BetweennessCentrality" : 0.0,
        "t_RNA_motif_Root" : "NO",
        "EdgeCount" : 6,
        "Causal_PLAMOM" : "no",
        "name" : "AT5G60520",
        "Causal_tRNA" : "no",
        "IsSingleNode" : false,
        "briefDescription" : "Late embryogenesis abundant (LEA) protein-like protein",
        "NeighborhoodConnectivity" : 9.0
      },
      "position" : {
        "x" : -5788.44140625,
        "y" : -391.71710205078125
      },
      "selected" : false
    }, {
      "data" : {
        "id" : "1261",
        "symbol" : "TPS8",
        "Shoot_Descriptions" : "",
        "PLAMOM_Match_Shoot" : "",
        "Causal_TF" : "ATTPSC",
        "Root_Descriptions" : "ATTPS8, ATTPSC, TPS8, trehalose-6-phosphatase synthase S8",
        "Eccentricity" : 12,
        "Outdegree" : 4,
        "Target_PLAMOM" : "",
        "PLAMOM_Match_Root" : "YES",
        "shared_name" : "AT1G70290",
        "Indegree" : 0,
        "Target_tRNA" : "",
        "t_RNA_motif_shoot" : "",
        "SUID" : 1261,
        "SelfLoops" : 0,
        "Target_TF" : "",
        "MolecularAttributes" : "protein_coding",
        "AverageShortestPathLength" : 6.89375,
        "selected" : false,
        "Name2" : "TPS8",
        "ClosenessCentrality" : 0.14505893,
        "Casual_Name" : "\"ATTPS8",
        "PartnerOfMultiEdgedNodePairs" : 0,
        "ClusteringCoefficient" : 0.0,
        "Target_Name" : "",
        "Stress" : 0,
        "BetweennessCentrality" : 0.0,
        "t_RNA_motif_Root" : "YES",
        "EdgeCount" : 4,
        "Causal_PLAMOM" : "AT1G70290",
        "name" : "AT1G70290",
        "Causal_tRNA" : "AT1G70290",
        "IsSingleNode" : false,
        "briefDescription" : "trehalose-6-phosphatase synthase S8",
        "NeighborhoodConnectivity" : 7.75
      },
      "position" : {
        "x" : -6000.1015625,
        "y" : -177.875732421875
      },
      "selected" : false
    }, {
      "data" : {
        "id" : "1260",
        "symbol" : "AT3G46880",
        "Shoot_Descriptions" : "",
        "PLAMOM_Match_Shoot" : "",
        "Causal_TF" : "no",
        "Root_Descriptions" : "unknown protein; FUNCTIONS IN: molecular_function unknown; INVOLVED IN: biological_process unknown; LOCATED IN: chloroplast; EXPRESSED IN: 14 plant structures; EXPRESSED DURING: 9 growth stages; BEST Arabidopsis thaliana protein match is: unknown protein (TAIR:AT5G59080.1); Has 51 Blast hits to 51 proteins in 8 species: Archae - 0; Bacteria - 0; Metazoa - 0; Fungi - 0; Plants - 51; Viruses - 0; Other Eukaryotes - 0 (source: NCBI BLink).",
        "Eccentricity" : 10,
        "Outdegree" : 4,
        "Target_PLAMOM" : "AT3G46880",
        "PLAMOM_Match_Root" : "YES",
        "shared_name" : "AT3G46880",
        "Indegree" : 1,
        "Target_tRNA" : "no",
        "t_RNA_motif_shoot" : "",
        "SUID" : 1260,
        "SelfLoops" : 0,
        "Target_TF" : "no",
        "MolecularAttributes" : "protein_coding",
        "AverageShortestPathLength" : 5.62577963,
        "selected" : false,
        "Name2" : "AT3G46880",
        "ClosenessCentrality" : 0.17775314,
        "Casual_Name" : "unknown protein; FUNCTIONS IN: molecular_function unknown; INVOLVED IN: biological_process unknown; LOCATED IN: chloroplast; EXPRESSED IN: 14 plant structures; EXPRESSED DURING: 9 growth stages; BEST Arabidopsis thaliana protein match is: unknown protein (TAIR:AT5G59080.1); Has 51 Blast hits to 51 proteins in 8 species: Archae - 0; Bacteria - 0; Metazoa - 0; Fungi - 0; Plants - 51; Viruses - 0; Other Eukaryotes - 0 (source: NCBI BLink).",
        "PartnerOfMultiEdgedNodePairs" : 0,
        "ClusteringCoefficient" : 0.0,
        "Target_Name" : "unknown protein; FUNCTIONS IN: molecular_function unknown; INVOLVED IN: biological_process unknown; LOCATED IN: chloroplast; EXPRESSED IN: 14 plant structures; EXPRESSED DURING: 9 growth stages; BEST Arabidopsis thaliana protein match is: unknown protein (TAIR:AT5G59080.1); Has 51 Blast hits to 51 proteins in 8 species: Archae - 0; Bacteria - 0; Metazoa - 0; Fungi - 0; Plants - 51; Viruses - 0; Other Eukaryotes - 0 (source: NCBI BLink).",
        "Stress" : 1155,
        "BetweennessCentrality" : 1.321E-4,
        "t_RNA_motif_Root" : "NO",
        "EdgeCount" : 5,
        "Causal_PLAMOM" : "AT3G46880",
        "name" : "AT3G46880",
        "Causal_tRNA" : "no",
        "IsSingleNode" : false,
        "briefDescription" : "hypothetical protein",
        "NeighborhoodConnectivity" : 9.2
      },
      "position" : {
        "x" : -6088.63232421875,
        "y" : 3716.388427734375
      },
      "selected" : false
    }, {
      "data" : {
        "id" : "1259",
        "symbol" : "RPL27AB",
        "Shoot_Descriptions" : "RPL27A, RPL27AB, Ribosomal protein L18e/L15 superfamily protein",
        "PLAMOM_Match_Shoot" : "YES",
        "Causal_TF" : "",
        "Root_Descriptions" : "",
        "Eccentricity" : 0,
        "Outdegree" : 0,
        "Target_PLAMOM" : "cytokinin oxidase 4\"",
        "PLAMOM_Match_Root" : "",
        "shared_name" : "AT1G23290",
        "Indegree" : 2,
        "Target_tRNA" : "no",
        "t_RNA_motif_shoot" : "NO",
        "SUID" : 1259,
        "SelfLoops" : 0,
        "Target_TF" : "no",
        "MolecularAttributes" : "protein_coding",
        "AverageShortestPathLength" : 0.0,
        "selected" : false,
        "Name2" : "RPL27AB",
        "ClosenessCentrality" : 0.0,
        "Casual_Name" : "",
        "PartnerOfMultiEdgedNodePairs" : 0,
        "ClusteringCoefficient" : 0.0,
        "Target_Name" : "AT1G23290",
        "Stress" : 0,
        "BetweennessCentrality" : 0.0,
        "t_RNA_motif_Root" : "",
        "EdgeCount" : 2,
        "Causal_PLAMOM" : "",
        "name" : "AT1G23290",
        "Causal_tRNA" : "",
        "IsSingleNode" : false,
        "briefDescription" : "Ribosomal protein L18e/L15 superfamily protein",
        "NeighborhoodConnectivity" : 6.0
      },
      "position" : {
        "x" : 4491.868114876921,
        "y" : 928.78662109375
      },
      "selected" : false
    }, {
      "data" : {
        "id" : "1258",
        "symbol" : "AT5G08350",
        "Shoot_Descriptions" : "",
        "PLAMOM_Match_Shoot" : "",
        "Causal_TF" : "no",
        "Root_Descriptions" : "GRAM domain-containing protein / ABA-responsive protein-related",
        "Eccentricity" : 12,
        "Outdegree" : 3,
        "Target_PLAMOM" : "no",
        "PLAMOM_Match_Root" : "NO",
        "shared_name" : "AT5G08350",
        "Indegree" : 7,
        "Target_tRNA" : "no",
        "t_RNA_motif_shoot" : "",
        "SUID" : 1258,
        "SelfLoops" : 0,
        "Target_TF" : "no",
        "MolecularAttributes" : "protein_coding",
        "AverageShortestPathLength" : 6.9916318,
        "selected" : false,
        "Name2" : "AT5G08350",
        "ClosenessCentrality" : 0.14302813,
        "Casual_Name" : "GRAM domain-containing protein / ABA-responsive protein-related",
        "PartnerOfMultiEdgedNodePairs" : 0,
        "ClusteringCoefficient" : 0.0,
        "Target_Name" : "GRAM domain-containing protein / ABA-responsive protein-related",
        "Stress" : 4629,
        "BetweennessCentrality" : 0.00133817,
        "t_RNA_motif_Root" : "NO",
        "EdgeCount" : 10,
        "Causal_PLAMOM" : "no",
        "name" : "AT5G08350",
        "Causal_tRNA" : "no",
        "IsSingleNode" : false,
        "briefDescription" : "GRAM domain-containing protein / ABA-responsive protein-like protein",
        "NeighborhoodConnectivity" : 7.6
      },
      "position" : {
        "x" : -6328.06494140625,
        "y" : 3285.77001953125
      },
      "selected" : false
    }, {
      "data" : {
        "id" : "1257",
        "symbol" : "RSZ21",
        "Shoot_Descriptions" : "RSZP21, SRZ-21, SRZ21, RS-containing zinc finger protein 21",
        "PLAMOM_Match_Shoot" : "NO",
        "Causal_TF" : "",
        "Root_Descriptions" : "",
        "Eccentricity" : 0,
        "Outdegree" : 0,
        "Target_PLAMOM" : "no",
        "PLAMOM_Match_Root" : "",
        "shared_name" : "AT1G23860",
        "Indegree" : 1,
        "Target_tRNA" : "no",
        "t_RNA_motif_shoot" : "NO",
        "SUID" : 1257,
        "SelfLoops" : 0,
        "Target_TF" : "SRZ-21",
        "MolecularAttributes" : "protein_coding",
        "AverageShortestPathLength" : 0.0,
        "selected" : false,
        "Name2" : "RSZ21",
        "ClosenessCentrality" : 0.0,
        "Casual_Name" : "",
        "PartnerOfMultiEdgedNodePairs" : 0,
        "ClusteringCoefficient" : 0.0,
        "Target_Name" : "\"RSZP21",
        "Stress" : 0,
        "BetweennessCentrality" : 0.0,
        "t_RNA_motif_Root" : "",
        "EdgeCount" : 1,
        "Causal_PLAMOM" : "",
        "name" : "AT1G23860",
        "Causal_tRNA" : "",
        "IsSingleNode" : false,
        "briefDescription" : "RS-containing zinc finger protein 21",
        "NeighborhoodConnectivity" : 10.0
      },
      "position" : {
        "x" : 4416.568369802832,
        "y" : 207.65994262695312
      },
      "selected" : false
    }, {
      "data" : {
        "id" : "1256",
        "symbol" : "SULTR3;5",
        "Shoot_Descriptions" : "",
        "PLAMOM_Match_Shoot" : "",
        "Causal_TF" : "sulfate transporter 3;5\"",
        "Root_Descriptions" : "SULTR3;5, sulfate transporter 3;5",
        "Eccentricity" : 3,
        "Outdegree" : 3,
        "Target_PLAMOM" : "no",
        "PLAMOM_Match_Root" : "NO",
        "shared_name" : "AT5G19600",
        "Indegree" : 2,
        "Target_tRNA" : "no",
        "t_RNA_motif_shoot" : "",
        "SUID" : 1256,
        "SelfLoops" : 0,
        "Target_TF" : "\"SULTR3;5",
        "MolecularAttributes" : "protein_coding",
        "AverageShortestPathLength" : 1.66666667,
        "selected" : false,
        "Name2" : "SULTR3;5",
        "ClosenessCentrality" : 0.6,
        "Casual_Name" : "\"SULTR3;5",
        "PartnerOfMultiEdgedNodePairs" : 0,
        "ClusteringCoefficient" : 0.0,
        "Target_Name" : "no",
        "Stress" : 12,
        "BetweennessCentrality" : 7.16E-6,
        "t_RNA_motif_Root" : "NO",
        "EdgeCount" : 5,
        "Causal_PLAMOM" : "no",
        "name" : "AT5G19600",
        "Causal_tRNA" : "no",
        "IsSingleNode" : false,
        "briefDescription" : "sulfate transporter 3;5",
        "NeighborhoodConnectivity" : 1.8
      },
      "position" : {
        "x" : -7521.0439453125,
        "y" : 3564.28857421875
      },
      "selected" : false
    }, {
      "data" : {
        "id" : "1255",
        "symbol" : "AT1G24270",
        "Shoot_Descriptions" : "unknown protein; BEST Arabidopsis thaliana protein match is: unknown protein (TAIR:AT5G13090.1); Has 84 Blast hits to 83 proteins in 13 species: Archae - 0; Bacteria - 0; Metazoa - 0; Fungi - 0; Plants - 84; Viruses - 0; Other Eukaryotes - 0 (source: NCBI BLink).",
        "PLAMOM_Match_Shoot" : "NO",
        "Causal_TF" : "",
        "Root_Descriptions" : "",
        "Eccentricity" : 0,
        "Outdegree" : 0,
        "Target_PLAMOM" : "no",
        "PLAMOM_Match_Root" : "",
        "shared_name" : "AT1G24270",
        "Indegree" : 2,
        "Target_tRNA" : "no",
        "t_RNA_motif_shoot" : "NO",
        "SUID" : 1255,
        "SelfLoops" : 0,
        "Target_TF" : "unknown protein; BEST Arabidopsis thaliana protein match is: unknown protein (TAIR:AT5G13090.1); Has 84 Blast hits to 83 proteins in 13 species: Archae - 0; Bacteria - 0; Metazoa - 0; Fungi - 0; Plants - 84; Viruses - 0; Other Eukaryotes - 0 (source: NCBI BLink).",
        "MolecularAttributes" : "protein_coding",
        "AverageShortestPathLength" : 0.0,
        "selected" : false,
        "Name2" : "AT1G24270",
        "ClosenessCentrality" : 0.0,
        "Casual_Name" : "",
        "PartnerOfMultiEdgedNodePairs" : 0,
        "ClusteringCoefficient" : 0.0,
        "Target_Name" : "no",
        "Stress" : 0,
        "BetweennessCentrality" : 0.0,
        "t_RNA_motif_Root" : "",
        "EdgeCount" : 2,
        "Causal_PLAMOM" : "",
        "name" : "AT1G24270",
        "Causal_tRNA" : "",
        "IsSingleNode" : false,
        "briefDescription" : "hypothetical protein",
        "NeighborhoodConnectivity" : 16.0
      },
      "position" : {
        "x" : 3247.712841439421,
        "y" : 383.9328308105469
      },
      "selected" : false
    }, {
      "data" : {
        "id" : "1254",
        "symbol" : "AT1G24440",
        "Shoot_Descriptions" : "RING/U-box superfamily protein",
        "PLAMOM_Match_Shoot" : "NO",
        "Causal_TF" : "no",
        "Root_Descriptions" : "",
        "Eccentricity" : 11,
        "Outdegree" : 5,
        "Target_PLAMOM" : "no",
        "PLAMOM_Match_Root" : "",
        "shared_name" : "AT1G24440",
        "Indegree" : 3,
        "Target_tRNA" : "no",
        "t_RNA_motif_shoot" : "NO",
        "SUID" : 1254,
        "SelfLoops" : 0,
        "Target_TF" : "no",
        "MolecularAttributes" : "protein_coding",
        "AverageShortestPathLength" : 6.01464435,
        "selected" : false,
        "Name2" : "AT1G24440",
        "ClosenessCentrality" : 0.16626087,
        "Casual_Name" : "RING/U-box superfamily protein",
        "PartnerOfMultiEdgedNodePairs" : 0,
        "ClusteringCoefficient" : 0.0,
        "Target_Name" : "RING/U-box superfamily protein",
        "Stress" : 5472,
        "BetweennessCentrality" : 0.00100463,
        "t_RNA_motif_Root" : "",
        "EdgeCount" : 8,
        "Causal_PLAMOM" : "no",
        "name" : "AT1G24440",
        "Causal_tRNA" : "no",
        "IsSingleNode" : false,
        "briefDescription" : "RING/U-box superfamily protein",
        "NeighborhoodConnectivity" : 10.875
      },
      "position" : {
        "x" : 3258.664306640625,
        "y" : 2722.33349609375
      },
      "selected" : false
    }, {
      "data" : {
        "id" : "1253",
        "symbol" : "AT1G76700",
        "Shoot_Descriptions" : "",
        "PLAMOM_Match_Shoot" : "",
        "Causal_TF" : "no",
        "Root_Descriptions" : "DNAJ heat shock N-terminal domain-containing protein",
        "Eccentricity" : 9,
        "Outdegree" : 7,
        "Target_PLAMOM" : "DOF zinc finger protein 1\"",
        "PLAMOM_Match_Root" : "NO",
        "shared_name" : "AT1G76700",
        "Indegree" : 1,
        "Target_tRNA" : "AT1G51700",
        "t_RNA_motif_shoot" : "",
        "SUID" : 1253,
        "SelfLoops" : 0,
        "Target_TF" : "no",
        "MolecularAttributes" : "protein_coding",
        "AverageShortestPathLength" : 5.12734864,
        "selected" : false,
        "Name2" : "AT1G76700",
        "ClosenessCentrality" : 0.19503257,
        "Casual_Name" : "DNAJ heat shock N-terminal domain-containing protein",
        "PartnerOfMultiEdgedNodePairs" : 0,
        "ClusteringCoefficient" : 0.0,
        "Target_Name" : "no",
        "Stress" : 948,
        "BetweennessCentrality" : 2.1759E-4,
        "t_RNA_motif_Root" : "NO",
        "EdgeCount" : 8,
        "Causal_PLAMOM" : "no",
        "name" : "AT1G76700",
        "Causal_tRNA" : "no",
        "IsSingleNode" : false,
        "briefDescription" : "DNAJ heat shock N-terminal domain-containing protein",
        "NeighborhoodConnectivity" : 14.25
      },
      "position" : {
        "x" : -6659.2978515625,
        "y" : 3169.6357421875
      },
      "selected" : false
    }, {
      "data" : {
        "id" : "1252",
        "symbol" : "AT3G13560",
        "Shoot_Descriptions" : "O-Glycosyl hydrolases family 17 protein",
        "PLAMOM_Match_Shoot" : "NO",
        "Causal_TF" : "no",
        "Root_Descriptions" : "O-Glycosyl hydrolases family 17 protein",
        "Eccentricity" : 12,
        "Outdegree" : 1,
        "Target_PLAMOM" : "no",
        "PLAMOM_Match_Root" : "NO",
        "shared_name" : "AT3G13560",
        "Indegree" : 2,
        "Target_tRNA" : "no",
        "t_RNA_motif_shoot" : "NO",
        "SUID" : 1252,
        "SelfLoops" : 0,
        "Target_TF" : "no",
        "MolecularAttributes" : "protein_coding",
        "AverageShortestPathLength" : 7.00208768,
        "selected" : false,
        "Name2" : "AT3G13560",
        "ClosenessCentrality" : 0.14281455,
        "Casual_Name" : "O-Glycosyl hydrolases family 17 protein",
        "PartnerOfMultiEdgedNodePairs" : 0,
        "ClusteringCoefficient" : 0.0,
        "Target_Name" : "O-Glycosyl hydrolases family 17 protein",
        "Stress" : 195,
        "BetweennessCentrality" : 5.167E-5,
        "t_RNA_motif_Root" : "NO",
        "EdgeCount" : 3,
        "Causal_PLAMOM" : "no",
        "name" : "AT3G13560",
        "Causal_tRNA" : "no",
        "IsSingleNode" : false,
        "briefDescription" : "O-Glycosyl hydrolases family 17 protein",
        "NeighborhoodConnectivity" : 7.333333333333333
      },
      "position" : {
        "x" : -1543.720703125,
        "y" : 2808.7822161346676
      },
      "selected" : false
    }, {
      "data" : {
        "id" : "1251",
        "symbol" : "NHL3",
        "Shoot_Descriptions" : "",
        "PLAMOM_Match_Shoot" : "",
        "Causal_TF" : "NDR1/HIN1-like 3\"",
        "Root_Descriptions" : "NHL3, NDR1/HIN1-like 3",
        "Eccentricity" : 1,
        "Outdegree" : 1,
        "Target_PLAMOM" : "no",
        "PLAMOM_Match_Root" : "YES",
        "shared_name" : "AT5G06320",
        "Indegree" : 2,
        "Target_tRNA" : "AT5G06320",
        "t_RNA_motif_shoot" : "",
        "SUID" : 1251,
        "SelfLoops" : 0,
        "Target_TF" : "\"NHL3",
        "MolecularAttributes" : "protein_coding",
        "AverageShortestPathLength" : 1.0,
        "selected" : false,
        "Name2" : "NHL3",
        "ClosenessCentrality" : 1.0,
        "Casual_Name" : "\"NHL3",
        "PartnerOfMultiEdgedNodePairs" : 0,
        "ClusteringCoefficient" : 0.0,
        "Target_Name" : "no",
        "Stress" : 3,
        "BetweennessCentrality" : 1.49E-6,
        "t_RNA_motif_Root" : "NO",
        "EdgeCount" : 3,
        "Causal_PLAMOM" : "AT5G06320",
        "name" : "AT5G06320",
        "Causal_tRNA" : "no",
        "IsSingleNode" : false,
        "briefDescription" : "NDR1/HIN1-like 3",
        "NeighborhoodConnectivity" : 2.3333333333333335
      },
      "position" : {
        "x" : -6578.533203125,
        "y" : 3848.76611328125
      },
      "selected" : false
    }, {
      "data" : {
        "id" : "1250",
        "symbol" : "AT1G24530",
        "Shoot_Descriptions" : "Transducin/WD40 repeat-like superfamily protein",
        "PLAMOM_Match_Shoot" : "YES",
        "Causal_TF" : "",
        "Root_Descriptions" : "",
        "Eccentricity" : 0,
        "Outdegree" : 0,
        "Target_PLAMOM" : "L-fucokinase/GDP-L-fucose pyrophosphorylase\"",
        "PLAMOM_Match_Root" : "",
        "shared_name" : "AT1G24530",
        "Indegree" : 3,
        "Target_tRNA" : "no",
        "t_RNA_motif_shoot" : "NO",
        "SUID" : 1250,
        "SelfLoops" : 0,
        "Target_TF" : "no",
        "MolecularAttributes" : "protein_coding",
        "AverageShortestPathLength" : 0.0,
        "selected" : false,
        "Name2" : "AT1G24530",
        "ClosenessCentrality" : 0.0,
        "Casual_Name" : "",
        "PartnerOfMultiEdgedNodePairs" : 0,
        "ClusteringCoefficient" : 0.0,
        "Target_Name" : "AT1G24530",
        "Stress" : 0,
        "BetweennessCentrality" : 0.0,
        "t_RNA_motif_Root" : "",
        "EdgeCount" : 3,
        "Causal_PLAMOM" : "",
        "name" : "AT1G24530",
        "Causal_tRNA" : "",
        "IsSingleNode" : false,
        "briefDescription" : "Transducin/WD40 repeat-like superfamily protein",
        "NeighborhoodConnectivity" : 10.666666666666666
      },
      "position" : {
        "x" : 3817.322704720671,
        "y" : 583.71240234375
      },
      "selected" : false
    }, {
      "data" : {
        "id" : "1249",
        "symbol" : "AT5G42150",
        "Shoot_Descriptions" : "",
        "PLAMOM_Match_Shoot" : "",
        "Causal_TF" : "no",
        "Root_Descriptions" : "Glutathione S-transferase family protein",
        "Eccentricity" : 12,
        "Outdegree" : 4,
        "Target_PLAMOM" : "no",
        "PLAMOM_Match_Root" : "NO",
        "shared_name" : "AT5G42150",
        "Indegree" : 6,
        "Target_tRNA" : "no",
        "t_RNA_motif_shoot" : "",
        "SUID" : 1249,
        "SelfLoops" : 0,
        "Target_TF" : "Glutathione S-transferase family protein",
        "MolecularAttributes" : "protein_coding",
        "AverageShortestPathLength" : 6.73640167,
        "selected" : false,
        "Name2" : "AT5G42150",
        "ClosenessCentrality" : 0.1484472,
        "Casual_Name" : "Glutathione S-transferase family protein",
        "PartnerOfMultiEdgedNodePairs" : 0,
        "ClusteringCoefficient" : 0.0,
        "Target_Name" : "AT5G42150",
        "Stress" : 6658,
        "BetweennessCentrality" : 0.00151456,
        "t_RNA_motif_Root" : "YES",
        "EdgeCount" : 10,
        "Causal_PLAMOM" : "no",
        "name" : "AT5G42150",
        "Causal_tRNA" : "AT5G42150",
        "IsSingleNode" : false,
        "briefDescription" : "Glutathione S-transferase family protein",
        "NeighborhoodConnectivity" : 4.2
      },
      "position" : {
        "x" : -6971.501953125,
        "y" : 3138.7841796875
      },
      "selected" : false
    }, {
      "data" : {
        "id" : "1248",
        "symbol" : "AT1G21000",
        "Shoot_Descriptions" : "",
        "PLAMOM_Match_Shoot" : "",
        "Causal_TF" : "no",
        "Root_Descriptions" : "PLATZ transcription factor family protein",
        "Eccentricity" : 11,
        "Outdegree" : 7,
        "Target_PLAMOM" : "",
        "PLAMOM_Match_Root" : "YES",
        "shared_name" : "AT1G21000",
        "Indegree" : 0,
        "Target_tRNA" : "",
        "t_RNA_motif_shoot" : "",
        "SUID" : 1248,
        "SelfLoops" : 0,
        "Target_TF" : "",
        "MolecularAttributes" : "protein_coding",
        "AverageShortestPathLength" : 6.64315353,
        "selected" : false,
        "Name2" : "AT1G21000",
        "ClosenessCentrality" : 0.15053092,
        "Casual_Name" : "PLATZ transcription factor family protein",
        "PartnerOfMultiEdgedNodePairs" : 0,
        "ClusteringCoefficient" : 0.0,
        "Target_Name" : "",
        "Stress" : 0,
        "BetweennessCentrality" : 0.0,
        "t_RNA_motif_Root" : "NO",
        "EdgeCount" : 7,
        "Causal_PLAMOM" : "AT1G21000",
        "name" : "AT1G21000",
        "Causal_tRNA" : "no",
        "IsSingleNode" : false,
        "briefDescription" : "PLATZ transcription factor family protein",
        "NeighborhoodConnectivity" : 12.428571428571429
      },
      "position" : {
        "x" : -5569.06005859375,
        "y" : -149.5401153564453
      },
      "selected" : false
    }, {
      "data" : {
        "id" : "1247",
        "symbol" : "AT1G26300",
        "Shoot_Descriptions" : "BSD domain-containing protein",
        "PLAMOM_Match_Shoot" : "NO",
        "Causal_TF" : "no",
        "Root_Descriptions" : "",
        "Eccentricity" : 2,
        "Outdegree" : 1,
        "Target_PLAMOM" : "no",
        "PLAMOM_Match_Root" : "",
        "shared_name" : "AT1G26300",
        "Indegree" : 5,
        "Target_tRNA" : "AT1G26300",
        "t_RNA_motif_shoot" : "YES",
        "SUID" : 1247,
        "SelfLoops" : 0,
        "Target_TF" : "no",
        "MolecularAttributes" : "protein_coding",
        "AverageShortestPathLength" : 1.5,
        "selected" : false,
        "Name2" : "AT1G26300",
        "ClosenessCentrality" : 0.66666667,
        "Casual_Name" : "BSD domain-containing protein",
        "PartnerOfMultiEdgedNodePairs" : 0,
        "ClusteringCoefficient" : 0.0,
        "Target_Name" : "BSD domain-containing protein",
        "Stress" : 506,
        "BetweennessCentrality" : 1.1536E-4,
        "t_RNA_motif_Root" : "",
        "EdgeCount" : 6,
        "Causal_PLAMOM" : "no",
        "name" : "AT1G26300",
        "Causal_tRNA" : "AT1G26300",
        "IsSingleNode" : false,
        "briefDescription" : "BSD domain-containing protein",
        "NeighborhoodConnectivity" : 9.833333333333334
      },
      "position" : {
        "x" : 4012.874950814421,
        "y" : 3013.682373046875
      },
      "selected" : false
    }, {
      "data" : {
        "id" : "1246",
        "symbol" : "SKOR",
        "Shoot_Descriptions" : "",
        "PLAMOM_Match_Shoot" : "",
        "Causal_TF" : "STELAR K+ outward rectifier\"",
        "Root_Descriptions" : "SKOR, STELAR K+ outward rectifier",
        "Eccentricity" : 3,
        "Outdegree" : 1,
        "Target_PLAMOM" : "",
        "PLAMOM_Match_Root" : "NO",
        "shared_name" : "AT3G02850",
        "Indegree" : 0,
        "Target_tRNA" : "",
        "t_RNA_motif_shoot" : "",
        "SUID" : 1246,
        "SelfLoops" : 0,
        "Target_TF" : "",
        "MolecularAttributes" : "protein_coding",
        "AverageShortestPathLength" : 2.0,
        "selected" : false,
        "Name2" : "SKOR",
        "ClosenessCentrality" : 0.5,
        "Casual_Name" : "\"SKOR",
        "PartnerOfMultiEdgedNodePairs" : 0,
        "ClusteringCoefficient" : 0.0,
        "Target_Name" : "",
        "Stress" : 0,
        "BetweennessCentrality" : 0.0,
        "t_RNA_motif_Root" : "NO",
        "EdgeCount" : 1,
        "Causal_PLAMOM" : "no",
        "name" : "AT3G02850",
        "Causal_tRNA" : "no",
        "IsSingleNode" : false,
        "briefDescription" : "STELAR K  outward rectifier",
        "NeighborhoodConnectivity" : 6.0
      },
      "position" : {
        "x" : -5312.80078125,
        "y" : -178.6326904296875
      },
      "selected" : false
    }, {
      "data" : {
        "id" : "1245",
        "symbol" : "AT3G47980",
        "degree_layout" : 1,
        "Shoot_Descriptions" : "",
        "PLAMOM_Match_Shoot" : "",
        "Causal_TF" : "no",
        "Root_Descriptions" : "Integral membrane HPP family protein",
        "Eccentricity" : 14,
        "Outdegree" : 3,
        "Target_PLAMOM" : "AT5G25160",
        "PLAMOM_Match_Root" : "NO",
        "shared_name" : "AT3G47980",
        "Indegree" : 8,
        "Target_tRNA" : "no",
        "t_RNA_motif_shoot" : "",
        "SUID" : 1245,
        "SelfLoops" : 0,
        "Target_TF" : "Integral membrane HPP family protein",
        "MolecularAttributes" : "protein_coding",
        "AverageShortestPathLength" : 9.38912134,
        "selected" : false,
        "Name2" : "AT3G47980",
        "ClosenessCentrality" : 0.10650624,
        "Casual_Name" : "Integral membrane HPP family protein",
        "PartnerOfMultiEdgedNodePairs" : 0,
        "ClusteringCoefficient" : 0.0,
        "Target_Name" : "no",
        "Stress" : 9062,
        "BetweennessCentrality" : 0.00261228,
        "t_RNA_motif_Root" : "NO",
        "EdgeCount" : 11,
        "Causal_PLAMOM" : "no",
        "name" : "AT3G47980",
        "Causal_tRNA" : "no",
        "IsSingleNode" : false,
        "briefDescription" : "Integral membrane HPP family protein",
        "NeighborhoodConnectivity" : 7.363636363636363
      },
      "position" : {
        "x" : -6286.841796875,
        "y" : 3178.18896484375
      },
      "selected" : false
    }, {
      "data" : {
        "id" : "1244",
        "symbol" : "AT1G56230",
        "degree_layout" : 11,
        "Shoot_Descriptions" : "",
        "PLAMOM_Match_Shoot" : "",
        "Causal_TF" : "no",
        "Root_Descriptions" : "Protein of unknown function (DUF1399)",
        "Eccentricity" : 8,
        "Outdegree" : 14,
        "Target_PLAMOM" : "no",
        "PLAMOM_Match_Root" : "NO",
        "shared_name" : "AT1G56230",
        "Indegree" : 3,
        "Target_tRNA" : "no",
        "t_RNA_motif_shoot" : "",
        "SUID" : 1244,
        "SelfLoops" : 0,
        "Target_TF" : "no",
        "MolecularAttributes" : "protein_coding",
        "AverageShortestPathLength" : 4.11087866,
        "selected" : false,
        "Name2" : "AT1G56230",
        "ClosenessCentrality" : 0.243257,
        "Casual_Name" : "Protein of unknown function (DUF1399)",
        "PartnerOfMultiEdgedNodePairs" : 0,
        "ClusteringCoefficient" : 0.0,
        "Target_Name" : "Protein of unknown function (DUF1399)",
        "Stress" : 24303,
        "BetweennessCentrality" : 0.00462494,
        "t_RNA_motif_Root" : "NO",
        "EdgeCount" : 17,
        "Causal_PLAMOM" : "no",
        "name" : "AT1G56230",
        "Causal_tRNA" : "no",
        "IsSingleNode" : false,
        "briefDescription" : "enolase (DUF1399)",
        "NeighborhoodConnectivity" : 12.470588235294118
      },
      "position" : {
        "x" : -6618.5029296875,
        "y" : 2952.635009765625
      },
      "selected" : false
    }, {
      "data" : {
        "id" : "1243",
        "symbol" : "UPS5",
        "degree_layout" : 7,
        "Shoot_Descriptions" : "ATUPS5, UPS5, UPS5, ureide permease 5",
        "PLAMOM_Match_Shoot" : "NO",
        "Causal_TF" : "UPS5",
        "Root_Descriptions" : "",
        "Eccentricity" : 9,
        "Outdegree" : 6,
        "Target_PLAMOM" : "no",
        "PLAMOM_Match_Root" : "",
        "shared_name" : "AT1G26440",
        "Indegree" : 1,
        "Target_tRNA" : "no",
        "t_RNA_motif_shoot" : "NO",
        "SUID" : 1243,
        "SelfLoops" : 0,
        "Target_TF" : "UPS5",
        "MolecularAttributes" : "protein_coding",
        "AverageShortestPathLength" : 4.29707113,
        "selected" : false,
        "Name2" : "UPS5",
        "ClosenessCentrality" : 0.23271665,
        "Casual_Name" : "\"ATUPS5",
        "PartnerOfMultiEdgedNodePairs" : 0,
        "ClusteringCoefficient" : 0.11904762,
        "Target_Name" : "\"ATUPS5",
        "Stress" : 3709,
        "BetweennessCentrality" : 9.1476E-4,
        "t_RNA_motif_Root" : "",
        "EdgeCount" : 7,
        "Causal_PLAMOM" : "no",
        "name" : "AT1G26440",
        "Causal_tRNA" : "no",
        "IsSingleNode" : false,
        "briefDescription" : "ureide permease 5",
        "NeighborhoodConnectivity" : 29.428571428571427
      },
      "position" : {
        "x" : 3499.810009408171,
        "y" : 2835.330322265625
      },
      "selected" : false
    }, {
      "data" : {
        "id" : "1242",
        "symbol" : "AT1G47395",
        "Shoot_Descriptions" : "",
        "PLAMOM_Match_Shoot" : "",
        "Causal_TF" : "no",
        "Root_Descriptions" : "unknown protein; FUNCTIONS IN: molecular_function unknown; INVOLVED IN: biological_process unknown; LOCATED IN: endomembrane system; BEST Arabidopsis thaliana protein match is: unknown protein (TAIR:AT1G47400.1); Has 9 Blast hits to 9 proteins in 2 species: Archae - 0; Bacteria - 0; Metazoa - 0; Fungi - 0; Plants - 9; Viruses - 0; Other Eukaryotes - 0 (source: NCBI BLink).",
        "Eccentricity" : 13,
        "Outdegree" : 4,
        "Target_PLAMOM" : "no",
        "PLAMOM_Match_Root" : "NO",
        "shared_name" : "AT1G47395",
        "Indegree" : 2,
        "Target_tRNA" : "no",
        "t_RNA_motif_shoot" : "",
        "SUID" : 1242,
        "SelfLoops" : 0,
        "Target_TF" : "no",
        "MolecularAttributes" : "protein_coding",
        "AverageShortestPathLength" : 7.7219917,
        "selected" : false,
        "Name2" : "AT1G47395",
        "ClosenessCentrality" : 0.12950027,
        "Casual_Name" : "unknown protein; FUNCTIONS IN: molecular_function unknown; INVOLVED IN: biological_process unknown; LOCATED IN: endomembrane system; BEST Arabidopsis thaliana protein match is: unknown protein (TAIR:AT1G47400.1); Has 9 Blast hits to 9 proteins in 2 species: Archae - 0; Bacteria - 0; Metazoa - 0; Fungi - 0; Plants - 9; Viruses - 0; Other Eukaryotes - 0 (source: NCBI BLink).",
        "PartnerOfMultiEdgedNodePairs" : 0,
        "ClusteringCoefficient" : 0.0,
        "Target_Name" : "unknown protein; FUNCTIONS IN: molecular_function unknown; INVOLVED IN: biological_process unknown; LOCATED IN: endomembrane system; BEST Arabidopsis thaliana protein match is: unknown protein (TAIR:AT1G47400.1); Has 9 Blast hits to 9 proteins in 2 species: Archae - 0; Bacteria - 0; Metazoa - 0; Fungi - 0; Plants - 9; Viruses - 0; Other Eukaryotes - 0 (source: NCBI BLink).",
        "Stress" : 1091,
        "BetweennessCentrality" : 3.1155E-4,
        "t_RNA_motif_Root" : "NO",
        "EdgeCount" : 6,
        "Causal_PLAMOM" : "no",
        "name" : "AT1G47395",
        "Causal_tRNA" : "no",
        "IsSingleNode" : false,
        "briefDescription" : "hypothetical protein",
        "NeighborhoodConnectivity" : 5.833333333333333
      },
      "position" : {
        "x" : -6765.248046875,
        "y" : 3767.348876953125
      },
      "selected" : false
    }, {
      "data" : {
        "id" : "1241",
        "symbol" : "AT1G27400",
        "Shoot_Descriptions" : "Ribosomal protein L22p/L17e family protein",
        "PLAMOM_Match_Shoot" : "YES",
        "Causal_TF" : "no",
        "Root_Descriptions" : "",
        "Eccentricity" : 12,
        "Outdegree" : 1,
        "Target_PLAMOM" : "cytokinin oxidase 4\"",
        "PLAMOM_Match_Root" : "",
        "shared_name" : "AT1G27400",
        "Indegree" : 2,
        "Target_tRNA" : "no",
        "t_RNA_motif_shoot" : "NO",
        "SUID" : 1241,
        "SelfLoops" : 0,
        "Target_TF" : "no",
        "MolecularAttributes" : "protein_coding",
        "AverageShortestPathLength" : 7.08977035,
        "selected" : false,
        "Name2" : "AT1G27400",
        "ClosenessCentrality" : 0.14104829,
        "Casual_Name" : "Ribosomal protein L22p/L17e family protein",
        "PartnerOfMultiEdgedNodePairs" : 0,
        "ClusteringCoefficient" : 0.0,
        "Target_Name" : "AT1G27400",
        "Stress" : 1586,
        "BetweennessCentrality" : 4.5177E-4,
        "t_RNA_motif_Root" : "",
        "EdgeCount" : 3,
        "Causal_PLAMOM" : "AT1G27400",
        "name" : "AT1G27400",
        "Causal_tRNA" : "no",
        "IsSingleNode" : false,
        "briefDescription" : "Ribosomal protein L22p/L17e family protein",
        "NeighborhoodConnectivity" : 15.0
      },
      "position" : {
        "x" : 3591.589306283171,
        "y" : 3757.120361328125
      },
      "selected" : false
    }, {
      "data" : {
        "id" : "1240",
        "symbol" : "AT3G60750",
        "Shoot_Descriptions" : "",
        "PLAMOM_Match_Shoot" : "",
        "Causal_TF" : "no",
        "Root_Descriptions" : "Transketolase",
        "Eccentricity" : 14,
        "Outdegree" : 5,
        "Target_PLAMOM" : "AT3G60750",
        "PLAMOM_Match_Root" : "YES",
        "shared_name" : "AT3G60750",
        "Indegree" : 1,
        "Target_tRNA" : "no",
        "t_RNA_motif_shoot" : "",
        "SUID" : 1240,
        "SelfLoops" : 0,
        "Target_TF" : "no",
        "MolecularAttributes" : "METABOLIC",
        "AverageShortestPathLength" : 8.82987552,
        "selected" : false,
        "Name2" : "AT3G60750",
        "ClosenessCentrality" : 0.11325188,
        "Casual_Name" : "Transketolase",
        "PartnerOfMultiEdgedNodePairs" : 0,
        "ClusteringCoefficient" : 0.0,
        "Target_Name" : "Transketolase",
        "Stress" : 1236,
        "BetweennessCentrality" : 2.8764E-4,
        "t_RNA_motif_Root" : "NO",
        "EdgeCount" : 6,
        "Causal_PLAMOM" : "AT3G60750",
        "name" : "AT3G60750",
        "Causal_tRNA" : "no",
        "IsSingleNode" : false,
        "briefDescription" : "Transketolase",
        "NeighborhoodConnectivity" : 1.8333333333333333
      },
      "position" : {
        "x" : -7362.458984375,
        "y" : 3717.571533203125
      },
      "selected" : false
    }, {
      "data" : {
        "id" : "1239",
        "symbol" : "ATU2AF35A",
        "Shoot_Descriptions" : "ATU2AF35A, U2 snRNP auxiliary factor small subunit, putative",
        "PLAMOM_Match_Shoot" : "NO",
        "Causal_TF" : "",
        "Root_Descriptions" : "",
        "Eccentricity" : 0,
        "Outdegree" : 0,
        "Target_PLAMOM" : "no",
        "PLAMOM_Match_Root" : "",
        "shared_name" : "AT1G27650",
        "Indegree" : 1,
        "Target_tRNA" : "no",
        "t_RNA_motif_shoot" : "NO",
        "SUID" : 1239,
        "SelfLoops" : 0,
        "Target_TF" : "U2 snRNP auxiliary factor small subunit",
        "MolecularAttributes" : "protein_coding",
        "AverageShortestPathLength" : 0.0,
        "selected" : false,
        "Name2" : "ATU2AF35A",
        "ClosenessCentrality" : 0.0,
        "Casual_Name" : "",
        "PartnerOfMultiEdgedNodePairs" : 0,
        "ClusteringCoefficient" : 0.0,
        "Target_Name" : "\"ATU2AF35A",
        "Stress" : 0,
        "BetweennessCentrality" : 0.0,
        "t_RNA_motif_Root" : "",
        "EdgeCount" : 1,
        "Causal_PLAMOM" : "",
        "name" : "AT1G27650",
        "Causal_tRNA" : "",
        "IsSingleNode" : false,
        "briefDescription" : "U2 snRNP auxiliary factor small subunit",
        "NeighborhoodConnectivity" : 6.0
      },
      "position" : {
        "x" : 3028.013622689421,
        "y" : 881.9261474609375
      },
      "selected" : false
    }, {
      "data" : {
        "id" : "1238",
        "symbol" : "UKL3",
        "Shoot_Descriptions" : "",
        "PLAMOM_Match_Shoot" : "",
        "Causal_TF" : "uridine kinase-like 3\"",
        "Root_Descriptions" : "UKL3, uridine kinase-like 3",
        "Eccentricity" : 11,
        "Outdegree" : 6,
        "Target_PLAMOM" : "",
        "PLAMOM_Match_Root" : "NO",
        "shared_name" : "AT1G55810",
        "Indegree" : 0,
        "Target_tRNA" : "",
        "t_RNA_motif_shoot" : "",
        "SUID" : 1238,
        "SelfLoops" : 0,
        "Target_TF" : "",
        "MolecularAttributes" : "METABOLIC",
        "AverageShortestPathLength" : 6.51552795,
        "selected" : false,
        "Name2" : "UKL3",
        "ClosenessCentrality" : 0.1534795,
        "Casual_Name" : "\"UKL3",
        "PartnerOfMultiEdgedNodePairs" : 0,
        "ClusteringCoefficient" : 0.0,
        "Target_Name" : "",
        "Stress" : 0,
        "BetweennessCentrality" : 0.0,
        "t_RNA_motif_Root" : "NO",
        "EdgeCount" : 6,
        "Causal_PLAMOM" : "no",
        "name" : "AT1G55810",
        "Causal_tRNA" : "no",
        "IsSingleNode" : false,
        "briefDescription" : "uridine kinase-like 3",
        "NeighborhoodConnectivity" : 8.333333333333334
      },
      "position" : {
        "x" : -5929.05712890625,
        "y" : 706.24462890625
      },
      "selected" : false
    }, {
      "data" : {
        "id" : "1237",
        "symbol" : "AT1G28060",
        "degree_layout" : 5,
        "Shoot_Descriptions" : "Pre-mRNA-splicing factor 3",
        "PLAMOM_Match_Shoot" : "NO",
        "Causal_TF" : "no",
        "Root_Descriptions" : "",
        "Eccentricity" : 12,
        "Outdegree" : 1,
        "Target_PLAMOM" : "no",
        "PLAMOM_Match_Root" : "",
        "shared_name" : "AT1G28060",
        "Indegree" : 6,
        "Target_tRNA" : "no",
        "t_RNA_motif_shoot" : "NO",
        "SUID" : 1237,
        "SelfLoops" : 0,
        "Target_TF" : "no",
        "MolecularAttributes" : "protein_coding",
        "AverageShortestPathLength" : 7.09414226,
        "selected" : false,
        "Name2" : "AT1G28060",
        "ClosenessCentrality" : 0.14096137,
        "Casual_Name" : "Pre-mRNA-splicing factor 3",
        "PartnerOfMultiEdgedNodePairs" : 0,
        "ClusteringCoefficient" : 0.04761905,
        "Target_Name" : "Pre-mRNA-splicing factor 3",
        "Stress" : 10396,
        "BetweennessCentrality" : 0.00141288,
        "t_RNA_motif_Root" : "",
        "EdgeCount" : 7,
        "Causal_PLAMOM" : "no",
        "name" : "AT1G28060",
        "Causal_tRNA" : "no",
        "IsSingleNode" : false,
        "briefDescription" : "Pre-mRNA-splicing factor 3",
        "NeighborhoodConnectivity" : 27.571428571428573
      },
      "position" : {
        "x" : 3367.478466439421,
        "y" : 3238.36669921875
      },
      "selected" : false
    }, {
      "data" : {
        "id" : "1236",
        "symbol" : "AT3G03790",
        "degree_layout" : 10,
        "Shoot_Descriptions" : "ankyrin repeat family protein / regulator of chromosome condensation (RCC1) family protein",
        "PLAMOM_Match_Shoot" : "NO",
        "Causal_TF" : "no",
        "Root_Descriptions" : "ankyrin repeat family protein / regulator of chromosome condensation (RCC1) family protein",
        "Eccentricity" : 12,
        "Outdegree" : 4,
        "Target_PLAMOM" : "no",
        "PLAMOM_Match_Root" : "NO",
        "shared_name" : "AT3G03790",
        "Indegree" : 20,
        "Target_tRNA" : "no",
        "t_RNA_motif_shoot" : "NO",
        "SUID" : 1236,
        "SelfLoops" : 0,
        "Target_TF" : "no",
        "MolecularAttributes" : "protein_coding",
        "AverageShortestPathLength" : 7.33891213,
        "selected" : false,
        "Name2" : "AT3G03790",
        "ClosenessCentrality" : 0.13625998,
        "Casual_Name" : "ankyrin repeat family protein / regulator of chromosome condensation (RCC1) family protein",
        "PartnerOfMultiEdgedNodePairs" : 0,
        "ClusteringCoefficient" : 0.01449275,
        "Target_Name" : "ankyrin repeat family protein / regulator of chromosome condensation (RCC1) family protein",
        "Stress" : 18487,
        "BetweennessCentrality" : 0.00384235,
        "t_RNA_motif_Root" : "NO",
        "EdgeCount" : 24,
        "Causal_PLAMOM" : "no",
        "name" : "AT3G03790",
        "Causal_tRNA" : "no",
        "IsSingleNode" : false,
        "briefDescription" : "ankyrin repeat family protein / regulator of chromosome condensation (RCC1) family protein",
        "NeighborhoodConnectivity" : 16.5
      },
      "position" : {
        "x" : -768.2392459629227,
        "y" : 354.3074645996094
      },
      "selected" : false
    }, {
      "data" : {
        "id" : "1235",
        "symbol" : "RD21A",
        "degree_layout" : 44,
        "Shoot_Descriptions" : "",
        "PLAMOM_Match_Shoot" : "",
        "Causal_TF" : "RD21A",
        "Root_Descriptions" : "RD21, RD21A, Granulin repeat cysteine protease family protein",
        "Eccentricity" : 9,
        "Outdegree" : 26,
        "Target_PLAMOM" : "AT1G47128",
        "PLAMOM_Match_Root" : "YES",
        "shared_name" : "AT1G47128",
        "Indegree" : 48,
        "Target_tRNA" : "no",
        "t_RNA_motif_shoot" : "",
        "SUID" : 1235,
        "SelfLoops" : 0,
        "Target_TF" : "RD21A",
        "MolecularAttributes" : "protein_coding",
        "AverageShortestPathLength" : 4.76359833,
        "selected" : false,
        "Name2" : "RD21A",
        "ClosenessCentrality" : 0.20992534,
        "Casual_Name" : "\"RD21",
        "PartnerOfMultiEdgedNodePairs" : 0,
        "ClusteringCoefficient" : 0.00740466,
        "Target_Name" : "\"RD21",
        "Stress" : 316856,
        "BetweennessCentrality" : 0.05960338,
        "t_RNA_motif_Root" : "NO",
        "EdgeCount" : 74,
        "Causal_PLAMOM" : "AT1G47128",
        "name" : "AT1G47128",
        "Causal_tRNA" : "no",
        "IsSingleNode" : false,
        "briefDescription" : "Granulin repeat cysteine protease family protein",
        "NeighborhoodConnectivity" : 9.256756756756756
      },
      "position" : {
        "x" : -6580.9560546875,
        "y" : 2921.198486328125
      },
      "selected" : false
    }, {
      "data" : {
        "id" : "1234",
        "symbol" : "AT5G10770",
        "degree_layout" : 14,
        "Shoot_Descriptions" : "Eukaryotic aspartyl protease family protein",
        "PLAMOM_Match_Shoot" : "NO",
        "Causal_TF" : "no",
        "Root_Descriptions" : "Eukaryotic aspartyl protease family protein",
        "Eccentricity" : 10,
        "Outdegree" : 14,
        "Target_PLAMOM" : "no",
        "PLAMOM_Match_Root" : "NO",
        "shared_name" : "AT5G10770",
        "Indegree" : 21,
        "Target_tRNA" : "no",
        "t_RNA_motif_shoot" : "NO",
        "SUID" : 1234,
        "SelfLoops" : 1,
        "Target_TF" : "no",
        "MolecularAttributes" : "protein_coding",
        "AverageShortestPathLength" : 5.72175732,
        "selected" : false,
        "Name2" : "AT5G10770",
        "ClosenessCentrality" : 0.17477148,
        "Casual_Name" : "Eukaryotic aspartyl protease family protein",
        "PartnerOfMultiEdgedNodePairs" : 0,
        "ClusteringCoefficient" : 0.00189394,
        "Target_Name" : "Eukaryotic aspartyl protease family protein",
        "Stress" : 54313,
        "BetweennessCentrality" : 0.01054585,
        "t_RNA_motif_Root" : "NO",
        "EdgeCount" : 35,
        "Causal_PLAMOM" : "no",
        "name" : "AT5G10770",
        "Causal_tRNA" : "no",
        "IsSingleNode" : false,
        "briefDescription" : "Eukaryotic aspartyl protease family protein",
        "NeighborhoodConnectivity" : 6.090909090909091
      },
      "position" : {
        "x" : -1647.474365234375,
        "y" : 2862.8268938690426
      },
      "selected" : false
    }, {
      "data" : {
        "id" : "1233",
        "symbol" : "RLP4",
        "Shoot_Descriptions" : "AtRLP4, RLP4, receptor like protein 4",
        "PLAMOM_Match_Shoot" : "NO",
        "Causal_TF" : "RLP4",
        "Root_Descriptions" : "",
        "Eccentricity" : 9,
        "Outdegree" : 3,
        "Target_PLAMOM" : "no",
        "PLAMOM_Match_Root" : "",
        "shared_name" : "AT1G28340",
        "Indegree" : 1,
        "Target_tRNA" : "no",
        "t_RNA_motif_shoot" : "NO",
        "SUID" : 1233,
        "SelfLoops" : 0,
        "Target_TF" : "RLP4",
        "MolecularAttributes" : "protein_coding",
        "AverageShortestPathLength" : 5.06694561,
        "selected" : false,
        "Name2" : "RLP4",
        "ClosenessCentrality" : 0.19735756,
        "Casual_Name" : "\"AtRLP4",
        "PartnerOfMultiEdgedNodePairs" : 0,
        "ClusteringCoefficient" : 0.0,
        "Target_Name" : "\"AtRLP4",
        "Stress" : 5154,
        "BetweennessCentrality" : 0.00113793,
        "t_RNA_motif_Root" : "",
        "EdgeCount" : 4,
        "Causal_PLAMOM" : "no",
        "name" : "AT1G28340",
        "Causal_tRNA" : "no",
        "IsSingleNode" : false,
        "briefDescription" : "receptor like protein 4",
        "NeighborhoodConnectivity" : 31.25
      },
      "position" : {
        "x" : 3530.438427376921,
        "y" : 2976.453857421875
      },
      "selected" : false
    }, {
      "data" : {
        "id" : "1232",
        "symbol" : "ACO2",
        "Shoot_Descriptions" : "",
        "PLAMOM_Match_Shoot" : "",
        "Causal_TF" : "aconitase 2\"",
        "Root_Descriptions" : "ACO2, aconitase 2",
        "Eccentricity" : 9,
        "Outdegree" : 13,
        "Target_PLAMOM" : "AT4G26970",
        "PLAMOM_Match_Root" : "YES",
        "shared_name" : "AT4G26970",
        "Indegree" : 1,
        "Target_tRNA" : "no",
        "t_RNA_motif_shoot" : "",
        "SUID" : 1232,
        "SelfLoops" : 0,
        "Target_TF" : "aconitase 2\"",
        "MolecularAttributes" : "METABOLIC",
        "AverageShortestPathLength" : 4.56485356,
        "selected" : false,
        "Name2" : "ACO2",
        "ClosenessCentrality" : 0.21906508,
        "Casual_Name" : "\"ACO2",
        "PartnerOfMultiEdgedNodePairs" : 0,
        "ClusteringCoefficient" : 0.0,
        "Target_Name" : "\"ACO2",
        "Stress" : 41208,
        "BetweennessCentrality" : 0.00816416,
        "t_RNA_motif_Root" : "NO",
        "EdgeCount" : 14,
        "Causal_PLAMOM" : "AT4G26970",
        "name" : "AT4G26970",
        "Causal_tRNA" : "no",
        "IsSingleNode" : false,
        "briefDescription" : "aconitase 2",
        "NeighborhoodConnectivity" : 6.142857142857143
      },
      "position" : {
        "x" : -6603.099609375,
        "y" : 3372.84912109375
      },
      "selected" : false
    }, {
      "data" : {
        "id" : "1231",
        "symbol" : "AT1G28510",
        "Shoot_Descriptions" : "Optic atrophy 3 protein (OPA3)",
        "PLAMOM_Match_Shoot" : "NO",
        "Causal_TF" : "no",
        "Root_Descriptions" : "",
        "Eccentricity" : 14,
        "Outdegree" : 1,
        "Target_PLAMOM" : "no",
        "PLAMOM_Match_Root" : "",
        "shared_name" : "AT1G28510",
        "Indegree" : 1,
        "Target_tRNA" : "no",
        "t_RNA_motif_shoot" : "NO",
        "SUID" : 1231,
        "SelfLoops" : 0,
        "Target_TF" : "Optic atrophy 3 protein (OPA3)",
        "MolecularAttributes" : "protein_coding",
        "AverageShortestPathLength" : 9.29288703,
        "selected" : false,
        "Name2" : "AT1G28510",
        "ClosenessCentrality" : 0.10760919,
        "Casual_Name" : "Optic atrophy 3 protein (OPA3)",
        "PartnerOfMultiEdgedNodePairs" : 0,
        "ClusteringCoefficient" : 0.0,
        "Target_Name" : "no",
        "Stress" : 2572,
        "BetweennessCentrality" : 4.8636E-4,
        "t_RNA_motif_Root" : "",
        "EdgeCount" : 2,
        "Causal_PLAMOM" : "no",
        "name" : "AT1G28510",
        "Causal_tRNA" : "no",
        "IsSingleNode" : false,
        "briefDescription" : "Optic atrophy 3 protein (OPA3)",
        "NeighborhoodConnectivity" : 8.0
      },
      "position" : {
        "x" : 3180.824657845671,
        "y" : 3367.69189453125
      },
      "selected" : false
    }, {
      "data" : {
        "id" : "1230",
        "symbol" : "AT1G20380",
        "Shoot_Descriptions" : "",
        "PLAMOM_Match_Shoot" : "",
        "Causal_TF" : "no",
        "Root_Descriptions" : "Prolyl oligopeptidase family protein",
        "Eccentricity" : 11,
        "Outdegree" : 5,
        "Target_PLAMOM" : "",
        "PLAMOM_Match_Root" : "NO",
        "shared_name" : "AT1G20380",
        "Indegree" : 0,
        "Target_tRNA" : "",
        "t_RNA_motif_shoot" : "",
        "SUID" : 1230,
        "SelfLoops" : 0,
        "Target_TF" : "",
        "MolecularAttributes" : "protein_coding",
        "AverageShortestPathLength" : 6.98547718,
        "selected" : false,
        "Name2" : "AT1G20380",
        "ClosenessCentrality" : 0.14315414,
        "Casual_Name" : "Prolyl oligopeptidase family protein",
        "PartnerOfMultiEdgedNodePairs" : 0,
        "ClusteringCoefficient" : 0.0,
        "Target_Name" : "",
        "Stress" : 0,
        "BetweennessCentrality" : 0.0,
        "t_RNA_motif_Root" : "NO",
        "EdgeCount" : 5,
        "Causal_PLAMOM" : "no",
        "name" : "AT1G20380",
        "Causal_tRNA" : "no",
        "IsSingleNode" : false,
        "briefDescription" : "Prolyl oligopeptidase family protein",
        "NeighborhoodConnectivity" : 5.2
      },
      "position" : {
        "x" : -6081.55322265625,
        "y" : -404.29742431640625
      },
      "selected" : false
    }, {
      "data" : {
        "id" : "1229",
        "symbol" : "ML5",
        "Shoot_Descriptions" : "AML5, ML5, MEI2-like protein 5",
        "PLAMOM_Match_Shoot" : "YES",
        "Causal_TF" : "ML5",
        "Root_Descriptions" : "",
        "Eccentricity" : 10,
        "Outdegree" : 2,
        "Target_PLAMOM" : "AT1G29400",
        "PLAMOM_Match_Root" : "",
        "shared_name" : "AT1G29400",
        "Indegree" : 2,
        "Target_tRNA" : "no",
        "t_RNA_motif_shoot" : "NO",
        "SUID" : 1229,
        "SelfLoops" : 0,
        "Target_TF" : "ML5",
        "MolecularAttributes" : "protein_coding",
        "AverageShortestPathLength" : 6.0460251,
        "selected" : false,
        "Name2" : "ML5",
        "ClosenessCentrality" : 0.16539792,
        "Casual_Name" : "\"AML5",
        "PartnerOfMultiEdgedNodePairs" : 0,
        "ClusteringCoefficient" : 0.0,
        "Target_Name" : "\"AML5",
        "Stress" : 1989,
        "BetweennessCentrality" : 5.6236E-4,
        "t_RNA_motif_Root" : "",
        "EdgeCount" : 4,
        "Causal_PLAMOM" : "AT1G29400",
        "name" : "AT1G29400",
        "Causal_tRNA" : "no",
        "IsSingleNode" : false,
        "briefDescription" : "MEI2-like protein 5",
        "NeighborhoodConnectivity" : 12.0
      },
      "position" : {
        "x" : 3635.232860970671,
        "y" : 2921.533935546875
      },
      "selected" : false
    }, {
      "data" : {
        "id" : "1228",
        "symbol" : "AT1G29465",
        "degree_layout" : 3,
        "Shoot_Descriptions" : "unknown protein; FUNCTIONS IN: molecular_function unknown; INVOLVED IN: biological_process unknown; LOCATED IN: endomembrane system; Has 30201 Blast hits to 17322 proteins in 780 species: Archae - 12; Bacteria - 1396; Metazoa - 17338; Fungi - 3422; Plants - 5037; Viruses - 0; Other Eukaryotes - 2996 (source: NCBI BLink).",
        "PLAMOM_Match_Shoot" : "NO",
        "Causal_TF" : "no",
        "Root_Descriptions" : "",
        "Eccentricity" : 10,
        "Outdegree" : 1,
        "Target_PLAMOM" : "no",
        "PLAMOM_Match_Root" : "",
        "shared_name" : "AT1G29465",
        "Indegree" : 12,
        "Target_tRNA" : "no",
        "t_RNA_motif_shoot" : "NO",
        "SUID" : 1228,
        "SelfLoops" : 0,
        "Target_TF" : "unknown protein; FUNCTIONS IN: molecular_function unknown; INVOLVED IN: biological_process unknown; LOCATED IN: endomembrane system; Has 30201 Blast hits to 17322 proteins in 780 species: Archae - 12; Bacteria - 1396; Metazoa - 17338; Fungi - 3422; Plants - 5037; Viruses - 0; Other Eukaryotes - 2996 (source: NCBI BLink).",
        "MolecularAttributes" : "protein_coding",
        "AverageShortestPathLength" : 5.38075314,
        "selected" : false,
        "Name2" : "AT1G29465",
        "ClosenessCentrality" : 0.18584759,
        "Casual_Name" : "unknown protein; FUNCTIONS IN: molecular_function unknown; INVOLVED IN: biological_process unknown; LOCATED IN: endomembrane system; Has 30201 Blast hits to 17322 proteins in 780 species: Archae - 12; Bacteria - 1396; Metazoa - 17338; Fungi - 3422; Plants - 5037; Viruses - 0; Other Eukaryotes - 2996 (source: NCBI BLink).",
        "PartnerOfMultiEdgedNodePairs" : 0,
        "ClusteringCoefficient" : 0.0,
        "Target_Name" : "no",
        "Stress" : 10291,
        "BetweennessCentrality" : 0.00237106,
        "t_RNA_motif_Root" : "",
        "EdgeCount" : 13,
        "Causal_PLAMOM" : "no",
        "name" : "AT1G29465",
        "Causal_tRNA" : "no",
        "IsSingleNode" : false,
        "briefDescription" : "transmembrane protein",
        "NeighborhoodConnectivity" : 13.0
      },
      "position" : {
        "x" : 3594.895946908171,
        "y" : 3265.39990234375
      },
      "selected" : false
    }, {
      "data" : {
        "id" : "1227",
        "symbol" : "AT5G24890",
        "Shoot_Descriptions" : "",
        "PLAMOM_Match_Shoot" : "",
        "Causal_TF" : "no",
        "Root_Descriptions" : "unknown protein; FUNCTIONS IN: molecular_function unknown; INVOLVED IN: biological_process unknown; LOCATED IN: cellular_component unknown; EXPRESSED IN: 24 plant structures; EXPRESSED DURING: 13 growth stages; BEST Arabidopsis thaliana protein match is: unknown protein (TAIR:AT2G24550.1); Has 1807 Blast hits to 1807 proteins in 277 species: Archae - 0; Bacteria - 0; Metazoa - 736; Fungi - 347; Plants - 385; Viruses - 0; Other Eukaryotes - 339 (source: NCBI BLink).",
        "Eccentricity" : 9,
        "Outdegree" : 7,
        "Target_PLAMOM" : "NAC domain containing protein 96\"",
        "PLAMOM_Match_Root" : "NO",
        "shared_name" : "AT5G24890",
        "Indegree" : 2,
        "Target_tRNA" : "AT5G46590",
        "t_RNA_motif_shoot" : "",
        "SUID" : 1227,
        "SelfLoops" : 0,
        "Target_TF" : "no",
        "MolecularAttributes" : "protein_coding",
        "AverageShortestPathLength" : 4.84551148,
        "selected" : false,
        "Name2" : "AT5G24890",
        "ClosenessCentrality" : 0.20637656,
        "Casual_Name" : "unknown protein; FUNCTIONS IN: molecular_function unknown; INVOLVED IN: biological_process unknown; LOCATED IN: cellular_component unknown; EXPRESSED IN: 24 plant structures; EXPRESSED DURING: 13 growth stages; BEST Arabidopsis thaliana protein match is: unknown protein (TAIR:AT2G24550.1); Has 1807 Blast hits to 1807 proteins in 277 species: Archae - 0; Bacteria - 0; Metazoa - 736; Fungi - 347; Plants - 385; Viruses - 0; Other Eukaryotes - 339 (source: NCBI BLink).",
        "PartnerOfMultiEdgedNodePairs" : 0,
        "ClusteringCoefficient" : 0.0,
        "Target_Name" : "no",
        "Stress" : 501,
        "BetweennessCentrality" : 1.4564E-4,
        "t_RNA_motif_Root" : "NO",
        "EdgeCount" : 9,
        "Causal_PLAMOM" : "no",
        "name" : "AT5G24890",
        "Causal_tRNA" : "no",
        "IsSingleNode" : false,
        "briefDescription" : "stress response NST1-like protein",
        "NeighborhoodConnectivity" : 8.555555555555555
      },
      "position" : {
        "x" : -6473.44287109375,
        "y" : 3210.504150390625
      },
      "selected" : false
    }, {
      "data" : {
        "id" : "1226",
        "symbol" : "APC2",
        "Shoot_Descriptions" : "Mitochondrial substrate carrier family protein",
        "PLAMOM_Match_Shoot" : "NO",
        "Causal_TF" : "no",
        "Root_Descriptions" : "Mitochondrial substrate carrier family protein",
        "Eccentricity" : 10,
        "Outdegree" : 8,
        "Target_PLAMOM" : "no",
        "PLAMOM_Match_Root" : "NO",
        "shared_name" : "AT5G51050",
        "Indegree" : 1,
        "Target_tRNA" : "no",
        "t_RNA_motif_shoot" : "NO",
        "SUID" : 1226,
        "SelfLoops" : 0,
        "Target_TF" : "no",
        "MolecularAttributes" : "protein_coding",
        "AverageShortestPathLength" : 5.76041667,
        "selected" : false,
        "Name2" : "APC2",
        "ClosenessCentrality" : 0.17359855,
        "Casual_Name" : "Mitochondrial substrate carrier family protein",
        "PartnerOfMultiEdgedNodePairs" : 0,
        "ClusteringCoefficient" : 0.0,
        "Target_Name" : "Mitochondrial substrate carrier family protein",
        "Stress" : 972,
        "BetweennessCentrality" : 2.8644E-4,
        "t_RNA_motif_Root" : "NO",
        "EdgeCount" : 9,
        "Causal_PLAMOM" : "no",
        "name" : "AT5G51050",
        "Causal_tRNA" : "no",
        "IsSingleNode" : false,
        "briefDescription" : "Mitochondrial substrate carrier family protein",
        "NeighborhoodConnectivity" : 9.555555555555555
      },
      "position" : {
        "x" : -978.8383684784176,
        "y" : -172.53237915039062
      },
      "selected" : false
    }, {
      "data" : {
        "id" : "1225",
        "symbol" : "ABF3",
        "Shoot_Descriptions" : "",
        "PLAMOM_Match_Shoot" : "",
        "Causal_TF" : "DPBF5",
        "Root_Descriptions" : "ABF3, DPBF5, abscisic acid responsive elements-binding factor 3",
        "Eccentricity" : 8,
        "Outdegree" : 11,
        "Target_PLAMOM" : "no",
        "PLAMOM_Match_Root" : "NO",
        "shared_name" : "AT4G34000",
        "Indegree" : 6,
        "Target_tRNA" : "no",
        "t_RNA_motif_shoot" : "",
        "SUID" : 1225,
        "SelfLoops" : 0,
        "Target_TF" : "DPBF5",
        "MolecularAttributes" : "TXNFACTOR",
        "AverageShortestPathLength" : 4.69456067,
        "selected" : false,
        "Name2" : "ABF3",
        "ClosenessCentrality" : 0.21301248,
        "Casual_Name" : "\"ABF3",
        "PartnerOfMultiEdgedNodePairs" : 0,
        "ClusteringCoefficient" : 0.00367647,
        "Target_Name" : "\"ABF3",
        "Stress" : 26699,
        "BetweennessCentrality" : 0.00431481,
        "t_RNA_motif_Root" : "NO",
        "EdgeCount" : 17,
        "Causal_PLAMOM" : "no",
        "name" : "AT4G34000",
        "Causal_tRNA" : "no",
        "IsSingleNode" : false,
        "briefDescription" : "abscisic acid responsive elements-binding factor 3",
        "NeighborhoodConnectivity" : 6.9411764705882355
      },
      "position" : {
        "x" : -6299.02880859375,
        "y" : 3181.927734375
      },
      "selected" : false
    }, {
      "data" : {
        "id" : "1224",
        "symbol" : "AT1G15640",
        "Shoot_Descriptions" : "",
        "PLAMOM_Match_Shoot" : "",
        "Causal_TF" : "no",
        "Root_Descriptions" : "unknown protein; BEST Arabidopsis thaliana protein match is: unknown protein (TAIR:AT1G15620.1); Has 154 Blast hits to 146 proteins in 13 species: Archae - 0; Bacteria - 0; Metazoa - 0; Fungi - 0; Plants - 154; Viruses - 0; Other Eukaryotes - 0 (source: NCBI BLink).",
        "Eccentricity" : 9,
        "Outdegree" : 10,
        "Target_PLAMOM" : "",
        "PLAMOM_Match_Root" : "NO",
        "shared_name" : "AT1G15640",
        "Indegree" : 0,
        "Target_tRNA" : "",
        "t_RNA_motif_shoot" : "",
        "SUID" : 1224,
        "SelfLoops" : 0,
        "Target_TF" : "",
        "MolecularAttributes" : "protein_coding",
        "AverageShortestPathLength" : 4.975,
        "selected" : false,
        "Name2" : "AT1G15640",
        "ClosenessCentrality" : 0.20100503,
        "Casual_Name" : "unknown protein; BEST Arabidopsis thaliana protein match is: unknown protein (TAIR:AT1G15620.1); Has 154 Blast hits to 146 proteins in 13 species: Archae - 0; Bacteria - 0; Metazoa - 0; Fungi - 0; Plants - 154; Viruses - 0; Other Eukaryotes - 0 (source: NCBI BLink).",
        "PartnerOfMultiEdgedNodePairs" : 0,
        "ClusteringCoefficient" : 0.0,
        "Target_Name" : "",
        "Stress" : 0,
        "BetweennessCentrality" : 0.0,
        "t_RNA_motif_Root" : "YES",
        "EdgeCount" : 10,
        "Causal_PLAMOM" : "no",
        "name" : "AT1G15640",
        "Causal_tRNA" : "AT1G15640",
        "IsSingleNode" : false,
        "briefDescription" : "transmembrane protein",
        "NeighborhoodConnectivity" : 11.8
      },
      "position" : {
        "x" : -6025.3515625,
        "y" : 125.48240661621094
      },
      "selected" : false
    }, {
      "data" : {
        "id" : "1223",
        "symbol" : "RLP24",
        "Shoot_Descriptions" : "",
        "PLAMOM_Match_Shoot" : "",
        "Causal_TF" : "RLP24",
        "Root_Descriptions" : "AtRLP24, RLP24, receptor like protein 24",
        "Eccentricity" : 9,
        "Outdegree" : 3,
        "Target_PLAMOM" : "",
        "PLAMOM_Match_Root" : "NO",
        "shared_name" : "AT2G33020",
        "Indegree" : 0,
        "Target_tRNA" : "",
        "t_RNA_motif_shoot" : "",
        "SUID" : 1223,
        "SelfLoops" : 0,
        "Target_TF" : "",
        "MolecularAttributes" : "protein_coding",
        "AverageShortestPathLength" : 5.35416667,
        "selected" : false,
        "Name2" : "RLP24",
        "ClosenessCentrality" : 0.18677043,
        "Casual_Name" : "\"AtRLP24",
        "PartnerOfMultiEdgedNodePairs" : 0,
        "ClusteringCoefficient" : 0.0,
        "Target_Name" : "",
        "Stress" : 0,
        "BetweennessCentrality" : 0.0,
        "t_RNA_motif_Root" : "NO",
        "EdgeCount" : 3,
        "Causal_PLAMOM" : "no",
        "name" : "AT2G33020",
        "Causal_tRNA" : "no",
        "IsSingleNode" : false,
        "briefDescription" : "receptor like protein 24",
        "NeighborhoodConnectivity" : 10.666666666666666
      },
      "position" : {
        "x" : -5827.65673828125,
        "y" : 372.9738464355469
      },
      "selected" : false
    }, {
      "data" : {
        "id" : "1222",
        "symbol" : "ROT3",
        "Shoot_Descriptions" : "ROT3, Cytochrome P450 superfamily protein",
        "PLAMOM_Match_Shoot" : "NO",
        "Causal_TF" : "Cytochrome P450 superfamily protein\"",
        "Root_Descriptions" : "ROT3, Cytochrome P450 superfamily protein",
        "Eccentricity" : 11,
        "Outdegree" : 4,
        "Target_PLAMOM" : "no",
        "PLAMOM_Match_Root" : "NO",
        "shared_name" : "AT4G36380",
        "Indegree" : 1,
        "Target_tRNA" : "no",
        "t_RNA_motif_shoot" : "NO",
        "SUID" : 1222,
        "SelfLoops" : 0,
        "Target_TF" : "Cytochrome P450 superfamily protein\"",
        "MolecularAttributes" : "METABOLIC",
        "AverageShortestPathLength" : 6.01461378,
        "selected" : false,
        "Name2" : "ROT3",
        "ClosenessCentrality" : 0.16626171,
        "Casual_Name" : "\"ROT3",
        "PartnerOfMultiEdgedNodePairs" : 0,
        "ClusteringCoefficient" : 0.0,
        "Target_Name" : "\"ROT3",
        "Stress" : 287,
        "BetweennessCentrality" : 9.956E-5,
        "t_RNA_motif_Root" : "NO",
        "EdgeCount" : 5,
        "Causal_PLAMOM" : "no",
        "name" : "AT4G36380",
        "Causal_tRNA" : "no",
        "IsSingleNode" : false,
        "briefDescription" : "Cytochrome P450 superfamily protein",
        "NeighborhoodConnectivity" : 10.4
      },
      "position" : {
        "x" : -511.1280399082352,
        "y" : 116.91905212402344
      },
      "selected" : false
    }, {
      "data" : {
        "id" : "1221",
        "symbol" : "NUDT18",
        "Shoot_Descriptions" : "",
        "PLAMOM_Match_Shoot" : "",
        "Causal_TF" : "NUDT18",
        "Root_Descriptions" : "atnudt18, NUDT18, nudix hydrolase homolog 18",
        "Eccentricity" : 12,
        "Outdegree" : 11,
        "Target_PLAMOM" : "no",
        "PLAMOM_Match_Root" : "NO",
        "shared_name" : "AT1G14860",
        "Indegree" : 7,
        "Target_tRNA" : "no",
        "t_RNA_motif_shoot" : "",
        "SUID" : 1221,
        "SelfLoops" : 0,
        "Target_TF" : "NUDT18",
        "MolecularAttributes" : "protein_coding",
        "AverageShortestPathLength" : 7.28661088,
        "selected" : false,
        "Name2" : "NUDT18",
        "ClosenessCentrality" : 0.13723801,
        "Casual_Name" : "\"atnudt18",
        "PartnerOfMultiEdgedNodePairs" : 0,
        "ClusteringCoefficient" : 0.00980392,
        "Target_Name" : "\"atnudt18",
        "Stress" : 19555,
        "BetweennessCentrality" : 0.00406331,
        "t_RNA_motif_Root" : "NO",
        "EdgeCount" : 18,
        "Causal_PLAMOM" : "no",
        "name" : "AT1G14860",
        "Causal_tRNA" : "no",
        "IsSingleNode" : false,
        "briefDescription" : "nudix hydrolase homolog 18",
        "NeighborhoodConnectivity" : 9.666666666666666
      },
      "position" : {
        "x" : -6957.4716796875,
        "y" : 3126.82421875
      },
      "selected" : false
    }, {
      "data" : {
        "id" : "1220",
        "symbol" : "CIPK23",
        "degree_layout" : 3,
        "Shoot_Descriptions" : "ATCIPK23, CIPK23, LKS1, SnRK3.23, CBL-interacting protein kinase 23",
        "PLAMOM_Match_Shoot" : "YES",
        "Causal_TF" : "CIPK23",
        "Root_Descriptions" : "ATCIPK23, CIPK23, LKS1, SnRK3.23, CBL-interacting protein kinase 23",
        "Eccentricity" : 13,
        "Outdegree" : 3,
        "Target_PLAMOM" : "AT1G30270",
        "PLAMOM_Match_Root" : "YES",
        "shared_name" : "AT1G30270",
        "Indegree" : 13,
        "Target_tRNA" : "no",
        "t_RNA_motif_shoot" : "NO",
        "SUID" : 1220,
        "SelfLoops" : 0,
        "Target_TF" : "CIPK23",
        "MolecularAttributes" : "METABOLIC",
        "AverageShortestPathLength" : 8.4665272,
        "selected" : false,
        "Name2" : "CIPK23",
        "ClosenessCentrality" : 0.11811218,
        "Casual_Name" : "\"ATCIPK23",
        "PartnerOfMultiEdgedNodePairs" : 0,
        "ClusteringCoefficient" : 0.00833333,
        "Target_Name" : "\"ATCIPK23",
        "Stress" : 6650,
        "BetweennessCentrality" : 0.00121311,
        "t_RNA_motif_Root" : "NO",
        "EdgeCount" : 16,
        "Causal_PLAMOM" : "AT1G30270",
        "name" : "AT1G30270",
        "Causal_tRNA" : "no",
        "IsSingleNode" : false,
        "briefDescription" : "CBL-interacting protein kinase 23",
        "NeighborhoodConnectivity" : 10.375
      },
      "position" : {
        "x" : -1261.034717154329,
        "y" : 3245.541015625
      },
      "selected" : false
    }, {
      "data" : {
        "id" : "1219",
        "symbol" : "MEE14",
        "Shoot_Descriptions" : "",
        "PLAMOM_Match_Shoot" : "",
        "Causal_TF" : "maternal effect embryo arrest 14\"",
        "Root_Descriptions" : "MEE14, maternal effect embryo arrest 14",
        "Eccentricity" : 12,
        "Outdegree" : 4,
        "Target_PLAMOM" : "",
        "PLAMOM_Match_Root" : "YES",
        "shared_name" : "AT2G15890",
        "Indegree" : 0,
        "Target_tRNA" : "",
        "t_RNA_motif_shoot" : "",
        "SUID" : 1219,
        "SelfLoops" : 0,
        "Target_TF" : "",
        "MolecularAttributes" : "protein_coding",
        "AverageShortestPathLength" : 7.32985386,
        "selected" : false,
        "Name2" : "MEE14",
        "ClosenessCentrality" : 0.13642837,
        "Casual_Name" : "\"MEE14",
        "PartnerOfMultiEdgedNodePairs" : 0,
        "ClusteringCoefficient" : 0.0,
        "Target_Name" : "",
        "Stress" : 0,
        "BetweennessCentrality" : 0.0,
        "t_RNA_motif_Root" : "NO",
        "EdgeCount" : 4,
        "Causal_PLAMOM" : "AT2G15890",
        "name" : "AT2G15890",
        "Causal_tRNA" : "no",
        "IsSingleNode" : false,
        "briefDescription" : "maternal effect embryo arrest 14",
        "NeighborhoodConnectivity" : 11.0
      },
      "position" : {
        "x" : -6415.45947265625,
        "y" : 247.41543579101562
      },
      "selected" : false
    }, {
      "data" : {
        "id" : "1218",
        "symbol" : "AT2G37750",
        "Shoot_Descriptions" : "",
        "PLAMOM_Match_Shoot" : "",
        "Causal_TF" : "no",
        "Root_Descriptions" : "unknown protein; Has 21 Blast hits to 21 proteins in 7 species: Archae - 0; Bacteria - 0; Metazoa - 0; Fungi - 0; Plants - 21; Viruses - 0; Other Eukaryotes - 0 (source: NCBI BLink).",
        "Eccentricity" : 10,
        "Outdegree" : 4,
        "Target_PLAMOM" : "",
        "PLAMOM_Match_Root" : "NO",
        "shared_name" : "AT2G37750",
        "Indegree" : 0,
        "Target_tRNA" : "",
        "t_RNA_motif_shoot" : "",
        "SUID" : 1218,
        "SelfLoops" : 0,
        "Target_TF" : "",
        "MolecularAttributes" : "protein_coding",
        "AverageShortestPathLength" : 6.30416667,
        "selected" : false,
        "Name2" : "AT2G37750",
        "ClosenessCentrality" : 0.15862525,
        "Casual_Name" : "unknown protein; Has 21 Blast hits to 21 proteins in 7 species: Archae - 0; Bacteria - 0; Metazoa - 0; Fungi - 0; Plants - 21; Viruses - 0; Other Eukaryotes - 0 (source: NCBI BLink).",
        "PartnerOfMultiEdgedNodePairs" : 0,
        "ClusteringCoefficient" : 0.0,
        "Target_Name" : "",
        "Stress" : 0,
        "BetweennessCentrality" : 0.0,
        "t_RNA_motif_Root" : "NO",
        "EdgeCount" : 4,
        "Causal_PLAMOM" : "no",
        "name" : "AT2G37750",
        "Causal_tRNA" : "no",
        "IsSingleNode" : false,
        "briefDescription" : "hypothetical protein",
        "NeighborhoodConnectivity" : 15.25
      },
      "position" : {
        "x" : -6270.26318359375,
        "y" : 294.83148193359375
      },
      "selected" : false
    }, {
      "data" : {
        "id" : "1217",
        "symbol" : "AT5G27920",
        "Shoot_Descriptions" : "",
        "PLAMOM_Match_Shoot" : "",
        "Causal_TF" : "no",
        "Root_Descriptions" : "F-box family protein",
        "Eccentricity" : 12,
        "Outdegree" : 3,
        "Target_PLAMOM" : "dihydrodipicolinate synthase 1\"",
        "PLAMOM_Match_Root" : "NO",
        "shared_name" : "AT5G27920",
        "Indegree" : 3,
        "Target_tRNA" : "no",
        "t_RNA_motif_shoot" : "",
        "SUID" : 1217,
        "SelfLoops" : 0,
        "Target_TF" : "no",
        "MolecularAttributes" : "protein_coding",
        "AverageShortestPathLength" : 7.50627615,
        "selected" : false,
        "Name2" : "AT5G27920",
        "ClosenessCentrality" : 0.13322185,
        "Casual_Name" : "F-box family protein",
        "PartnerOfMultiEdgedNodePairs" : 0,
        "ClusteringCoefficient" : 0.0,
        "Target_Name" : "no",
        "Stress" : 14595,
        "BetweennessCentrality" : 0.003324,
        "t_RNA_motif_Root" : "NO",
        "EdgeCount" : 6,
        "Causal_PLAMOM" : "no",
        "name" : "AT5G27920",
        "Causal_tRNA" : "no",
        "IsSingleNode" : false,
        "briefDescription" : "F-box family protein",
        "NeighborhoodConnectivity" : 5.833333333333333
      },
      "position" : {
        "x" : -6482.67822265625,
        "y" : 3407.08642578125
      },
      "selected" : false
    }, {
      "data" : {
        "id" : "1216",
        "symbol" : "AT5G54170",
        "degree_layout" : 15,
        "Shoot_Descriptions" : "Polyketide cyclase/dehydrase and lipid transport superfamily protein",
        "PLAMOM_Match_Shoot" : "NO",
        "Causal_TF" : "no",
        "Root_Descriptions" : "Polyketide cyclase/dehydrase and lipid transport superfamily protein",
        "Eccentricity" : 9,
        "Outdegree" : 9,
        "Target_PLAMOM" : "no",
        "PLAMOM_Match_Root" : "NO",
        "shared_name" : "AT5G54170",
        "Indegree" : 6,
        "Target_tRNA" : "no",
        "t_RNA_motif_shoot" : "NO",
        "SUID" : 1216,
        "SelfLoops" : 0,
        "Target_TF" : "no",
        "MolecularAttributes" : "protein_coding",
        "AverageShortestPathLength" : 4.86401674,
        "selected" : false,
        "Name2" : "AT5G54170",
        "ClosenessCentrality" : 0.2055914,
        "Casual_Name" : "Polyketide cyclase/dehydrase and lipid transport superfamily protein",
        "PartnerOfMultiEdgedNodePairs" : 0,
        "ClusteringCoefficient" : 0.02380952,
        "Target_Name" : "Polyketide cyclase/dehydrase and lipid transport superfamily protein",
        "Stress" : 50323,
        "BetweennessCentrality" : 0.01084954,
        "t_RNA_motif_Root" : "NO",
        "EdgeCount" : 15,
        "Causal_PLAMOM" : "no",
        "name" : "AT5G54170",
        "Causal_tRNA" : "no",
        "IsSingleNode" : false,
        "briefDescription" : "Polyketide cyclase/dehydrase and lipid transport superfamily protein",
        "NeighborhoodConnectivity" : 29.733333333333334
      },
      "position" : {
        "x" : -1153.330859732454,
        "y" : 3144.85498046875
      },
      "selected" : false
    }, {
      "data" : {
        "id" : "1215",
        "symbol" : "IRT3",
        "Shoot_Descriptions" : "",
        "PLAMOM_Match_Shoot" : "",
        "Causal_TF" : "IRT3",
        "Root_Descriptions" : "ATIRT3, IRT3, iron regulated transporter 3",
        "Eccentricity" : 12,
        "Outdegree" : 7,
        "Target_PLAMOM" : "",
        "PLAMOM_Match_Root" : "NO",
        "shared_name" : "AT1G60960",
        "Indegree" : 0,
        "Target_tRNA" : "",
        "t_RNA_motif_shoot" : "",
        "SUID" : 1215,
        "SelfLoops" : 0,
        "Target_TF" : "",
        "MolecularAttributes" : "protein_coding",
        "AverageShortestPathLength" : 6.95824635,
        "selected" : false,
        "Name2" : "IRT3",
        "ClosenessCentrality" : 0.14371437,
        "Casual_Name" : "\"ATIRT3",
        "PartnerOfMultiEdgedNodePairs" : 0,
        "ClusteringCoefficient" : 0.0,
        "Target_Name" : "",
        "Stress" : 0,
        "BetweennessCentrality" : 0.0,
        "t_RNA_motif_Root" : "NO",
        "EdgeCount" : 7,
        "Causal_PLAMOM" : "no",
        "name" : "AT1G60960",
        "Causal_tRNA" : "no",
        "IsSingleNode" : false,
        "briefDescription" : "iron regulated transporter 3",
        "NeighborhoodConnectivity" : 10.428571428571429
      },
      "position" : {
        "x" : -6392.291015625,
        "y" : 448.0351257324219
      },
      "selected" : false
    }, {
      "data" : {
        "id" : "1214",
        "symbol" : "AT3G13070",
        "Shoot_Descriptions" : "CBS domain-containing protein / transporter associated domain-containing protein",
        "PLAMOM_Match_Shoot" : "YES",
        "Causal_TF" : "no",
        "Root_Descriptions" : "CBS domain-containing protein / transporter associated domain-containing protein",
        "Eccentricity" : 11,
        "Outdegree" : 7,
        "Target_PLAMOM" : "",
        "PLAMOM_Match_Root" : "YES",
        "shared_name" : "AT3G13070",
        "Indegree" : 0,
        "Target_tRNA" : "",
        "t_RNA_motif_shoot" : "NO",
        "SUID" : 1214,
        "SelfLoops" : 0,
        "Target_TF" : "",
        "MolecularAttributes" : "protein_coding",
        "AverageShortestPathLength" : 6.11458333,
        "selected" : false,
        "Name2" : "AT3G13070",
        "ClosenessCentrality" : 0.16354344,
        "Casual_Name" : "CBS domain-containing protein / transporter associated domain-containing protein",
        "PartnerOfMultiEdgedNodePairs" : 0,
        "ClusteringCoefficient" : 0.0,
        "Target_Name" : "",
        "Stress" : 0,
        "BetweennessCentrality" : 0.0,
        "t_RNA_motif_Root" : "NO",
        "EdgeCount" : 7,
        "Causal_PLAMOM" : "AT3G13070",
        "name" : "AT3G13070",
        "Causal_tRNA" : "no",
        "IsSingleNode" : false,
        "briefDescription" : "CBS domain-containing protein / transporter associated domain-containing protein",
        "NeighborhoodConnectivity" : 15.857142857142858
      },
      "position" : {
        "x" : -1049.1327396152665,
        "y" : 3364.22412109375
      },
      "selected" : false
    }, {
      "data" : {
        "id" : "1213",
        "symbol" : "IMD3",
        "Shoot_Descriptions" : "ATIMD3, IMD3, IPMDH1, isopropylmalate dehydrogenase 3",
        "PLAMOM_Match_Shoot" : "YES",
        "Causal_TF" : "",
        "Root_Descriptions" : "",
        "Eccentricity" : 0,
        "Outdegree" : 0,
        "Target_PLAMOM" : "AT1G31180",
        "PLAMOM_Match_Root" : "",
        "shared_name" : "AT1G31180",
        "Indegree" : 1,
        "Target_tRNA" : "no",
        "t_RNA_motif_shoot" : "NO",
        "SUID" : 1213,
        "SelfLoops" : 0,
        "Target_TF" : "IMD3",
        "MolecularAttributes" : "METABOLIC",
        "AverageShortestPathLength" : 0.0,
        "selected" : false,
        "Name2" : "IMD3",
        "ClosenessCentrality" : 0.0,
        "Casual_Name" : "",
        "PartnerOfMultiEdgedNodePairs" : 0,
        "ClusteringCoefficient" : 0.0,
        "Target_Name" : "\"ATIMD3",
        "Stress" : 0,
        "BetweennessCentrality" : 0.0,
        "t_RNA_motif_Root" : "",
        "EdgeCount" : 1,
        "Causal_PLAMOM" : "",
        "name" : "AT1G31180",
        "Causal_tRNA" : "",
        "IsSingleNode" : false,
        "briefDescription" : "isopropylmalate dehydrogenase 3",
        "NeighborhoodConnectivity" : 7.0
      },
      "position" : {
        "x" : 4280.211924490332,
        "y" : 247.08761596679688
      },
      "selected" : false
    }, {
      "data" : {
        "id" : "1212",
        "symbol" : "AK-HSDH",
        "Shoot_Descriptions" : "AK-HSDH, AK-HSDH I, aspartate kinase-homoserine dehydrogenase i",
        "PLAMOM_Match_Shoot" : "NO",
        "Causal_TF" : "AK-HSDH I",
        "Root_Descriptions" : "",
        "Eccentricity" : 9,
        "Outdegree" : 1,
        "Target_PLAMOM" : "no",
        "PLAMOM_Match_Root" : "",
        "shared_name" : "AT1G31230",
        "Indegree" : 4,
        "Target_tRNA" : "no",
        "t_RNA_motif_shoot" : "YES",
        "SUID" : 1212,
        "SelfLoops" : 0,
        "Target_TF" : "\"AK-HSDH",
        "MolecularAttributes" : "METABOLIC",
        "AverageShortestPathLength" : 4.43933054,
        "selected" : false,
        "Name2" : "AK-HSDH I",
        "ClosenessCentrality" : 0.22525919,
        "Casual_Name" : "\"AK-HSDH",
        "PartnerOfMultiEdgedNodePairs" : 0,
        "ClusteringCoefficient" : 0.0,
        "Target_Name" : "AT1G31230",
        "Stress" : 11511,
        "BetweennessCentrality" : 0.00229918,
        "t_RNA_motif_Root" : "",
        "EdgeCount" : 5,
        "Causal_PLAMOM" : "no",
        "name" : "AT1G31230",
        "Causal_tRNA" : "AT1G31230",
        "IsSingleNode" : false,
        "briefDescription" : "aspartate kinase-homoserine dehydrogenase i",
        "NeighborhoodConnectivity" : 23.0
      },
      "position" : {
        "x" : 3512.223095345671,
        "y" : 3521.635009765625
      },
      "selected" : false
    }, {
      "data" : {
        "id" : "1211",
        "symbol" : "AT4G31320",
        "Shoot_Descriptions" : "",
        "PLAMOM_Match_Shoot" : "",
        "Causal_TF" : "no",
        "Root_Descriptions" : "SAUR-like auxin-responsive protein family",
        "Eccentricity" : 8,
        "Outdegree" : 16,
        "Target_PLAMOM" : "",
        "PLAMOM_Match_Root" : "NO",
        "shared_name" : "AT4G31320",
        "Indegree" : 0,
        "Target_tRNA" : "",
        "t_RNA_motif_shoot" : "",
        "SUID" : 1211,
        "SelfLoops" : 0,
        "Target_TF" : "",
        "MolecularAttributes" : "protein_coding",
        "AverageShortestPathLength" : 3.8946281,
        "selected" : false,
        "Name2" : "AT4G31320",
        "ClosenessCentrality" : 0.25676393,
        "Casual_Name" : "SAUR-like auxin-responsive protein family",
        "PartnerOfMultiEdgedNodePairs" : 0,
        "ClusteringCoefficient" : 0.0,
        "Target_Name" : "",
        "Stress" : 0,
        "BetweennessCentrality" : 0.0,
        "t_RNA_motif_Root" : "NO",
        "EdgeCount" : 16,
        "Causal_PLAMOM" : "no",
        "name" : "AT4G31320",
        "Causal_tRNA" : "no",
        "IsSingleNode" : false,
        "briefDescription" : "SAUR-like auxin-responsive protein family",
        "NeighborhoodConnectivity" : 8.5625
      },
      "position" : {
        "x" : -5859.22412109375,
        "y" : 275.9677429199219
      },
      "selected" : false
    }, {
      "data" : {
        "id" : "1210",
        "symbol" : "NIC3",
        "Shoot_Descriptions" : "",
        "PLAMOM_Match_Shoot" : "",
        "Causal_TF" : "nicotinamidase 3\"",
        "Root_Descriptions" : "NIC3, nicotinamidase 3",
        "Eccentricity" : 10,
        "Outdegree" : 1,
        "Target_PLAMOM" : "response regulator 9\"",
        "PLAMOM_Match_Root" : "NO",
        "shared_name" : "AT5G23220",
        "Indegree" : 3,
        "Target_tRNA" : "no",
        "t_RNA_motif_shoot" : "",
        "SUID" : 1210,
        "SelfLoops" : 0,
        "Target_TF" : "no",
        "MolecularAttributes" : "METABOLIC",
        "AverageShortestPathLength" : 5.43006263,
        "selected" : false,
        "Name2" : "NIC3",
        "ClosenessCentrality" : 0.18415994,
        "Casual_Name" : "\"NIC3",
        "PartnerOfMultiEdgedNodePairs" : 0,
        "ClusteringCoefficient" : 0.0,
        "Target_Name" : "no",
        "Stress" : 3008,
        "BetweennessCentrality" : 7.9209E-4,
        "t_RNA_motif_Root" : "NO",
        "EdgeCount" : 4,
        "Causal_PLAMOM" : "no",
        "name" : "AT5G23220",
        "Causal_tRNA" : "no",
        "IsSingleNode" : false,
        "briefDescription" : "nicotinamidase 3",
        "NeighborhoodConnectivity" : 2.75
      },
      "position" : {
        "x" : -6663.630859375,
        "y" : 4052.203857421875
      },
      "selected" : false
    }, {
      "data" : {
        "id" : "1209",
        "symbol" : "AT5G54970",
        "Shoot_Descriptions" : "",
        "PLAMOM_Match_Shoot" : "",
        "Causal_TF" : "no",
        "Root_Descriptions" : "unknown protein; BEST Arabidopsis thaliana protein match is: unknown protein (TAIR:AT4G26960.1); Has 1807 Blast hits to 1807 proteins in 277 species: Archae - 0; Bacteria - 0; Metazoa - 736; Fungi - 347; Plants - 385; Viruses - 0; Other Eukaryotes - 339 (source: NCBI BLink).",
        "Eccentricity" : 10,
        "Outdegree" : 3,
        "Target_PLAMOM" : "",
        "PLAMOM_Match_Root" : "NO",
        "shared_name" : "AT5G54970",
        "Indegree" : 0,
        "Target_tRNA" : "",
        "t_RNA_motif_shoot" : "",
        "SUID" : 1209,
        "SelfLoops" : 0,
        "Target_TF" : "",
        "MolecularAttributes" : "protein_coding",
        "AverageShortestPathLength" : 6.04573805,
        "selected" : false,
        "Name2" : "AT5G54970",
        "ClosenessCentrality" : 0.16540578,
        "Casual_Name" : "unknown protein; BEST Arabidopsis thaliana protein match is: unknown protein (TAIR:AT4G26960.1); Has 1807 Blast hits to 1807 proteins in 277 species: Archae - 0; Bacteria - 0; Metazoa - 736; Fungi - 347; Plants - 385; Viruses - 0; Other Eukaryotes - 339 (source: NCBI BLink).",
        "PartnerOfMultiEdgedNodePairs" : 0,
        "ClusteringCoefficient" : 0.0,
        "Target_Name" : "",
        "Stress" : 0,
        "BetweennessCentrality" : 0.0,
        "t_RNA_motif_Root" : "NO",
        "EdgeCount" : 3,
        "Causal_PLAMOM" : "no",
        "name" : "AT5G54970",
        "Causal_tRNA" : "no",
        "IsSingleNode" : false,
        "briefDescription" : "hypothetical protein",
        "NeighborhoodConnectivity" : 2.3333333333333335
      },
      "position" : {
        "x" : -5954.76806640625,
        "y" : 823.5914306640625
      },
      "selected" : false
    }, {
      "data" : {
        "id" : "1208",
        "symbol" : "NAC13",
        "Shoot_Descriptions" : "ANAC013, ANAC13, NAC13, NAC domain protein 13",
        "PLAMOM_Match_Shoot" : "NO",
        "Causal_TF" : "",
        "Root_Descriptions" : "",
        "Eccentricity" : 0,
        "Outdegree" : 0,
        "Target_PLAMOM" : "no",
        "PLAMOM_Match_Root" : "",
        "shared_name" : "AT1G32870",
        "Indegree" : 1,
        "Target_tRNA" : "no",
        "t_RNA_motif_shoot" : "NO",
        "SUID" : 1208,
        "SelfLoops" : 0,
        "Target_TF" : "ANAC13",
        "MolecularAttributes" : "TXNFACTOR",
        "AverageShortestPathLength" : 0.0,
        "selected" : false,
        "Name2" : "NAC13",
        "ClosenessCentrality" : 0.0,
        "Casual_Name" : "",
        "PartnerOfMultiEdgedNodePairs" : 0,
        "ClusteringCoefficient" : 0.0,
        "Target_Name" : "\"ANAC013",
        "Stress" : 0,
        "BetweennessCentrality" : 0.0,
        "t_RNA_motif_Root" : "",
        "EdgeCount" : 1,
        "Causal_PLAMOM" : "",
        "name" : "AT1G32870",
        "Causal_tRNA" : "",
        "IsSingleNode" : false,
        "briefDescription" : "NAC domain protein 13",
        "NeighborhoodConnectivity" : 3.0
      },
      "position" : {
        "x" : 4132.303721365332,
        "y" : 862.50244140625
      },
      "selected" : false
    }, {
      "data" : {
        "id" : "1207",
        "symbol" : "AT1G32920",
        "degree_layout" : 4,
        "Shoot_Descriptions" : "unknown protein; FUNCTIONS IN: molecular_function unknown; INVOLVED IN: response to wounding; LOCATED IN: endomembrane system; EXPRESSED IN: 23 plant structures; EXPRESSED DURING: 13 growth stages; BEST Arabidopsis thaliana protein match is: unknown protein (TAIR:AT1G32928.1); Has 42 Blast hits to 42 proteins in 8 species: Archae - 0; Bacteria - 0; Metazoa - 0; Fungi - 0; Plants - 42; Viruses - 0; Other Eukaryotes - 0 (source: NCBI BLink).",
        "PLAMOM_Match_Shoot" : "YES",
        "Causal_TF" : "no",
        "Root_Descriptions" : "",
        "Eccentricity" : 9,
        "Outdegree" : 3,
        "Target_PLAMOM" : "AT1G32920",
        "PLAMOM_Match_Root" : "",
        "shared_name" : "AT1G32920",
        "Indegree" : 4,
        "Target_tRNA" : "no",
        "t_RNA_motif_shoot" : "NO",
        "SUID" : 1207,
        "SelfLoops" : 0,
        "Target_TF" : "no",
        "MolecularAttributes" : "protein_coding",
        "AverageShortestPathLength" : 4.33054393,
        "selected" : false,
        "Name2" : "AT1G32920",
        "ClosenessCentrality" : 0.23091787,
        "Casual_Name" : "unknown protein; FUNCTIONS IN: molecular_function unknown; INVOLVED IN: response to wounding; LOCATED IN: endomembrane system; EXPRESSED IN: 23 plant structures; EXPRESSED DURING: 13 growth stages; BEST Arabidopsis thaliana protein match is: unknown protein (TAIR:AT1G32928.1); Has 42 Blast hits to 42 proteins in 8 species: Archae - 0; Bacteria - 0; Metazoa - 0; Fungi - 0; Plants - 42; Viruses - 0; Other Eukaryotes - 0 (source: NCBI BLink).",
        "PartnerOfMultiEdgedNodePairs" : 0,
        "ClusteringCoefficient" : 0.02380952,
        "Target_Name" : "unknown protein; FUNCTIONS IN: molecular_function unknown; INVOLVED IN: response to wounding; LOCATED IN: endomembrane system; EXPRESSED IN: 23 plant structures; EXPRESSED DURING: 13 growth stages; BEST Arabidopsis thaliana protein match is: unknown protein (TAIR:AT1G32928.1); Has 42 Blast hits to 42 proteins in 8 species: Archae - 0; Bacteria - 0; Metazoa - 0; Fungi - 0; Plants - 42; Viruses - 0; Other Eukaryotes - 0 (source: NCBI BLink).",
        "Stress" : 8993,
        "BetweennessCentrality" : 0.00132674,
        "t_RNA_motif_Root" : "",
        "EdgeCount" : 7,
        "Causal_PLAMOM" : "AT1G32920",
        "name" : "AT1G32920",
        "Causal_tRNA" : "no",
        "IsSingleNode" : false,
        "briefDescription" : "hypothetical protein",
        "NeighborhoodConnectivity" : 33.714285714285715
      },
      "position" : {
        "x" : 3722.813183236296,
        "y" : 3456.357666015625
      },
      "selected" : false
    }, {
      "data" : {
        "id" : "1206",
        "symbol" : "AT2G41410",
        "Shoot_Descriptions" : "",
        "PLAMOM_Match_Shoot" : "",
        "Causal_TF" : "no",
        "Root_Descriptions" : "Calcium-binding EF-hand family protein",
        "Eccentricity" : 8,
        "Outdegree" : 17,
        "Target_PLAMOM" : "no",
        "PLAMOM_Match_Root" : "NO",
        "shared_name" : "AT2G41410",
        "Indegree" : 4,
        "Target_tRNA" : "no",
        "t_RNA_motif_shoot" : "",
        "SUID" : 1206,
        "SelfLoops" : 0,
        "Target_TF" : "no",
        "MolecularAttributes" : "protein_coding",
        "AverageShortestPathLength" : 4.01035197,
        "selected" : false,
        "Name2" : "AT2G41410",
        "ClosenessCentrality" : 0.24935467,
        "Casual_Name" : "Calcium-binding EF-hand family protein",
        "PartnerOfMultiEdgedNodePairs" : 0,
        "ClusteringCoefficient" : 0.00238095,
        "Target_Name" : "Calcium-binding EF-hand family protein",
        "Stress" : 3972,
        "BetweennessCentrality" : 7.6675E-4,
        "t_RNA_motif_Root" : "NO",
        "EdgeCount" : 21,
        "Causal_PLAMOM" : "no",
        "name" : "AT2G41410",
        "Causal_tRNA" : "no",
        "IsSingleNode" : false,
        "briefDescription" : "Calcium-binding EF-hand family protein",
        "NeighborhoodConnectivity" : 7.476190476190476
      },
      "position" : {
        "x" : -6419.693359375,
        "y" : 3329.7109375
      },
      "selected" : false
    }, {
      "data" : {
        "id" : "1205",
        "symbol" : "AXS2",
        "Shoot_Descriptions" : "",
        "PLAMOM_Match_Shoot" : "",
        "Causal_TF" : "UDP-D-apiose/UDP-D-xylose synthase 2\"",
        "Root_Descriptions" : "AXS2, UDP-D-apiose/UDP-D-xylose synthase 2",
        "Eccentricity" : 9,
        "Outdegree" : 13,
        "Target_PLAMOM" : "",
        "PLAMOM_Match_Root" : "YES",
        "shared_name" : "AT1G08200",
        "Indegree" : 0,
        "Target_tRNA" : "",
        "t_RNA_motif_shoot" : "",
        "SUID" : 1205,
        "SelfLoops" : 0,
        "Target_TF" : "",
        "MolecularAttributes" : "METABOLIC",
        "AverageShortestPathLength" : 5.05327869,
        "selected" : false,
        "Name2" : "AXS2",
        "ClosenessCentrality" : 0.19789132,
        "Casual_Name" : "\"AXS2",
        "PartnerOfMultiEdgedNodePairs" : 0,
        "ClusteringCoefficient" : 0.0,
        "Target_Name" : "",
        "Stress" : 0,
        "BetweennessCentrality" : 0.0,
        "t_RNA_motif_Root" : "NO",
        "EdgeCount" : 13,
        "Causal_PLAMOM" : "AT1G08200",
        "name" : "AT1G08200",
        "Causal_tRNA" : "no",
        "IsSingleNode" : false,
        "briefDescription" : "UDP-D-apiose/UDP-D-xylose synthase 2",
        "NeighborhoodConnectivity" : 7.3076923076923075
      },
      "position" : {
        "x" : -6307.7373046875,
        "y" : -167.7948760986328
      },
      "selected" : false
    }, {
      "data" : {
        "id" : "1204",
        "symbol" : "AT1G33120",
        "Shoot_Descriptions" : "Ribosomal protein L6 family",
        "PLAMOM_Match_Shoot" : "YES",
        "Causal_TF" : "",
        "Root_Descriptions" : "",
        "Eccentricity" : 0,
        "Outdegree" : 0,
        "Target_PLAMOM" : "AT1G33120",
        "PLAMOM_Match_Root" : "",
        "shared_name" : "AT1G33120",
        "Indegree" : 3,
        "Target_tRNA" : "no",
        "t_RNA_motif_shoot" : "NO",
        "SUID" : 1204,
        "SelfLoops" : 0,
        "Target_TF" : "no",
        "MolecularAttributes" : "protein_coding",
        "AverageShortestPathLength" : 0.0,
        "selected" : false,
        "Name2" : "AT1G33120",
        "ClosenessCentrality" : 0.0,
        "Casual_Name" : "",
        "PartnerOfMultiEdgedNodePairs" : 0,
        "ClusteringCoefficient" : 0.0,
        "Target_Name" : "Ribosomal protein L6 family",
        "Stress" : 0,
        "BetweennessCentrality" : 0.0,
        "t_RNA_motif_Root" : "",
        "EdgeCount" : 3,
        "Causal_PLAMOM" : "",
        "name" : "AT1G33120",
        "Causal_tRNA" : "",
        "IsSingleNode" : false,
        "briefDescription" : "Ribosomal protein L6 family",
        "NeighborhoodConnectivity" : 9.0
      },
      "position" : {
        "x" : 3926.234814095671,
        "y" : 6.652795791625977
      },
      "selected" : false
    }, {
      "data" : {
        "id" : "1203",
        "symbol" : "EMB1075",
        "Shoot_Descriptions" : "",
        "PLAMOM_Match_Shoot" : "",
        "Causal_TF" : "Pyridoxal phosphate (PLP)-dependent transferases superfamily protein\"",
        "Root_Descriptions" : "emb1075, Pyridoxal phosphate (PLP)-dependent transferases superfamily protein",
        "Eccentricity" : 11,
        "Outdegree" : 3,
        "Target_PLAMOM" : "",
        "PLAMOM_Match_Root" : "YES",
        "shared_name" : "AT1G43710",
        "Indegree" : 0,
        "Target_tRNA" : "",
        "t_RNA_motif_shoot" : "",
        "SUID" : 1203,
        "SelfLoops" : 0,
        "Target_TF" : "",
        "MolecularAttributes" : "METABOLIC",
        "AverageShortestPathLength" : 6.06263048,
        "selected" : false,
        "Name2" : "EMB1075",
        "ClosenessCentrality" : 0.1649449,
        "Casual_Name" : "\"emb1075",
        "PartnerOfMultiEdgedNodePairs" : 0,
        "ClusteringCoefficient" : 0.0,
        "Target_Name" : "",
        "Stress" : 0,
        "BetweennessCentrality" : 0.0,
        "t_RNA_motif_Root" : "NO",
        "EdgeCount" : 3,
        "Causal_PLAMOM" : "AT1G43710",
        "name" : "AT1G43710",
        "Causal_tRNA" : "no",
        "IsSingleNode" : false,
        "briefDescription" : "Pyridoxal phosphate (PLP)-dependent transferases superfamily protein",
        "NeighborhoodConnectivity" : 24.333333333333332
      },
      "position" : {
        "x" : -6342.63916015625,
        "y" : -276.4936218261719
      },
      "selected" : false
    }, {
      "data" : {
        "id" : "1202",
        "symbol" : "AT3G59040",
        "Shoot_Descriptions" : "",
        "PLAMOM_Match_Shoot" : "",
        "Causal_TF" : "no",
        "Root_Descriptions" : "Tetratricopeptide repeat (TPR)-like superfamily protein",
        "Eccentricity" : 9,
        "Outdegree" : 9,
        "Target_PLAMOM" : "signal responsive 1\"",
        "PLAMOM_Match_Root" : "NO",
        "shared_name" : "AT3G59040",
        "Indegree" : 2,
        "Target_tRNA" : "AT2G22300",
        "t_RNA_motif_shoot" : "",
        "SUID" : 1202,
        "SelfLoops" : 0,
        "Target_TF" : "no",
        "MolecularAttributes" : "protein_coding",
        "AverageShortestPathLength" : 4.71338912,
        "selected" : false,
        "Name2" : "AT3G59040",
        "ClosenessCentrality" : 0.21216156,
        "Casual_Name" : "Tetratricopeptide repeat (TPR)-like superfamily protein",
        "PartnerOfMultiEdgedNodePairs" : 0,
        "ClusteringCoefficient" : 0.0,
        "Target_Name" : "no",
        "Stress" : 70313,
        "BetweennessCentrality" : 0.01264619,
        "t_RNA_motif_Root" : "NO",
        "EdgeCount" : 11,
        "Causal_PLAMOM" : "no",
        "name" : "AT3G59040",
        "Causal_tRNA" : "no",
        "IsSingleNode" : false,
        "briefDescription" : "Tetratricopeptide repeat (TPR)-like superfamily protein",
        "NeighborhoodConnectivity" : 19.818181818181817
      },
      "position" : {
        "x" : -6344.96484375,
        "y" : 2934.997802734375
      },
      "selected" : false
    }, {
      "data" : {
        "id" : "1201",
        "symbol" : "bHLH38",
        "Shoot_Descriptions" : "",
        "PLAMOM_Match_Shoot" : "",
        "Causal_TF" : "ORG2",
        "Root_Descriptions" : "BHLH038, ORG2, basic helix-loop-helix (bHLH) DNA-binding superfamily protein",
        "Eccentricity" : 12,
        "Outdegree" : 3,
        "Target_PLAMOM" : "",
        "PLAMOM_Match_Root" : "NO",
        "shared_name" : "AT3G56970",
        "Indegree" : 0,
        "Target_tRNA" : "",
        "t_RNA_motif_shoot" : "",
        "SUID" : 1201,
        "SelfLoops" : 0,
        "Target_TF" : "",
        "MolecularAttributes" : "TXNFACTOR",
        "AverageShortestPathLength" : 7.23382046,
        "selected" : false,
        "Name2" : "bHLH38",
        "ClosenessCentrality" : 0.13823954,
        "Casual_Name" : "\"BHLH038",
        "PartnerOfMultiEdgedNodePairs" : 0,
        "ClusteringCoefficient" : 0.0,
        "Target_Name" : "",
        "Stress" : 0,
        "BetweennessCentrality" : 0.0,
        "t_RNA_motif_Root" : "NO",
        "EdgeCount" : 3,
        "Causal_PLAMOM" : "no",
        "name" : "AT3G56970",
        "Causal_tRNA" : "no",
        "IsSingleNode" : false,
        "briefDescription" : "basic helix-loop-helix (bHLH) DNA-binding superfamily protein",
        "NeighborhoodConnectivity" : 9.333333333333334
      },
      "position" : {
        "x" : -6502.79833984375,
        "y" : 496.80694580078125
      },
      "selected" : false
    }, {
      "data" : {
        "id" : "1200",
        "symbol" : "FAS4",
        "Shoot_Descriptions" : "ATFAS4, FAS4, RNA helicase family protein",
        "PLAMOM_Match_Shoot" : "NO",
        "Causal_TF" : "FAS4",
        "Root_Descriptions" : "",
        "Eccentricity" : 1,
        "Outdegree" : 1,
        "Target_PLAMOM" : "WR3",
        "PLAMOM_Match_Root" : "",
        "shared_name" : "AT1G33390",
        "Indegree" : 8,
        "Target_tRNA" : "nitrate transmembrane transporters\"",
        "t_RNA_motif_shoot" : "NO",
        "SUID" : 1200,
        "SelfLoops" : 0,
        "Target_TF" : "no",
        "MolecularAttributes" : "protein_coding",
        "AverageShortestPathLength" : 1.0,
        "selected" : false,
        "Name2" : "FAS4",
        "ClosenessCentrality" : 1.0,
        "Casual_Name" : "\"ATFAS4",
        "PartnerOfMultiEdgedNodePairs" : 0,
        "ClusteringCoefficient" : 0.0,
        "Target_Name" : "no",
        "Stress" : 1328,
        "BetweennessCentrality" : 4.4553E-4,
        "t_RNA_motif_Root" : "",
        "EdgeCount" : 9,
        "Causal_PLAMOM" : "no",
        "name" : "AT1G33390",
        "Causal_tRNA" : "no",
        "IsSingleNode" : false,
        "briefDescription" : "RNA helicase family protein",
        "NeighborhoodConnectivity" : 10.11111111111111
      },
      "position" : {
        "x" : 2988.208691048796,
        "y" : 3389.427978515625
      },
      "selected" : false
    }, {
      "data" : {
        "id" : "1199",
        "symbol" : "AT1G47400",
        "Shoot_Descriptions" : "unknown protein; BEST Arabidopsis thaliana protein match is: unknown protein (TAIR:AT1G47395.1); Has 11 Blast hits to 11 proteins in 2 species: Archae - 0; Bacteria - 0; Metazoa - 0; Fungi - 0; Plants - 11; Viruses - 0; Other Eukaryotes - 0 (source: NCBI BLink).",
        "PLAMOM_Match_Shoot" : "NO",
        "Causal_TF" : "no",
        "Root_Descriptions" : "unknown protein; BEST Arabidopsis thaliana protein match is: unknown protein (TAIR:AT1G47395.1); Has 11 Blast hits to 11 proteins in 2 species: Archae - 0; Bacteria - 0; Metazoa - 0; Fungi - 0; Plants - 11; Viruses - 0; Other Eukaryotes - 0 (source: NCBI BLink).",
        "Eccentricity" : 2,
        "Outdegree" : 1,
        "Target_PLAMOM" : "no",
        "PLAMOM_Match_Root" : "NO",
        "shared_name" : "AT1G47400",
        "Indegree" : 2,
        "Target_tRNA" : "no",
        "t_RNA_motif_shoot" : "NO",
        "SUID" : 1199,
        "SelfLoops" : 0,
        "Target_TF" : "unknown protein; BEST Arabidopsis thaliana protein match is: unknown protein (TAIR:AT1G47395.1); Has 11 Blast hits to 11 proteins in 2 species: Archae - 0; Bacteria - 0; Metazoa - 0; Fungi - 0; Plants - 11; Viruses - 0; Other Eukaryotes - 0 (source: NCBI BLink).",
        "MolecularAttributes" : "protein_coding",
        "AverageShortestPathLength" : 1.5,
        "selected" : false,
        "Name2" : "AT1G47400",
        "ClosenessCentrality" : 0.66666667,
        "Casual_Name" : "unknown protein; BEST Arabidopsis thaliana protein match is: unknown protein (TAIR:AT1G47395.1); Has 11 Blast hits to 11 proteins in 2 species: Archae - 0; Bacteria - 0; Metazoa - 0; Fungi - 0; Plants - 11; Viruses - 0; Other Eukaryotes - 0 (source: NCBI BLink).",
        "PartnerOfMultiEdgedNodePairs" : 0,
        "ClusteringCoefficient" : 0.0,
        "Target_Name" : "no",
        "Stress" : 130,
        "BetweennessCentrality" : 2.272E-5,
        "t_RNA_motif_Root" : "NO",
        "EdgeCount" : 3,
        "Causal_PLAMOM" : "no",
        "name" : "AT1G47400",
        "Causal_tRNA" : "no",
        "IsSingleNode" : false,
        "briefDescription" : "hypothetical protein",
        "NeighborhoodConnectivity" : 8.666666666666666
      },
      "position" : {
        "x" : -803.0593143222977,
        "y" : 739.0307006835938
      },
      "selected" : false
    }, {
      "data" : {
        "id" : "1198",
        "symbol" : "PNC1",
        "Shoot_Descriptions" : "",
        "PLAMOM_Match_Shoot" : "",
        "Causal_TF" : "peroxisomal adenine nucleotide carrier 1\"",
        "Root_Descriptions" : "PNC1, peroxisomal adenine nucleotide carrier 1",
        "Eccentricity" : 2,
        "Outdegree" : 2,
        "Target_PLAMOM" : "no",
        "PLAMOM_Match_Root" : "NO",
        "shared_name" : "AT3G05290",
        "Indegree" : 1,
        "Target_tRNA" : "AT3G05290",
        "t_RNA_motif_shoot" : "",
        "SUID" : 1198,
        "SelfLoops" : 0,
        "Target_TF" : "peroxisomal adenine nucleotide carrier 1\"",
        "MolecularAttributes" : "protein_coding",
        "AverageShortestPathLength" : 1.33333333,
        "selected" : false,
        "Name2" : "PNC1",
        "ClosenessCentrality" : 0.75,
        "Casual_Name" : "\"PNC1",
        "PartnerOfMultiEdgedNodePairs" : 0,
        "ClusteringCoefficient" : 0.0,
        "Target_Name" : "\"PNC1",
        "Stress" : 2590,
        "BetweennessCentrality" : 8.7554E-4,
        "t_RNA_motif_Root" : "YES",
        "EdgeCount" : 3,
        "Causal_PLAMOM" : "no",
        "name" : "AT3G05290",
        "Causal_tRNA" : "AT3G05290",
        "IsSingleNode" : false,
        "briefDescription" : "peroxisomal adenine nucleotide carrier 1",
        "NeighborhoodConnectivity" : 5.666666666666667
      },
      "position" : {
        "x" : -6659.6884765625,
        "y" : 3240.55126953125
      },
      "selected" : false
    }, {
      "data" : {
        "id" : "1197",
        "symbol" : "WR3",
        "Shoot_Descriptions" : "",
        "PLAMOM_Match_Shoot" : "",
        "Causal_TF" : "NRT3.1",
        "Root_Descriptions" : "ATNRT3.1, NRT3.1, WR3, nitrate transmembrane transporters",
        "Eccentricity" : 12,
        "Outdegree" : 3,
        "Target_PLAMOM" : "no",
        "PLAMOM_Match_Root" : "NO",
        "shared_name" : "AT5G50200",
        "Indegree" : 23,
        "Target_tRNA" : "no",
        "t_RNA_motif_shoot" : "",
        "SUID" : 1197,
        "SelfLoops" : 0,
        "Target_TF" : "\"ATNRT3.1",
        "MolecularAttributes" : "protein_coding",
        "AverageShortestPathLength" : 6.9874477,
        "selected" : false,
        "Name2" : "WR3",
        "ClosenessCentrality" : 0.14311377,
        "Casual_Name" : "\"ATNRT3.1",
        "PartnerOfMultiEdgedNodePairs" : 0,
        "ClusteringCoefficient" : 0.0,
        "Target_Name" : "no",
        "Stress" : 32501,
        "BetweennessCentrality" : 0.00732203,
        "t_RNA_motif_Root" : "NO",
        "EdgeCount" : 26,
        "Causal_PLAMOM" : "no",
        "name" : "AT5G50200",
        "Causal_tRNA" : "no",
        "IsSingleNode" : false,
        "briefDescription" : "nitrate transmembrane transporter",
        "NeighborhoodConnectivity" : 4.1923076923076925
      },
      "position" : {
        "x" : -7287.4853515625,
        "y" : 3252.66552734375
      },
      "selected" : false
    }, {
      "data" : {
        "id" : "1196",
        "symbol" : "AT2G17150",
        "degree_layout" : 9,
        "Shoot_Descriptions" : "Plant regulator RWP-RK family protein",
        "PLAMOM_Match_Shoot" : "NO",
        "Causal_TF" : "AT2G17150",
        "Root_Descriptions" : "Plant regulator RWP-RK family protein",
        "Eccentricity" : 8,
        "Outdegree" : 8,
        "Target_PLAMOM" : "no",
        "PLAMOM_Match_Root" : "NO",
        "shared_name" : "AT2G17150",
        "Indegree" : 1,
        "Target_tRNA" : "AT2G17150",
        "t_RNA_motif_shoot" : "YES",
        "SUID" : 1196,
        "SelfLoops" : 0,
        "Target_TF" : "AT2G17150",
        "MolecularAttributes" : "TXNFACTOR",
        "AverageShortestPathLength" : 4.67782427,
        "selected" : false,
        "Name2" : "AT2G17150",
        "ClosenessCentrality" : 0.2137746,
        "Casual_Name" : "Plant regulator RWP-RK family protein",
        "PartnerOfMultiEdgedNodePairs" : 0,
        "ClusteringCoefficient" : 0.0,
        "Target_Name" : "Plant regulator RWP-RK family protein",
        "Stress" : 29137,
        "BetweennessCentrality" : 0.00662056,
        "t_RNA_motif_Root" : "YES",
        "EdgeCount" : 9,
        "Causal_PLAMOM" : "no",
        "name" : "AT2G17150",
        "Causal_tRNA" : "AT2G17150",
        "IsSingleNode" : false,
        "briefDescription" : "Plant regulator RWP-RK family protein",
        "NeighborhoodConnectivity" : 20.88888888888889
      },
      "position" : {
        "x" : -893.1072879551102,
        "y" : 3251.33447265625
      },
      "selected" : false
    }, {
      "data" : {
        "id" : "1195",
        "symbol" : "AT1G33780",
        "degree_layout" : 1,
        "Shoot_Descriptions" : "Protein of unknown function (DUF179)",
        "PLAMOM_Match_Shoot" : "NO",
        "Causal_TF" : "",
        "Root_Descriptions" : "",
        "Eccentricity" : 0,
        "Outdegree" : 0,
        "Target_PLAMOM" : "no",
        "PLAMOM_Match_Root" : "",
        "shared_name" : "AT1G33780",
        "Indegree" : 3,
        "Target_tRNA" : "no",
        "t_RNA_motif_shoot" : "NO",
        "SUID" : 1195,
        "SelfLoops" : 0,
        "Target_TF" : "no",
        "MolecularAttributes" : "protein_coding",
        "AverageShortestPathLength" : 0.0,
        "selected" : false,
        "Name2" : "AT1G33780",
        "ClosenessCentrality" : 0.0,
        "Casual_Name" : "",
        "PartnerOfMultiEdgedNodePairs" : 0,
        "ClusteringCoefficient" : 0.0,
        "Target_Name" : "Protein of unknown function (DUF179)",
        "Stress" : 0,
        "BetweennessCentrality" : 0.0,
        "t_RNA_motif_Root" : "",
        "EdgeCount" : 3,
        "Causal_PLAMOM" : "",
        "name" : "AT1G33780",
        "Causal_tRNA" : "",
        "IsSingleNode" : false,
        "briefDescription" : "electron transporter",
        "NeighborhoodConnectivity" : 5.333333333333333
      },
      "position" : {
        "x" : 4522.120068001921,
        "y" : 224.7782440185547
      },
      "selected" : false
    }, {
      "data" : {
        "id" : "1194",
        "symbol" : "AO",
        "Shoot_Descriptions" : "",
        "PLAMOM_Match_Shoot" : "",
        "Causal_TF" : "L-aspartate oxidase\"",
        "Root_Descriptions" : "AO, L-aspartate oxidase",
        "Eccentricity" : 1,
        "Outdegree" : 1,
        "Target_PLAMOM" : "",
        "PLAMOM_Match_Root" : "NO",
        "shared_name" : "AT5G14760",
        "Indegree" : 0,
        "Target_tRNA" : "",
        "t_RNA_motif_shoot" : "",
        "SUID" : 1194,
        "SelfLoops" : 0,
        "Target_TF" : "",
        "MolecularAttributes" : "METABOLIC",
        "AverageShortestPathLength" : 1.0,
        "selected" : false,
        "Name2" : "AO",
        "ClosenessCentrality" : 1.0,
        "Casual_Name" : "\"AO",
        "PartnerOfMultiEdgedNodePairs" : 0,
        "ClusteringCoefficient" : 0.0,
        "Target_Name" : "",
        "Stress" : 0,
        "BetweennessCentrality" : 0.0,
        "t_RNA_motif_Root" : "NO",
        "EdgeCount" : 1,
        "Causal_PLAMOM" : "no",
        "name" : "AT5G14760",
        "Causal_tRNA" : "no",
        "IsSingleNode" : false,
        "briefDescription" : "L-aspartate oxidase",
        "NeighborhoodConnectivity" : 3.0
      },
      "position" : {
        "x" : -5214.544921875,
        "y" : 356.9801940917969
      },
      "selected" : false
    }, {
      "data" : {
        "id" : "1193",
        "symbol" : "AT3G54530",
        "Shoot_Descriptions" : "",
        "PLAMOM_Match_Shoot" : "",
        "Causal_TF" : "no",
        "Root_Descriptions" : "unknown protein; BEST Arabidopsis thaliana protein match is: unknown protein (TAIR:AT3G54520.1); Has 159 Blast hits to 159 proteins in 50 species: Archae - 2; Bacteria - 26; Metazoa - 55; Fungi - 5; Plants - 20; Viruses - 2; Other Eukaryotes - 49 (source: NCBI BLink).",
        "Eccentricity" : 9,
        "Outdegree" : 6,
        "Target_PLAMOM" : "",
        "PLAMOM_Match_Root" : "NO",
        "shared_name" : "AT3G54530",
        "Indegree" : 0,
        "Target_tRNA" : "",
        "t_RNA_motif_shoot" : "",
        "SUID" : 1193,
        "SelfLoops" : 0,
        "Target_TF" : "",
        "MolecularAttributes" : "protein_coding",
        "AverageShortestPathLength" : 5.05010438,
        "selected" : false,
        "Name2" : "AT3G54530",
        "ClosenessCentrality" : 0.19801571,
        "Casual_Name" : "unknown protein; BEST Arabidopsis thaliana protein match is: unknown protein (TAIR:AT3G54520.1); Has 159 Blast hits to 159 proteins in 50 species: Archae - 2; Bacteria - 26; Metazoa - 55; Fungi - 5; Plants - 20; Viruses - 2; Other Eukaryotes - 49 (source: NCBI BLink).",
        "PartnerOfMultiEdgedNodePairs" : 0,
        "ClusteringCoefficient" : 0.0,
        "Target_Name" : "",
        "Stress" : 0,
        "BetweennessCentrality" : 0.0,
        "t_RNA_motif_Root" : "NO",
        "EdgeCount" : 6,
        "Causal_PLAMOM" : "no",
        "name" : "AT3G54530",
        "Causal_tRNA" : "no",
        "IsSingleNode" : false,
        "briefDescription" : "hypothetical protein",
        "NeighborhoodConnectivity" : 15.166666666666666
      },
      "position" : {
        "x" : -5783.326171875,
        "y" : 5.505334854125977
      },
      "selected" : false
    }, {
      "data" : {
        "id" : "1192",
        "symbol" : "AT1G35560",
        "degree_layout" : 3,
        "Shoot_Descriptions" : "TCP family transcription factor",
        "PLAMOM_Match_Shoot" : "NO",
        "Causal_TF" : "AT1G35560",
        "Root_Descriptions" : "",
        "Eccentricity" : 10,
        "Outdegree" : 2,
        "Target_PLAMOM" : "no",
        "PLAMOM_Match_Root" : "",
        "shared_name" : "AT1G35560",
        "Indegree" : 2,
        "Target_tRNA" : "no",
        "t_RNA_motif_shoot" : "NO",
        "SUID" : 1192,
        "SelfLoops" : 0,
        "Target_TF" : "AT1G35560",
        "MolecularAttributes" : "TXNFACTOR",
        "AverageShortestPathLength" : 5.72803347,
        "selected" : false,
        "Name2" : "AT1G35560",
        "ClosenessCentrality" : 0.17457999,
        "Casual_Name" : "TCP family transcription factor",
        "PartnerOfMultiEdgedNodePairs" : 0,
        "ClusteringCoefficient" : 0.08333333,
        "Target_Name" : "TCP family transcription factor",
        "Stress" : 574,
        "BetweennessCentrality" : 6.051E-5,
        "t_RNA_motif_Root" : "",
        "EdgeCount" : 4,
        "Causal_PLAMOM" : "no",
        "name" : "AT1G35560",
        "Causal_tRNA" : "no",
        "IsSingleNode" : false,
        "briefDescription" : "TCP family transcription factor",
        "NeighborhoodConnectivity" : 42.75
      },
      "position" : {
        "x" : 3812.919872689421,
        "y" : 3164.9140625
      },
      "selected" : false
    }, {
      "data" : {
        "id" : "1191",
        "symbol" : "AT1G36730",
        "Shoot_Descriptions" : "Translation initiation factor IF2/IF5",
        "PLAMOM_Match_Shoot" : "NO",
        "Causal_TF" : "no",
        "Root_Descriptions" : "",
        "Eccentricity" : 9,
        "Outdegree" : 4,
        "Target_PLAMOM" : "no",
        "PLAMOM_Match_Root" : "",
        "shared_name" : "AT1G36730",
        "Indegree" : 3,
        "Target_tRNA" : "no",
        "t_RNA_motif_shoot" : "NO",
        "SUID" : 1191,
        "SelfLoops" : 0,
        "Target_TF" : "no",
        "MolecularAttributes" : "protein_coding",
        "AverageShortestPathLength" : 4.58577406,
        "selected" : false,
        "Name2" : "AT1G36730",
        "ClosenessCentrality" : 0.21806569,
        "Casual_Name" : "Translation initiation factor IF2/IF5",
        "PartnerOfMultiEdgedNodePairs" : 0,
        "ClusteringCoefficient" : 0.0,
        "Target_Name" : "Translation initiation factor IF2/IF5",
        "Stress" : 6762,
        "BetweennessCentrality" : 0.0010429,
        "t_RNA_motif_Root" : "",
        "EdgeCount" : 7,
        "Causal_PLAMOM" : "no",
        "name" : "AT1G36730",
        "Causal_tRNA" : "no",
        "IsSingleNode" : false,
        "briefDescription" : "Translation initiation factor IF2/IF5",
        "NeighborhoodConnectivity" : 22.0
      },
      "position" : {
        "x" : 3424.645946908171,
        "y" : 2952.873779296875
      },
      "selected" : false
    }, {
      "data" : {
        "id" : "1190",
        "symbol" : "AT1G44160",
        "degree_layout" : 5,
        "Shoot_Descriptions" : "",
        "PLAMOM_Match_Shoot" : "",
        "Causal_TF" : "no",
        "Root_Descriptions" : "HSP40/DnaJ peptide-binding protein",
        "Eccentricity" : 8,
        "Outdegree" : 11,
        "Target_PLAMOM" : "no",
        "PLAMOM_Match_Root" : "NO",
        "shared_name" : "AT1G44160",
        "Indegree" : 3,
        "Target_tRNA" : "no",
        "t_RNA_motif_shoot" : "",
        "SUID" : 1190,
        "SelfLoops" : 0,
        "Target_TF" : "no",
        "MolecularAttributes" : "protein_coding",
        "AverageShortestPathLength" : 4.39121339,
        "selected" : false,
        "Name2" : "AT1G44160",
        "ClosenessCentrality" : 0.22772749,
        "Casual_Name" : "HSP40/DnaJ peptide-binding protein",
        "PartnerOfMultiEdgedNodePairs" : 0,
        "ClusteringCoefficient" : 0.0,
        "Target_Name" : "HSP40/DnaJ peptide-binding protein",
        "Stress" : 79854,
        "BetweennessCentrality" : 0.01648291,
        "t_RNA_motif_Root" : "NO",
        "EdgeCount" : 14,
        "Causal_PLAMOM" : "no",
        "name" : "AT1G44160",
        "Causal_tRNA" : "no",
        "IsSingleNode" : false,
        "briefDescription" : "HSP40/DnaJ peptide-binding protein",
        "NeighborhoodConnectivity" : 10.571428571428571
      },
      "position" : {
        "x" : -6612.818359375,
        "y" : 2881.919189453125
      },
      "selected" : false
    }, {
      "data" : {
        "id" : "1189",
        "symbol" : "AT1G41880",
        "Shoot_Descriptions" : "Ribosomal protein L35Ae family protein",
        "PLAMOM_Match_Shoot" : "YES",
        "Causal_TF" : "",
        "Root_Descriptions" : "",
        "Eccentricity" : 0,
        "Outdegree" : 0,
        "Target_PLAMOM" : "AT1G41880",
        "PLAMOM_Match_Root" : "",
        "shared_name" : "AT1G41880",
        "Indegree" : 2,
        "Target_tRNA" : "AT1G41880",
        "t_RNA_motif_shoot" : "YES",
        "SUID" : 1189,
        "SelfLoops" : 0,
        "Target_TF" : "no",
        "MolecularAttributes" : "protein_coding",
        "AverageShortestPathLength" : 0.0,
        "selected" : false,
        "Name2" : "AT1G41880",
        "ClosenessCentrality" : 0.0,
        "Casual_Name" : "",
        "PartnerOfMultiEdgedNodePairs" : 0,
        "ClusteringCoefficient" : 0.0,
        "Target_Name" : "Ribosomal protein L35Ae family protein",
        "Stress" : 0,
        "BetweennessCentrality" : 0.0,
        "t_RNA_motif_Root" : "",
        "EdgeCount" : 2,
        "Causal_PLAMOM" : "",
        "name" : "AT1G41880",
        "Causal_tRNA" : "",
        "IsSingleNode" : false,
        "briefDescription" : "Ribosomal protein L35Ae family protein",
        "NeighborhoodConnectivity" : 6.5
      },
      "position" : {
        "x" : 3778.9694928497074,
        "y" : -397.6512451171875
      },
      "selected" : false
    }, {
      "data" : {
        "id" : "1188",
        "symbol" : "UMAMIT11",
        "Shoot_Descriptions" : "",
        "PLAMOM_Match_Shoot" : "",
        "Causal_TF" : "no",
        "Root_Descriptions" : "nodulin MtN21 /EamA-like transporter family protein",
        "Eccentricity" : 9,
        "Outdegree" : 8,
        "Target_PLAMOM" : "",
        "PLAMOM_Match_Root" : "NO",
        "shared_name" : "AT2G40900",
        "Indegree" : 0,
        "Target_tRNA" : "",
        "t_RNA_motif_shoot" : "",
        "SUID" : 1188,
        "SelfLoops" : 0,
        "Target_TF" : "",
        "MolecularAttributes" : "protein_coding",
        "AverageShortestPathLength" : 4.53541667,
        "selected" : false,
        "Name2" : "UMAMIT11",
        "ClosenessCentrality" : 0.22048691,
        "Casual_Name" : "nodulin MtN21 /EamA-like transporter family protein",
        "PartnerOfMultiEdgedNodePairs" : 0,
        "ClusteringCoefficient" : 0.0,
        "Target_Name" : "",
        "Stress" : 0,
        "BetweennessCentrality" : 0.0,
        "t_RNA_motif_Root" : "NO",
        "EdgeCount" : 8,
        "Causal_PLAMOM" : "no",
        "name" : "AT2G40900",
        "Causal_tRNA" : "no",
        "IsSingleNode" : false,
        "briefDescription" : "nodulin MtN21 /EamA-like transporter family protein",
        "NeighborhoodConnectivity" : 11.875
      },
      "position" : {
        "x" : -6176.47607421875,
        "y" : 153.86131286621094
      },
      "selected" : false
    }, {
      "data" : {
        "id" : "1187",
        "symbol" : "RP1",
        "degree_layout" : 2,
        "Shoot_Descriptions" : "ARP1, emb2207, RP1, RPL3A, ribosomal protein 1",
        "PLAMOM_Match_Shoot" : "YES",
        "Causal_TF" : "emb2207",
        "Root_Descriptions" : "",
        "Eccentricity" : 10,
        "Outdegree" : 1,
        "Target_PLAMOM" : "no",
        "PLAMOM_Match_Root" : "",
        "shared_name" : "AT1G43170",
        "Indegree" : 12,
        "Target_tRNA" : "AT1G43170",
        "t_RNA_motif_shoot" : "NO",
        "SUID" : 1187,
        "SelfLoops" : 0,
        "Target_TF" : "\"ARP1",
        "MolecularAttributes" : "protein_coding",
        "AverageShortestPathLength" : 5.77824268,
        "selected" : false,
        "Name2" : "RP1",
        "ClosenessCentrality" : 0.173063,
        "Casual_Name" : "\"ARP1",
        "PartnerOfMultiEdgedNodePairs" : 0,
        "ClusteringCoefficient" : 0.0,
        "Target_Name" : "no",
        "Stress" : 47901,
        "BetweennessCentrality" : 0.01086673,
        "t_RNA_motif_Root" : "",
        "EdgeCount" : 13,
        "Causal_PLAMOM" : "AT1G43170",
        "name" : "AT1G43170",
        "Causal_tRNA" : "no",
        "IsSingleNode" : false,
        "briefDescription" : "ribosomal protein 1",
        "NeighborhoodConnectivity" : 18.076923076923077
      },
      "position" : {
        "x" : 3497.652782845671,
        "y" : 3120.9091796875
      },
      "selected" : false
    }, {
      "data" : {
        "id" : "1186",
        "symbol" : "AT3G52950",
        "Shoot_Descriptions" : "",
        "PLAMOM_Match_Shoot" : "",
        "Causal_TF" : "no",
        "Root_Descriptions" : "CBS / octicosapeptide/Phox/Bemp1 (PB1) domains-containing protein",
        "Eccentricity" : 10,
        "Outdegree" : 3,
        "Target_PLAMOM" : "",
        "PLAMOM_Match_Root" : "NO",
        "shared_name" : "AT3G52950",
        "Indegree" : 0,
        "Target_tRNA" : "",
        "t_RNA_motif_shoot" : "",
        "SUID" : 1186,
        "SelfLoops" : 0,
        "Target_TF" : "",
        "MolecularAttributes" : "protein_coding",
        "AverageShortestPathLength" : 6.36382536,
        "selected" : false,
        "Name2" : "AT3G52950",
        "ClosenessCentrality" : 0.15713819,
        "Casual_Name" : "CBS / octicosapeptide/Phox/Bemp1 (PB1) domains-containing protein",
        "PartnerOfMultiEdgedNodePairs" : 0,
        "ClusteringCoefficient" : 0.0,
        "Target_Name" : "",
        "Stress" : 0,
        "BetweennessCentrality" : 0.0,
        "t_RNA_motif_Root" : "NO",
        "EdgeCount" : 3,
        "Causal_PLAMOM" : "no",
        "name" : "AT3G52950",
        "Causal_tRNA" : "no",
        "IsSingleNode" : false,
        "briefDescription" : "CBS / octicosapeptide/Phox/Bemp1 (PB1) domains-containing protein",
        "NeighborhoodConnectivity" : 4.0
      },
      "position" : {
        "x" : -6022.73681640625,
        "y" : 989.9899291992188
      },
      "selected" : false
    }, {
      "data" : {
        "id" : "1185",
        "symbol" : "AT1G44770",
        "Shoot_Descriptions" : "unknown protein; BEST Arabidopsis thaliana protein match is: unknown protein (TAIR:AT5G49710.3); Has 81 Blast hits to 81 proteins in 17 species: Archae - 0; Bacteria - 0; Metazoa - 0; Fungi - 0; Plants - 81; Viruses - 0; Other Eukaryotes - 0 (source: NCBI BLink).",
        "PLAMOM_Match_Shoot" : "NO",
        "Causal_TF" : "",
        "Root_Descriptions" : "",
        "Eccentricity" : 0,
        "Outdegree" : 0,
        "Target_PLAMOM" : "no",
        "PLAMOM_Match_Root" : "",
        "shared_name" : "AT1G44770",
        "Indegree" : 1,
        "Target_tRNA" : "no",
        "t_RNA_motif_shoot" : "NO",
        "SUID" : 1185,
        "SelfLoops" : 0,
        "Target_TF" : "no",
        "MolecularAttributes" : "protein_coding",
        "AverageShortestPathLength" : 0.0,
        "selected" : false,
        "Name2" : "AT1G44770",
        "ClosenessCentrality" : 0.0,
        "Casual_Name" : "",
        "PartnerOfMultiEdgedNodePairs" : 0,
        "ClusteringCoefficient" : 0.0,
        "Target_Name" : "unknown protein; BEST Arabidopsis thaliana protein match is: unknown protein (TAIR:AT5G49710.3); Has 81 Blast hits to 81 proteins in 17 species: Archae - 0; Bacteria - 0; Metazoa - 0; Fungi - 0; Plants - 81; Viruses - 0; Other Eukaryotes - 0 (source: NCBI BLink).",
        "Stress" : 0,
        "BetweennessCentrality" : 0.0,
        "t_RNA_motif_Root" : "",
        "EdgeCount" : 1,
        "Causal_PLAMOM" : "",
        "name" : "AT1G44770",
        "Causal_tRNA" : "",
        "IsSingleNode" : false,
        "briefDescription" : "elongation factor",
        "NeighborhoodConnectivity" : 3.0
      },
      "position" : {
        "x" : 3908.6140240997074,
        "y" : 1100.7724609375
      },
      "selected" : false
    }, {
      "data" : {
        "id" : "1184",
        "symbol" : "ABF2",
        "Shoot_Descriptions" : "ABF2, AREB1, ATAREB1, abscisic acid responsive elements-binding factor 2",
        "PLAMOM_Match_Shoot" : "NO",
        "Causal_TF" : "",
        "Root_Descriptions" : "",
        "Eccentricity" : 0,
        "Outdegree" : 0,
        "Target_PLAMOM" : "no",
        "PLAMOM_Match_Root" : "",
        "shared_name" : "AT1G45249",
        "Indegree" : 1,
        "Target_tRNA" : "no",
        "t_RNA_motif_shoot" : "NO",
        "SUID" : 1184,
        "SelfLoops" : 0,
        "Target_TF" : "\"ABF2",
        "MolecularAttributes" : "TXNFACTOR",
        "AverageShortestPathLength" : 0.0,
        "selected" : false,
        "Name2" : "ABF2",
        "ClosenessCentrality" : 0.0,
        "Casual_Name" : "",
        "PartnerOfMultiEdgedNodePairs" : 0,
        "ClusteringCoefficient" : 0.0,
        "Target_Name" : "no",
        "Stress" : 0,
        "BetweennessCentrality" : 0.0,
        "t_RNA_motif_Root" : "",
        "EdgeCount" : 1,
        "Causal_PLAMOM" : "",
        "name" : "AT1G45249",
        "Causal_tRNA" : "",
        "IsSingleNode" : false,
        "briefDescription" : "abscisic acid responsive elements-binding factor 2",
        "NeighborhoodConnectivity" : 6.0
      },
      "position" : {
        "x" : 4052.1938580840824,
        "y" : 769.341064453125
      },
      "selected" : false
    }, {
      "data" : {
        "id" : "1183",
        "symbol" : "AT5G13400",
        "Shoot_Descriptions" : "",
        "PLAMOM_Match_Shoot" : "",
        "Causal_TF" : "no",
        "Root_Descriptions" : "Major facilitator superfamily protein",
        "Eccentricity" : 1,
        "Outdegree" : 4,
        "Target_PLAMOM" : "",
        "PLAMOM_Match_Root" : "NO",
        "shared_name" : "AT5G13400",
        "Indegree" : 0,
        "Target_tRNA" : "",
        "t_RNA_motif_shoot" : "",
        "SUID" : 1183,
        "SelfLoops" : 0,
        "Target_TF" : "",
        "MolecularAttributes" : "protein_coding",
        "AverageShortestPathLength" : 1.0,
        "selected" : false,
        "Name2" : "AT5G13400",
        "ClosenessCentrality" : 1.0,
        "Casual_Name" : "Major facilitator superfamily protein",
        "PartnerOfMultiEdgedNodePairs" : 0,
        "ClusteringCoefficient" : 0.0,
        "Target_Name" : "",
        "Stress" : 0,
        "BetweennessCentrality" : 0.0,
        "t_RNA_motif_Root" : "NO",
        "EdgeCount" : 4,
        "Causal_PLAMOM" : "no",
        "name" : "AT5G13400",
        "Causal_tRNA" : "no",
        "IsSingleNode" : false,
        "briefDescription" : "Major facilitator superfamily protein",
        "NeighborhoodConnectivity" : 4.75
      },
      "position" : {
        "x" : -6300.4814453125,
        "y" : -263.35589599609375
      },
      "selected" : false
    }, {
      "data" : {
        "id" : "1182",
        "symbol" : "AT1G47310",
        "degree_layout" : 2,
        "Shoot_Descriptions" : "unknown protein; FUNCTIONS IN: molecular_function unknown; INVOLVED IN: biological_process unknown; LOCATED IN: endomembrane system; EXPRESSED IN: 24 plant structures; EXPRESSED DURING: 13 growth stages; Has 45 Blast hits to 45 proteins in 11 species: Archae - 0; Bacteria - 0; Metazoa - 0; Fungi - 0; Plants - 45; Viruses - 0; Other Eukaryotes - 0 (source: NCBI BLink).",
        "PLAMOM_Match_Shoot" : "NO",
        "Causal_TF" : "",
        "Root_Descriptions" : "",
        "Eccentricity" : 0,
        "Outdegree" : 0,
        "Target_PLAMOM" : "no",
        "PLAMOM_Match_Root" : "",
        "shared_name" : "AT1G47310",
        "Indegree" : 5,
        "Target_tRNA" : "no",
        "t_RNA_motif_shoot" : "YES",
        "SUID" : 1182,
        "SelfLoops" : 0,
        "Target_TF" : "unknown protein; FUNCTIONS IN: molecular_function unknown; INVOLVED IN: biological_process unknown; LOCATED IN: endomembrane system; EXPRESSED IN: 24 plant structures; EXPRESSED DURING: 13 growth stages; Has 45 Blast hits to 45 proteins in 11 species: Archae - 0; Bacteria - 0; Metazoa - 0; Fungi - 0; Plants - 45; Viruses - 0; Other Eukaryotes - 0 (source: NCBI BLink).",
        "MolecularAttributes" : "protein_coding",
        "AverageShortestPathLength" : 0.0,
        "selected" : false,
        "Name2" : "AT1G47310",
        "ClosenessCentrality" : 0.0,
        "Casual_Name" : "",
        "PartnerOfMultiEdgedNodePairs" : 0,
        "ClusteringCoefficient" : 0.0,
        "Target_Name" : "AT1G47310",
        "Stress" : 0,
        "BetweennessCentrality" : 0.0,
        "t_RNA_motif_Root" : "",
        "EdgeCount" : 5,
        "Causal_PLAMOM" : "",
        "name" : "AT1G47310",
        "Causal_tRNA" : "",
        "IsSingleNode" : false,
        "briefDescription" : "signal peptidase I",
        "NeighborhoodConnectivity" : 19.6
      },
      "position" : {
        "x" : 3900.640819955046,
        "y" : 57.34459686279297
      },
      "selected" : false
    }, {
      "data" : {
        "id" : "1181",
        "symbol" : "AT2G44380",
        "Shoot_Descriptions" : "",
        "PLAMOM_Match_Shoot" : "",
        "Causal_TF" : "no",
        "Root_Descriptions" : "Cysteine/Histidine-rich C1 domain family protein",
        "Eccentricity" : 11,
        "Outdegree" : 9,
        "Target_PLAMOM" : "",
        "PLAMOM_Match_Root" : "NO",
        "shared_name" : "AT2G44380",
        "Indegree" : 0,
        "Target_tRNA" : "",
        "t_RNA_motif_shoot" : "",
        "SUID" : 1181,
        "SelfLoops" : 0,
        "Target_TF" : "",
        "MolecularAttributes" : "protein_coding",
        "AverageShortestPathLength" : 6.41478439,
        "selected" : false,
        "Name2" : "AT2G44380",
        "ClosenessCentrality" : 0.15588988,
        "Casual_Name" : "Cysteine/Histidine-rich C1 domain family protein",
        "PartnerOfMultiEdgedNodePairs" : 0,
        "ClusteringCoefficient" : 0.0,
        "Target_Name" : "",
        "Stress" : 0,
        "BetweennessCentrality" : 0.0,
        "t_RNA_motif_Root" : "NO",
        "EdgeCount" : 9,
        "Causal_PLAMOM" : "no",
        "name" : "AT2G44380",
        "Causal_tRNA" : "no",
        "IsSingleNode" : false,
        "briefDescription" : "Cysteine/Histidine-rich C1 domain family protein",
        "NeighborhoodConnectivity" : 2.6666666666666665
      },
      "position" : {
        "x" : -6207.5859375,
        "y" : 887.1748046875
      },
      "selected" : false
    }, {
      "data" : {
        "id" : "1180",
        "symbol" : "AT1G48040",
        "Shoot_Descriptions" : "Protein phosphatase 2C family protein",
        "PLAMOM_Match_Shoot" : "NO",
        "Causal_TF" : "",
        "Root_Descriptions" : "",
        "Eccentricity" : 0,
        "Outdegree" : 0,
        "Target_PLAMOM" : "no",
        "PLAMOM_Match_Root" : "",
        "shared_name" : "AT1G48040",
        "Indegree" : 1,
        "Target_tRNA" : "AT1G48040",
        "t_RNA_motif_shoot" : "YES",
        "SUID" : 1180,
        "SelfLoops" : 0,
        "Target_TF" : "no",
        "MolecularAttributes" : "protein_coding",
        "AverageShortestPathLength" : 0.0,
        "selected" : false,
        "Name2" : "AT1G48040",
        "ClosenessCentrality" : 0.0,
        "Casual_Name" : "",
        "PartnerOfMultiEdgedNodePairs" : 0,
        "ClusteringCoefficient" : 0.0,
        "Target_Name" : "Protein phosphatase 2C family protein",
        "Stress" : 0,
        "BetweennessCentrality" : 0.0,
        "t_RNA_motif_Root" : "",
        "EdgeCount" : 1,
        "Causal_PLAMOM" : "",
        "name" : "AT1G48040",
        "Causal_tRNA" : "",
        "IsSingleNode" : false,
        "briefDescription" : "Protein phosphatase 2C family protein",
        "NeighborhoodConnectivity" : 9.0
      },
      "position" : {
        "x" : 3774.853466439421,
        "y" : 1219.2364501953125
      },
      "selected" : false
    }, {
      "data" : {
        "id" : "1179",
        "symbol" : "PP2-B1",
        "Shoot_Descriptions" : "",
        "PLAMOM_Match_Shoot" : "",
        "Causal_TF" : "PP2-B1",
        "Root_Descriptions" : "AtPP2-B1, PP2-B1, phloem protein 2-B1",
        "Eccentricity" : 15,
        "Outdegree" : 4,
        "Target_PLAMOM" : "no",
        "PLAMOM_Match_Root" : "NO",
        "shared_name" : "AT2G02230",
        "Indegree" : 2,
        "Target_tRNA" : "no",
        "t_RNA_motif_shoot" : "",
        "SUID" : 1179,
        "SelfLoops" : 0,
        "Target_TF" : "PP2-B1",
        "MolecularAttributes" : "protein_coding",
        "AverageShortestPathLength" : 9.84567901,
        "selected" : false,
        "Name2" : "PP2-B1",
        "ClosenessCentrality" : 0.1015674,
        "Casual_Name" : "\"AtPP2-B1",
        "PartnerOfMultiEdgedNodePairs" : 0,
        "ClusteringCoefficient" : 0.0,
        "Target_Name" : "\"AtPP2-B1",
        "Stress" : 3644,
        "BetweennessCentrality" : 8.6649E-4,
        "t_RNA_motif_Root" : "NO",
        "EdgeCount" : 6,
        "Causal_PLAMOM" : "no",
        "name" : "AT2G02230",
        "Causal_tRNA" : "no",
        "IsSingleNode" : false,
        "briefDescription" : "phloem protein 2-B1",
        "NeighborhoodConnectivity" : 2.1666666666666665
      },
      "position" : {
        "x" : -7376.841796875,
        "y" : 3085.50439453125
      },
      "selected" : false
    }, {
      "data" : {
        "id" : "1178",
        "symbol" : "AT1G49560",
        "Shoot_Descriptions" : "Homeodomain-like superfamily protein",
        "PLAMOM_Match_Shoot" : "NO",
        "Causal_TF" : "",
        "Root_Descriptions" : "",
        "Eccentricity" : 0,
        "Outdegree" : 0,
        "Target_PLAMOM" : "phloem protein 2-B1\"",
        "PLAMOM_Match_Root" : "",
        "shared_name" : "AT1G49560",
        "Indegree" : 1,
        "Target_tRNA" : "no",
        "t_RNA_motif_shoot" : "NO",
        "SUID" : 1178,
        "SelfLoops" : 0,
        "Target_TF" : "no",
        "MolecularAttributes" : "TXNFACTOR",
        "AverageShortestPathLength" : 0.0,
        "selected" : false,
        "Name2" : "AT1G49560",
        "ClosenessCentrality" : 0.0,
        "Casual_Name" : "",
        "PartnerOfMultiEdgedNodePairs" : 0,
        "ClusteringCoefficient" : 0.0,
        "Target_Name" : "no",
        "Stress" : 0,
        "BetweennessCentrality" : 0.0,
        "t_RNA_motif_Root" : "",
        "EdgeCount" : 1,
        "Causal_PLAMOM" : "",
        "name" : "AT1G49560",
        "Causal_tRNA" : "",
        "IsSingleNode" : false,
        "briefDescription" : "Homeodomain-like superfamily protein",
        "NeighborhoodConnectivity" : 6.0
      },
      "position" : {
        "x" : 2988.291454720671,
        "y" : 36.59449005126953
      },
      "selected" : false
    }, {
      "data" : {
        "id" : "1177",
        "symbol" : "AT1G50020",
        "Shoot_Descriptions" : "unknown protein; FUNCTIONS IN: molecular_function unknown; INVOLVED IN: biological_process unknown; LOCATED IN: chloroplast thylakoid membrane, chloroplast; EXPRESSED IN: 24 plant structures; EXPRESSED DURING: 15 growth stages; Has 72 Blast hits to 72 proteins in 27 species: Archae - 0; Bacteria - 0; Metazoa - 0; Fungi - 0; Plants - 72; Viruses - 0; Other Eukaryotes - 0 (source: NCBI BLink).",
        "PLAMOM_Match_Shoot" : "YES",
        "Causal_TF" : "",
        "Root_Descriptions" : "",
        "Eccentricity" : 0,
        "Outdegree" : 0,
        "Target_PLAMOM" : "no",
        "PLAMOM_Match_Root" : "",
        "shared_name" : "AT1G50020",
        "Indegree" : 7,
        "Target_tRNA" : "AT1G50020",
        "t_RNA_motif_shoot" : "NO",
        "SUID" : 1177,
        "SelfLoops" : 0,
        "Target_TF" : "\"unknown protein; FUNCTIONS IN: molecular_function unknown; INVOLVED IN: biological_process unknown; LOCATED IN: chloroplast thylakoid membrane",
        "MolecularAttributes" : "protein_coding",
        "AverageShortestPathLength" : 0.0,
        "selected" : false,
        "Name2" : "AT1G50020",
        "ClosenessCentrality" : 0.0,
        "Casual_Name" : "",
        "PartnerOfMultiEdgedNodePairs" : 0,
        "ClusteringCoefficient" : 0.0,
        "Target_Name" : "no",
        "Stress" : 0,
        "BetweennessCentrality" : 0.0,
        "t_RNA_motif_Root" : "",
        "EdgeCount" : 7,
        "Causal_PLAMOM" : "",
        "name" : "AT1G50020",
        "Causal_tRNA" : "",
        "IsSingleNode" : false,
        "briefDescription" : "tubulin alpha-6 chain",
        "NeighborhoodConnectivity" : 5.0
      },
      "position" : {
        "x" : 3440.239696908171,
        "y" : 559.2396240234375
      },
      "selected" : false
    }, {
      "data" : {
        "id" : "1176",
        "symbol" : "ENO1",
        "Shoot_Descriptions" : "",
        "PLAMOM_Match_Shoot" : "",
        "Causal_TF" : "enolase 1\"",
        "Root_Descriptions" : "ENO1, enolase 1",
        "Eccentricity" : 1,
        "Outdegree" : 1,
        "Target_PLAMOM" : "",
        "PLAMOM_Match_Root" : "NO",
        "shared_name" : "AT1G74030",
        "Indegree" : 0,
        "Target_tRNA" : "",
        "t_RNA_motif_shoot" : "",
        "SUID" : 1176,
        "SelfLoops" : 0,
        "Target_TF" : "",
        "MolecularAttributes" : "METABOLIC",
        "AverageShortestPathLength" : 1.0,
        "selected" : false,
        "Name2" : "ENO1",
        "ClosenessCentrality" : 1.0,
        "Casual_Name" : "\"ENO1",
        "PartnerOfMultiEdgedNodePairs" : 0,
        "ClusteringCoefficient" : 0.0,
        "Target_Name" : "",
        "Stress" : 0,
        "BetweennessCentrality" : 0.0,
        "t_RNA_motif_Root" : "NO",
        "EdgeCount" : 1,
        "Causal_PLAMOM" : "no",
        "name" : "AT1G74030",
        "Causal_tRNA" : "no",
        "IsSingleNode" : false,
        "briefDescription" : "enolase 1",
        "NeighborhoodConnectivity" : 7.0
      },
      "position" : {
        "x" : -6741.14013671875,
        "y" : 672.086669921875
      },
      "selected" : false
    }, {
      "data" : {
        "id" : "1175",
        "symbol" : "Kin3",
        "Shoot_Descriptions" : "",
        "PLAMOM_Match_Shoot" : "",
        "Causal_TF" : "no",
        "Root_Descriptions" : "Protein kinase superfamily protein",
        "Eccentricity" : 13,
        "Outdegree" : 3,
        "Target_PLAMOM" : "no",
        "PLAMOM_Match_Root" : "NO",
        "shared_name" : "AT2G17220",
        "Indegree" : 2,
        "Target_tRNA" : "no",
        "t_RNA_motif_shoot" : "",
        "SUID" : 1175,
        "SelfLoops" : 0,
        "Target_TF" : "Protein kinase superfamily protein",
        "MolecularAttributes" : "protein_coding",
        "AverageShortestPathLength" : 8.17364017,
        "selected" : false,
        "Name2" : "Kin3",
        "ClosenessCentrality" : 0.12234451,
        "Casual_Name" : "Protein kinase superfamily protein",
        "PartnerOfMultiEdgedNodePairs" : 0,
        "ClusteringCoefficient" : 0.0,
        "Target_Name" : "no",
        "Stress" : 344,
        "BetweennessCentrality" : 7.787E-5,
        "t_RNA_motif_Root" : "NO",
        "EdgeCount" : 5,
        "Causal_PLAMOM" : "no",
        "name" : "AT2G17220",
        "Causal_tRNA" : "no",
        "IsSingleNode" : false,
        "briefDescription" : "Protein kinase superfamily protein",
        "NeighborhoodConnectivity" : 11.6
      },
      "position" : {
        "x" : -7128.26171875,
        "y" : 3061.21533203125
      },
      "selected" : false
    }, {
      "data" : {
        "id" : "1174",
        "symbol" : "AT4G09770",
        "Shoot_Descriptions" : "",
        "PLAMOM_Match_Shoot" : "",
        "Causal_TF" : "no",
        "Root_Descriptions" : "TRAF-like family protein",
        "Eccentricity" : 1,
        "Outdegree" : 4,
        "Target_PLAMOM" : "",
        "PLAMOM_Match_Root" : "NO",
        "shared_name" : "AT4G09770",
        "Indegree" : 0,
        "Target_tRNA" : "",
        "t_RNA_motif_shoot" : "",
        "SUID" : 1174,
        "SelfLoops" : 0,
        "Target_TF" : "",
        "MolecularAttributes" : "protein_coding",
        "AverageShortestPathLength" : 1.0,
        "selected" : false,
        "Name2" : "AT4G09770",
        "ClosenessCentrality" : 1.0,
        "Casual_Name" : "TRAF-like family protein",
        "PartnerOfMultiEdgedNodePairs" : 0,
        "ClusteringCoefficient" : 0.0,
        "Target_Name" : "",
        "Stress" : 0,
        "BetweennessCentrality" : 0.0,
        "t_RNA_motif_Root" : "NO",
        "EdgeCount" : 4,
        "Causal_PLAMOM" : "no",
        "name" : "AT4G09770",
        "Causal_tRNA" : "no",
        "IsSingleNode" : false,
        "briefDescription" : "TRAF-like family protein",
        "NeighborhoodConnectivity" : 14.0
      },
      "position" : {
        "x" : -6447.72119140625,
        "y" : 520.7867431640625
      },
      "selected" : false
    }, {
      "data" : {
        "id" : "1173",
        "symbol" : "AT1G63430",
        "Shoot_Descriptions" : "",
        "PLAMOM_Match_Shoot" : "",
        "Causal_TF" : "no",
        "Root_Descriptions" : "Leucine-rich repeat protein kinase family protein",
        "Eccentricity" : 12,
        "Outdegree" : 7,
        "Target_PLAMOM" : "no",
        "PLAMOM_Match_Root" : "NO",
        "shared_name" : "AT1G63430",
        "Indegree" : 4,
        "Target_tRNA" : "no",
        "t_RNA_motif_shoot" : "",
        "SUID" : 1173,
        "SelfLoops" : 0,
        "Target_TF" : "no",
        "MolecularAttributes" : "protein_coding",
        "AverageShortestPathLength" : 7.05416667,
        "selected" : false,
        "Name2" : "AT1G63430",
        "ClosenessCentrality" : 0.14176019,
        "Casual_Name" : "Leucine-rich repeat protein kinase family protein",
        "PartnerOfMultiEdgedNodePairs" : 0,
        "ClusteringCoefficient" : 0.0,
        "Target_Name" : "Leucine-rich repeat protein kinase family protein",
        "Stress" : 2309,
        "BetweennessCentrality" : 5.6761E-4,
        "t_RNA_motif_Root" : "NO",
        "EdgeCount" : 11,
        "Causal_PLAMOM" : "no",
        "name" : "AT1G63430",
        "Causal_tRNA" : "no",
        "IsSingleNode" : false,
        "briefDescription" : "Leucine-rich repeat protein kinase family protein",
        "NeighborhoodConnectivity" : 7.363636363636363
      },
      "position" : {
        "x" : -7050.318359375,
        "y" : 3360.86279296875
      },
      "selected" : false
    }, {
      "data" : {
        "id" : "1172",
        "symbol" : "emb2742",
        "Shoot_Descriptions" : "emb2742, CTP synthase family protein",
        "PLAMOM_Match_Shoot" : "NO",
        "Causal_TF" : "CTP synthase family protein\"",
        "Root_Descriptions" : "emb2742, CTP synthase family protein",
        "Eccentricity" : 11,
        "Outdegree" : 3,
        "Target_PLAMOM" : "no",
        "PLAMOM_Match_Root" : "NO",
        "shared_name" : "AT3G12670",
        "Indegree" : 1,
        "Target_tRNA" : "no",
        "t_RNA_motif_shoot" : "NO",
        "SUID" : 1172,
        "SelfLoops" : 0,
        "Target_TF" : "CTP synthase family protein\"",
        "MolecularAttributes" : "METABOLIC",
        "AverageShortestPathLength" : 6.70292887,
        "selected" : false,
        "Name2" : "emb2742",
        "ClosenessCentrality" : 0.14918851,
        "Casual_Name" : "\"emb2742",
        "PartnerOfMultiEdgedNodePairs" : 0,
        "ClusteringCoefficient" : 0.0,
        "Target_Name" : "\"emb2742",
        "Stress" : 13427,
        "BetweennessCentrality" : 0.00266901,
        "t_RNA_motif_Root" : "NO",
        "EdgeCount" : 4,
        "Causal_PLAMOM" : "no",
        "name" : "AT3G12670",
        "Causal_tRNA" : "no",
        "IsSingleNode" : false,
        "briefDescription" : "CTP synthase family protein",
        "NeighborhoodConnectivity" : 6.0
      },
      "position" : {
        "x" : -731.3821902988602,
        "y" : 659.169677734375
      },
      "selected" : false
    }, {
      "data" : {
        "id" : "1171",
        "symbol" : "VIIIA",
        "Shoot_Descriptions" : "ATVIIIA, VIIIA, P-loop containing nucleoside triphosphate hydrolases superfamily protein",
        "PLAMOM_Match_Shoot" : "NO",
        "Causal_TF" : "",
        "Root_Descriptions" : "",
        "Eccentricity" : 0,
        "Outdegree" : 0,
        "Target_PLAMOM" : "cytokinin oxidase 4\"",
        "PLAMOM_Match_Root" : "",
        "shared_name" : "AT1G50360",
        "Indegree" : 1,
        "Target_tRNA" : "no",
        "t_RNA_motif_shoot" : "YES",
        "SUID" : 1171,
        "SelfLoops" : 0,
        "Target_TF" : "AT1G50360",
        "MolecularAttributes" : "protein_coding",
        "AverageShortestPathLength" : 0.0,
        "selected" : false,
        "Name2" : "VIIIA",
        "ClosenessCentrality" : 0.0,
        "Casual_Name" : "",
        "PartnerOfMultiEdgedNodePairs" : 0,
        "ClusteringCoefficient" : 0.0,
        "Target_Name" : "no",
        "Stress" : 0,
        "BetweennessCentrality" : 0.0,
        "t_RNA_motif_Root" : "",
        "EdgeCount" : 1,
        "Causal_PLAMOM" : "",
        "name" : "AT1G50360",
        "Causal_tRNA" : "",
        "IsSingleNode" : false,
        "briefDescription" : "P-loop containing nucleoside triphosphate hydrolases superfamily protein",
        "NeighborhoodConnectivity" : 7.0
      },
      "position" : {
        "x" : 4460.620068001921,
        "y" : 1075.4876708984375
      },
      "selected" : false
    }, {
      "data" : {
        "id" : "1170",
        "symbol" : "AT1G50920",
        "degree_layout" : 2,
        "Shoot_Descriptions" : "Nucleolar GTP-binding protein",
        "PLAMOM_Match_Shoot" : "YES",
        "Causal_TF" : "",
        "Root_Descriptions" : "",
        "Eccentricity" : 0,
        "Outdegree" : 0,
        "Target_PLAMOM" : "AT1G50920",
        "PLAMOM_Match_Root" : "",
        "shared_name" : "AT1G50920",
        "Indegree" : 3,
        "Target_tRNA" : "no",
        "t_RNA_motif_shoot" : "NO",
        "SUID" : 1170,
        "SelfLoops" : 0,
        "Target_TF" : "no",
        "MolecularAttributes" : "protein_coding",
        "AverageShortestPathLength" : 0.0,
        "selected" : false,
        "Name2" : "AT1G50920",
        "ClosenessCentrality" : 0.0,
        "Casual_Name" : "",
        "PartnerOfMultiEdgedNodePairs" : 0,
        "ClusteringCoefficient" : 0.0,
        "Target_Name" : "Nucleolar GTP-binding protein",
        "Stress" : 0,
        "BetweennessCentrality" : 0.0,
        "t_RNA_motif_Root" : "",
        "EdgeCount" : 3,
        "Causal_PLAMOM" : "",
        "name" : "AT1G50920",
        "Causal_tRNA" : "",
        "IsSingleNode" : false,
        "briefDescription" : "Nucleolar GTP-binding protein",
        "NeighborhoodConnectivity" : 39.0
      },
      "position" : {
        "x" : 3636.8972272247074,
        "y" : -411.65118408203125
      },
      "selected" : false
    }, {
      "data" : {
        "id" : "1169",
        "symbol" : "AT4G36010",
        "Shoot_Descriptions" : "",
        "PLAMOM_Match_Shoot" : "",
        "Causal_TF" : "no",
        "Root_Descriptions" : "Pathogenesis-related thaumatin superfamily protein",
        "Eccentricity" : 1,
        "Outdegree" : 3,
        "Target_PLAMOM" : "",
        "PLAMOM_Match_Root" : "NO",
        "shared_name" : "AT4G36010",
        "Indegree" : 0,
        "Target_tRNA" : "",
        "t_RNA_motif_shoot" : "",
        "SUID" : 1169,
        "SelfLoops" : 0,
        "Target_TF" : "",
        "MolecularAttributes" : "protein_coding",
        "AverageShortestPathLength" : 1.0,
        "selected" : false,
        "Name2" : "AT4G36010",
        "ClosenessCentrality" : 1.0,
        "Casual_Name" : "Pathogenesis-related thaumatin superfamily protein",
        "PartnerOfMultiEdgedNodePairs" : 0,
        "ClusteringCoefficient" : 0.0,
        "Target_Name" : "",
        "Stress" : 0,
        "BetweennessCentrality" : 0.0,
        "t_RNA_motif_Root" : "NO",
        "EdgeCount" : 3,
        "Causal_PLAMOM" : "no",
        "name" : "AT4G36010",
        "Causal_tRNA" : "no",
        "IsSingleNode" : false,
        "briefDescription" : "Pathogenesis-related thaumatin superfamily protein",
        "NeighborhoodConnectivity" : 20.0
      },
      "position" : {
        "x" : -5409.4453125,
        "y" : 198.36155700683594
      },
      "selected" : false
    }, {
      "data" : {
        "id" : "1168",
        "symbol" : "AT1G51380",
        "Shoot_Descriptions" : "DEA(D/H)-box RNA helicase family protein",
        "PLAMOM_Match_Shoot" : "NO",
        "Causal_TF" : "",
        "Root_Descriptions" : "",
        "Eccentricity" : 0,
        "Outdegree" : 0,
        "Target_PLAMOM" : "no",
        "PLAMOM_Match_Root" : "",
        "shared_name" : "AT1G51380",
        "Indegree" : 1,
        "Target_tRNA" : "no",
        "t_RNA_motif_shoot" : "NO",
        "SUID" : 1168,
        "SelfLoops" : 0,
        "Target_TF" : "no",
        "MolecularAttributes" : "protein_coding",
        "AverageShortestPathLength" : 0.0,
        "selected" : false,
        "Name2" : "AT1G51380",
        "ClosenessCentrality" : 0.0,
        "Casual_Name" : "",
        "PartnerOfMultiEdgedNodePairs" : 0,
        "ClusteringCoefficient" : 0.0,
        "Target_Name" : "DEA(D/H)-box RNA helicase family protein",
        "Stress" : 0,
        "BetweennessCentrality" : 0.0,
        "t_RNA_motif_Root" : "",
        "EdgeCount" : 1,
        "Causal_PLAMOM" : "",
        "name" : "AT1G51380",
        "Causal_tRNA" : "",
        "IsSingleNode" : false,
        "briefDescription" : "DEA(D/H)-box RNA helicase family protein",
        "NeighborhoodConnectivity" : 3.0
      },
      "position" : {
        "x" : 4881.304638314421,
        "y" : 244.59121704101562
      },
      "selected" : false
    }, {
      "data" : {
        "id" : "1167",
        "symbol" : "Y14",
        "Shoot_Descriptions" : "Y14, RNA-binding (RRM/RBD/RNP motifs) family protein",
        "PLAMOM_Match_Shoot" : "YES",
        "Causal_TF" : "",
        "Root_Descriptions" : "",
        "Eccentricity" : 0,
        "Outdegree" : 0,
        "Target_PLAMOM" : "AT1G51510",
        "PLAMOM_Match_Root" : "",
        "shared_name" : "AT1G51510",
        "Indegree" : 7,
        "Target_tRNA" : "no",
        "t_RNA_motif_shoot" : "NO",
        "SUID" : 1167,
        "SelfLoops" : 0,
        "Target_TF" : "RNA-binding (RRM/RBD/RNP motifs) family protein\"",
        "MolecularAttributes" : "protein_coding",
        "AverageShortestPathLength" : 0.0,
        "selected" : false,
        "Name2" : "Y14",
        "ClosenessCentrality" : 0.0,
        "Casual_Name" : "",
        "PartnerOfMultiEdgedNodePairs" : 0,
        "ClusteringCoefficient" : 0.0,
        "Target_Name" : "\"Y14",
        "Stress" : 0,
        "BetweennessCentrality" : 0.0,
        "t_RNA_motif_Root" : "",
        "EdgeCount" : 7,
        "Causal_PLAMOM" : "",
        "name" : "AT1G51510",
        "Causal_tRNA" : "",
        "IsSingleNode" : false,
        "briefDescription" : "RNA-binding (RRM/RBD/RNP motifs) family protein",
        "NeighborhoodConnectivity" : 7.0
      },
      "position" : {
        "x" : 4496.132763314421,
        "y" : -37.835784912109375
      },
      "selected" : false
    }, {
      "data" : {
        "id" : "1166",
        "symbol" : "HMA2",
        "Shoot_Descriptions" : "",
        "PLAMOM_Match_Shoot" : "",
        "Causal_TF" : "HMA2",
        "Root_Descriptions" : "ATHMA2, HMA2, heavy metal atpase 2",
        "Eccentricity" : 1,
        "Outdegree" : 4,
        "Target_PLAMOM" : "",
        "PLAMOM_Match_Root" : "NO",
        "shared_name" : "AT4G30110",
        "Indegree" : 0,
        "Target_tRNA" : "",
        "t_RNA_motif_shoot" : "",
        "SUID" : 1166,
        "SelfLoops" : 0,
        "Target_TF" : "",
        "MolecularAttributes" : "protein_coding",
        "AverageShortestPathLength" : 1.0,
        "selected" : false,
        "Name2" : "HMA2",
        "ClosenessCentrality" : 1.0,
        "Casual_Name" : "\"ATHMA2",
        "PartnerOfMultiEdgedNodePairs" : 0,
        "ClusteringCoefficient" : 0.0,
        "Target_Name" : "",
        "Stress" : 0,
        "BetweennessCentrality" : 0.0,
        "t_RNA_motif_Root" : "NO",
        "EdgeCount" : 4,
        "Causal_PLAMOM" : "no",
        "name" : "AT4G30110",
        "Causal_tRNA" : "no",
        "IsSingleNode" : false,
        "briefDescription" : "heavy metal atpase 2",
        "NeighborhoodConnectivity" : 7.0
      },
      "position" : {
        "x" : -5485.822265625,
        "y" : -214.20375061035156
      },
      "selected" : false
    }, {
      "data" : {
        "id" : "1165",
        "symbol" : "AT5G62710",
        "Shoot_Descriptions" : "",
        "PLAMOM_Match_Shoot" : "",
        "Causal_TF" : "no",
        "Root_Descriptions" : "Leucine-rich repeat protein kinase family protein",
        "Eccentricity" : 12,
        "Outdegree" : 5,
        "Target_PLAMOM" : "",
        "PLAMOM_Match_Root" : "NO",
        "shared_name" : "AT5G62710",
        "Indegree" : 0,
        "Target_tRNA" : "",
        "t_RNA_motif_shoot" : "",
        "SUID" : 1165,
        "SelfLoops" : 0,
        "Target_TF" : "",
        "MolecularAttributes" : "protein_coding",
        "AverageShortestPathLength" : 6.90248963,
        "selected" : false,
        "Name2" : "AT5G62710",
        "ClosenessCentrality" : 0.14487526,
        "Casual_Name" : "Leucine-rich repeat protein kinase family protein",
        "PartnerOfMultiEdgedNodePairs" : 0,
        "ClusteringCoefficient" : 0.0,
        "Target_Name" : "",
        "Stress" : 0,
        "BetweennessCentrality" : 0.0,
        "t_RNA_motif_Root" : "NO",
        "EdgeCount" : 5,
        "Causal_PLAMOM" : "no",
        "name" : "AT5G62710",
        "Causal_tRNA" : "no",
        "IsSingleNode" : false,
        "briefDescription" : "Leucine-rich repeat protein kinase family protein",
        "NeighborhoodConnectivity" : 5.0
      },
      "position" : {
        "x" : -5359.21435546875,
        "y" : 20.028356552124023
      },
      "selected" : false
    }, {
      "data" : {
        "id" : "1164",
        "symbol" : "MUR4",
        "Shoot_Descriptions" : "",
        "PLAMOM_Match_Shoot" : "",
        "Causal_TF" : "MUR4",
        "Root_Descriptions" : "HSR8, MUR4, UXE1, NAD(P)-binding Rossmann-fold superfamily protein",
        "Eccentricity" : 11,
        "Outdegree" : 6,
        "Target_PLAMOM" : "",
        "PLAMOM_Match_Root" : "NO",
        "shared_name" : "AT1G30620",
        "Indegree" : 0,
        "Target_tRNA" : "",
        "t_RNA_motif_shoot" : "",
        "SUID" : 1164,
        "SelfLoops" : 0,
        "Target_TF" : "",
        "MolecularAttributes" : "METABOLIC",
        "AverageShortestPathLength" : 6.58178054,
        "selected" : false,
        "Name2" : "MUR4",
        "ClosenessCentrality" : 0.15193457,
        "Casual_Name" : "\"HSR8",
        "PartnerOfMultiEdgedNodePairs" : 0,
        "ClusteringCoefficient" : 0.0,
        "Target_Name" : "",
        "Stress" : 0,
        "BetweennessCentrality" : 0.0,
        "t_RNA_motif_Root" : "NO",
        "EdgeCount" : 6,
        "Causal_PLAMOM" : "no",
        "name" : "AT1G30620",
        "Causal_tRNA" : "no",
        "IsSingleNode" : false,
        "briefDescription" : "NAD(P)-binding Rossmann-fold superfamily protein",
        "NeighborhoodConnectivity" : 9.0
      },
      "position" : {
        "x" : -5669.4736328125,
        "y" : -209.97718811035156
      },
      "selected" : false
    }, {
      "data" : {
        "id" : "1163",
        "symbol" : "HK1",
        "Shoot_Descriptions" : "",
        "PLAMOM_Match_Shoot" : "",
        "Causal_TF" : "ATHK1",
        "Root_Descriptions" : "AHK1, ATHK1, HK1, histidine kinase 1",
        "Eccentricity" : 13,
        "Outdegree" : 2,
        "Target_PLAMOM" : "",
        "PLAMOM_Match_Root" : "YES",
        "shared_name" : "AT2G17820",
        "Indegree" : 0,
        "Target_tRNA" : "",
        "t_RNA_motif_shoot" : "",
        "SUID" : 1163,
        "SelfLoops" : 0,
        "Target_TF" : "",
        "MolecularAttributes" : "protein_coding",
        "AverageShortestPathLength" : 8.06263048,
        "selected" : false,
        "Name2" : "HK1",
        "ClosenessCentrality" : 0.124029,
        "Casual_Name" : "\"AHK1",
        "PartnerOfMultiEdgedNodePairs" : 0,
        "ClusteringCoefficient" : 0.0,
        "Target_Name" : "",
        "Stress" : 0,
        "BetweennessCentrality" : 0.0,
        "t_RNA_motif_Root" : "NO",
        "EdgeCount" : 2,
        "Causal_PLAMOM" : "AT2G17820",
        "name" : "AT2G17820",
        "Causal_tRNA" : "no",
        "IsSingleNode" : false,
        "briefDescription" : "histidine kinase 1",
        "NeighborhoodConnectivity" : 6.5
      },
      "position" : {
        "x" : -5587.15283203125,
        "y" : -143.9663848876953
      },
      "selected" : false
    }, {
      "data" : {
        "id" : "1162",
        "symbol" : "AT3G22530",
        "Shoot_Descriptions" : "",
        "PLAMOM_Match_Shoot" : "",
        "Causal_TF" : "no",
        "Root_Descriptions" : "unknown protein; FUNCTIONS IN: molecular_function unknown; INVOLVED IN: biological_process unknown; LOCATED IN: plasma membrane; EXPRESSED IN: 25 plant structures; EXPRESSED DURING: 15 growth stages; BEST Arabidopsis thaliana protein match is: unknown protein (TAIR:AT4G14830.1); Has 77 Blast hits to 77 proteins in 12 species: Archae - 0; Bacteria - 0; Metazoa - 0; Fungi - 0; Plants - 77; Viruses - 0; Other Eukaryotes - 0 (source: NCBI BLink).",
        "Eccentricity" : 13,
        "Outdegree" : 3,
        "Target_PLAMOM" : "",
        "PLAMOM_Match_Root" : "NO",
        "shared_name" : "AT3G22530",
        "Indegree" : 0,
        "Target_tRNA" : "",
        "t_RNA_motif_shoot" : "",
        "SUID" : 1162,
        "SelfLoops" : 0,
        "Target_TF" : "",
        "MolecularAttributes" : "protein_coding",
        "AverageShortestPathLength" : 8.01666667,
        "selected" : false,
        "Name2" : "AT3G22530",
        "ClosenessCentrality" : 0.12474012,
        "Casual_Name" : "unknown protein; FUNCTIONS IN: molecular_function unknown; INVOLVED IN: biological_process unknown; LOCATED IN: plasma membrane; EXPRESSED IN: 25 plant structures; EXPRESSED DURING: 15 growth stages; BEST Arabidopsis thaliana protein match is: unknown protein (TAIR:AT4G14830.1); Has 77 Blast hits to 77 proteins in 12 species: Archae - 0; Bacteria - 0; Metazoa - 0; Fungi - 0; Plants - 77; Viruses - 0; Other Eukaryotes - 0 (source: NCBI BLink).",
        "PartnerOfMultiEdgedNodePairs" : 0,
        "ClusteringCoefficient" : 0.0,
        "Target_Name" : "",
        "Stress" : 0,
        "BetweennessCentrality" : 0.0,
        "t_RNA_motif_Root" : "NO",
        "EdgeCount" : 3,
        "Causal_PLAMOM" : "no",
        "name" : "AT3G22530",
        "Causal_tRNA" : "no",
        "IsSingleNode" : false,
        "briefDescription" : "heat shock protein",
        "NeighborhoodConnectivity" : 8.0
      },
      "position" : {
        "x" : -5707.64794921875,
        "y" : -315.2616271972656
      },
      "selected" : false
    }, {
      "data" : {
        "id" : "1161",
        "symbol" : "AT1G51805",
        "Shoot_Descriptions" : "Leucine-rich repeat protein kinase family protein",
        "PLAMOM_Match_Shoot" : "NO",
        "Causal_TF" : "no",
        "Root_Descriptions" : "",
        "Eccentricity" : 13,
        "Outdegree" : 1,
        "Target_PLAMOM" : "no",
        "PLAMOM_Match_Root" : "",
        "shared_name" : "AT1G51805",
        "Indegree" : 1,
        "Target_tRNA" : "no",
        "t_RNA_motif_shoot" : "NO",
        "SUID" : 1161,
        "SelfLoops" : 0,
        "Target_TF" : "Leucine-rich repeat protein kinase family protein",
        "MolecularAttributes" : "protein_coding",
        "AverageShortestPathLength" : 8.0125261,
        "selected" : false,
        "Name2" : "AT1G51805",
        "ClosenessCentrality" : 0.12480459,
        "Casual_Name" : "Leucine-rich repeat protein kinase family protein",
        "PartnerOfMultiEdgedNodePairs" : 0,
        "ClusteringCoefficient" : 0.0,
        "Target_Name" : "no",
        "Stress" : 9,
        "BetweennessCentrality" : 1.05E-6,
        "t_RNA_motif_Root" : "",
        "EdgeCount" : 2,
        "Causal_PLAMOM" : "no",
        "name" : "AT1G51805",
        "Causal_tRNA" : "no",
        "IsSingleNode" : false,
        "briefDescription" : "Leucine-rich repeat protein kinase family protein",
        "NeighborhoodConnectivity" : 43.5
      },
      "position" : {
        "x" : 3230.439403939421,
        "y" : 3511.722412109375
      },
      "selected" : false
    }, {
      "data" : {
        "id" : "1160",
        "symbol" : "DGK3",
        "Shoot_Descriptions" : "",
        "PLAMOM_Match_Shoot" : "",
        "Causal_TF" : "DGK3",
        "Root_Descriptions" : "ATDGK3, DGK3, diacylglycerol kinase 3",
        "Eccentricity" : 10,
        "Outdegree" : 6,
        "Target_PLAMOM" : "NAC domain containing protein 96\"",
        "PLAMOM_Match_Root" : "YES",
        "shared_name" : "AT2G18730",
        "Indegree" : 8,
        "Target_tRNA" : "AT5G46590",
        "t_RNA_motif_shoot" : "",
        "SUID" : 1160,
        "SelfLoops" : 0,
        "Target_TF" : "no",
        "MolecularAttributes" : "protein_coding",
        "AverageShortestPathLength" : 5.4623431,
        "selected" : false,
        "Name2" : "DGK3",
        "ClosenessCentrality" : 0.18307162,
        "Casual_Name" : "\"ATDGK3",
        "PartnerOfMultiEdgedNodePairs" : 0,
        "ClusteringCoefficient" : 0.0,
        "Target_Name" : "AT2G18730",
        "Stress" : 12465,
        "BetweennessCentrality" : 0.00285615,
        "t_RNA_motif_Root" : "NO",
        "EdgeCount" : 14,
        "Causal_PLAMOM" : "AT2G18730",
        "name" : "AT2G18730",
        "Causal_tRNA" : "no",
        "IsSingleNode" : false,
        "briefDescription" : "diacylglycerol kinase 3",
        "NeighborhoodConnectivity" : 7.785714285714286
      },
      "position" : {
        "x" : -6392.90771484375,
        "y" : 3297.40771484375
      },
      "selected" : false
    }, {
      "data" : {
        "id" : "1159",
        "symbol" : "NAM",
        "Shoot_Descriptions" : "ANAC018, ATNAM, NAM, NARS2, NAC (No Apical Meristem) domain transcriptional regulator superfamily protein",
        "PLAMOM_Match_Shoot" : "NO",
        "Causal_TF" : "",
        "Root_Descriptions" : "",
        "Eccentricity" : 0,
        "Outdegree" : 0,
        "Target_PLAMOM" : "no",
        "PLAMOM_Match_Root" : "",
        "shared_name" : "AT1G52880",
        "Indegree" : 12,
        "Target_tRNA" : "no",
        "t_RNA_motif_shoot" : "NO",
        "SUID" : 1159,
        "SelfLoops" : 0,
        "Target_TF" : "\"ANAC018",
        "MolecularAttributes" : "TXNFACTOR",
        "AverageShortestPathLength" : 0.0,
        "selected" : false,
        "Name2" : "NAM",
        "ClosenessCentrality" : 0.0,
        "Casual_Name" : "",
        "PartnerOfMultiEdgedNodePairs" : 0,
        "ClusteringCoefficient" : 0.0,
        "Target_Name" : "no",
        "Stress" : 0,
        "BetweennessCentrality" : 0.0,
        "t_RNA_motif_Root" : "",
        "EdgeCount" : 12,
        "Causal_PLAMOM" : "",
        "name" : "AT1G52880",
        "Causal_tRNA" : "",
        "IsSingleNode" : false,
        "briefDescription" : "NAC (No Apical Meristem) domain transcriptional regulator superfamily protein",
        "NeighborhoodConnectivity" : 17.0
      },
      "position" : {
        "x" : 4084.972607064421,
        "y" : 147.33982849121094
      },
      "selected" : false
    }, {
      "data" : {
        "id" : "1158",
        "symbol" : "PFA-DSP2",
        "degree_layout" : 9,
        "Shoot_Descriptions" : "",
        "PLAMOM_Match_Shoot" : "",
        "Causal_TF" : "no",
        "Root_Descriptions" : "Phosphotyrosine protein phosphatases superfamily protein",
        "Eccentricity" : 12,
        "Outdegree" : 5,
        "Target_PLAMOM" : "no",
        "PLAMOM_Match_Root" : "NO",
        "shared_name" : "AT2G32960",
        "Indegree" : 67,
        "Target_tRNA" : "no",
        "t_RNA_motif_shoot" : "",
        "SUID" : 1158,
        "SelfLoops" : 0,
        "Target_TF" : "no",
        "MolecularAttributes" : "protein_coding",
        "AverageShortestPathLength" : 7.02719665,
        "selected" : false,
        "Name2" : "PFA-DSP2",
        "ClosenessCentrality" : 0.14230426,
        "Casual_Name" : "Phosphotyrosine protein phosphatases superfamily protein",
        "PartnerOfMultiEdgedNodePairs" : 0,
        "ClusteringCoefficient" : 5.8685E-4,
        "Target_Name" : "Phosphotyrosine protein phosphatases superfamily protein",
        "Stress" : 80523,
        "BetweennessCentrality" : 0.01707671,
        "t_RNA_motif_Root" : "NO",
        "EdgeCount" : 72,
        "Causal_PLAMOM" : "no",
        "name" : "AT2G32960",
        "Causal_tRNA" : "no",
        "IsSingleNode" : false,
        "briefDescription" : "Phosphotyrosine protein phosphatases superfamily protein",
        "NeighborhoodConnectivity" : 4.833333333333333
      },
      "position" : {
        "x" : -6728.970703125,
        "y" : 3374.19970703125
      },
      "selected" : false
    }, {
      "data" : {
        "id" : "1157",
        "symbol" : "AT4G22820",
        "Shoot_Descriptions" : "",
        "PLAMOM_Match_Shoot" : "",
        "Causal_TF" : "no",
        "Root_Descriptions" : "A20/AN1-like zinc finger family protein",
        "Eccentricity" : 1,
        "Outdegree" : 2,
        "Target_PLAMOM" : "",
        "PLAMOM_Match_Root" : "NO",
        "shared_name" : "AT4G22820",
        "Indegree" : 0,
        "Target_tRNA" : "",
        "t_RNA_motif_shoot" : "",
        "SUID" : 1157,
        "SelfLoops" : 0,
        "Target_TF" : "",
        "MolecularAttributes" : "protein_coding",
        "AverageShortestPathLength" : 1.0,
        "selected" : false,
        "Name2" : "AT4G22820",
        "ClosenessCentrality" : 1.0,
        "Casual_Name" : "A20/AN1-like zinc finger family protein",
        "PartnerOfMultiEdgedNodePairs" : 0,
        "ClusteringCoefficient" : 0.0,
        "Target_Name" : "",
        "Stress" : 0,
        "BetweennessCentrality" : 0.0,
        "t_RNA_motif_Root" : "NO",
        "EdgeCount" : 2,
        "Causal_PLAMOM" : "no",
        "name" : "AT4G22820",
        "Causal_tRNA" : "no",
        "IsSingleNode" : false,
        "briefDescription" : "A20/AN1-like zinc finger family protein",
        "NeighborhoodConnectivity" : 6.5
      },
      "position" : {
        "x" : -5471.10546875,
        "y" : 370.8018493652344
      },
      "selected" : false
    }, {
      "data" : {
        "id" : "1156",
        "symbol" : "UMAMIT33",
        "Shoot_Descriptions" : "",
        "PLAMOM_Match_Shoot" : "",
        "Causal_TF" : "no",
        "Root_Descriptions" : "nodulin MtN21 /EamA-like transporter family protein",
        "Eccentricity" : 8,
        "Outdegree" : 6,
        "Target_PLAMOM" : "",
        "PLAMOM_Match_Root" : "NO",
        "shared_name" : "AT4G28040",
        "Indegree" : 0,
        "Target_tRNA" : "",
        "t_RNA_motif_shoot" : "",
        "SUID" : 1156,
        "SelfLoops" : 0,
        "Target_TF" : "",
        "MolecularAttributes" : "protein_coding",
        "AverageShortestPathLength" : 4.61875,
        "selected" : false,
        "Name2" : "UMAMIT33",
        "ClosenessCentrality" : 0.2165088,
        "Casual_Name" : "nodulin MtN21 /EamA-like transporter family protein",
        "PartnerOfMultiEdgedNodePairs" : 0,
        "ClusteringCoefficient" : 0.0,
        "Target_Name" : "",
        "Stress" : 0,
        "BetweennessCentrality" : 0.0,
        "t_RNA_motif_Root" : "NO",
        "EdgeCount" : 6,
        "Causal_PLAMOM" : "no",
        "name" : "AT4G28040",
        "Causal_tRNA" : "no",
        "IsSingleNode" : false,
        "briefDescription" : "nodulin MtN21 /EamA-like transporter family protein",
        "NeighborhoodConnectivity" : 19.666666666666668
      },
      "position" : {
        "x" : -5781.02099609375,
        "y" : 155.91990661621094
      },
      "selected" : false
    }, {
      "data" : {
        "id" : "1155",
        "symbol" : "BT5",
        "degree_layout" : 2,
        "Shoot_Descriptions" : "",
        "PLAMOM_Match_Shoot" : "",
        "Causal_TF" : "BTB and TAZ domain protein 5\"",
        "Root_Descriptions" : "BT5, BTB and TAZ domain protein 5",
        "Eccentricity" : 9,
        "Outdegree" : 8,
        "Target_PLAMOM" : "NAC domain containing protein 96\"",
        "PLAMOM_Match_Root" : "NO",
        "shared_name" : "AT4G37610",
        "Indegree" : 3,
        "Target_tRNA" : "AT5G46590",
        "t_RNA_motif_shoot" : "",
        "SUID" : 1155,
        "SelfLoops" : 0,
        "Target_TF" : "no",
        "MolecularAttributes" : "protein_coding",
        "AverageShortestPathLength" : 4.98117155,
        "selected" : false,
        "Name2" : "BT5",
        "ClosenessCentrality" : 0.20075598,
        "Casual_Name" : "\"BT5",
        "PartnerOfMultiEdgedNodePairs" : 0,
        "ClusteringCoefficient" : 0.00909091,
        "Target_Name" : "no",
        "Stress" : 9433,
        "BetweennessCentrality" : 0.00153529,
        "t_RNA_motif_Root" : "NO",
        "EdgeCount" : 11,
        "Causal_PLAMOM" : "no",
        "name" : "AT4G37610",
        "Causal_tRNA" : "no",
        "IsSingleNode" : false,
        "briefDescription" : "BTB and TAZ domain protein 5",
        "NeighborhoodConnectivity" : 10.090909090909092
      },
      "position" : {
        "x" : -6563.05078125,
        "y" : 3309.33056640625
      },
      "selected" : false
    }, {
      "data" : {
        "id" : "1154",
        "symbol" : "ASN2",
        "Shoot_Descriptions" : "",
        "PLAMOM_Match_Shoot" : "",
        "Causal_TF" : "asparagine synthetase 2\"",
        "Root_Descriptions" : "ASN2, asparagine synthetase 2",
        "Eccentricity" : 11,
        "Outdegree" : 3,
        "Target_PLAMOM" : "",
        "PLAMOM_Match_Root" : "YES",
        "shared_name" : "AT5G65010",
        "Indegree" : 0,
        "Target_tRNA" : "",
        "t_RNA_motif_shoot" : "",
        "SUID" : 1154,
        "SelfLoops" : 0,
        "Target_TF" : "",
        "MolecularAttributes" : "METABOLIC",
        "AverageShortestPathLength" : 6.69583333,
        "selected" : false,
        "Name2" : "ASN2",
        "ClosenessCentrality" : 0.14934661,
        "Casual_Name" : "\"ASN2",
        "PartnerOfMultiEdgedNodePairs" : 0,
        "ClusteringCoefficient" : 0.0,
        "Target_Name" : "",
        "Stress" : 0,
        "BetweennessCentrality" : 0.0,
        "t_RNA_motif_Root" : "NO",
        "EdgeCount" : 3,
        "Causal_PLAMOM" : "AT5G65010",
        "name" : "AT5G65010",
        "Causal_tRNA" : "no",
        "IsSingleNode" : false,
        "briefDescription" : "asparagine synthetase 2",
        "NeighborhoodConnectivity" : 11.333333333333334
      },
      "position" : {
        "x" : -5734.85205078125,
        "y" : -156.81642150878906
      },
      "selected" : false
    }, {
      "data" : {
        "id" : "1153",
        "symbol" : "SAG21",
        "Shoot_Descriptions" : "AtLEA5, SAG21, senescence-associated gene 21",
        "PLAMOM_Match_Shoot" : "YES",
        "Causal_TF" : "SAG21",
        "Root_Descriptions" : "AtLEA5, SAG21, senescence-associated gene 21",
        "Eccentricity" : 17,
        "Outdegree" : 8,
        "Target_PLAMOM" : "",
        "PLAMOM_Match_Root" : "YES",
        "shared_name" : "AT4G02380",
        "Indegree" : 0,
        "Target_tRNA" : "",
        "t_RNA_motif_shoot" : "NO",
        "SUID" : 1153,
        "SelfLoops" : 0,
        "Target_TF" : "",
        "MolecularAttributes" : "protein_coding",
        "AverageShortestPathLength" : 11.6097561,
        "selected" : false,
        "Name2" : "SAG21",
        "ClosenessCentrality" : 0.08613445,
        "Casual_Name" : "\"AtLEA5",
        "PartnerOfMultiEdgedNodePairs" : 0,
        "ClusteringCoefficient" : 0.0,
        "Target_Name" : "",
        "Stress" : 0,
        "BetweennessCentrality" : 0.0,
        "t_RNA_motif_Root" : "NO",
        "EdgeCount" : 8,
        "Causal_PLAMOM" : "AT4G02380",
        "name" : "AT4G02380",
        "Causal_tRNA" : "no",
        "IsSingleNode" : false,
        "briefDescription" : "senescence-associated gene 21",
        "NeighborhoodConnectivity" : 7.125
      },
      "position" : {
        "x" : -579.2701907871415,
        "y" : 3512.470947265625
      },
      "selected" : false
    }, {
      "data" : {
        "id" : "1152",
        "symbol" : "AT1G52930",
        "Shoot_Descriptions" : "Ribosomal RNA processing Brix domain protein",
        "PLAMOM_Match_Shoot" : "YES",
        "Causal_TF" : "",
        "Root_Descriptions" : "",
        "Eccentricity" : 0,
        "Outdegree" : 0,
        "Target_PLAMOM" : "senescence-associated gene 21\"",
        "PLAMOM_Match_Root" : "",
        "shared_name" : "AT1G52930",
        "Indegree" : 1,
        "Target_tRNA" : "no",
        "t_RNA_motif_shoot" : "NO",
        "SUID" : 1152,
        "SelfLoops" : 0,
        "Target_TF" : "no",
        "MolecularAttributes" : "protein_coding",
        "AverageShortestPathLength" : 0.0,
        "selected" : false,
        "Name2" : "AT1G52930",
        "ClosenessCentrality" : 0.0,
        "Casual_Name" : "",
        "PartnerOfMultiEdgedNodePairs" : 0,
        "ClusteringCoefficient" : 0.0,
        "Target_Name" : "AT1G52930",
        "Stress" : 0,
        "BetweennessCentrality" : 0.0,
        "t_RNA_motif_Root" : "",
        "EdgeCount" : 1,
        "Causal_PLAMOM" : "",
        "name" : "AT1G52930",
        "Causal_tRNA" : "",
        "IsSingleNode" : false,
        "briefDescription" : "Ribosomal RNA processing Brix domain protein",
        "NeighborhoodConnectivity" : 8.0
      },
      "position" : {
        "x" : 4661.310069021582,
        "y" : 380.41534423828125
      },
      "selected" : false
    }, {
      "data" : {
        "id" : "1151",
        "symbol" : "SPA4",
        "degree_layout" : 1,
        "Shoot_Descriptions" : "SPA4, SPA1-related 4",
        "PLAMOM_Match_Shoot" : "NO",
        "Causal_TF" : "",
        "Root_Descriptions" : "",
        "Eccentricity" : 0,
        "Outdegree" : 0,
        "Target_PLAMOM" : "no",
        "PLAMOM_Match_Root" : "",
        "shared_name" : "AT1G53090",
        "Indegree" : 3,
        "Target_tRNA" : "no",
        "t_RNA_motif_shoot" : "NO",
        "SUID" : 1151,
        "SelfLoops" : 0,
        "Target_TF" : "SPA1-related 4\"",
        "MolecularAttributes" : "protein_coding",
        "AverageShortestPathLength" : 0.0,
        "selected" : false,
        "Name2" : "SPA4",
        "ClosenessCentrality" : 0.0,
        "Casual_Name" : "",
        "PartnerOfMultiEdgedNodePairs" : 0,
        "ClusteringCoefficient" : 0.0,
        "Target_Name" : "\"SPA4",
        "Stress" : 0,
        "BetweennessCentrality" : 0.0,
        "t_RNA_motif_Root" : "",
        "EdgeCount" : 3,
        "Causal_PLAMOM" : "",
        "name" : "AT1G53090",
        "Causal_tRNA" : "",
        "IsSingleNode" : false,
        "briefDescription" : "SPA1-related 4",
        "NeighborhoodConnectivity" : 29.666666666666668
      },
      "position" : {
        "x" : 3931.111278939421,
        "y" : 337.8841247558594
      },
      "selected" : false
    }, {
      "data" : {
        "id" : "1150",
        "symbol" : "AT3G18773",
        "degree_layout" : 2,
        "Shoot_Descriptions" : "",
        "PLAMOM_Match_Shoot" : "",
        "Causal_TF" : "no",
        "Root_Descriptions" : "RING/U-box superfamily protein",
        "Eccentricity" : 1,
        "Outdegree" : 4,
        "Target_PLAMOM" : "no",
        "PLAMOM_Match_Root" : "NO",
        "shared_name" : "AT3G18773",
        "Indegree" : 2,
        "Target_tRNA" : "no",
        "t_RNA_motif_shoot" : "",
        "SUID" : 1150,
        "SelfLoops" : 0,
        "Target_TF" : "no",
        "MolecularAttributes" : "protein_coding",
        "AverageShortestPathLength" : 1.0,
        "selected" : false,
        "Name2" : "AT3G18773",
        "ClosenessCentrality" : 1.0,
        "Casual_Name" : "RING/U-box superfamily protein",
        "PartnerOfMultiEdgedNodePairs" : 0,
        "ClusteringCoefficient" : 0.0,
        "Target_Name" : "RING/U-box superfamily protein",
        "Stress" : 11156,
        "BetweennessCentrality" : 0.00140879,
        "t_RNA_motif_Root" : "NO",
        "EdgeCount" : 6,
        "Causal_PLAMOM" : "no",
        "name" : "AT3G18773",
        "Causal_tRNA" : "no",
        "IsSingleNode" : false,
        "briefDescription" : "RING/U-box superfamily protein",
        "NeighborhoodConnectivity" : 4.333333333333333
      },
      "position" : {
        "x" : -6560.099609375,
        "y" : 2450.251953125
      },
      "selected" : false
    }, {
      "data" : {
        "id" : "1149",
        "symbol" : "AT1G53190",
        "Shoot_Descriptions" : "RING/U-box superfamily protein",
        "PLAMOM_Match_Shoot" : "YES",
        "Causal_TF" : "",
        "Root_Descriptions" : "",
        "Eccentricity" : 0,
        "Outdegree" : 0,
        "Target_PLAMOM" : "AT1G53190",
        "PLAMOM_Match_Root" : "",
        "shared_name" : "AT1G53190",
        "Indegree" : 1,
        "Target_tRNA" : "no",
        "t_RNA_motif_shoot" : "NO",
        "SUID" : 1149,
        "SelfLoops" : 0,
        "Target_TF" : "no",
        "MolecularAttributes" : "protein_coding",
        "AverageShortestPathLength" : 0.0,
        "selected" : false,
        "Name2" : "AT1G53190",
        "ClosenessCentrality" : 0.0,
        "Casual_Name" : "",
        "PartnerOfMultiEdgedNodePairs" : 0,
        "ClusteringCoefficient" : 0.0,
        "Target_Name" : "RING/U-box superfamily protein",
        "Stress" : 0,
        "BetweennessCentrality" : 0.0,
        "t_RNA_motif_Root" : "",
        "EdgeCount" : 1,
        "Causal_PLAMOM" : "",
        "name" : "AT1G53190",
        "Causal_tRNA" : "",
        "IsSingleNode" : false,
        "briefDescription" : "RING/U-box superfamily protein",
        "NeighborhoodConnectivity" : 6.0
      },
      "position" : {
        "x" : 3817.756298470671,
        "y" : -793.99072265625
      },
      "selected" : false
    }, {
      "data" : {
        "id" : "1148",
        "symbol" : "ZFP1",
        "Shoot_Descriptions" : "",
        "PLAMOM_Match_Shoot" : "",
        "Causal_TF" : "ZFP1",
        "Root_Descriptions" : "ATZFP1, ZFP1, zinc-finger protein 1",
        "Eccentricity" : 8,
        "Outdegree" : 7,
        "Target_PLAMOM" : "",
        "PLAMOM_Match_Root" : "NO",
        "shared_name" : "AT1G80730",
        "Indegree" : 0,
        "Target_tRNA" : "",
        "t_RNA_motif_shoot" : "",
        "SUID" : 1148,
        "SelfLoops" : 0,
        "Target_TF" : "",
        "MolecularAttributes" : "TXNFACTOR",
        "AverageShortestPathLength" : 4.775,
        "selected" : false,
        "Name2" : "ZFP1",
        "ClosenessCentrality" : 0.20942408,
        "Casual_Name" : "\"ATZFP1",
        "PartnerOfMultiEdgedNodePairs" : 0,
        "ClusteringCoefficient" : 0.0,
        "Target_Name" : "",
        "Stress" : 0,
        "BetweennessCentrality" : 0.0,
        "t_RNA_motif_Root" : "NO",
        "EdgeCount" : 7,
        "Causal_PLAMOM" : "no",
        "name" : "AT1G80730",
        "Causal_tRNA" : "no",
        "IsSingleNode" : false,
        "briefDescription" : "zinc-finger protein 1",
        "NeighborhoodConnectivity" : 8.428571428571429
      },
      "position" : {
        "x" : -5856.14794921875,
        "y" : 254.74124145507812
      },
      "selected" : false
    }, {
      "data" : {
        "id" : "1147",
        "symbol" : "GLY3",
        "Shoot_Descriptions" : "ETHE1, GLX2-3, GLY3, glyoxalase II  3",
        "PLAMOM_Match_Shoot" : "NO",
        "Causal_TF" : "GLX2-3",
        "Root_Descriptions" : "",
        "Eccentricity" : 9,
        "Outdegree" : 2,
        "Target_PLAMOM" : "zinc-finger protein 1\"",
        "PLAMOM_Match_Root" : "",
        "shared_name" : "AT1G53580",
        "Indegree" : 1,
        "Target_tRNA" : "AT1G80730",
        "t_RNA_motif_shoot" : "NO",
        "SUID" : 1147,
        "SelfLoops" : 0,
        "Target_TF" : "no",
        "MolecularAttributes" : "METABOLIC",
        "AverageShortestPathLength" : 4.73068894,
        "selected" : false,
        "Name2" : "GLY3",
        "ClosenessCentrality" : 0.2113857,
        "Casual_Name" : "\"ETHE1",
        "PartnerOfMultiEdgedNodePairs" : 0,
        "ClusteringCoefficient" : 0.0,
        "Target_Name" : "no",
        "Stress" : 349,
        "BetweennessCentrality" : 8.099E-5,
        "t_RNA_motif_Root" : "",
        "EdgeCount" : 3,
        "Causal_PLAMOM" : "no",
        "name" : "AT1G53580",
        "Causal_tRNA" : "no",
        "IsSingleNode" : false,
        "briefDescription" : "glyoxalase II 3",
        "NeighborhoodConnectivity" : 26.666666666666668
      },
      "position" : {
        "x" : 3464.047314095671,
        "y" : 3155.0107421875
      },
      "selected" : false
    }, {
      "data" : {
        "id" : "1146",
        "symbol" : "AT1G53760",
        "Shoot_Descriptions" : "unknown protein; FUNCTIONS IN: molecular_function unknown; INVOLVED IN: biological_process unknown; LOCATED IN: mitochondrion, plasma membrane, plastid, membrane; EXPRESSED IN: 25 plant structures; EXPRESSED DURING: 13 growth stages; CONTAINS InterPro DOMAIN/s: Protein of unknown function DUF2343 (InterPro:IPR018786); Has 171 Blast hits to 171 proteins in 90 species: Archae - 0; Bacteria - 0; Metazoa - 0; Fungi - 127; Plants - 41; Viruses - 0; Other Eukaryotes - 3 (source: NCBI BLink).",
        "PLAMOM_Match_Shoot" : "NO",
        "Causal_TF" : "plasma membrane",
        "Root_Descriptions" : "",
        "Eccentricity" : 12,
        "Outdegree" : 1,
        "Target_PLAMOM" : "ATGLB2",
        "PLAMOM_Match_Root" : "",
        "shared_name" : "AT1G53760",
        "Indegree" : 1,
        "Target_tRNA" : "GLB2",
        "t_RNA_motif_shoot" : "YES",
        "SUID" : 1146,
        "SelfLoops" : 0,
        "Target_TF" : "NSHB2",
        "MolecularAttributes" : "protein_coding",
        "AverageShortestPathLength" : 7.68410042,
        "selected" : false,
        "Name2" : "AT1G53760",
        "ClosenessCentrality" : 0.13013885,
        "Casual_Name" : "\"unknown protein; FUNCTIONS IN: molecular_function unknown; INVOLVED IN: biological_process unknown; LOCATED IN: mitochondrion",
        "PartnerOfMultiEdgedNodePairs" : 0,
        "ClusteringCoefficient" : 0.0,
        "Target_Name" : "HB2",
        "Stress" : 17,
        "BetweennessCentrality" : 2.7E-6,
        "t_RNA_motif_Root" : "",
        "EdgeCount" : 2,
        "Causal_PLAMOM" : "no",
        "name" : "AT1G53760",
        "Causal_tRNA" : "AT1G53760",
        "IsSingleNode" : false,
        "briefDescription" : "K -H  exchange-like protein",
        "NeighborhoodConnectivity" : 17.0
      },
      "position" : {
        "x" : 3550.53466796875,
        "y" : 2565.56298828125
      },
      "selected" : false
    }, {
      "data" : {
        "id" : "1145",
        "symbol" : "GAMMA-H2AX",
        "Shoot_Descriptions" : "G-H2AX, GAMMA-H2AX, H2AXB, HTA3, gamma histone variant H2AX",
        "PLAMOM_Match_Shoot" : "YES",
        "Causal_TF" : "GAMMA-H2AX",
        "Root_Descriptions" : "G-H2AX, GAMMA-H2AX, H2AXB, HTA3, gamma histone variant H2AX",
        "Eccentricity" : 1,
        "Outdegree" : 5,
        "Target_PLAMOM" : "nudix hydrolase homolog 18\"",
        "PLAMOM_Match_Root" : "YES",
        "shared_name" : "AT1G54690",
        "Indegree" : 2,
        "Target_tRNA" : "no",
        "t_RNA_motif_shoot" : "NO",
        "SUID" : 1145,
        "SelfLoops" : 0,
        "Target_TF" : "no",
        "MolecularAttributes" : "protein_coding",
        "AverageShortestPathLength" : 1.0,
        "selected" : false,
        "Name2" : "GAMMA-H2AX",
        "ClosenessCentrality" : 1.0,
        "Casual_Name" : "\"G-H2AX",
        "PartnerOfMultiEdgedNodePairs" : 0,
        "ClusteringCoefficient" : 0.02380952,
        "Target_Name" : "AT1G54690",
        "Stress" : 5366,
        "BetweennessCentrality" : 0.00147561,
        "t_RNA_motif_Root" : "NO",
        "EdgeCount" : 7,
        "Causal_PLAMOM" : "AT1G54690",
        "name" : "AT1G54690",
        "Causal_tRNA" : "no",
        "IsSingleNode" : false,
        "briefDescription" : "gamma histone variant H2AX",
        "NeighborhoodConnectivity" : 9.142857142857142
      },
      "position" : {
        "x" : -373.378467154329,
        "y" : -209.30239868164062
      },
      "selected" : false
    }, {
      "data" : {
        "id" : "1144",
        "symbol" : "RSH3",
        "Shoot_Descriptions" : "AT-RSH3, ATRSH3, RSH3, RELA/SPOT homolog 3",
        "PLAMOM_Match_Shoot" : "YES",
        "Causal_TF" : "",
        "Root_Descriptions" : "",
        "Eccentricity" : 0,
        "Outdegree" : 0,
        "Target_PLAMOM" : "H2AXB",
        "PLAMOM_Match_Root" : "",
        "shared_name" : "AT1G54130",
        "Indegree" : 1,
        "Target_tRNA" : "HTA3",
        "t_RNA_motif_shoot" : "NO",
        "SUID" : 1144,
        "SelfLoops" : 0,
        "Target_TF" : "no",
        "MolecularAttributes" : "protein_coding",
        "AverageShortestPathLength" : 0.0,
        "selected" : false,
        "Name2" : "RSH3",
        "ClosenessCentrality" : 0.0,
        "Casual_Name" : "",
        "PartnerOfMultiEdgedNodePairs" : 0,
        "ClusteringCoefficient" : 0.0,
        "Target_Name" : "gamma histone variant H2AX\"",
        "Stress" : 0,
        "BetweennessCentrality" : 0.0,
        "t_RNA_motif_Root" : "",
        "EdgeCount" : 1,
        "Causal_PLAMOM" : "",
        "name" : "AT1G54130",
        "Causal_tRNA" : "",
        "IsSingleNode" : false,
        "briefDescription" : "RELA/SPOT homolog 3",
        "NeighborhoodConnectivity" : 7.0
      },
      "position" : {
        "x" : 4237.837901052832,
        "y" : -695.41259765625
      },
      "selected" : false
    }, {
      "data" : {
        "id" : "1143",
        "symbol" : "TAFII21",
        "Shoot_Descriptions" : "TAF9, TAFII21, TATA binding protein associated factor 21kDa subunit",
        "PLAMOM_Match_Shoot" : "NO",
        "Causal_TF" : "",
        "Root_Descriptions" : "",
        "Eccentricity" : 0,
        "Outdegree" : 0,
        "Target_PLAMOM" : "no",
        "PLAMOM_Match_Root" : "",
        "shared_name" : "AT1G54140",
        "Indegree" : 3,
        "Target_tRNA" : "no",
        "t_RNA_motif_shoot" : "NO",
        "SUID" : 1143,
        "SelfLoops" : 0,
        "Target_TF" : "TAFII21",
        "MolecularAttributes" : "protein_coding",
        "AverageShortestPathLength" : 0.0,
        "selected" : false,
        "Name2" : "TAFII21",
        "ClosenessCentrality" : 0.0,
        "Casual_Name" : "",
        "PartnerOfMultiEdgedNodePairs" : 0,
        "ClusteringCoefficient" : 0.0,
        "Target_Name" : "\"TAF9",
        "Stress" : 0,
        "BetweennessCentrality" : 0.0,
        "t_RNA_motif_Root" : "",
        "EdgeCount" : 3,
        "Causal_PLAMOM" : "",
        "name" : "AT1G54140",
        "Causal_tRNA" : "",
        "IsSingleNode" : false,
        "briefDescription" : "TATA binding protein associated factor 21kDa subunit",
        "NeighborhoodConnectivity" : 7.333333333333333
      },
      "position" : {
        "x" : 4517.780224251921,
        "y" : 73.24890899658203
      },
      "selected" : false
    }, {
      "data" : {
        "id" : "1142",
        "symbol" : "AT5G64550",
        "degree_layout" : 8,
        "Shoot_Descriptions" : "",
        "PLAMOM_Match_Shoot" : "",
        "Causal_TF" : "no",
        "Root_Descriptions" : "loricrin-related",
        "Eccentricity" : 10,
        "Outdegree" : 4,
        "Target_PLAMOM" : "no",
        "PLAMOM_Match_Root" : "NO",
        "shared_name" : "AT5G64550",
        "Indegree" : 9,
        "Target_tRNA" : "AT5G64550",
        "t_RNA_motif_shoot" : "",
        "SUID" : 1142,
        "SelfLoops" : 0,
        "Target_TF" : "no",
        "MolecularAttributes" : "protein_coding",
        "AverageShortestPathLength" : 5.80334728,
        "selected" : false,
        "Name2" : "AT5G64550",
        "ClosenessCentrality" : 0.17231435,
        "Casual_Name" : "loricrin-related",
        "PartnerOfMultiEdgedNodePairs" : 0,
        "ClusteringCoefficient" : 0.0,
        "Target_Name" : "loricrin-related",
        "Stress" : 22884,
        "BetweennessCentrality" : 0.00532333,
        "t_RNA_motif_Root" : "YES",
        "EdgeCount" : 13,
        "Causal_PLAMOM" : "no",
        "name" : "AT5G64550",
        "Causal_tRNA" : "AT5G64550",
        "IsSingleNode" : false,
        "briefDescription" : "loricrin-like protein",
        "NeighborhoodConnectivity" : 7.6923076923076925
      },
      "position" : {
        "x" : -6334.4345703125,
        "y" : 3040.224853515625
      },
      "selected" : false
    }, {
      "data" : {
        "id" : "1141",
        "symbol" : "AT1G54380",
        "Shoot_Descriptions" : "spliceosome protein-related",
        "PLAMOM_Match_Shoot" : "NO",
        "Causal_TF" : "",
        "Root_Descriptions" : "",
        "Eccentricity" : 0,
        "Outdegree" : 0,
        "Target_PLAMOM" : "no",
        "PLAMOM_Match_Root" : "",
        "shared_name" : "AT1G54380",
        "Indegree" : 1,
        "Target_tRNA" : "no",
        "t_RNA_motif_shoot" : "NO",
        "SUID" : 1141,
        "SelfLoops" : 0,
        "Target_TF" : "no",
        "MolecularAttributes" : "protein_coding",
        "AverageShortestPathLength" : 0.0,
        "selected" : false,
        "Name2" : "AT1G54380",
        "ClosenessCentrality" : 0.0,
        "Casual_Name" : "",
        "PartnerOfMultiEdgedNodePairs" : 0,
        "ClusteringCoefficient" : 0.0,
        "Target_Name" : "spliceosome protein-related",
        "Stress" : 0,
        "BetweennessCentrality" : 0.0,
        "t_RNA_motif_Root" : "",
        "EdgeCount" : 1,
        "Causal_PLAMOM" : "",
        "name" : "AT1G54380",
        "Causal_tRNA" : "",
        "IsSingleNode" : false,
        "briefDescription" : "spliceosome protein-like protein",
        "NeighborhoodConnectivity" : 5.0
      },
      "position" : {
        "x" : 3898.813671517546,
        "y" : -664.0924072265625
      },
      "selected" : false
    }, {
      "data" : {
        "id" : "1140",
        "symbol" : "AT1G49500",
        "Shoot_Descriptions" : "",
        "PLAMOM_Match_Shoot" : "",
        "Causal_TF" : "no",
        "Root_Descriptions" : "unknown protein; FUNCTIONS IN: molecular_function unknown; INVOLVED IN: biological_process unknown; LOCATED IN: endomembrane system; EXPRESSED IN: 19 plant structures; EXPRESSED DURING: 10 growth stages; BEST Arabidopsis thaliana protein match is: unknown protein (TAIR:AT3G19030.1); Has 24 Blast hits to 24 proteins in 2 species: Archae - 0; Bacteria - 0; Metazoa - 0; Fungi - 0; Plants - 24; Viruses - 0; Other Eukaryotes - 0 (source: NCBI BLink).",
        "Eccentricity" : 14,
        "Outdegree" : 2,
        "Target_PLAMOM" : "",
        "PLAMOM_Match_Root" : "YES",
        "shared_name" : "AT1G49500",
        "Indegree" : 0,
        "Target_tRNA" : "",
        "t_RNA_motif_shoot" : "",
        "SUID" : 1140,
        "SelfLoops" : 0,
        "Target_TF" : "",
        "MolecularAttributes" : "protein_coding",
        "AverageShortestPathLength" : 8.82952183,
        "selected" : false,
        "Name2" : "AT1G49500",
        "ClosenessCentrality" : 0.11325642,
        "Casual_Name" : "unknown protein; FUNCTIONS IN: molecular_function unknown; INVOLVED IN: biological_process unknown; LOCATED IN: endomembrane system; EXPRESSED IN: 19 plant structures; EXPRESSED DURING: 10 growth stages; BEST Arabidopsis thaliana protein match is: unknown protein (TAIR:AT3G19030.1); Has 24 Blast hits to 24 proteins in 2 species: Archae - 0; Bacteria - 0; Metazoa - 0; Fungi - 0; Plants - 24; Viruses - 0; Other Eukaryotes - 0 (source: NCBI BLink).",
        "PartnerOfMultiEdgedNodePairs" : 0,
        "ClusteringCoefficient" : 0.0,
        "Target_Name" : "",
        "Stress" : 0,
        "BetweennessCentrality" : 0.0,
        "t_RNA_motif_Root" : "YES",
        "EdgeCount" : 2,
        "Causal_PLAMOM" : "AT1G49500",
        "name" : "AT1G49500",
        "Causal_tRNA" : "AT1G49500",
        "IsSingleNode" : false,
        "briefDescription" : "transcription initiation factor TFIID subunit 1b-like protein",
        "NeighborhoodConnectivity" : 6.0
      },
      "position" : {
        "x" : -5492.48779296875,
        "y" : -366.09765625
      },
      "selected" : false
    }, {
      "data" : {
        "id" : "1139",
        "symbol" : "ERF5",
        "degree_layout" : 2,
        "Shoot_Descriptions" : "",
        "PLAMOM_Match_Shoot" : "",
        "Causal_TF" : "ATERF5",
        "Root_Descriptions" : "ATERF-5, ATERF5, ERF5, ethylene responsive element binding factor 5",
        "Eccentricity" : 9,
        "Outdegree" : 12,
        "Target_PLAMOM" : "NAC (No Apical Meristem) domain transcriptional regulator superfamily protein\"",
        "PLAMOM_Match_Root" : "YES",
        "shared_name" : "AT5G47230",
        "Indegree" : 3,
        "Target_tRNA" : "AT4G27410",
        "t_RNA_motif_shoot" : "",
        "SUID" : 1139,
        "SelfLoops" : 0,
        "Target_TF" : "no",
        "MolecularAttributes" : "TXNFACTOR",
        "AverageShortestPathLength" : 4.52301255,
        "selected" : false,
        "Name2" : "ERF5",
        "ClosenessCentrality" : 0.22109158,
        "Casual_Name" : "\"ATERF-5",
        "PartnerOfMultiEdgedNodePairs" : 0,
        "ClusteringCoefficient" : 0.0,
        "Target_Name" : "AT5G47230",
        "Stress" : 28836,
        "BetweennessCentrality" : 0.00577637,
        "t_RNA_motif_Root" : "NO",
        "EdgeCount" : 15,
        "Causal_PLAMOM" : "AT5G47230",
        "name" : "AT5G47230",
        "Causal_tRNA" : "no",
        "IsSingleNode" : false,
        "briefDescription" : "ethylene responsive element binding factor 5",
        "NeighborhoodConnectivity" : 6.133333333333334
      },
      "position" : {
        "x" : -6339.6728515625,
        "y" : 3352.49853515625
      },
      "selected" : false
    }, {
      "data" : {
        "id" : "1138",
        "symbol" : "TMN6",
        "Shoot_Descriptions" : "AtTMN6, TMN6, Endomembrane protein 70 protein family",
        "PLAMOM_Match_Shoot" : "NO",
        "Causal_TF" : "TMN6",
        "Root_Descriptions" : "",
        "Eccentricity" : 8,
        "Outdegree" : 8,
        "Target_PLAMOM" : "metacaspase 8\"",
        "PLAMOM_Match_Root" : "",
        "shared_name" : "AT1G55130",
        "Indegree" : 2,
        "Target_tRNA" : "no",
        "t_RNA_motif_shoot" : "NO",
        "SUID" : 1138,
        "SelfLoops" : 0,
        "Target_TF" : "no",
        "MolecularAttributes" : "protein_coding",
        "AverageShortestPathLength" : 3.94769874,
        "selected" : false,
        "Name2" : "TMN6",
        "ClosenessCentrality" : 0.25331214,
        "Casual_Name" : "\"AtTMN6",
        "PartnerOfMultiEdgedNodePairs" : 0,
        "ClusteringCoefficient" : 0.0,
        "Target_Name" : "no",
        "Stress" : 14945,
        "BetweennessCentrality" : 0.00337228,
        "t_RNA_motif_Root" : "",
        "EdgeCount" : 10,
        "Causal_PLAMOM" : "no",
        "name" : "AT1G55130",
        "Causal_tRNA" : "no",
        "IsSingleNode" : false,
        "briefDescription" : "Endomembrane protein 70 protein family",
        "NeighborhoodConnectivity" : 24.1
      },
      "position" : {
        "x" : 3872.904491830046,
        "y" : 3229.1142578125
      },
      "selected" : false
    }, {
      "data" : {
        "id" : "1137",
        "symbol" : "AT3G22550",
        "Shoot_Descriptions" : "",
        "PLAMOM_Match_Shoot" : "",
        "Causal_TF" : "no",
        "Root_Descriptions" : "Protein of unknown function (DUF581)",
        "Eccentricity" : 1,
        "Outdegree" : 2,
        "Target_PLAMOM" : "no",
        "PLAMOM_Match_Root" : "NO",
        "shared_name" : "AT3G22550",
        "Indegree" : 2,
        "Target_tRNA" : "no",
        "t_RNA_motif_shoot" : "",
        "SUID" : 1137,
        "SelfLoops" : 0,
        "Target_TF" : "no",
        "MolecularAttributes" : "protein_coding",
        "AverageShortestPathLength" : 1.0,
        "selected" : false,
        "Name2" : "AT3G22550",
        "ClosenessCentrality" : 1.0,
        "Casual_Name" : "Protein of unknown function (DUF581)",
        "PartnerOfMultiEdgedNodePairs" : 0,
        "ClusteringCoefficient" : 0.0,
        "Target_Name" : "Protein of unknown function (DUF581)",
        "Stress" : 4,
        "BetweennessCentrality" : 2.09E-6,
        "t_RNA_motif_Root" : "NO",
        "EdgeCount" : 4,
        "Causal_PLAMOM" : "no",
        "name" : "AT3G22550",
        "Causal_tRNA" : "no",
        "IsSingleNode" : false,
        "briefDescription" : "NAD(P)H-quinone oxidoreductase subunit",
        "NeighborhoodConnectivity" : 6.0
      },
      "position" : {
        "x" : -6939.060546875,
        "y" : 3882.147705078125
      },
      "selected" : false
    }, {
      "data" : {
        "id" : "1136",
        "symbol" : "AT1G55820",
        "Shoot_Descriptions" : "Kinase-related protein of unknown function (DUF1296)",
        "PLAMOM_Match_Shoot" : "NO",
        "Causal_TF" : "",
        "Root_Descriptions" : "",
        "Eccentricity" : 0,
        "Outdegree" : 0,
        "Target_PLAMOM" : "PSK4",
        "PLAMOM_Match_Root" : "",
        "shared_name" : "AT1G55820",
        "Indegree" : 2,
        "Target_tRNA" : "phytosulfokine 4 precursor\"",
        "t_RNA_motif_shoot" : "NO",
        "SUID" : 1136,
        "SelfLoops" : 0,
        "Target_TF" : "no",
        "MolecularAttributes" : "protein_coding",
        "AverageShortestPathLength" : 0.0,
        "selected" : false,
        "Name2" : "AT1G55820",
        "ClosenessCentrality" : 0.0,
        "Casual_Name" : "",
        "PartnerOfMultiEdgedNodePairs" : 0,
        "ClusteringCoefficient" : 0.0,
        "Target_Name" : "no",
        "Stress" : 0,
        "BetweennessCentrality" : 0.0,
        "t_RNA_motif_Root" : "",
        "EdgeCount" : 2,
        "Causal_PLAMOM" : "",
        "name" : "AT1G55820",
        "Causal_tRNA" : "",
        "IsSingleNode" : false,
        "briefDescription" : "lysine-specific demethylase",
        "NeighborhoodConnectivity" : 5.0
      },
      "position" : {
        "x" : 3509.414989876921,
        "y" : 1059.6131591796875
      },
      "selected" : false
    }, {
      "data" : {
        "id" : "1135",
        "symbol" : "PSK4",
        "Shoot_Descriptions" : "",
        "PLAMOM_Match_Shoot" : "",
        "Causal_TF" : "ATPSK4",
        "Root_Descriptions" : "ATPSK3 (FORMER SYMBOL), ATPSK4, PSK4, phytosulfokine 4 precursor",
        "Eccentricity" : 12,
        "Outdegree" : 5,
        "Target_PLAMOM" : "no",
        "PLAMOM_Match_Root" : "NO",
        "shared_name" : "AT3G49780",
        "Indegree" : 1,
        "Target_tRNA" : "no",
        "t_RNA_motif_shoot" : "",
        "SUID" : 1135,
        "SelfLoops" : 0,
        "Target_TF" : "ATPSK4",
        "MolecularAttributes" : "protein_coding",
        "AverageShortestPathLength" : 7.56790123,
        "selected" : false,
        "Name2" : "PSK4",
        "ClosenessCentrality" : 0.13213703,
        "Casual_Name" : "\"ATPSK3 (FORMER SYMBOL)",
        "PartnerOfMultiEdgedNodePairs" : 0,
        "ClusteringCoefficient" : 0.0,
        "Target_Name" : "\"ATPSK3 (FORMER SYMBOL)",
        "Stress" : 967,
        "BetweennessCentrality" : 2.9002E-4,
        "t_RNA_motif_Root" : "NO",
        "EdgeCount" : 6,
        "Causal_PLAMOM" : "no",
        "name" : "AT3G49780",
        "Causal_tRNA" : "no",
        "IsSingleNode" : false,
        "briefDescription" : "phytosulfokine 4 precursor",
        "NeighborhoodConnectivity" : 4.5
      },
      "position" : {
        "x" : -7011.7431640625,
        "y" : 3837.571044921875
      },
      "selected" : false
    }, {
      "data" : {
        "id" : "1134",
        "symbol" : "CSLE1",
        "Shoot_Descriptions" : "ATCSLE1, CSLE1, cellulose synthase like E1",
        "PLAMOM_Match_Shoot" : "YES",
        "Causal_TF" : "CSLE1",
        "Root_Descriptions" : "",
        "Eccentricity" : 9,
        "Outdegree" : 5,
        "Target_PLAMOM" : "phloem protein 2-A9\"",
        "PLAMOM_Match_Root" : "",
        "shared_name" : "AT1G55850",
        "Indegree" : 2,
        "Target_tRNA" : "no",
        "t_RNA_motif_shoot" : "YES",
        "SUID" : 1134,
        "SelfLoops" : 0,
        "Target_TF" : "AT1G55850",
        "MolecularAttributes" : "METABOLIC",
        "AverageShortestPathLength" : 4.61087866,
        "selected" : false,
        "Name2" : "CSLE1",
        "ClosenessCentrality" : 0.2168784,
        "Casual_Name" : "\"ATCSLE1",
        "PartnerOfMultiEdgedNodePairs" : 0,
        "ClusteringCoefficient" : 0.0,
        "Target_Name" : "AT1G55850",
        "Stress" : 3120,
        "BetweennessCentrality" : 4.7704E-4,
        "t_RNA_motif_Root" : "",
        "EdgeCount" : 7,
        "Causal_PLAMOM" : "AT1G55850",
        "name" : "AT1G55850",
        "Causal_tRNA" : "AT1G55850",
        "IsSingleNode" : false,
        "briefDescription" : "cellulose synthase like E1",
        "NeighborhoodConnectivity" : 31.714285714285715
      },
      "position" : {
        "x" : 3875.750194955046,
        "y" : 2898.589111328125
      },
      "selected" : false
    }, {
      "data" : {
        "id" : "1133",
        "symbol" : "PP2-A9",
        "Shoot_Descriptions" : "",
```
