## Supplemental Network for "Granger-Causal Testing for Irregularly Sampled Time Series with Application to Nitrogen Signaling in Arabidopsis": styles.js

```
var styles = [ {
  "format_version" : "1.0",
  "generated_by" : "cytoscape-3.7.2",
  "target_cytoscapejs_version" : "~2.1",
  "title" : "Diffany - Differential",
  "style" : [ {
    "selector" : "node",
    "css" : {
      "text-opacity" : 1.0,
      "background-opacity" : 1.0,
      "border-width" : 0.0,
      "color" : "rgb(0,0,0)",
      "border-color" : "rgb(0,0,0)",
      "font-size" : 11,
      "font-family" : "SansSerif.plain",
      "font-weight" : "normal",
      "text-valign" : "center",
      "text-halign" : "center",
      "border-opacity" : 1.0,
      "shape" : "ellipse",
      "width" : 80.0,
      "background-color" : "rgb(128,128,128)",
      "height" : 55.0,
      "content" : "data(official_symbol)"
    }
  }, {
    "selector" : "node[differentially_expressed = 'up-regulated']",
    "css" : {
      "background-color" : "rgb(124,124,7)"
    }
  }, {
    "selector" : "node[differentially_expressed = 'down-regulated']",
    "css" : {
      "background-color" : "rgb(30,88,124)"
    }
  }, {
    "selector" : "node[kinase_function = 'yes']",
    "css" : {
      "border-width" : 2.0
    }
  }, {
    "selector" : "node[differentially_expressed = 'up-regulated']",
    "css" : {
      "background-color" : "rgb(255,255,15)"
    }
  }, {
    "selector" : "node[differentially_expressed = 'down-regulated']",
    "css" : {
      "background-color" : "rgb(62,181,255)"
    }
  }, {
    "selector" : "node[phosphorylation_site = 'yes']",
    "css" : {
      "shape" : "diamond"
    }
  }, {
    "selector" : "node[kinase_function = 'yes']",
    "css" : {
      "border-color" : "rgb(0,0,0)"
    }
  }, {
    "selector" : "node:selected",
    "css" : {
      "background-color" : "rgb(62,62,62)"
    }
  }, {
    "selector" : "edge",
    "css" : {
      "font-size" : 10,
      "line-color" : "rgb(64,64,64)",
      "opacity" : 1.0,
      "source-arrow-shape" : "none",
      "target-arrow-color" : "rgb(0,0,0)",
      "text-opacity" : 1.0,
      "line-style" : "solid",
      "color" : "rgb(0,0,0)",
      "font-family" : "SansSerif.plain",
      "font-weight" : "normal",
      "content" : "",
      "source-arrow-color" : "rgb(0,0,0)",
      "target-arrow-shape" : "none",
      "width" : 1.0
    }
  }, {
    "selector" : "edge[negated]",
    "css" : {
      "line-style" : "dotted"
    }
  }, {
    "selector" : "edge[!negated]",
    "css" : {
      "line-style" : "solid"
    }
  }, {
    "selector" : "edge[weight > 20]",
    "css" : {
      "width" : 25.0
    }
  }, {
    "selector" : "edge[weight = 20]",
    "css" : {
      "width" : 25.0
    }
  }, {
    "selector" : "edge[weight > 0][weight < 20]",
    "css" : {
      "width" : "mapData(weight,0,20,1.0,25.0)"
    }
  }, {
    "selector" : "edge[weight = 0]",
    "css" : {
      "width" : 1.0
    }
  }, {
    "selector" : "edge[weight < 0]",
    "css" : {
      "width" : 1.0
    }
  }, {
    "selector" : "edge:selected",
    "css" : {
      "line-color" : "rgb(255,0,0)"
    }
  } ]
}, {
  "format_version" : "1.0",
  "generated_by" : "cytoscape-3.7.2",
  "target_cytoscapejs_version" : "~2.1",
  "title" : "Marquee",
  "style" : [ {
    "selector" : "node",
    "css" : {
      "text-opacity" : 1.0,
      "background-opacity" : 1.0,
      "border-width" : 10.0,
      "color" : "rgb(102,102,102)",
      "border-color" : "rgb(255,255,255)",
      "font-size" : 12,
      "font-family" : "Dialog.plain",
      "font-weight" : "normal",
      "text-valign" : "bottom",
      "text-halign" : "center",
      "border-opacity" : 1.0,
      "shape" : "ellipse",
      "width" : 20.0,
      "background-color" : "rgb(0,204,255)",
      "height" : 20.0,
      "content" : "data(name)"
    }
  }, {
    "selector" : "node:selected",
    "css" : {
      "background-color" : "rgb(255,0,102)"
    }
  }, {
    "selector" : "edge",
    "css" : {
      "font-size" : 8,
      "line-color" : "rgb(255,255,255)",
      "opacity" : 1.0,
      "source-arrow-shape" : "none",
      "target-arrow-color" : "rgb(255,255,255)",
      "text-opacity" : 1.0,
      "line-style" : "dashed",
      "color" : "rgb(102,102,102)",
      "font-family" : "Dialog.plain",
      "font-weight" : "normal",
      "source-arrow-color" : "rgb(255,255,255)",
      "target-arrow-shape" : "triangle",
      "width" : 2.0,
      "content" : "data(interaction)"
    }
  }, {
    "selector" : "edge:selected",
    "css" : {
      "line-color" : "rgb(255,0,0)"
    }
  } ]
}, {
  "format_version" : "1.0",
  "generated_by" : "cytoscape-3.7.2",
  "target_cytoscapejs_version" : "~2.1",
  "title" : "size_rank_0_0_0_0",
  "style" : [ {
    "selector" : "node",
    "css" : {
      "text-opacity" : 1.0,
      "background-opacity" : 1.0,
      "border-width" : 0.0,
      "color" : "rgb(51,51,51)",
      "border-color" : "rgb(0,0,0)",
      "font-size" : 9,
      "font-family" : "Dialog.plain",
      "font-weight" : "normal",
      "text-valign" : "center",
      "text-halign" : "center",
      "border-opacity" : 1.0,
      "shape" : "rectangle",
      "width" : 12.0,
      "background-color" : "rgb(204,204,255)",
      "height" : 12.0,
      "content" : "data(name)"
    }
  }, {
    "selector" : "node:selected",
    "css" : {
      "background-color" : "rgb(255,255,0)"
    }
  }, {
    "selector" : "edge",
    "css" : {
      "font-size" : 10,
      "line-color" : "rgb(76,76,76)",
      "opacity" : 1.0,
      "source-arrow-shape" : "none",
      "target-arrow-color" : "rgb(0,0,0)",
      "text-opacity" : 1.0,
      "line-style" : "solid",
      "color" : "rgb(0,0,0)",
      "font-family" : "Dialog.plain",
      "font-weight" : "normal",
      "content" : "",
      "source-arrow-color" : "rgb(0,0,0)",
      "target-arrow-shape" : "none",
      "width" : 2.0
    }
  }, {
    "selector" : "edge:selected",
    "css" : {
      "line-color" : "rgb(255,0,0)"
    }
  } ]
}, {
  "format_version" : "1.0",
  "generated_by" : "cytoscape-3.7.2",
  "target_cytoscapejs_version" : "~2.1",
  "title" : "Minimal",
  "style" : [ {
    "selector" : "node",
    "css" : {
      "text-opacity" : 1.0,
      "background-opacity" : 1.0,
      "border-width" : 0.0,
      "color" : "rgb(51,51,51)",
      "border-color" : "rgb(0,0,0)",
      "font-size" : 9,
      "font-family" : "Dialog.plain",
      "font-weight" : "normal",
      "text-valign" : "center",
      "text-halign" : "center",
      "border-opacity" : 1.0,
      "shape" : "rectangle",
      "width" : 42.0,
      "background-color" : "rgb(255,255,255)",
      "height" : 42.0,
      "content" : "data(name)"
    }
  }, {
    "selector" : "node:selected",
    "css" : {
      "background-color" : "rgb(255,255,0)"
    }
  }, {
    "selector" : "edge",
    "css" : {
      "font-size" : 10,
      "line-color" : "rgb(76,76,76)",
      "opacity" : 1.0,
      "source-arrow-shape" : "none",
      "target-arrow-color" : "rgb(0,0,0)",
      "text-opacity" : 1.0,
      "line-style" : "solid",
      "color" : "rgb(0,0,0)",
      "font-family" : "Dialog.plain",
      "font-weight" : "normal",
      "content" : "",
      "source-arrow-color" : "rgb(0,0,0)",
      "target-arrow-shape" : "none",
      "width" : 2.0
    }
  }, {
    "selector" : "edge:selected",
    "css" : {
      "line-color" : "rgb(255,0,0)"
    }
  } ]
}, {
  "format_version" : "1.0",
  "generated_by" : "cytoscape-3.7.2",
  "target_cytoscapejs_version" : "~2.1",
  "title" : "Big Labels",
  "style" : [ {
    "selector" : "node",
    "css" : {
      "text-opacity" : 1.0,
      "background-opacity" : 1.0,
      "border-width" : 0.0,
      "color" : "rgb(51,51,51)",
      "border-color" : "rgb(0,0,0)",
      "font-size" : 24,
      "font-family" : "Dialog.plain",
      "font-weight" : "normal",
      "text-valign" : "center",
      "text-halign" : "center",
      "border-opacity" : 1.0,
      "shape" : "ellipse",
      "width" : 5.0,
      "background-color" : "rgb(255,255,255)",
      "height" : 5.0,
      "content" : "data(name)"
    }
  }, {
    "selector" : "node:selected",
    "css" : {
      "background-color" : "rgb(255,0,102)"
    }
  }, {
    "selector" : "edge",
    "css" : {
      "font-size" : 10,
      "line-color" : "rgb(183,183,183)",
      "opacity" : 1.0,
      "source-arrow-shape" : "none",
      "target-arrow-color" : "rgb(0,0,0)",
      "text-opacity" : 1.0,
      "line-style" : "solid",
      "color" : "rgb(0,0,0)",
      "font-family" : "Dialog.plain",
      "font-weight" : "normal",
      "content" : "",
      "source-arrow-color" : "rgb(0,0,0)",
      "target-arrow-shape" : "none",
      "width" : 1.0
    }
  }, {
    "selector" : "edge:selected",
    "css" : {
      "line-color" : "rgb(255,0,0)"
    }
  } ]
}, {
  "format_version" : "1.0",
  "generated_by" : "cytoscape-3.7.2",
  "target_cytoscapejs_version" : "~2.1",
  "title" : "Directed",
  "style" : [ {
    "selector" : "node",
    "css" : {
      "text-opacity" : 1.0,
      "background-opacity" : 1.0,
      "border-width" : 5.0,
      "color" : "rgb(51,153,255)",
      "border-color" : "rgb(145,145,145)",
      "font-size" : 8,
      "font-family" : "Dialog.plain",
      "font-weight" : "normal",
      "text-valign" : "center",
      "text-halign" : "center",
      "border-opacity" : 1.0,
      "shape" : "ellipse",
      "width" : 45.0,
      "background-color" : "rgb(255,255,255)",
      "height" : 45.0,
      "content" : "data(name)"
    }
  }, {
    "selector" : "node:selected",
    "css" : {
      "background-color" : "rgb(255,0,102)"
    }
  }, {
    "selector" : "edge",
    "css" : {
      "font-size" : 12,
      "line-color" : "rgb(204,204,204)",
      "opacity" : 1.0,
      "source-arrow-shape" : "none",
      "target-arrow-color" : "rgb(204,204,204)",
      "text-opacity" : 1.0,
      "line-style" : "solid",
      "color" : "rgb(51,153,255)",
      "font-family" : "Dialog.plain",
      "font-weight" : "normal",
      "source-arrow-color" : "rgb(204,204,204)",
      "target-arrow-shape" : "triangle",
      "width" : 5.0,
      "content" : "data(interaction)"
    }
  }, {
    "selector" : "edge:selected",
    "css" : {
      "line-color" : "rgb(255,0,0)"
    }
  } ]
}, {
  "format_version" : "1.0",
  "generated_by" : "cytoscape-3.7.2",
  "target_cytoscapejs_version" : "~2.1",
  "title" : "bingo Style for Sheet 1_0",
  "style" : [ {
    "selector" : "node",
    "css" : {
      "text-opacity" : 1.0,
      "background-opacity" : 0.8235294117647058,
      "border-width" : 2.0,
      "color" : "rgb(0,0,0)",
      "border-color" : "rgb(64,64,64)",
      "font-size" : 11,
      "font-family" : "Dialog.plain",
      "font-weight" : "normal",
      "text-valign" : "center",
      "text-halign" : "center",
      "border-opacity" : 1.0,
      "shape" : "ellipse",
      "width" : 40.0,
      "background-color" : "rgb(200,0,0)",
      "height" : 40.0,
      "content" : "data(description_Sheet_1)"
    }
  }, {
    "selector" : "node[nodeFillColor_Sheet_1 > 6.30103]",
    "css" : {
      "background-color" : "rgb(255,127,0)"
    }
  }, {
    "selector" : "node[nodeFillColor_Sheet_1 = 6.30103]",
    "css" : {
      "background-color" : "rgb(255,127,0)"
    }
  }, {
    "selector" : "node[nodeFillColor_Sheet_1 > 1.30103][nodeFillColor_Sheet_1 < 6.30103]",
    "css" : {
      "background-color" : "mapData(nodeFillColor_Sheet_1,1.30103,6.30103,rgb(255,255,0),rgb(255,127,0))"
    }
  }, {
    "selector" : "node[nodeFillColor_Sheet_1 = 1.30103]",
    "css" : {
      "background-color" : "rgb(255,255,0)"
    }
  }, {
    "selector" : "node[nodeFillColor_Sheet_1 < 1.30103]",
    "css" : {
      "background-color" : "rgb(255,255,255)"
    }
  }, {
    "selector" : "node[nodeSize_Sheet_1 > 100]",
    "css" : {
      "width" : 400.0
    }
  }, {
    "selector" : "node[nodeSize_Sheet_1 = 100]",
    "css" : {
      "width" : 400.0
    }
  }, {
    "selector" : "node[nodeSize_Sheet_1 > 1][nodeSize_Sheet_1 < 100]",
    "css" : {
      "width" : "mapData(nodeSize_Sheet_1,1,100,20.0,400.0)"
    }
  }, {
    "selector" : "node[nodeSize_Sheet_1 = 1]",
    "css" : {
      "width" : 20.0
    }
  }, {
    "selector" : "node[nodeSize_Sheet_1 < 1]",
    "css" : {
      "width" : 20.0
    }
  }, {
    "selector" : "node[nodeSize_Sheet_1 > 100]",
    "css" : {
      "height" : 400.0
    }
  }, {
    "selector" : "node[nodeSize_Sheet_1 = 100]",
    "css" : {
      "height" : 400.0
    }
  }, {
    "selector" : "node[nodeSize_Sheet_1 > 1][nodeSize_Sheet_1 < 100]",
    "css" : {
      "height" : "mapData(nodeSize_Sheet_1,1,100,20.0,400.0)"
    }
  }, {
    "selector" : "node[nodeSize_Sheet_1 = 1]",
    "css" : {
      "height" : 20.0
    }
  }, {
    "selector" : "node[nodeSize_Sheet_1 < 1]",
    "css" : {
      "height" : 20.0
    }
  }, {
    "selector" : "node:selected",
    "css" : {
      "background-color" : "rgb(255,255,0)"
    }
  }, {
    "selector" : "edge",
    "css" : {
      "font-size" : 10,
      "line-color" : "rgb(64,64,64)",
      "opacity" : 0.7450980392156863,
      "source-arrow-shape" : "none",
      "target-arrow-color" : "rgb(0,0,0)",
      "text-opacity" : 1.0,
      "line-style" : "solid",
      "color" : "rgb(0,0,0)",
      "font-family" : "Dialog.plain",
      "font-weight" : "normal",
      "content" : "",
      "source-arrow-color" : "rgb(0,0,0)",
      "target-arrow-shape" : "triangle",
      "width" : 2.0
    }
  }, {
    "selector" : "edge:selected",
    "css" : {
      "line-color" : "rgb(255,0,0)"
    }
  } ]
}, {
  "format_version" : "1.0",
  "generated_by" : "cytoscape-3.7.2",
  "target_cytoscapejs_version" : "~2.1",
  "title" : "BioPAX_0",
  "style" : [ {
    "selector" : "node",
    "css" : {
      "text-opacity" : 1.0,
      "background-opacity" : 1.0,
      "border-width" : 2.0,
      "color" : "rgb(0,0,0)",
      "border-color" : "rgb(0,102,102)",
      "font-size" : 12,
      "font-family" : "Dialog.plain",
      "font-weight" : "normal",
      "text-valign" : "center",
      "text-halign" : "center",
      "border-opacity" : 1.0,
      "shape" : "ellipse",
      "width" : 20.0,
      "background-color" : "rgb(255,255,255)",
      "height" : 20.0,
      "content" : "data(name)"
    }
  }, {
    "selector" : "node[BIOPAX_TYPE = 'Complex']",
    "css" : {
      "background-color" : "rgb(255,255,255)"
    }
  }, {
    "selector" : "node[BIOPAX_TYPE = 'SimplePhysicalEntity']",
    "css" : {
      "shape" : "ellipse"
    }
  }, {
    "selector" : "node[BIOPAX_TYPE = 'Rna']",
    "css" : {
      "shape" : "ellipse"
    }
  }, {
    "selector" : "node[BIOPAX_TYPE = 'BiochemicalReaction']",
    "css" : {
      "shape" : "rectangle"
    }
  }, {
    "selector" : "node[BIOPAX_TYPE = 'GeneticInteraction']",
    "css" : {
      "shape" : "rectangle"
    }
  }, {
    "selector" : "node[BIOPAX_TYPE = 'Interaction']",
    "css" : {
      "shape" : "rectangle"
    }
  }, {
    "selector" : "node[BIOPAX_TYPE = 'TransportWithBiochemicalReaction']",
    "css" : {
      "shape" : "rectangle"
    }
  }, {
    "selector" : "node[BIOPAX_TYPE = 'Conversion']",
    "css" : {
      "shape" : "rectangle"
    }
  }, {
    "selector" : "node[BIOPAX_TYPE = 'ComplexAssembly']",
    "css" : {
      "shape" : "rectangle"
    }
  }, {
    "selector" : "node[BIOPAX_TYPE = 'Protein']",
    "css" : {
      "shape" : "ellipse"
    }
  }, {
    "selector" : "node[BIOPAX_TYPE = 'Complex']",
    "css" : {
      "shape" : "diamond"
    }
  }, {
    "selector" : "node[BIOPAX_TYPE = 'RnaRegion']",
    "css" : {
      "shape" : "ellipse"
    }
  }, {
    "selector" : "node[BIOPAX_TYPE = 'Degradation']",
    "css" : {
      "shape" : "rectangle"
    }
  }, {
    "selector" : "node[BIOPAX_TYPE = 'TemplateReactionRegulation']",
    "css" : {
      "shape" : "triangle"
    }
  }, {
    "selector" : "node[BIOPAX_TYPE = 'Control']",
    "css" : {
      "shape" : "triangle"
    }
  }, {
    "selector" : "node[BIOPAX_TYPE = 'PhysicalEntity']",
    "css" : {
      "shape" : "ellipse"
    }
  }, {
    "selector" : "node[BIOPAX_TYPE = 'DnaRegion']",
    "css" : {
      "shape" : "ellipse"
    }
  }, {
    "selector" : "node[BIOPAX_TYPE = 'SmallMolecule']",
    "css" : {
      "shape" : "ellipse"
    }
  }, {
    "selector" : "node[BIOPAX_TYPE = 'Dna']",
    "css" : {
      "shape" : "ellipse"
    }
  }, {
    "selector" : "node[BIOPAX_TYPE = 'Modulation']",
    "css" : {
      "shape" : "triangle"
    }
  }, {
    "selector" : "node[BIOPAX_TYPE = 'TemplateReaction']",
    "css" : {
      "shape" : "rectangle"
    }
  }, {
    "selector" : "node[BIOPAX_TYPE = 'MolecularInteraction']",
    "css" : {
      "shape" : "rectangle"
    }
  }, {
    "selector" : "node[BIOPAX_TYPE = 'Catalysis']",
    "css" : {
      "shape" : "triangle"
    }
  }, {
    "selector" : "node[BIOPAX_TYPE = 'Transport']",
    "css" : {
      "shape" : "rectangle"
    }
  }, {
    "selector" : "node[BIOPAX_TYPE = 'Protein-phosphorylated']",
    "css" : {
      "shape" : "ellipse"
    }
  }, {
    "selector" : "node[BIOPAX_TYPE = 'Complex']",
    "css" : {
      "border-color" : "rgb(0,102,102)"
    }
  }, {
    "selector" : "node[BIOPAX_TYPE = 'BiochemicalReaction']",
    "css" : {
      "width" : 13.4
    }
  }, {
    "selector" : "node[BIOPAX_TYPE = 'GeneticInteraction']",
    "css" : {
      "width" : 13.4
    }
  }, {
    "selector" : "node[BIOPAX_TYPE = 'Interaction']",
    "css" : {
      "width" : 13.4
    }
  }, {
    "selector" : "node[BIOPAX_TYPE = 'TransportWithBiochemicalReaction']",
    "css" : {
      "width" : 13.4
    }
  }, {
    "selector" : "node[BIOPAX_TYPE = 'Conversion']",
    "css" : {
      "width" : 13.4
    }
  }, {
    "selector" : "node[BIOPAX_TYPE = 'ComplexAssembly']",
    "css" : {
      "width" : 13.4
    }
  }, {
    "selector" : "node[BIOPAX_TYPE = 'Complex']",
    "css" : {
      "width" : 13.4
    }
  }, {
    "selector" : "node[BIOPAX_TYPE = 'Degradation']",
    "css" : {
      "width" : 13.4
    }
  }, {
    "selector" : "node[BIOPAX_TYPE = 'TemplateReactionRegulation']",
    "css" : {
      "width" : 13.4
    }
  }, {
    "selector" : "node[BIOPAX_TYPE = 'Control']",
    "css" : {
      "width" : 13.4
    }
  }, {
    "selector" : "node[BIOPAX_TYPE = 'Modulation']",
    "css" : {
      "width" : 13.4
    }
  }, {
    "selector" : "node[BIOPAX_TYPE = 'TemplateReaction']",
    "css" : {
      "width" : 13.4
    }
  }, {
    "selector" : "node[BIOPAX_TYPE = 'MolecularInteraction']",
    "css" : {
      "width" : 13.4
    }
  }, {
    "selector" : "node[BIOPAX_TYPE = 'Catalysis']",
    "css" : {
      "width" : 13.4
    }
  }, {
    "selector" : "node[BIOPAX_TYPE = 'Transport']",
    "css" : {
      "width" : 13.4
    }
  }, {
    "selector" : "node[BIOPAX_TYPE = 'BiochemicalReaction']",
    "css" : {
      "height" : 13.4
    }
  }, {
    "selector" : "node[BIOPAX_TYPE = 'GeneticInteraction']",
    "css" : {
      "height" : 13.4
    }
  }, {
    "selector" : "node[BIOPAX_TYPE = 'Interaction']",
    "css" : {
      "height" : 13.4
    }
  }, {
    "selector" : "node[BIOPAX_TYPE = 'TransportWithBiochemicalReaction']",
    "css" : {
      "height" : 13.4
    }
  }, {
    "selector" : "node[BIOPAX_TYPE = 'Conversion']",
    "css" : {
      "height" : 13.4
    }
  }, {
    "selector" : "node[BIOPAX_TYPE = 'ComplexAssembly']",
    "css" : {
      "height" : 13.4
    }
  }, {
    "selector" : "node[BIOPAX_TYPE = 'Complex']",
    "css" : {
      "height" : 13.4
    }
  }, {
    "selector" : "node[BIOPAX_TYPE = 'Degradation']",
    "css" : {
      "height" : 13.4
    }
  }, {
    "selector" : "node[BIOPAX_TYPE = 'TemplateReactionRegulation']",
    "css" : {
      "height" : 13.4
    }
  }, {
    "selector" : "node[BIOPAX_TYPE = 'Control']",
    "css" : {
      "height" : 13.4
    }
  }, {
    "selector" : "node[BIOPAX_TYPE = 'Modulation']",
    "css" : {
      "height" : 13.4
    }
  }, {
    "selector" : "node[BIOPAX_TYPE = 'TemplateReaction']",
    "css" : {
      "height" : 13.4
    }
  }, {
    "selector" : "node[BIOPAX_TYPE = 'MolecularInteraction']",
    "css" : {
      "height" : 13.4
    }
  }, {
    "selector" : "node[BIOPAX_TYPE = 'Catalysis']",
    "css" : {
      "height" : 13.4
    }
  }, {
    "selector" : "node[BIOPAX_TYPE = 'Transport']",
    "css" : {
      "height" : 13.4
    }
  }, {
    "selector" : "node:selected",
    "css" : {
      "background-color" : "rgb(255,255,0)"
    }
  }, {
    "selector" : "edge",
    "css" : {
      "font-size" : 10,
      "line-color" : "rgb(64,64,64)",
      "opacity" : 1.0,
      "source-arrow-shape" : "none",
      "target-arrow-color" : "rgb(64,64,64)",
      "text-opacity" : 1.0,
      "line-style" : "solid",
      "color" : "rgb(0,0,0)",
      "font-family" : "Dialog.plain",
      "font-weight" : "normal",
      "content" : "",
      "source-arrow-color" : "rgb(64,64,64)",
      "target-arrow-shape" : "none",
      "width" : 1.0
    }
  }, {
    "selector" : "edge[interaction = 'INHIBITION_NONCOMPETITIVE']",
    "css" : {
      "target-arrow-shape" : "tee"
    }
  }, {
    "selector" : "edge[interaction = 'INHIBITION_OTHER']",
    "css" : {
      "target-arrow-shape" : "tee"
    }
  }, {
    "selector" : "edge[interaction = 'ACTIVATION']",
    "css" : {
      "target-arrow-shape" : "triangle"
    }
  }, {
    "selector" : "edge[interaction = 'INHIBITION_UNCOMPETITIVE']",
    "css" : {
      "target-arrow-shape" : "tee"
    }
  }, {
    "selector" : "edge[interaction = 'cofactor']",
    "css" : {
      "target-arrow-shape" : "triangle"
    }
  }, {
    "selector" : "edge[interaction = 'ACTIVATION_ALLOSTERIC']",
    "css" : {
      "target-arrow-shape" : "triangle"
    }
  }, {
    "selector" : "edge[interaction = 'right']",
    "css" : {
      "target-arrow-shape" : "triangle"
    }
  }, {
    "selector" : "edge[interaction = 'INHIBITION_ALLOSTERIC']",
    "css" : {
      "target-arrow-shape" : "tee"
    }
  }, {
    "selector" : "edge[interaction = 'controlled']",
    "css" : {
      "target-arrow-shape" : "triangle"
    }
  }, {
    "selector" : "edge[interaction = 'contains']",
    "css" : {
      "target-arrow-shape" : "circle"
    }
  }, {
    "selector" : "edge[interaction = 'INHIBITION']",
    "css" : {
      "target-arrow-shape" : "tee"
    }
  }, {
    "selector" : "edge[interaction = 'INHIBITION_UNKMECH']",
    "css" : {
      "target-arrow-shape" : "tee"
    }
  }, {
    "selector" : "edge[interaction = 'INHIBITION_IRREVERSIBLE']",
    "css" : {
      "target-arrow-shape" : "tee"
    }
  }, {
    "selector" : "edge[interaction = 'INHIBITION_COMPETITIVE']",
    "css" : {
      "target-arrow-shape" : "tee"
    }
  }, {
    "selector" : "edge[interaction = 'ACTIVATION_UNKMECH']",
    "css" : {
      "target-arrow-shape" : "triangle"
    }
  }, {
    "selector" : "edge[interaction = 'ACTIVATION_NONALLOSTERIC']",
    "css" : {
      "target-arrow-shape" : "triangle"
    }
  }, {
    "selector" : "edge:selected",
    "css" : {
      "line-color" : "rgb(255,0,0)"
    }
  } ]
}, {
  "format_version" : "1.0",
  "generated_by" : "cytoscape-3.7.2",
  "target_cytoscapejs_version" : "~2.1",
  "title" : "size_rank_0_0",
  "style" : [ {
    "selector" : "node",
    "css" : {
      "text-opacity" : 1.0,
      "background-opacity" : 1.0,
      "border-width" : 0.0,
      "color" : "rgb(51,51,51)",
      "border-color" : "rgb(0,0,0)",
      "font-size" : 9,
      "font-family" : "Dialog.plain",
      "font-weight" : "normal",
      "text-valign" : "center",
      "text-halign" : "center",
      "border-opacity" : 1.0,
      "shape" : "rectangle",
      "width" : 12.0,
      "background-color" : "rgb(204,204,255)",
      "height" : 12.0,
      "content" : "data(name)"
    }
  }, {
    "selector" : "node:selected",
    "css" : {
      "background-color" : "rgb(255,255,0)"
    }
  }, {
    "selector" : "edge",
    "css" : {
      "font-size" : 10,
      "line-color" : "rgb(76,76,76)",
      "opacity" : 1.0,
      "source-arrow-shape" : "none",
      "target-arrow-color" : "rgb(0,0,0)",
      "text-opacity" : 1.0,
      "line-style" : "solid",
      "color" : "rgb(0,0,0)",
      "font-family" : "Dialog.plain",
      "font-weight" : "normal",
      "content" : "",
      "source-arrow-color" : "rgb(0,0,0)",
      "target-arrow-shape" : "none",
      "width" : 2.0
    }
  }, {
    "selector" : "edge:selected",
    "css" : {
      "line-color" : "rgb(255,0,0)"
    }
  } ]
}, {
  "format_version" : "1.0",
  "generated_by" : "cytoscape-3.7.2",
  "target_cytoscapejs_version" : "~2.1",
  "title" : "Diffany - Source_0",
  "style" : [ {
    "selector" : "node",
    "css" : {
      "text-opacity" : 1.0,
      "background-opacity" : 1.0,
      "border-width" : 0.0,
      "color" : "rgb(0,0,0)",
      "border-color" : "rgb(0,0,0)",
      "font-size" : 11,
      "font-family" : "SansSerif.plain",
      "font-weight" : "normal",
      "text-valign" : "center",
      "text-halign" : "center",
      "border-opacity" : 1.0,
      "shape" : "ellipse",
      "width" : 80.0,
      "background-color" : "rgb(128,128,128)",
      "height" : 55.0,
      "content" : "data(official_symbol)"
    }
  }, {
    "selector" : "node[differentially_expressed = 'up-regulated']",
    "css" : {
      "background-color" : "rgb(124,124,7)"
    }
  }, {
    "selector" : "node[differentially_expressed = 'down-regulated']",
    "css" : {
      "background-color" : "rgb(30,88,124)"
    }
  }, {
    "selector" : "node[kinase_function = 'yes']",
    "css" : {
      "border-width" : 2.0
    }
  }, {
    "selector" : "node[differentially_expressed = 'up-regulated']",
    "css" : {
      "background-color" : "rgb(255,255,15)"
    }
  }, {
    "selector" : "node[differentially_expressed = 'down-regulated']",
    "css" : {
      "background-color" : "rgb(62,181,255)"
    }
  }, {
    "selector" : "node[phosphorylation_site = 'yes']",
    "css" : {
      "shape" : "diamond"
    }
  }, {
    "selector" : "node[kinase_function = 'yes']",
    "css" : {
      "border-color" : "rgb(0,0,0)"
    }
  }, {
    "selector" : "node:selected",
    "css" : {
      "background-color" : "rgb(62,62,62)"
    }
  }, {
    "selector" : "edge",
    "css" : {
      "font-size" : 10,
      "line-color" : "rgb(64,64,64)",
      "opacity" : 1.0,
      "source-arrow-shape" : "none",
      "target-arrow-color" : "rgb(0,0,0)",
      "text-opacity" : 1.0,
      "line-style" : "solid",
      "color" : "rgb(0,0,0)",
      "font-family" : "SansSerif.plain",
      "font-weight" : "normal",
      "content" : "",
      "source-arrow-color" : "rgb(0,0,0)",
      "target-arrow-shape" : "none",
      "width" : 1.0
    }
  }, {
    "selector" : "edge[negated]",
    "css" : {
      "line-style" : "dotted"
    }
  }, {
    "selector" : "edge[!negated]",
    "css" : {
      "line-style" : "solid"
    }
  }, {
    "selector" : "edge[weight > 20]",
    "css" : {
      "width" : 25.0
    }
  }, {
    "selector" : "edge[weight = 20]",
    "css" : {
      "width" : 25.0
    }
  }, {
    "selector" : "edge[weight > 0][weight < 20]",
    "css" : {
      "width" : "mapData(weight,0,20,1.0,25.0)"
    }
  }, {
    "selector" : "edge[weight = 0]",
    "css" : {
      "width" : 1.0
    }
  }, {
    "selector" : "edge[weight < 0]",
    "css" : {
      "width" : 1.0
    }
  }, {
    "selector" : "edge:selected",
    "css" : {
      "line-color" : "rgb(255,0,0)"
    }
  } ]
}, {
  "format_version" : "1.0",
  "generated_by" : "cytoscape-3.7.2",
  "target_cytoscapejs_version" : "~2.1",
  "title" : "bingo Style for sheet 2_0",
  "style" : [ {
    "selector" : "node",
    "css" : {
      "text-opacity" : 1.0,
      "background-opacity" : 0.8235294117647058,
      "border-width" : 2.0,
      "color" : "rgb(0,0,0)",
      "border-color" : "rgb(64,64,64)",
      "font-size" : 11,
      "font-family" : "Dialog.plain",
      "font-weight" : "normal",
      "text-valign" : "center",
      "text-halign" : "center",
      "border-opacity" : 1.0,
      "shape" : "ellipse",
      "width" : 40.0,
      "background-color" : "rgb(200,0,0)",
      "height" : 40.0,
      "content" : "data(description_sheet_2)"
    }
  }, {
    "selector" : "node[nodeFillColor_sheet_2 > 6.30103]",
    "css" : {
      "background-color" : "rgb(255,127,0)"
    }
  }, {
    "selector" : "node[nodeFillColor_sheet_2 = 6.30103]",
    "css" : {
      "background-color" : "rgb(255,127,0)"
    }
  }, {
    "selector" : "node[nodeFillColor_sheet_2 > 1.30103][nodeFillColor_sheet_2 < 6.30103]",
    "css" : {
      "background-color" : "mapData(nodeFillColor_sheet_2,1.30103,6.30103,rgb(255,255,0),rgb(255,127,0))"
    }
  }, {
    "selector" : "node[nodeFillColor_sheet_2 = 1.30103]",
    "css" : {
      "background-color" : "rgb(255,255,0)"
    }
  }, {
    "selector" : "node[nodeFillColor_sheet_2 < 1.30103]",
    "css" : {
      "background-color" : "rgb(255,255,255)"
    }
  }, {
    "selector" : "node[nodeSize_sheet_2 > 100]",
    "css" : {
      "width" : 400.0
    }
  }, {
    "selector" : "node[nodeSize_sheet_2 = 100]",
    "css" : {
      "width" : 400.0
    }
  }, {
    "selector" : "node[nodeSize_sheet_2 > 1][nodeSize_sheet_2 < 100]",
    "css" : {
      "width" : "mapData(nodeSize_sheet_2,1,100,20.0,400.0)"
    }
  }, {
    "selector" : "node[nodeSize_sheet_2 = 1]",
    "css" : {
      "width" : 20.0
    }
  }, {
    "selector" : "node[nodeSize_sheet_2 < 1]",
    "css" : {
      "width" : 20.0
    }
  }, {
    "selector" : "node[nodeSize_sheet_2 > 100]",
    "css" : {
      "height" : 400.0
    }
  }, {
    "selector" : "node[nodeSize_sheet_2 = 100]",
    "css" : {
      "height" : 400.0
    }
  }, {
    "selector" : "node[nodeSize_sheet_2 > 1][nodeSize_sheet_2 < 100]",
    "css" : {
      "height" : "mapData(nodeSize_sheet_2,1,100,20.0,400.0)"
    }
  }, {
    "selector" : "node[nodeSize_sheet_2 = 1]",
    "css" : {
      "height" : 20.0
    }
  }, {
    "selector" : "node[nodeSize_sheet_2 < 1]",
    "css" : {
      "height" : 20.0
    }
  }, {
    "selector" : "node:selected",
    "css" : {
      "background-color" : "rgb(255,255,0)"
    }
  }, {
    "selector" : "edge",
    "css" : {
      "font-size" : 10,
      "line-color" : "rgb(64,64,64)",
      "opacity" : 0.7450980392156863,
      "source-arrow-shape" : "none",
      "target-arrow-color" : "rgb(0,0,0)",
      "text-opacity" : 1.0,
      "line-style" : "solid",
      "color" : "rgb(0,0,0)",
      "font-family" : "Dialog.plain",
      "font-weight" : "normal",
      "content" : "",
      "source-arrow-color" : "rgb(0,0,0)",
      "target-arrow-shape" : "triangle",
      "width" : 2.0
    }
  }, {
    "selector" : "edge:selected",
    "css" : {
      "line-color" : "rgb(255,0,0)"
    }
  } ]
}, {
  "format_version" : "1.0",
  "generated_by" : "cytoscape-3.7.2",
  "target_cytoscapejs_version" : "~2.1",
  "title" : "default black",
  "style" : [ {
    "selector" : "node",
    "css" : {
      "text-opacity" : 1.0,
      "background-opacity" : 1.0,
      "border-width" : 0.0,
      "color" : "rgb(204,204,204)",
      "border-color" : "rgb(0,153,0)",
      "font-size" : 12,
      "font-family" : "Dialog.plain",
      "font-weight" : "normal",
      "text-valign" : "bottom",
      "text-halign" : "right",
      "border-opacity" : 1.0,
      "shape" : "ellipse",
      "width" : 15.0,
      "background-color" : "rgb(255,255,255)",
      "height" : 15.0,
      "content" : "data(name)"
    }
  }, {
    "selector" : "node:selected",
    "css" : {
      "background-color" : "rgb(255,255,0)"
    }
  }, {
    "selector" : "edge",
    "css" : {
      "font-size" : 10,
      "line-color" : "rgb(0,153,0)",
      "opacity" : 1.0,
      "source-arrow-shape" : "none",
      "target-arrow-color" : "rgb(0,0,0)",
      "text-opacity" : 1.0,
      "line-style" : "solid",
      "color" : "rgb(0,0,0)",
      "font-family" : "Dialog.plain",
      "font-weight" : "normal",
      "content" : "",
      "source-arrow-color" : "rgb(0,0,0)",
      "target-arrow-shape" : "none",
      "width" : 2.0
    }
  }, {
    "selector" : "edge:selected",
    "css" : {
      "line-color" : "rgb(255,0,0)"
    }
  } ]
}, {
  "format_version" : "1.0",
  "generated_by" : "cytoscape-3.7.2",
  "target_cytoscapejs_version" : "~2.1",
  "title" : "bingo Style for sheet 2",
  "style" : [ {
    "selector" : "node",
    "css" : {
      "text-opacity" : 1.0,
      "background-opacity" : 0.8235294117647058,
      "border-width" : 2.0,
      "color" : "rgb(0,0,0)",
      "border-color" : "rgb(64,64,64)",
      "font-size" : 11,
      "font-family" : "Dialog.plain",
      "font-weight" : "normal",
      "text-valign" : "center",
      "text-halign" : "center",
      "border-opacity" : 1.0,
      "shape" : "ellipse",
      "width" : 40.0,
      "background-color" : "rgb(200,0,0)",
      "height" : 40.0,
      "content" : "data(description_sheet_2)"
    }
  }, {
    "selector" : "node[nodeFillColor_sheet_2 > 6.30103]",
    "css" : {
      "background-color" : "rgb(255,127,0)"
    }
  }, {
    "selector" : "node[nodeFillColor_sheet_2 = 6.30103]",
    "css" : {
      "background-color" : "rgb(255,127,0)"
    }
  }, {
    "selector" : "node[nodeFillColor_sheet_2 > 1.30103][nodeFillColor_sheet_2 < 6.30103]",
    "css" : {
      "background-color" : "mapData(nodeFillColor_sheet_2,1.30103,6.30103,rgb(255,255,0),rgb(255,127,0))"
    }
  }, {
    "selector" : "node[nodeFillColor_sheet_2 = 1.30103]",
    "css" : {
      "background-color" : "rgb(255,255,0)"
    }
  }, {
    "selector" : "node[nodeFillColor_sheet_2 < 1.30103]",
    "css" : {
      "background-color" : "rgb(255,255,255)"
    }
  }, {
    "selector" : "node[nodeSize_sheet_2 > 100]",
    "css" : {
      "width" : 400.0
    }
  }, {
    "selector" : "node[nodeSize_sheet_2 = 100]",
    "css" : {
      "width" : 400.0
    }
  }, {
    "selector" : "node[nodeSize_sheet_2 > 1][nodeSize_sheet_2 < 100]",
    "css" : {
      "width" : "mapData(nodeSize_sheet_2,1,100,20.0,400.0)"
    }
  }, {
    "selector" : "node[nodeSize_sheet_2 = 1]",
    "css" : {
      "width" : 20.0
    }
  }, {
    "selector" : "node[nodeSize_sheet_2 < 1]",
    "css" : {
      "width" : 20.0
    }
  }, {
    "selector" : "node[nodeSize_sheet_2 > 100]",
    "css" : {
      "height" : 400.0
    }
  }, {
    "selector" : "node[nodeSize_sheet_2 = 100]",
    "css" : {
      "height" : 400.0
    }
  }, {
    "selector" : "node[nodeSize_sheet_2 > 1][nodeSize_sheet_2 < 100]",
    "css" : {
      "height" : "mapData(nodeSize_sheet_2,1,100,20.0,400.0)"
    }
  }, {
    "selector" : "node[nodeSize_sheet_2 = 1]",
    "css" : {
      "height" : 20.0
    }
  }, {
    "selector" : "node[nodeSize_sheet_2 < 1]",
    "css" : {
      "height" : 20.0
    }
  }, {
    "selector" : "node:selected",
    "css" : {
      "background-color" : "rgb(255,255,0)"
    }
  }, {
    "selector" : "edge",
    "css" : {
      "font-size" : 10,
      "line-color" : "rgb(64,64,64)",
      "opacity" : 0.7450980392156863,
      "source-arrow-shape" : "none",
      "target-arrow-color" : "rgb(0,0,0)",
      "text-opacity" : 1.0,
      "line-style" : "solid",
      "color" : "rgb(0,0,0)",
      "font-family" : "Dialog.plain",
      "font-weight" : "normal",
      "content" : "",
      "source-arrow-color" : "rgb(0,0,0)",
      "target-arrow-shape" : "triangle",
      "width" : 2.0
    }
  }, {
    "selector" : "edge:selected",
    "css" : {
      "line-color" : "rgb(255,0,0)"
    }
  } ]
}, {
  "format_version" : "1.0",
  "generated_by" : "cytoscape-3.7.2",
  "target_cytoscapejs_version" : "~2.1",
  "title" : "size_rank_0",
  "style" : [ {
    "selector" : "node",
    "css" : {
      "text-opacity" : 1.0,
      "background-opacity" : 1.0,
      "border-width" : 0.0,
      "color" : "rgb(51,51,51)",
      "border-color" : "rgb(0,0,0)",
      "font-size" : 9,
      "font-family" : "SansSerif.plain",
      "font-weight" : "normal",
      "text-valign" : "center",
      "text-halign" : "center",
      "border-opacity" : 1.0,
      "shape" : "rectangle",
      "width" : 12.0,
      "background-color" : "rgb(204,204,255)",
      "height" : 12.0,
      "content" : "data(name)"
    }
  }, {
    "selector" : "node:selected",
    "css" : {
      "background-color" : "rgb(255,255,0)"
    }
  }, {
    "selector" : "edge",
    "css" : {
      "font-size" : 10,
      "line-color" : "rgb(76,76,76)",
      "opacity" : 1.0,
      "source-arrow-shape" : "none",
      "target-arrow-color" : "rgb(0,0,0)",
      "text-opacity" : 1.0,
      "line-style" : "solid",
      "color" : "rgb(0,0,0)",
      "font-family" : "SansSerif.plain",
      "font-weight" : "normal",
      "content" : "",
      "source-arrow-color" : "rgb(0,0,0)",
      "target-arrow-shape" : "none",
      "width" : 2.0
    }
  }, {
    "selector" : "edge:selected",
    "css" : {
      "line-color" : "rgb(255,0,0)"
    }
  } ]
}, {
  "format_version" : "1.0",
  "generated_by" : "cytoscape-3.7.2",
  "target_cytoscapejs_version" : "~2.1",
  "title" : "Gradient1",
  "style" : [ {
    "selector" : "node",
    "css" : {
      "text-opacity" : 1.0,
      "background-opacity" : 1.0,
      "border-width" : 0.0,
      "color" : "rgb(204,204,204)",
      "border-color" : "rgb(0,0,0)",
      "font-size" : 8,
      "font-family" : "Dialog.plain",
      "font-weight" : "normal",
      "text-valign" : "bottom",
      "text-halign" : "right",
      "border-opacity" : 1.0,
      "shape" : "ellipse",
      "width" : 30.0,
      "background-color" : "rgb(0,0,0)",
      "height" : 30.0,
      "content" : "data(name)"
    }
  }, {
    "selector" : "node:selected",
    "css" : {
      "background-color" : "rgb(255,255,0)"
    }
  }, {
    "selector" : "edge",
    "css" : {
      "font-size" : 10,
      "line-color" : "rgb(102,102,102)",
      "opacity" : 1.0,
      "source-arrow-shape" : "none",
      "target-arrow-color" : "rgb(0,0,0)",
      "text-opacity" : 1.0,
      "line-style" : "solid",
      "color" : "rgb(0,0,0)",
      "font-family" : "Dialog.plain",
      "font-weight" : "normal",
      "content" : "",
      "source-arrow-color" : "rgb(0,0,0)",
      "target-arrow-shape" : "none",
      "width" : 1.0
    }
  }, {
    "selector" : "edge:selected",
    "css" : {
      "line-color" : "rgb(255,0,0)"
    }
  } ]
}, {
  "format_version" : "1.0",
  "generated_by" : "cytoscape-3.7.2",
  "target_cytoscapejs_version" : "~2.1",
  "title" : "default",
  "style" : [ {
    "selector" : "node",
    "css" : {
      "text-opacity" : 1.0,
      "background-opacity" : 1.0,
      "border-width" : 0.0,
      "color" : "rgb(0,0,0)",
      "border-color" : "rgb(204,204,204)",
      "font-size" : 12,
      "font-family" : "Dialog.plain",
      "font-weight" : "normal",
      "text-valign" : "center",
      "text-halign" : "center",
      "border-opacity" : 1.0,
      "shape" : "roundrectangle",
      "width" : 35.0,
      "background-color" : "rgb(137,208,245)",
      "height" : 35.0,
      "content" : "data(symbol11)"
    }
  }, {
    "selector" : "node[MolecularAttributes = 'METABOLIC']",
    "css" : {
      "width" : 60.0,
      "height" : 60.0
    }
  }, {
    "selector" : "node[MolecularAttributes = 'TXNFACTOR']",
    "css" : {
      "width" : 60.0,
      "height" : 60.0
    }
  }, {
    "selector" : "node[MolecularAttributes = 'small_nucleolar_rna']",
    "css" : {
      "width" : 40.0,
      "height" : 40.0
    }
  }, {
    "selector" : "node[MolecularAttributes = 'protein_coding']",
    "css" : {
      "width" : 60.0,
      "height" : 60.0
    }
  }, {
    "selector" : "node[MolecularAttributes = 'other_rna']",
    "css" : {
      "width" : 40.0,
      "height" : 40.0
    }
  }, {
    "selector" : "node[MolecularAttributes = 'METABOLIC']",
    "css" : {
      "background-color" : "rgb(51,51,255)"
    }
  }, {
    "selector" : "node[MolecularAttributes = 'TXNFACTOR']",
    "css" : {
      "background-color" : "rgb(0,153,51)"
    }
  }, {
    "selector" : "node[MolecularAttributes = 'small_nucleolar_rna']",
    "css" : {
      "background-color" : "rgb(255,0,255)"
    }
  }, {
    "selector" : "node[MolecularAttributes = 'protein_coding']",
    "css" : {
      "background-color" : "rgb(102,204,255)"
    }
  }, {
    "selector" : "node[MolecularAttributes = 'other_rna']",
    "css" : {
      "background-color" : "rgb(255,0,255)"
    }
  }, {
    "selector" : "node[MolecularAttributes = 'METABOLIC']",
    "css" : {
      "shape" : "roundrectangle"
    }
  }, {
    "selector" : "node[MolecularAttributes = 'TXNFACTOR']",
    "css" : {
      "shape" : "triangle"
    }
  }, {
    "selector" : "node[MolecularAttributes = 'small_nucleolar_rna']",
    "css" : {
      "shape" : "ellipse"
    }
  }, {
    "selector" : "node[MolecularAttributes = 'protein_coding']",
    "css" : {
      "shape" : "roundrectangle"
    }
  }, {
    "selector" : "node[MolecularAttributes = 'other_rna']",
    "css" : {
      "shape" : "ellipse"
    }
  }, {
    "selector" : "node:selected",
    "css" : {
      "background-color" : "rgb(255,255,0)"
    }
  }, {
    "selector" : "edge",
    "css" : {
      "font-size" : 10,
      "line-color" : "rgb(64,64,64)",
      "opacity" : 1.0,
      "source-arrow-shape" : "none",
      "target-arrow-color" : "rgb(64,64,64)",
      "text-opacity" : 1.0,
      "line-style" : "solid",
      "color" : "rgb(0,0,0)",
      "font-family" : "Dialog.plain",
      "font-weight" : "normal",
      "content" : "",
      "source-arrow-color" : "rgb(64,64,64)",
      "target-arrow-shape" : "none",
      "width" : 2.0
    }
  }, {
    "selector" : "edge[Relationship = 'Negative']",
    "css" : {
      "line-color" : "rgb(0,204,0)",
      "target-arrow-color" : "rgb(0,204,0)",
      "source-arrow-color" : "rgb(0,204,0)"
    }
  }, {
    "selector" : "edge[Relationship = 'Positive']",
    "css" : {
      "line-color" : "rgb(255,0,0)",
      "target-arrow-color" : "rgb(255,0,0)",
      "source-arrow-color" : "rgb(255,0,0)"
    }
  }, {
    "selector" : "edge[Direction = 'R->S']",
    "css" : {
      "line-color" : "rgb(51,51,255)"
    }
  }, {
    "selector" : "edge[Direction = 'R<-S']",
    "css" : {
      "line-color" : "rgb(0,204,0)"
    }
  }, {
    "selector" : "edge[Direction = 'R->S']",
    "css" : {
      "source-arrow-shape" : "none"
    }
  }, {
    "selector" : "edge[Direction = 'R<-S']",
    "css" : {
      "source-arrow-shape" : "triangle"
    }
  }, {
    "selector" : "edge[Direction = 'R->S']",
    "css" : {
      "line-style" : "dashed"
    }
  }, {
    "selector" : "edge[Direction = 'R->S']",
    "css" : {
      "target-arrow-shape" : "triangle"
    }
  }, {
    "selector" : "edge[Direction = 'R<-S']",
    "css" : {
      "target-arrow-shape" : "none"
    }
  }, {
    "selector" : "edge:selected",
    "css" : {
      "line-color" : "rgb(255,0,0)"
    }
  } ]
}, {
  "format_version" : "1.0",
  "generated_by" : "cytoscape-3.7.2",
  "target_cytoscapejs_version" : "~2.1",
  "title" : "Ripple",
  "style" : [ {
    "selector" : "node",
    "css" : {
      "text-opacity" : 1.0,
      "background-opacity" : 1.0,
      "border-width" : 20.0,
      "color" : "rgb(19,58,96)",
      "border-color" : "rgb(51,153,255)",
      "font-size" : 8,
      "font-family" : "Dialog.plain",
      "font-weight" : "normal",
      "text-valign" : "center",
      "text-halign" : "center",
      "border-opacity" : 1.0,
      "shape" : "ellipse",
      "width" : 50.0,
      "background-color" : "rgb(255,255,255)",
      "height" : 50.0,
      "content" : "data(name)"
    }
  }, {
    "selector" : "node:selected",
    "css" : {
      "background-color" : "rgb(255,255,204)"
    }
  }, {
    "selector" : "edge",
    "css" : {
      "font-size" : 10,
      "line-color" : "rgb(51,153,255)",
      "opacity" : 1.0,
      "source-arrow-shape" : "none",
      "target-arrow-color" : "rgb(0,0,0)",
      "text-opacity" : 1.0,
      "line-style" : "solid",
      "color" : "rgb(0,0,0)",
      "font-family" : "Dialog.plain",
      "font-weight" : "normal",
      "content" : "",
      "source-arrow-color" : "rgb(0,0,0)",
      "target-arrow-shape" : "none",
      "width" : 3.0
    }
  }, {
    "selector" : "edge:selected",
    "css" : {
      "line-color" : "rgb(255,0,0)"
    }
  } ]
}, {
  "format_version" : "1.0",
  "generated_by" : "cytoscape-3.7.2",
  "target_cytoscapejs_version" : "~2.1",
  "title" : "Diffany - Source_1",
  "style" : [ {
    "selector" : "node",
    "css" : {
      "text-opacity" : 1.0,
      "background-opacity" : 1.0,
      "border-width" : 0.0,
      "color" : "rgb(0,0,0)",
      "border-color" : "rgb(0,0,0)",
      "font-size" : 11,
      "font-family" : "SansSerif.plain",
      "font-weight" : "normal",
      "text-valign" : "center",
      "text-halign" : "center",
      "border-opacity" : 1.0,
      "shape" : "ellipse",
      "width" : 80.0,
      "background-color" : "rgb(128,128,128)",
      "height" : 55.0,
      "content" : "data(official_symbol)"
    }
  }, {
    "selector" : "node[differentially_expressed = 'up-regulated']",
    "css" : {
      "background-color" : "rgb(124,124,7)"
    }
  }, {
    "selector" : "node[differentially_expressed = 'down-regulated']",
    "css" : {
      "background-color" : "rgb(30,88,124)"
    }
  }, {
    "selector" : "node[kinase_function = 'yes']",
    "css" : {
      "border-width" : 2.0
    }
  }, {
    "selector" : "node[differentially_expressed = 'up-regulated']",
    "css" : {
      "background-color" : "rgb(255,255,15)"
    }
  }, {
    "selector" : "node[differentially_expressed = 'down-regulated']",
    "css" : {
      "background-color" : "rgb(62,181,255)"
    }
  }, {
    "selector" : "node[phosphorylation_site = 'yes']",
    "css" : {
      "shape" : "diamond"
    }
  }, {
    "selector" : "node[kinase_function = 'yes']",
    "css" : {
      "border-color" : "rgb(0,0,0)"
    }
  }, {
    "selector" : "node:selected",
    "css" : {
      "background-color" : "rgb(62,62,62)"
    }
  }, {
    "selector" : "edge",
    "css" : {
      "font-size" : 10,
      "line-color" : "rgb(64,64,64)",
      "opacity" : 1.0,
      "source-arrow-shape" : "none",
      "target-arrow-color" : "rgb(0,0,0)",
      "text-opacity" : 1.0,
      "line-style" : "solid",
      "color" : "rgb(0,0,0)",
      "font-family" : "SansSerif.plain",
      "font-weight" : "normal",
      "content" : "",
      "source-arrow-color" : "rgb(0,0,0)",
      "target-arrow-shape" : "none",
      "width" : 1.0
    }
  }, {
    "selector" : "edge[negated]",
    "css" : {
      "line-style" : "dotted"
    }
  }, {
    "selector" : "edge[!negated]",
    "css" : {
      "line-style" : "solid"
    }
  }, {
    "selector" : "edge[weight > 20]",
    "css" : {
      "width" : 25.0
    }
  }, {
    "selector" : "edge[weight = 20]",
    "css" : {
      "width" : 25.0
    }
  }, {
    "selector" : "edge[weight > 0][weight < 20]",
    "css" : {
      "width" : "mapData(weight,0,20,1.0,25.0)"
    }
  }, {
    "selector" : "edge[weight = 0]",
    "css" : {
      "width" : 1.0
    }
  }, {
    "selector" : "edge[weight < 0]",
    "css" : {
      "width" : 1.0
    }
  }, {
    "selector" : "edge:selected",
    "css" : {
      "line-color" : "rgb(255,0,0)"
    }
  } ]
}, {
  "format_version" : "1.0",
  "generated_by" : "cytoscape-3.7.2",
  "target_cytoscapejs_version" : "~2.1",
  "title" : "Stuti's magical network",
  "style" : [ {
    "selector" : "node",
    "css" : {
      "text-opacity" : 1.0,
      "background-opacity" : 1.0,
      "border-width" : 0.0,
      "color" : "rgb(0,0,0)",
      "border-color" : "rgb(204,204,204)",
      "font-size" : 12,
      "font-family" : "Dialog.plain",
      "font-weight" : "normal",
      "text-valign" : "center",
      "text-halign" : "center",
      "border-opacity" : 1.0,
      "shape" : "roundrectangle",
      "width" : 35.0,
      "background-color" : "rgb(137,208,245)",
      "height" : 35.0,
      "content" : "data(Name2)"
    }
  }, {
    "selector" : "node[MolecularAttributes = 'METABOLIC']",
    "css" : {
      "width" : 60.0,
      "height" : 60.0
    }
  }, {
    "selector" : "node[MolecularAttributes = 'TXNFACTOR']",
    "css" : {
      "width" : 60.0,
      "height" : 60.0
    }
  }, {
    "selector" : "node[MolecularAttributes = 'small_nucleolar_rna']",
    "css" : {
      "width" : 40.0,
      "height" : 40.0
    }
  }, {
    "selector" : "node[MolecularAttributes = 'protein_coding']",
    "css" : {
      "width" : 60.0,
      "height" : 60.0
    }
  }, {
    "selector" : "node[MolecularAttributes = 'other_rna']",
    "css" : {
      "width" : 40.0,
      "height" : 40.0
    }
  }, {
    "selector" : "node[MolecularAttributes = 'METABOLIC']",
    "css" : {
      "background-color" : "rgb(51,51,255)"
    }
  }, {
    "selector" : "node[MolecularAttributes = 'TXNFACTOR']",
    "css" : {
      "background-color" : "rgb(0,153,51)"
    }
  }, {
    "selector" : "node[MolecularAttributes = 'small_nucleolar_rna']",
    "css" : {
      "background-color" : "rgb(255,0,255)"
    }
  }, {
    "selector" : "node[MolecularAttributes = 'protein_coding']",
    "css" : {
      "background-color" : "rgb(102,204,255)"
    }
  }, {
    "selector" : "node[MolecularAttributes = 'other_rna']",
    "css" : {
      "background-color" : "rgb(255,0,255)"
    }
  }, {
    "selector" : "node[MolecularAttributes = 'METABOLIC']",
    "css" : {
      "shape" : "roundrectangle"
    }
  }, {
    "selector" : "node[MolecularAttributes = 'TXNFACTOR']",
    "css" : {
      "shape" : "triangle"
    }
  }, {
    "selector" : "node[MolecularAttributes = 'small_nucleolar_rna']",
    "css" : {
      "shape" : "ellipse"
    }
  }, {
    "selector" : "node[MolecularAttributes = 'protein_coding']",
    "css" : {
      "shape" : "roundrectangle"
    }
  }, {
    "selector" : "node[MolecularAttributes = 'other_rna']",
    "css" : {
      "shape" : "ellipse"
    }
  }, {
    "selector" : "node:selected",
    "css" : {
      "background-color" : "rgb(255,255,0)"
    }
  }, {
    "selector" : "edge",
    "css" : {
      "font-size" : 12,
      "line-color" : "rgb(64,64,64)",
      "opacity" : 1.0,
      "source-arrow-shape" : "none",
      "target-arrow-color" : "rgb(64,64,64)",
      "text-opacity" : 1.0,
      "line-style" : "solid",
      "color" : "rgb(0,0,0)",
      "font-family" : "Dialog.plain",
      "font-weight" : "normal",
      "content" : "",
      "source-arrow-color" : "rgb(64,64,64)",
      "target-arrow-shape" : "none",
      "width" : 2.0
    }
  }, {
    "selector" : "edge[Relationship = 'Negative']",
    "css" : {
      "line-color" : "rgb(0,204,0)",
      "target-arrow-color" : "rgb(0,204,0)",
      "source-arrow-color" : "rgb(0,204,0)"
    }
  }, {
    "selector" : "edge[Relationship = 'Positive']",
    "css" : {
      "line-color" : "rgb(255,0,51)",
      "target-arrow-color" : "rgb(255,0,51)",
      "source-arrow-color" : "rgb(255,0,51)"
    }
  }, {
    "selector" : "edge[Direction = 'shoot-to-root']",
    "css" : {
      "content" : "shoot-to-root"
    }
  }, {
    "selector" : "edge[Direction = 'root-to-shoot']",
    "css" : {
      "content" : "root-to-shoot"
    }
  }, {
    "selector" : "edge[Direction = 'R->S']",
    "css" : {
      "line-color" : "rgb(51,51,255)"
    }
  }, {
    "selector" : "edge[Direction = 'R<-S']",
    "css" : {
      "line-color" : "rgb(0,204,0)"
    }
  }, {
    "selector" : "edge[Direction = 'shoot-to-root']",
    "css" : {
      "line-style" : "solid"
    }
  }, {
    "selector" : "edge[Direction = 'root-to-shoot']",
    "css" : {
      "line-style" : "dashed"
    }
  }, {
    "selector" : "edge[Relationship = 'Negative']",
    "css" : {
      "target-arrow-shape" : "tee"
    }
  }, {
    "selector" : "edge[Relationship = 'Positive']",
    "css" : {
      "target-arrow-shape" : "triangle"
    }
  }, {
    "selector" : "edge:selected",
    "css" : {
      "line-color" : "rgb(255,0,0)"
    }
  } ]
}, {
  "format_version" : "1.0",
  "generated_by" : "cytoscape-3.7.2",
  "target_cytoscapejs_version" : "~2.1",
  "title" : "Universe",
  "style" : [ {
    "selector" : "node",
    "css" : {
      "text-opacity" : 1.0,
      "background-opacity" : 1.0,
      "border-width" : 0.0,
      "color" : "rgb(255,255,255)",
      "border-color" : "rgb(0,0,0)",
      "font-size" : 18,
      "font-family" : "Dialog.plain",
      "font-weight" : "normal",
      "text-valign" : "center",
      "text-halign" : "center",
      "border-opacity" : 1.0,
      "shape" : "ellipse",
      "width" : 40.0,
      "background-color" : "rgb(0,0,0)",
      "height" : 40.0,
      "content" : "data(name)"
    }
  }, {
    "selector" : "node:selected",
    "css" : {
      "background-color" : "rgb(255,255,0)"
    }
  }, {
    "selector" : "edge",
    "css" : {
      "font-size" : 10,
      "line-color" : "rgb(153,153,153)",
      "opacity" : 1.0,
      "source-arrow-shape" : "none",
      "target-arrow-color" : "rgb(0,0,0)",
      "text-opacity" : 1.0,
      "line-style" : "dashed",
      "color" : "rgb(0,0,0)",
      "font-family" : "Dialog.plain",
      "font-weight" : "normal",
      "content" : "",
      "source-arrow-color" : "rgb(0,0,0)",
      "target-arrow-shape" : "none",
      "width" : 2.0
    }
  }, {
    "selector" : "edge:selected",
    "css" : {
      "line-color" : "rgb(255,0,0)"
    }
  } ]
}, {
  "format_version" : "1.0",
  "generated_by" : "cytoscape-3.7.2",
  "target_cytoscapejs_version" : "~2.1",
  "title" : "Solid",
  "style" : [ {
    "selector" : "node",
    "css" : {
      "text-opacity" : 1.0,
      "background-opacity" : 1.0,
      "border-width" : 0.0,
      "color" : "rgb(0,0,0)",
      "border-color" : "rgb(0,0,0)",
      "font-size" : 14,
      "font-family" : "Dialog.plain",
      "font-weight" : "normal",
      "text-valign" : "center",
      "text-halign" : "center",
      "border-opacity" : 1.0,
      "shape" : "ellipse",
      "width" : 40.0,
      "background-color" : "rgb(102,102,102)",
      "height" : 40.0,
      "content" : "data(name)"
    }
  }, {
    "selector" : "node:selected",
    "css" : {
      "background-color" : "rgb(255,255,0)"
    }
  }, {
    "selector" : "edge",
    "css" : {
      "font-size" : 10,
      "line-color" : "rgb(204,204,204)",
      "opacity" : 1.0,
      "source-arrow-shape" : "none",
      "target-arrow-color" : "rgb(0,0,0)",
      "text-opacity" : 1.0,
      "line-style" : "solid",
      "color" : "rgb(0,0,0)",
      "font-family" : "Dialog.plain",
      "font-weight" : "normal",
      "source-arrow-color" : "rgb(0,0,0)",
      "target-arrow-shape" : "none",
      "width" : 12.0,
      "content" : "data(interaction)"
    }
  }, {
    "selector" : "edge:selected",
    "css" : {
      "line-color" : "rgb(255,0,0)"
    }
  } ]
}, {
  "format_version" : "1.0",
  "generated_by" : "cytoscape-3.7.2",
  "target_cytoscapejs_version" : "~2.1",
  "title" : "size_rank_0_1",
  "style" : [ {
    "selector" : "node",
    "css" : {
      "text-opacity" : 1.0,
      "background-opacity" : 1.0,
      "border-width" : 0.0,
      "color" : "rgb(51,51,51)",
      "border-color" : "rgb(0,0,0)",
      "font-size" : 9,
      "font-family" : "Dialog.plain",
      "font-weight" : "normal",
      "text-valign" : "center",
      "text-halign" : "center",
      "border-opacity" : 1.0,
      "shape" : "rectangle",
      "width" : 12.0,
      "background-color" : "rgb(204,204,255)",
      "height" : 12.0,
      "content" : "data(name)"
    }
  }, {
    "selector" : "node:selected",
    "css" : {
      "background-color" : "rgb(255,255,0)"
    }
  }, {
    "selector" : "edge",
    "css" : {
      "font-size" : 10,
      "line-color" : "rgb(76,76,76)",
      "opacity" : 1.0,
      "source-arrow-shape" : "none",
      "target-arrow-color" : "rgb(0,0,0)",
      "text-opacity" : 1.0,
      "line-style" : "solid",
      "color" : "rgb(0,0,0)",
      "font-family" : "Dialog.plain",
      "font-weight" : "normal",
      "content" : "",
      "source-arrow-color" : "rgb(0,0,0)",
      "target-arrow-shape" : "none",
      "width" : 2.0
    }
  }, {
    "selector" : "edge:selected",
    "css" : {
      "line-color" : "rgb(255,0,0)"
    }
  } ]
}, {
  "format_version" : "1.0",
  "generated_by" : "cytoscape-3.7.2",
  "target_cytoscapejs_version" : "~2.1",
  "title" : "BioPAX",
  "style" : [ {
    "selector" : "node",
    "css" : {
      "text-opacity" : 1.0,
      "background-opacity" : 1.0,
      "border-width" : 2.0,
      "color" : "rgb(0,0,0)",
      "border-color" : "rgb(0,102,102)",
      "font-size" : 12,
      "font-family" : "SansSerif.plain",
      "font-weight" : "normal",
      "text-valign" : "center",
      "text-halign" : "center",
      "border-opacity" : 1.0,
      "shape" : "ellipse",
      "width" : 20.0,
      "background-color" : "rgb(255,255,255)",
      "height" : 20.0,
      "content" : "data(name)"
    }
  }, {
    "selector" : "node[BIOPAX_TYPE = 'Complex']",
    "css" : {
      "background-color" : "rgb(255,255,255)"
    }
  }, {
    "selector" : "node[BIOPAX_TYPE = 'SimplePhysicalEntity']",
    "css" : {
      "shape" : "ellipse"
    }
  }, {
    "selector" : "node[BIOPAX_TYPE = 'Rna']",
    "css" : {
      "shape" : "ellipse"
    }
  }, {
    "selector" : "node[BIOPAX_TYPE = 'GeneticInteraction']",
    "css" : {
      "shape" : "rectangle"
    }
  }, {
    "selector" : "node[BIOPAX_TYPE = 'BiochemicalReaction']",
    "css" : {
      "shape" : "rectangle"
    }
  }, {
    "selector" : "node[BIOPAX_TYPE = 'Interaction']",
    "css" : {
      "shape" : "rectangle"
    }
  }, {
    "selector" : "node[BIOPAX_TYPE = 'TransportWithBiochemicalReaction']",
    "css" : {
      "shape" : "rectangle"
    }
  }, {
    "selector" : "node[BIOPAX_TYPE = 'Conversion']",
    "css" : {
      "shape" : "rectangle"
    }
  }, {
    "selector" : "node[BIOPAX_TYPE = 'ComplexAssembly']",
    "css" : {
      "shape" : "rectangle"
    }
  }, {
    "selector" : "node[BIOPAX_TYPE = 'Protein']",
    "css" : {
      "shape" : "ellipse"
    }
  }, {
    "selector" : "node[BIOPAX_TYPE = 'RnaRegion']",
    "css" : {
      "shape" : "ellipse"
    }
  }, {
    "selector" : "node[BIOPAX_TYPE = 'Complex']",
    "css" : {
      "shape" : "diamond"
    }
  }, {
    "selector" : "node[BIOPAX_TYPE = 'Degradation']",
    "css" : {
      "shape" : "rectangle"
    }
  }, {
    "selector" : "node[BIOPAX_TYPE = 'TemplateReactionRegulation']",
    "css" : {
      "shape" : "triangle"
    }
  }, {
    "selector" : "node[BIOPAX_TYPE = 'Control']",
    "css" : {
      "shape" : "triangle"
    }
  }, {
    "selector" : "node[BIOPAX_TYPE = 'DnaRegion']",
    "css" : {
      "shape" : "ellipse"
    }
  }, {
    "selector" : "node[BIOPAX_TYPE = 'PhysicalEntity']",
    "css" : {
      "shape" : "ellipse"
    }
  }, {
    "selector" : "node[BIOPAX_TYPE = 'SmallMolecule']",
    "css" : {
      "shape" : "ellipse"
    }
  }, {
    "selector" : "node[BIOPAX_TYPE = 'Dna']",
    "css" : {
      "shape" : "ellipse"
    }
  }, {
    "selector" : "node[BIOPAX_TYPE = 'Modulation']",
    "css" : {
      "shape" : "triangle"
    }
  }, {
    "selector" : "node[BIOPAX_TYPE = 'TemplateReaction']",
    "css" : {
      "shape" : "rectangle"
    }
  }, {
    "selector" : "node[BIOPAX_TYPE = 'MolecularInteraction']",
    "css" : {
      "shape" : "rectangle"
    }
  }, {
    "selector" : "node[BIOPAX_TYPE = 'Catalysis']",
    "css" : {
      "shape" : "triangle"
    }
  }, {
    "selector" : "node[BIOPAX_TYPE = 'Transport']",
    "css" : {
      "shape" : "rectangle"
    }
  }, {
    "selector" : "node[BIOPAX_TYPE = 'Protein-phosphorylated']",
    "css" : {
      "shape" : "ellipse"
    }
  }, {
    "selector" : "node[BIOPAX_TYPE = 'Complex']",
    "css" : {
      "border-color" : "rgb(0,102,102)"
    }
  }, {
    "selector" : "node[BIOPAX_TYPE = 'GeneticInteraction']",
    "css" : {
      "width" : 13.4
    }
  }, {
    "selector" : "node[BIOPAX_TYPE = 'BiochemicalReaction']",
    "css" : {
      "width" : 13.4
    }
  }, {
    "selector" : "node[BIOPAX_TYPE = 'Interaction']",
    "css" : {
      "width" : 13.4
    }
  }, {
    "selector" : "node[BIOPAX_TYPE = 'TransportWithBiochemicalReaction']",
    "css" : {
      "width" : 13.4
    }
  }, {
    "selector" : "node[BIOPAX_TYPE = 'Conversion']",
    "css" : {
      "width" : 13.4
    }
  }, {
    "selector" : "node[BIOPAX_TYPE = 'ComplexAssembly']",
    "css" : {
      "width" : 13.4
    }
  }, {
    "selector" : "node[BIOPAX_TYPE = 'Complex']",
    "css" : {
      "width" : 13.4
    }
  }, {
    "selector" : "node[BIOPAX_TYPE = 'Degradation']",
    "css" : {
      "width" : 13.4
    }
  }, {
    "selector" : "node[BIOPAX_TYPE = 'TemplateReactionRegulation']",
    "css" : {
      "width" : 13.4
    }
  }, {
    "selector" : "node[BIOPAX_TYPE = 'Control']",
    "css" : {
      "width" : 13.4
    }
  }, {
    "selector" : "node[BIOPAX_TYPE = 'Modulation']",
    "css" : {
      "width" : 13.4
    }
  }, {
    "selector" : "node[BIOPAX_TYPE = 'TemplateReaction']",
    "css" : {
      "width" : 13.4
    }
  }, {
    "selector" : "node[BIOPAX_TYPE = 'MolecularInteraction']",
    "css" : {
      "width" : 13.4
    }
  }, {
    "selector" : "node[BIOPAX_TYPE = 'Catalysis']",
    "css" : {
      "width" : 13.4
    }
  }, {
    "selector" : "node[BIOPAX_TYPE = 'Transport']",
    "css" : {
      "width" : 13.4
    }
  }, {
    "selector" : "node[BIOPAX_TYPE = 'GeneticInteraction']",
    "css" : {
      "height" : 13.4
    }
  }, {
    "selector" : "node[BIOPAX_TYPE = 'BiochemicalReaction']",
    "css" : {
      "height" : 13.4
    }
  }, {
    "selector" : "node[BIOPAX_TYPE = 'Interaction']",
    "css" : {
      "height" : 13.4
    }
  }, {
    "selector" : "node[BIOPAX_TYPE = 'TransportWithBiochemicalReaction']",
    "css" : {
      "height" : 13.4
    }
  }, {
    "selector" : "node[BIOPAX_TYPE = 'Conversion']",
    "css" : {
      "height" : 13.4
    }
  }, {
    "selector" : "node[BIOPAX_TYPE = 'ComplexAssembly']",
    "css" : {
      "height" : 13.4
    }
  }, {
    "selector" : "node[BIOPAX_TYPE = 'Complex']",
    "css" : {
      "height" : 13.4
    }
  }, {
    "selector" : "node[BIOPAX_TYPE = 'Degradation']",
    "css" : {
      "height" : 13.4
    }
  }, {
    "selector" : "node[BIOPAX_TYPE = 'TemplateReactionRegulation']",
    "css" : {
      "height" : 13.4
    }
  }, {
    "selector" : "node[BIOPAX_TYPE = 'Control']",
    "css" : {
      "height" : 13.4
    }
  }, {
    "selector" : "node[BIOPAX_TYPE = 'Modulation']",
    "css" : {
      "height" : 13.4
    }
  }, {
    "selector" : "node[BIOPAX_TYPE = 'TemplateReaction']",
    "css" : {
      "height" : 13.4
    }
  }, {
    "selector" : "node[BIOPAX_TYPE = 'MolecularInteraction']",
    "css" : {
      "height" : 13.4
    }
  }, {
    "selector" : "node[BIOPAX_TYPE = 'Catalysis']",
    "css" : {
      "height" : 13.4
    }
  }, {
    "selector" : "node[BIOPAX_TYPE = 'Transport']",
    "css" : {
      "height" : 13.4
    }
  }, {
    "selector" : "node:selected",
    "css" : {
      "background-color" : "rgb(255,255,0)"
    }
  }, {
    "selector" : "edge",
    "css" : {
      "font-size" : 10,
      "line-color" : "rgb(64,64,64)",
      "opacity" : 1.0,
      "source-arrow-shape" : "none",
      "target-arrow-color" : "rgb(64,64,64)",
      "text-opacity" : 1.0,
      "line-style" : "solid",
      "color" : "rgb(0,0,0)",
      "font-family" : "SansSerif.plain",
      "font-weight" : "normal",
      "content" : "",
      "source-arrow-color" : "rgb(64,64,64)",
      "target-arrow-shape" : "none",
      "width" : 1.0
    }
  }, {
    "selector" : "edge[interaction = 'INHIBITION_NONCOMPETITIVE']",
    "css" : {
      "target-arrow-shape" : "tee"
    }
  }, {
    "selector" : "edge[interaction = 'INHIBITION_OTHER']",
    "css" : {
      "target-arrow-shape" : "tee"
    }
  }, {
    "selector" : "edge[interaction = 'ACTIVATION']",
    "css" : {
      "target-arrow-shape" : "triangle"
    }
  }, {
    "selector" : "edge[interaction = 'INHIBITION_UNCOMPETITIVE']",
    "css" : {
      "target-arrow-shape" : "tee"
    }
  }, {
    "selector" : "edge[interaction = 'cofactor']",
    "css" : {
      "target-arrow-shape" : "triangle"
    }
  }, {
    "selector" : "edge[interaction = 'ACTIVATION_ALLOSTERIC']",
    "css" : {
      "target-arrow-shape" : "triangle"
    }
  }, {
    "selector" : "edge[interaction = 'right']",
    "css" : {
      "target-arrow-shape" : "triangle"
    }
  }, {
    "selector" : "edge[interaction = 'INHIBITION_ALLOSTERIC']",
    "css" : {
      "target-arrow-shape" : "tee"
    }
  }, {
    "selector" : "edge[interaction = 'controlled']",
    "css" : {
      "target-arrow-shape" : "triangle"
    }
  }, {
    "selector" : "edge[interaction = 'contains']",
    "css" : {
      "target-arrow-shape" : "circle"
    }
  }, {
    "selector" : "edge[interaction = 'INHIBITION']",
    "css" : {
      "target-arrow-shape" : "tee"
    }
  }, {
    "selector" : "edge[interaction = 'INHIBITION_UNKMECH']",
    "css" : {
      "target-arrow-shape" : "tee"
    }
  }, {
    "selector" : "edge[interaction = 'INHIBITION_IRREVERSIBLE']",
    "css" : {
      "target-arrow-shape" : "tee"
    }
  }, {
    "selector" : "edge[interaction = 'INHIBITION_COMPETITIVE']",
    "css" : {
      "target-arrow-shape" : "tee"
    }
  }, {
    "selector" : "edge[interaction = 'ACTIVATION_UNKMECH']",
    "css" : {
      "target-arrow-shape" : "triangle"
    }
  }, {
    "selector" : "edge[interaction = 'ACTIVATION_NONALLOSTERIC']",
    "css" : {
      "target-arrow-shape" : "triangle"
    }
  }, {
    "selector" : "edge:selected",
    "css" : {
      "line-color" : "rgb(255,0,0)"
    }
  } ]
}, {
  "format_version" : "1.0",
  "generated_by" : "cytoscape-3.7.2",
  "target_cytoscapejs_version" : "~2.1",
  "title" : "ISMAGS",
  "style" : [ {
    "selector" : "node",
    "css" : {
      "text-opacity" : 1.0,
      "background-opacity" : 1.0,
      "border-width" : 1.0,
      "color" : "rgb(0,0,0)",
      "border-color" : "rgb(62,62,62)",
      "font-size" : 12,
      "font-family" : "SansSerif.plain",
      "font-weight" : "normal",
      "text-valign" : "center",
      "text-halign" : "center",
      "border-opacity" : 1.0,
      "shape" : "roundrectangle",
      "width" : 60.0,
      "background-color" : "rgb(128,128,128)",
      "height" : 40.0,
      "content" : "data(name)"
    }
  }, {
    "selector" : "node:selected",
    "css" : {
      "background-color" : "rgb(255,255,0)"
    }
  }, {
    "selector" : "edge",
    "css" : {
      "font-size" : 10,
      "line-color" : "rgb(64,64,64)",
      "opacity" : 1.0,
      "source-arrow-shape" : "none",
      "target-arrow-color" : "rgb(0,0,0)",
      "text-opacity" : 1.0,
      "line-style" : "solid",
      "color" : "rgb(0,0,0)",
      "font-family" : "SansSerif.plain",
      "font-weight" : "normal",
      "source-arrow-color" : "rgb(0,0,0)",
      "target-arrow-shape" : "none",
      "width" : 2.0,
      "content" : "data(interaction)"
    }
  }, {
    "selector" : "edge:selected",
    "css" : {
      "line-color" : "rgb(255,0,0)"
    }
  } ]
}, {
  "format_version" : "1.0",
  "generated_by" : "cytoscape-3.7.2",
  "target_cytoscapejs_version" : "~2.1",
  "title" : "size_rank_0_0_0",
  "style" : [ {
    "selector" : "node",
    "css" : {
      "text-opacity" : 1.0,
      "background-opacity" : 1.0,
      "border-width" : 0.0,
      "color" : "rgb(51,51,51)",
      "border-color" : "rgb(0,0,0)",
      "font-size" : 9,
      "font-family" : "Dialog.plain",
      "font-weight" : "normal",
      "text-valign" : "center",
      "text-halign" : "center",
      "border-opacity" : 1.0,
      "shape" : "rectangle",
      "width" : 12.0,
      "background-color" : "rgb(204,204,255)",
      "height" : 12.0,
      "content" : "data(name)"
    }
  }, {
    "selector" : "node:selected",
    "css" : {
      "background-color" : "rgb(255,255,0)"
    }
  }, {
    "selector" : "edge",
    "css" : {
      "font-size" : 10,
      "line-color" : "rgb(76,76,76)",
      "opacity" : 1.0,
      "source-arrow-shape" : "none",
      "target-arrow-color" : "rgb(0,0,0)",
      "text-opacity" : 1.0,
      "line-style" : "solid",
      "color" : "rgb(0,0,0)",
      "font-family" : "Dialog.plain",
      "font-weight" : "normal",
      "content" : "",
      "source-arrow-color" : "rgb(0,0,0)",
      "target-arrow-shape" : "none",
      "width" : 2.0
    }
  }, {
    "selector" : "edge:selected",
    "css" : {
      "line-color" : "rgb(255,0,0)"
    }
  } ]
}, {
  "format_version" : "1.0",
  "generated_by" : "cytoscape-3.7.2",
  "target_cytoscapejs_version" : "~2.1",
  "title" : "ISMAGS_1",
  "style" : [ {
    "selector" : "node",
    "css" : {
      "text-opacity" : 1.0,
      "background-opacity" : 1.0,
      "border-width" : 1.0,
      "color" : "rgb(0,0,0)",
      "border-color" : "rgb(62,62,62)",
      "font-size" : 12,
      "font-family" : "SansSerif.plain",
      "font-weight" : "normal",
      "text-valign" : "center",
      "text-halign" : "center",
      "border-opacity" : 1.0,
      "shape" : "roundrectangle",
      "width" : 60.0,
      "background-color" : "rgb(128,128,128)",
      "height" : 40.0,
      "content" : "data(name)"
    }
  }, {
    "selector" : "node:selected",
    "css" : {
      "background-color" : "rgb(255,255,0)"
    }
  }, {
    "selector" : "edge",
    "css" : {
      "font-size" : 10,
      "line-color" : "rgb(64,64,64)",
      "opacity" : 1.0,
      "source-arrow-shape" : "none",
      "target-arrow-color" : "rgb(0,0,0)",
      "text-opacity" : 1.0,
      "line-style" : "solid",
      "color" : "rgb(0,0,0)",
      "font-family" : "SansSerif.plain",
      "font-weight" : "normal",
      "source-arrow-color" : "rgb(0,0,0)",
      "target-arrow-shape" : "none",
      "width" : 2.0,
      "content" : "data(interaction)"
    }
  }, {
    "selector" : "edge:selected",
    "css" : {
      "line-color" : "rgb(255,0,0)"
    }
  } ]
}, {
  "format_version" : "1.0",
  "generated_by" : "cytoscape-3.7.2",
  "target_cytoscapejs_version" : "~2.1",
  "title" : "bingo Style for Sheet 1",
  "style" : [ {
    "selector" : "node",
    "css" : {
      "text-opacity" : 1.0,
      "background-opacity" : 0.8235294117647058,
      "border-width" : 2.0,
      "color" : "rgb(0,0,0)",
      "border-color" : "rgb(64,64,64)",
      "font-size" : 11,
      "font-family" : "Dialog.plain",
      "font-weight" : "normal",
      "text-valign" : "center",
      "text-halign" : "center",
      "border-opacity" : 1.0,
      "shape" : "ellipse",
      "width" : 40.0,
      "background-color" : "rgb(200,0,0)",
      "height" : 40.0,
      "content" : "data(description_Sheet_1)"
    }
  }, {
    "selector" : "node[nodeFillColor_Sheet_1 > 6.30103]",
    "css" : {
      "background-color" : "rgb(255,127,0)"
    }
  }, {
    "selector" : "node[nodeFillColor_Sheet_1 = 6.30103]",
    "css" : {
      "background-color" : "rgb(255,127,0)"
    }
  }, {
    "selector" : "node[nodeFillColor_Sheet_1 > 1.30103][nodeFillColor_Sheet_1 < 6.30103]",
    "css" : {
      "background-color" : "mapData(nodeFillColor_Sheet_1,1.30103,6.30103,rgb(255,255,0),rgb(255,127,0))"
    }
  }, {
    "selector" : "node[nodeFillColor_Sheet_1 = 1.30103]",
    "css" : {
      "background-color" : "rgb(255,255,0)"
    }
  }, {
    "selector" : "node[nodeFillColor_Sheet_1 < 1.30103]",
    "css" : {
      "background-color" : "rgb(255,255,255)"
    }
  }, {
    "selector" : "node[nodeSize_Sheet_1 > 100]",
    "css" : {
      "width" : 400.0
    }
  }, {
    "selector" : "node[nodeSize_Sheet_1 = 100]",
    "css" : {
      "width" : 400.0
    }
  }, {
    "selector" : "node[nodeSize_Sheet_1 > 1][nodeSize_Sheet_1 < 100]",
    "css" : {
      "width" : "mapData(nodeSize_Sheet_1,1,100,20.0,400.0)"
    }
  }, {
    "selector" : "node[nodeSize_Sheet_1 = 1]",
    "css" : {
      "width" : 20.0
    }
  }, {
    "selector" : "node[nodeSize_Sheet_1 < 1]",
    "css" : {
      "width" : 20.0
    }
  }, {
    "selector" : "node[nodeSize_Sheet_1 > 100]",
    "css" : {
      "height" : 400.0
    }
  }, {
    "selector" : "node[nodeSize_Sheet_1 = 100]",
    "css" : {
      "height" : 400.0
    }
  }, {
    "selector" : "node[nodeSize_Sheet_1 > 1][nodeSize_Sheet_1 < 100]",
    "css" : {
      "height" : "mapData(nodeSize_Sheet_1,1,100,20.0,400.0)"
    }
  }, {
    "selector" : "node[nodeSize_Sheet_1 = 1]",
    "css" : {
      "height" : 20.0
    }
  }, {
    "selector" : "node[nodeSize_Sheet_1 < 1]",
    "css" : {
      "height" : 20.0
    }
  }, {
    "selector" : "node:selected",
    "css" : {
      "background-color" : "rgb(255,255,0)"
    }
  }, {
    "selector" : "edge",
    "css" : {
      "font-size" : 10,
      "line-color" : "rgb(64,64,64)",
      "opacity" : 0.7450980392156863,
      "source-arrow-shape" : "none",
      "target-arrow-color" : "rgb(0,0,0)",
      "text-opacity" : 1.0,
      "line-style" : "solid",
      "color" : "rgb(0,0,0)",
      "font-family" : "Dialog.plain",
      "font-weight" : "normal",
      "content" : "",
      "source-arrow-color" : "rgb(0,0,0)",
      "target-arrow-shape" : "triangle",
      "width" : 2.0
    }
  }, {
    "selector" : "edge:selected",
    "css" : {
      "line-color" : "rgb(255,0,0)"
    }
  } ]
}, {
  "format_version" : "1.0",
  "generated_by" : "cytoscape-3.7.2",
  "target_cytoscapejs_version" : "~2.1",
  "title" : "size_rank_0_1_0",
  "style" : [ {
    "selector" : "node",
    "css" : {
      "text-opacity" : 1.0,
      "background-opacity" : 1.0,
      "border-width" : 0.0,
      "color" : "rgb(51,51,51)",
      "border-color" : "rgb(0,0,0)",
      "font-size" : 9,
      "font-family" : "Dialog.plain",
      "font-weight" : "normal",
      "text-valign" : "center",
      "text-halign" : "center",
      "border-opacity" : 1.0,
      "shape" : "rectangle",
      "width" : 12.0,
      "background-color" : "rgb(204,204,255)",
      "height" : 12.0,
      "content" : "data(name)"
    }
  }, {
    "selector" : "node:selected",
    "css" : {
      "background-color" : "rgb(255,255,0)"
    }
  }, {
    "selector" : "edge",
    "css" : {
      "font-size" : 10,
      "line-color" : "rgb(76,76,76)",
      "opacity" : 1.0,
      "source-arrow-shape" : "none",
      "target-arrow-color" : "rgb(0,0,0)",
      "text-opacity" : 1.0,
      "line-style" : "solid",
      "color" : "rgb(0,0,0)",
      "font-family" : "Dialog.plain",
      "font-weight" : "normal",
      "content" : "",
      "source-arrow-color" : "rgb(0,0,0)",
      "target-arrow-shape" : "none",
      "width" : 2.0
    }
  }, {
    "selector" : "edge:selected",
    "css" : {
      "line-color" : "rgb(255,0,0)"
    }
  } ]
}, {
  "format_version" : "1.0",
  "generated_by" : "cytoscape-3.7.2",
  "target_cytoscapejs_version" : "~2.1",
  "title" : "Diffany - Source",
  "style" : [ {
    "selector" : "node",
    "css" : {
      "text-opacity" : 1.0,
      "background-opacity" : 1.0,
      "border-width" : 0.0,
      "color" : "rgb(0,0,0)",
      "border-color" : "rgb(0,0,0)",
      "font-size" : 11,
      "font-family" : "SansSerif.plain",
      "font-weight" : "normal",
      "text-valign" : "center",
      "text-halign" : "center",
      "border-opacity" : 1.0,
      "shape" : "ellipse",
      "width" : 80.0,
      "background-color" : "rgb(128,128,128)",
      "height" : 55.0,
      "content" : "data(official_symbol)"
    }
  }, {
    "selector" : "node[differentially_expressed = 'up-regulated']",
    "css" : {
      "background-color" : "rgb(124,124,7)"
    }
  }, {
    "selector" : "node[differentially_expressed = 'down-regulated']",
    "css" : {
      "background-color" : "rgb(30,88,124)"
    }
  }, {
    "selector" : "node[kinase_function = 'yes']",
    "css" : {
      "border-width" : 2.0
    }
  }, {
    "selector" : "node[differentially_expressed = 'up-regulated']",
    "css" : {
      "background-color" : "rgb(255,255,15)"
    }
  }, {
    "selector" : "node[differentially_expressed = 'down-regulated']",
    "css" : {
      "background-color" : "rgb(62,181,255)"
    }
  }, {
    "selector" : "node[phosphorylation_site = 'yes']",
    "css" : {
      "shape" : "diamond"
    }
  }, {
    "selector" : "node[kinase_function = 'yes']",
    "css" : {
      "border-color" : "rgb(0,0,0)"
    }
  }, {
    "selector" : "node:selected",
    "css" : {
      "background-color" : "rgb(62,62,62)"
    }
  }, {
    "selector" : "edge",
    "css" : {
      "font-size" : 10,
      "line-color" : "rgb(64,64,64)",
      "opacity" : 1.0,
      "source-arrow-shape" : "none",
      "target-arrow-color" : "rgb(0,0,0)",
      "text-opacity" : 1.0,
      "line-style" : "solid",
      "color" : "rgb(0,0,0)",
      "font-family" : "SansSerif.plain",
      "font-weight" : "normal",
      "content" : "",
      "source-arrow-color" : "rgb(0,0,0)",
      "target-arrow-shape" : "none",
      "width" : 1.0
    }
  }, {
    "selector" : "edge[negated]",
    "css" : {
      "line-style" : "dotted"
    }
  }, {
    "selector" : "edge[!negated]",
    "css" : {
      "line-style" : "solid"
    }
  }, {
    "selector" : "edge[weight > 20]",
    "css" : {
      "width" : 25.0
    }
  }, {
    "selector" : "edge[weight = 20]",
    "css" : {
      "width" : 25.0
    }
  }, {
    "selector" : "edge[weight > 0][weight < 20]",
    "css" : {
      "width" : "mapData(weight,0,20,1.0,25.0)"
    }
  }, {
    "selector" : "edge[weight = 0]",
    "css" : {
      "width" : 1.0
    }
  }, {
    "selector" : "edge[weight < 0]",
    "css" : {
      "width" : 1.0
    }
  }, {
    "selector" : "edge:selected",
    "css" : {
      "line-color" : "rgb(255,0,0)"
    }
  } ]
}, {
  "format_version" : "1.0",
  "generated_by" : "cytoscape-3.7.2",
  "target_cytoscapejs_version" : "~2.1",
  "title" : "Sample2",
  "style" : [ {
    "selector" : "node",
    "css" : {
      "text-opacity" : 1.0,
      "background-opacity" : 1.0,
      "border-width" : 15.0,
      "color" : "rgb(102,102,102)",
      "border-color" : "rgb(255,255,255)",
      "font-size" : 20,
      "font-family" : "Dialog.plain",
      "font-weight" : "normal",
      "text-valign" : "center",
      "text-halign" : "right",
      "border-opacity" : 1.0,
      "shape" : "ellipse",
      "width" : 50.0,
      "background-color" : "rgb(58,127,182)",
      "height" : 50.0,
      "content" : "data(name)"
    }
  }, {
    "selector" : "node:selected",
    "css" : {
      "background-color" : "rgb(255,255,0)"
    }
  }, {
    "selector" : "edge",
    "css" : {
      "font-size" : 10,
      "line-color" : "rgb(255,255,255)",
      "opacity" : 1.0,
      "source-arrow-shape" : "none",
      "target-arrow-color" : "rgb(0,0,0)",
      "text-opacity" : 1.0,
      "line-style" : "solid",
      "color" : "rgb(0,0,0)",
      "font-family" : "Dialog.plain",
      "font-weight" : "normal",
      "content" : "",
      "source-arrow-color" : "rgb(0,0,0)",
      "target-arrow-shape" : "none",
      "width" : 20.0
    }
  }, {
    "selector" : "edge:selected",
    "css" : {
      "line-color" : "rgb(255,0,0)"
    }
  } ]
}, {
  "format_version" : "1.0",
  "generated_by" : "cytoscape-3.7.2",
  "target_cytoscapejs_version" : "~2.1",
  "title" : "Diffany - Differential_0",
  "style" : [ {
    "selector" : "node",
    "css" : {
      "text-opacity" : 1.0,
      "background-opacity" : 1.0,
      "border-width" : 0.0,
      "color" : "rgb(0,0,0)",
      "border-color" : "rgb(0,0,0)",
      "font-size" : 11,
      "font-family" : "SansSerif.plain",
      "font-weight" : "normal",
      "text-valign" : "center",
      "text-halign" : "center",
      "border-opacity" : 1.0,
      "shape" : "ellipse",
      "width" : 80.0,
      "background-color" : "rgb(128,128,128)",
      "height" : 55.0,
      "content" : "data(official_symbol)"
    }
  }, {
    "selector" : "node[differentially_expressed = 'up-regulated']",
    "css" : {
      "background-color" : "rgb(124,124,7)"
    }
  }, {
    "selector" : "node[differentially_expressed = 'down-regulated']",
    "css" : {
      "background-color" : "rgb(30,88,124)"
    }
  }, {
    "selector" : "node[kinase_function = 'yes']",
    "css" : {
      "border-width" : 2.0
    }
  }, {
    "selector" : "node[differentially_expressed = 'up-regulated']",
    "css" : {
      "background-color" : "rgb(255,255,15)"
    }
  }, {
    "selector" : "node[differentially_expressed = 'down-regulated']",
    "css" : {
      "background-color" : "rgb(62,181,255)"
    }
  }, {
    "selector" : "node[phosphorylation_site = 'yes']",
    "css" : {
      "shape" : "diamond"
    }
  }, {
    "selector" : "node[kinase_function = 'yes']",
    "css" : {
      "border-color" : "rgb(0,0,0)"
    }
  }, {
    "selector" : "node:selected",
    "css" : {
      "background-color" : "rgb(62,62,62)"
    }
  }, {
    "selector" : "edge",
    "css" : {
      "font-size" : 10,
      "line-color" : "rgb(64,64,64)",
      "opacity" : 1.0,
      "source-arrow-shape" : "none",
      "target-arrow-color" : "rgb(0,0,0)",
      "text-opacity" : 1.0,
      "line-style" : "solid",
      "color" : "rgb(0,0,0)",
      "font-family" : "SansSerif.plain",
      "font-weight" : "normal",
      "content" : "",
      "source-arrow-color" : "rgb(0,0,0)",
      "target-arrow-shape" : "none",
      "width" : 1.0
    }
  }, {
    "selector" : "edge[negated]",
    "css" : {
      "line-style" : "dotted"
    }
  }, {
    "selector" : "edge[!negated]",
    "css" : {
      "line-style" : "solid"
    }
  }, {
    "selector" : "edge[weight > 20]",
    "css" : {
      "width" : 25.0
    }
  }, {
    "selector" : "edge[weight = 20]",
    "css" : {
      "width" : 25.0
    }
  }, {
    "selector" : "edge[weight > 0][weight < 20]",
    "css" : {
      "width" : "mapData(weight,0,20,1.0,25.0)"
    }
  }, {
    "selector" : "edge[weight = 0]",
    "css" : {
      "width" : 1.0
    }
  }, {
    "selector" : "edge[weight < 0]",
    "css" : {
      "width" : 1.0
    }
  }, {
    "selector" : "edge:selected",
    "css" : {
      "line-color" : "rgb(255,0,0)"
    }
  } ]
}, {
  "format_version" : "1.0",
  "generated_by" : "cytoscape-3.7.2",
  "target_cytoscapejs_version" : "~2.1",
  "title" : "Diffany - Differential_1",
  "style" : [ {
    "selector" : "node",
    "css" : {
      "text-opacity" : 1.0,
      "background-opacity" : 1.0,
      "border-width" : 0.0,
      "color" : "rgb(0,0,0)",
      "border-color" : "rgb(0,0,0)",
      "font-size" : 11,
      "font-family" : "SansSerif.plain",
      "font-weight" : "normal",
      "text-valign" : "center",
      "text-halign" : "center",
      "border-opacity" : 1.0,
      "shape" : "ellipse",
      "width" : 80.0,
      "background-color" : "rgb(128,128,128)",
      "height" : 55.0,
      "content" : "data(official_symbol)"
    }
  }, {
    "selector" : "node[differentially_expressed = 'up-regulated']",
    "css" : {
      "background-color" : "rgb(124,124,7)"
    }
  }, {
    "selector" : "node[differentially_expressed = 'down-regulated']",
    "css" : {
      "background-color" : "rgb(30,88,124)"
    }
  }, {
    "selector" : "node[kinase_function = 'yes']",
    "css" : {
      "border-width" : 2.0
    }
  }, {
    "selector" : "node[differentially_expressed = 'up-regulated']",
    "css" : {
      "background-color" : "rgb(255,255,15)"
    }
  }, {
    "selector" : "node[differentially_expressed = 'down-regulated']",
    "css" : {
      "background-color" : "rgb(62,181,255)"
    }
  }, {
    "selector" : "node[phosphorylation_site = 'yes']",
    "css" : {
      "shape" : "diamond"
    }
  }, {
    "selector" : "node[kinase_function = 'yes']",
    "css" : {
      "border-color" : "rgb(0,0,0)"
    }
  }, {
    "selector" : "node:selected",
    "css" : {
      "background-color" : "rgb(62,62,62)"
    }
  }, {
    "selector" : "edge",
    "css" : {
      "font-size" : 10,
      "line-color" : "rgb(64,64,64)",
      "opacity" : 1.0,
      "source-arrow-shape" : "none",
      "target-arrow-color" : "rgb(0,0,0)",
      "text-opacity" : 1.0,
      "line-style" : "solid",
      "color" : "rgb(0,0,0)",
      "font-family" : "SansSerif.plain",
      "font-weight" : "normal",
      "content" : "",
      "source-arrow-color" : "rgb(0,0,0)",
      "target-arrow-shape" : "none",
      "width" : 1.0
    }
  }, {
    "selector" : "edge[negated]",
    "css" : {
      "line-style" : "dotted"
    }
  }, {
    "selector" : "edge[!negated]",
    "css" : {
      "line-style" : "solid"
    }
  }, {
    "selector" : "edge[weight > 20]",
    "css" : {
      "width" : 25.0
    }
  }, {
    "selector" : "edge[weight = 20]",
    "css" : {
      "width" : 25.0
    }
  }, {
    "selector" : "edge[weight > 0][weight < 20]",
    "css" : {
      "width" : "mapData(weight,0,20,1.0,25.0)"
    }
  }, {
    "selector" : "edge[weight = 0]",
    "css" : {
      "width" : 1.0
    }
  }, {
    "selector" : "edge[weight < 0]",
    "css" : {
      "width" : 1.0
    }
  }, {
    "selector" : "edge:selected",
    "css" : {
      "line-color" : "rgb(255,0,0)"
    }
  } ]
}, {
  "format_version" : "1.0",
  "generated_by" : "cytoscape-3.7.2",
  "target_cytoscapejs_version" : "~2.1",
  "title" : "Sample1",
  "style" : [ {
    "selector" : "node",
    "css" : {
      "text-opacity" : 1.0,
      "background-opacity" : 1.0,
      "border-width" : 0.0,
      "color" : "rgb(51,51,51)",
      "border-color" : "rgb(0,0,0)",
      "font-size" : 10,
      "font-family" : "Dialog.plain",
      "font-weight" : "normal",
      "text-valign" : "center",
      "text-halign" : "center",
      "border-opacity" : 1.0,
      "shape" : "ellipse",
      "width" : 25.0,
      "background-color" : "rgb(127,205,187)",
      "height" : 25.0,
      "content" : "data(name)"
    }
  }, {
    "selector" : "node:selected",
    "css" : {
      "background-color" : "rgb(255,255,0)"
    }
  }, {
    "selector" : "edge",
    "css" : {
      "font-size" : 10,
      "line-color" : "rgb(153,153,153)",
      "opacity" : 1.0,
      "source-arrow-shape" : "none",
      "target-arrow-color" : "rgb(0,0,0)",
      "text-opacity" : 1.0,
      "line-style" : "solid",
      "color" : "rgb(51,51,51)",
      "font-family" : "Dialog.plain",
      "font-weight" : "normal",
      "source-arrow-color" : "rgb(0,0,0)",
      "target-arrow-shape" : "none",
      "width" : 1.0,
      "content" : "data(interaction)"
    }
  }, {
    "selector" : "edge[interaction = 'pp']",
    "css" : {
      "line-style" : "solid"
    }
  }, {
    "selector" : "edge[interaction = 'pd']",
    "css" : {
      "line-style" : "dashed"
    }
  }, {
    "selector" : "edge:selected",
    "css" : {
      "line-color" : "rgb(255,0,0)"
    }
  } ]
}, {
  "format_version" : "1.0",
  "generated_by" : "cytoscape-3.7.2",
  "target_cytoscapejs_version" : "~2.1",
  "title" : "size_rank",
  "style" : [ {
    "selector" : "node",
    "css" : {
      "text-opacity" : 1.0,
      "background-opacity" : 1.0,
      "border-width" : 0.0,
      "color" : "rgb(51,51,51)",
      "border-color" : "rgb(0,0,0)",
      "font-size" : 9,
      "font-family" : "SansSerif.plain",
      "font-weight" : "normal",
      "text-valign" : "center",
      "text-halign" : "center",
      "border-opacity" : 1.0,
      "shape" : "rectangle",
      "width" : 12.0,
      "background-color" : "rgb(204,204,255)",
      "height" : 12.0,
      "content" : "data(name)"
    }
  }, {
    "selector" : "node:selected",
    "css" : {
      "background-color" : "rgb(255,255,0)"
    }
  }, {
    "selector" : "edge",
    "css" : {
      "font-size" : 10,
      "line-color" : "rgb(76,76,76)",
      "opacity" : 1.0,
      "source-arrow-shape" : "none",
      "target-arrow-color" : "rgb(0,0,0)",
      "text-opacity" : 1.0,
      "line-style" : "solid",
      "color" : "rgb(0,0,0)",
      "font-family" : "SansSerif.plain",
      "font-weight" : "normal",
      "content" : "",
      "source-arrow-color" : "rgb(0,0,0)",
      "target-arrow-shape" : "none",
      "width" : 2.0
    }
  }, {
    "selector" : "edge:selected",
    "css" : {
      "line-color" : "rgb(255,0,0)"
    }
  } ]
}, {
  "format_version" : "1.0",
  "generated_by" : "cytoscape-3.7.2",
  "target_cytoscapejs_version" : "~2.1",
  "title" : "size_rank_0_2_0",
  "style" : [ {
    "selector" : "node",
    "css" : {
      "text-opacity" : 1.0,
      "background-opacity" : 1.0,
      "border-width" : 0.0,
      "color" : "rgb(51,51,51)",
      "border-color" : "rgb(0,0,0)",
      "font-size" : 9,
      "font-family" : "Dialog.plain",
      "font-weight" : "normal",
      "text-valign" : "center",
      "text-halign" : "center",
      "border-opacity" : 1.0,
      "shape" : "rectangle",
      "width" : 12.0,
      "background-color" : "rgb(204,204,255)",
      "height" : 12.0,
      "content" : "data(name)"
    }
  }, {
    "selector" : "node:selected",
    "css" : {
      "background-color" : "rgb(255,255,0)"
    }
  }, {
    "selector" : "edge",
    "css" : {
      "font-size" : 10,
      "line-color" : "rgb(76,76,76)",
      "opacity" : 1.0,
      "source-arrow-shape" : "none",
      "target-arrow-color" : "rgb(0,0,0)",
      "text-opacity" : 1.0,
      "line-style" : "solid",
      "color" : "rgb(0,0,0)",
      "font-family" : "Dialog.plain",
      "font-weight" : "normal",
      "content" : "",
      "source-arrow-color" : "rgb(0,0,0)",
      "target-arrow-shape" : "none",
      "width" : 2.0
    }
  }, {
    "selector" : "edge:selected",
    "css" : {
      "line-color" : "rgb(255,0,0)"
    }
  } ]
}, {
  "format_version" : "1.0",
  "generated_by" : "cytoscape-3.7.2",
  "target_cytoscapejs_version" : "~2.1",
  "title" : "Nested Network Style",
  "style" : [ {
    "selector" : "node",
    "css" : {
      "text-opacity" : 1.0,
      "background-opacity" : 1.0,
      "border-width" : 2.0,
      "color" : "rgb(0,0,0)",
      "border-color" : "rgb(0,0,0)",
      "font-size" : 12,
      "font-family" : "Dialog.plain",
      "font-weight" : "normal",
      "text-valign" : "center",
      "text-halign" : "center",
      "border-opacity" : 1.0,
      "shape" : "ellipse",
      "width" : 60.0,
      "background-color" : "rgb(255,255,255)",
      "height" : 40.0,
      "content" : "data(shared_name)"
    }
  }, {
    "selector" : "node[has_nested_network]",
    "css" : {
      "text-valign" : "bottom"
    }
  }, {
    "selector" : "node[has_nested_network]",
    "css" : {
      "color" : "rgb(0,102,204)"
    }
  }, {
    "selector" : "node[has_nested_network]",
    "css" : {
      "background-color" : "rgb(255,255,255)"
    }
  }, {
    "selector" : "node[has_nested_network]",
    "css" : {
      "shape" : "rectangle"
    }
  }, {
    "selector" : "node[has_nested_network]",
    "css" : {
      "border-color" : "rgb(0,102,204)"
    }
  }, {
    "selector" : "node:selected",
    "css" : {
      "background-color" : "rgb(255,255,0)"
    }
  }, {
    "selector" : "edge",
    "css" : {
      "font-size" : 10,
      "line-color" : "rgb(64,64,64)",
      "opacity" : 1.0,
      "source-arrow-shape" : "none",
      "target-arrow-color" : "rgb(0,0,0)",
      "text-opacity" : 1.0,
      "line-style" : "solid",
      "color" : "rgb(0,0,0)",
      "font-family" : "Dialog.plain",
      "font-weight" : "normal",
      "content" : "",
      "source-arrow-color" : "rgb(0,0,0)",
      "target-arrow-shape" : "none",
      "width" : 1.0
    }
  }, {
    "selector" : "edge:selected",
    "css" : {
      "line-color" : "rgb(255,0,0)"
    }
  } ]
}, {
  "format_version" : "1.0",
  "generated_by" : "cytoscape-3.7.2",
  "target_cytoscapejs_version" : "~2.1",
  "title" : "bingo Style for NRT3.1_0",
  "style" : [ {
    "selector" : "node",
    "css" : {
      "text-opacity" : 1.0,
      "background-opacity" : 0.8235294117647058,
      "border-width" : 2.0,
      "color" : "rgb(0,0,0)",
      "border-color" : "rgb(64,64,64)",
      "font-size" : 11,
      "font-family" : "Dialog.plain",
      "font-weight" : "normal",
      "text-valign" : "center",
      "text-halign" : "center",
      "border-opacity" : 1.0,
      "shape" : "ellipse",
      "width" : 40.0,
      "background-color" : "rgb(200,0,0)",
      "height" : 40.0,
      "content" : "data(description_NRT3_1)"
    }
  }, {
    "selector" : "node[nodeFillColor_NRT3_1 > 6.30103]",
    "css" : {
      "background-color" : "rgb(255,127,0)"
    }
  }, {
    "selector" : "node[nodeFillColor_NRT3_1 = 6.30103]",
    "css" : {
      "background-color" : "rgb(255,127,0)"
    }
  }, {
    "selector" : "node[nodeFillColor_NRT3_1 > 1.30103][nodeFillColor_NRT3_1 < 6.30103]",
    "css" : {
      "background-color" : "mapData(nodeFillColor_NRT3_1,1.30103,6.30103,rgb(255,255,0),rgb(255,127,0))"
    }
  }, {
    "selector" : "node[nodeFillColor_NRT3_1 = 1.30103]",
    "css" : {
      "background-color" : "rgb(255,255,0)"
    }
  }, {
    "selector" : "node[nodeFillColor_NRT3_1 < 1.30103]",
    "css" : {
      "background-color" : "rgb(255,255,255)"
    }
  }, {
    "selector" : "node[nodeSize_NRT3_1 > 100]",
    "css" : {
      "width" : 400.0
    }
  }, {
    "selector" : "node[nodeSize_NRT3_1 = 100]",
    "css" : {
      "width" : 400.0
    }
  }, {
    "selector" : "node[nodeSize_NRT3_1 > 1][nodeSize_NRT3_1 < 100]",
    "css" : {
      "width" : "mapData(nodeSize_NRT3_1,1,100,20.0,400.0)"
    }
  }, {
    "selector" : "node[nodeSize_NRT3_1 = 1]",
    "css" : {
      "width" : 20.0
    }
  }, {
    "selector" : "node[nodeSize_NRT3_1 < 1]",
    "css" : {
      "width" : 20.0
    }
  }, {
    "selector" : "node[nodeSize_NRT3_1 > 100]",
    "css" : {
      "height" : 400.0
    }
  }, {
    "selector" : "node[nodeSize_NRT3_1 = 100]",
    "css" : {
      "height" : 400.0
    }
  }, {
    "selector" : "node[nodeSize_NRT3_1 > 1][nodeSize_NRT3_1 < 100]",
    "css" : {
      "height" : "mapData(nodeSize_NRT3_1,1,100,20.0,400.0)"
    }
  }, {
    "selector" : "node[nodeSize_NRT3_1 = 1]",
    "css" : {
      "height" : 20.0
    }
  }, {
    "selector" : "node[nodeSize_NRT3_1 < 1]",
    "css" : {
      "height" : 20.0
    }
  }, {
    "selector" : "node:selected",
    "css" : {
      "background-color" : "rgb(255,255,0)"
    }
  }, {
    "selector" : "edge",
    "css" : {
      "font-size" : 10,
      "line-color" : "rgb(64,64,64)",
      "opacity" : 0.7450980392156863,
      "source-arrow-shape" : "none",
      "target-arrow-color" : "rgb(0,0,0)",
      "text-opacity" : 1.0,
      "line-style" : "solid",
      "color" : "rgb(0,0,0)",
      "font-family" : "Dialog.plain",
      "font-weight" : "normal",
      "content" : "",
      "source-arrow-color" : "rgb(0,0,0)",
      "target-arrow-shape" : "triangle",
      "width" : 2.0
    }
  }, {
    "selector" : "edge:selected",
    "css" : {
      "line-color" : "rgb(255,0,0)"
    }
  } ]
}, {
  "format_version" : "1.0",
  "generated_by" : "cytoscape-3.7.2",
  "target_cytoscapejs_version" : "~2.1",
  "title" : "bingo Style for NRT3.1",
  "style" : [ {
    "selector" : "node",
    "css" : {
      "text-opacity" : 1.0,
      "background-opacity" : 0.8235294117647058,
      "border-width" : 2.0,
      "color" : "rgb(0,0,0)",
      "border-color" : "rgb(64,64,64)",
      "font-size" : 11,
      "font-family" : "Dialog.plain",
      "font-weight" : "normal",
      "text-valign" : "center",
      "text-halign" : "center",
      "border-opacity" : 1.0,
      "shape" : "ellipse",
      "width" : 40.0,
      "background-color" : "rgb(200,0,0)",
      "height" : 40.0,
      "content" : "data(description_NRT3_1)"
    }
  }, {
    "selector" : "node[nodeFillColor_NRT3_1 > 6.30103]",
    "css" : {
      "background-color" : "rgb(255,127,0)"
    }
  }, {
    "selector" : "node[nodeFillColor_NRT3_1 = 6.30103]",
    "css" : {
      "background-color" : "rgb(255,127,0)"
    }
  }, {
    "selector" : "node[nodeFillColor_NRT3_1 > 1.30103][nodeFillColor_NRT3_1 < 6.30103]",
    "css" : {
      "background-color" : "mapData(nodeFillColor_NRT3_1,1.30103,6.30103,rgb(255,255,0),rgb(255,127,0))"
    }
  }, {
    "selector" : "node[nodeFillColor_NRT3_1 = 1.30103]",
    "css" : {
      "background-color" : "rgb(255,255,0)"
    }
  }, {
    "selector" : "node[nodeFillColor_NRT3_1 < 1.30103]",
    "css" : {
      "background-color" : "rgb(255,255,255)"
    }
  }, {
    "selector" : "node[nodeSize_NRT3_1 > 100]",
    "css" : {
      "width" : 400.0
    }
  }, {
    "selector" : "node[nodeSize_NRT3_1 = 100]",
    "css" : {
      "width" : 400.0
    }
  }, {
    "selector" : "node[nodeSize_NRT3_1 > 1][nodeSize_NRT3_1 < 100]",
    "css" : {
      "width" : "mapData(nodeSize_NRT3_1,1,100,20.0,400.0)"
    }
  }, {
    "selector" : "node[nodeSize_NRT3_1 = 1]",
    "css" : {
      "width" : 20.0
    }
  }, {
    "selector" : "node[nodeSize_NRT3_1 < 1]",
    "css" : {
      "width" : 20.0
    }
  }, {
    "selector" : "node[nodeSize_NRT3_1 > 100]",
    "css" : {
      "height" : 400.0
    }
  }, {
    "selector" : "node[nodeSize_NRT3_1 = 100]",
    "css" : {
      "height" : 400.0
    }
  }, {
    "selector" : "node[nodeSize_NRT3_1 > 1][nodeSize_NRT3_1 < 100]",
    "css" : {
      "height" : "mapData(nodeSize_NRT3_1,1,100,20.0,400.0)"
    }
  }, {
    "selector" : "node[nodeSize_NRT3_1 = 1]",
    "css" : {
      "height" : 20.0
    }
  }, {
    "selector" : "node[nodeSize_NRT3_1 < 1]",
    "css" : {
      "height" : 20.0
    }
  }, {
    "selector" : "node:selected",
    "css" : {
      "background-color" : "rgb(255,255,0)"
    }
  }, {
    "selector" : "edge",
    "css" : {
      "font-size" : 10,
      "line-color" : "rgb(64,64,64)",
      "opacity" : 0.7450980392156863,
      "source-arrow-shape" : "none",
      "target-arrow-color" : "rgb(0,0,0)",
      "text-opacity" : 1.0,
      "line-style" : "solid",
      "color" : "rgb(0,0,0)",
      "font-family" : "Dialog.plain",
      "font-weight" : "normal",
      "content" : "",
      "source-arrow-color" : "rgb(0,0,0)",
      "target-arrow-shape" : "triangle",
      "width" : 2.0
    }
  }, {
    "selector" : "edge:selected",
    "css" : {
      "line-color" : "rgb(255,0,0)"
    }
  } ]
}, {
  "format_version" : "1.0",
  "generated_by" : "cytoscape-3.7.2",
  "target_cytoscapejs_version" : "~2.1",
  "title" : "size_rank_0_2",
  "style" : [ {
    "selector" : "node",
    "css" : {
      "text-opacity" : 1.0,
      "background-opacity" : 1.0,
      "border-width" : 0.0,
      "color" : "rgb(51,51,51)",
      "border-color" : "rgb(0,0,0)",
      "font-size" : 9,
      "font-family" : "Dialog.plain",
      "font-weight" : "normal",
      "text-valign" : "center",
      "text-halign" : "center",
      "border-opacity" : 1.0,
      "shape" : "rectangle",
      "width" : 12.0,
      "background-color" : "rgb(204,204,255)",
      "height" : 12.0,
      "content" : "data(name)"
    }
  }, {
    "selector" : "node:selected",
    "css" : {
      "background-color" : "rgb(255,255,0)"
    }
  }, {
    "selector" : "edge",
    "css" : {
      "font-size" : 10,
      "line-color" : "rgb(76,76,76)",
      "opacity" : 1.0,
      "source-arrow-shape" : "none",
      "target-arrow-color" : "rgb(0,0,0)",
      "text-opacity" : 1.0,
      "line-style" : "solid",
      "color" : "rgb(0,0,0)",
      "font-family" : "Dialog.plain",
      "font-weight" : "normal",
      "content" : "",
      "source-arrow-color" : "rgb(0,0,0)",
      "target-arrow-shape" : "none",
      "width" : 2.0
    }
  }, {
    "selector" : "edge:selected",
    "css" : {
      "line-color" : "rgb(255,0,0)"
    }
  } ]
}, {
  "format_version" : "1.0",
  "generated_by" : "cytoscape-3.7.2",
  "target_cytoscapejs_version" : "~2.1",
  "title" : "ISMAGS_0",
  "style" : [ {
    "selector" : "node",
    "css" : {
      "text-opacity" : 1.0,
      "background-opacity" : 1.0,
      "border-width" : 1.0,
      "color" : "rgb(0,0,0)",
      "border-color" : "rgb(62,62,62)",
      "font-size" : 12,
      "font-family" : "SansSerif.plain",
      "font-weight" : "normal",
      "text-valign" : "center",
      "text-halign" : "center",
      "border-opacity" : 1.0,
      "shape" : "roundrectangle",
      "width" : 60.0,
      "background-color" : "rgb(128,128,128)",
      "height" : 40.0,
      "content" : "data(name)"
    }
  }, {
    "selector" : "node:selected",
    "css" : {
      "background-color" : "rgb(255,255,0)"
    }
  }, {
    "selector" : "edge",
    "css" : {
      "font-size" : 10,
      "line-color" : "rgb(64,64,64)",
      "opacity" : 1.0,
      "source-arrow-shape" : "none",
      "target-arrow-color" : "rgb(0,0,0)",
      "text-opacity" : 1.0,
      "line-style" : "solid",
      "color" : "rgb(0,0,0)",
      "font-family" : "SansSerif.plain",
      "font-weight" : "normal",
      "source-arrow-color" : "rgb(0,0,0)",
      "target-arrow-shape" : "none",
      "width" : 2.0,
      "content" : "data(interaction)"
    }
  }, {
    "selector" : "edge:selected",
    "css" : {
      "line-color" : "rgb(255,0,0)"
    }
  } ]
}, {
  "format_version" : "1.0",
  "generated_by" : "cytoscape-3.7.2",
  "target_cytoscapejs_version" : "~2.1",
  "title" : "Sample3",
  "style" : [ {
    "selector" : "node",
    "css" : {
      "text-opacity" : 1.0,
      "background-opacity" : 1.0,
      "border-width" : 8.0,
      "color" : "rgb(206,206,206)",
      "border-color" : "rgb(255,255,255)",
      "font-size" : 14,
      "font-family" : "Dialog.plain",
      "font-weight" : "normal",
      "text-valign" : "bottom",
      "text-halign" : "right",
      "border-opacity" : 1.0,
      "shape" : "ellipse",
      "width" : 20.0,
      "background-color" : "rgb(61,154,255)",
      "height" : 20.0,
      "content" : "data(name)"
    }
  }, {
    "selector" : "node:selected",
    "css" : {
      "background-color" : "rgb(255,255,0)"
    }
  }, {
    "selector" : "edge",
    "css" : {
      "font-size" : 10,
      "line-color" : "rgb(255,255,255)",
      "opacity" : 1.0,
      "source-arrow-shape" : "none",
      "target-arrow-color" : "rgb(0,0,0)",
      "text-opacity" : 1.0,
      "line-style" : "solid",
      "color" : "rgb(0,0,0)",
      "font-family" : "Dialog.plain",
      "font-weight" : "normal",
      "content" : "",
      "source-arrow-color" : "rgb(0,0,0)",
      "target-arrow-shape" : "none",
      "width" : 2.0
    }
  }, {
    "selector" : "edge:selected",
    "css" : {
      "line-color" : "rgb(255,0,0)"
    }
  } ]
}, {
  "format_version" : "1.0",
  "generated_by" : "cytoscape-3.7.2",
  "target_cytoscapejs_version" : "~2.1",
  "title" : "BioPAX_SIF",
  "style" : [ {
    "selector" : "node",
    "css" : {
      "text-opacity" : 1.0,
      "background-opacity" : 0.49019607843137253,
      "border-width" : 2.0,
      "color" : "rgb(0,0,0)",
      "border-color" : "rgb(0,0,0)",
      "font-size" : 12,
      "font-family" : "SansSerif.plain",
      "font-weight" : "normal",
      "text-valign" : "center",
      "text-halign" : "center",
      "border-opacity" : 1.0,
      "shape" : "ellipse",
      "width" : 60.0,
      "background-color" : "rgb(255,153,153)",
      "height" : 40.0,
      "content" : "data(name)"
    }
  }, {
    "selector" : "node[BIOPAX_TYPE = 'Complex']",
    "css" : {
      "background-color" : "rgb(153,204,255)"
    }
  }, {
    "selector" : "node[BIOPAX_TYPE = 'Complex']",
    "css" : {
      "shape" : "hexagon"
    }
  }, {
    "selector" : "node:selected",
    "css" : {
      "background-color" : "rgb(255,255,0)"
    }
  }, {
    "selector" : "edge",
    "css" : {
      "font-size" : 10,
      "line-color" : "rgb(0,0,0)",
      "opacity" : 1.0,
      "source-arrow-shape" : "none",
      "target-arrow-color" : "rgb(0,0,0)",
      "text-opacity" : 1.0,
      "line-style" : "solid",
      "color" : "rgb(0,0,0)",
      "font-family" : "SansSerif.plain",
      "font-weight" : "normal",
      "content" : "",
      "source-arrow-color" : "rgb(0,0,0)",
      "target-arrow-shape" : "none",
      "width" : 4.0
    }
  }, {
    "selector" : "edge[interaction = 'interacts-with']",
    "css" : {
      "line-color" : "rgb(0,85,0)",
      "target-arrow-color" : "rgb(0,85,0)",
      "source-arrow-color" : "rgb(0,85,0)"
    }
  }, {
    "selector" : "edge[interaction = 'chemical-affects']",
    "css" : {
      "line-color" : "rgb(240,144,0)",
      "target-arrow-color" : "rgb(240,144,0)",
      "source-arrow-color" : "rgb(240,144,0)"
    }
  }, {
    "selector" : "edge[interaction = 'controls-state-change-of']",
    "css" : {
      "line-color" : "rgb(0,0,192)",
      "target-arrow-color" : "rgb(0,0,192)",
      "source-arrow-color" : "rgb(0,0,192)"
    }
  }, {
    "selector" : "edge[interaction = 'controls-transport-of']",
    "css" : {
      "line-color" : "rgb(112,0,0)",
      "target-arrow-color" : "rgb(112,0,0)",
      "source-arrow-color" : "rgb(112,0,0)"
    }
  }, {
    "selector" : "edge[interaction = 'consumption-controled-by']",
    "css" : {
      "line-color" : "rgb(255,51,0)",
      "target-arrow-color" : "rgb(255,51,0)",
      "source-arrow-color" : "rgb(255,51,0)"
    }
  }, {
    "selector" : "edge[interaction = 'reacts-with']",
    "css" : {
      "line-color" : "rgb(0,255,0)",
      "target-arrow-color" : "rgb(0,255,0)",
      "source-arrow-color" : "rgb(0,255,0)"
    }
  }, {
    "selector" : "edge[interaction = 'neighbor-of']",
    "css" : {
      "line-color" : "rgb(0,170,0)",
      "target-arrow-color" : "rgb(0,170,0)",
      "source-arrow-color" : "rgb(0,170,0)"
    }
  }, {
    "selector" : "edge[interaction = 'controls-expression-of']",
    "css" : {
      "line-color" : "rgb(0,160,160)",
      "target-arrow-color" : "rgb(0,160,160)",
      "source-arrow-color" : "rgb(0,160,160)"
    }
  }, {
    "selector" : "edge[interaction = 'controls-phosphorylation-of']",
    "css" : {
      "line-color" : "rgb(0,0,255)",
      "target-arrow-color" : "rgb(0,0,255)",
      "source-arrow-color" : "rgb(0,0,255)"
    }
  }, {
    "selector" : "edge[interaction = 'controls-transport-of-chemical']",
    "css" : {
      "line-color" : "rgb(160,0,0)",
      "target-arrow-color" : "rgb(160,0,0)",
      "source-arrow-color" : "rgb(160,0,0)"
    }
  }, {
    "selector" : "edge[interaction = 'used-to-produce']",
    "css" : {
      "line-color" : "rgb(247,85,0)",
      "target-arrow-color" : "rgb(247,85,0)",
      "source-arrow-color" : "rgb(247,85,0)"
    }
  }, {
    "selector" : "edge[interaction = 'in-complex-with']",
    "css" : {
      "line-color" : "rgb(240,0,160)",
      "target-arrow-color" : "rgb(240,0,160)",
      "source-arrow-color" : "rgb(240,0,160)"
    }
  }, {
    "selector" : "edge[interaction = 'controls-production-of']",
    "css" : {
      "line-color" : "rgb(0,204,240)",
      "target-arrow-color" : "rgb(0,204,240)",
      "source-arrow-color" : "rgb(0,204,240)"
    }
  }, {
    "selector" : "edge[interaction = 'catalysis-precedes']",
    "css" : {
      "line-color" : "rgb(112,0,160)",
      "target-arrow-color" : "rgb(112,0,160)",
      "source-arrow-color" : "rgb(112,0,160)"
    }
  }, {
    "selector" : "edge[interaction = 'interacts-with']",
    "css" : {
      "line-color" : "rgb(0,85,0)"
    }
  }, {
    "selector" : "edge[interaction = 'chemical-affects']",
    "css" : {
      "line-color" : "rgb(240,144,0)"
    }
  }, {
    "selector" : "edge[interaction = 'controls-state-change-of']",
    "css" : {
      "line-color" : "rgb(0,0,192)"
    }
  }, {
    "selector" : "edge[interaction = 'controls-transport-of']",
    "css" : {
      "line-color" : "rgb(112,0,0)"
    }
  }, {
    "selector" : "edge[interaction = 'consumption-controled-by']",
    "css" : {
      "line-color" : "rgb(255,51,0)"
    }
  }, {
    "selector" : "edge[interaction = 'reacts-with']",
    "css" : {
      "line-color" : "rgb(0,255,0)"
    }
  }, {
    "selector" : "edge[interaction = 'neighbor-of']",
    "css" : {
      "line-color" : "rgb(0,170,0)"
    }
  }, {
    "selector" : "edge[interaction = 'controls-expression-of']",
    "css" : {
      "line-color" : "rgb(0,160,160)"
    }
  }, {
    "selector" : "edge[interaction = 'controls-phosphorylation-of']",
    "css" : {
      "line-color" : "rgb(0,0,255)"
    }
  }, {
    "selector" : "edge[interaction = 'controls-transport-of-chemical']",
    "css" : {
      "line-color" : "rgb(160,0,0)"
    }
  }, {
    "selector" : "edge[interaction = 'used-to-produce']",
    "css" : {
      "line-color" : "rgb(247,85,0)"
    }
  }, {
    "selector" : "edge[interaction = 'in-complex-with']",
    "css" : {
      "line-color" : "rgb(240,0,160)"
    }
  }, {
    "selector" : "edge[interaction = 'controls-production-of']",
    "css" : {
      "line-color" : "rgb(0,204,240)"
    }
  }, {
    "selector" : "edge[interaction = 'catalysis-precedes']",
    "css" : {
      "line-color" : "rgb(112,0,160)"
    }
  }, {
    "selector" : "edge[interaction = 'controls-expression-of']",
    "css" : {
      "target-arrow-shape" : "triangle"
    }
  }, {
    "selector" : "edge[interaction = 'chemical-affects']",
    "css" : {
      "target-arrow-shape" : "triangle"
    }
  }, {
    "selector" : "edge[interaction = 'controls-state-change-of']",
    "css" : {
      "target-arrow-shape" : "triangle"
    }
  }, {
    "selector" : "edge[interaction = 'controls-phosphorylation-of']",
    "css" : {
      "target-arrow-shape" : "triangle"
    }
  }, {
    "selector" : "edge[interaction = 'controls-transport-of-chemical']",
    "css" : {
      "target-arrow-shape" : "triangle"
    }
  }, {
    "selector" : "edge[interaction = 'used-to-produce']",
    "css" : {
      "target-arrow-shape" : "triangle"
    }
  }, {
    "selector" : "edge[interaction = 'controls-transport-of']",
    "css" : {
      "target-arrow-shape" : "triangle"
    }
  }, {
    "selector" : "edge[interaction = 'consumption-controled-by']",
    "css" : {
      "target-arrow-shape" : "triangle"
    }
  }, {
    "selector" : "edge[interaction = 'controls-production-of']",
    "css" : {
      "target-arrow-shape" : "triangle"
    }
  }, {
    "selector" : "edge[interaction = 'catalysis-precedes']",
    "css" : {
      "target-arrow-shape" : "triangle"
    }
  }, {
    "selector" : "edge:selected",
    "css" : {
      "line-color" : "rgb(255,0,0)"
    }
  } ]
}, {
  "format_version" : "1.0",
  "generated_by" : "cytoscape-3.7.2",
  "target_cytoscapejs_version" : "~2.1",
  "title" : "Curved",
  "style" : [ {
    "selector" : "node",
    "css" : {
      "text-opacity" : 1.0,
      "background-opacity" : 1.0,
      "border-width" : 7.0,
      "color" : "rgb(102,102,102)",
      "border-color" : "rgb(255,255,255)",
      "font-size" : 14,
      "font-family" : "Dialog.plain",
      "font-weight" : "normal",
      "text-valign" : "bottom",
      "text-halign" : "right",
      "border-opacity" : 1.0,
      "shape" : "ellipse",
      "width" : 18.0,
      "background-color" : "rgb(254,196,79)",
      "height" : 18.0,
      "content" : "data(name)"
    }
  }, {
    "selector" : "node:selected",
    "css" : {
      "background-color" : "rgb(255,255,0)"
    }
  }, {
    "selector" : "edge",
    "css" : {
      "font-size" : 10,
      "line-color" : "rgb(255,255,255)",
      "opacity" : 1.0,
      "source-arrow-shape" : "none",
      "target-arrow-color" : "rgb(255,255,255)",
      "text-opacity" : 1.0,
      "line-style" : "solid",
      "color" : "rgb(0,0,0)",
      "font-family" : "Dialog.plain",
      "font-weight" : "normal",
      "content" : "",
      "source-arrow-color" : "rgb(255,255,255)",
      "target-arrow-shape" : "triangle",
      "width" : 3.0
    }
  }, {
    "selector" : "edge:selected",
    "css" : {
      "line-color" : "rgb(255,0,0)"
    }
  } ]
} ]
```
