## Supplemental Network for "Granger-Causal Testing for Irregularly Sampled Time Series with Application to Nitrogen Signaling in Arabidopsis": 1d327552.scripts.js

```
"use strict";
angular.module("cyViewerApp", ["ngCookies", "ngResource", "ngSanitize", "ngRoute", "ngAnimate", "ui.bootstrap", "angular-underscore", "colorpicker.module", "angularSpinner"]).config(["$routeProvider", function(a) {
    a.when("/", {
        templateUrl: "main.html",
        controller: "MainCtrl"
    }).otherwise({
        redirectTo: "/"
    })
}]), angular.module("cyViewerApp").controller("MainCtrl", ["$scope", "$http", "$location", "$routeParams", "$window", "Network", "VisualStyles", function(a, b, c, d, e, f, g) {
    function h() {
        var b = angular.element(q);
        b.on("dragenter", function(a) {
            a.stopPropagation(), a.preventDefault()
        }), b.on("dragover", function(a) {
            a.stopPropagation(), a.preventDefault()
        }), b.on("drop", function(b) {
            b.preventDefault();
            var c = b.originalEvent.dataTransfer.files,
                d = c[0],
                e = new FileReader;
            e.onload = function(b) {
                var json = b.target.result
                if (!json.startsWith('{'))
                    json = json.substr(json.indexOf('{'))
                
                var c = JSON.parse(json)
                if (c.hasOwnProperty('format_version')){
                    c = {[c.data.name]: c}
                }
                d = null
                Object.keys(c).forEach(function(name){
                    a.networks[name] = c[name];
                    if (a.networkNames.indexOf(name) === -1)
                        a.networkNames.push(name);
                    (d === null && (d = c[name], a.currentNetwork = name));
                    z()
                }), a.cy.load(d.elements), k()
            }, e.readAsText(d)
        })
    }

    function i(b) {

        a.nodes = o.elements.nodes, a.edges = o.elements.edges, m(b);
        var c = o.data.name;
        a.networks[c] || (a.networks[c] = o, a.networkNames.push(c), a.currentNetwork = o.data.name), j(), d.bgcolor && (a.bg.color = d.bgcolor)
    }

    function j() {
        a.columnNames = [], a.edgeColumnNames = [], a.networkColumnNames = [];
        var b = a.nodes[0];
        for (var c in b.data) a.columnNames.push(c);
        var d = a.edges[0];
        for (var e in d.data) a.edgeColumnNames.push(e);
        for (var f in o.data) a.networkColumnNames.push(f)
    }

    function k() {
        a.selectedNodes = {}, a.selectedEdges = {}
    }

    function l() {
        a.selectedNodes = {}, a.selectedEdges = {};
        var b = !1;
        a.cy.on("select", "node", function(c) {
            var d = c.cyTarget.id();
            a.selectedNodes[d] = c.cyTarget, b = !0
        }), a.cy.on("select", "edge", function(c) {
            var d = c.cyTarget.id();
            a.selectedEdges[d] = c.cyTarget, b = !0
        }), a.cy.on("unselect", "node", function(c) {
            var d = c.cyTarget.id();
            delete a.selectedNodes[d], b = !0
        }), a.cy.on("unselect", "edge", function(c) {
            var d = c.cyTarget.id();
            delete a.selectedEdges[d], b = !0
        }), setInterval(function() {
            b && a.browserState.show && (a.$apply(), b = !1)
        }, 300)
    }

    function m(b) {
        _.each(b, function(b) {
            a.visualStyles[b.title] = b, a.visualStyleNames.push(b.title)
        }), a.currentVS = r
    }
    function z(){
        var el = document.getElementsByClassName('glyphicon')[0]
            el.click()
            el.click()
    }
    var n, o,
        q = "#network",
        r = "default";
    a.LAYOUTS = ["preset", "cola", "random", "grid", "circle", "concentric", "breadthfirst", "cose"], a.networks = {}, a.currentVS = null, a.visualStyles = [], a.visualStyleNames = [], a.networkNames = [], a.currentNetworkData = null, a.browserState = {
        show: !1
    }, a.overlayState = {
        show: !0
    }, a.toolbarState = {
        show: !0
    }, a.bg = {
        color: "#FAFAFA"
    }, a.columnNames = [], a.edgeColumnNames = [], a.networkColumnNames = [];
    var s = {
        showOverlay: !1,
        minZoom: .01,
        maxZoom: 200,
        boxSelectionEnabled: !0,
        layout: {
            name: "preset"
        },
        ready: function() {
            a.cy = this,

            a.cy.load(o.elements),
            i(n),
            h(),
            l(),
            a.currentVS = r,
            a.currentLayout = "preset",
            a.cy.style().fromJson(a.visualStyles[r].style).update(),
            angular.element(".loading").remove()
            var b = {
	            name: a.currentLayout
	        };
	        a.cy.layout(b)
            
            z()
        }
    };
    a.toggleTableBrowser = function() {
        a.browserState.show = !a.browserState.show
    }, a.toggleOverlay = function() {
        a.overlayState.show = !a.overlayState.show
    }, a.toggleToolbar = function() {
        a.toolbarState.show = !a.toolbarState.show
    }, a.fit = function() {
        a.cy.fit()
    }, a.switchVS = function() {
        var b = a.currentVS.trim(),
            c = a.visualStyles[b].style;
        a.cy.style().fromJson(c).update()
    }, a.switchNetwork = function() {
        var b = a.networks[a.currentNetwork]
        a.cy.load(b.elements),
        a.currentNetworkData = o,
        k(), a.nodes = b.elements.nodes,
        a.edges = b.elements.edges,
        j()
        //a.switchLayout()
    }, a.switchLayout = function() {
        var b = {
            name: a.currentLayout
        };
        a.cy.layout(b)
    }, function(){
        var b = networks;
    	a.networks = b;
        a.networkNames = Object.keys(b);
        c = a.networkNames[0];
        n = styles;
        o = b[c], a.currentNetworkData = o.data, a.currentNetwork = c
        ,angular.element(q).cytoscape(s)
    }()
}]), angular.module("cyViewerApp").factory("Network", ["$resource", function(a) {
	return a("data/:filename", {
        filename: "@filename"
    })
    return {get: function(){}}
}]), angular.module("cyViewerApp").factory("VisualStyles", ["$resource", function(a) {
	return a("data/:filename", {
        filename: styles
    })
}]);
```
