## Supplemental Network for "Granger-Causal Testing for Irregularly Sampled Time Series with Application to Nitrogen Signaling in Arabidopsis": 3e4be17b.app.js

```
// Sample project for visualizing JSON files.
//
//   by Keiichiro Ono
//
'use strict';

angular.module('cyViewerApp', [
    'ngCookies',
    'ngResource',
    'ngSanitize',
    'ngRoute',
    'ngAnimate',
    'ui.bootstrap',
    'angular-underscore',
    'colorpicker.module',
    'angularSpinner'
])
    .config(function($routeProvider) {
        // Routing
        $routeProvider
            .when('/', {
                templateUrl: 'main.html',
                controller: 'MainCtrl'
            })
            .otherwise({
                 redirectTo: '/'
            });
    });
```
