## Supplemental Network for "Granger-Causal Testing for Irregularly Sampled Time Series with Application to Nitrogen Signaling in Arabidopsis": 65d77273.vendor.js

```
if (function(a, b) {
        "object" == typeof module && "object" == typeof module.exports ? module.exports = a.document ? b(a, !0) : function(a) {
            if (!a.document) throw new Error("jQuery requires a window with a document");
            return b(a)
        } : b(a)
    }("undefined" != typeof window ? window : this, function(a, b) {
        function c(a) {
            var b = "length" in a && a.length,
                c = _.type(a);
            return "function" === c || _.isWindow(a) ? !1 : 1 === a.nodeType && b ? !0 : "array" === c || 0 === b || "number" == typeof b && b > 0 && b - 1 in a
        }

        function d(a, b, c) {
            if (_.isFunction(b)) return _.grep(a, function(a, d) {
                return !!b.call(a, d, a) !== c
            });
            if (b.nodeType) return _.grep(a, function(a) {
                return a === b !== c
            });
            if ("string" == typeof b) {
                if (ha.test(b)) return _.filter(b, a, c);
                b = _.filter(b, a)
            }
            return _.grep(a, function(a) {
                return U.call(b, a) >= 0 !== c
            })
        }

        function e(a, b) {
            for (;
                (a = a[b]) && 1 !== a.nodeType;);
            return a
        }

        function f(a) {
            var b = oa[a] = {};
            return _.each(a.match(na) || [], function(a, c) {
                b[c] = !0
            }), b
        }

        function g() {
            Z.removeEventListener("DOMContentLoaded", g, !1), a.removeEventListener("load", g, !1), _.ready()
        }

        function h() {
            Object.defineProperty(this.cache = {}, 0, {
                get: function() {
                    return {}
                }
            }), this.expando = _.expando + h.uid++
        }

        function i(a, b, c) {
            var d;
            if (void 0 === c && 1 === a.nodeType)
                if (d = "data-" + b.replace(ua, "-$1").toLowerCase(), c = a.getAttribute(d), "string" == typeof c) {
                    try {
                        c = "true" === c ? !0 : "false" === c ? !1 : "null" === c ? null : +c + "" === c ? +c : ta.test(c) ? _.parseJSON(c) : c
                    } catch (e) {}
                    sa.set(a, b, c)
                } else c = void 0;
            return c
        }

        function j() {
            return !0
        }

        function k() {
            return !1
        }

        function l() {
            try {
                return Z.activeElement
            } catch (a) {}
        }

        function m(a, b) {
            return _.nodeName(a, "table") && _.nodeName(11 !== b.nodeType ? b : b.firstChild, "tr") ? a.getElementsByTagName("tbody")[0] || a.appendChild(a.ownerDocument.createElement("tbody")) : a
        }

        function n(a) {
            return a.type = (null !== a.getAttribute("type")) + "/" + a.type, a
        }

        function o(a) {
            var b = Ka.exec(a.type);
            return b ? a.type = b[1] : a.removeAttribute("type"), a
        }

        function p(a, b) {
            for (var c = 0, d = a.length; d > c; c++) ra.set(a[c], "globalEval", !b || ra.get(b[c], "globalEval"))
        }

        function q(a, b) {
            var c, d, e, f, g, h, i, j;
            if (1 === b.nodeType) {
                if (ra.hasData(a) && (f = ra.access(a), g = ra.set(b, f), j = f.events)) {
                    delete g.handle, g.events = {};
                    for (e in j)
                        for (c = 0, d = j[e].length; d > c; c++) _.event.add(b, e, j[e][c])
                }
                sa.hasData(a) && (h = sa.access(a), i = _.extend({}, h), sa.set(b, i))
            }
        }

        function r(a, b) {
            var c = a.getElementsByTagName ? a.getElementsByTagName(b || "*") : a.querySelectorAll ? a.querySelectorAll(b || "*") : [];
            return void 0 === b || b && _.nodeName(a, b) ? _.merge([a], c) : c
        }

        function s(a, b) {
            var c = b.nodeName.toLowerCase();
            "input" === c && ya.test(a.type) ? b.checked = a.checked : ("input" === c || "textarea" === c) && (b.defaultValue = a.defaultValue)
        }

        function t(b, c) {
            var d, e = _(c.createElement(b)).appendTo(c.body),
                f = a.getDefaultComputedStyle && (d = a.getDefaultComputedStyle(e[0])) ? d.display : _.css(e[0], "display");
            return e.detach(), f
        }

        function u(a) {
            var b = Z,
                c = Oa[a];
            return c || (c = t(a, b), "none" !== c && c || (Na = (Na || _("<iframe frameborder='0' width='0' height='0'/>")).appendTo(b.documentElement), b = Na[0].contentDocument, b.write(), b.close(), c = t(a, b), Na.detach()), Oa[a] = c), c
        }

        function v(a, b, c) {
            var d, e, f, g, h = a.style;
            return c = c || Ra(a), c && (g = c.getPropertyValue(b) || c[b]), c && ("" !== g || _.contains(a.ownerDocument, a) || (g = _.style(a, b)), Qa.test(g) && Pa.test(b) && (d = h.width, e = h.minWidth, f = h.maxWidth, h.minWidth = h.maxWidth = h.width = g, g = c.width, h.width = d, h.minWidth = e, h.maxWidth = f)), void 0 !== g ? g + "" : g
        }

        function w(a, b) {
            return {
                get: function() {
                    return a() ? void delete this.get : (this.get = b).apply(this, arguments)
                }
            }
        }

        function x(a, b) {
            if (b in a) return b;
            for (var c = b[0].toUpperCase() + b.slice(1), d = b, e = Xa.length; e--;)
                if (b = Xa[e] + c, b in a) return b;
            return d
        }

        function y(a, b, c) {
            var d = Ta.exec(b);
            return d ? Math.max(0, d[1] - (c || 0)) + (d[2] || "px") : b
        }

        function z(a, b, c, d, e) {
            for (var f = c === (d ? "border" : "content") ? 4 : "width" === b ? 1 : 0, g = 0; 4 > f; f += 2) "margin" === c && (g += _.css(a, c + wa[f], !0, e)), d ? ("content" === c && (g -= _.css(a, "padding" + wa[f], !0, e)), "margin" !== c && (g -= _.css(a, "border" + wa[f] + "Width", !0, e))) : (g += _.css(a, "padding" + wa[f], !0, e), "padding" !== c && (g += _.css(a, "border" + wa[f] + "Width", !0, e)));
            return g
        }

        function A(a, b, c) {
            var d = !0,
                e = "width" === b ? a.offsetWidth : a.offsetHeight,
                f = Ra(a),
                g = "border-box" === _.css(a, "boxSizing", !1, f);
            if (0 >= e || null == e) {
                if (e = v(a, b, f), (0 > e || null == e) && (e = a.style[b]), Qa.test(e)) return e;
                d = g && (Y.boxSizingReliable() || e === a.style[b]), e = parseFloat(e) || 0
            }
            return e + z(a, b, c || (g ? "border" : "content"), d, f) + "px"
        }

        function B(a, b) {
            for (var c, d, e, f = [], g = 0, h = a.length; h > g; g++) d = a[g], d.style && (f[g] = ra.get(d, "olddisplay"), c = d.style.display, b ? (f[g] || "none" !== c || (d.style.display = ""), "" === d.style.display && xa(d) && (f[g] = ra.access(d, "olddisplay", u(d.nodeName)))) : (e = xa(d), "none" === c && e || ra.set(d, "olddisplay", e ? c : _.css(d, "display"))));
            for (g = 0; h > g; g++) d = a[g], d.style && (b && "none" !== d.style.display && "" !== d.style.display || (d.style.display = b ? f[g] || "" : "none"));
            return a
        }

        function C(a, b, c, d, e) {
            return new C.prototype.init(a, b, c, d, e)
        }

        function D() {
            return setTimeout(function() {
                Ya = void 0
            }), Ya = _.now()
        }

        function E(a, b) {
            var c, d = 0,
                e = {
                    height: a
                };
            for (b = b ? 1 : 0; 4 > d; d += 2 - b) c = wa[d], e["margin" + c] = e["padding" + c] = a;
            return b && (e.opacity = e.width = a), e
        }

        function F(a, b, c) {
            for (var d, e = (cb[b] || []).concat(cb["*"]), f = 0, g = e.length; g > f; f++)
                if (d = e[f].call(c, b, a)) return d
        }

        function G(a, b, c) {
            var d, e, f, g, h, i, j, k, l = this,
                m = {},
                n = a.style,
                o = a.nodeType && xa(a),
                p = ra.get(a, "fxshow");
            c.queue || (h = _._queueHooks(a, "fx"), null == h.unqueued && (h.unqueued = 0, i = h.empty.fire, h.empty.fire = function() {
                h.unqueued || i()
            }), h.unqueued++, l.always(function() {
                l.always(function() {
                    h.unqueued--, _.queue(a, "fx").length || h.empty.fire()
                })
            })), 1 === a.nodeType && ("height" in b || "width" in b) && (c.overflow = [n.overflow, n.overflowX, n.overflowY], j = _.css(a, "display"), k = "none" === j ? ra.get(a, "olddisplay") || u(a.nodeName) : j, "inline" === k && "none" === _.css(a, "float") && (n.display = "inline-block")), c.overflow && (n.overflow = "hidden", l.always(function() {
                n.overflow = c.overflow[0], n.overflowX = c.overflow[1], n.overflowY = c.overflow[2]
            }));
            for (d in b)
                if (e = b[d], $a.exec(e)) {
                    if (delete b[d], f = f || "toggle" === e, e === (o ? "hide" : "show")) {
                        if ("show" !== e || !p || void 0 === p[d]) continue;
                        o = !0
                    }
                    m[d] = p && p[d] || _.style(a, d)
                } else j = void 0;
            if (_.isEmptyObject(m)) "inline" === ("none" === j ? u(a.nodeName) : j) && (n.display = j);
            else {
                p ? "hidden" in p && (o = p.hidden) : p = ra.access(a, "fxshow", {}), f && (p.hidden = !o), o ? _(a).show() : l.done(function() {
                    _(a).hide()
                }), l.done(function() {
                    var b;
                    ra.remove(a, "fxshow");
                    for (b in m) _.style(a, b, m[b])
                });
                for (d in m) g = F(o ? p[d] : 0, d, l), d in p || (p[d] = g.start, o && (g.end = g.start, g.start = "width" === d || "height" === d ? 1 : 0))
            }
        }

        function H(a, b) {
            var c, d, e, f, g;
            for (c in a)
                if (d = _.camelCase(c), e = b[d], f = a[c], _.isArray(f) && (e = f[1], f = a[c] = f[0]), c !== d && (a[d] = f, delete a[c]), g = _.cssHooks[d], g && "expand" in g) {
                    f = g.expand(f), delete a[d];
                    for (c in f) c in a || (a[c] = f[c], b[c] = e)
                } else b[d] = e
        }

        function I(a, b, c) {
            var d, e, f = 0,
                g = bb.length,
                h = _.Deferred().always(function() {
                    delete i.elem
                }),
                i = function() {
                    if (e) return !1;
                    for (var b = Ya || D(), c = Math.max(0, j.startTime + j.duration - b), d = c / j.duration || 0, f = 1 - d, g = 0, i = j.tweens.length; i > g; g++) j.tweens[g].run(f);
                    return h.notifyWith(a, [j, f, c]), 1 > f && i ? c : (h.resolveWith(a, [j]), !1)
                },
                j = h.promise({
                    elem: a,
                    props: _.extend({}, b),
                    opts: _.extend(!0, {
                        specialEasing: {}
                    }, c),
                    originalProperties: b,
                    originalOptions: c,
                    startTime: Ya || D(),
                    duration: c.duration,
                    tweens: [],
                    createTween: function(b, c) {
                        var d = _.Tween(a, j.opts, b, c, j.opts.specialEasing[b] || j.opts.easing);
                        return j.tweens.push(d), d
                    },
                    stop: function(b) {
                        var c = 0,
                            d = b ? j.tweens.length : 0;
                        if (e) return this;
                        for (e = !0; d > c; c++) j.tweens[c].run(1);
                        return b ? h.resolveWith(a, [j, b]) : h.rejectWith(a, [j, b]), this
                    }
                }),
                k = j.props;
            for (H(k, j.opts.specialEasing); g > f; f++)
                if (d = bb[f].call(j, a, k, j.opts)) return d;
            return _.map(k, F, j), _.isFunction(j.opts.start) && j.opts.start.call(a, j), _.fx.timer(_.extend(i, {
                elem: a,
                anim: j,
                queue: j.opts.queue
            })), j.progress(j.opts.progress).done(j.opts.done, j.opts.complete).fail(j.opts.fail).always(j.opts.always)
        }

        function J(a) {
            return function(b, c) {
                "string" != typeof b && (c = b, b = "*");
                var d, e = 0,
                    f = b.toLowerCase().match(na) || [];
                if (_.isFunction(c))
                    for (; d = f[e++];) "+" === d[0] ? (d = d.slice(1) || "*", (a[d] = a[d] || []).unshift(c)) : (a[d] = a[d] || []).push(c)
            }
        }

        function K(a, b, c, d) {
            function e(h) {
                var i;
                return f[h] = !0, _.each(a[h] || [], function(a, h) {
                    var j = h(b, c, d);
                    return "string" != typeof j || g || f[j] ? g ? !(i = j) : void 0 : (b.dataTypes.unshift(j), e(j), !1)
                }), i
            }
            var f = {},
                g = a === tb;
            return e(b.dataTypes[0]) || !f["*"] && e("*")
        }

        function L(a, b) {
            var c, d, e = _.ajaxSettings.flatOptions || {};
            for (c in b) void 0 !== b[c] && ((e[c] ? a : d || (d = {}))[c] = b[c]);
            return d && _.extend(!0, a, d), a
        }

        function M(a, b, c) {
            for (var d, e, f, g, h = a.contents, i = a.dataTypes;
                "*" === i[0];) i.shift(), void 0 === d && (d = a.mimeType || b.getResponseHeader("Content-Type"));
            if (d)
                for (e in h)
                    if (h[e] && h[e].test(d)) {
                        i.unshift(e);
                        break
                    }
            if (i[0] in c) f = i[0];
            else {
                for (e in c) {
                    if (!i[0] || a.converters[e + " " + i[0]]) {
                        f = e;
                        break
                    }
                    g || (g = e)
                }
                f = f || g
            }
            return f ? (f !== i[0] && i.unshift(f), c[f]) : void 0
        }

        function N(a, b, c, d) {
            var e, f, g, h, i, j = {},
                k = a.dataTypes.slice();
            if (k[1])
                for (g in a.converters) j[g.toLowerCase()] = a.converters[g];
            for (f = k.shift(); f;)
                if (a.responseFields[f] && (c[a.responseFields[f]] = b), !i && d && a.dataFilter && (b = a.dataFilter(b, a.dataType)), i = f, f = k.shift())
                    if ("*" === f) f = i;
                    else if ("*" !== i && i !== f) {
                if (g = j[i + " " + f] || j["* " + f], !g)
                    for (e in j)
                        if (h = e.split(" "), h[1] === f && (g = j[i + " " + h[0]] || j["* " + h[0]])) {
                            g === !0 ? g = j[e] : j[e] !== !0 && (f = h[0], k.unshift(h[1]));
                            break
                        }
                if (g !== !0)
                    if (g && a["throws"]) b = g(b);
                    else try {
                        b = g(b)
                    } catch (l) {
                        return {
                            state: "parsererror",
                            error: g ? l : "No conversion from " + i + " to " + f
                        }
                    }
            }
            return {
                state: "success",
                data: b
            }
        }

        function O(a, b, c, d) {
            var e;
            if (_.isArray(b)) _.each(b, function(b, e) {
                c || yb.test(a) ? d(a, e) : O(a + "[" + ("object" == typeof e ? b : "") + "]", e, c, d)
            });
            else if (c || "object" !== _.type(b)) d(a, b);
            else
                for (e in b) O(a + "[" + e + "]", b[e], c, d)
        }

        function P(a) {
            return _.isWindow(a) ? a : 9 === a.nodeType && a.defaultView
        }
        var Q = [],
            R = Q.slice,
            S = Q.concat,
            T = Q.push,
            U = Q.indexOf,
            V = {},
            W = V.toString,
            X = V.hasOwnProperty,
            Y = {},
            Z = a.document,
            $ = "2.1.4",
            _ = function(a, b) {
                return new _.fn.init(a, b)
            },
            aa = /^[\s\uFEFF\xA0]+|[\s\uFEFF\xA0]+$/g,
            ba = /^-ms-/,
            ca = /-([\da-z])/gi,
            da = function(a, b) {
                return b.toUpperCase()
            };
        _.fn = _.prototype = {
            jquery: $,
            constructor: _,
            selector: "",
            length: 0,
            toArray: function() {
                return R.call(this)
            },
            get: function(a) {
                return null != a ? 0 > a ? this[a + this.length] : this[a] : R.call(this)
            },
            pushStack: function(a) {
                var b = _.merge(this.constructor(), a);
                return b.prevObject = this, b.context = this.context, b
            },
            each: function(a, b) {
                return _.each(this, a, b)
            },
            map: function(a) {
                return this.pushStack(_.map(this, function(b, c) {
                    return a.call(b, c, b)
                }))
            },
            slice: function() {
                return this.pushStack(R.apply(this, arguments))
            },
            first: function() {
                return this.eq(0)
            },
            last: function() {
                return this.eq(-1)
            },
            eq: function(a) {
                var b = this.length,
                    c = +a + (0 > a ? b : 0);
                return this.pushStack(c >= 0 && b > c ? [this[c]] : [])
            },
            end: function() {
                return this.prevObject || this.constructor(null)
            },
            push: T,
            sort: Q.sort,
            splice: Q.splice
        }, _.extend = _.fn.extend = function() {
            var a, b, c, d, e, f, g = arguments[0] || {},
                h = 1,
                i = arguments.length,
                j = !1;
            for ("boolean" == typeof g && (j = g, g = arguments[h] || {}, h++), "object" == typeof g || _.isFunction(g) || (g = {}), h === i && (g = this, h--); i > h; h++)
                if (null != (a = arguments[h]))
                    for (b in a) c = g[b], d = a[b], g !== d && (j && d && (_.isPlainObject(d) || (e = _.isArray(d))) ? (e ? (e = !1, f = c && _.isArray(c) ? c : []) : f = c && _.isPlainObject(c) ? c : {}, g[b] = _.extend(j, f, d)) : void 0 !== d && (g[b] = d));
            return g
        }, _.extend({
            expando: "jQuery" + ($ + Math.random()).replace(/\D/g, ""),
            isReady: !0,
            error: function(a) {
                throw new Error(a)
            },
            noop: function() {},
            isFunction: function(a) {
                return "function" === _.type(a)
            },
            isArray: Array.isArray,
            isWindow: function(a) {
                return null != a && a === a.window
            },
            isNumeric: function(a) {
                return !_.isArray(a) && a - parseFloat(a) + 1 >= 0
            },
            isPlainObject: function(a) {
                return "object" !== _.type(a) || a.nodeType || _.isWindow(a) ? !1 : a.constructor && !X.call(a.constructor.prototype, "isPrototypeOf") ? !1 : !0
            },
            isEmptyObject: function(a) {
                var b;
                for (b in a) return !1;
                return !0
            },
            type: function(a) {
                return null == a ? a + "" : "object" == typeof a || "function" == typeof a ? V[W.call(a)] || "object" : typeof a
            },
            globalEval: function(a) {
                var b, c = eval;
                a = _.trim(a), a && (1 === a.indexOf("use strict") ? (b = Z.createElement("script"), b.text = a, Z.head.appendChild(b).parentNode.removeChild(b)) : c(a))
            },
            camelCase: function(a) {
                return a.replace(ba, "ms-").replace(ca, da)
            },
            nodeName: function(a, b) {
                return a.nodeName && a.nodeName.toLowerCase() === b.toLowerCase()
            },
            each: function(a, b, d) {
                var e, f = 0,
                    g = a.length,
                    h = c(a);
                if (d) {
                    if (h)
                        for (; g > f && (e = b.apply(a[f], d), e !== !1); f++);
                    else
                        for (f in a)
                            if (e = b.apply(a[f], d), e === !1) break
                } else if (h)
                    for (; g > f && (e = b.call(a[f], f, a[f]), e !== !1); f++);
                else
                    for (f in a)
                        if (e = b.call(a[f], f, a[f]), e === !1) break;
                return a
            },
            trim: function(a) {
                return null == a ? "" : (a + "").replace(aa, "")
            },
            makeArray: function(a, b) {
                var d = b || [];
                return null != a && (c(Object(a)) ? _.merge(d, "string" == typeof a ? [a] : a) : T.call(d, a)), d
            },
            inArray: function(a, b, c) {
                return null == b ? -1 : U.call(b, a, c)
            },
            merge: function(a, b) {
                for (var c = +b.length, d = 0, e = a.length; c > d; d++) a[e++] = b[d];
                return a.length = e, a
            },
            grep: function(a, b, c) {
                for (var d, e = [], f = 0, g = a.length, h = !c; g > f; f++) d = !b(a[f], f), d !== h && e.push(a[f]);
                return e
            },
            map: function(a, b, d) {
                var e, f = 0,
                    g = a.length,
                    h = c(a),
                    i = [];
                if (h)
                    for (; g > f; f++) e = b(a[f], f, d), null != e && i.push(e);
                else
                    for (f in a) e = b(a[f], f, d), null != e && i.push(e);
                return S.apply([], i)
            },
            guid: 1,
            proxy: function(a, b) {
                var c, d, e;
                return "string" == typeof b && (c = a[b], b = a, a = c), _.isFunction(a) ? (d = R.call(arguments, 2), e = function() {
                    return a.apply(b || this, d.concat(R.call(arguments)))
                }, e.guid = a.guid = a.guid || _.guid++, e) : void 0
            },
            now: Date.now,
            support: Y
        }), _.each("Boolean Number String Function Array Date RegExp Object Error".split(" "), function(a, b) {
            V["[object " + b + "]"] = b.toLowerCase()
        });
        var ea = function(a) {
            function b(a, b, c, d) {
                var e, f, g, h, i, j, l, n, o, p;
                if ((b ? b.ownerDocument || b : O) !== G && F(b), b = b || G, c = c || [], h = b.nodeType, "string" != typeof a || !a || 1 !== h && 9 !== h && 11 !== h) return c;
                if (!d && I) {
                    if (11 !== h && (e = sa.exec(a)))
                        if (g = e[1]) {
                            if (9 === h) {
                                if (f = b.getElementById(g), !f || !f.parentNode) return c;
                                if (f.id === g) return c.push(f), c
                            } else if (b.ownerDocument && (f = b.ownerDocument.getElementById(g)) && M(b, f) && f.id === g) return c.push(f), c
                        } else {
                            if (e[2]) return $.apply(c, b.getElementsByTagName(a)), c;
                            if ((g = e[3]) && v.getElementsByClassName) return $.apply(c, b.getElementsByClassName(g)), c
                        }
                    if (v.qsa && (!J || !J.test(a))) {
                        if (n = l = N, o = b, p = 1 !== h && a, 1 === h && "object" !== b.nodeName.toLowerCase()) {
                            for (j = z(a), (l = b.getAttribute("id")) ? n = l.replace(ua, "\\$&") : b.setAttribute("id", n), n = "[id='" + n + "'] ", i = j.length; i--;) j[i] = n + m(j[i]);
                            o = ta.test(a) && k(b.parentNode) || b, p = j.join(",")
                        }
                        if (p) try {
                            return $.apply(c, o.querySelectorAll(p)), c
                        } catch (q) {} finally {
                            l || b.removeAttribute("id")
                        }
                    }
                }
                return B(a.replace(ia, "$1"), b, c, d)
            }

            function c() {
                function a(c, d) {
                    return b.push(c + " ") > w.cacheLength && delete a[b.shift()], a[c + " "] = d
                }
                var b = [];
                return a
            }

            function d(a) {
                return a[N] = !0, a
            }

            function e(a) {
                var b = G.createElement("div");
                try {
                    return !!a(b)
                } catch (c) {
                    return !1
                } finally {
                    b.parentNode && b.parentNode.removeChild(b), b = null
                }
            }

            function f(a, b) {
                for (var c = a.split("|"), d = a.length; d--;) w.attrHandle[c[d]] = b
            }

            function g(a, b) {
                var c = b && a,
                    d = c && 1 === a.nodeType && 1 === b.nodeType && (~b.sourceIndex || V) - (~a.sourceIndex || V);
                if (d) return d;
                if (c)
                    for (; c = c.nextSibling;)
                        if (c === b) return -1;
                return a ? 1 : -1
            }

            function h(a) {
                return function(b) {
                    var c = b.nodeName.toLowerCase();
                    return "input" === c && b.type === a
                }
            }

            function i(a) {
                return function(b) {
                    var c = b.nodeName.toLowerCase();
                    return ("input" === c || "button" === c) && b.type === a
                }
            }

            function j(a) {
                return d(function(b) {
                    return b = +b, d(function(c, d) {
                        for (var e, f = a([], c.length, b), g = f.length; g--;) c[e = f[g]] && (c[e] = !(d[e] = c[e]))
                    })
                })
            }

            function k(a) {
                return a && "undefined" != typeof a.getElementsByTagName && a
            }

            function l() {}

            function m(a) {
                for (var b = 0, c = a.length, d = ""; c > b; b++) d += a[b].value;
                return d
            }

            function n(a, b, c) {
                var d = b.dir,
                    e = c && "parentNode" === d,
                    f = Q++;
                return b.first ? function(b, c, f) {
                    for (; b = b[d];)
                        if (1 === b.nodeType || e) return a(b, c, f)
                } : function(b, c, g) {
                    var h, i, j = [P, f];
                    if (g) {
                        for (; b = b[d];)
                            if ((1 === b.nodeType || e) && a(b, c, g)) return !0
                    } else
                        for (; b = b[d];)
                            if (1 === b.nodeType || e) {
                                if (i = b[N] || (b[N] = {}), (h = i[d]) && h[0] === P && h[1] === f) return j[2] = h[2];
                                if (i[d] = j, j[2] = a(b, c, g)) return !0
                            }
                }
            }

            function o(a) {
                return a.length > 1 ? function(b, c, d) {
                    for (var e = a.length; e--;)
                        if (!a[e](b, c, d)) return !1;
                    return !0
                } : a[0]
            }

            function p(a, c, d) {
                for (var e = 0, f = c.length; f > e; e++) b(a, c[e], d);
                return d
            }

            function q(a, b, c, d, e) {
                for (var f, g = [], h = 0, i = a.length, j = null != b; i > h; h++)(f = a[h]) && (!c || c(f, d, e)) && (g.push(f), j && b.push(h));
                return g
            }

            function r(a, b, c, e, f, g) {
                return e && !e[N] && (e = r(e)), f && !f[N] && (f = r(f, g)), d(function(d, g, h, i) {
                    var j, k, l, m = [],
                        n = [],
                        o = g.length,
                        r = d || p(b || "*", h.nodeType ? [h] : h, []),
                        s = !a || !d && b ? r : q(r, m, a, h, i),
                        t = c ? f || (d ? a : o || e) ? [] : g : s;
                    if (c && c(s, t, h, i), e)
                        for (j = q(t, n), e(j, [], h, i), k = j.length; k--;)(l = j[k]) && (t[n[k]] = !(s[n[k]] = l));
                    if (d) {
                        if (f || a) {
                            if (f) {
                                for (j = [], k = t.length; k--;)(l = t[k]) && j.push(s[k] = l);
                                f(null, t = [], j, i)
                            }
                            for (k = t.length; k--;)(l = t[k]) && (j = f ? aa(d, l) : m[k]) > -1 && (d[j] = !(g[j] = l))
                        }
                    } else t = q(t === g ? t.splice(o, t.length) : t), f ? f(null, g, t, i) : $.apply(g, t)
                })
            }

            function s(a) {
                for (var b, c, d, e = a.length, f = w.relative[a[0].type], g = f || w.relative[" "], h = f ? 1 : 0, i = n(function(a) {
                        return a === b
                    }, g, !0), j = n(function(a) {
                        return aa(b, a) > -1
                    }, g, !0), k = [function(a, c, d) {
                        var e = !f && (d || c !== C) || ((b = c).nodeType ? i(a, c, d) : j(a, c, d));
                        return b = null, e
                    }]; e > h; h++)
                    if (c = w.relative[a[h].type]) k = [n(o(k), c)];
                    else {
                        if (c = w.filter[a[h].type].apply(null, a[h].matches), c[N]) {
                            for (d = ++h; e > d && !w.relative[a[d].type]; d++);
                            return r(h > 1 && o(k), h > 1 && m(a.slice(0, h - 1).concat({
                                value: " " === a[h - 2].type ? "*" : ""
                            })).replace(ia, "$1"), c, d > h && s(a.slice(h, d)), e > d && s(a = a.slice(d)), e > d && m(a))
                        }
                        k.push(c)
                    }
                return o(k)
            }

            function t(a, c) {
                var e = c.length > 0,
                    f = a.length > 0,
                    g = function(d, g, h, i, j) {
                        var k, l, m, n = 0,
                            o = "0",
                            p = d && [],
                            r = [],
                            s = C,
                            t = d || f && w.find.TAG("*", j),
                            u = P += null == s ? 1 : Math.random() || .1,
                            v = t.length;
                        for (j && (C = g !== G && g); o !== v && null != (k = t[o]); o++) {
                            if (f && k) {
                                for (l = 0; m = a[l++];)
                                    if (m(k, g, h)) {
                                        i.push(k);
                                        break
                                    }
                                j && (P = u)
                            }
                            e && ((k = !m && k) && n--, d && p.push(k))
                        }
                        if (n += o, e && o !== n) {
                            for (l = 0; m = c[l++];) m(p, r, g, h);
                            if (d) {
                                if (n > 0)
                                    for (; o--;) p[o] || r[o] || (r[o] = Y.call(i));
                                r = q(r)
                            }
                            $.apply(i, r), j && !d && r.length > 0 && n + c.length > 1 && b.uniqueSort(i)
                        }
                        return j && (P = u, C = s), p
                    };
                return e ? d(g) : g
            }
            var u, v, w, x, y, z, A, B, C, D, E, F, G, H, I, J, K, L, M, N = "sizzle" + 1 * new Date,
                O = a.document,
                P = 0,
                Q = 0,
                R = c(),
                S = c(),
                T = c(),
                U = function(a, b) {
                    return a === b && (E = !0), 0
                },
                V = 1 << 31,
                W = {}.hasOwnProperty,
                X = [],
                Y = X.pop,
                Z = X.push,
                $ = X.push,
                _ = X.slice,
                aa = function(a, b) {
                    for (var c = 0, d = a.length; d > c; c++)
                        if (a[c] === b) return c;
                    return -1
                },
                ba = "checked|selected|async|autofocus|autoplay|controls|defer|disabled|hidden|ismap|loop|multiple|open|readonly|required|scoped",
                ca = "[\\x20\\t\\r\\n\\f]",
                da = "(?:\\\\.|[\\w-]|[^\\x00-\\xa0])+",
                ea = da.replace("w", "w#"),
                fa = "\\[" + ca + "*(" + da + ")(?:" + ca + "*([*^$|!~]?=)" + ca + "*(?:'((?:\\\\.|[^\\\\'])*)'|\"((?:\\\\.|[^\\\\\"])*)\"|(" + ea + "))|)" + ca + "*\\]",
                ga = ":(" + da + ")(?:\\((('((?:\\\\.|[^\\\\'])*)'|\"((?:\\\\.|[^\\\\\"])*)\")|((?:\\\\.|[^\\\\()[\\]]|" + fa + ")*)|.*)\\)|)",
                ha = new RegExp(ca + "+", "g"),
                ia = new RegExp("^" + ca + "+|((?:^|[^\\\\])(?:\\\\.)*)" + ca + "+$", "g"),
                ja = new RegExp("^" + ca + "*," + ca + "*"),
                ka = new RegExp("^" + ca + "*([>+~]|" + ca + ")" + ca + "*"),
                la = new RegExp("=" + ca + "*([^\\]'\"]*?)" + ca + "*\\]", "g"),
                ma = new RegExp(ga),
                na = new RegExp("^" + ea + "$"),
                oa = {
                    ID: new RegExp("^#(" + da + ")"),
                    CLASS: new RegExp("^\\.(" + da + ")"),
                    TAG: new RegExp("^(" + da.replace("w", "w*") + ")"),
                    ATTR: new RegExp("^" + fa),
                    PSEUDO: new RegExp("^" + ga),
                    CHILD: new RegExp("^:(only|first|last|nth|nth-last)-(child|of-type)(?:\\(" + ca + "*(even|odd|(([+-]|)(\\d*)n|)" + ca + "*(?:([+-]|)" + ca + "*(\\d+)|))" + ca + "*\\)|)", "i"),
                    bool: new RegExp("^(?:" + ba + ")$", "i"),
                    needsContext: new RegExp("^" + ca + "*[>+~]|:(even|odd|eq|gt|lt|nth|first|last)(?:\\(" + ca + "*((?:-\\d)?\\d*)" + ca + "*\\)|)(?=[^-]|$)", "i")
                },
                pa = /^(?:input|select|textarea|button)$/i,
                qa = /^h\d$/i,
                ra = /^[^{]+\{\s*\[native \w/,
                sa = /^(?:#([\w-]+)|(\w+)|\.([\w-]+))$/,
                ta = /[+~]/,
                ua = /'|\\/g,
                va = new RegExp("\\\\([\\da-f]{1,6}" + ca + "?|(" + ca + ")|.)", "ig"),
                wa = function(a, b, c) {
                    var d = "0x" + b - 65536;
                    return d !== d || c ? b : 0 > d ? String.fromCharCode(d + 65536) : String.fromCharCode(d >> 10 | 55296, 1023 & d | 56320)
                },
                xa = function() {
                    F()
                };
            try {
                $.apply(X = _.call(O.childNodes), O.childNodes), X[O.childNodes.length].nodeType
            } catch (ya) {
                $ = {
                    apply: X.length ? function(a, b) {
                        Z.apply(a, _.call(b))
                    } : function(a, b) {
                        for (var c = a.length, d = 0; a[c++] = b[d++];);
                        a.length = c - 1
                    }
                }
            }
            v = b.support = {}, y = b.isXML = function(a) {
                var b = a && (a.ownerDocument || a).documentElement;
                return b ? "HTML" !== b.nodeName : !1
            }, F = b.setDocument = function(a) {
                var b, c, d = a ? a.ownerDocument || a : O;
                return d !== G && 9 === d.nodeType && d.documentElement ? (G = d, H = d.documentElement, c = d.defaultView, c && c !== c.top && (c.addEventListener ? c.addEventListener("unload", xa, !1) : c.attachEvent && c.attachEvent("onunload", xa)), I = !y(d), v.attributes = e(function(a) {
                    return a.className = "i", !a.getAttribute("className")
                }), v.getElementsByTagName = e(function(a) {
                    return a.appendChild(d.createComment("")), !a.getElementsByTagName("*").length
                }), v.getElementsByClassName = ra.test(d.getElementsByClassName), v.getById = e(function(a) {
                    return H.appendChild(a).id = N, !d.getElementsByName || !d.getElementsByName(N).length
                }), v.getById ? (w.find.ID = function(a, b) {
                    if ("undefined" != typeof b.getElementById && I) {
                        var c = b.getElementById(a);
                        return c && c.parentNode ? [c] : []
                    }
                }, w.filter.ID = function(a) {
                    var b = a.replace(va, wa);
                    return function(a) {
                        return a.getAttribute("id") === b
                    }
                }) : (delete w.find.ID, w.filter.ID = function(a) {
                    var b = a.replace(va, wa);
                    return function(a) {
                        var c = "undefined" != typeof a.getAttributeNode && a.getAttributeNode("id");
                        return c && c.value === b
                    }
                }), w.find.TAG = v.getElementsByTagName ? function(a, b) {
                    return "undefined" != typeof b.getElementsByTagName ? b.getElementsByTagName(a) : v.qsa ? b.querySelectorAll(a) : void 0
                } : function(a, b) {
                    var c, d = [],
                        e = 0,
                        f = b.getElementsByTagName(a);
                    if ("*" === a) {
                        for (; c = f[e++];) 1 === c.nodeType && d.push(c);
                        return d
                    }
                    return f
                }, w.find.CLASS = v.getElementsByClassName && function(a, b) {
                    return I ? b.getElementsByClassName(a) : void 0
                }, K = [], J = [], (v.qsa = ra.test(d.querySelectorAll)) && (e(function(a) {
                    H.appendChild(a).innerHTML = "<a id='" + N + "'></a><select id='" + N + "-\f]' msallowcapture=''><option selected=''></option></select>", a.querySelectorAll("[msallowcapture^='']").length && J.push("[*^$]=" + ca + "*(?:''|\"\")"), a.querySelectorAll("[selected]").length || J.push("\\[" + ca + "*(?:value|" + ba + ")"), a.querySelectorAll("[id~=" + N + "-]").length || J.push("~="), a.querySelectorAll(":checked").length || J.push(":checked"), a.querySelectorAll("a#" + N + "+*").length || J.push(".#.+[+~]")
                }), e(function(a) {
                    var b = d.createElement("input");
                    b.setAttribute("type", "hidden"), a.appendChild(b).setAttribute("name", "D"), a.querySelectorAll("[name=d]").length && J.push("name" + ca + "*[*^$|!~]?="), a.querySelectorAll(":enabled").length || J.push(":enabled", ":disabled"), a.querySelectorAll("*,:x"), J.push(",.*:")
                })), (v.matchesSelector = ra.test(L = H.matches || H.webkitMatchesSelector || H.mozMatchesSelector || H.oMatchesSelector || H.msMatchesSelector)) && e(function(a) {
                    v.disconnectedMatch = L.call(a, "div"), L.call(a, "[s!='']:x"), K.push("!=", ga)
                }), J = J.length && new RegExp(J.join("|")), K = K.length && new RegExp(K.join("|")), b = ra.test(H.compareDocumentPosition), M = b || ra.test(H.contains) ? function(a, b) {
                    var c = 9 === a.nodeType ? a.documentElement : a,
                        d = b && b.parentNode;
                    return a === d || !(!d || 1 !== d.nodeType || !(c.contains ? c.contains(d) : a.compareDocumentPosition && 16 & a.compareDocumentPosition(d)))
                } : function(a, b) {
                    if (b)
                        for (; b = b.parentNode;)
                            if (b === a) return !0;
                    return !1
                }, U = b ? function(a, b) {
                    if (a === b) return E = !0, 0;
                    var c = !a.compareDocumentPosition - !b.compareDocumentPosition;
                    return c ? c : (c = (a.ownerDocument || a) === (b.ownerDocument || b) ? a.compareDocumentPosition(b) : 1, 1 & c || !v.sortDetached && b.compareDocumentPosition(a) === c ? a === d || a.ownerDocument === O && M(O, a) ? -1 : b === d || b.ownerDocument === O && M(O, b) ? 1 : D ? aa(D, a) - aa(D, b) : 0 : 4 & c ? -1 : 1)
                } : function(a, b) {
                    if (a === b) return E = !0, 0;
                    var c, e = 0,
                        f = a.parentNode,
                        h = b.parentNode,
                        i = [a],
                        j = [b];
                    if (!f || !h) return a === d ? -1 : b === d ? 1 : f ? -1 : h ? 1 : D ? aa(D, a) - aa(D, b) : 0;
                    if (f === h) return g(a, b);
                    for (c = a; c = c.parentNode;) i.unshift(c);
                    for (c = b; c = c.parentNode;) j.unshift(c);
                    for (; i[e] === j[e];) e++;
                    return e ? g(i[e], j[e]) : i[e] === O ? -1 : j[e] === O ? 1 : 0
                }, d) : G
            }, b.matches = function(a, c) {
                return b(a, null, null, c)
            }, b.matchesSelector = function(a, c) {
                if ((a.ownerDocument || a) !== G && F(a), c = c.replace(la, "='$1']"), v.matchesSelector && I && (!K || !K.test(c)) && (!J || !J.test(c))) try {
                    var d = L.call(a, c);
                    if (d || v.disconnectedMatch || a.document && 11 !== a.document.nodeType) return d
                } catch (e) {}
                return b(c, G, null, [a]).length > 0
            }, b.contains = function(a, b) {
                return (a.ownerDocument || a) !== G && F(a), M(a, b)
            }, b.attr = function(a, b) {
                (a.ownerDocument || a) !== G && F(a);
                var c = w.attrHandle[b.toLowerCase()],
                    d = c && W.call(w.attrHandle, b.toLowerCase()) ? c(a, b, !I) : void 0;
                return void 0 !== d ? d : v.attributes || !I ? a.getAttribute(b) : (d = a.getAttributeNode(b)) && d.specified ? d.value : null
            }, b.error = function(a) {
                throw new Error("Syntax error, unrecognized expression: " + a)
            }, b.uniqueSort = function(a) {
                var b, c = [],
                    d = 0,
                    e = 0;
                if (E = !v.detectDuplicates, D = !v.sortStable && a.slice(0), a.sort(U), E) {
                    for (; b = a[e++];) b === a[e] && (d = c.push(e));
                    for (; d--;) a.splice(c[d], 1)
                }
                return D = null, a
            }, x = b.getText = function(a) {
                var b, c = "",
                    d = 0,
                    e = a.nodeType;
                if (e) {
                    if (1 === e || 9 === e || 11 === e) {
                        if ("string" == typeof a.textContent) return a.textContent;
                        for (a = a.firstChild; a; a = a.nextSibling) c += x(a)
                    } else if (3 === e || 4 === e) return a.nodeValue
                } else
                    for (; b = a[d++];) c += x(b);
                return c
            }, w = b.selectors = {
                cacheLength: 50,
                createPseudo: d,
                match: oa,
                attrHandle: {},
                find: {},
                relative: {
                    ">": {
                        dir: "parentNode",
                        first: !0
                    },
                    " ": {
                        dir: "parentNode"
                    },
                    "+": {
                        dir: "previousSibling",
                        first: !0
                    },
                    "~": {
                        dir: "previousSibling"
                    }
                },
                preFilter: {
                    ATTR: function(a) {
                        return a[1] = a[1].replace(va, wa), a[3] = (a[3] || a[4] || a[5] || "").replace(va, wa), "~=" === a[2] && (a[3] = " " + a[3] + " "), a.slice(0, 4)
                    },
                    CHILD: function(a) {
                        return a[1] = a[1].toLowerCase(), "nth" === a[1].slice(0, 3) ? (a[3] || b.error(a[0]), a[4] = +(a[4] ? a[5] + (a[6] || 1) : 2 * ("even" === a[3] || "odd" === a[3])), a[5] = +(a[7] + a[8] || "odd" === a[3])) : a[3] && b.error(a[0]), a
                    },
                    PSEUDO: function(a) {
                        var b, c = !a[6] && a[2];
                        return oa.CHILD.test(a[0]) ? null : (a[3] ? a[2] = a[4] || a[5] || "" : c && ma.test(c) && (b = z(c, !0)) && (b = c.indexOf(")", c.length - b) - c.length) && (a[0] = a[0].slice(0, b), a[2] = c.slice(0, b)), a.slice(0, 3))
                    }
                },
                filter: {
                    TAG: function(a) {
                        var b = a.replace(va, wa).toLowerCase();
                        return "*" === a ? function() {
                            return !0
                        } : function(a) {
                            return a.nodeName && a.nodeName.toLowerCase() === b
                        }
                    },
                    CLASS: function(a) {
                        var b = R[a + " "];
                        return b || (b = new RegExp("(^|" + ca + ")" + a + "(" + ca + "|$)")) && R(a, function(a) {
                            return b.test("string" == typeof a.className && a.className || "undefined" != typeof a.getAttribute && a.getAttribute("class") || "")
                        })
                    },
                    ATTR: function(a, c, d) {
                        return function(e) {
                            var f = b.attr(e, a);
                            return null == f ? "!=" === c : c ? (f += "", "=" === c ? f === d : "!=" === c ? f !== d : "^=" === c ? d && 0 === f.indexOf(d) : "*=" === c ? d && f.indexOf(d) > -1 : "$=" === c ? d && f.slice(-d.length) === d : "~=" === c ? (" " + f.replace(ha, " ") + " ").indexOf(d) > -1 : "|=" === c ? f === d || f.slice(0, d.length + 1) === d + "-" : !1) : !0
                        }
                    },
                    CHILD: function(a, b, c, d, e) {
                        var f = "nth" !== a.slice(0, 3),
                            g = "last" !== a.slice(-4),
                            h = "of-type" === b;
                        return 1 === d && 0 === e ? function(a) {
                            return !!a.parentNode
                        } : function(b, c, i) {
                            var j, k, l, m, n, o, p = f !== g ? "nextSibling" : "previousSibling",
                                q = b.parentNode,
                                r = h && b.nodeName.toLowerCase(),
                                s = !i && !h;
                            if (q) {
                                if (f) {
                                    for (; p;) {
                                        for (l = b; l = l[p];)
                                            if (h ? l.nodeName.toLowerCase() === r : 1 === l.nodeType) return !1;
                                        o = p = "only" === a && !o && "nextSibling"
                                    }
                                    return !0
                                }
                                if (o = [g ? q.firstChild : q.lastChild], g && s) {
                                    for (k = q[N] || (q[N] = {}), j = k[a] || [], n = j[0] === P && j[1], m = j[0] === P && j[2], l = n && q.childNodes[n]; l = ++n && l && l[p] || (m = n = 0) || o.pop();)
                                        if (1 === l.nodeType && ++m && l === b) {
                                            k[a] = [P, n, m];
                                            break
                                        }
                                } else if (s && (j = (b[N] || (b[N] = {}))[a]) && j[0] === P) m = j[1];
                                else
                                    for (;
                                        (l = ++n && l && l[p] || (m = n = 0) || o.pop()) && ((h ? l.nodeName.toLowerCase() !== r : 1 !== l.nodeType) || !++m || (s && ((l[N] || (l[N] = {}))[a] = [P, m]), l !== b)););
                                return m -= e, m === d || m % d === 0 && m / d >= 0
                            }
                        }
                    },
                    PSEUDO: function(a, c) {
                        var e, f = w.pseudos[a] || w.setFilters[a.toLowerCase()] || b.error("unsupported pseudo: " + a);
                        return f[N] ? f(c) : f.length > 1 ? (e = [a, a, "", c], w.setFilters.hasOwnProperty(a.toLowerCase()) ? d(function(a, b) {
                            for (var d, e = f(a, c), g = e.length; g--;) d = aa(a, e[g]), a[d] = !(b[d] = e[g])
                        }) : function(a) {
                            return f(a, 0, e)
                        }) : f
                    }
                },
                pseudos: {
                    not: d(function(a) {
                        var b = [],
                            c = [],
                            e = A(a.replace(ia, "$1"));
                        return e[N] ? d(function(a, b, c, d) {
                            for (var f, g = e(a, null, d, []), h = a.length; h--;)(f = g[h]) && (a[h] = !(b[h] = f))
                        }) : function(a, d, f) {
                            return b[0] = a, e(b, null, f, c), b[0] = null, !c.pop()
                        }
                    }),
                    has: d(function(a) {
                        return function(c) {
                            return b(a, c).length > 0
                        }
                    }),
                    contains: d(function(a) {
                        return a = a.replace(va, wa),
                            function(b) {
                                return (b.textContent || b.innerText || x(b)).indexOf(a) > -1
                            }
                    }),
                    lang: d(function(a) {
                        return na.test(a || "") || b.error("unsupported lang: " + a), a = a.replace(va, wa).toLowerCase(),
                            function(b) {
                                var c;
                                do
                                    if (c = I ? b.lang : b.getAttribute("xml:lang") || b.getAttribute("lang")) return c = c.toLowerCase(), c === a || 0 === c.indexOf(a + "-"); while ((b = b.parentNode) && 1 === b.nodeType);
                                return !1
                            }
                    }),
                    target: function(b) {
                        var c = a.location && a.location.hash;
                        return c && c.slice(1) === b.id
                    },
                    root: function(a) {
                        return a === H
                    },
                    focus: function(a) {
                        return a === G.activeElement && (!G.hasFocus || G.hasFocus()) && !!(a.type || a.href || ~a.tabIndex)
                    },
                    enabled: function(a) {
                        return a.disabled === !1
                    },
                    disabled: function(a) {
                        return a.disabled === !0
                    },
                    checked: function(a) {
                        var b = a.nodeName.toLowerCase();
                        return "input" === b && !!a.checked || "option" === b && !!a.selected
                    },
                    selected: function(a) {
                        return a.parentNode && a.parentNode.selectedIndex, a.selected === !0
                    },
                    empty: function(a) {
                        for (a = a.firstChild; a; a = a.nextSibling)
                            if (a.nodeType < 6) return !1;
                        return !0
                    },
                    parent: function(a) {
                        return !w.pseudos.empty(a)
                    },
                    header: function(a) {
                        return qa.test(a.nodeName)
                    },
                    input: function(a) {
                        return pa.test(a.nodeName)
                    },
                    button: function(a) {
                        var b = a.nodeName.toLowerCase();
                        return "input" === b && "button" === a.type || "button" === b
                    },
                    text: function(a) {
                        var b;
                        return "input" === a.nodeName.toLowerCase() && "text" === a.type && (null == (b = a.getAttribute("type")) || "text" === b.toLowerCase())
                    },
                    first: j(function() {
                        return [0]
                    }),
                    last: j(function(a, b) {
                        return [b - 1]
                    }),
                    eq: j(function(a, b, c) {
                        return [0 > c ? c + b : c]
                    }),
                    even: j(function(a, b) {
                        for (var c = 0; b > c; c += 2) a.push(c);
                        return a
                    }),
                    odd: j(function(a, b) {
                        for (var c = 1; b > c; c += 2) a.push(c);
                        return a
                    }),
                    lt: j(function(a, b, c) {
                        for (var d = 0 > c ? c + b : c; --d >= 0;) a.push(d);
                        return a
                    }),
                    gt: j(function(a, b, c) {
                        for (var d = 0 > c ? c + b : c; ++d < b;) a.push(d);
                        return a
                    })
                }
            }, w.pseudos.nth = w.pseudos.eq;
            for (u in {
                    radio: !0,
                    checkbox: !0,
                    file: !0,
                    password: !0,
                    image: !0
                }) w.pseudos[u] = h(u);
            for (u in {
                    submit: !0,
                    reset: !0
                }) w.pseudos[u] = i(u);
            return l.prototype = w.filters = w.pseudos, w.setFilters = new l, z = b.tokenize = function(a, c) {
                var d, e, f, g, h, i, j, k = S[a + " "];
                if (k) return c ? 0 : k.slice(0);
                for (h = a, i = [], j = w.preFilter; h;) {
                    (!d || (e = ja.exec(h))) && (e && (h = h.slice(e[0].length) || h), i.push(f = [])), d = !1, (e = ka.exec(h)) && (d = e.shift(), f.push({
                        value: d,
                        type: e[0].replace(ia, " ")
                    }), h = h.slice(d.length));
                    for (g in w.filter) !(e = oa[g].exec(h)) || j[g] && !(e = j[g](e)) || (d = e.shift(), f.push({
                        value: d,
                        type: g,
                        matches: e
                    }), h = h.slice(d.length));
                    if (!d) break
                }
                return c ? h.length : h ? b.error(a) : S(a, i).slice(0)
            }, A = b.compile = function(a, b) {
                var c, d = [],
                    e = [],
                    f = T[a + " "];
                if (!f) {
                    for (b || (b = z(a)), c = b.length; c--;) f = s(b[c]), f[N] ? d.push(f) : e.push(f);
                    f = T(a, t(e, d)), f.selector = a
                }
                return f
            }, B = b.select = function(a, b, c, d) {
                var e, f, g, h, i, j = "function" == typeof a && a,
                    l = !d && z(a = j.selector || a);
                if (c = c || [], 1 === l.length) {
                    if (f = l[0] = l[0].slice(0), f.length > 2 && "ID" === (g = f[0]).type && v.getById && 9 === b.nodeType && I && w.relative[f[1].type]) {
                        if (b = (w.find.ID(g.matches[0].replace(va, wa), b) || [])[0], !b) return c;
                        j && (b = b.parentNode), a = a.slice(f.shift().value.length)
                    }
                    for (e = oa.needsContext.test(a) ? 0 : f.length; e-- && (g = f[e], !w.relative[h = g.type]);)
                        if ((i = w.find[h]) && (d = i(g.matches[0].replace(va, wa), ta.test(f[0].type) && k(b.parentNode) || b))) {
                            if (f.splice(e, 1), a = d.length && m(f), !a) return $.apply(c, d), c;
                            break
                        }
                }
                return (j || A(a, l))(d, b, !I, c, ta.test(a) && k(b.parentNode) || b), c
            }, v.sortStable = N.split("").sort(U).join("") === N, v.detectDuplicates = !!E, F(), v.sortDetached = e(function(a) {
                return 1 & a.compareDocumentPosition(G.createElement("div"))
            }), e(function(a) {
                return a.innerHTML = "<a href='#'></a>", "#" === a.firstChild.getAttribute("href")
            }) || f("type|href|height|width", function(a, b, c) {
                return c ? void 0 : a.getAttribute(b, "type" === b.toLowerCase() ? 1 : 2)
            }), v.attributes && e(function(a) {
                return a.innerHTML = "<input/>", a.firstChild.setAttribute("value", ""), "" === a.firstChild.getAttribute("value")
            }) || f("value", function(a, b, c) {
                return c || "input" !== a.nodeName.toLowerCase() ? void 0 : a.defaultValue
            }), e(function(a) {
                return null == a.getAttribute("disabled")
            }) || f(ba, function(a, b, c) {
                var d;
                return c ? void 0 : a[b] === !0 ? b.toLowerCase() : (d = a.getAttributeNode(b)) && d.specified ? d.value : null
            }), b
        }(a);
        _.find = ea, _.expr = ea.selectors, _.expr[":"] = _.expr.pseudos, _.unique = ea.uniqueSort, _.text = ea.getText, _.isXMLDoc = ea.isXML, _.contains = ea.contains;
        var fa = _.expr.match.needsContext,
            ga = /^<(\w+)\s*\/?>(?:<\/\1>|)$/,
            ha = /^.[^:#\[\.,]*$/;
        _.filter = function(a, b, c) {
            var d = b[0];
            return c && (a = ":not(" + a + ")"), 1 === b.length && 1 === d.nodeType ? _.find.matchesSelector(d, a) ? [d] : [] : _.find.matches(a, _.grep(b, function(a) {
                return 1 === a.nodeType
            }))
        }, _.fn.extend({
            find: function(a) {
                var b, c = this.length,
                    d = [],
                    e = this;
                if ("string" != typeof a) return this.pushStack(_(a).filter(function() {
                    for (b = 0; c > b; b++)
                        if (_.contains(e[b], this)) return !0
                }));
                for (b = 0; c > b; b++) _.find(a, e[b], d);
                return d = this.pushStack(c > 1 ? _.unique(d) : d), d.selector = this.selector ? this.selector + " " + a : a, d
            },
            filter: function(a) {
                return this.pushStack(d(this, a || [], !1))
            },
            not: function(a) {
                return this.pushStack(d(this, a || [], !0))
            },
            is: function(a) {
                return !!d(this, "string" == typeof a && fa.test(a) ? _(a) : a || [], !1).length
            }
        });
        var ia, ja = /^(?:\s*(<[\w\W]+>)[^>]*|#([\w-]*))$/,
            ka = _.fn.init = function(a, b) {
                var c, d;
                if (!a) return this;
                if ("string" == typeof a) {
                    if (c = "<" === a[0] && ">" === a[a.length - 1] && a.length >= 3 ? [null, a, null] : ja.exec(a), !c || !c[1] && b) return !b || b.jquery ? (b || ia).find(a) : this.constructor(b).find(a);
                    if (c[1]) {
                        if (b = b instanceof _ ? b[0] : b, _.merge(this, _.parseHTML(c[1], b && b.nodeType ? b.ownerDocument || b : Z, !0)), ga.test(c[1]) && _.isPlainObject(b))
                            for (c in b) _.isFunction(this[c]) ? this[c](b[c]) : this.attr(c, b[c]);
                        return this
                    }
                    return d = Z.getElementById(c[2]), d && d.parentNode && (this.length = 1, this[0] = d), this.context = Z, this.selector = a, this
                }
                return a.nodeType ? (this.context = this[0] = a, this.length = 1, this) : _.isFunction(a) ? "undefined" != typeof ia.ready ? ia.ready(a) : a(_) : (void 0 !== a.selector && (this.selector = a.selector, this.context = a.context), _.makeArray(a, this))
            };
        ka.prototype = _.fn, ia = _(Z);
        var la = /^(?:parents|prev(?:Until|All))/,
            ma = {
                children: !0,
                contents: !0,
                next: !0,
                prev: !0
            };
        _.extend({
            dir: function(a, b, c) {
                for (var d = [], e = void 0 !== c;
                    (a = a[b]) && 9 !== a.nodeType;)
                    if (1 === a.nodeType) {
                        if (e && _(a).is(c)) break;
                        d.push(a)
                    }
                return d
            },
            sibling: function(a, b) {
                for (var c = []; a; a = a.nextSibling) 1 === a.nodeType && a !== b && c.push(a);
                return c
            }
        }), _.fn.extend({
            has: function(a) {
                var b = _(a, this),
                    c = b.length;
                return this.filter(function() {
                    for (var a = 0; c > a; a++)
                        if (_.contains(this, b[a])) return !0
                })
            },
            closest: function(a, b) {
                for (var c, d = 0, e = this.length, f = [], g = fa.test(a) || "string" != typeof a ? _(a, b || this.context) : 0; e > d; d++)
                    for (c = this[d]; c && c !== b; c = c.parentNode)
                        if (c.nodeType < 11 && (g ? g.index(c) > -1 : 1 === c.nodeType && _.find.matchesSelector(c, a))) {
                            f.push(c);
                            break
                        }
                return this.pushStack(f.length > 1 ? _.unique(f) : f)
            },
            index: function(a) {
                return a ? "string" == typeof a ? U.call(_(a), this[0]) : U.call(this, a.jquery ? a[0] : a) : this[0] && this[0].parentNode ? this.first().prevAll().length : -1
            },
            add: function(a, b) {
                return this.pushStack(_.unique(_.merge(this.get(), _(a, b))))
            },
            addBack: function(a) {
                return this.add(null == a ? this.prevObject : this.prevObject.filter(a))
            }
        }), _.each({
            parent: function(a) {
                var b = a.parentNode;
                return b && 11 !== b.nodeType ? b : null
            },
            parents: function(a) {
                return _.dir(a, "parentNode")
            },
            parentsUntil: function(a, b, c) {
                return _.dir(a, "parentNode", c)
            },
            next: function(a) {
                return e(a, "nextSibling")
            },
            prev: function(a) {
                return e(a, "previousSibling")
            },
            nextAll: function(a) {
                return _.dir(a, "nextSibling")
            },
            prevAll: function(a) {
                return _.dir(a, "previousSibling")
            },
            nextUntil: function(a, b, c) {
                return _.dir(a, "nextSibling", c)
            },
            prevUntil: function(a, b, c) {
                return _.dir(a, "previousSibling", c)
            },
            siblings: function(a) {
                return _.sibling((a.parentNode || {}).firstChild, a)
            },
            children: function(a) {
                return _.sibling(a.firstChild)
            },
            contents: function(a) {
                return a.contentDocument || _.merge([], a.childNodes)
            }
        }, function(a, b) {
            _.fn[a] = function(c, d) {
                var e = _.map(this, b, c);
                return "Until" !== a.slice(-5) && (d = c), d && "string" == typeof d && (e = _.filter(d, e)), this.length > 1 && (ma[a] || _.unique(e), la.test(a) && e.reverse()), this.pushStack(e)
            }
        });
        var na = /\S+/g,
            oa = {};
        _.Callbacks = function(a) {
            a = "string" == typeof a ? oa[a] || f(a) : _.extend({}, a);
            var b, c, d, e, g, h, i = [],
                j = !a.once && [],
                k = function(f) {
                    for (b = a.memory && f, c = !0, h = e || 0, e = 0, g = i.length, d = !0; i && g > h; h++)
                        if (i[h].apply(f[0], f[1]) === !1 && a.stopOnFalse) {
                            b = !1;
                            break
                        }
                    d = !1, i && (j ? j.length && k(j.shift()) : b ? i = [] : l.disable())
                },
                l = {
                    add: function() {
                        if (i) {
                            var c = i.length;
                            ! function f(b) {
                                _.each(b, function(b, c) {
                                    var d = _.type(c);
                                    "function" === d ? a.unique && l.has(c) || i.push(c) : c && c.length && "string" !== d && f(c)
                                })
                            }(arguments), d ? g = i.length : b && (e = c, k(b))
                        }
                        return this
                    },
                    remove: function() {
                        return i && _.each(arguments, function(a, b) {
                            for (var c;
                                (c = _.inArray(b, i, c)) > -1;) i.splice(c, 1), d && (g >= c && g--, h >= c && h--)
                        }), this
                    },
                    has: function(a) {
                        return a ? _.inArray(a, i) > -1 : !(!i || !i.length)
                    },
                    empty: function() {
                        return i = [], g = 0, this
                    },
                    disable: function() {
                        return i = j = b = void 0, this
                    },
                    disabled: function() {
                        return !i
                    },
                    lock: function() {
                        return j = void 0, b || l.disable(), this
                    },
                    locked: function() {
                        return !j
                    },
                    fireWith: function(a, b) {
                        return !i || c && !j || (b = b || [], b = [a, b.slice ? b.slice() : b], d ? j.push(b) : k(b)), this
                    },
                    fire: function() {
                        return l.fireWith(this, arguments), this
                    },
                    fired: function() {
                        return !!c
                    }
                };
            return l
        }, _.extend({
            Deferred: function(a) {
                var b = [
                        ["resolve", "done", _.Callbacks("once memory"), "resolved"],
                        ["reject", "fail", _.Callbacks("once memory"), "rejected"],
                        ["notify", "progress", _.Callbacks("memory")]
                    ],
                    c = "pending",
                    d = {
                        state: function() {
                            return c
                        },
                        always: function() {
                            return e.done(arguments).fail(arguments), this
                        },
                        then: function() {
                            var a = arguments;
                            return _.Deferred(function(c) {
                                _.each(b, function(b, f) {
                                    var g = _.isFunction(a[b]) && a[b];
                                    e[f[1]](function() {
                                        var a = g && g.apply(this, arguments);
                                        a && _.isFunction(a.promise) ? a.promise().done(c.resolve).fail(c.reject).progress(c.notify) : c[f[0] + "With"](this === d ? c.promise() : this, g ? [a] : arguments)
                                    })
                                }), a = null
                            }).promise()
                        },
                        promise: function(a) {
                            return null != a ? _.extend(a, d) : d
                        }
                    },
                    e = {};
                return d.pipe = d.then, _.each(b, function(a, f) {
                    var g = f[2],
                        h = f[3];
                    d[f[1]] = g.add, h && g.add(function() {
                        c = h
                    }, b[1 ^ a][2].disable, b[2][2].lock), e[f[0]] = function() {
                        return e[f[0] + "With"](this === e ? d : this, arguments), this
                    }, e[f[0] + "With"] = g.fireWith
                }), d.promise(e), a && a.call(e, e), e
            },
            when: function(a) {
                var b, c, d, e = 0,
                    f = R.call(arguments),
                    g = f.length,
                    h = 1 !== g || a && _.isFunction(a.promise) ? g : 0,
                    i = 1 === h ? a : _.Deferred(),
                    j = function(a, c, d) {
                        return function(e) {
                            c[a] = this, d[a] = arguments.length > 1 ? R.call(arguments) : e, d === b ? i.notifyWith(c, d) : --h || i.resolveWith(c, d)
                        }
                    };
                if (g > 1)
                    for (b = new Array(g), c = new Array(g), d = new Array(g); g > e; e++) f[e] && _.isFunction(f[e].promise) ? f[e].promise().done(j(e, d, f)).fail(i.reject).progress(j(e, c, b)) : --h;
                return h || i.resolveWith(d, f), i.promise()
            }
        });
        var pa;
        _.fn.ready = function(a) {
            return _.ready.promise().done(a), this
        }, _.extend({
            isReady: !1,
            readyWait: 1,
            holdReady: function(a) {
                a ? _.readyWait++ : _.ready(!0)
            },
            ready: function(a) {
                (a === !0 ? --_.readyWait : _.isReady) || (_.isReady = !0, a !== !0 && --_.readyWait > 0 || (pa.resolveWith(Z, [_]), _.fn.triggerHandler && (_(Z).triggerHandler("ready"), _(Z).off("ready"))))
            }
        }), _.ready.promise = function(b) {
            return pa || (pa = _.Deferred(), "complete" === Z.readyState ? setTimeout(_.ready) : (Z.addEventListener("DOMContentLoaded", g, !1), a.addEventListener("load", g, !1))), pa.promise(b)
        }, _.ready.promise();
        var qa = _.access = function(a, b, c, d, e, f, g) {
            var h = 0,
                i = a.length,
                j = null == c;
            if ("object" === _.type(c)) {
                e = !0;
                for (h in c) _.access(a, b, h, c[h], !0, f, g)
            } else if (void 0 !== d && (e = !0, _.isFunction(d) || (g = !0), j && (g ? (b.call(a, d), b = null) : (j = b, b = function(a, b, c) {
                    return j.call(_(a), c)
                })), b))
                for (; i > h; h++) b(a[h], c, g ? d : d.call(a[h], h, b(a[h], c)));
            return e ? a : j ? b.call(a) : i ? b(a[0], c) : f
        };
        _.acceptData = function(a) {
            return 1 === a.nodeType || 9 === a.nodeType || !+a.nodeType
        }, h.uid = 1, h.accepts = _.acceptData, h.prototype = {
            key: function(a) {
                if (!h.accepts(a)) return 0;
                var b = {},
                    c = a[this.expando];
                if (!c) {
                    c = h.uid++;
                    try {
                        b[this.expando] = {
                            value: c
                        }, Object.defineProperties(a, b)
                    } catch (d) {
                        b[this.expando] = c, _.extend(a, b)
                    }
                }
                return this.cache[c] || (this.cache[c] = {}), c
            },
            set: function(a, b, c) {
                var d, e = this.key(a),
                    f = this.cache[e];
                if ("string" == typeof b) f[b] = c;
                else if (_.isEmptyObject(f)) _.extend(this.cache[e], b);
                else
                    for (d in b) f[d] = b[d];
                return f
            },
            get: function(a, b) {
                var c = this.cache[this.key(a)];
                return void 0 === b ? c : c[b]
            },
            access: function(a, b, c) {
                var d;
                return void 0 === b || b && "string" == typeof b && void 0 === c ? (d = this.get(a, b), void 0 !== d ? d : this.get(a, _.camelCase(b))) : (this.set(a, b, c), void 0 !== c ? c : b)
            },
            remove: function(a, b) {
                var c, d, e, f = this.key(a),
                    g = this.cache[f];
                if (void 0 === b) this.cache[f] = {};
                else {
                    _.isArray(b) ? d = b.concat(b.map(_.camelCase)) : (e = _.camelCase(b), b in g ? d = [b, e] : (d = e, d = d in g ? [d] : d.match(na) || [])), c = d.length;
                    for (; c--;) delete g[d[c]]
                }
            },
            hasData: function(a) {
                return !_.isEmptyObject(this.cache[a[this.expando]] || {})
            },
            discard: function(a) {
                a[this.expando] && delete this.cache[a[this.expando]]
            }
        };
        var ra = new h,
            sa = new h,
            ta = /^(?:\{[\w\W]*\}|\[[\w\W]*\])$/,
            ua = /([A-Z])/g;
        _.extend({
            hasData: function(a) {
                return sa.hasData(a) || ra.hasData(a)
            },
            data: function(a, b, c) {
                return sa.access(a, b, c)
            },
            removeData: function(a, b) {
                sa.remove(a, b)
            },
            _data: function(a, b, c) {
                return ra.access(a, b, c)
            },
            _removeData: function(a, b) {
                ra.remove(a, b)
            }
        }), _.fn.extend({
            data: function(a, b) {
                var c, d, e, f = this[0],
                    g = f && f.attributes;
                if (void 0 === a) {
                    if (this.length && (e = sa.get(f), 1 === f.nodeType && !ra.get(f, "hasDataAttrs"))) {
                        for (c = g.length; c--;) g[c] && (d = g[c].name, 0 === d.indexOf("data-") && (d = _.camelCase(d.slice(5)), i(f, d, e[d])));
                        ra.set(f, "hasDataAttrs", !0)
                    }
                    return e
                }
                return "object" == typeof a ? this.each(function() {
                    sa.set(this, a)
                }) : qa(this, function(b) {
                    var c, d = _.camelCase(a);
                    if (f && void 0 === b) {
                        if (c = sa.get(f, a), void 0 !== c) return c;
                        if (c = sa.get(f, d), void 0 !== c) return c;
                        if (c = i(f, d, void 0), void 0 !== c) return c
                    } else this.each(function() {
                        var c = sa.get(this, d);
                        sa.set(this, d, b), -1 !== a.indexOf("-") && void 0 !== c && sa.set(this, a, b)
                    })
                }, null, b, arguments.length > 1, null, !0)
            },
            removeData: function(a) {
                return this.each(function() {
                    sa.remove(this, a)
                })
            }
        }), _.extend({
            queue: function(a, b, c) {
                var d;
                return a ? (b = (b || "fx") + "queue", d = ra.get(a, b), c && (!d || _.isArray(c) ? d = ra.access(a, b, _.makeArray(c)) : d.push(c)), d || []) : void 0
            },
            dequeue: function(a, b) {
                b = b || "fx";
                var c = _.queue(a, b),
                    d = c.length,
                    e = c.shift(),
                    f = _._queueHooks(a, b),
                    g = function() {
                        _.dequeue(a, b)
                    };
                "inprogress" === e && (e = c.shift(), d--), e && ("fx" === b && c.unshift("inprogress"), delete f.stop, e.call(a, g, f)), !d && f && f.empty.fire()
            },
            _queueHooks: function(a, b) {
                var c = b + "queueHooks";
                return ra.get(a, c) || ra.access(a, c, {
                    empty: _.Callbacks("once memory").add(function() {
                        ra.remove(a, [b + "queue", c])
                    })
                })
            }
        }), _.fn.extend({
            queue: function(a, b) {
                var c = 2;
                return "string" != typeof a && (b = a, a = "fx", c--), arguments.length < c ? _.queue(this[0], a) : void 0 === b ? this : this.each(function() {
                    var c = _.queue(this, a, b);
                    _._queueHooks(this, a), "fx" === a && "inprogress" !== c[0] && _.dequeue(this, a)
                })
            },
            dequeue: function(a) {
                return this.each(function() {
                    _.dequeue(this, a)
                })
            },
            clearQueue: function(a) {
                return this.queue(a || "fx", [])
            },
            promise: function(a, b) {
                var c, d = 1,
                    e = _.Deferred(),
                    f = this,
                    g = this.length,
                    h = function() {
                        --d || e.resolveWith(f, [f])
                    };
                for ("string" != typeof a && (b = a, a = void 0), a = a || "fx"; g--;) c = ra.get(f[g], a + "queueHooks"), c && c.empty && (d++, c.empty.add(h));
                return h(), e.promise(b)
            }
        });
        var va = /[+-]?(?:\d*\.|)\d+(?:[eE][+-]?\d+|)/.source,
            wa = ["Top", "Right", "Bottom", "Left"],
            xa = function(a, b) {
                return a = b || a, "none" === _.css(a, "display") || !_.contains(a.ownerDocument, a)
            },
            ya = /^(?:checkbox|radio)$/i;
        ! function() {
            var a = Z.createDocumentFragment(),
                b = a.appendChild(Z.createElement("div")),
                c = Z.createElement("input");
            c.setAttribute("type", "radio"), c.setAttribute("checked", "checked"), c.setAttribute("name", "t"), b.appendChild(c), Y.checkClone = b.cloneNode(!0).cloneNode(!0).lastChild.checked, b.innerHTML = "<textarea>x</textarea>", Y.noCloneChecked = !!b.cloneNode(!0).lastChild.defaultValue
        }();
        var za = "undefined";
        Y.focusinBubbles = "onfocusin" in a;
        var Aa = /^key/,
            Ba = /^(?:mouse|pointer|contextmenu)|click/,
            Ca = /^(?:focusinfocus|focusoutblur)$/,
            Da = /^([^.]*)(?:\.(.+)|)$/;
        _.event = {
            global: {},
            add: function(a, b, c, d, e) {
                var f, g, h, i, j, k, l, m, n, o, p, q = ra.get(a);
                if (q)
                    for (c.handler && (f = c, c = f.handler, e = f.selector), c.guid || (c.guid = _.guid++), (i = q.events) || (i = q.events = {}), (g = q.handle) || (g = q.handle = function(b) {
                            return typeof _ !== za && _.event.triggered !== b.type ? _.event.dispatch.apply(a, arguments) : void 0
                        }), b = (b || "").match(na) || [""], j = b.length; j--;) h = Da.exec(b[j]) || [], n = p = h[1], o = (h[2] || "").split(".").sort(), n && (l = _.event.special[n] || {}, n = (e ? l.delegateType : l.bindType) || n, l = _.event.special[n] || {}, k = _.extend({
                        type: n,
                        origType: p,
                        data: d,
                        handler: c,
                        guid: c.guid,
                        selector: e,
                        needsContext: e && _.expr.match.needsContext.test(e),
                        namespace: o.join(".")
                    }, f), (m = i[n]) || (m = i[n] = [], m.delegateCount = 0, l.setup && l.setup.call(a, d, o, g) !== !1 || a.addEventListener && a.addEventListener(n, g, !1)), l.add && (l.add.call(a, k), k.handler.guid || (k.handler.guid = c.guid)), e ? m.splice(m.delegateCount++, 0, k) : m.push(k), _.event.global[n] = !0)
            },
            remove: function(a, b, c, d, e) {
                var f, g, h, i, j, k, l, m, n, o, p, q = ra.hasData(a) && ra.get(a);
                if (q && (i = q.events)) {
                    for (b = (b || "").match(na) || [""], j = b.length; j--;)
                        if (h = Da.exec(b[j]) || [], n = p = h[1], o = (h[2] || "").split(".").sort(), n) {
                            for (l = _.event.special[n] || {}, n = (d ? l.delegateType : l.bindType) || n, m = i[n] || [], h = h[2] && new RegExp("(^|\\.)" + o.join("\\.(?:.*\\.|)") + "(\\.|$)"), g = f = m.length; f--;) k = m[f], !e && p !== k.origType || c && c.guid !== k.guid || h && !h.test(k.namespace) || d && d !== k.selector && ("**" !== d || !k.selector) || (m.splice(f, 1), k.selector && m.delegateCount--, l.remove && l.remove.call(a, k));
                            g && !m.length && (l.teardown && l.teardown.call(a, o, q.handle) !== !1 || _.removeEvent(a, n, q.handle), delete i[n])
                        } else
                            for (n in i) _.event.remove(a, n + b[j], c, d, !0);
                    _.isEmptyObject(i) && (delete q.handle, ra.remove(a, "events"))
                }
            },
            trigger: function(b, c, d, e) {
                var f, g, h, i, j, k, l, m = [d || Z],
                    n = X.call(b, "type") ? b.type : b,
                    o = X.call(b, "namespace") ? b.namespace.split(".") : [];
                if (g = h = d = d || Z, 3 !== d.nodeType && 8 !== d.nodeType && !Ca.test(n + _.event.triggered) && (n.indexOf(".") >= 0 && (o = n.split("."), n = o.shift(), o.sort()), j = n.indexOf(":") < 0 && "on" + n, b = b[_.expando] ? b : new _.Event(n, "object" == typeof b && b), b.isTrigger = e ? 2 : 3, b.namespace = o.join("."), b.namespace_re = b.namespace ? new RegExp("(^|\\.)" + o.join("\\.(?:.*\\.|)") + "(\\.|$)") : null, b.result = void 0, b.target || (b.target = d), c = null == c ? [b] : _.makeArray(c, [b]), l = _.event.special[n] || {}, e || !l.trigger || l.trigger.apply(d, c) !== !1)) {
                    if (!e && !l.noBubble && !_.isWindow(d)) {
                        for (i = l.delegateType || n, Ca.test(i + n) || (g = g.parentNode); g; g = g.parentNode) m.push(g), h = g;
                        h === (d.ownerDocument || Z) && m.push(h.defaultView || h.parentWindow || a)
                    }
                    for (f = 0;
                        (g = m[f++]) && !b.isPropagationStopped();) b.type = f > 1 ? i : l.bindType || n, k = (ra.get(g, "events") || {})[b.type] && ra.get(g, "handle"), k && k.apply(g, c), k = j && g[j], k && k.apply && _.acceptData(g) && (b.result = k.apply(g, c), b.result === !1 && b.preventDefault());
                    return b.type = n, e || b.isDefaultPrevented() || l._default && l._default.apply(m.pop(), c) !== !1 || !_.acceptData(d) || j && _.isFunction(d[n]) && !_.isWindow(d) && (h = d[j], h && (d[j] = null), _.event.triggered = n, d[n](), _.event.triggered = void 0, h && (d[j] = h)), b.result
                }
            },
            dispatch: function(a) {
                a = _.event.fix(a);
                var b, c, d, e, f, g = [],
                    h = R.call(arguments),
                    i = (ra.get(this, "events") || {})[a.type] || [],
                    j = _.event.special[a.type] || {};
                if (h[0] = a, a.delegateTarget = this, !j.preDispatch || j.preDispatch.call(this, a) !== !1) {
                    for (g = _.event.handlers.call(this, a, i), b = 0;
                        (e = g[b++]) && !a.isPropagationStopped();)
                        for (a.currentTarget = e.elem, c = 0;
                            (f = e.handlers[c++]) && !a.isImmediatePropagationStopped();)(!a.namespace_re || a.namespace_re.test(f.namespace)) && (a.handleObj = f, a.data = f.data, d = ((_.event.special[f.origType] || {}).handle || f.handler).apply(e.elem, h), void 0 !== d && (a.result = d) === !1 && (a.preventDefault(), a.stopPropagation()));
                    return j.postDispatch && j.postDispatch.call(this, a), a.result
                }
            },
            handlers: function(a, b) {
                var c, d, e, f, g = [],
                    h = b.delegateCount,
                    i = a.target;
                if (h && i.nodeType && (!a.button || "click" !== a.type))
                    for (; i !== this; i = i.parentNode || this)
                        if (i.disabled !== !0 || "click" !== a.type) {
                            for (d = [], c = 0; h > c; c++) f = b[c], e = f.selector + " ", void 0 === d[e] && (d[e] = f.needsContext ? _(e, this).index(i) >= 0 : _.find(e, this, null, [i]).length), d[e] && d.push(f);
                            d.length && g.push({
                                elem: i,
                                handlers: d
                            })
                        }
                return h < b.length && g.push({
                    elem: this,
                    handlers: b.slice(h)
                }), g
            },
            props: "altKey bubbles cancelable ctrlKey currentTarget eventPhase metaKey relatedTarget shiftKey target timeStamp view which".split(" "),
            fixHooks: {},
            keyHooks: {
                props: "char charCode key keyCode".split(" "),
                filter: function(a, b) {
                    return null == a.which && (a.which = null != b.charCode ? b.charCode : b.keyCode), a
                }
            },
            mouseHooks: {
                props: "button buttons clientX clientY offsetX offsetY pageX pageY screenX screenY toElement".split(" "),
                filter: function(a, b) {
                    var c, d, e, f = b.button;
                    return null == a.pageX && null != b.clientX && (c = a.target.ownerDocument || Z, d = c.documentElement, e = c.body, a.pageX = b.clientX + (d && d.scrollLeft || e && e.scrollLeft || 0) - (d && d.clientLeft || e && e.clientLeft || 0), a.pageY = b.clientY + (d && d.scrollTop || e && e.scrollTop || 0) - (d && d.clientTop || e && e.clientTop || 0)), a.which || void 0 === f || (a.which = 1 & f ? 1 : 2 & f ? 3 : 4 & f ? 2 : 0), a
                }
            },
            fix: function(a) {
                if (a[_.expando]) return a;
                var b, c, d, e = a.type,
                    f = a,
                    g = this.fixHooks[e];
                for (g || (this.fixHooks[e] = g = Ba.test(e) ? this.mouseHooks : Aa.test(e) ? this.keyHooks : {}), d = g.props ? this.props.concat(g.props) : this.props, a = new _.Event(f), b = d.length; b--;) c = d[b], a[c] = f[c];
                return a.target || (a.target = Z), 3 === a.target.nodeType && (a.target = a.target.parentNode), g.filter ? g.filter(a, f) : a
            },
            special: {
                load: {
                    noBubble: !0
                },
                focus: {
                    trigger: function() {
                        return this !== l() && this.focus ? (this.focus(), !1) : void 0
                    },
                    delegateType: "focusin"
                },
                blur: {
                    trigger: function() {
                        return this === l() && this.blur ? (this.blur(), !1) : void 0
                    },
                    delegateType: "focusout"
                },
                click: {
                    trigger: function() {
                        return "checkbox" === this.type && this.click && _.nodeName(this, "input") ? (this.click(), !1) : void 0
                    },
                    _default: function(a) {
                        return _.nodeName(a.target, "a")
                    }
                },
                beforeunload: {
                    postDispatch: function(a) {
                        void 0 !== a.result && a.originalEvent && (a.originalEvent.returnValue = a.result)
                    }
                }
            },
            simulate: function(a, b, c, d) {
                var e = _.extend(new _.Event, c, {
                    type: a,
                    isSimulated: !0,
                    originalEvent: {}
                });
                d ? _.event.trigger(e, null, b) : _.event.dispatch.call(b, e), e.isDefaultPrevented() && c.preventDefault()
            }
        }, _.removeEvent = function(a, b, c) {
            a.removeEventListener && a.removeEventListener(b, c, !1)
        }, _.Event = function(a, b) {
            return this instanceof _.Event ? (a && a.type ? (this.originalEvent = a, this.type = a.type, this.isDefaultPrevented = a.defaultPrevented || void 0 === a.defaultPrevented && a.returnValue === !1 ? j : k) : this.type = a, b && _.extend(this, b), this.timeStamp = a && a.timeStamp || _.now(), void(this[_.expando] = !0)) : new _.Event(a, b)
        }, _.Event.prototype = {
            isDefaultPrevented: k,
            isPropagationStopped: k,
            isImmediatePropagationStopped: k,
            preventDefault: function() {
                var a = this.originalEvent;
                this.isDefaultPrevented = j, a && a.preventDefault && a.preventDefault()
            },
            stopPropagation: function() {
                var a = this.originalEvent;
                this.isPropagationStopped = j, a && a.stopPropagation && a.stopPropagation()
            },
            stopImmediatePropagation: function() {
                var a = this.originalEvent;
                this.isImmediatePropagationStopped = j, a && a.stopImmediatePropagation && a.stopImmediatePropagation(), this.stopPropagation()
            }
        }, _.each({
            mouseenter: "mouseover",
            mouseleave: "mouseout",
            pointerenter: "pointerover",
            pointerleave: "pointerout"
        }, function(a, b) {
            _.event.special[a] = {
                delegateType: b,
                bindType: b,
                handle: function(a) {
                    var c, d = this,
                        e = a.relatedTarget,
                        f = a.handleObj;
                    return (!e || e !== d && !_.contains(d, e)) && (a.type = f.origType, c = f.handler.apply(this, arguments), a.type = b), c
                }
            }
        }), Y.focusinBubbles || _.each({
            focus: "focusin",
            blur: "focusout"
        }, function(a, b) {
            var c = function(a) {
                _.event.simulate(b, a.target, _.event.fix(a), !0)
            };
            _.event.special[b] = {
                setup: function() {
                    var d = this.ownerDocument || this,
                        e = ra.access(d, b);
                    e || d.addEventListener(a, c, !0), ra.access(d, b, (e || 0) + 1)
                },
                teardown: function() {
                    var d = this.ownerDocument || this,
                        e = ra.access(d, b) - 1;
                    e ? ra.access(d, b, e) : (d.removeEventListener(a, c, !0), ra.remove(d, b))
                }
            }
        }), _.fn.extend({
            on: function(a, b, c, d, e) {
                var f, g;
                if ("object" == typeof a) {
                    "string" != typeof b && (c = c || b, b = void 0);
                    for (g in a) this.on(g, b, c, a[g], e);
                    return this
                }
                if (null == c && null == d ? (d = b, c = b = void 0) : null == d && ("string" == typeof b ? (d = c, c = void 0) : (d = c, c = b, b = void 0)), d === !1) d = k;
                else if (!d) return this;
                return 1 === e && (f = d, d = function(a) {
                    return _().off(a), f.apply(this, arguments)
                }, d.guid = f.guid || (f.guid = _.guid++)), this.each(function() {
                    _.event.add(this, a, d, c, b)
                })
            },
            one: function(a, b, c, d) {
                return this.on(a, b, c, d, 1)
            },
            off: function(a, b, c) {
                var d, e;
                if (a && a.preventDefault && a.handleObj) return d = a.handleObj, _(a.delegateTarget).off(d.namespace ? d.origType + "." + d.namespace : d.origType, d.selector, d.handler), this;
                if ("object" == typeof a) {
                    for (e in a) this.off(e, b, a[e]);
                    return this
                }
                return (b === !1 || "function" == typeof b) && (c = b, b = void 0), c === !1 && (c = k), this.each(function() {
                    _.event.remove(this, a, c, b)
                })
            },
            trigger: function(a, b) {
                return this.each(function() {
                    _.event.trigger(a, b, this)
                })
            },
            triggerHandler: function(a, b) {
                var c = this[0];
                return c ? _.event.trigger(a, b, c, !0) : void 0
            }
        });
        var Ea = /<(?!area|br|col|embed|hr|img|input|link|meta|param)(([\w:]+)[^>]*)\/>/gi,
            Fa = /<([\w:]+)/,
            Ga = /<|&#?\w+;/,
            Ha = /<(?:script|style|link)/i,
            Ia = /checked\s*(?:[^=]|=\s*.checked.)/i,
            Ja = /^$|\/(?:java|ecma)script/i,
            Ka = /^true\/(.*)/,
            La = /^\s*<!(?:\[CDATA\[|--)|(?:\]\]|--)>\s*$/g,
            Ma = {
                option: [1, "<select multiple='multiple'>", "</select>"],
                thead: [1, "<table>", "</table>"],
                col: [2, "<table><colgroup>", "</colgroup></table>"],
                tr: [2, "<table><tbody>", "</tbody></table>"],
                td: [3, "<table><tbody><tr>", "</tr></tbody></table>"],
                _default: [0, "", ""]
            };
        Ma.optgroup = Ma.option, Ma.tbody = Ma.tfoot = Ma.colgroup = Ma.caption = Ma.thead, Ma.th = Ma.td, _.extend({
            clone: function(a, b, c) {
                var d, e, f, g, h = a.cloneNode(!0),
                    i = _.contains(a.ownerDocument, a);
                if (!(Y.noCloneChecked || 1 !== a.nodeType && 11 !== a.nodeType || _.isXMLDoc(a)))
                    for (g = r(h), f = r(a), d = 0, e = f.length; e > d; d++) s(f[d], g[d]);
                if (b)
                    if (c)
                        for (f = f || r(a), g = g || r(h), d = 0, e = f.length; e > d; d++) q(f[d], g[d]);
                    else q(a, h);
                return g = r(h, "script"), g.length > 0 && p(g, !i && r(a, "script")), h
            },
            buildFragment: function(a, b, c, d) {
                for (var e, f, g, h, i, j, k = b.createDocumentFragment(), l = [], m = 0, n = a.length; n > m; m++)
                    if (e = a[m], e || 0 === e)
                        if ("object" === _.type(e)) _.merge(l, e.nodeType ? [e] : e);
                        else if (Ga.test(e)) {
                    for (f = f || k.appendChild(b.createElement("div")), g = (Fa.exec(e) || ["", ""])[1].toLowerCase(), h = Ma[g] || Ma._default, f.innerHTML = h[1] + e.replace(Ea, "<$1></$2>") + h[2], j = h[0]; j--;) f = f.lastChild;
                    _.merge(l, f.childNodes), f = k.firstChild, f.textContent = ""
                } else l.push(b.createTextNode(e));
                for (k.textContent = "", m = 0; e = l[m++];)
                    if ((!d || -1 === _.inArray(e, d)) && (i = _.contains(e.ownerDocument, e), f = r(k.appendChild(e), "script"), i && p(f), c))
                        for (j = 0; e = f[j++];) Ja.test(e.type || "") && c.push(e);
                return k
            },
            cleanData: function(a) {
                for (var b, c, d, e, f = _.event.special, g = 0; void 0 !== (c = a[g]); g++) {
                    if (_.acceptData(c) && (e = c[ra.expando], e && (b = ra.cache[e]))) {
                        if (b.events)
                            for (d in b.events) f[d] ? _.event.remove(c, d) : _.removeEvent(c, d, b.handle);
                        ra.cache[e] && delete ra.cache[e]
                    }
                    delete sa.cache[c[sa.expando]]
                }
            }
        }), _.fn.extend({
            text: function(a) {
                return qa(this, function(a) {
                    return void 0 === a ? _.text(this) : this.empty().each(function() {
                        (1 === this.nodeType || 11 === this.nodeType || 9 === this.nodeType) && (this.textContent = a)
                    })
                }, null, a, arguments.length)
            },
            append: function() {
                return this.domManip(arguments, function(a) {
                    if (1 === this.nodeType || 11 === this.nodeType || 9 === this.nodeType) {
                        var b = m(this, a);
                        b.appendChild(a)
                    }
                })
            },
            prepend: function() {
                return this.domManip(arguments, function(a) {
                    if (1 === this.nodeType || 11 === this.nodeType || 9 === this.nodeType) {
                        var b = m(this, a);
                        b.insertBefore(a, b.firstChild)
                    }
                })
            },
            before: function() {
                return this.domManip(arguments, function(a) {
                    this.parentNode && this.parentNode.insertBefore(a, this)
                })
            },
            after: function() {
                return this.domManip(arguments, function(a) {
                    this.parentNode && this.parentNode.insertBefore(a, this.nextSibling)
                })
            },
            remove: function(a, b) {
                for (var c, d = a ? _.filter(a, this) : this, e = 0; null != (c = d[e]); e++) b || 1 !== c.nodeType || _.cleanData(r(c)), c.parentNode && (b && _.contains(c.ownerDocument, c) && p(r(c, "script")), c.parentNode.removeChild(c));
                return this
            },
            empty: function() {
                for (var a, b = 0; null != (a = this[b]); b++) 1 === a.nodeType && (_.cleanData(r(a, !1)), a.textContent = "");
                return this
            },
            clone: function(a, b) {
                return a = null == a ? !1 : a, b = null == b ? a : b, this.map(function() {
                    return _.clone(this, a, b)
                })
            },
            html: function(a) {
                return qa(this, function(a) {
                    var b = this[0] || {},
                        c = 0,
                        d = this.length;
                    if (void 0 === a && 1 === b.nodeType) return b.innerHTML;
                    if ("string" == typeof a && !Ha.test(a) && !Ma[(Fa.exec(a) || ["", ""])[1].toLowerCase()]) {
                        a = a.replace(Ea, "<$1></$2>");
                        try {
                            for (; d > c; c++) b = this[c] || {}, 1 === b.nodeType && (_.cleanData(r(b, !1)), b.innerHTML = a);
                            b = 0
                        } catch (e) {}
                    }
                    b && this.empty().append(a)
                }, null, a, arguments.length)
            },
            replaceWith: function() {
                var a = arguments[0];
                return this.domManip(arguments, function(b) {
                    a = this.parentNode, _.cleanData(r(this)), a && a.replaceChild(b, this)
                }), a && (a.length || a.nodeType) ? this : this.remove()
            },
            detach: function(a) {
                return this.remove(a, !0)
            },
            domManip: function(a, b) {
                a = S.apply([], a);
                var c, d, e, f, g, h, i = 0,
                    j = this.length,
                    k = this,
                    l = j - 1,
                    m = a[0],
                    p = _.isFunction(m);
                if (p || j > 1 && "string" == typeof m && !Y.checkClone && Ia.test(m)) return this.each(function(c) {
                    var d = k.eq(c);
                    p && (a[0] = m.call(this, c, d.html())), d.domManip(a, b)
                });
                if (j && (c = _.buildFragment(a, this[0].ownerDocument, !1, this), d = c.firstChild, 1 === c.childNodes.length && (c = d), d)) {
                    for (e = _.map(r(c, "script"), n), f = e.length; j > i; i++) g = c, i !== l && (g = _.clone(g, !0, !0), f && _.merge(e, r(g, "script"))), b.call(this[i], g, i);
                    if (f)
                        for (h = e[e.length - 1].ownerDocument, _.map(e, o), i = 0; f > i; i++) g = e[i], Ja.test(g.type || "") && !ra.access(g, "globalEval") && _.contains(h, g) && (g.src ? _._evalUrl && _._evalUrl(g.src) : _.globalEval(g.textContent.replace(La, "")))
                }
                return this
            }
        }), _.each({
            appendTo: "append",
            prependTo: "prepend",
            insertBefore: "before",
            insertAfter: "after",
            replaceAll: "replaceWith"
        }, function(a, b) {
            _.fn[a] = function(a) {
                for (var c, d = [], e = _(a), f = e.length - 1, g = 0; f >= g; g++) c = g === f ? this : this.clone(!0), _(e[g])[b](c), T.apply(d, c.get());
                return this.pushStack(d)
            }
        });
        var Na, Oa = {},
            Pa = /^margin/,
            Qa = new RegExp("^(" + va + ")(?!px)[a-z%]+$", "i"),
            Ra = function(b) {
                return b.ownerDocument.defaultView.opener ? b.ownerDocument.defaultView.getComputedStyle(b, null) : a.getComputedStyle(b, null)
            };
        ! function() {
            function b() {
                g.style.cssText = "-webkit-box-sizing:border-box;-moz-box-sizing:border-box;box-sizing:border-box;display:block;margin-top:1%;top:1%;border:1px;padding:1px;width:4px;position:absolute", g.innerHTML = "", e.appendChild(f);
                var b = a.getComputedStyle(g, null);
                c = "1%" !== b.top, d = "4px" === b.width, e.removeChild(f)
            }
            var c, d, e = Z.documentElement,
                f = Z.createElement("div"),
                g = Z.createElement("div");
            g.style && (g.style.backgroundClip = "content-box", g.cloneNode(!0).style.backgroundClip = "", Y.clearCloneStyle = "content-box" === g.style.backgroundClip, f.style.cssText = "border:0;width:0;height:0;top:0;left:-9999px;margin-top:1px;position:absolute", f.appendChild(g), a.getComputedStyle && _.extend(Y, {
                pixelPosition: function() {
                    return b(), c
                },
                boxSizingReliable: function() {
                    return null == d && b(), d
                },
                reliableMarginRight: function() {
                    var b, c = g.appendChild(Z.createElement("div"));
                    return c.style.cssText = g.style.cssText = "-webkit-box-sizing:content-box;-moz-box-sizing:content-box;box-sizing:content-box;display:block;margin:0;border:0;padding:0", c.style.marginRight = c.style.width = "0", g.style.width = "1px", e.appendChild(f), b = !parseFloat(a.getComputedStyle(c, null).marginRight), e.removeChild(f), g.removeChild(c), b
                }
            }))
        }(), _.swap = function(a, b, c, d) {
            var e, f, g = {};
            for (f in b) g[f] = a.style[f], a.style[f] = b[f];
            e = c.apply(a, d || []);
            for (f in b) a.style[f] = g[f];
            return e
        };
        var Sa = /^(none|table(?!-c[ea]).+)/,
            Ta = new RegExp("^(" + va + ")(.*)$", "i"),
            Ua = new RegExp("^([+-])=(" + va + ")", "i"),
            Va = {
                position: "absolute",
                visibility: "hidden",
                display: "block"
            },
            Wa = {
                letterSpacing: "0",
                fontWeight: "400"
            },
            Xa = ["Webkit", "O", "Moz", "ms"];
        _.extend({
            cssHooks: {
                opacity: {
                    get: function(a, b) {
                        if (b) {
                            var c = v(a, "opacity");
                            return "" === c ? "1" : c
                        }
                    }
                }
            },
            cssNumber: {
                columnCount: !0,
                fillOpacity: !0,
                flexGrow: !0,
                flexShrink: !0,
                fontWeight: !0,
                lineHeight: !0,
                opacity: !0,
                order: !0,
                orphans: !0,
                widows: !0,
                zIndex: !0,
                zoom: !0
            },
            cssProps: {
                "float": "cssFloat"
            },
            style: function(a, b, c, d) {
                if (a && 3 !== a.nodeType && 8 !== a.nodeType && a.style) {
                    var e, f, g, h = _.camelCase(b),
                        i = a.style;
                    return b = _.cssProps[h] || (_.cssProps[h] = x(i, h)), g = _.cssHooks[b] || _.cssHooks[h], void 0 === c ? g && "get" in g && void 0 !== (e = g.get(a, !1, d)) ? e : i[b] : (f = typeof c, "string" === f && (e = Ua.exec(c)) && (c = (e[1] + 1) * e[2] + parseFloat(_.css(a, b)), f = "number"), null != c && c === c && ("number" !== f || _.cssNumber[h] || (c += "px"), Y.clearCloneStyle || "" !== c || 0 !== b.indexOf("background") || (i[b] = "inherit"), g && "set" in g && void 0 === (c = g.set(a, c, d)) || (i[b] = c)), void 0)
                }
            },
            css: function(a, b, c, d) {
                var e, f, g, h = _.camelCase(b);
                return b = _.cssProps[h] || (_.cssProps[h] = x(a.style, h)), g = _.cssHooks[b] || _.cssHooks[h], g && "get" in g && (e = g.get(a, !0, c)), void 0 === e && (e = v(a, b, d)), "normal" === e && b in Wa && (e = Wa[b]), "" === c || c ? (f = parseFloat(e), c === !0 || _.isNumeric(f) ? f || 0 : e) : e
            }
        }), _.each(["height", "width"], function(a, b) {
            _.cssHooks[b] = {
                get: function(a, c, d) {
                    return c ? Sa.test(_.css(a, "display")) && 0 === a.offsetWidth ? _.swap(a, Va, function() {
                        return A(a, b, d)
                    }) : A(a, b, d) : void 0
                },
                set: function(a, c, d) {
                    var e = d && Ra(a);
                    return y(a, c, d ? z(a, b, d, "border-box" === _.css(a, "boxSizing", !1, e), e) : 0)
                }
            }
        }), _.cssHooks.marginRight = w(Y.reliableMarginRight, function(a, b) {
            return b ? _.swap(a, {
                display: "inline-block"
            }, v, [a, "marginRight"]) : void 0
        }), _.each({
            margin: "",
            padding: "",
            border: "Width"
        }, function(a, b) {
            _.cssHooks[a + b] = {
                expand: function(c) {
                    for (var d = 0, e = {}, f = "string" == typeof c ? c.split(" ") : [c]; 4 > d; d++) e[a + wa[d] + b] = f[d] || f[d - 2] || f[0];
                    return e
                }
            }, Pa.test(a) || (_.cssHooks[a + b].set = y)
        }), _.fn.extend({
            css: function(a, b) {
                return qa(this, function(a, b, c) {
                    var d, e, f = {},
                        g = 0;
                    if (_.isArray(b)) {
                        for (d = Ra(a), e = b.length; e > g; g++) f[b[g]] = _.css(a, b[g], !1, d);
                        return f
                    }
                    return void 0 !== c ? _.style(a, b, c) : _.css(a, b)
                }, a, b, arguments.length > 1)
            },
            show: function() {
                return B(this, !0)
            },
            hide: function() {
                return B(this)
            },
            toggle: function(a) {
                return "boolean" == typeof a ? a ? this.show() : this.hide() : this.each(function() {
                    xa(this) ? _(this).show() : _(this).hide()
                })
            }
        }), _.Tween = C, C.prototype = {
            constructor: C,
            init: function(a, b, c, d, e, f) {
                this.elem = a, this.prop = c, this.easing = e || "swing", this.options = b, this.start = this.now = this.cur(), this.end = d, this.unit = f || (_.cssNumber[c] ? "" : "px")
            },
            cur: function() {
                var a = C.propHooks[this.prop];
                return a && a.get ? a.get(this) : C.propHooks._default.get(this)
            },
            run: function(a) {
                var b, c = C.propHooks[this.prop];
                return this.options.duration ? this.pos = b = _.easing[this.easing](a, this.options.duration * a, 0, 1, this.options.duration) : this.pos = b = a, this.now = (this.end - this.start) * b + this.start, this.options.step && this.options.step.call(this.elem, this.now, this), c && c.set ? c.set(this) : C.propHooks._default.set(this), this
            }
        }, C.prototype.init.prototype = C.prototype, C.propHooks = {
            _default: {
                get: function(a) {
                    var b;
                    return null == a.elem[a.prop] || a.elem.style && null != a.elem.style[a.prop] ? (b = _.css(a.elem, a.prop, ""), b && "auto" !== b ? b : 0) : a.elem[a.prop]
                },
                set: function(a) {
                    _.fx.step[a.prop] ? _.fx.step[a.prop](a) : a.elem.style && (null != a.elem.style[_.cssProps[a.prop]] || _.cssHooks[a.prop]) ? _.style(a.elem, a.prop, a.now + a.unit) : a.elem[a.prop] = a.now
                }
            }
        }, C.propHooks.scrollTop = C.propHooks.scrollLeft = {
            set: function(a) {
                a.elem.nodeType && a.elem.parentNode && (a.elem[a.prop] = a.now)
            }
        }, _.easing = {
            linear: function(a) {
                return a
            },
            swing: function(a) {
                return .5 - Math.cos(a * Math.PI) / 2
            }
        }, _.fx = C.prototype.init, _.fx.step = {};
        var Ya, Za, $a = /^(?:toggle|show|hide)$/,
            _a = new RegExp("^(?:([+-])=|)(" + va + ")([a-z%]*)$", "i"),
            ab = /queueHooks$/,
            bb = [G],
            cb = {
                "*": [function(a, b) {
                    var c = this.createTween(a, b),
                        d = c.cur(),
                        e = _a.exec(b),
                        f = e && e[3] || (_.cssNumber[a] ? "" : "px"),
                        g = (_.cssNumber[a] || "px" !== f && +d) && _a.exec(_.css(c.elem, a)),
                        h = 1,
                        i = 20;
                    if (g && g[3] !== f) {
                        f = f || g[3], e = e || [], g = +d || 1;
                        do h = h || ".5", g /= h, _.style(c.elem, a, g + f); while (h !== (h = c.cur() / d) && 1 !== h && --i)
                    }
                    return e && (g = c.start = +g || +d || 0, c.unit = f, c.end = e[1] ? g + (e[1] + 1) * e[2] : +e[2]), c
                }]
            };
        _.Animation = _.extend(I, {
                tweener: function(a, b) {
                    _.isFunction(a) ? (b = a, a = ["*"]) : a = a.split(" ");
                    for (var c, d = 0, e = a.length; e > d; d++) c = a[d], cb[c] = cb[c] || [], cb[c].unshift(b)
                },
                prefilter: function(a, b) {
                    b ? bb.unshift(a) : bb.push(a)
                }
            }), _.speed = function(a, b, c) {
                var d = a && "object" == typeof a ? _.extend({}, a) : {
                    complete: c || !c && b || _.isFunction(a) && a,
                    duration: a,
                    easing: c && b || b && !_.isFunction(b) && b
                };
                return d.duration = _.fx.off ? 0 : "number" == typeof d.duration ? d.duration : d.duration in _.fx.speeds ? _.fx.speeds[d.duration] : _.fx.speeds._default, (null == d.queue || d.queue === !0) && (d.queue = "fx"), d.old = d.complete, d.complete = function() {
                    _.isFunction(d.old) && d.old.call(this), d.queue && _.dequeue(this, d.queue)
                }, d
            }, _.fn.extend({
                fadeTo: function(a, b, c, d) {
                    return this.filter(xa).css("opacity", 0).show().end().animate({
                        opacity: b
                    }, a, c, d)
                },
                animate: function(a, b, c, d) {
                    var e = _.isEmptyObject(a),
                        f = _.speed(b, c, d),
                        g = function() {
                            var b = I(this, _.extend({}, a), f);
                            (e || ra.get(this, "finish")) && b.stop(!0)
                        };
                    return g.finish = g, e || f.queue === !1 ? this.each(g) : this.queue(f.queue, g)
                },
                stop: function(a, b, c) {
                    var d = function(a) {
                        var b = a.stop;
                        delete a.stop, b(c)
                    };
                    return "string" != typeof a && (c = b, b = a, a = void 0), b && a !== !1 && this.queue(a || "fx", []), this.each(function() {
                        var b = !0,
                            e = null != a && a + "queueHooks",
                            f = _.timers,
                            g = ra.get(this);
                        if (e) g[e] && g[e].stop && d(g[e]);
                        else
                            for (e in g) g[e] && g[e].stop && ab.test(e) && d(g[e]);
                        for (e = f.length; e--;) f[e].elem !== this || null != a && f[e].queue !== a || (f[e].anim.stop(c), b = !1, f.splice(e, 1));
                        (b || !c) && _.dequeue(this, a)
                    })
                },
                finish: function(a) {
                    return a !== !1 && (a = a || "fx"), this.each(function() {
                        var b, c = ra.get(this),
                            d = c[a + "queue"],
                            e = c[a + "queueHooks"],
                            f = _.timers,
                            g = d ? d.length : 0;
                        for (c.finish = !0, _.queue(this, a, []),
                            e && e.stop && e.stop.call(this, !0), b = f.length; b--;) f[b].elem === this && f[b].queue === a && (f[b].anim.stop(!0), f.splice(b, 1));
                        for (b = 0; g > b; b++) d[b] && d[b].finish && d[b].finish.call(this);
                        delete c.finish
                    })
                }
            }), _.each(["toggle", "show", "hide"], function(a, b) {
                var c = _.fn[b];
                _.fn[b] = function(a, d, e) {
                    return null == a || "boolean" == typeof a ? c.apply(this, arguments) : this.animate(E(b, !0), a, d, e)
                }
            }), _.each({
                slideDown: E("show"),
                slideUp: E("hide"),
                slideToggle: E("toggle"),
                fadeIn: {
                    opacity: "show"
                },
                fadeOut: {
                    opacity: "hide"
                },
                fadeToggle: {
                    opacity: "toggle"
                }
            }, function(a, b) {
                _.fn[a] = function(a, c, d) {
                    return this.animate(b, a, c, d)
                }
            }), _.timers = [], _.fx.tick = function() {
                var a, b = 0,
                    c = _.timers;
                for (Ya = _.now(); b < c.length; b++) a = c[b], a() || c[b] !== a || c.splice(b--, 1);
                c.length || _.fx.stop(), Ya = void 0
            }, _.fx.timer = function(a) {
                _.timers.push(a), a() ? _.fx.start() : _.timers.pop()
            }, _.fx.interval = 13, _.fx.start = function() {
                Za || (Za = setInterval(_.fx.tick, _.fx.interval))
            }, _.fx.stop = function() {
                clearInterval(Za), Za = null
            }, _.fx.speeds = {
                slow: 600,
                fast: 200,
                _default: 400
            }, _.fn.delay = function(a, b) {
                return a = _.fx ? _.fx.speeds[a] || a : a, b = b || "fx", this.queue(b, function(b, c) {
                    var d = setTimeout(b, a);
                    c.stop = function() {
                        clearTimeout(d)
                    }
                })
            },
            function() {
                var a = Z.createElement("input"),
                    b = Z.createElement("select"),
                    c = b.appendChild(Z.createElement("option"));
                a.type = "checkbox", Y.checkOn = "" !== a.value, Y.optSelected = c.selected, b.disabled = !0, Y.optDisabled = !c.disabled, a = Z.createElement("input"), a.value = "t", a.type = "radio", Y.radioValue = "t" === a.value
            }();
        var db, eb, fb = _.expr.attrHandle;
        _.fn.extend({
            attr: function(a, b) {
                return qa(this, _.attr, a, b, arguments.length > 1)
            },
            removeAttr: function(a) {
                return this.each(function() {
                    _.removeAttr(this, a)
                })
            }
        }), _.extend({
            attr: function(a, b, c) {
                var d, e, f = a.nodeType;
                if (a && 3 !== f && 8 !== f && 2 !== f) return typeof a.getAttribute === za ? _.prop(a, b, c) : (1 === f && _.isXMLDoc(a) || (b = b.toLowerCase(), d = _.attrHooks[b] || (_.expr.match.bool.test(b) ? eb : db)), void 0 === c ? d && "get" in d && null !== (e = d.get(a, b)) ? e : (e = _.find.attr(a, b), null == e ? void 0 : e) : null !== c ? d && "set" in d && void 0 !== (e = d.set(a, c, b)) ? e : (a.setAttribute(b, c + ""), c) : void _.removeAttr(a, b))
            },
            removeAttr: function(a, b) {
                var c, d, e = 0,
                    f = b && b.match(na);
                if (f && 1 === a.nodeType)
                    for (; c = f[e++];) d = _.propFix[c] || c, _.expr.match.bool.test(c) && (a[d] = !1), a.removeAttribute(c)
            },
            attrHooks: {
                type: {
                    set: function(a, b) {
                        if (!Y.radioValue && "radio" === b && _.nodeName(a, "input")) {
                            var c = a.value;
                            return a.setAttribute("type", b), c && (a.value = c), b
                        }
                    }
                }
            }
        }), eb = {
            set: function(a, b, c) {
                return b === !1 ? _.removeAttr(a, c) : a.setAttribute(c, c), c
            }
        }, _.each(_.expr.match.bool.source.match(/\w+/g), function(a, b) {
            var c = fb[b] || _.find.attr;
            fb[b] = function(a, b, d) {
                var e, f;
                return d || (f = fb[b], fb[b] = e, e = null != c(a, b, d) ? b.toLowerCase() : null, fb[b] = f), e
            }
        });
        var gb = /^(?:input|select|textarea|button)$/i;
        _.fn.extend({
            prop: function(a, b) {
                return qa(this, _.prop, a, b, arguments.length > 1)
            },
            removeProp: function(a) {
                return this.each(function() {
                    delete this[_.propFix[a] || a]
                })
            }
        }), _.extend({
            propFix: {
                "for": "htmlFor",
                "class": "className"
            },
            prop: function(a, b, c) {
                var d, e, f, g = a.nodeType;
                if (a && 3 !== g && 8 !== g && 2 !== g) return f = 1 !== g || !_.isXMLDoc(a), f && (b = _.propFix[b] || b, e = _.propHooks[b]), void 0 !== c ? e && "set" in e && void 0 !== (d = e.set(a, c, b)) ? d : a[b] = c : e && "get" in e && null !== (d = e.get(a, b)) ? d : a[b]
            },
            propHooks: {
                tabIndex: {
                    get: function(a) {
                        return a.hasAttribute("tabindex") || gb.test(a.nodeName) || a.href ? a.tabIndex : -1
                    }
                }
            }
        }), Y.optSelected || (_.propHooks.selected = {
            get: function(a) {
                var b = a.parentNode;
                return b && b.parentNode && b.parentNode.selectedIndex, null
            }
        }), _.each(["tabIndex", "readOnly", "maxLength", "cellSpacing", "cellPadding", "rowSpan", "colSpan", "useMap", "frameBorder", "contentEditable"], function() {
            _.propFix[this.toLowerCase()] = this
        });
        var hb = /[\t\r\n\f]/g;
        _.fn.extend({
            addClass: function(a) {
                var b, c, d, e, f, g, h = "string" == typeof a && a,
                    i = 0,
                    j = this.length;
                if (_.isFunction(a)) return this.each(function(b) {
                    _(this).addClass(a.call(this, b, this.className))
                });
                if (h)
                    for (b = (a || "").match(na) || []; j > i; i++)
                        if (c = this[i], d = 1 === c.nodeType && (c.className ? (" " + c.className + " ").replace(hb, " ") : " ")) {
                            for (f = 0; e = b[f++];) d.indexOf(" " + e + " ") < 0 && (d += e + " ");
                            g = _.trim(d), c.className !== g && (c.className = g)
                        }
                return this
            },
            removeClass: function(a) {
                var b, c, d, e, f, g, h = 0 === arguments.length || "string" == typeof a && a,
                    i = 0,
                    j = this.length;
                if (_.isFunction(a)) return this.each(function(b) {
                    _(this).removeClass(a.call(this, b, this.className))
                });
                if (h)
                    for (b = (a || "").match(na) || []; j > i; i++)
                        if (c = this[i], d = 1 === c.nodeType && (c.className ? (" " + c.className + " ").replace(hb, " ") : "")) {
                            for (f = 0; e = b[f++];)
                                for (; d.indexOf(" " + e + " ") >= 0;) d = d.replace(" " + e + " ", " ");
                            g = a ? _.trim(d) : "", c.className !== g && (c.className = g)
                        }
                return this
            },
            toggleClass: function(a, b) {
                var c = typeof a;
                return "boolean" == typeof b && "string" === c ? b ? this.addClass(a) : this.removeClass(a) : _.isFunction(a) ? this.each(function(c) {
                    _(this).toggleClass(a.call(this, c, this.className, b), b)
                }) : this.each(function() {
                    if ("string" === c)
                        for (var b, d = 0, e = _(this), f = a.match(na) || []; b = f[d++];) e.hasClass(b) ? e.removeClass(b) : e.addClass(b);
                    else(c === za || "boolean" === c) && (this.className && ra.set(this, "__className__", this.className), this.className = this.className || a === !1 ? "" : ra.get(this, "__className__") || "")
                })
            },
            hasClass: function(a) {
                for (var b = " " + a + " ", c = 0, d = this.length; d > c; c++)
                    if (1 === this[c].nodeType && (" " + this[c].className + " ").replace(hb, " ").indexOf(b) >= 0) return !0;
                return !1
            }
        });
        var ib = /\r/g;
        _.fn.extend({
            val: function(a) {
                var b, c, d, e = this[0]; {
                    if (arguments.length) return d = _.isFunction(a), this.each(function(c) {
                        var e;
                        1 === this.nodeType && (e = d ? a.call(this, c, _(this).val()) : a, null == e ? e = "" : "number" == typeof e ? e += "" : _.isArray(e) && (e = _.map(e, function(a) {
                            return null == a ? "" : a + ""
                        })), b = _.valHooks[this.type] || _.valHooks[this.nodeName.toLowerCase()], b && "set" in b && void 0 !== b.set(this, e, "value") || (this.value = e))
                    });
                    if (e) return b = _.valHooks[e.type] || _.valHooks[e.nodeName.toLowerCase()], b && "get" in b && void 0 !== (c = b.get(e, "value")) ? c : (c = e.value, "string" == typeof c ? c.replace(ib, "") : null == c ? "" : c)
                }
            }
        }), _.extend({
            valHooks: {
                option: {
                    get: function(a) {
                        var b = _.find.attr(a, "value");
                        return null != b ? b : _.trim(_.text(a))
                    }
                },
                select: {
                    get: function(a) {
                        for (var b, c, d = a.options, e = a.selectedIndex, f = "select-one" === a.type || 0 > e, g = f ? null : [], h = f ? e + 1 : d.length, i = 0 > e ? h : f ? e : 0; h > i; i++)
                            if (c = d[i], (c.selected || i === e) && (Y.optDisabled ? !c.disabled : null === c.getAttribute("disabled")) && (!c.parentNode.disabled || !_.nodeName(c.parentNode, "optgroup"))) {
                                if (b = _(c).val(), f) return b;
                                g.push(b)
                            }
                        return g
                    },
                    set: function(a, b) {
                        for (var c, d, e = a.options, f = _.makeArray(b), g = e.length; g--;) d = e[g], (d.selected = _.inArray(d.value, f) >= 0) && (c = !0);
                        return c || (a.selectedIndex = -1), f
                    }
                }
            }
        }), _.each(["radio", "checkbox"], function() {
            _.valHooks[this] = {
                set: function(a, b) {
                    return _.isArray(b) ? a.checked = _.inArray(_(a).val(), b) >= 0 : void 0
                }
            }, Y.checkOn || (_.valHooks[this].get = function(a) {
                return null === a.getAttribute("value") ? "on" : a.value
            })
        }), _.each("blur focus focusin focusout load resize scroll unload click dblclick mousedown mouseup mousemove mouseover mouseout mouseenter mouseleave change select submit keydown keypress keyup error contextmenu".split(" "), function(a, b) {
            _.fn[b] = function(a, c) {
                return arguments.length > 0 ? this.on(b, null, a, c) : this.trigger(b)
            }
        }), _.fn.extend({
            hover: function(a, b) {
                return this.mouseenter(a).mouseleave(b || a)
            },
            bind: function(a, b, c) {
                return this.on(a, null, b, c)
            },
            unbind: function(a, b) {
                return this.off(a, null, b)
            },
            delegate: function(a, b, c, d) {
                return this.on(b, a, c, d)
            },
            undelegate: function(a, b, c) {
                return 1 === arguments.length ? this.off(a, "**") : this.off(b, a || "**", c)
            }
        });
        var jb = _.now(),
            kb = /\?/;
        _.parseJSON = function(a) {
            return JSON.parse(a + "")
        }, _.parseXML = function(a) {
            var b, c;
            if (!a || "string" != typeof a) return null;
            try {
                c = new DOMParser, b = c.parseFromString(a, "text/xml")
            } catch (d) {
                b = void 0
            }
            return (!b || b.getElementsByTagName("parsererror").length) && _.error("Invalid XML: " + a), b
        };
        var lb = /#.*$/,
            mb = /([?&])_=[^&]*/,
            nb = /^(.*?):[ \t]*([^\r\n]*)$/gm,
            ob = /^(?:about|app|app-storage|.+-extension|file|res|widget):$/,
            pb = /^(?:GET|HEAD)$/,
            qb = /^\/\//,
            rb = /^([\w.+-]+:)(?:\/\/(?:[^\/?#]*@|)([^\/?#:]*)(?::(\d+)|)|)/,
            sb = {},
            tb = {},
            ub = "*/".concat("*"),
            vb = a.location.href,
            wb = rb.exec(vb.toLowerCase()) || [];
        _.extend({
            active: 0,
            lastModified: {},
            etag: {},
            ajaxSettings: {
                url: vb,
                type: "GET",
                isLocal: ob.test(wb[1]),
                global: !0,
                processData: !0,
                async: !0,
                contentType: "application/x-www-form-urlencoded; charset=UTF-8",
                accepts: {
                    "*": ub,
                    text: "text/plain",
                    html: "text/html",
                    xml: "application/xml, text/xml",
                    json: "application/json, text/javascript"
                },
                contents: {
                    xml: /xml/,
                    html: /html/,
                    json: /json/
                },
                responseFields: {
                    xml: "responseXML",
                    text: "responseText",
                    json: "responseJSON"
                },
                converters: {
                    "* text": String,
                    "text html": !0,
                    "text json": _.parseJSON,
                    "text xml": _.parseXML
                },
                flatOptions: {
                    url: !0,
                    context: !0
                }
            },
            ajaxSetup: function(a, b) {
                return b ? L(L(a, _.ajaxSettings), b) : L(_.ajaxSettings, a)
            },
            ajaxPrefilter: J(sb),
            ajaxTransport: J(tb),
            ajax: function(a, b) {
                function c(a, b, c, g) {
                    var i, k, r, s, u, w = b;
                    2 !== t && (t = 2, h && clearTimeout(h), d = void 0, f = g || "", v.readyState = a > 0 ? 4 : 0, i = a >= 200 && 300 > a || 304 === a, c && (s = M(l, v, c)), s = N(l, s, v, i), i ? (l.ifModified && (u = v.getResponseHeader("Last-Modified"), u && (_.lastModified[e] = u), u = v.getResponseHeader("etag"), u && (_.etag[e] = u)), 204 === a || "HEAD" === l.type ? w = "nocontent" : 304 === a ? w = "notmodified" : (w = s.state, k = s.data, r = s.error, i = !r)) : (r = w, (a || !w) && (w = "error", 0 > a && (a = 0))), v.status = a, v.statusText = (b || w) + "", i ? o.resolveWith(m, [k, w, v]) : o.rejectWith(m, [v, w, r]), v.statusCode(q), q = void 0, j && n.trigger(i ? "ajaxSuccess" : "ajaxError", [v, l, i ? k : r]), p.fireWith(m, [v, w]), j && (n.trigger("ajaxComplete", [v, l]), --_.active || _.event.trigger("ajaxStop")))
                }
                "object" == typeof a && (b = a, a = void 0), b = b || {};
                var d, e, f, g, h, i, j, k, l = _.ajaxSetup({}, b),
                    m = l.context || l,
                    n = l.context && (m.nodeType || m.jquery) ? _(m) : _.event,
                    o = _.Deferred(),
                    p = _.Callbacks("once memory"),
                    q = l.statusCode || {},
                    r = {},
                    s = {},
                    t = 0,
                    u = "canceled",
                    v = {
                        readyState: 0,
                        getResponseHeader: function(a) {
                            var b;
                            if (2 === t) {
                                if (!g)
                                    for (g = {}; b = nb.exec(f);) g[b[1].toLowerCase()] = b[2];
                                b = g[a.toLowerCase()]
                            }
                            return null == b ? null : b
                        },
                        getAllResponseHeaders: function() {
                            return 2 === t ? f : null
                        },
                        setRequestHeader: function(a, b) {
                            var c = a.toLowerCase();
                            return t || (a = s[c] = s[c] || a, r[a] = b), this
                        },
                        overrideMimeType: function(a) {
                            return t || (l.mimeType = a), this
                        },
                        statusCode: function(a) {
                            var b;
                            if (a)
                                if (2 > t)
                                    for (b in a) q[b] = [q[b], a[b]];
                                else v.always(a[v.status]);
                            return this
                        },
                        abort: function(a) {
                            var b = a || u;
                            return d && d.abort(b), c(0, b), this
                        }
                    };
                if (o.promise(v).complete = p.add, v.success = v.done, v.error = v.fail, l.url = ((a || l.url || vb) + "").replace(lb, "").replace(qb, wb[1] + "//"), l.type = b.method || b.type || l.method || l.type, l.dataTypes = _.trim(l.dataType || "*").toLowerCase().match(na) || [""], null == l.crossDomain && (i = rb.exec(l.url.toLowerCase()), l.crossDomain = !(!i || i[1] === wb[1] && i[2] === wb[2] && (i[3] || ("http:" === i[1] ? "80" : "443")) === (wb[3] || ("http:" === wb[1] ? "80" : "443")))), l.data && l.processData && "string" != typeof l.data && (l.data = _.param(l.data, l.traditional)), K(sb, l, b, v), 2 === t) return v;
                j = _.event && l.global, j && 0 === _.active++ && _.event.trigger("ajaxStart"), l.type = l.type.toUpperCase(), l.hasContent = !pb.test(l.type), e = l.url, l.hasContent || (l.data && (e = l.url += (kb.test(e) ? "&" : "?") + l.data, delete l.data), l.cache === !1 && (l.url = mb.test(e) ? e.replace(mb, "$1_=" + jb++) : e + (kb.test(e) ? "&" : "?") + "_=" + jb++)), l.ifModified && (_.lastModified[e] && v.setRequestHeader("If-Modified-Since", _.lastModified[e]), _.etag[e] && v.setRequestHeader("If-None-Match", _.etag[e])), (l.data && l.hasContent && l.contentType !== !1 || b.contentType) && v.setRequestHeader("Content-Type", l.contentType), v.setRequestHeader("Accept", l.dataTypes[0] && l.accepts[l.dataTypes[0]] ? l.accepts[l.dataTypes[0]] + ("*" !== l.dataTypes[0] ? ", " + ub + "; q=0.01" : "") : l.accepts["*"]);
                for (k in l.headers) v.setRequestHeader(k, l.headers[k]);
                if (l.beforeSend && (l.beforeSend.call(m, v, l) === !1 || 2 === t)) return v.abort();
                u = "abort";
                for (k in {
                        success: 1,
                        error: 1,
                        complete: 1
                    }) v[k](l[k]);
                if (d = K(tb, l, b, v)) {
                    v.readyState = 1, j && n.trigger("ajaxSend", [v, l]), l.async && l.timeout > 0 && (h = setTimeout(function() {
                        v.abort("timeout")
                    }, l.timeout));
                    try {
                        t = 1, d.send(r, c)
                    } catch (w) {
                        if (!(2 > t)) throw w;
                        c(-1, w)
                    }
                } else c(-1, "No Transport");
                return v
            },
            getJSON: function(a, b, c) {
                return _.get(a, b, c, "json")
            },
            getScript: function(a, b) {
                return _.get(a, void 0, b, "script")
            }
        }), _.each(["get", "post"], function(a, b) {
            _[b] = function(a, c, d, e) {
                return _.isFunction(c) && (e = e || d, d = c, c = void 0), _.ajax({
                    url: a,
                    type: b,
                    dataType: e,
                    data: c,
                    success: d
                })
            }
        }), _._evalUrl = function(a) {
            return _.ajax({
                url: a,
                type: "GET",
                dataType: "script",
                async: !1,
                global: !1,
                "throws": !0
            })
        }, _.fn.extend({
            wrapAll: function(a) {
                var b;
                return _.isFunction(a) ? this.each(function(b) {
                    _(this).wrapAll(a.call(this, b))
                }) : (this[0] && (b = _(a, this[0].ownerDocument).eq(0).clone(!0), this[0].parentNode && b.insertBefore(this[0]), b.map(function() {
                    for (var a = this; a.firstElementChild;) a = a.firstElementChild;
                    return a
                }).append(this)), this)
            },
            wrapInner: function(a) {
                return _.isFunction(a) ? this.each(function(b) {
                    _(this).wrapInner(a.call(this, b))
                }) : this.each(function() {
                    var b = _(this),
                        c = b.contents();
                    c.length ? c.wrapAll(a) : b.append(a)
                })
            },
            wrap: function(a) {
                var b = _.isFunction(a);
                return this.each(function(c) {
                    _(this).wrapAll(b ? a.call(this, c) : a)
                })
            },
            unwrap: function() {
                return this.parent().each(function() {
                    _.nodeName(this, "body") || _(this).replaceWith(this.childNodes)
                }).end()
            }
        }), _.expr.filters.hidden = function(a) {
            return a.offsetWidth <= 0 && a.offsetHeight <= 0
        }, _.expr.filters.visible = function(a) {
            return !_.expr.filters.hidden(a)
        };
        var xb = /%20/g,
            yb = /\[\]$/,
            zb = /\r?\n/g,
            Ab = /^(?:submit|button|image|reset|file)$/i,
            Bb = /^(?:input|select|textarea|keygen)/i;
        _.param = function(a, b) {
            var c, d = [],
                e = function(a, b) {
                    b = _.isFunction(b) ? b() : null == b ? "" : b, d[d.length] = encodeURIComponent(a) + "=" + encodeURIComponent(b)
                };
            if (void 0 === b && (b = _.ajaxSettings && _.ajaxSettings.traditional), _.isArray(a) || a.jquery && !_.isPlainObject(a)) _.each(a, function() {
                e(this.name, this.value)
            });
            else
                for (c in a) O(c, a[c], b, e);
            return d.join("&").replace(xb, "+")
        }, _.fn.extend({
            serialize: function() {
                return _.param(this.serializeArray())
            },
            serializeArray: function() {
                return this.map(function() {
                    var a = _.prop(this, "elements");
                    return a ? _.makeArray(a) : this
                }).filter(function() {
                    var a = this.type;
                    return this.name && !_(this).is(":disabled") && Bb.test(this.nodeName) && !Ab.test(a) && (this.checked || !ya.test(a))
                }).map(function(a, b) {
                    var c = _(this).val();
                    return null == c ? null : _.isArray(c) ? _.map(c, function(a) {
                        return {
                            name: b.name,
                            value: a.replace(zb, "\r\n")
                        }
                    }) : {
                        name: b.name,
                        value: c.replace(zb, "\r\n")
                    }
                }).get()
            }
        }), _.ajaxSettings.xhr = function() {
            try {
                return new XMLHttpRequest
            } catch (a) {}
        };
        var Cb = 0,
            Db = {},
            Eb = {
                0: 200,
                1223: 204
            },
            Fb = _.ajaxSettings.xhr();
        a.attachEvent && a.attachEvent("onunload", function() {
            for (var a in Db) Db[a]()
        }), Y.cors = !!Fb && "withCredentials" in Fb, Y.ajax = Fb = !!Fb, _.ajaxTransport(function(a) {
            var b;
            return Y.cors || Fb && !a.crossDomain ? {
                send: function(c, d) {
                    var e, f = a.xhr(),
                        g = ++Cb;
                    if (f.open(a.type, a.url, a.async, a.username, a.password), a.xhrFields)
                        for (e in a.xhrFields) f[e] = a.xhrFields[e];
                    a.mimeType && f.overrideMimeType && f.overrideMimeType(a.mimeType), a.crossDomain || c["X-Requested-With"] || (c["X-Requested-With"] = "XMLHttpRequest");
                    for (e in c) f.setRequestHeader(e, c[e]);
                    b = function(a) {
                        return function() {
                            b && (delete Db[g], b = f.onload = f.onerror = null, "abort" === a ? f.abort() : "error" === a ? d(f.status, f.statusText) : d(Eb[f.status] || f.status, f.statusText, "string" == typeof f.responseText ? {
                                text: f.responseText
                            } : void 0, f.getAllResponseHeaders()))
                        }
                    }, f.onload = b(), f.onerror = b("error"), b = Db[g] = b("abort");
                    try {
                        f.send(a.hasContent && a.data || null)
                    } catch (h) {
                        if (b) throw h
                    }
                },
                abort: function() {
                    b && b()
                }
            } : void 0
        }), _.ajaxSetup({
            accepts: {
                script: "text/javascript, application/javascript, application/ecmascript, application/x-ecmascript"
            },
            contents: {
                script: /(?:java|ecma)script/
            },
            converters: {
                "text script": function(a) {
                    return _.globalEval(a), a
                }
            }
        }), _.ajaxPrefilter("script", function(a) {
            void 0 === a.cache && (a.cache = !1), a.crossDomain && (a.type = "GET")
        }), _.ajaxTransport("script", function(a) {
            if (a.crossDomain) {
                var b, c;
                return {
                    send: function(d, e) {
                        b = _("<script>").prop({
                            async: !0,
                            charset: a.scriptCharset,
                            src: a.url
                        }).on("load error", c = function(a) {
                            b.remove(), c = null, a && e("error" === a.type ? 404 : 200, a.type)
                        }), Z.head.appendChild(b[0])
                    },
                    abort: function() {
                        c && c()
                    }
                }
            }
        });
        var Gb = [],
            Hb = /(=)\?(?=&|$)|\?\?/;
        _.ajaxSetup({
            jsonp: "callback",
            jsonpCallback: function() {
                var a = Gb.pop() || _.expando + "_" + jb++;
                return this[a] = !0, a
            }
        }), _.ajaxPrefilter("json jsonp", function(b, c, d) {
            var e, f, g, h = b.jsonp !== !1 && (Hb.test(b.url) ? "url" : "string" == typeof b.data && !(b.contentType || "").indexOf("application/x-www-form-urlencoded") && Hb.test(b.data) && "data");
            return h || "jsonp" === b.dataTypes[0] ? (e = b.jsonpCallback = _.isFunction(b.jsonpCallback) ? b.jsonpCallback() : b.jsonpCallback, h ? b[h] = b[h].replace(Hb, "$1" + e) : b.jsonp !== !1 && (b.url += (kb.test(b.url) ? "&" : "?") + b.jsonp + "=" + e), b.converters["script json"] = function() {
                return g || _.error(e + " was not called"), g[0]
            }, b.dataTypes[0] = "json", f = a[e], a[e] = function() {
                g = arguments
            }, d.always(function() {
                a[e] = f, b[e] && (b.jsonpCallback = c.jsonpCallback, Gb.push(e)), g && _.isFunction(f) && f(g[0]), g = f = void 0
            }), "script") : void 0
        }), _.parseHTML = function(a, b, c) {
            if (!a || "string" != typeof a) return null;
            "boolean" == typeof b && (c = b, b = !1), b = b || Z;
            var d = ga.exec(a),
                e = !c && [];
            return d ? [b.createElement(d[1])] : (d = _.buildFragment([a], b, e), e && e.length && _(e).remove(), _.merge([], d.childNodes))
        };
        var Ib = _.fn.load;
        _.fn.load = function(a, b, c) {
            if ("string" != typeof a && Ib) return Ib.apply(this, arguments);
            var d, e, f, g = this,
                h = a.indexOf(" ");
            return h >= 0 && (d = _.trim(a.slice(h)), a = a.slice(0, h)), _.isFunction(b) ? (c = b, b = void 0) : b && "object" == typeof b && (e = "POST"), g.length > 0 && _.ajax({
                url: a,
                type: e,
                dataType: "html",
                data: b
            }).done(function(a) {
                f = arguments, g.html(d ? _("<div>").append(_.parseHTML(a)).find(d) : a)
            }).complete(c && function(a, b) {
                g.each(c, f || [a.responseText, b, a])
            }), this
        }, _.each(["ajaxStart", "ajaxStop", "ajaxComplete", "ajaxError", "ajaxSuccess", "ajaxSend"], function(a, b) {
            _.fn[b] = function(a) {
                return this.on(b, a)
            }
        }), _.expr.filters.animated = function(a) {
            return _.grep(_.timers, function(b) {
                return a === b.elem
            }).length
        };
        var Jb = a.document.documentElement;
        _.offset = {
            setOffset: function(a, b, c) {
                var d, e, f, g, h, i, j, k = _.css(a, "position"),
                    l = _(a),
                    m = {};
                "static" === k && (a.style.position = "relative"), h = l.offset(), f = _.css(a, "top"), i = _.css(a, "left"), j = ("absolute" === k || "fixed" === k) && (f + i).indexOf("auto") > -1, j ? (d = l.position(), g = d.top, e = d.left) : (g = parseFloat(f) || 0, e = parseFloat(i) || 0), _.isFunction(b) && (b = b.call(a, c, h)), null != b.top && (m.top = b.top - h.top + g), null != b.left && (m.left = b.left - h.left + e), "using" in b ? b.using.call(a, m) : l.css(m)
            }
        }, _.fn.extend({
            offset: function(a) {
                if (arguments.length) return void 0 === a ? this : this.each(function(b) {
                    _.offset.setOffset(this, a, b)
                });
                var b, c, d = this[0],
                    e = {
                        top: 0,
                        left: 0
                    },
                    f = d && d.ownerDocument;
                if (f) return b = f.documentElement, _.contains(b, d) ? (typeof d.getBoundingClientRect !== za && (e = d.getBoundingClientRect()), c = P(f), {
                    top: e.top + c.pageYOffset - b.clientTop,
                    left: e.left + c.pageXOffset - b.clientLeft
                }) : e
            },
            position: function() {
                if (this[0]) {
                    var a, b, c = this[0],
                        d = {
                            top: 0,
                            left: 0
                        };
                    return "fixed" === _.css(c, "position") ? b = c.getBoundingClientRect() : (a = this.offsetParent(), b = this.offset(), _.nodeName(a[0], "html") || (d = a.offset()), d.top += _.css(a[0], "borderTopWidth", !0), d.left += _.css(a[0], "borderLeftWidth", !0)), {
                        top: b.top - d.top - _.css(c, "marginTop", !0),
                        left: b.left - d.left - _.css(c, "marginLeft", !0)
                    }
                }
            },
            offsetParent: function() {
                return this.map(function() {
                    for (var a = this.offsetParent || Jb; a && !_.nodeName(a, "html") && "static" === _.css(a, "position");) a = a.offsetParent;
                    return a || Jb
                })
            }
        }), _.each({
            scrollLeft: "pageXOffset",
            scrollTop: "pageYOffset"
        }, function(b, c) {
            var d = "pageYOffset" === c;
            _.fn[b] = function(e) {
                return qa(this, function(b, e, f) {
                    var g = P(b);
                    return void 0 === f ? g ? g[c] : b[e] : void(g ? g.scrollTo(d ? a.pageXOffset : f, d ? f : a.pageYOffset) : b[e] = f)
                }, b, e, arguments.length, null)
            }
        }), _.each(["top", "left"], function(a, b) {
            _.cssHooks[b] = w(Y.pixelPosition, function(a, c) {
                return c ? (c = v(a, b), Qa.test(c) ? _(a).position()[b] + "px" : c) : void 0
            })
        }), _.each({
            Height: "height",
            Width: "width"
        }, function(a, b) {
            _.each({
                padding: "inner" + a,
                content: b,
                "": "outer" + a
            }, function(c, d) {
                _.fn[d] = function(d, e) {
                    var f = arguments.length && (c || "boolean" != typeof d),
                        g = c || (d === !0 || e === !0 ? "margin" : "border");
                    return qa(this, function(b, c, d) {
                        var e;
                        return _.isWindow(b) ? b.document.documentElement["client" + a] : 9 === b.nodeType ? (e = b.documentElement, Math.max(b.body["scroll" + a], e["scroll" + a], b.body["offset" + a], e["offset" + a], e["client" + a])) : void 0 === d ? _.css(b, c, g) : _.style(b, c, d, g)
                    }, b, f ? d : void 0, f, null)
                }
            })
        }), _.fn.size = function() {
            return this.length
        }, _.fn.andSelf = _.fn.addBack, "function" == typeof define && define.amd && define("jquery", [], function() {
            return _
        });
        var Kb = a.jQuery,
            Lb = a.$;
        return _.noConflict = function(b) {
            return a.$ === _ && (a.$ = Lb), b && a.jQuery === _ && (a.jQuery = Kb), _
        }, typeof b === za && (a.jQuery = a.$ = _), _
    }), function() {
        function a(a) {
            function b(b, c, d, e, f, g) {
                for (; f >= 0 && g > f; f += a) {
                    var h = e ? e[f] : f;
                    d = c(d, b[h], h, b)
                }
                return d
            }
            return function(c, d, e, f) {
                d = t(d, f, 4);
                var g = !A(c) && s.keys(c),
                    h = (g || c).length,
                    i = a > 0 ? 0 : h - 1;
                return arguments.length < 3 && (e = c[g ? g[i] : i], i += a), b(c, d, e, g, i, h)
            }
        }

        function b(a) {
            return function(b, c, d) {
                c = u(c, d);
                for (var e = z(b), f = a > 0 ? 0 : e - 1; f >= 0 && e > f; f += a)
                    if (c(b[f], f, b)) return f;
                return -1
            }
        }

        function c(a, b, c) {
            return function(d, e, f) {
                var g = 0,
                    h = z(d);
                if ("number" == typeof f) a > 0 ? g = f >= 0 ? f : Math.max(f + h, g) : h = f >= 0 ? Math.min(f + 1, h) : f + h + 1;
                else if (c && f && h) return f = c(d, e), d[f] === e ? f : -1;
                if (e !== e) return f = b(k.call(d, g, h), s.isNaN), f >= 0 ? f + g : -1;
                for (f = a > 0 ? g : h - 1; f >= 0 && h > f; f += a)
                    if (d[f] === e) return f;
                return -1
            }
        }

        function d(a, b) {
            var c = F.length,
                d = a.constructor,
                e = s.isFunction(d) && d.prototype || h,
                f = "constructor";
            for (s.has(a, f) && !s.contains(b, f) && b.push(f); c--;) f = F[c], f in a && a[f] !== e[f] && !s.contains(b, f) && b.push(f)
        }
        var e = this,
            f = e._,
            g = Array.prototype,
            h = Object.prototype,
            i = Function.prototype,
            j = g.push,
            k = g.slice,
            l = h.toString,
            m = h.hasOwnProperty,
            n = Array.isArray,
            o = Object.keys,
            p = i.bind,
            q = Object.create,
            r = function() {},
            s = function(a) {
                return a instanceof s ? a : this instanceof s ? void(this._wrapped = a) : new s(a)
            };
        "undefined" != typeof exports ? ("undefined" != typeof module && module.exports && (exports = module.exports = s), exports._ = s) : e._ = s, s.VERSION = "1.8.3";
        var t = function(a, b, c) {
                if (void 0 === b) return a;
                switch (null == c ? 3 : c) {
                    case 1:
                        return function(c) {
                            return a.call(b, c)
                        };
                    case 2:
                        return function(c, d) {
                            return a.call(b, c, d)
                        };
                    case 3:
                        return function(c, d, e) {
                            return a.call(b, c, d, e)
                        };
                    case 4:
                        return function(c, d, e, f) {
                            return a.call(b, c, d, e, f)
                        }
                }
                return function() {
                    return a.apply(b, arguments)
                }
            },
            u = function(a, b, c) {
                return null == a ? s.identity : s.isFunction(a) ? t(a, b, c) : s.isObject(a) ? s.matcher(a) : s.property(a)
            };
        s.iteratee = function(a, b) {
            return u(a, b, 1 / 0)
        };
        var v = function(a, b) {
                return function(c) {
                    var d = arguments.length;
                    if (2 > d || null == c) return c;
                    for (var e = 1; d > e; e++)
                        for (var f = arguments[e], g = a(f), h = g.length, i = 0; h > i; i++) {
                            var j = g[i];
                            b && void 0 !== c[j] || (c[j] = f[j])
                        }
                    return c
                }
            },
            w = function(a) {
                if (!s.isObject(a)) return {};
                if (q) return q(a);
                r.prototype = a;
                var b = new r;
                return r.prototype = null, b
            },
            x = function(a) {
                return function(b) {
                    return null == b ? void 0 : b[a]
                }
            },
            y = Math.pow(2, 53) - 1,
            z = x("length"),
            A = function(a) {
                var b = z(a);
                return "number" == typeof b && b >= 0 && y >= b
            };
        s.each = s.forEach = function(a, b, c) {
            b = t(b, c);
            var d, e;
            if (A(a))
                for (d = 0, e = a.length; e > d; d++) b(a[d], d, a);
            else {
                var f = s.keys(a);
                for (d = 0, e = f.length; e > d; d++) b(a[f[d]], f[d], a)
            }
            return a
        }, s.map = s.collect = function(a, b, c) {
            b = u(b, c);
            for (var d = !A(a) && s.keys(a), e = (d || a).length, f = Array(e), g = 0; e > g; g++) {
                var h = d ? d[g] : g;
                f[g] = b(a[h], h, a)
            }
            return f
        }, s.reduce = s.foldl = s.inject = a(1), s.reduceRight = s.foldr = a(-1), s.find = s.detect = function(a, b, c) {
            var d;
            return d = A(a) ? s.findIndex(a, b, c) : s.findKey(a, b, c), void 0 !== d && -1 !== d ? a[d] : void 0
        }, s.filter = s.select = function(a, b, c) {
            var d = [];
            return b = u(b, c), s.each(a, function(a, c, e) {
                b(a, c, e) && d.push(a)
            }), d
        }, s.reject = function(a, b, c) {
            return s.filter(a, s.negate(u(b)), c)
        }, s.every = s.all = function(a, b, c) {
            b = u(b, c);
            for (var d = !A(a) && s.keys(a), e = (d || a).length, f = 0; e > f; f++) {
                var g = d ? d[f] : f;
                if (!b(a[g], g, a)) return !1
            }
            return !0
        }, s.some = s.any = function(a, b, c) {
            b = u(b, c);
            for (var d = !A(a) && s.keys(a), e = (d || a).length, f = 0; e > f; f++) {
                var g = d ? d[f] : f;
                if (b(a[g], g, a)) return !0
            }
            return !1
        }, s.contains = s.includes = s.include = function(a, b, c, d) {
            return A(a) || (a = s.values(a)), ("number" != typeof c || d) && (c = 0), s.indexOf(a, b, c) >= 0
        }, s.invoke = function(a, b) {
            var c = k.call(arguments, 2),
                d = s.isFunction(b);
            return s.map(a, function(a) {
                var e = d ? b : a[b];
                return null == e ? e : e.apply(a, c)
            })
        }, s.pluck = function(a, b) {
            return s.map(a, s.property(b))
        }, s.where = function(a, b) {
            return s.filter(a, s.matcher(b))
        }, s.findWhere = function(a, b) {
            return s.find(a, s.matcher(b))
        }, s.max = function(a, b, c) {
            var d, e, f = -(1 / 0),
                g = -(1 / 0);
            if (null == b && null != a) {
                a = A(a) ? a : s.values(a);
                for (var h = 0, i = a.length; i > h; h++) d = a[h], d > f && (f = d)
            } else b = u(b, c), s.each(a, function(a, c, d) {
                e = b(a, c, d), (e > g || e === -(1 / 0) && f === -(1 / 0)) && (f = a, g = e)
            });
            return f
        }, s.min = function(a, b, c) {
            var d, e, f = 1 / 0,
                g = 1 / 0;
            if (null == b && null != a) {
                a = A(a) ? a : s.values(a);
                for (var h = 0, i = a.length; i > h; h++) d = a[h], f > d && (f = d)
            } else b = u(b, c), s.each(a, function(a, c, d) {
                e = b(a, c, d), (g > e || e === 1 / 0 && f === 1 / 0) && (f = a, g = e)
            });
            return f
        }, s.shuffle = function(a) {
            for (var b, c = A(a) ? a : s.values(a), d = c.length, e = Array(d), f = 0; d > f; f++) b = s.random(0, f), b !== f && (e[f] = e[b]), e[b] = c[f];
            return e
        }, s.sample = function(a, b, c) {
            return null == b || c ? (A(a) || (a = s.values(a)), a[s.random(a.length - 1)]) : s.shuffle(a).slice(0, Math.max(0, b))
        }, s.sortBy = function(a, b, c) {
            return b = u(b, c), s.pluck(s.map(a, function(a, c, d) {
                return {
                    value: a,
                    index: c,
                    criteria: b(a, c, d)
                }
            }).sort(function(a, b) {
                var c = a.criteria,
                    d = b.criteria;
                if (c !== d) {
                    if (c > d || void 0 === c) return 1;
                    if (d > c || void 0 === d) return -1
                }
                return a.index - b.index
            }), "value")
        };
        var B = function(a) {
            return function(b, c, d) {
                var e = {};
                return c = u(c, d), s.each(b, function(d, f) {
                    var g = c(d, f, b);
                    a(e, d, g)
                }), e
            }
        };
        s.groupBy = B(function(a, b, c) {
            s.has(a, c) ? a[c].push(b) : a[c] = [b]
        }), s.indexBy = B(function(a, b, c) {
            a[c] = b
        }), s.countBy = B(function(a, b, c) {
            s.has(a, c) ? a[c]++ : a[c] = 1
        }), s.toArray = function(a) {
            return a ? s.isArray(a) ? k.call(a) : A(a) ? s.map(a, s.identity) : s.values(a) : []
        }, s.size = function(a) {
            return null == a ? 0 : A(a) ? a.length : s.keys(a).length
        }, s.partition = function(a, b, c) {
            b = u(b, c);
            var d = [],
                e = [];
            return s.each(a, function(a, c, f) {
                (b(a, c, f) ? d : e).push(a)
            }), [d, e]
        }, s.first = s.head = s.take = function(a, b, c) {
            return null == a ? void 0 : null == b || c ? a[0] : s.initial(a, a.length - b)
        }, s.initial = function(a, b, c) {
            return k.call(a, 0, Math.max(0, a.length - (null == b || c ? 1 : b)))
        }, s.last = function(a, b, c) {
            return null == a ? void 0 : null == b || c ? a[a.length - 1] : s.rest(a, Math.max(0, a.length - b))
        }, s.rest = s.tail = s.drop = function(a, b, c) {
            return k.call(a, null == b || c ? 1 : b)
        }, s.compact = function(a) {
            return s.filter(a, s.identity)
        };
        var C = function(a, b, c, d) {
            for (var e = [], f = 0, g = d || 0, h = z(a); h > g; g++) {
                var i = a[g];
                if (A(i) && (s.isArray(i) || s.isArguments(i))) {
                    b || (i = C(i, b, c));
                    var j = 0,
                        k = i.length;
                    for (e.length += k; k > j;) e[f++] = i[j++]
                } else c || (e[f++] = i)
            }
            return e
        };
        s.flatten = function(a, b) {
            return C(a, b, !1)
        }, s.without = function(a) {
            return s.difference(a, k.call(arguments, 1))
        }, s.uniq = s.unique = function(a, b, c, d) {
            s.isBoolean(b) || (d = c, c = b, b = !1), null != c && (c = u(c, d));
            for (var e = [], f = [], g = 0, h = z(a); h > g; g++) {
                var i = a[g],
                    j = c ? c(i, g, a) : i;
                b ? (g && f === j || e.push(i), f = j) : c ? s.contains(f, j) || (f.push(j), e.push(i)) : s.contains(e, i) || e.push(i)
            }
            return e
        }, s.union = function() {
            return s.uniq(C(arguments, !0, !0))
        }, s.intersection = function(a) {
            for (var b = [], c = arguments.length, d = 0, e = z(a); e > d; d++) {
                var f = a[d];
                if (!s.contains(b, f)) {
                    for (var g = 1; c > g && s.contains(arguments[g], f); g++);
                    g === c && b.push(f)
                }
            }
            return b
        }, s.difference = function(a) {
            var b = C(arguments, !0, !0, 1);
            return s.filter(a, function(a) {
                return !s.contains(b, a)
            })
        }, s.zip = function() {
            return s.unzip(arguments)
        }, s.unzip = function(a) {
            for (var b = a && s.max(a, z).length || 0, c = Array(b), d = 0; b > d; d++) c[d] = s.pluck(a, d);
            return c
        }, s.object = function(a, b) {
            for (var c = {}, d = 0, e = z(a); e > d; d++) b ? c[a[d]] = b[d] : c[a[d][0]] = a[d][1];
            return c
        }, s.findIndex = b(1), s.findLastIndex = b(-1), s.sortedIndex = function(a, b, c, d) {
            c = u(c, d, 1);
            for (var e = c(b), f = 0, g = z(a); g > f;) {
                var h = Math.floor((f + g) / 2);
                c(a[h]) < e ? f = h + 1 : g = h
            }
            return f
        }, s.indexOf = c(1, s.findIndex, s.sortedIndex), s.lastIndexOf = c(-1, s.findLastIndex), s.range = function(a, b, c) {
            null == b && (b = a || 0, a = 0), c = c || 1;
            for (var d = Math.max(Math.ceil((b - a) / c), 0), e = Array(d), f = 0; d > f; f++, a += c) e[f] = a;
            return e
        };
        var D = function(a, b, c, d, e) {
            if (!(d instanceof b)) return a.apply(c, e);
            var f = w(a.prototype),
                g = a.apply(f, e);
            return s.isObject(g) ? g : f
        };
        s.bind = function(a, b) {
            if (p && a.bind === p) return p.apply(a, k.call(arguments, 1));
            if (!s.isFunction(a)) throw new TypeError("Bind must be called on a function");
            var c = k.call(arguments, 2),
                d = function() {
                    return D(a, d, b, this, c.concat(k.call(arguments)))
                };
            return d
        }, s.partial = function(a) {
            var b = k.call(arguments, 1),
                c = function() {
                    for (var d = 0, e = b.length, f = Array(e), g = 0; e > g; g++) f[g] = b[g] === s ? arguments[d++] : b[g];
                    for (; d < arguments.length;) f.push(arguments[d++]);
                    return D(a, c, this, this, f)
                };
            return c
        }, s.bindAll = function(a) {
            var b, c, d = arguments.length;
            if (1 >= d) throw new Error("bindAll must be passed function names");
            for (b = 1; d > b; b++) c = arguments[b], a[c] = s.bind(a[c], a);
            return a
        }, s.memoize = function(a, b) {
            var c = function(d) {
                var e = c.cache,
                    f = "" + (b ? b.apply(this, arguments) : d);
                return s.has(e, f) || (e[f] = a.apply(this, arguments)), e[f]
            };
            return c.cache = {}, c
        }, s.delay = function(a, b) {
            var c = k.call(arguments, 2);
            return setTimeout(function() {
                return a.apply(null, c)
            }, b)
        }, s.defer = s.partial(s.delay, s, 1), s.throttle = function(a, b, c) {
            var d, e, f, g = null,
                h = 0;
            c || (c = {});
            var i = function() {
                h = c.leading === !1 ? 0 : s.now(), g = null, f = a.apply(d, e), g || (d = e = null)
            };
            return function() {
                var j = s.now();
                h || c.leading !== !1 || (h = j);
                var k = b - (j - h);
                return d = this, e = arguments, 0 >= k || k > b ? (g && (clearTimeout(g), g = null), h = j, f = a.apply(d, e), g || (d = e = null)) : g || c.trailing === !1 || (g = setTimeout(i, k)), f
            }
        }, s.debounce = function(a, b, c) {
            var d, e, f, g, h, i = function() {
                var j = s.now() - g;
                b > j && j >= 0 ? d = setTimeout(i, b - j) : (d = null, c || (h = a.apply(f, e), d || (f = e = null)))
            };
            return function() {
                f = this, e = arguments, g = s.now();
                var j = c && !d;
                return d || (d = setTimeout(i, b)), j && (h = a.apply(f, e), f = e = null), h
            }
        }, s.wrap = function(a, b) {
            return s.partial(b, a)
        }, s.negate = function(a) {
            return function() {
                return !a.apply(this, arguments)
            }
        }, s.compose = function() {
            var a = arguments,
                b = a.length - 1;
            return function() {
                for (var c = b, d = a[b].apply(this, arguments); c--;) d = a[c].call(this, d);
                return d
            }
        }, s.after = function(a, b) {
            return function() {
                return --a < 1 ? b.apply(this, arguments) : void 0
            }
        }, s.before = function(a, b) {
            var c;
            return function() {
                return --a > 0 && (c = b.apply(this, arguments)), 1 >= a && (b = null), c
            }
        }, s.once = s.partial(s.before, 2);
        var E = !{
                toString: null
            }.propertyIsEnumerable("toString"),
            F = ["valueOf", "isPrototypeOf", "toString", "propertyIsEnumerable", "hasOwnProperty", "toLocaleString"];
        s.keys = function(a) {
            if (!s.isObject(a)) return [];
            if (o) return o(a);
            var b = [];
            for (var c in a) s.has(a, c) && b.push(c);
            return E && d(a, b), b
        }, s.allKeys = function(a) {
            if (!s.isObject(a)) return [];
            var b = [];
            for (var c in a) b.push(c);
            return E && d(a, b), b
        }, s.values = function(a) {
            for (var b = s.keys(a), c = b.length, d = Array(c), e = 0; c > e; e++) d[e] = a[b[e]];
            return d
        }, s.mapObject = function(a, b, c) {
            b = u(b, c);
            for (var d, e = s.keys(a), f = e.length, g = {}, h = 0; f > h; h++) d = e[h], g[d] = b(a[d], d, a);
            return g
        }, s.pairs = function(a) {
            for (var b = s.keys(a), c = b.length, d = Array(c), e = 0; c > e; e++) d[e] = [b[e], a[b[e]]];
            return d
        }, s.invert = function(a) {
            for (var b = {}, c = s.keys(a), d = 0, e = c.length; e > d; d++) b[a[c[d]]] = c[d];
            return b
        }, s.functions = s.methods = function(a) {
            var b = [];
            for (var c in a) s.isFunction(a[c]) && b.push(c);
            return b.sort()
        }, s.extend = v(s.allKeys), s.extendOwn = s.assign = v(s.keys), s.findKey = function(a, b, c) {
            b = u(b, c);
            for (var d, e = s.keys(a), f = 0, g = e.length; g > f; f++)
                if (d = e[f], b(a[d], d, a)) return d
        }, s.pick = function(a, b, c) {
            var d, e, f = {},
                g = a;
            if (null == g) return f;
            s.isFunction(b) ? (e = s.allKeys(g), d = t(b, c)) : (e = C(arguments, !1, !1, 1), d = function(a, b, c) {
                return b in c
            }, g = Object(g));
            for (var h = 0, i = e.length; i > h; h++) {
                var j = e[h],
                    k = g[j];
                d(k, j, g) && (f[j] = k)
            }
            return f
        }, s.omit = function(a, b, c) {
            if (s.isFunction(b)) b = s.negate(b);
            else {
                var d = s.map(C(arguments, !1, !1, 1), String);
                b = function(a, b) {
                    return !s.contains(d, b)
                }
            }
            return s.pick(a, b, c)
        }, s.defaults = v(s.allKeys, !0), s.create = function(a, b) {
            var c = w(a);
            return b && s.extendOwn(c, b), c
        }, s.clone = function(a) {
            return s.isObject(a) ? s.isArray(a) ? a.slice() : s.extend({}, a) : a
        }, s.tap = function(a, b) {
            return b(a), a
        }, s.isMatch = function(a, b) {
            var c = s.keys(b),
                d = c.length;
            if (null == a) return !d;
            for (var e = Object(a), f = 0; d > f; f++) {
                var g = c[f];
                if (b[g] !== e[g] || !(g in e)) return !1
            }
            return !0
        };
        var G = function(a, b, c, d) {
            if (a === b) return 0 !== a || 1 / a === 1 / b;
            if (null == a || null == b) return a === b;
            a instanceof s && (a = a._wrapped), b instanceof s && (b = b._wrapped);
            var e = l.call(a);
            if (e !== l.call(b)) return !1;
            switch (e) {
                case "[object RegExp]":
                case "[object String]":
                    return "" + a == "" + b;
                case "[object Number]":
                    return +a !== +a ? +b !== +b : 0 === +a ? 1 / +a === 1 / b : +a === +b;
                case "[object Date]":
                case "[object Boolean]":
                    return +a === +b
            }
            var f = "[object Array]" === e;
            if (!f) {
                if ("object" != typeof a || "object" != typeof b) return !1;
                var g = a.constructor,
                    h = b.constructor;
                if (g !== h && !(s.isFunction(g) && g instanceof g && s.isFunction(h) && h instanceof h) && "constructor" in a && "constructor" in b) return !1;
            }
            c = c || [], d = d || [];
            for (var i = c.length; i--;)
                if (c[i] === a) return d[i] === b;
            if (c.push(a), d.push(b), f) {
                if (i = a.length, i !== b.length) return !1;
                for (; i--;)
                    if (!G(a[i], b[i], c, d)) return !1
            } else {
                var j, k = s.keys(a);
                if (i = k.length, s.keys(b).length !== i) return !1;
                for (; i--;)
                    if (j = k[i], !s.has(b, j) || !G(a[j], b[j], c, d)) return !1
            }
            return c.pop(), d.pop(), !0
        };
        s.isEqual = function(a, b) {
            return G(a, b)
        }, s.isEmpty = function(a) {
            return null == a ? !0 : A(a) && (s.isArray(a) || s.isString(a) || s.isArguments(a)) ? 0 === a.length : 0 === s.keys(a).length
        }, s.isElement = function(a) {
            return !(!a || 1 !== a.nodeType)
        }, s.isArray = n || function(a) {
            return "[object Array]" === l.call(a)
        }, s.isObject = function(a) {
            var b = typeof a;
            return "function" === b || "object" === b && !!a
        }, s.each(["Arguments", "Function", "String", "Number", "Date", "RegExp", "Error"], function(a) {
            s["is" + a] = function(b) {
                return l.call(b) === "[object " + a + "]"
            }
        }), s.isArguments(arguments) || (s.isArguments = function(a) {
            return s.has(a, "callee")
        }), "function" != typeof /./ && "object" != typeof Int8Array && (s.isFunction = function(a) {
            return "function" == typeof a || !1
        }), s.isFinite = function(a) {
            return isFinite(a) && !isNaN(parseFloat(a))
        }, s.isNaN = function(a) {
            return s.isNumber(a) && a !== +a
        }, s.isBoolean = function(a) {
            return a === !0 || a === !1 || "[object Boolean]" === l.call(a)
        }, s.isNull = function(a) {
            return null === a
        }, s.isUndefined = function(a) {
            return void 0 === a
        }, s.has = function(a, b) {
            return null != a && m.call(a, b)
        }, s.noConflict = function() {
            return e._ = f, this
        }, s.identity = function(a) {
            return a
        }, s.constant = function(a) {
            return function() {
                return a
            }
        }, s.noop = function() {}, s.property = x, s.propertyOf = function(a) {
            return null == a ? function() {} : function(b) {
                return a[b]
            }
        }, s.matcher = s.matches = function(a) {
            return a = s.extendOwn({}, a),
                function(b) {
                    return s.isMatch(b, a)
                }
        }, s.times = function(a, b, c) {
            var d = Array(Math.max(0, a));
            b = t(b, c, 1);
            for (var e = 0; a > e; e++) d[e] = b(e);
            return d
        }, s.random = function(a, b) {
            return null == b && (b = a, a = 0), a + Math.floor(Math.random() * (b - a + 1))
        }, s.now = Date.now || function() {
            return (new Date).getTime()
        };
        var H = {
                "&": "&amp;",
                "<": "&lt;",
                ">": "&gt;",
                '"': "&quot;",
                "'": "&#x27;",
                "`": "&#x60;"
            },
            I = s.invert(H),
            J = function(a) {
                var b = function(b) {
                        return a[b]
                    },
                    c = "(?:" + s.keys(a).join("|") + ")",
                    d = RegExp(c),
                    e = RegExp(c, "g");
                return function(a) {
                    return a = null == a ? "" : "" + a, d.test(a) ? a.replace(e, b) : a
                }
            };
        s.escape = J(H), s.unescape = J(I), s.result = function(a, b, c) {
            var d = null == a ? void 0 : a[b];
            return void 0 === d && (d = c), s.isFunction(d) ? d.call(a) : d
        };
        var K = 0;
        s.uniqueId = function(a) {
            var b = ++K + "";
            return a ? a + b : b
        }, s.templateSettings = {
            evaluate: /<%([\s\S]+?)%>/g,
            interpolate: /<%=([\s\S]+?)%>/g,
            escape: /<%-([\s\S]+?)%>/g
        };
        var L = /(.)^/,
            M = {
                "'": "'",
                "\\": "\\",
                "\r": "r",
                "\n": "n",
                "\u2028": "u2028",
                "\u2029": "u2029"
            },
            N = /\\|'|\r|\n|\u2028|\u2029/g,
            O = function(a) {
                return "\\" + M[a]
            };
        s.template = function(a, b, c) {
            !b && c && (b = c), b = s.defaults({}, b, s.templateSettings);
            var d = RegExp([(b.escape || L).source, (b.interpolate || L).source, (b.evaluate || L).source].join("|") + "|$", "g"),
                e = 0,
                f = "__p+='";
            a.replace(d, function(b, c, d, g, h) {
                return f += a.slice(e, h).replace(N, O), e = h + b.length, c ? f += "'+\n((__t=(" + c + "))==null?'':_.escape(__t))+\n'" : d ? f += "'+\n((__t=(" + d + "))==null?'':__t)+\n'" : g && (f += "';\n" + g + "\n__p+='"), b
            }), f += "';\n", b.variable || (f = "with(obj||{}){\n" + f + "}\n"), f = "var __t,__p='',__j=Array.prototype.join,print=function(){__p+=__j.call(arguments,'');};\n" + f + "return __p;\n";
            try {
                var g = new Function(b.variable || "obj", "_", f)
            } catch (h) {
                throw h.source = f, h
            }
            var i = function(a) {
                    return g.call(this, a, s)
                },
                j = b.variable || "obj";
            return i.source = "function(" + j + "){\n" + f + "}", i
        }, s.chain = function(a) {
            var b = s(a);
            return b._chain = !0, b
        };
        var P = function(a, b) {
            return a._chain ? s(b).chain() : b
        };
        s.mixin = function(a) {
            s.each(s.functions(a), function(b) {
                var c = s[b] = a[b];
                s.prototype[b] = function() {
                    var a = [this._wrapped];
                    return j.apply(a, arguments), P(this, c.apply(s, a))
                }
            })
        }, s.mixin(s), s.each(["pop", "push", "reverse", "shift", "sort", "splice", "unshift"], function(a) {
            var b = g[a];
            s.prototype[a] = function() {
                var c = this._wrapped;
                return b.apply(c, arguments), "shift" !== a && "splice" !== a || 0 !== c.length || delete c[0], P(this, c)
            }
        }), s.each(["concat", "join", "slice"], function(a) {
            var b = g[a];
            s.prototype[a] = function() {
                return P(this, b.apply(this._wrapped, arguments))
            }
        }), s.prototype.value = function() {
            return this._wrapped
        }, s.prototype.valueOf = s.prototype.toJSON = s.prototype.value, s.prototype.toString = function() {
            return "" + this._wrapped
        }, "function" == typeof define && define.amd && define("underscore", [], function() {
            return s
        })
    }.call(this), function(a, b, c) {
        "use strict";

        function d(a) {
            return function() {
                var b, c, d = arguments[0],
                    e = "[" + (a ? a + ":" : "") + d + "] ",
                    f = arguments[1],
                    g = arguments,
                    h = function(a) {
                        return "function" == typeof a ? a.toString().replace(/ \{[\s\S]*$/, "") : "undefined" == typeof a ? "undefined" : "string" != typeof a ? JSON.stringify(a) : a
                    };
                for (b = e + f.replace(/\{\d+\}/g, function(a) {
                        var b, c = +a.slice(1, -1);
                        return c + 2 < g.length ? (b = g[c + 2], "function" == typeof b ? b.toString().replace(/ ?\{[\s\S]*$/, "") : "undefined" == typeof b ? "undefined" : "string" != typeof b ? Q(b) : b) : a
                    }), b = b + "\nhttp://errors.angularjs.org/1.2.28/" + (a ? a + "/" : "") + d, c = 2; c < arguments.length; c++) b = b + (2 == c ? "?" : "&") + "p" + (c - 2) + "=" + encodeURIComponent(h(arguments[c]));
                return new Error(b)
            }
        }

        function e(a) {
            if (null == a || z(a)) return !1;
            var b = a.length;
            return 1 === a.nodeType && b ? !0 : u(a) || Ec(a) || 0 === b || "number" == typeof b && b > 0 && b - 1 in a
        }

        function f(a, b, c) {
            var d;
            if (a)
                if (x(a))
                    for (d in a) "prototype" == d || "length" == d || "name" == d || a.hasOwnProperty && !a.hasOwnProperty(d) || b.call(c, a[d], d);
                else if (Ec(a) || e(a))
                for (d = 0; d < a.length; d++) b.call(c, a[d], d);
            else if (a.forEach && a.forEach !== f) a.forEach(b, c);
            else
                for (d in a) a.hasOwnProperty(d) && b.call(c, a[d], d);
            return a
        }

        function g(a) {
            var b = [];
            for (var c in a) a.hasOwnProperty(c) && b.push(c);
            return b.sort()
        }

        function h(a, b, c) {
            for (var d = g(a), e = 0; e < d.length; e++) b.call(c, a[d[e]], d[e]);
            return d
        }

        function i(a) {
            return function(b, c) {
                a(c, b)
            }
        }

        function j() {
            for (var a, b = Dc.length; b;) {
                if (b--, a = Dc[b].charCodeAt(0), 57 == a) return Dc[b] = "A", Dc.join("");
                if (90 != a) return Dc[b] = String.fromCharCode(a + 1), Dc.join("");
                Dc[b] = "0"
            }
            return Dc.unshift("0"), Dc.join("")
        }

        function k(a, b) {
            b ? a.$$hashKey = b : delete a.$$hashKey
        }

        function l(a) {
            var b = a.$$hashKey;
            return f(arguments, function(b) {
                b !== a && f(b, function(b, c) {
                    a[c] = b
                })
            }), k(a, b), a
        }

        function m(a) {
            return parseInt(a, 10)
        }

        function n(a, b) {
            return l(new(l(function() {}, {
                prototype: a
            })), b)
        }

        function o() {}

        function p(a) {
            return a
        }

        function q(a) {
            return function() {
                return a
            }
        }

        function r(a) {
            return "undefined" == typeof a
        }

        function s(a) {
            return "undefined" != typeof a
        }

        function t(a) {
            return null != a && "object" == typeof a
        }

        function u(a) {
            return "string" == typeof a
        }

        function v(a) {
            return "number" == typeof a
        }

        function w(a) {
            return "[object Date]" === Ac.call(a)
        }

        function x(a) {
            return "function" == typeof a
        }

        function y(a) {
            return "[object RegExp]" === Ac.call(a)
        }

        function z(a) {
            return a && a.document && a.location && a.alert && a.setInterval
        }

        function A(a) {
            return a && a.$evalAsync && a.$watch
        }

        function B(a) {
            return "[object File]" === Ac.call(a)
        }

        function C(a) {
            return "[object Blob]" === Ac.call(a)
        }

        function D(a) {
            return a && x(a.then)
        }

        function E(a) {
            return !(!a || !(a.nodeName || a.prop && a.attr && a.find))
        }

        function F(a, b, c) {
            var d = [];
            return f(a, function(a, e, f) {
                d.push(b.call(c, a, e, f))
            }), d
        }

        function G(a, b) {
            return -1 != H(a, b)
        }

        function H(a, b) {
            if (a.indexOf) return a.indexOf(b);
            for (var c = 0; c < a.length; c++)
                if (b === a[c]) return c;
            return -1
        }

        function I(a, b) {
            var c = H(a, b);
            return c >= 0 && a.splice(c, 1), b
        }

        function J(a, b, c, d) {
            if (z(a) || A(a)) throw Bc("cpws", "Can't copy! Making copies of Window or Scope instances is not supported.");
            if (b) {
                if (a === b) throw Bc("cpi", "Can't copy! Source and destination are identical.");
                if (c = c || [], d = d || [], t(a)) {
                    var e = H(c, a);
                    if (-1 !== e) return d[e];
                    c.push(a), d.push(b)
                }
                var g;
                if (Ec(a)) {
                    b.length = 0;
                    for (var h = 0; h < a.length; h++) g = J(a[h], null, c, d), t(a[h]) && (c.push(a[h]), d.push(g)), b.push(g)
                } else {
                    var i = b.$$hashKey;
                    Ec(b) ? b.length = 0 : f(b, function(a, c) {
                        delete b[c]
                    });
                    for (var j in a) g = J(a[j], null, c, d), t(a[j]) && (c.push(a[j]), d.push(g)), b[j] = g;
                    k(b, i)
                }
            } else b = a, a && (Ec(a) ? b = J(a, [], c, d) : w(a) ? b = new Date(a.getTime()) : y(a) ? (b = new RegExp(a.source, a.toString().match(/[^\/]*$/)[0]), b.lastIndex = a.lastIndex) : t(a) && (b = J(a, {}, c, d)));
            return b
        }

        function K(a, b) {
            if (Ec(a)) {
                b = b || [];
                for (var c = 0; c < a.length; c++) b[c] = a[c]
            } else if (t(a)) {
                b = b || {};
                for (var d in a) !pc.call(a, d) || "$" === d.charAt(0) && "$" === d.charAt(1) || (b[d] = a[d])
            }
            return b || a
        }

        function L(a, b) {
            if (a === b) return !0;
            if (null === a || null === b) return !1;
            if (a !== a && b !== b) return !0;
            var d, e, f, g = typeof a,
                h = typeof b;
            if (g == h && "object" == g) {
                if (!Ec(a)) {
                    if (w(a)) return w(b) ? isNaN(a.getTime()) && isNaN(b.getTime()) || a.getTime() === b.getTime() : !1;
                    if (y(a) && y(b)) return a.toString() == b.toString();
                    if (A(a) || A(b) || z(a) || z(b) || Ec(b)) return !1;
                    f = {};
                    for (e in a)
                        if ("$" !== e.charAt(0) && !x(a[e])) {
                            if (!L(a[e], b[e])) return !1;
                            f[e] = !0
                        }
                    for (e in b)
                        if (!f.hasOwnProperty(e) && "$" !== e.charAt(0) && b[e] !== c && !x(b[e])) return !1;
                    return !0
                }
                if (!Ec(b)) return !1;
                if ((d = a.length) == b.length) {
                    for (e = 0; d > e; e++)
                        if (!L(a[e], b[e])) return !1;
                    return !0
                }
            }
            return !1
        }

        function M(a, b, c) {
            return a.concat(yc.call(b, c))
        }

        function N(a, b) {
            return yc.call(a, b || 0)
        }

        function O(a, b) {
            var c = arguments.length > 2 ? N(arguments, 2) : [];
            return !x(b) || b instanceof RegExp ? b : c.length ? function() {
                return arguments.length ? b.apply(a, c.concat(yc.call(arguments, 0))) : b.apply(a, c)
            } : function() {
                return arguments.length ? b.apply(a, arguments) : b.call(a)
            }
        }

        function P(a, d) {
            var e = d;
            return "string" == typeof a && "$" === a.charAt(0) ? e = c : z(d) ? e = "$WINDOW" : d && b === d ? e = "$DOCUMENT" : A(d) && (e = "$SCOPE"), e
        }

        function Q(a, b) {
            return "undefined" == typeof a ? c : JSON.stringify(a, P, b ? "  " : null)
        }

        function R(a) {
            return u(a) ? JSON.parse(a) : a
        }

        function S(a) {
            if ("function" == typeof a) a = !0;
            else if (a && 0 !== a.length) {
                var b = oc("" + a);
                a = !("f" == b || "0" == b || "false" == b || "no" == b || "n" == b || "[]" == b)
            } else a = !1;
            return a
        }

        function T(a) {
            a = uc(a).clone();
            try {
                a.empty()
            } catch (b) {}
            var c = 3,
                d = uc("<div>").append(a).html();
            try {
                return a[0].nodeType === c ? oc(d) : d.match(/^(<[^>]+>)/)[1].replace(/^<([\w\-]+)/, function(a, b) {
                    return "<" + oc(b)
                })
            } catch (b) {
                return oc(d)
            }
        }

        function U(a) {
            try {
                return decodeURIComponent(a)
            } catch (b) {}
        }

        function V(a) {
            var b, c, d = {};
            return f((a || "").split("&"), function(a) {
                if (a && (b = a.replace(/\+/g, "%20").split("="), c = U(b[0]), s(c))) {
                    var e = s(b[1]) ? U(b[1]) : !0;
                    pc.call(d, c) ? Ec(d[c]) ? d[c].push(e) : d[c] = [d[c], e] : d[c] = e
                }
            }), d
        }

        function W(a) {
            var b = [];
            return f(a, function(a, c) {
                Ec(a) ? f(a, function(a) {
                    b.push(Y(c, !0) + (a === !0 ? "" : "=" + Y(a, !0)))
                }) : b.push(Y(c, !0) + (a === !0 ? "" : "=" + Y(a, !0)))
            }), b.length ? b.join("&") : ""
        }

        function X(a) {
            return Y(a, !0).replace(/%26/gi, "&").replace(/%3D/gi, "=").replace(/%2B/gi, "+")
        }

        function Y(a, b) {
            return encodeURIComponent(a).replace(/%40/gi, "@").replace(/%3A/gi, ":").replace(/%24/g, "$").replace(/%2C/gi, ",").replace(/%20/g, b ? "%20" : "+")
        }

        function Z(a, c) {
            function d(a) {
                a && h.push(a)
            }
            var e, g, h = [a],
                i = ["ng:app", "ng-app", "x-ng-app", "data-ng-app"],
                j = /\sng[:\-]app(:\s*([\w\d_]+);?)?\s/;
            f(i, function(c) {
                i[c] = !0, d(b.getElementById(c)), c = c.replace(":", "\\:"), a.querySelectorAll && (f(a.querySelectorAll("." + c), d), f(a.querySelectorAll("." + c + "\\:"), d), f(a.querySelectorAll("[" + c + "]"), d))
            }), f(h, function(a) {
                if (!e) {
                    var b = " " + a.className + " ",
                        c = j.exec(b);
                    c ? (e = a, g = (c[2] || "").replace(/\s+/g, ",")) : f(a.attributes, function(b) {
                        !e && i[b.name] && (e = a, g = b.value)
                    })
                }
            }), e && c(e, g ? [g] : [])
        }

        function $(c, d) {
            var e = function() {
                    if (c = uc(c), c.injector()) {
                        var a = c[0] === b ? "document" : T(c);
                        throw Bc("btstrpd", "App Already Bootstrapped with this Element '{0}'", a.replace(/</, "&lt;").replace(/>/, "&gt;"))
                    }
                    d = d || [], d.unshift(["$provide", function(a) {
                        a.value("$rootElement", c)
                    }]), d.unshift("ng");
                    var e = Ha(d);
                    return e.invoke(["$rootScope", "$rootElement", "$compile", "$injector", "$animate", function(a, b, c, d, e) {
                        a.$apply(function() {
                            b.data("$injector", d), c(b)(a)
                        })
                    }]), e
                },
                g = /^NG_DEFER_BOOTSTRAP!/;
            return a && !g.test(a.name) ? e() : (a.name = a.name.replace(g, ""), void(Cc.resumeBootstrap = function(a) {
                f(a, function(a) {
                    d.push(a)
                }), e()
            }))
        }

        function _(a, b) {
            return b = b || "_", a.replace(Hc, function(a, c) {
                return (c ? b : "") + a.toLowerCase()
            })
        }

        function aa() {
            vc = a.jQuery, vc && vc.fn.on ? (uc = vc, l(vc.fn, {
                scope: Vc.scope,
                isolateScope: Vc.isolateScope,
                controller: Vc.controller,
                injector: Vc.injector,
                inheritedData: Vc.inheritedData
            }), ka("remove", !0, !0, !1), ka("empty", !1, !1, !1), ka("html", !1, !1, !0)) : uc = oa, Cc.element = uc
        }

        function ba(a, b, c) {
            if (!a) throw Bc("areq", "Argument '{0}' is {1}", b || "?", c || "required");
            return a
        }

        function ca(a, b, c) {
            return c && Ec(a) && (a = a[a.length - 1]), ba(x(a), b, "not a function, got " + (a && "object" == typeof a ? a.constructor.name || "Object" : typeof a)), a
        }

        function da(a, b) {
            if ("hasOwnProperty" === a) throw Bc("badname", "hasOwnProperty is not a valid {0} name", b)
        }

        function ea(a, b, c) {
            if (!b) return a;
            for (var d, e = b.split("."), f = a, g = e.length, h = 0; g > h; h++) d = e[h], a && (a = (f = a)[d]);
            return !c && x(a) ? O(f, a) : a
        }

        function fa(a) {
            var b = a[0],
                c = a[a.length - 1];
            if (b === c) return uc(b);
            var d = b,
                e = [d];
            do {
                if (d = d.nextSibling, !d) break;
                e.push(d)
            } while (d !== c);
            return uc(e)
        }

        function ga(a) {
            function b(a, b, c) {
                return a[b] || (a[b] = c())
            }
            var c = d("$injector"),
                e = d("ng"),
                f = b(a, "angular", Object);
            return f.$$minErr = f.$$minErr || d, b(f, "module", function() {
                var a = {};
                return function(d, f, g) {
                    var h = function(a, b) {
                        if ("hasOwnProperty" === a) throw e("badname", "hasOwnProperty is not a valid {0} name", b)
                    };
                    return h(d, "module"), f && a.hasOwnProperty(d) && (a[d] = null), b(a, d, function() {
                        function a(a, c, d) {
                            return function() {
                                return b[d || "push"]([a, c, arguments]), i
                            }
                        }
                        if (!f) throw c("nomod", "Module '{0}' is not available! You either misspelled the module name or forgot to load it. If registering a module ensure that you specify the dependencies as the second argument.", d);
                        var b = [],
                            e = [],
                            h = a("$injector", "invoke"),
                            i = {
                                _invokeQueue: b,
                                _runBlocks: e,
                                requires: f,
                                name: d,
                                provider: a("$provide", "provider"),
                                factory: a("$provide", "factory"),
                                service: a("$provide", "service"),
                                value: a("$provide", "value"),
                                constant: a("$provide", "constant", "unshift"),
                                animation: a("$animateProvider", "register"),
                                filter: a("$filterProvider", "register"),
                                controller: a("$controllerProvider", "register"),
                                directive: a("$compileProvider", "directive"),
                                config: h,
                                run: function(a) {
                                    return e.push(a), this
                                }
                            };
                        return g && h(g), i
                    })
                }
            })
        }

        function ha(b) {
            l(b, {
                bootstrap: $,
                copy: J,
                extend: l,
                equals: L,
                element: uc,
                forEach: f,
                injector: Ha,
                noop: o,
                bind: O,
                toJson: Q,
                fromJson: R,
                identity: p,
                isUndefined: r,
                isDefined: s,
                isString: u,
                isFunction: x,
                isObject: t,
                isNumber: v,
                isElement: E,
                isArray: Ec,
                version: Ic,
                isDate: w,
                lowercase: oc,
                uppercase: qc,
                callbacks: {
                    counter: 0
                },
                $$minErr: d,
                $$csp: Gc
            }), wc = ga(a);
            try {
                wc("ngLocale")
            } catch (c) {
                wc("ngLocale", []).provider("$locale", cb)
            }
            wc("ng", ["ngLocale"], ["$provide", function(a) {
                a.provider({
                    $$sanitizeUri: Eb
                }), a.provider("$compile", Oa).directive({
                    a: Fd,
                    input: Qd,
                    textarea: Qd,
                    form: Jd,
                    script: ye,
                    select: Be,
                    style: De,
                    option: Ce,
                    ngBind: ae,
                    ngBindHtml: ce,
                    ngBindTemplate: be,
                    ngClass: de,
                    ngClassEven: fe,
                    ngClassOdd: ee,
                    ngCloak: ge,
                    ngController: he,
                    ngForm: Kd,
                    ngHide: se,
                    ngIf: ke,
                    ngInclude: le,
                    ngInit: ne,
                    ngNonBindable: oe,
                    ngPluralize: pe,
                    ngRepeat: qe,
                    ngShow: re,
                    ngStyle: te,
                    ngSwitch: ue,
                    ngSwitchWhen: ve,
                    ngSwitchDefault: we,
                    ngOptions: Ae,
                    ngTransclude: xe,
                    ngModel: Wd,
                    ngList: Zd,
                    ngChange: Xd,
                    required: Yd,
                    ngRequired: Yd,
                    ngValue: _d
                }).directive({
                    ngInclude: me
                }).directive(Gd).directive(ie), a.provider({
                    $anchorScroll: Ia,
                    $animate: cd,
                    $browser: La,
                    $cacheFactory: Ma,
                    $controller: Ra,
                    $document: Sa,
                    $exceptionHandler: Ta,
                    $filter: Pb,
                    $interpolate: ab,
                    $interval: bb,
                    $http: Ya,
                    $httpBackend: $a,
                    $location: pb,
                    $log: qb,
                    $parse: zb,
                    $rootScope: Db,
                    $q: Ab,
                    $sce: Jb,
                    $sceDelegate: Ib,
                    $sniffer: Kb,
                    $templateCache: Na,
                    $timeout: Lb,
                    $window: Ob,
                    $$rAF: Cb,
                    $$asyncCallback: Ja
                })
            }])
        }

        function ia() {
            return ++Kc
        }

        function ja(a) {
            return a.replace(Nc, function(a, b, c, d) {
                return d ? c.toUpperCase() : c
            }).replace(Oc, "Moz$1")
        }

        function ka(a, b, c, d) {
            function e(a) {
                var e, g, h, i, j, k, l, m = c && a ? [this.filter(a)] : [this],
                    n = b;
                if (!d || null != a)
                    for (; m.length;)
                        for (e = m.shift(), g = 0, h = e.length; h > g; g++)
                            for (i = uc(e[g]), n ? i.triggerHandler("$destroy") : n = !n, j = 0, k = (l = i.children()).length; k > j; j++) m.push(vc(l[j]));
                return f.apply(this, arguments)
            }
            var f = vc.fn[a];
            f = f.$original || f, e.$original = f, vc.fn[a] = e
        }

        function la(a) {
            return !Rc.test(a)
        }

        function ma(a, b) {
            var c, d, e, f, g, h, i = b.createDocumentFragment(),
                j = [];
            if (la(a)) j.push(b.createTextNode(a));
            else {
                for (c = i.appendChild(b.createElement("div")), d = (Sc.exec(a) || ["", ""])[1].toLowerCase(), e = Uc[d] || Uc._default, c.innerHTML = "<div>&#160;</div>" + e[1] + a.replace(Tc, "<$1></$2>") + e[2], c.removeChild(c.firstChild), f = e[0]; f--;) c = c.lastChild;
                for (g = 0, h = c.childNodes.length; h > g; ++g) j.push(c.childNodes[g]);
                c = i.firstChild, c.textContent = ""
            }
            return i.textContent = "", i.innerHTML = "", j
        }

        function na(a, c) {
            c = c || b;
            var d;
            return (d = Qc.exec(a)) ? [c.createElement(d[1])] : ma(a, c)
        }

        function oa(a) {
            if (a instanceof oa) return a;
            if (u(a) && (a = Fc(a)), !(this instanceof oa)) {
                if (u(a) && "<" != a.charAt(0)) throw Pc("nosel", "Looking up elements via selectors is not supported by jqLite! See: http://docs.angularjs.org/api/angular.element");
                return new oa(a)
            }
            if (u(a)) {
                ya(this, na(a));
                var c = uc(b.createDocumentFragment());
                c.append(this)
            } else ya(this, a)
        }

        function pa(a) {
            return a.cloneNode(!0)
        }

        function qa(a) {
            sa(a);
            for (var b = 0, c = a.childNodes || []; b < c.length; b++) qa(c[b])
        }

        function ra(a, b, c, d) {
            if (s(d)) throw Pc("offargs", "jqLite#off() does not support the `selector` argument");
            var e = ta(a, "events"),
                g = ta(a, "handle");
            g && (r(b) ? f(e, function(b, c) {
                Mc(a, c, b), delete e[c]
            }) : f(b.split(" "), function(b) {
                r(c) ? (Mc(a, b, e[b]), delete e[b]) : I(e[b] || [], c)
            }))
        }

        function sa(a, b) {
            var d = a.ng339,
                e = Jc[d];
            if (e) {
                if (b) return void delete Jc[d].data[b];
                e.handle && (e.events.$destroy && e.handle({}, "$destroy"), ra(a)), delete Jc[d], a.ng339 = c
            }
        }

        function ta(a, b, c) {
            var d = a.ng339,
                e = Jc[d || -1];
            return s(c) ? (e || (a.ng339 = d = ia(), e = Jc[d] = {}), void(e[b] = c)) : e && e[b]
        }

        function ua(a, b, c) {
            var d = ta(a, "data"),
                e = s(c),
                f = !e && s(b),
                g = f && !t(b);
            if (d || g || ta(a, "data", d = {}), e) d[b] = c;
            else {
                if (!f) return d;
                if (g) return d && d[b];
                l(d, b)
            }
        }

        function va(a, b) {
            return a.getAttribute ? (" " + (a.getAttribute("class") || "") + " ").replace(/[\n\t]/g, " ").indexOf(" " + b + " ") > -1 : !1
        }

        function wa(a, b) {
            b && a.setAttribute && f(b.split(" "), function(b) {
                a.setAttribute("class", Fc((" " + (a.getAttribute("class") || "") + " ").replace(/[\n\t]/g, " ").replace(" " + Fc(b) + " ", " ")))
            })
        }

        function xa(a, b) {
            if (b && a.setAttribute) {
                var c = (" " + (a.getAttribute("class") || "") + " ").replace(/[\n\t]/g, " ");
                f(b.split(" "), function(a) {
                    a = Fc(a), -1 === c.indexOf(" " + a + " ") && (c += a + " ")
                }), a.setAttribute("class", Fc(c))
            }
        }

        function ya(a, b) {
            if (b) {
                b = b.nodeName || !s(b.length) || z(b) ? [b] : b;
                for (var c = 0; c < b.length; c++) a.push(b[c])
            }
        }

        function za(a, b) {
            return Aa(a, "$" + (b || "ngController") + "Controller")
        }

        function Aa(a, b, d) {
            9 == a.nodeType && (a = a.documentElement);
            for (var e = Ec(b) ? b : [b]; a;) {
                for (var f = 0, g = e.length; g > f; f++)
                    if ((d = uc.data(a, e[f])) !== c) return d;
                a = a.parentNode || 11 === a.nodeType && a.host
            }
        }

        function Ba(a) {
            for (var b = 0, c = a.childNodes; b < c.length; b++) qa(c[b]);
            for (; a.firstChild;) a.removeChild(a.firstChild)
        }

        function Ca(a, b) {
            var c = Wc[b.toLowerCase()];
            return c && Xc[a.nodeName] && c
        }

        function Da(a, c) {
            var d = function(d, e) {
                if (d.preventDefault || (d.preventDefault = function() {
                        d.returnValue = !1
                    }), d.stopPropagation || (d.stopPropagation = function() {
                        d.cancelBubble = !0
                    }), d.target || (d.target = d.srcElement || b), r(d.defaultPrevented)) {
                    var g = d.preventDefault;
                    d.preventDefault = function() {
                        d.defaultPrevented = !0, g.call(d)
                    }, d.defaultPrevented = !1
                }
                d.isDefaultPrevented = function() {
                    return d.defaultPrevented || d.returnValue === !1
                };
                var h = K(c[e || d.type] || []);
                f(h, function(b) {
                    b.call(a, d)
                }), 8 >= tc ? (d.preventDefault = null, d.stopPropagation = null, d.isDefaultPrevented = null) : (delete d.preventDefault, delete d.stopPropagation, delete d.isDefaultPrevented)
            };
            return d.elem = a, d
        }

        function Ea(a, b) {
            var d, e = typeof a;
            return "function" == e || "object" == e && null !== a ? "function" == typeof(d = a.$$hashKey) ? d = a.$$hashKey() : d === c && (d = a.$$hashKey = (b || j)()) : d = a, e + ":" + d
        }

        function Fa(a, b) {
            if (b) {
                var c = 0;
                this.nextUid = function() {
                    return ++c
                }
            }
            f(a, this.put, this)
        }

        function Ga(a) {
            var b, c, d, e;
            return "function" == typeof a ? (b = a.$inject) || (b = [], a.length && (c = a.toString().replace(_c, ""), d = c.match(Yc), f(d[1].split(Zc), function(a) {
                a.replace($c, function(a, c, d) {
                    b.push(d)
                })
            })), a.$inject = b) : Ec(a) ? (e = a.length - 1, ca(a[e], "fn"), b = a.slice(0, e)) : ca(a, "fn", !0), b
        }

        function Ha(a) {
            function b(a) {
                return function(b, c) {
                    return t(b) ? void f(b, i(a)) : a(b, c)
                }
            }

            function c(a, b) {
                if (da(a, "service"), (x(b) || Ec(b)) && (b = v.instantiate(b)), !b.$get) throw ad("pget", "Provider '{0}' must define $get factory method.", a);
                return s[a + n] = b
            }

            function d(a, b) {
                return c(a, {
                    $get: b
                })
            }

            function e(a, b) {
                return d(a, ["$injector", function(a) {
                    return a.instantiate(b)
                }])
            }

            function g(a, b) {
                return d(a, q(b))
            }

            function h(a, b) {
                da(a, "constant"), s[a] = b, w[a] = b
            }

            function j(a, b) {
                var c = v.get(a + n),
                    d = c.$get;
                c.$get = function() {
                    var a = y.invoke(d, c);
                    return y.invoke(b, null, {
                        $delegate: a
                    })
                }
            }

            function k(a) {
                var b, c, d, e, g = [];
                return f(a, function(a) {
                    if (!r.get(a)) {
                        r.put(a, !0);
                        try {
                            if (u(a))
                                for (b = wc(a), g = g.concat(k(b.requires)).concat(b._runBlocks), c = b._invokeQueue, d = 0, e = c.length; e > d; d++) {
                                    var f = c[d],
                                        h = v.get(f[0]);
                                    h[f[1]].apply(h, f[2])
                                } else x(a) ? g.push(v.invoke(a)) : Ec(a) ? g.push(v.invoke(a)) : ca(a, "module")
                        } catch (i) {
                            throw Ec(a) && (a = a[a.length - 1]), i.message && i.stack && -1 == i.stack.indexOf(i.message) && (i = i.message + "\n" + i.stack), ad("modulerr", "Failed to instantiate module {0} due to:\n{1}", a, i.stack || i.message || i)
                        }
                    }
                }), g
            }

            function l(a, b) {
                function c(c) {
                    if (a.hasOwnProperty(c)) {
                        if (a[c] === m) throw ad("cdep", "Circular dependency found: {0}", c + " <- " + p.join(" <- "));
                        return a[c]
                    }
                    try {
                        return p.unshift(c), a[c] = m, a[c] = b(c)
                    } catch (d) {
                        throw a[c] === m && delete a[c], d
                    } finally {
                        p.shift()
                    }
                }

                function d(a, b, d) {
                    var e, f, g, h = [],
                        i = Ga(a);
                    for (f = 0, e = i.length; e > f; f++) {
                        if (g = i[f], "string" != typeof g) throw ad("itkn", "Incorrect injection token! Expected service name as string, got {0}", g);
                        h.push(d && d.hasOwnProperty(g) ? d[g] : c(g))
                    }
                    return Ec(a) && (a = a[e]), a.apply(b, h)
                }

                function e(a, b) {
                    var c, e, f = function() {};
                    return f.prototype = (Ec(a) ? a[a.length - 1] : a).prototype, c = new f, e = d(a, c, b), t(e) || x(e) ? e : c
                }
                return {
                    invoke: d,
                    instantiate: e,
                    get: c,
                    annotate: Ga,
                    has: function(b) {
                        return s.hasOwnProperty(b + n) || a.hasOwnProperty(b)
                    }
                }
            }
            var m = {},
                n = "Provider",
                p = [],
                r = new Fa([], !0),
                s = {
                    $provide: {
                        provider: b(c),
                        factory: b(d),
                        service: b(e),
                        value: b(g),
                        constant: b(h),
                        decorator: j
                    }
                },
                v = s.$injector = l(s, function() {
                    throw ad("unpr", "Unknown provider: {0}", p.join(" <- "))
                }),
                w = {},
                y = w.$injector = l(w, function(a) {
                    var b = v.get(a + n);
                    return y.invoke(b.$get, b)
                });
            return f(k(a), function(a) {
                y.invoke(a || o)
            }), y
        }

        function Ia() {
            var a = !0;
            this.disableAutoScrolling = function() {
                a = !1
            }, this.$get = ["$window", "$location", "$rootScope", function(b, c, d) {
                function e(a) {
                    var b = null;
                    return f(a, function(a) {
                        b || "a" !== oc(a.nodeName) || (b = a)
                    }), b
                }

                function g() {
                    var a, d = c.hash();
                    d ? (a = h.getElementById(d)) ? a.scrollIntoView() : (a = e(h.getElementsByName(d))) ? a.scrollIntoView() : "top" === d && b.scrollTo(0, 0) : b.scrollTo(0, 0)
                }
                var h = b.document;
                return a && d.$watch(function() {
                    return c.hash()
                }, function() {
                    d.$evalAsync(g)
                }), g
            }]
        }

        function Ja() {
            this.$get = ["$$rAF", "$timeout", function(a, b) {
                return a.supported ? function(b) {
                    return a(b)
                } : function(a) {
                    return b(a, 0, !1)
                }
            }]
        }

        function Ka(a, b, d, e) {
            function g(a) {
                try {
                    a.apply(null, N(arguments, 1))
                } finally {
                    if (s--, 0 === s)
                        for (; t.length;) try {
                            t.pop()()
                        } catch (b) {
                            d.error(b)
                        }
                }
            }

            function h(a, b) {
                ! function c() {
                    f(w, function(a) {
                        a()
                    }), v = b(c, a)
                }()
            }

            function i() {
                x != j.url() && (x = j.url(), f(A, function(a) {
                    a(j.url())
                }))
            }
            var j = this,
                k = b[0],
                l = a.location,
                m = a.history,
                n = a.setTimeout,
                p = a.clearTimeout,
                q = {};
            j.isMock = !1;
            var s = 0,
                t = [];
            j.$$completeOutstandingRequest = g, j.$$incOutstandingRequestCount = function() {
                s++
            }, j.notifyWhenNoOutstandingRequests = function(a) {
                f(w, function(a) {
                    a()
                }), 0 === s ? a() : t.push(a)
            };
            var v, w = [];
            j.addPollFn = function(a) {
                return r(v) && h(100, n), w.push(a), a
            };
            var x = l.href,
                y = b.find("base"),
                z = null;
            j.url = function(b, c) {
                if (l !== a.location && (l = a.location), m !== a.history && (m = a.history), b) {
                    if (x == b) return;
                    var d = x && hb(x) === hb(b);
                    return x = b, !d && e.history ? c ? m.replaceState(null, "", b) : (m.pushState(null, "", b), y.attr("href", y.attr("href"))) : (d || (z = b), c ? l.replace(b) : l.href = b), j
                }
                return z || l.href.replace(/%27/g, "'")
            };
            var A = [],
                B = !1;
            j.onUrlChange = function(b) {
                return B || (e.history && uc(a).on("popstate", i), e.hashchange ? uc(a).on("hashchange", i) : j.addPollFn(i), B = !0), A.push(b), b
            }, j.$$checkUrlChange = i, j.baseHref = function() {
                var a = y.attr("href");
                return a ? a.replace(/^(https?\:)?\/\/[^\/]*/, "") : ""
            };
            var C = {},
                D = "",
                E = j.baseHref();
            j.cookies = function(a, b) {
                var e, f, g, h, i;
                if (!a) {
                    if (k.cookie !== D)
                        for (D = k.cookie, f = D.split("; "), C = {}, h = 0; h < f.length; h++) g = f[h], i = g.indexOf("="), i > 0 && (a = unescape(g.substring(0, i)), C[a] === c && (C[a] = unescape(g.substring(i + 1))));
                    return C
                }
                b === c ? k.cookie = escape(a) + "=;path=" + E + ";expires=Thu, 01 Jan 1970 00:00:00 GMT" : u(b) && (e = (k.cookie = escape(a) + "=" + escape(b) + ";path=" + E).length + 1, e > 4096 && d.warn("Cookie '" + a + "' possibly not set or overflowed because it was too large (" + e + " > 4096 bytes)!"))
            }, j.defer = function(a, b) {
                var c;
                return s++, c = n(function() {
                    delete q[c], g(a)
                }, b || 0), q[c] = !0, c
            }, j.defer.cancel = function(a) {
                return q[a] ? (delete q[a], p(a), g(o), !0) : !1
            }
        }

        function La() {
            this.$get = ["$window", "$log", "$sniffer", "$document", function(a, b, c, d) {
                return new Ka(a, d, b, c)
            }]
        }

        function Ma() {
            this.$get = function() {
                function a(a, c) {
                    function e(a) {
                        a != m && (n ? n == a && (n = a.n) : n = a, f(a.n, a.p), f(a, m), m = a, m.n = null)
                    }

                    function f(a, b) {
                        a != b && (a && (a.p = b), b && (b.n = a))
                    }
                    if (a in b) throw d("$cacheFactory")("iid", "CacheId '{0}' is already taken!", a);
                    var g = 0,
                        h = l({}, c, {
                            id: a
                        }),
                        i = {},
                        j = c && c.capacity || Number.MAX_VALUE,
                        k = {},
                        m = null,
                        n = null;
                    return b[a] = {
                        put: function(a, b) {
                            if (j < Number.MAX_VALUE) {
                                var c = k[a] || (k[a] = {
                                    key: a
                                });
                                e(c)
                            }
                            if (!r(b)) return a in i || g++, i[a] = b, g > j && this.remove(n.key), b
                        },
                        get: function(a) {
                            if (j < Number.MAX_VALUE) {
                                var b = k[a];
                                if (!b) return;
                                e(b)
                            }
                            return i[a]
                        },
                        remove: function(a) {
                            if (j < Number.MAX_VALUE) {
                                var b = k[a];
                                if (!b) return;
                                b == m && (m = b.p), b == n && (n = b.n), f(b.n, b.p), delete k[a]
                            }
                            delete i[a], g--
                        },
                        removeAll: function() {
                            i = {}, g = 0, k = {}, m = n = null
                        },
                        destroy: function() {
                            i = null, h = null, k = null, delete b[a]
                        },
                        info: function() {
                            return l({}, h, {
                                size: g
                            })
                        }
                    }
                }
                var b = {};
                return a.info = function() {
                    var a = {};
                    return f(b, function(b, c) {
                        a[c] = b.info()
                    }), a
                }, a.get = function(a) {
                    return b[a]
                }, a
            }
        }

        function Na() {
            this.$get = ["$cacheFactory", function(a) {
                return a("templates")
            }]
        }

        function Oa(a, d) {
            var e = {},
                g = "Directive",
                h = /^\s*directive\:\s*([\d\w_\-]+)\s+(.*)$/,
                j = /(([\d\w_\-]+)(?:\:([^;]+))?;?)/,
                k = /^(on[a-z]+|formaction)$/;
            this.directive = function m(b, c) {
                return da(b, "directive"), u(b) ? (ba(c, "directiveFactory"), e.hasOwnProperty(b) || (e[b] = [], a.factory(b + g, ["$injector", "$exceptionHandler", function(a, c) {
                    var d = [];
                    return f(e[b], function(e, f) {
                        try {
                            var g = a.invoke(e);
                            x(g) ? g = {
                                compile: q(g)
                            } : !g.compile && g.link && (g.compile = q(g.link)), g.priority = g.priority || 0, g.index = f, g.name = g.name || b, g.require = g.require || g.controller && g.name, g.restrict = g.restrict || "A", d.push(g)
                        } catch (h) {
                            c(h)
                        }
                    }), d
                }])), e[b].push(c)) : f(b, i(m)), this
            }, this.aHrefSanitizationWhitelist = function(a) {
                return s(a) ? (d.aHrefSanitizationWhitelist(a), this) : d.aHrefSanitizationWhitelist()
            }, this.imgSrcSanitizationWhitelist = function(a) {
                return s(a) ? (d.imgSrcSanitizationWhitelist(a), this) : d.imgSrcSanitizationWhitelist()
            }, this.$get = ["$injector", "$interpolate", "$exceptionHandler", "$http", "$templateCache", "$parse", "$controller", "$rootScope", "$document", "$sce", "$animate", "$$sanitizeUri", function(a, d, i, m, o, q, r, s, v, w, y, z) {
                function A(a, b, c, d, e) {
                    a instanceof uc || (a = uc(a)), f(a, function(b, c) {
                        3 == b.nodeType && b.nodeValue.match(/\S+/) && (a[c] = b = uc(b).wrap("<span></span>").parent()[0])
                    });
                    var g = C(a, b, a, c, d, e);
                    return B(a, "ng-scope"),
                        function(b, c, d, e) {
                            ba(b, "scope");
                            var h = c ? Vc.clone.call(a) : a;
                            f(d, function(a, b) {
                                h.data("$" + b + "Controller", a)
                            });
                            for (var i = 0, j = h.length; j > i; i++) {
                                var k = h[i],
                                    l = k.nodeType;
                                (1 === l || 9 === l) && h.eq(i).data("$scope", b)
                            }
                            return c && c(h, b), g && g(b, h, h, e), h
                        }
                }

                function B(a, b) {
                    try {
                        a.addClass(b)
                    } catch (c) {}
                }

                function C(a, b, d, e, f, g) {
                    function h(a, d, e, f) {
                        var g, h, i, j, k, l, m, n, p = d.length,
                            q = new Array(p);
                        for (k = 0; p > k; k++) q[k] = d[k];
                        for (k = 0, m = 0, l = o.length; l > k; m++) i = q[m], g = o[k++], h = o[k++], g ? (g.scope ? (j = a.$new(), uc.data(i, "$scope", j)) : j = a, n = g.transcludeOnThisElement ? D(a, g.transclude, f) : !g.templateOnThisElement && f ? f : !f && b ? D(a, b) : null, g(h, j, i, e, n)) : h && h(a, i.childNodes, c, f)
                    }
                    for (var i, j, k, l, m, n, o = [], p = 0; p < a.length; p++) i = new X, j = E(a[p], [], i, 0 === p ? e : c, f), k = j.length ? H(j, a[p], i, b, d, null, [], [], g) : null, k && k.scope && B(i.$$element, "ng-scope"), m = k && k.terminal || !(l = a[p].childNodes) || !l.length ? null : C(l, k ? (k.transcludeOnThisElement || !k.templateOnThisElement) && k.transclude : b), o.push(k, m), n = n || k || m, g = null;
                    return n ? h : null
                }

                function D(a, b, c) {
                    var d = function(d, e, f) {
                        var g = !1;
                        d || (d = a.$new(), d.$$transcluded = !0, g = !0);
                        var h = b(d, e, f, c);
                        return g && h.on("$destroy", function() {
                            d.$destroy()
                        }), h
                    };
                    return d
                }

                function E(a, b, c, d, e) {
                    var f, g, i = a.nodeType,
                        k = c.$attr;
                    switch (i) {
                        case 1:
                            J(b, Pa(xc(a).toLowerCase()), "E", d, e);
                            for (var l, m, n, o, p, q, r = a.attributes, s = 0, t = r && r.length; t > s; s++) {
                                var v = !1,
                                    w = !1;
                                if (l = r[s], !tc || tc >= 8 || l.specified) {
                                    m = l.name, p = Fc(l.value), o = Pa(m), (q = aa.test(o)) && (m = _(o.substr(6), "-"));
                                    var x = o.replace(/(Start|End)$/, "");
                                    o === x + "Start" && (v = m, w = m.substr(0, m.length - 5) + "end", m = m.substr(0, m.length - 6)), n = Pa(m.toLowerCase()), k[n] = m, (q || !c.hasOwnProperty(n)) && (c[n] = p, Ca(a, n) && (c[n] = !0)), U(a, b, p, n), J(b, n, "A", d, e, v, w)
                                }
                            }
                            if (g = a.className, u(g) && "" !== g)
                                for (; f = j.exec(g);) n = Pa(f[2]), J(b, n, "C", d, e) && (c[n] = Fc(f[3])), g = g.substr(f.index + f[0].length);
                            break;
                        case 3:
                            R(b, a.nodeValue);
                            break;
                        case 8:
                            try {
                                f = h.exec(a.nodeValue), f && (n = Pa(f[1]), J(b, n, "M", d, e) && (c[n] = Fc(f[2])))
                            } catch (y) {}
                    }
                    return b.sort(P), b
                }

                function F(a, b, c) {
                    var d = [],
                        e = 0;
                    if (b && a.hasAttribute && a.hasAttribute(b)) {
                        do {
                            if (!a) throw dd("uterdir", "Unterminated attribute, found '{0}' but no matching '{1}' found.", b, c);
                            1 == a.nodeType && (a.hasAttribute(b) && e++, a.hasAttribute(c) && e--), d.push(a), a = a.nextSibling
                        } while (e > 0)
                    } else d.push(a);
                    return uc(d)
                }

                function G(a, b, c) {
                    return function(d, e, f, g, h) {
                        return e = F(e[0], b, c), a(d, e, f, g, h)
                    }
                }

                function H(a, e, g, h, j, k, l, m, n) {
                    function o(a, b, c, d) {
                        a && (c && (a = G(a, c, d)), a.require = w.require, a.directiveName = y, (P === w || w.$$isolateScope) && (a = W(a, {
                            isolateScope: !0
                        })), l.push(a)), b && (c && (b = G(b, c, d)), b.require = w.require, b.directiveName = y, (P === w || w.$$isolateScope) && (b = W(b, {
                            isolateScope: !0
                        })), m.push(b))
                    }

                    function p(a, b, c, d) {
                        var e, g = "data",
                            h = !1;
                        if (u(b)) {
                            for (;
                                "^" == (e = b.charAt(0)) || "?" == e;) b = b.substr(1), "^" == e && (g = "inheritedData"), h = h || "?" == e;
                            if (e = null, d && "data" === g && (e = d[b]), e = e || c[g]("$" + b + "Controller"), !e && !h) throw dd("ctreq", "Controller '{0}', required by directive '{1}', can't be found!", b, a);
                            return e
                        }
                        return Ec(b) && (e = [], f(b, function(b) {
                            e.push(p(a, b, c, d))
                        })), e
                    }

                    function s(a, b, h, j, k) {
                        function n(a, b) {
                            var d;
                            return arguments.length < 2 && (b = a, a = c), Z && (d = z), k(a, b, d)
                        }
                        var o, s, t, u, v, w, x, y, z = {};
                        if (o = e === h ? g : K(g, new X(uc(h), g.$attr)), s = o.$$element, P) {
                            var A = /^\s*([@=&])(\??)\s*(\w*)\s*$/;
                            x = b.$new(!0), !R || R !== P && R !== P.$$originalDirective ? s.data("$isolateScopeNoTemplate", x) : s.data("$isolateScope", x), B(s, "ng-isolate-scope"), f(P.scope, function(a, c) {
                                var e, f, g, h, i = a.match(A) || [],
                                    j = i[3] || c,
                                    k = "?" == i[2],
                                    l = i[1];
                                switch (x.$$isolateBindings[c] = l + j, l) {
                                    case "@":
                                        o.$observe(j, function(a) {
                                            x[c] = a
                                        }), o.$$observers[j].$$scope = b, o[j] && (x[c] = d(o[j])(b));
                                        break;
                                    case "=":
                                        if (k && !o[j]) return;
                                        f = q(o[j]), h = f.literal ? L : function(a, b) {
                                            return a === b || a !== a && b !== b
                                        }, g = f.assign || function() {
                                            throw e = x[c] = f(b), dd("nonassign", "Expression '{0}' used with directive '{1}' is non-assignable!", o[j], P.name)
                                        }, e = x[c] = f(b), x.$watch(function() {
                                            var a = f(b);
                                            return h(a, x[c]) || (h(a, e) ? g(b, a = x[c]) : x[c] = a), e = a
                                        }, null, f.literal);
                                        break;
                                    case "&":
                                        f = q(o[j]), x[c] = function(a) {
                                            return f(b, a)
                                        };
                                        break;
                                    default:
                                        throw dd("iscp", "Invalid isolate scope definition for directive '{0}'. Definition: {... {1}: '{2}' ...}", P.name, c, a)
                                }
                            })
                        }
                        for (y = k && n, J && f(J, function(a) {
                                var c, d = {
                                    $scope: a === P || a.$$isolateScope ? x : b,
                                    $element: s,
                                    $attrs: o,
                                    $transclude: y
                                };
                                w = a.controller, "@" == w && (w = o[a.name]), c = r(w, d), z[a.name] = c, Z || s.data("$" + a.name + "Controller", c), a.controllerAs && (d.$scope[a.controllerAs] = c)
                            }), t = 0, u = l.length; u > t; t++) try {
                            v = l[t], v(v.isolateScope ? x : b, s, o, v.require && p(v.directiveName, v.require, s, z), y)
                        } catch (C) {
                            i(C, T(s))
                        }
                        var D = b;
                        for (P && (P.template || null === P.templateUrl) && (D = x), a && a(D, h.childNodes, c, k), t = m.length - 1; t >= 0; t--) try {
                            v = m[t], v(v.isolateScope ? x : b, s, o, v.require && p(v.directiveName, v.require, s, z), y)
                        } catch (C) {
                            i(C, T(s))
                        }
                    }
                    n = n || {};
                    for (var v, w, y, z, C, D, H = -Number.MAX_VALUE, J = n.controllerDirectives, P = n.newIsolateScopeDirective, R = n.templateDirective, S = n.nonTlbTranscludeDirective, U = !1, Y = !1, Z = n.hasElementTranscludeDirective, _ = g.$$element = uc(e), aa = k, ba = h, ca = 0, da = a.length; da > ca; ca++) {
                        w = a[ca];
                        var ea = w.$$start,
                            fa = w.$$end;
                        if (ea && (_ = F(e, ea, fa)), z = c, H > w.priority) break;
                        if ((D = w.scope) && (v = v || w, w.templateUrl || (Q("new/isolated scope", P, w, _), t(D) && (P = w))), y = w.name, !w.templateUrl && w.controller && (D = w.controller, J = J || {}, Q("'" + y + "' controller", J[y], w, _), J[y] = w), (D = w.transclude) && (U = !0, w.$$tlb || (Q("transclusion", S, w, _), S = w), "element" == D ? (Z = !0, H = w.priority, z = _, _ = g.$$element = uc(b.createComment(" " + y + ": " + g[y] + " ")), e = _[0], V(j, N(z), e),
                                ba = A(z, h, H, aa && aa.name, {
                                    nonTlbTranscludeDirective: S
                                })) : (z = uc(pa(e)).contents(), _.empty(), ba = A(z, h))), w.template)
                            if (Y = !0, Q("template", R, w, _), R = w, D = x(w.template) ? w.template(_, g) : w.template, D = $(D), w.replace) {
                                if (aa = w, z = la(D) ? [] : uc(Fc(D)), e = z[0], 1 != z.length || 1 !== e.nodeType) throw dd("tplrt", "Template for directive '{0}' must have exactly one root element. {1}", y, "");
                                V(j, _, e);
                                var ga = {
                                        $attr: {}
                                    },
                                    ha = E(e, [], ga),
                                    ia = a.splice(ca + 1, a.length - (ca + 1));
                                P && I(ha), a = a.concat(ha).concat(ia), M(g, ga), da = a.length
                            } else _.html(D);
                        if (w.templateUrl) Y = !0, Q("template", R, w, _), R = w, w.replace && (aa = w), s = O(a.splice(ca, a.length - ca), _, g, j, U && ba, l, m, {
                            controllerDirectives: J,
                            newIsolateScopeDirective: P,
                            templateDirective: R,
                            nonTlbTranscludeDirective: S
                        }), da = a.length;
                        else if (w.compile) try {
                            C = w.compile(_, g, ba), x(C) ? o(null, C, ea, fa) : C && o(C.pre, C.post, ea, fa)
                        } catch (ja) {
                            i(ja, T(_))
                        }
                        w.terminal && (s.terminal = !0, H = Math.max(H, w.priority))
                    }
                    return s.scope = v && v.scope === !0, s.transcludeOnThisElement = U, s.templateOnThisElement = Y, s.transclude = ba, n.hasElementTranscludeDirective = Z, s
                }

                function I(a) {
                    for (var b = 0, c = a.length; c > b; b++) a[b] = n(a[b], {
                        $$isolateScope: !0
                    })
                }

                function J(b, d, f, h, j, k, l) {
                    if (d === j) return null;
                    var m = null;
                    if (e.hasOwnProperty(d))
                        for (var o, p = a.get(d + g), q = 0, r = p.length; r > q; q++) try {
                            o = p[q], (h === c || h > o.priority) && -1 != o.restrict.indexOf(f) && (k && (o = n(o, {
                                $$start: k,
                                $$end: l
                            })), b.push(o), m = o)
                        } catch (s) {
                            i(s)
                        }
                    return m
                }

                function M(a, b) {
                    var c = b.$attr,
                        d = a.$attr,
                        e = a.$$element;
                    f(a, function(d, e) {
                        "$" != e.charAt(0) && (b[e] && b[e] !== d && (d += ("style" === e ? ";" : " ") + b[e]), a.$set(e, d, !0, c[e]))
                    }), f(b, function(b, f) {
                        "class" == f ? (B(e, b), a["class"] = (a["class"] ? a["class"] + " " : "") + b) : "style" == f ? (e.attr("style", e.attr("style") + ";" + b), a.style = (a.style ? a.style + ";" : "") + b) : "$" == f.charAt(0) || a.hasOwnProperty(f) || (a[f] = b, d[f] = c[f])
                    })
                }

                function O(a, b, c, d, e, g, h, i) {
                    var j, k, n = [],
                        p = b[0],
                        q = a.shift(),
                        r = l({}, q, {
                            templateUrl: null,
                            transclude: null,
                            replace: null,
                            $$originalDirective: q
                        }),
                        s = x(q.templateUrl) ? q.templateUrl(b, c) : q.templateUrl;
                    return b.empty(), m.get(w.getTrustedResourceUrl(s), {
                            cache: o
                        }).success(function(l) {
                            var m, o, u, v;
                            if (l = $(l), q.replace) {
                                if (u = la(l) ? [] : uc(Fc(l)), m = u[0], 1 != u.length || 1 !== m.nodeType) throw dd("tplrt", "Template for directive '{0}' must have exactly one root element. {1}", q.name, s);
                                o = {
                                    $attr: {}
                                }, V(d, b, m);
                                var w = E(m, [], o);
                                t(q.scope) && I(w), a = w.concat(a), M(c, o)
                            } else m = p, b.html(l);
                            for (a.unshift(r), j = H(a, m, c, e, b, q, g, h, i), f(d, function(a, c) {
                                    a == m && (d[c] = b[0])
                                }), k = C(b[0].childNodes, e); n.length;) {
                                var x = n.shift(),
                                    y = n.shift(),
                                    z = n.shift(),
                                    A = n.shift(),
                                    F = b[0];
                                if (y !== p) {
                                    var G = y.className;
                                    i.hasElementTranscludeDirective && q.replace || (F = pa(m)), V(z, uc(y), F), B(uc(F), G)
                                }
                                v = j.transcludeOnThisElement ? D(x, j.transclude, A) : A, j(k, x, F, d, v)
                            }
                            n = null
                        }).error(function(a, b, c, d) {
                            throw dd("tpload", "Failed to load template: {0}", d.url)
                        }),
                        function(a, b, c, d, e) {
                            var f = e;
                            n ? (n.push(b), n.push(c), n.push(d), n.push(f)) : (j.transcludeOnThisElement && (f = D(b, j.transclude, e)), j(k, b, c, d, f))
                        }
                }

                function P(a, b) {
                    var c = b.priority - a.priority;
                    return 0 !== c ? c : a.name !== b.name ? a.name < b.name ? -1 : 1 : a.index - b.index
                }

                function Q(a, b, c, d) {
                    if (b) throw dd("multidir", "Multiple directives [{0}, {1}] asking for {2} on: {3}", b.name, c.name, a, T(d))
                }

                function R(a, b) {
                    var c = d(b, !0);
                    c && a.push({
                        priority: 0,
                        compile: function(a) {
                            var b = a.parent(),
                                d = b.length;
                            return d && B(a.parent(), "ng-binding"),
                                function(a, b) {
                                    var e = b.parent(),
                                        f = e.data("$binding") || [];
                                    f.push(c), e.data("$binding", f), d || B(e, "ng-binding"), a.$watch(c, function(a) {
                                        b[0].nodeValue = a
                                    })
                                }
                        }
                    })
                }

                function S(a, b) {
                    if ("srcdoc" == b) return w.HTML;
                    var c = xc(a);
                    return "xlinkHref" == b || "FORM" == c && "action" == b || "IMG" != c && ("src" == b || "ngSrc" == b) ? w.RESOURCE_URL : void 0
                }

                function U(a, b, c, e) {
                    var f = d(c, !0);
                    if (f) {
                        if ("multiple" === e && "SELECT" === xc(a)) throw dd("selmulti", "Binding to the 'multiple' attribute is not supported. Element: {0}", T(a));
                        b.push({
                            priority: 100,
                            compile: function() {
                                return {
                                    pre: function(b, c, g) {
                                        var h = g.$$observers || (g.$$observers = {});
                                        if (k.test(e)) throw dd("nodomevents", "Interpolations for HTML DOM event attributes are disallowed.  Please use the ng- versions (such as ng-click instead of onclick) instead.");
                                        f = d(g[e], !0, S(a, e)), f && (g[e] = f(b), (h[e] || (h[e] = [])).$$inter = !0, (g.$$observers && g.$$observers[e].$$scope || b).$watch(f, function(a, b) {
                                            "class" === e && a != b ? g.$updateClass(a, b) : g.$set(e, a)
                                        }))
                                    }
                                }
                            }
                        })
                    }
                }

                function V(a, c, d) {
                    var e, f, g = c[0],
                        h = c.length,
                        i = g.parentNode;
                    if (a)
                        for (e = 0, f = a.length; f > e; e++)
                            if (a[e] == g) {
                                a[e++] = d;
                                for (var j = e, k = j + h - 1, l = a.length; l > j; j++, k++) l > k ? a[j] = a[k] : delete a[j];
                                a.length -= h - 1;
                                break
                            }
                    i && i.replaceChild(d, g);
                    var m = b.createDocumentFragment();
                    m.appendChild(g), d[uc.expando] = g[uc.expando];
                    for (var n = 1, o = c.length; o > n; n++) {
                        var p = c[n];
                        uc(p).remove(), m.appendChild(p), delete c[n]
                    }
                    c[0] = d, c.length = 1
                }

                function W(a, b) {
                    return l(function() {
                        return a.apply(null, arguments)
                    }, a, b)
                }
                var X = function(a, b) {
                    this.$$element = a, this.$attr = b || {}
                };
                X.prototype = {
                    $normalize: Pa,
                    $addClass: function(a) {
                        a && a.length > 0 && y.addClass(this.$$element, a)
                    },
                    $removeClass: function(a) {
                        a && a.length > 0 && y.removeClass(this.$$element, a)
                    },
                    $updateClass: function(a, b) {
                        var c = Qa(a, b),
                            d = Qa(b, a);
                        0 === c.length ? y.removeClass(this.$$element, d) : 0 === d.length ? y.addClass(this.$$element, c) : y.setClass(this.$$element, c, d)
                    },
                    $set: function(a, b, d, e) {
                        var g, h = Ca(this.$$element[0], a);
                        h && (this.$$element.prop(a, b), e = h), this[a] = b, e ? this.$attr[a] = e : (e = this.$attr[a], e || (this.$attr[a] = e = _(a, "-"))), g = xc(this.$$element), ("A" === g && "href" === a || "IMG" === g && "src" === a) && (this[a] = b = z(b, "src" === a)), d !== !1 && (null === b || b === c ? this.$$element.removeAttr(e) : this.$$element.attr(e, b));
                        var j = this.$$observers;
                        j && f(j[a], function(a) {
                            try {
                                a(b)
                            } catch (c) {
                                i(c)
                            }
                        })
                    },
                    $observe: function(a, b) {
                        var c = this,
                            d = c.$$observers || (c.$$observers = {}),
                            e = d[a] || (d[a] = []);
                        return e.push(b), s.$evalAsync(function() {
                            e.$$inter || b(c[a])
                        }), b
                    }
                };
                var Y = d.startSymbol(),
                    Z = d.endSymbol(),
                    $ = "{{" == Y || "}}" == Z ? p : function(a) {
                        return a.replace(/\{\{/g, Y).replace(/}}/g, Z)
                    },
                    aa = /^ngAttr[A-Z]/;
                return A
            }]
        }

        function Pa(a) {
            return ja(a.replace(ed, ""))
        }

        function Qa(a, b) {
            var c = "",
                d = a.split(/\s+/),
                e = b.split(/\s+/);
            a: for (var f = 0; f < d.length; f++) {
                for (var g = d[f], h = 0; h < e.length; h++)
                    if (g == e[h]) continue a;
                c += (c.length > 0 ? " " : "") + g
            }
            return c
        }

        function Ra() {
            var a = {},
                b = /^(\S+)(\s+as\s+(\w+))?$/;
            this.register = function(b, c) {
                da(b, "controller"), t(b) ? l(a, b) : a[b] = c
            }, this.$get = ["$injector", "$window", function(c, e) {
                return function(f, g) {
                    var h, i, j, k;
                    if (u(f) && (i = f.match(b), j = i[1], k = i[3], f = a.hasOwnProperty(j) ? a[j] : ea(g.$scope, j, !0) || ea(e, j, !0), ca(f, j, !0)), h = c.instantiate(f, g), k) {
                        if (!g || "object" != typeof g.$scope) throw d("$controller")("noscp", "Cannot export controller '{0}' as '{1}'! No $scope object provided via `locals`.", j || f.name, k);
                        g.$scope[k] = h
                    }
                    return h
                }
            }]
        }

        function Sa() {
            this.$get = ["$window", function(a) {
                return uc(a.document)
            }]
        }

        function Ta() {
            this.$get = ["$log", function(a) {
                return function(b, c) {
                    a.error.apply(a, arguments)
                }
            }]
        }

        function Ua(a) {
            var b, c, d, e = {};
            return a ? (f(a.split("\n"), function(a) {
                d = a.indexOf(":"), b = oc(Fc(a.substr(0, d))), c = Fc(a.substr(d + 1)), b && (e[b] = e[b] ? e[b] + ", " + c : c)
            }), e) : e
        }

        function Va(a) {
            var b = t(a) ? a : c;
            return function(c) {
                return b || (b = Ua(a)), c ? b[oc(c)] || null : b
            }
        }

        function Wa(a, b, c) {
            return x(c) ? c(a, b) : (f(c, function(c) {
                a = c(a, b)
            }), a)
        }

        function Xa(a) {
            return a >= 200 && 300 > a
        }

        function Ya() {
            var a = /^\s*(\[|\{[^\{])/,
                b = /[\}\]]\s*$/,
                d = /^\)\]\}',?\n/,
                e = {
                    "Content-Type": "application/json;charset=utf-8"
                },
                g = this.defaults = {
                    transformResponse: [function(c) {
                        return u(c) && (c = c.replace(d, ""), a.test(c) && b.test(c) && (c = R(c))), c
                    }],
                    transformRequest: [function(a) {
                        return !t(a) || B(a) || C(a) ? a : Q(a)
                    }],
                    headers: {
                        common: {
                            Accept: "application/json, text/plain, */*"
                        },
                        post: K(e),
                        put: K(e),
                        patch: K(e)
                    },
                    xsrfCookieName: "XSRF-TOKEN",
                    xsrfHeaderName: "X-XSRF-TOKEN"
                },
                i = this.interceptors = [],
                j = this.responseInterceptors = [];
            this.$get = ["$httpBackend", "$browser", "$cacheFactory", "$rootScope", "$q", "$injector", function(a, b, d, e, k, m) {
                function n(a) {
                    function b(a) {
                        var b = l({}, a, {
                            data: Wa(a.data, a.headers, e.transformResponse)
                        });
                        return Xa(a.status) ? b : k.reject(b)
                    }

                    function d(a) {
                        function b(a) {
                            var b;
                            f(a, function(c, d) {
                                x(c) && (b = c(), null != b ? a[d] = b : delete a[d])
                            })
                        }
                        var c, d, e, h = g.headers,
                            i = l({}, a.headers);
                        h = l({}, h.common, h[oc(a.method)]);
                        a: for (c in h) {
                            d = oc(c);
                            for (e in i)
                                if (oc(e) === d) continue a;
                            i[c] = h[c]
                        }
                        return b(i), i
                    }
                    var e = {
                            method: "get",
                            transformRequest: g.transformRequest,
                            transformResponse: g.transformResponse
                        },
                        h = d(a);
                    l(e, a), e.headers = h, e.method = qc(e.method);
                    var i = function(a) {
                            h = a.headers;
                            var c = Wa(a.data, Va(h), a.transformRequest);
                            return r(c) && f(h, function(a, b) {
                                "content-type" === oc(b) && delete h[b]
                            }), r(a.withCredentials) && !r(g.withCredentials) && (a.withCredentials = g.withCredentials), q(a, c, h).then(b, b)
                        },
                        j = [i, c],
                        m = k.when(e);
                    for (f(z, function(a) {
                            (a.request || a.requestError) && j.unshift(a.request, a.requestError), (a.response || a.responseError) && j.push(a.response, a.responseError)
                        }); j.length;) {
                        var n = j.shift(),
                            o = j.shift();
                        m = m.then(n, o)
                    }
                    return m.success = function(a) {
                        return m.then(function(b) {
                            a(b.data, b.status, b.headers, e)
                        }), m
                    }, m.error = function(a) {
                        return m.then(null, function(b) {
                            a(b.data, b.status, b.headers, e)
                        }), m
                    }, m
                }

                function o(a) {
                    f(arguments, function(a) {
                        n[a] = function(b, c) {
                            return n(l(c || {}, {
                                method: a,
                                url: b
                            }))
                        }
                    })
                }

                function p(a) {
                    f(arguments, function(a) {
                        n[a] = function(b, c, d) {
                            return n(l(d || {}, {
                                method: a,
                                url: b,
                                data: c
                            }))
                        }
                    })
                }

                function q(d, f, h) {
                    function i(a, b, c, d) {
                        m && (Xa(a) ? m.put(u, [a, b, Ua(c), d]) : m.remove(u)), j(b, a, c, d), e.$$phase || e.$apply()
                    }

                    function j(a, b, c, e) {
                        b = Math.max(b, 0), (Xa(b) ? p.resolve : p.reject)({
                            data: a,
                            status: b,
                            headers: Va(c),
                            config: d,
                            statusText: e
                        })
                    }

                    function l() {
                        var a = H(n.pendingRequests, d); - 1 !== a && n.pendingRequests.splice(a, 1)
                    }
                    var m, o, p = k.defer(),
                        q = p.promise,
                        u = v(d.url, d.params);
                    if (n.pendingRequests.push(d), q.then(l, l), !d.cache && !g.cache || d.cache === !1 || "GET" !== d.method && "JSONP" !== d.method || (m = t(d.cache) ? d.cache : t(g.cache) ? g.cache : y), m)
                        if (o = m.get(u), s(o)) {
                            if (D(o)) return o.then(l, l), o;
                            Ec(o) ? j(o[1], o[0], K(o[2]), o[3]) : j(o, 200, {}, "OK")
                        } else m.put(u, q);
                    if (r(o)) {
                        var w = Nb(d.url) ? b.cookies()[d.xsrfCookieName || g.xsrfCookieName] : c;
                        w && (h[d.xsrfHeaderName || g.xsrfHeaderName] = w), a(d.method, u, f, i, h, d.timeout, d.withCredentials, d.responseType)
                    }
                    return q
                }

                function v(a, b) {
                    if (!b) return a;
                    var c = [];
                    return h(b, function(a, b) {
                        null === a || r(a) || (Ec(a) || (a = [a]), f(a, function(a) {
                            t(a) && (a = w(a) ? a.toISOString() : Q(a)), c.push(Y(b) + "=" + Y(a))
                        }))
                    }), c.length > 0 && (a += (-1 == a.indexOf("?") ? "?" : "&") + c.join("&")), a
                }
                var y = d("$http"),
                    z = [];
                return f(i, function(a) {
                    z.unshift(u(a) ? m.get(a) : m.invoke(a))
                }), f(j, function(a, b) {
                    var c = u(a) ? m.get(a) : m.invoke(a);
                    z.splice(b, 0, {
                        response: function(a) {
                            return c(k.when(a))
                        },
                        responseError: function(a) {
                            return c(k.reject(a))
                        }
                    })
                }), n.pendingRequests = [], o("get", "delete", "head", "jsonp"), p("post", "put", "patch"), n.defaults = g, n
            }]
        }

        function Za(b) {
            if (8 >= tc && (!b.match(/^(get|post|head|put|delete|options)$/i) || !a.XMLHttpRequest)) return new a.ActiveXObject("Microsoft.XMLHTTP");
            if (a.XMLHttpRequest) return new a.XMLHttpRequest;
            throw d("$httpBackend")("noxhr", "This browser does not support XMLHttpRequest.")
        }

        function $a() {
            this.$get = ["$browser", "$window", "$document", function(a, b, c) {
                return _a(a, Za, a.defer, b.angular.callbacks, c[0])
            }]
        }

        function _a(a, b, c, d, e) {
            function g(a, b, c) {
                var f = e.createElement("script"),
                    g = null;
                return f.type = "text/javascript", f.src = a, f.async = !0, g = function(a) {
                    Mc(f, "load", g), Mc(f, "error", g), e.body.removeChild(f), f = null;
                    var h = -1,
                        i = "unknown";
                    a && ("load" !== a.type || d[b].called || (a = {
                        type: "error"
                    }), i = a.type, h = "error" === a.type ? 404 : 200), c && c(h, i)
                }, Lc(f, "load", g), Lc(f, "error", g), 8 >= tc && (f.onreadystatechange = function() {
                    u(f.readyState) && /loaded|complete/.test(f.readyState) && (f.onreadystatechange = null, g({
                        type: "load"
                    }))
                }), e.body.appendChild(f), g
            }
            var h = -1;
            return function(e, i, j, k, l, m, n, p) {
                function q() {
                    t = h, v && v(), w && w.abort()
                }

                function r(b, d, e, f, g) {
                    y && c.cancel(y), v = w = null, 0 === d && (d = e ? 200 : "file" == Mb(i).protocol ? 404 : 0), d = 1223 === d ? 204 : d, g = g || "", b(d, e, f, g), a.$$completeOutstandingRequest(o)
                }
                var t;
                if (a.$$incOutstandingRequestCount(), i = i || a.url(), "jsonp" == oc(e)) {
                    var u = "_" + (d.counter++).toString(36);
                    d[u] = function(a) {
                        d[u].data = a, d[u].called = !0
                    };
                    var v = g(i.replace("JSON_CALLBACK", "angular.callbacks." + u), u, function(a, b) {
                        r(k, a, d[u].data, "", b), d[u] = o
                    })
                } else {
                    var w = b(e);
                    if (w.open(e, i, !0), f(l, function(a, b) {
                            s(a) && w.setRequestHeader(b, a)
                        }), w.onreadystatechange = function() {
                            if (w && 4 == w.readyState) {
                                var a = null,
                                    b = null,
                                    c = "";
                                t !== h && (a = w.getAllResponseHeaders(), b = "response" in w ? w.response : w.responseText), t === h && 10 > tc || (c = w.statusText), r(k, t || w.status, b, a, c)
                            }
                        }, n && (w.withCredentials = !0), p) try {
                        w.responseType = p
                    } catch (x) {
                        if ("json" !== p) throw x
                    }
                    w.send(j || null)
                }
                if (m > 0) var y = c(q, m);
                else D(m) && m.then(q)
            }
        }

        function ab() {
            var a = "{{",
                b = "}}";
            this.startSymbol = function(b) {
                return b ? (a = b, this) : a
            }, this.endSymbol = function(a) {
                return a ? (b = a, this) : b
            }, this.$get = ["$parse", "$exceptionHandler", "$sce", function(c, d, e) {
                function f(f, i, j) {
                    for (var k, l, m, n, o = 0, p = [], q = f.length, r = !1, s = []; q > o;) - 1 != (k = f.indexOf(a, o)) && -1 != (l = f.indexOf(b, k + g)) ? (o != k && p.push(f.substring(o, k)), p.push(m = c(n = f.substring(k + g, l))), m.exp = n, o = l + h, r = !0) : (o != q && p.push(f.substring(o)), o = q);
                    if ((q = p.length) || (p.push(""), q = 1), j && p.length > 1) throw fd("noconcat", "Error while interpolating: {0}\nStrict Contextual Escaping disallows interpolations that concatenate multiple expressions when a trusted value is required.  See http://docs.angularjs.org/api/ng.$sce", f);
                    return !i || r ? (s.length = q, m = function(a) {
                        try {
                            for (var b, c = 0, g = q; g > c; c++) {
                                if ("function" == typeof(b = p[c]))
                                    if (b = b(a), b = j ? e.getTrusted(j, b) : e.valueOf(b), null == b) b = "";
                                    else switch (typeof b) {
                                        case "string":
                                            break;
                                        case "number":
                                            b = "" + b;
                                            break;
                                        default:
                                            b = Q(b)
                                    }
                                s[c] = b
                            }
                            return s.join("")
                        } catch (h) {
                            var i = fd("interr", "Can't interpolate: {0}\n{1}", f, h.toString());
                            d(i)
                        }
                    }, m.exp = f, m.parts = p, m) : void 0
                }
                var g = a.length,
                    h = b.length;
                return f.startSymbol = function() {
                    return a
                }, f.endSymbol = function() {
                    return b
                }, f
            }]
        }

        function bb() {
            this.$get = ["$rootScope", "$window", "$q", function(a, b, c) {
                function d(d, f, g, h) {
                    var i = b.setInterval,
                        j = b.clearInterval,
                        k = c.defer(),
                        l = k.promise,
                        m = 0,
                        n = s(h) && !h;
                    return g = s(g) ? g : 0, l.then(null, null, d), l.$$intervalId = i(function() {
                        k.notify(m++), g > 0 && m >= g && (k.resolve(m), j(l.$$intervalId), delete e[l.$$intervalId]), n || a.$apply()
                    }, f), e[l.$$intervalId] = k, l
                }
                var e = {};
                return d.cancel = function(a) {
                    return a && a.$$intervalId in e ? (e[a.$$intervalId].reject("canceled"), b.clearInterval(a.$$intervalId), delete e[a.$$intervalId], !0) : !1
                }, d
            }]
        }

        function cb() {
            this.$get = function() {
                return {
                    id: "en-us",
                    NUMBER_FORMATS: {
                        DECIMAL_SEP: ".",
                        GROUP_SEP: ",",
                        PATTERNS: [{
                            minInt: 1,
                            minFrac: 0,
                            maxFrac: 3,
                            posPre: "",
                            posSuf: "",
                            negPre: "-",
                            negSuf: "",
                            gSize: 3,
                            lgSize: 3
                        }, {
                            minInt: 1,
                            minFrac: 2,
                            maxFrac: 2,
                            posPre: "¤",
                            posSuf: "",
                            negPre: "(¤",
                            negSuf: ")",
                            gSize: 3,
                            lgSize: 3
                        }],
                        CURRENCY_SYM: "$"
                    },
                    DATETIME_FORMATS: {
                        MONTH: "January,February,March,April,May,June,July,August,September,October,November,December".split(","),
                        SHORTMONTH: "Jan,Feb,Mar,Apr,May,Jun,Jul,Aug,Sep,Oct,Nov,Dec".split(","),
                        DAY: "Sunday,Monday,Tuesday,Wednesday,Thursday,Friday,Saturday".split(","),
                        SHORTDAY: "Sun,Mon,Tue,Wed,Thu,Fri,Sat".split(","),
                        AMPMS: ["AM", "PM"],
                        medium: "MMM d, y h:mm:ss a",
                        "short": "M/d/yy h:mm a",
                        fullDate: "EEEE, MMMM d, y",
                        longDate: "MMMM d, y",
                        mediumDate: "MMM d, y",
                        shortDate: "M/d/yy",
                        mediumTime: "h:mm:ss a",
                        shortTime: "h:mm a"
                    },
                    pluralCat: function(a) {
                        return 1 === a ? "one" : "other"
                    }
                }
            }
        }

        function db(a) {
            for (var b = a.split("/"), c = b.length; c--;) b[c] = X(b[c]);
            return b.join("/")
        }

        function eb(a, b, c) {
            var d = Mb(a, c);
            b.$$protocol = d.protocol, b.$$host = d.hostname, b.$$port = m(d.port) || hd[d.protocol] || null
        }

        function fb(a, b, c) {
            var d = "/" !== a.charAt(0);
            d && (a = "/" + a);
            var e = Mb(a, c);
            b.$$path = decodeURIComponent(d && "/" === e.pathname.charAt(0) ? e.pathname.substring(1) : e.pathname), b.$$search = V(e.search), b.$$hash = decodeURIComponent(e.hash), b.$$path && "/" != b.$$path.charAt(0) && (b.$$path = "/" + b.$$path)
        }

        function gb(a, b) {
            return 0 === b.indexOf(a) ? b.substr(a.length) : void 0
        }

        function hb(a) {
            var b = a.indexOf("#");
            return -1 == b ? a : a.substr(0, b)
        }

        function ib(a) {
            return a.substr(0, hb(a).lastIndexOf("/") + 1)
        }

        function jb(a) {
            return a.substring(0, a.indexOf("/", a.indexOf("//") + 2))
        }

        function kb(a, b) {
            this.$$html5 = !0, b = b || "";
            var d = ib(a);
            eb(a, this, a), this.$$parse = function(b) {
                var c = gb(d, b);
                if (!u(c)) throw id("ipthprfx", 'Invalid url "{0}", missing path prefix "{1}".', b, d);
                fb(c, this, a), this.$$path || (this.$$path = "/"), this.$$compose()
            }, this.$$compose = function() {
                var a = W(this.$$search),
                    b = this.$$hash ? "#" + X(this.$$hash) : "";
                this.$$url = db(this.$$path) + (a ? "?" + a : "") + b, this.$$absUrl = d + this.$$url.substr(1)
            }, this.$$parseLinkUrl = function(e, f) {
                var g, h, i;
                return (g = gb(a, e)) !== c ? (h = g, i = (g = gb(b, g)) !== c ? d + (gb("/", g) || g) : a + h) : (g = gb(d, e)) !== c ? i = d + g : d == e + "/" && (i = d), i && this.$$parse(i), !!i
            }
        }

        function lb(a, b) {
            var c = ib(a);
            eb(a, this, a), this.$$parse = function(d) {
                function e(a, b, c) {
                    var d, e = /^\/[A-Z]:(\/.*)/;
                    return 0 === b.indexOf(c) && (b = b.replace(c, "")), e.exec(b) ? a : (d = e.exec(a), d ? d[1] : a)
                }
                var f = gb(a, d) || gb(c, d),
                    g = "#" == f.charAt(0) ? gb(b, f) : this.$$html5 ? f : "";
                if (!u(g)) throw id("ihshprfx", 'Invalid url "{0}", missing hash prefix "{1}".', d, b);
                fb(g, this, a), this.$$path = e(this.$$path, g, a), this.$$compose()
            }, this.$$compose = function() {
                var c = W(this.$$search),
                    d = this.$$hash ? "#" + X(this.$$hash) : "";
                this.$$url = db(this.$$path) + (c ? "?" + c : "") + d, this.$$absUrl = a + (this.$$url ? b + this.$$url : "")
            }, this.$$parseLinkUrl = function(b, c) {
                return hb(a) == hb(b) ? (this.$$parse(b), !0) : !1
            }
        }

        function mb(a, b) {
            this.$$html5 = !0, lb.apply(this, arguments);
            var c = ib(a);
            this.$$parseLinkUrl = function(d, e) {
                var f, g;
                return a == hb(d) ? f = d : (g = gb(c, d)) ? f = a + b + g : c === d + "/" && (f = c), f && this.$$parse(f), !!f
            }, this.$$compose = function() {
                var c = W(this.$$search),
                    d = this.$$hash ? "#" + X(this.$$hash) : "";
                this.$$url = db(this.$$path) + (c ? "?" + c : "") + d, this.$$absUrl = a + b + this.$$url
            }
        }

        function nb(a) {
            return function() {
                return this[a]
            }
        }

        function ob(a, b) {
            return function(c) {
                return r(c) ? this[a] : (this[a] = b(c), this.$$compose(), this)
            }
        }

        function pb() {
            var b = "",
                c = !1;
            this.hashPrefix = function(a) {
                return s(a) ? (b = a, this) : b
            }, this.html5Mode = function(a) {
                return s(a) ? (c = a, this) : c
            }, this.$get = ["$rootScope", "$browser", "$sniffer", "$rootElement", function(d, e, f, g) {
                function h(a) {
                    d.$broadcast("$locationChangeSuccess", i.absUrl(), a)
                }
                var i, j, k, l = e.baseHref(),
                    m = e.url();
                c ? (k = jb(m) + (l || "/"), j = f.history ? kb : mb) : (k = hb(m), j = lb), i = new j(k, "#" + b), i.$$parseLinkUrl(m, m);
                var n = /^\s*(javascript|mailto):/i;
                g.on("click", function(b) {
                    if (!b.ctrlKey && !b.metaKey && 2 != b.which) {
                        for (var c = uc(b.target);
                            "a" !== oc(c[0].nodeName);)
                            if (c[0] === g[0] || !(c = c.parent())[0]) return;
                        var f = c.prop("href"),
                            h = c.attr("href") || c.attr("xlink:href");
                        t(f) && "[object SVGAnimatedString]" === f.toString() && (f = Mb(f.animVal).href), n.test(f) || !f || c.attr("target") || b.isDefaultPrevented() || i.$$parseLinkUrl(f, h) && (b.preventDefault(), i.absUrl() != e.url() && (d.$apply(), a.angular["ff-684208-preventDefault"] = !0))
                    }
                }), i.absUrl() != m && e.url(i.absUrl(), !0), e.onUrlChange(function(a) {
                    i.absUrl() != a && (d.$evalAsync(function() {
                        var b = i.absUrl();
                        i.$$parse(a), d.$broadcast("$locationChangeStart", a, b).defaultPrevented ? (i.$$parse(b), e.url(b)) : h(b)
                    }), d.$$phase || d.$digest())
                });
                var o = 0;
                return d.$watch(function() {
                    var a = e.url(),
                        b = i.$$replace;
                    return o && a == i.absUrl() || (o++, d.$evalAsync(function() {
                        d.$broadcast("$locationChangeStart", i.absUrl(), a).defaultPrevented ? i.$$parse(a) : (e.url(i.absUrl(), b), h(a))
                    })), i.$$replace = !1, o
                }), i
            }]
        }

        function qb() {
            var a = !0,
                b = this;
            this.debugEnabled = function(b) {
                return s(b) ? (a = b, this) : a
            }, this.$get = ["$window", function(c) {
                function d(a) {
                    return a instanceof Error && (a.stack ? a = a.message && -1 === a.stack.indexOf(a.message) ? "Error: " + a.message + "\n" + a.stack : a.stack : a.sourceURL && (a = a.message + "\n" + a.sourceURL + ":" + a.line)), a
                }

                function e(a) {
                    var b = c.console || {},
                        e = b[a] || b.log || o,
                        g = !1;
                    try {
                        g = !!e.apply
                    } catch (h) {}
                    return g ? function() {
                        var a = [];
                        return f(arguments, function(b) {
                            a.push(d(b))
                        }), e.apply(b, a)
                    } : function(a, b) {
                        e(a, null == b ? "" : b)
                    }
                }
                return {
                    log: e("log"),
                    info: e("info"),
                    warn: e("warn"),
                    error: e("error"),
                    debug: function() {
                        var c = e("debug");
                        return function() {
                            a && c.apply(b, arguments)
                        }
                    }()
                }
            }]
        }

        function rb(a, b) {
            if ("__defineGetter__" === a || "__defineSetter__" === a || "__lookupGetter__" === a || "__lookupSetter__" === a || "__proto__" === a) throw kd("isecfld", "Attempting to access a disallowed field in Angular expressions! Expression: {0}", b);
            return a
        }

        function sb(a, b) {
            if (a) {
                if (a.constructor === a) throw kd("isecfn", "Referencing Function in Angular expressions is disallowed! Expression: {0}", b);
                if (a.document && a.location && a.alert && a.setInterval) throw kd("isecwindow", "Referencing the Window in Angular expressions is disallowed! Expression: {0}", b);
                if (a.children && (a.nodeName || a.prop && a.attr && a.find)) throw kd("isecdom", "Referencing DOM nodes in Angular expressions is disallowed! Expression: {0}", b);
                if (a === Object) throw kd("isecobj", "Referencing Object in Angular expressions is disallowed! Expression: {0}", b)
            }
            return a
        }

        function tb(a, b) {
            if (a) {
                if (a.constructor === a) throw kd("isecfn", "Referencing Function in Angular expressions is disallowed! Expression: {0}", b);
                if (a === md || a === nd || od && a === od) throw kd("isecff", "Referencing call, apply or bind in Angular expressions is disallowed! Expression: {0}", b)
            }
        }

        function ub(a, b, d, e, f) {
            sb(a, e), f = f || {};
            for (var g, h = b.split("."), i = 0; h.length > 1; i++) {
                g = rb(h.shift(), e);
                var j = sb(a[g], e);
                j || (j = {}, a[g] = j), a = j, a.then && f.unwrapPromises && (jd(e), "$$v" in a || ! function(a) {
                    a.then(function(b) {
                        a.$$v = b
                    })
                }(a), a.$$v === c && (a.$$v = {}), a = a.$$v)
            }
            return g = rb(h.shift(), e), sb(a[g], e), a[g] = d, d
        }

        function vb(a) {
            return "constructor" == a
        }

        function wb(a, b, d, e, f, g, h) {
            rb(a, g), rb(b, g), rb(d, g), rb(e, g), rb(f, g);
            var i = function(a) {
                    return sb(a, g)
                },
                j = h.expensiveChecks,
                k = j || vb(a) ? i : p,
                l = j || vb(b) ? i : p,
                m = j || vb(d) ? i : p,
                n = j || vb(e) ? i : p,
                o = j || vb(f) ? i : p;
            return h.unwrapPromises ? function(h, i) {
                var j, p = i && i.hasOwnProperty(a) ? i : h;
                return null == p ? p : (p = k(p[a]), p && p.then && (jd(g), "$$v" in p || (j = p, j.$$v = c, j.then(function(a) {
                    j.$$v = k(a)
                })), p = k(p.$$v)), b ? null == p ? c : (p = l(p[b]), p && p.then && (jd(g), "$$v" in p || (j = p, j.$$v = c, j.then(function(a) {
                    j.$$v = l(a)
                })), p = l(p.$$v)), d ? null == p ? c : (p = m(p[d]), p && p.then && (jd(g), "$$v" in p || (j = p, j.$$v = c, j.then(function(a) {
                    j.$$v = m(a)
                })), p = m(p.$$v)), e ? null == p ? c : (p = n(p[e]), p && p.then && (jd(g), "$$v" in p || (j = p, j.$$v = c, j.then(function(a) {
                    j.$$v = n(a)
                })), p = n(p.$$v)), f ? null == p ? c : (p = o(p[f]), p && p.then && (jd(g), "$$v" in p || (j = p, j.$$v = c, j.then(function(a) {
                    j.$$v = o(a)
                })), p = o(p.$$v)), p) : p) : p) : p) : p)
            } : function(g, h) {
                var i = h && h.hasOwnProperty(a) ? h : g;
                return null == i ? i : (i = k(i[a]), b ? null == i ? c : (i = l(i[b]), d ? null == i ? c : (i = m(i[d]), e ? null == i ? c : (i = n(i[e]), f ? null == i ? c : i = o(i[f]) : i) : i) : i) : i)
            }
        }

        function xb(a, b) {
            return function(c, d) {
                return a(c, d, jd, sb, b)
            }
        }

        function yb(a, b, d) {
            var e = b.expensiveChecks,
                g = e ? ud : td;
            if (g.hasOwnProperty(a)) return g[a];
            var h, i = a.split("."),
                j = i.length;
            if (b.csp) h = 6 > j ? wb(i[0], i[1], i[2], i[3], i[4], d, b) : function(a, e) {
                var f, g = 0;
                do f = wb(i[g++], i[g++], i[g++], i[g++], i[g++], d, b)(a, e), e = c, a = f; while (j > g);
                return f
            };
            else {
                var k = "var p;\n";
                e && (k += "s = eso(s, fe);\nl = eso(l, fe);\n");
                var l = e;
                f(i, function(a, c) {
                    rb(a, d);
                    var f = (c ? "s" : '((l&&l.hasOwnProperty("' + a + '"))?l:s)') + '["' + a + '"]',
                        g = e || vb(a);
                    g && (f = "eso(" + f + ", fe)", l = !0), k += "if(s == null) return undefined;\ns=" + f + ";\n", b.unwrapPromises && (k += 'if (s && s.then) {\n pw("' + d.replace(/(["\r\n])/g, "\\$1") + '");\n if (!("$$v" in s)) {\n p=s;\n p.$$v = undefined;\n p.then(function(v) {p.$$v=' + (g ? "eso(v)" : "v") + ";});\n}\n s=" + (g ? "eso(s.$$v)" : "s.$$v") + "\n}\n")
                }), k += "return s;";
                var m = new Function("s", "l", "pw", "eso", "fe", k);
                m.toString = q(k), (l || b.unwrapPromises) && (m = xb(m, d)), h = m
            }
            return "hasOwnProperty" !== a && (g[a] = h), h
        }

        function zb() {
            var a = {},
                b = {},
                c = {
                    csp: !1,
                    unwrapPromises: !1,
                    logPromiseWarnings: !0,
                    expensiveChecks: !1
                };
            this.unwrapPromises = function(a) {
                return s(a) ? (c.unwrapPromises = !!a, this) : c.unwrapPromises
            }, this.logPromiseWarnings = function(a) {
                return s(a) ? (c.logPromiseWarnings = a, this) : c.logPromiseWarnings
            }, this.$get = ["$filter", "$sniffer", "$log", function(d, e, f) {
                c.csp = e.csp;
                var g = {
                    csp: c.csp,
                    unwrapPromises: c.unwrapPromises,
                    logPromiseWarnings: c.logPromiseWarnings,
                    expensiveChecks: !0
                };
                return jd = function(a) {
                        c.logPromiseWarnings && !ld.hasOwnProperty(a) && (ld[a] = !0, f.warn("[$parse] Promise found in the expression `" + a + "`. Automatic unwrapping of promises in Angular expressions is deprecated."))
                    },
                    function(e, f) {
                        var h;
                        switch (typeof e) {
                            case "string":
                                var i = f ? b : a;
                                if (i.hasOwnProperty(e)) return i[e];
                                var j = f ? g : c,
                                    k = new rd(j),
                                    l = new sd(k, d, j);
                                return h = l.parse(e), "hasOwnProperty" !== e && (i[e] = h), h;
                            case "function":
                                return e;
                            default:
                                return o
                        }
                    }
            }]
        }

        function Ab() {
            this.$get = ["$rootScope", "$exceptionHandler", function(a, b) {
                return Bb(function(b) {
                    a.$evalAsync(b)
                }, b)
            }]
        }

        function Bb(a, b) {
            function d(a) {
                return a
            }

            function e(a) {
                return j(a)
            }

            function g(a) {
                var b = h(),
                    c = 0,
                    d = Ec(a) ? [] : {};
                return f(a, function(a, e) {
                    c++, i(a).then(function(a) {
                        d.hasOwnProperty(e) || (d[e] = a, --c || b.resolve(d))
                    }, function(a) {
                        d.hasOwnProperty(e) || b.reject(a)
                    })
                }), 0 === c && b.resolve(d), b.promise
            }
            var h = function() {
                    var f, g, j = [];
                    return g = {
                        resolve: function(b) {
                            if (j) {
                                var d = j;
                                j = c, f = i(b), d.length && a(function() {
                                    for (var a, b = 0, c = d.length; c > b; b++) a = d[b], f.then(a[0], a[1], a[2])
                                })
                            }
                        },
                        reject: function(a) {
                            g.resolve(k(a))
                        },
                        notify: function(b) {
                            if (j) {
                                var c = j;
                                j.length && a(function() {
                                    for (var a, d = 0, e = c.length; e > d; d++) a = c[d], a[2](b)
                                })
                            }
                        },
                        promise: {
                            then: function(a, c, g) {
                                var i = h(),
                                    k = function(c) {
                                        try {
                                            i.resolve((x(a) ? a : d)(c))
                                        } catch (e) {
                                            i.reject(e), b(e)
                                        }
                                    },
                                    l = function(a) {
                                        try {
                                            i.resolve((x(c) ? c : e)(a))
                                        } catch (d) {
                                            i.reject(d), b(d)
                                        }
                                    },
                                    m = function(a) {
                                        try {
                                            i.notify((x(g) ? g : d)(a))
                                        } catch (c) {
                                            b(c)
                                        }
                                    };
                                return j ? j.push([k, l, m]) : f.then(k, l, m), i.promise
                            },
                            "catch": function(a) {
                                return this.then(null, a)
                            },
                            "finally": function(a) {
                                function b(a, b) {
                                    var c = h();
                                    return b ? c.resolve(a) : c.reject(a), c.promise
                                }

                                function c(c, e) {
                                    var f = null;
                                    try {
                                        f = (a || d)()
                                    } catch (g) {
                                        return b(g, !1)
                                    }
                                    return D(f) ? f.then(function() {
                                        return b(c, e)
                                    }, function(a) {
                                        return b(a, !1)
                                    }) : b(c, e)
                                }
                                return this.then(function(a) {
                                    return c(a, !0)
                                }, function(a) {
                                    return c(a, !1)
                                })
                            }
                        }
                    }
                },
                i = function(b) {
                    return D(b) ? b : {
                        then: function(c) {
                            var d = h();
                            return a(function() {
                                d.resolve(c(b))
                            }), d.promise
                        }
                    }
                },
                j = function(a) {
                    var b = h();
                    return b.reject(a), b.promise
                },
                k = function(c) {
                    return {
                        then: function(d, f) {
                            var g = h();
                            return a(function() {
                                try {
                                    g.resolve((x(f) ? f : e)(c))
                                } catch (a) {
                                    g.reject(a), b(a)
                                }
                            }), g.promise
                        }
                    }
                },
                l = function(c, f, g, k) {
                    var l, m = h(),
                        n = function(a) {
                            try {
                                return (x(f) ? f : d)(a)
                            } catch (c) {
                                return b(c), j(c)
                            }
                        },
                        o = function(a) {
                            try {
                                return (x(g) ? g : e)(a)
                            } catch (c) {
                                return b(c), j(c)
                            }
                        },
                        p = function(a) {
                            try {
                                return (x(k) ? k : d)(a)
                            } catch (c) {
                                b(c)
                            }
                        };
                    return a(function() {
                        i(c).then(function(a) {
                            l || (l = !0, m.resolve(i(a).then(n, o, p)))
                        }, function(a) {
                            l || (l = !0, m.resolve(o(a)))
                        }, function(a) {
                            l || m.notify(p(a))
                        })
                    }), m.promise
                };
            return {
                defer: h,
                reject: j,
                when: l,
                all: g
            }
        }

        function Cb() {
            this.$get = ["$window", "$timeout", function(a, b) {
                var c = a.requestAnimationFrame || a.webkitRequestAnimationFrame || a.mozRequestAnimationFrame,
                    d = a.cancelAnimationFrame || a.webkitCancelAnimationFrame || a.mozCancelAnimationFrame || a.webkitCancelRequestAnimationFrame,
                    e = !!c,
                    f = e ? function(a) {
                        var b = c(a);
                        return function() {
                            d(b)
                        }
                    } : function(a) {
                        var c = b(a, 16.66, !1);
                        return function() {
                            b.cancel(c)
                        }
                    };
                return f.supported = e, f
            }]
        }

        function Db() {
            var a = 10,
                b = d("$rootScope"),
                c = null;
            this.digestTtl = function(b) {
                return arguments.length && (a = b), a
            }, this.$get = ["$injector", "$exceptionHandler", "$parse", "$browser", function(d, g, h, i) {
                function k() {
                    this.$id = j(), this.$$phase = this.$parent = this.$$watchers = this.$$nextSibling = this.$$prevSibling = this.$$childHead = this.$$childTail = null, this["this"] = this.$root = this, this.$$destroyed = !1, this.$$asyncQueue = [], this.$$postDigestQueue = [], this.$$listeners = {}, this.$$listenerCount = {}, this.$$isolateBindings = {}
                }

                function l(a) {
                    if (r.$$phase) throw b("inprog", "{0} already in progress", r.$$phase);
                    r.$$phase = a
                }

                function m() {
                    r.$$phase = null
                }

                function n(a, b) {
                    var c = h(a);
                    return ca(c, b), c
                }

                function p(a, b, c) {
                    do a.$$listenerCount[c] -= b, 0 === a.$$listenerCount[c] && delete a.$$listenerCount[c]; while (a = a.$parent)
                }

                function q() {}
                k.prototype = {
                    constructor: k,
                    $new: function(a) {
                        var b;
                        return a ? (b = new k, b.$root = this.$root, b.$$asyncQueue = this.$$asyncQueue, b.$$postDigestQueue = this.$$postDigestQueue) : (this.$$childScopeClass || (this.$$childScopeClass = function() {
                            this.$$watchers = this.$$nextSibling = this.$$childHead = this.$$childTail = null, this.$$listeners = {}, this.$$listenerCount = {}, this.$id = j(), this.$$childScopeClass = null
                        }, this.$$childScopeClass.prototype = this), b = new this.$$childScopeClass), b["this"] = b, b.$parent = this, b.$$prevSibling = this.$$childTail, this.$$childHead ? (this.$$childTail.$$nextSibling = b, this.$$childTail = b) : this.$$childHead = this.$$childTail = b, b
                    },
                    $watch: function(a, b, d) {
                        var e = this,
                            f = n(a, "watch"),
                            g = e.$$watchers,
                            h = {
                                fn: b,
                                last: q,
                                get: f,
                                exp: a,
                                eq: !!d
                            };
                        if (c = null, !x(b)) {
                            var i = n(b || o, "listener");
                            h.fn = function(a, b, c) {
                                i(c)
                            }
                        }
                        if ("string" == typeof a && f.constant) {
                            var j = h.fn;
                            h.fn = function(a, b, c) {
                                j.call(this, a, b, c), I(g, h)
                            }
                        }
                        return g || (g = e.$$watchers = []), g.unshift(h),
                            function() {
                                I(g, h), c = null
                            }
                    },
                    $watchCollection: function(a, b) {
                        function c() {
                            f = m(j);
                            var a, b, c;
                            if (t(f))
                                if (e(f)) {
                                    g !== n && (g = n, q = g.length = 0, l++), a = f.length, q !== a && (l++, g.length = q = a);
                                    for (var d = 0; a > d; d++) c = g[d] !== g[d] && f[d] !== f[d], c || g[d] === f[d] || (l++, g[d] = f[d])
                                } else {
                                    g !== o && (g = o = {}, q = 0, l++), a = 0;
                                    for (b in f) f.hasOwnProperty(b) && (a++, g.hasOwnProperty(b) ? (c = g[b] !== g[b] && f[b] !== f[b], c || g[b] === f[b] || (l++, g[b] = f[b])) : (q++, g[b] = f[b], l++));
                                    if (q > a) {
                                        l++;
                                        for (b in g) g.hasOwnProperty(b) && !f.hasOwnProperty(b) && (q--, delete g[b])
                                    }
                                }
                            else g !== f && (g = f, l++);
                            return l
                        }

                        function d() {
                            if (p ? (p = !1, b(f, f, j)) : b(f, i, j), k)
                                if (t(f))
                                    if (e(f)) {
                                        i = new Array(f.length);
                                        for (var a = 0; a < f.length; a++) i[a] = f[a]
                                    } else {
                                        i = {};
                                        for (var c in f) pc.call(f, c) && (i[c] = f[c])
                                    }
                            else i = f
                        }
                        var f, g, i, j = this,
                            k = b.length > 1,
                            l = 0,
                            m = h(a),
                            n = [],
                            o = {},
                            p = !0,
                            q = 0;
                        return this.$watch(c, d)
                    },
                    $digest: function() {
                        var d, e, f, h, j, k, n, o, p, r, s, t = this.$$asyncQueue,
                            u = this.$$postDigestQueue,
                            v = a,
                            w = this,
                            y = [];
                        l("$digest"), i.$$checkUrlChange(), c = null;
                        do {
                            for (k = !1, o = w; t.length;) {
                                try {
                                    s = t.shift(), s.scope.$eval(s.expression)
                                } catch (z) {
                                    m(), g(z)
                                }
                                c = null
                            }
                            a: do {
                                if (h = o.$$watchers)
                                    for (j = h.length; j--;) try {
                                        if (d = h[j])
                                            if ((e = d.get(o)) === (f = d.last) || (d.eq ? L(e, f) : "number" == typeof e && "number" == typeof f && isNaN(e) && isNaN(f))) {
                                                if (d === c) {
                                                    k = !1;
                                                    break a
                                                }
                                            } else k = !0, c = d, d.last = d.eq ? J(e, null) : e, d.fn(e, f === q ? e : f, o), 5 > v && (p = 4 - v, y[p] || (y[p] = []), r = x(d.exp) ? "fn: " + (d.exp.name || d.exp.toString()) : d.exp, r += "; newVal: " + Q(e) + "; oldVal: " + Q(f), y[p].push(r))
                                    } catch (z) {
                                        m(), g(z)
                                    }
                                if (!(n = o.$$childHead || o !== w && o.$$nextSibling))
                                    for (; o !== w && !(n = o.$$nextSibling);) o = o.$parent
                            } while (o = n);
                            if ((k || t.length) && !v--) throw m(), b("infdig", "{0} $digest() iterations reached. Aborting!\nWatchers fired in the last 5 iterations: {1}", a, Q(y))
                        } while (k || t.length);
                        for (m(); u.length;) try {
                            u.shift()()
                        } catch (z) {
                            g(z)
                        }
                    },
                    $destroy: function() {
                        if (!this.$$destroyed) {
                            var a = this.$parent;
                            this.$broadcast("$destroy"), this.$$destroyed = !0, this !== r && (f(this.$$listenerCount, O(null, p, this)), a.$$childHead == this && (a.$$childHead = this.$$nextSibling), a.$$childTail == this && (a.$$childTail = this.$$prevSibling), this.$$prevSibling && (this.$$prevSibling.$$nextSibling = this.$$nextSibling), this.$$nextSibling && (this.$$nextSibling.$$prevSibling = this.$$prevSibling), this.$parent = this.$$nextSibling = this.$$prevSibling = this.$$childHead = this.$$childTail = this.$root = null, this.$$listeners = {}, this.$$watchers = this.$$asyncQueue = this.$$postDigestQueue = [], this.$destroy = this.$digest = this.$apply = o, this.$on = this.$watch = function() {
                                return o
                            })
                        }
                    },
                    $eval: function(a, b) {
                        return h(a)(this, b)
                    },
                    $evalAsync: function(a) {
                        r.$$phase || r.$$asyncQueue.length || i.defer(function() {
                            r.$$asyncQueue.length && r.$digest()
                        }), this.$$asyncQueue.push({
                            scope: this,
                            expression: a
                        })
                    },
                    $$postDigest: function(a) {
                        this.$$postDigestQueue.push(a)
                    },
                    $apply: function(a) {
                        try {
                            return l("$apply"), this.$eval(a)
                        } catch (b) {
                            g(b)
                        } finally {
                            m();
                            try {
                                r.$digest()
                            } catch (b) {
                                throw g(b), b
                            }
                        }
                    },
                    $on: function(a, b) {
                        var c = this.$$listeners[a];
                        c || (this.$$listeners[a] = c = []), c.push(b);
                        var d = this;
                        do d.$$listenerCount[a] || (d.$$listenerCount[a] = 0), d.$$listenerCount[a]++; while (d = d.$parent);
                        var e = this;
                        return function() {
                            var d = H(c, b); - 1 !== d && (c[d] = null, p(e, 1, a))
                        }
                    },
                    $emit: function(a, b) {
                        var c, d, e, f = [],
                            h = this,
                            i = !1,
                            j = {
                                name: a,
                                targetScope: h,
                                stopPropagation: function() {
                                    i = !0
                                },
                                preventDefault: function() {
                                    j.defaultPrevented = !0
                                },
                                defaultPrevented: !1
                            },
                            k = M([j], arguments, 1);
                        do {
                            for (c = h.$$listeners[a] || f, j.currentScope = h, d = 0, e = c.length; e > d; d++)
                                if (c[d]) try {
                                    c[d].apply(null, k)
                                } catch (l) {
                                    g(l)
                                } else c.splice(d, 1), d--, e--;
                            if (i) return j;
                            h = h.$parent
                        } while (h);
                        return j
                    },
                    $broadcast: function(a, b) {
                        for (var c, d, e, f = this, h = f, i = f, j = {
                                name: a,
                                targetScope: f,
                                preventDefault: function() {
                                    j.defaultPrevented = !0
                                },
                                defaultPrevented: !1
                            }, k = M([j], arguments, 1); h = i;) {
                            for (j.currentScope = h, c = h.$$listeners[a] || [], d = 0, e = c.length; e > d; d++)
                                if (c[d]) try {
                                    c[d].apply(null, k)
                                } catch (l) {
                                    g(l)
                                } else c.splice(d, 1), d--, e--;
                            if (!(i = h.$$listenerCount[a] && h.$$childHead || h !== f && h.$$nextSibling))
                                for (; h !== f && !(i = h.$$nextSibling);) h = h.$parent
                        }
                        return j
                    }
                };
                var r = new k;
                return r
            }]
        }

        function Eb() {
            var a = /^\s*(https?|ftp|mailto|tel|file):/,
                b = /^\s*((https?|ftp|file):|data:image\/)/;
            this.aHrefSanitizationWhitelist = function(b) {
                return s(b) ? (a = b, this) : a
            }, this.imgSrcSanitizationWhitelist = function(a) {
                return s(a) ? (b = a, this) : b
            }, this.$get = function() {
                return function(c, d) {
                    var e, f = d ? b : a;
                    return tc && !(tc >= 8) || (e = Mb(c).href, "" === e || e.match(f)) ? c : "unsafe:" + e
                }
            }
        }

        function Fb(a) {
            return a.replace(/([-()\[\]{}+?*.$\^|,:#<!\\])/g, "\\$1").replace(/\x08/g, "\\x08")
        }

        function Gb(a) {
            if ("self" === a) return a;
            if (u(a)) {
                if (a.indexOf("***") > -1) throw vd("iwcard", "Illegal sequence *** in string matcher.  String: {0}", a);
                return a = Fb(a).replace("\\*\\*", ".*").replace("\\*", "[^:/.?&;]*"), new RegExp("^" + a + "$")
            }
            if (y(a)) return new RegExp("^" + a.source + "$");
            throw vd("imatcher", 'Matchers may only be "self", string patterns or RegExp objects')
        }

        function Hb(a) {
            var b = [];
            return s(a) && f(a, function(a) {
                b.push(Gb(a))
            }), b
        }

        function Ib() {
            this.SCE_CONTEXTS = wd;
            var a = ["self"],
                b = [];
            this.resourceUrlWhitelist = function(b) {
                return arguments.length && (a = Hb(b)), a
            }, this.resourceUrlBlacklist = function(a) {
                return arguments.length && (b = Hb(a)), b
            }, this.$get = ["$injector", function(d) {
                function e(a, b) {
                    return "self" === a ? Nb(b) : !!a.exec(b.href)
                }

                function f(c) {
                    var d, f, g = Mb(c.toString()),
                        h = !1;
                    for (d = 0, f = a.length; f > d; d++)
                        if (e(a[d], g)) {
                            h = !0;
                            break
                        }
                    if (h)
                        for (d = 0, f = b.length; f > d; d++)
                            if (e(b[d], g)) {
                                h = !1;
                                break
                            }
                    return h
                }

                function g(a) {
                    var b = function(a) {
                        this.$$unwrapTrustedValue = function() {
                            return a
                        }
                    };
                    return a && (b.prototype = new a), b.prototype.valueOf = function() {
                        return this.$$unwrapTrustedValue()
                    }, b.prototype.toString = function() {
                        return this.$$unwrapTrustedValue().toString()
                    }, b
                }

                function h(a, b) {
                    var d = m.hasOwnProperty(a) ? m[a] : null;
                    if (!d) throw vd("icontext", "Attempted to trust a value in invalid context. Context: {0}; Value: {1}", a, b);
                    if (null === b || b === c || "" === b) return b;
                    if ("string" != typeof b) throw vd("itype", "Attempted to trust a non-string value in a content requiring a string: Context: {0}", a);
                    return new d(b)
                }

                function i(a) {
                    return a instanceof l ? a.$$unwrapTrustedValue() : a
                }

                function j(a, b) {
                    if (null === b || b === c || "" === b) return b;
                    var d = m.hasOwnProperty(a) ? m[a] : null;
                    if (d && b instanceof d) return b.$$unwrapTrustedValue();
                    if (a === wd.RESOURCE_URL) {
                        if (f(b)) return b;
                        throw vd("insecurl", "Blocked loading resource from url not allowed by $sceDelegate policy.  URL: {0}", b.toString())
                    }
                    if (a === wd.HTML) return k(b);
                    throw vd("unsafe", "Attempting to use an unsafe value in a safe context.")
                }
                var k = function(a) {
                    throw vd("unsafe", "Attempting to use an unsafe value in a safe context.")
                };
                d.has("$sanitize") && (k = d.get("$sanitize"));
                var l = g(),
                    m = {};
                return m[wd.HTML] = g(l), m[wd.CSS] = g(l), m[wd.URL] = g(l), m[wd.JS] = g(l), m[wd.RESOURCE_URL] = g(m[wd.URL]), {
                    trustAs: h,
                    getTrusted: j,
                    valueOf: i
                }
            }]
        }

        function Jb() {
            var a = !0;
            this.enabled = function(b) {
                return arguments.length && (a = !!b), a
            }, this.$get = ["$parse", "$sniffer", "$sceDelegate", function(b, c, d) {
                if (a && c.msie && c.msieDocumentMode < 8) throw vd("iequirks", "Strict Contextual Escaping does not support Internet Explorer version < 9 in quirks mode.  You can fix this by adding the text <!doctype html> to the top of your HTML document.  See http://docs.angularjs.org/api/ng.$sce for more information.");
                var e = K(wd);
                e.isEnabled = function() {
                    return a
                }, e.trustAs = d.trustAs, e.getTrusted = d.getTrusted, e.valueOf = d.valueOf, a || (e.trustAs = e.getTrusted = function(a, b) {
                    return b
                }, e.valueOf = p), e.parseAs = function(a, c) {
                    var d = b(c);
                    return d.literal && d.constant ? d : function(b, c) {
                        return e.getTrusted(a, d(b, c))
                    }
                };
                var g = e.parseAs,
                    h = e.getTrusted,
                    i = e.trustAs;
                return f(wd, function(a, b) {
                    var c = oc(b);
                    e[ja("parse_as_" + c)] = function(b) {
                        return g(a, b)
                    }, e[ja("get_trusted_" + c)] = function(b) {
                        return h(a, b)
                    }, e[ja("trust_as_" + c)] = function(b) {
                        return i(a, b)
                    }
                }), e
            }]
        }

        function Kb() {
            this.$get = ["$window", "$document", function(a, b) {
                var c, d, e = {},
                    f = m((/android (\d+)/.exec(oc((a.navigator || {}).userAgent)) || [])[1]),
                    g = /Boxee/i.test((a.navigator || {}).userAgent),
                    h = b[0] || {},
                    i = h.documentMode,
                    j = /^(Moz|webkit|O|ms)(?=[A-Z])/,
                    k = h.body && h.body.style,
                    l = !1,
                    n = !1;
                if (k) {
                    for (var o in k)
                        if (d = j.exec(o)) {
                            c = d[0], c = c.substr(0, 1).toUpperCase() + c.substr(1);
                            break
                        }
                    c || (c = "WebkitOpacity" in k && "webkit"), l = !!("transition" in k || c + "Transition" in k), n = !!("animation" in k || c + "Animation" in k), !f || l && n || (l = u(h.body.style.webkitTransition), n = u(h.body.style.webkitAnimation))
                }
                return {
                    history: !(!a.history || !a.history.pushState || 4 > f || g),
                    hashchange: "onhashchange" in a && (!i || i > 7),
                    hasEvent: function(a) {
                        if ("input" == a && 9 == tc) return !1;
                        if (r(e[a])) {
                            var b = h.createElement("div");
                            e[a] = "on" + a in b
                        }
                        return e[a]
                    },
                    csp: Gc(),
                    vendorPrefix: c,
                    transitions: l,
                    animations: n,
                    android: f,
                    msie: tc,
                    msieDocumentMode: i
                }
            }]
        }

        function Lb() {
            this.$get = ["$rootScope", "$browser", "$q", "$exceptionHandler", function(a, b, c, d) {
                function e(e, g, h) {
                    var i, j = c.defer(),
                        k = j.promise,
                        l = s(h) && !h;
                    return i = b.defer(function() {
                        try {
                            j.resolve(e())
                        } catch (b) {
                            j.reject(b), d(b)
                        } finally {
                            delete f[k.$$timeoutId]
                        }
                        l || a.$apply()
                    }, g), k.$$timeoutId = i, f[i] = j, k
                }
                var f = {};
                return e.cancel = function(a) {
                    return a && a.$$timeoutId in f ? (f[a.$$timeoutId].reject("canceled"), delete f[a.$$timeoutId], b.defer.cancel(a.$$timeoutId)) : !1
                }, e
            }]
        }

        function Mb(a, b) {
            var c = a;
            return tc && (xd.setAttribute("href", c), c = xd.href), xd.setAttribute("href", c), {
                href: xd.href,
                protocol: xd.protocol ? xd.protocol.replace(/:$/, "") : "",
                host: xd.host,
                search: xd.search ? xd.search.replace(/^\?/, "") : "",
                hash: xd.hash ? xd.hash.replace(/^#/, "") : "",
                hostname: xd.hostname,
                port: xd.port,
                pathname: "/" === xd.pathname.charAt(0) ? xd.pathname : "/" + xd.pathname
            }
        }

        function Nb(a) {
            var b = u(a) ? Mb(a) : a;
            return b.protocol === yd.protocol && b.host === yd.host
        }

        function Ob() {
            this.$get = q(a)
        }

        function Pb(a) {
            function b(d, e) {
                if (t(d)) {
                    var g = {};
                    return f(d, function(a, c) {
                        g[c] = b(c, a)
                    }), g
                }
                return a.factory(d + c, e)
            }
            var c = "Filter";
            this.register = b, this.$get = ["$injector", function(a) {
                return function(b) {
                    return a.get(b + c)
                }
            }], b("currency", Rb), b("date", Zb), b("filter", Qb), b("json", $b), b("limitTo", _b), b("lowercase", Dd), b("number", Sb), b("orderBy", ac), b("uppercase", Ed)
        }

        function Qb() {
            return function(a, b, c) {
                if (!Ec(a)) return a;
                var d = typeof c,
                    e = [];
                e.check = function(a) {
                    for (var b = 0; b < e.length; b++)
                        if (!e[b](a)) return !1;
                    return !0
                }, "function" !== d && (c = "boolean" === d && c ? function(a, b) {
                    return Cc.equals(a, b)
                } : function(a, b) {
                    if (a && b && "object" == typeof a && "object" == typeof b) {
                        for (var d in a)
                            if ("$" !== d.charAt(0) && pc.call(a, d) && c(a[d], b[d])) return !0;
                        return !1
                    }
                    return b = ("" + b).toLowerCase(), ("" + a).toLowerCase().indexOf(b) > -1
                });
                var f = function(a, b) {
                    if ("string" == typeof b && "!" === b.charAt(0)) return !f(a, b.substr(1));
                    switch (typeof a) {
                        case "boolean":
                        case "number":
                        case "string":
                            return c(a, b);
                        case "object":
                            switch (typeof b) {
                                case "object":
                                    return c(a, b);
                                default:
                                    for (var d in a)
                                        if ("$" !== d.charAt(0) && f(a[d], b)) return !0
                            }
                            return !1;
                        case "array":
                            for (var e = 0; e < a.length; e++)
                                if (f(a[e], b)) return !0;
                            return !1;
                        default:
                            return !1
                    }
                };
                switch (typeof b) {
                    case "boolean":
                    case "number":
                    case "string":
                        b = {
                            $: b
                        };
                    case "object":
                        for (var g in b) ! function(a) {
                            "undefined" != typeof b[a] && e.push(function(c) {
                                return f("$" == a ? c : c && c[a], b[a])
                            })
                        }(g);
                        break;
                    case "function":
                        e.push(b);
                        break;
                    default:
                        return a
                }
                for (var h = [], i = 0; i < a.length; i++) {
                    var j = a[i];
                    e.check(j) && h.push(j)
                }
                return h
            }
        }

        function Rb(a) {
            var b = a.NUMBER_FORMATS;
            return function(a, c) {
                return r(c) && (c = b.CURRENCY_SYM), Tb(a, b.PATTERNS[1], b.GROUP_SEP, b.DECIMAL_SEP, 2).replace(/\u00A4/g, c)
            }
        }

        function Sb(a) {
            var b = a.NUMBER_FORMATS;
            return function(a, c) {
                return Tb(a, b.PATTERNS[0], b.GROUP_SEP, b.DECIMAL_SEP, c)
            }
        }

        function Tb(a, b, c, d, e) {
            if (null == a || !isFinite(a) || t(a)) return "";
            var f = 0 > a;
            a = Math.abs(a);
            var g = a + "",
                h = "",
                i = [],
                j = !1;
            if (-1 !== g.indexOf("e")) {
                var k = g.match(/([\d\.]+)e(-?)(\d+)/);
                k && "-" == k[2] && k[3] > e + 1 ? (g = "0", a = 0) : (h = g, j = !0)
            }
            if (j) e > 0 && a > -1 && 1 > a && (h = a.toFixed(e));
            else {
                var l = (g.split(zd)[1] || "").length;
                r(e) && (e = Math.min(Math.max(b.minFrac, l), b.maxFrac)), a = +(Math.round(+(a.toString() + "e" + e)).toString() + "e" + -e), 0 === a && (f = !1);
                var m = ("" + a).split(zd),
                    n = m[0];
                m = m[1] || "";
                var o, p = 0,
                    q = b.lgSize,
                    s = b.gSize;
                if (n.length >= q + s)
                    for (p = n.length - q, o = 0; p > o; o++)(p - o) % s === 0 && 0 !== o && (h += c), h += n.charAt(o);
                for (o = p; o < n.length; o++)(n.length - o) % q === 0 && 0 !== o && (h += c), h += n.charAt(o);
                for (; m.length < e;) m += "0";
                e && "0" !== e && (h += d + m.substr(0, e))
            }
            return i.push(f ? b.negPre : b.posPre), i.push(h), i.push(f ? b.negSuf : b.posSuf), i.join("")
        }

        function Ub(a, b, c) {
            var d = "";
            for (0 > a && (d = "-", a = -a), a = "" + a; a.length < b;) a = "0" + a;
            return c && (a = a.substr(a.length - b)), d + a
        }

        function Vb(a, b, c, d) {
            return c = c || 0,
                function(e) {
                    var f = e["get" + a]();
                    return (c > 0 || f > -c) && (f += c), 0 === f && -12 == c && (f = 12), Ub(f, b, d)
                }
        }

        function Wb(a, b) {
            return function(c, d) {
                var e = c["get" + a](),
                    f = qc(b ? "SHORT" + a : a);
                return d[f][e]
            }
        }

        function Xb(a) {
            var b = -1 * a.getTimezoneOffset(),
                c = b >= 0 ? "+" : "";
            return c += Ub(Math[b > 0 ? "floor" : "ceil"](b / 60), 2) + Ub(Math.abs(b % 60), 2)
        }

        function Yb(a, b) {
            return a.getHours() < 12 ? b.AMPMS[0] : b.AMPMS[1]
        }

        function Zb(a) {
            function b(a) {
                var b;
                if (b = a.match(c)) {
                    var d = new Date(0),
                        e = 0,
                        f = 0,
                        g = b[8] ? d.setUTCFullYear : d.setFullYear,
                        h = b[8] ? d.setUTCHours : d.setHours;
                    b[9] && (e = m(b[9] + b[10]), f = m(b[9] + b[11])), g.call(d, m(b[1]), m(b[2]) - 1, m(b[3]));
                    var i = m(b[4] || 0) - e,
                        j = m(b[5] || 0) - f,
                        k = m(b[6] || 0),
                        l = Math.round(1e3 * parseFloat("0." + (b[7] || 0)));
                    return h.call(d, i, j, k, l), d
                }
                return a
            }
            var c = /^(\d{4})-?(\d\d)-?(\d\d)(?:T(\d\d)(?::?(\d\d)(?::?(\d\d)(?:\.(\d+))?)?)?(Z|([+-])(\d\d):?(\d\d))?)?$/;
            return function(c, d) {
                var e, g, h = "",
                    i = [];
                if (d = d || "mediumDate", d = a.DATETIME_FORMATS[d] || d, u(c) && (c = Cd.test(c) ? m(c) : b(c)), v(c) && (c = new Date(c)), !w(c)) return c;
                for (; d;) g = Bd.exec(d), g ? (i = M(i, g, 1), d = i.pop()) : (i.push(d), d = null);
                return f(i, function(b) {
                    e = Ad[b], h += e ? e(c, a.DATETIME_FORMATS) : b.replace(/(^'|'$)/g, "").replace(/''/g, "'")
                }), h
            }
        }

        function $b() {
            return function(a) {
                return Q(a, !0)
            }
        }

        function _b() {
            return function(a, b) {
                if (!Ec(a) && !u(a)) return a;
                if (b = Math.abs(Number(b)) === 1 / 0 ? Number(b) : m(b), u(a)) return b ? b >= 0 ? a.slice(0, b) : a.slice(b, a.length) : "";
                var c, d, e = [];
                for (b > a.length ? b = a.length : b < -a.length && (b = -a.length), b > 0 ? (c = 0, d = b) : (c = a.length + b, d = a.length); d > c; c++) e.push(a[c]);
                return e
            }
        }

        function ac(a) {
            return function(b, c, d) {
                function f(a, b) {
                    for (var d = 0; d < c.length; d++) {
                        var e = c[d](a, b);
                        if (0 !== e) return e
                    }
                    return 0
                }

                function g(a, b) {
                    return S(b) ? function(b, c) {
                        return a(c, b)
                    } : a
                }

                function h(a, b) {
                    var c = typeof a,
                        d = typeof b;
                    return c == d ? (w(a) && w(b) && (a = a.valueOf(), b = b.valueOf()), "string" == c && (a = a.toLowerCase(), b = b.toLowerCase()), a === b ? 0 : b > a ? -1 : 1) : d > c ? -1 : 1
                }
                return e(b) ? (c = Ec(c) ? c : [c], 0 === c.length && (c = ["+"]), c = F(c, function(b) {
                    var c = !1,
                        d = b || p;
                    if (u(b)) {
                        if (("+" == b.charAt(0) || "-" == b.charAt(0)) && (c = "-" == b.charAt(0), b = b.substring(1)), "" === b) return g(function(a, b) {
                            return h(a, b)
                        }, c);
                        if (d = a(b), d.constant) {
                            var e = d();
                            return g(function(a, b) {
                                return h(a[e], b[e])
                            }, c)
                        }
                    }
                    return g(function(a, b) {
                        return h(d(a), d(b))
                    }, c)
                }), yc.call(b).sort(g(f, d))) : b
            }
        }

        function bc(a) {
            return x(a) && (a = {
                link: a
            }), a.restrict = a.restrict || "AC", q(a)
        }

        function cc(a, b, c, d) {
            function e(b, c) {
                c = c ? "-" + _(c, "-") : "", d.setClass(a, (b ? Rd : Sd) + c, (b ? Sd : Rd) + c)
            }
            var g = this,
                h = a.parent().controller("form") || Hd,
                i = 0,
                j = g.$error = {},
                k = [];
            g.$name = b.name || b.ngForm, g.$dirty = !1, g.$pristine = !0, g.$valid = !0, g.$invalid = !1, h.$addControl(g), a.addClass(Td), e(!0), g.$addControl = function(a) {
                da(a.$name, "input"), k.push(a), a.$name && (g[a.$name] = a)
            }, g.$removeControl = function(a) {
                a.$name && g[a.$name] === a && delete g[a.$name], f(j, function(b, c) {
                    g.$setValidity(c, !0, a)
                }), I(k, a)
            }, g.$setValidity = function(a, b, c) {
                var d = j[a];
                if (b) d && (I(d, c), d.length || (i--, i || (e(b), g.$valid = !0, g.$invalid = !1), j[a] = !1, e(!0, a), h.$setValidity(a, !0, g)));
                else {
                    if (i || e(b), d) {
                        if (G(d, c)) return
                    } else j[a] = d = [], i++, e(!1, a), h.$setValidity(a, !1, g);
                    d.push(c), g.$valid = !1, g.$invalid = !0
                }
            }, g.$setDirty = function() {
                d.removeClass(a, Td), d.addClass(a, Ud), g.$dirty = !0, g.$pristine = !1, h.$setDirty()
            }, g.$setPristine = function() {
                d.removeClass(a, Ud), d.addClass(a, Td), g.$dirty = !1, g.$pristine = !0, f(k, function(a) {
                    a.$setPristine()
                })
            }
        }

        function dc(a, b, d, e) {
            return a.$setValidity(b, d), d ? e : c
        }

        function ec(a, b) {
            var c, d;
            if (b)
                for (c = 0; c < b.length; ++c)
                    if (d = b[c], a[d]) return !0;
            return !1
        }

        function fc(a, b, c, d, e) {
            if (t(e)) {
                a.$$hasNativeValidators = !0;
                var f = function(f) {
                    return a.$error[b] || ec(e, d) || !ec(e, c) ? f : void a.$setValidity(b, !1)
                };
                a.$parsers.push(f)
            }
        }

        function gc(a, b, c, e, f, g) {
            var h = b.prop(nc),
                i = b[0].placeholder,
                j = {},
                k = oc(b[0].type);
            if (e.$$validityState = h, !f.android) {
                var l = !1;
                b.on("compositionstart", function(a) {
                    l = !0
                }), b.on("compositionend", function() {
                    l = !1, n()
                })
            }
            var n = function(d) {
                if (!l) {
                    var f = b.val();
                    if (tc && "input" === (d || j).type && b[0].placeholder !== i) return void(i = b[0].placeholder);
                    "password" !== k && S(c.ngTrim || "T") && (f = Fc(f));
                    var g = h && e.$$hasNativeValidators;
                    (e.$viewValue !== f || "" === f && g) && (a.$root.$$phase ? e.$setViewValue(f) : a.$apply(function() {
                        e.$setViewValue(f)
                    }))
                }
            };
            if (f.hasEvent("input")) b.on("input", n);
            else {
                var o, p = function() {
                    o || (o = g.defer(function() {
                        n(), o = null
                    }))
                };
                b.on("keydown", function(a) {
                    var b = a.keyCode;
                    91 === b || b > 15 && 19 > b || b >= 37 && 40 >= b || p()
                }), f.hasEvent("paste") && b.on("paste cut", p)
            }
            b.on("change", n), e.$render = function() {
                b.val(e.$isEmpty(e.$viewValue) ? "" : e.$viewValue)
            };
            var q, r, s = c.ngPattern;
            if (s) {
                var t = function(a, b) {
                    return dc(e, "pattern", e.$isEmpty(b) || a.test(b), b)
                };
                r = s.match(/^\/(.*)\/([gim]*)$/), r ? (s = new RegExp(r[1], r[2]), q = function(a) {
                    return t(s, a)
                }) : q = function(c) {
                    var e = a.$eval(s);
                    if (!e || !e.test) throw d("ngPattern")("noregexp", "Expected {0} to be a RegExp but was {1}. Element: {2}", s, e, T(b));
                    return t(e, c)
                }, e.$formatters.push(q), e.$parsers.push(q)
            }
            if (c.ngMinlength) {
                var u = m(c.ngMinlength),
                    v = function(a) {
                        return dc(e, "minlength", e.$isEmpty(a) || a.length >= u, a)
                    };
                e.$parsers.push(v), e.$formatters.push(v)
            }
            if (c.ngMaxlength) {
                var w = m(c.ngMaxlength),
                    x = function(a) {
                        return dc(e, "maxlength", e.$isEmpty(a) || a.length <= w, a)
                    };
                e.$parsers.push(x), e.$formatters.push(x)
            }
        }

        function hc(a, b, d, e, f, g) {
            if (gc(a, b, d, e, f, g), e.$parsers.push(function(a) {
                    var b = e.$isEmpty(a);
                    return b || Nd.test(a) ? (e.$setValidity("number", !0), "" === a ? null : b ? a : parseFloat(a)) : (e.$setValidity("number", !1), c)
                }), fc(e, "number", Pd, null, e.$$validityState), e.$formatters.push(function(a) {
                    return e.$isEmpty(a) ? "" : "" + a
                }), d.min) {
                var h = function(a) {
                    var b = parseFloat(d.min);
                    return dc(e, "min", e.$isEmpty(a) || a >= b, a)
                };
                e.$parsers.push(h), e.$formatters.push(h)
            }
            if (d.max) {
                var i = function(a) {
                    var b = parseFloat(d.max);
                    return dc(e, "max", e.$isEmpty(a) || b >= a, a)
                };
                e.$parsers.push(i), e.$formatters.push(i)
            }
            e.$formatters.push(function(a) {
                return dc(e, "number", e.$isEmpty(a) || v(a), a)
            })
        }

        function ic(a, b, c, d, e, f) {
            gc(a, b, c, d, e, f);
            var g = function(a) {
                return dc(d, "url", d.$isEmpty(a) || Ld.test(a), a)
            };
            d.$formatters.push(g), d.$parsers.push(g)
        }

        function jc(a, b, c, d, e, f) {
            gc(a, b, c, d, e, f);
            var g = function(a) {
                return dc(d, "email", d.$isEmpty(a) || Md.test(a), a)
            };
            d.$formatters.push(g), d.$parsers.push(g)
        }

        function kc(a, b, c, d) {
            r(c.name) && b.attr("name", j()), b.on("click", function() {
                b[0].checked && a.$apply(function() {
                    d.$setViewValue(c.value)
                })
            }), d.$render = function() {
                var a = c.value;
                b[0].checked = a == d.$viewValue
            }, c.$observe("value", d.$render)
        }

        function lc(a, b, c, d) {
            var e = c.ngTrueValue,
                f = c.ngFalseValue;
            u(e) || (e = !0), u(f) || (f = !1), b.on("click", function() {
                a.$apply(function() {
                    d.$setViewValue(b[0].checked)
                })
            }), d.$render = function() {
                b[0].checked = d.$viewValue
            }, d.$isEmpty = function(a) {
                return a !== e
            }, d.$formatters.push(function(a) {
                return a === e
            }), d.$parsers.push(function(a) {
                return a ? e : f
            })
        }

        function mc(a, b) {
            return a = "ngClass" + a, ["$animate", function(c) {
                function d(a, b) {
                    var c = [];
                    a: for (var d = 0; d < a.length; d++) {
                        for (var e = a[d], f = 0; f < b.length; f++)
                            if (e == b[f]) continue a;
                        c.push(e)
                    }
                    return c
                }

                function e(a) {
                    if (Ec(a)) return a;
                    if (u(a)) return a.split(" ");
                    if (t(a)) {
                        var b = [];
                        return f(a, function(a, c) {
                            a && (b = b.concat(c.split(" ")))
                        }), b
                    }
                    return a
                }
                return {
                    restrict: "AC",
                    link: function(g, h, i) {
                        function j(a) {
                            var b = l(a, 1);
                            i.$addClass(b)
                        }

                        function k(a) {
                            var b = l(a, -1);
                            i.$removeClass(b)
                        }

                        function l(a, b) {
                            var c = h.data("$classCounts") || {},
                                d = [];
                            return f(a, function(a) {
                                (b > 0 || c[a]) && (c[a] = (c[a] || 0) + b, c[a] === +(b > 0) && d.push(a))
                            }), h.data("$classCounts", c), d.join(" ")
                        }

                        function m(a, b) {
                            var e = d(b, a),
                                f = d(a, b);
                            f = l(f, -1), e = l(e, 1), 0 === e.length ? c.removeClass(h, f) : 0 === f.length ? c.addClass(h, e) : c.setClass(h, e, f)
                        }

                        function n(a) {
                            if (b === !0 || g.$index % 2 === b) {
                                var c = e(a || []);
                                if (o) {
                                    if (!L(a, o)) {
                                        var d = e(o);
                                        m(d, c)
                                    }
                                } else j(c)
                            }
                            o = K(a)
                        }
                        var o;
                        g.$watch(i[a], n, !0), i.$observe("class", function(b) {
                            n(g.$eval(i[a]))
                        }), "ngClass" !== a && g.$watch("$index", function(c, d) {
                            var f = 1 & c;
                            if (f !== (1 & d)) {
                                var h = e(g.$eval(i[a]));
                                f === b ? j(h) : k(h)
                            }
                        })
                    }
                }
            }]
        }
        var nc = "validity",
            oc = function(a) {
                return u(a) ? a.toLowerCase() : a
            },
            pc = Object.prototype.hasOwnProperty,
            qc = function(a) {
                return u(a) ? a.toUpperCase() : a
            },
            rc = function(a) {
                return u(a) ? a.replace(/[A-Z]/g, function(a) {
                    return String.fromCharCode(32 | a.charCodeAt(0))
                }) : a
            },
            sc = function(a) {
                return u(a) ? a.replace(/[a-z]/g, function(a) {
                    return String.fromCharCode(-33 & a.charCodeAt(0))
                }) : a
            };
        "i" !== "I".toLowerCase() && (oc = rc, qc = sc);
        var tc, uc, vc, wc, xc, yc = [].slice,
            zc = [].push,
            Ac = Object.prototype.toString,
            Bc = d("ng"),
            Cc = a.angular || (a.angular = {}),
            Dc = ["0", "0", "0"];
        tc = m((/msie (\d+)/.exec(oc(navigator.userAgent)) || [])[1]), isNaN(tc) && (tc = m((/trident\/.*; rv:(\d+)/.exec(oc(navigator.userAgent)) || [])[1])), o.$inject = [], p.$inject = [];
        var Ec = function() {
                return x(Array.isArray) ? Array.isArray : function(a) {
                    return "[object Array]" === Ac.call(a)
                }
            }(),
            Fc = function() {
                return String.prototype.trim ? function(a) {
                    return u(a) ? a.trim() : a
                } : function(a) {
                    return u(a) ? a.replace(/^\s\s*/, "").replace(/\s\s*$/, "") : a
                }
            }();
        xc = 9 > tc ? function(a) {
            return a = a.nodeName ? a : a[0], a.scopeName && "HTML" != a.scopeName ? qc(a.scopeName + ":" + a.nodeName) : a.nodeName
        } : function(a) {
            return a.nodeName ? a.nodeName : a[0].nodeName
        };
        var Gc = function() {
                if (s(Gc.isActive_)) return Gc.isActive_;
                var a = !(!b.querySelector("[ng-csp]") && !b.querySelector("[data-ng-csp]"));
                if (!a) try {
                    new Function("")
                } catch (c) {
                    a = !0
                }
                return Gc.isActive_ = a
            },
            Hc = /[A-Z]/g,
            Ic = {
                full: "1.2.28",
                major: 1,
                minor: 2,
                dot: 28,
                codeName: "finnish-disembarkation"
            };
        oa.expando = "ng339";
        var Jc = oa.cache = {},
            Kc = 1,
            Lc = a.document.addEventListener ? function(a, b, c) {
                a.addEventListener(b, c, !1)
            } : function(a, b, c) {
                a.attachEvent("on" + b, c)
            },
            Mc = a.document.removeEventListener ? function(a, b, c) {
                a.removeEventListener(b, c, !1)
            } : function(a, b, c) {
                a.detachEvent("on" + b, c)
            },
            Nc = (oa._data = function(a) {
                return this.cache[a[this.expando]] || {}
            }, /([\:\-\_]+(.))/g),
            Oc = /^moz([A-Z])/,
            Pc = d("jqLite"),
            Qc = /^<(\w+)\s*\/?>(?:<\/\1>|)$/,
            Rc = /<|&#?\w+;/,
            Sc = /<([\w:]+)/,
            Tc = /<(?!area|br|col|embed|hr|img|input|link|meta|param)(([\w:]+)[^>]*)\/>/gi,
            Uc = {
                option: [1, '<select multiple="multiple">', "</select>"],
                thead: [1, "<table>", "</table>"],
                col: [2, "<table><colgroup>", "</colgroup></table>"],
                tr: [2, "<table><tbody>", "</tbody></table>"],
                td: [3, "<table><tbody><tr>", "</tr></tbody></table>"],
                _default: [0, "", ""]
            };
        Uc.optgroup = Uc.option, Uc.tbody = Uc.tfoot = Uc.colgroup = Uc.caption = Uc.thead, Uc.th = Uc.td;
        var Vc = oa.prototype = {
                ready: function(c) {
                    function d() {
                        e || (e = !0, c())
                    }
                    var e = !1;
                    "complete" === b.readyState ? setTimeout(d) : (this.on("DOMContentLoaded", d), oa(a).on("load", d))
                },
                toString: function() {
                    var a = [];
                    return f(this, function(b) {
                        a.push("" + b)
                    }), "[" + a.join(", ") + "]"
                },
                eq: function(a) {
                    return uc(a >= 0 ? this[a] : this[this.length + a])
                },
                length: 0,
                push: zc,
                sort: [].sort,
                splice: [].splice
            },
            Wc = {};
        f("multiple,selected,checked,disabled,readOnly,required,open".split(","), function(a) {
            Wc[oc(a)] = a
        });
        var Xc = {};
        f("input,select,option,textarea,button,form,details".split(","), function(a) {
            Xc[qc(a)] = !0
        }), f({
            data: ua,
            removeData: sa
        }, function(a, b) {
            oa[b] = a
        }), f({
            data: ua,
            inheritedData: Aa,
            scope: function(a) {
                return uc.data(a, "$scope") || Aa(a.parentNode || a, ["$isolateScope", "$scope"])
            },
            isolateScope: function(a) {
                return uc.data(a, "$isolateScope") || uc.data(a, "$isolateScopeNoTemplate")
            },
            controller: za,
            injector: function(a) {
                return Aa(a, "$injector")
            },
            removeAttr: function(a, b) {
                a.removeAttribute(b)
            },
            hasClass: va,
            css: function(a, b, d) {
                if (b = ja(b), !s(d)) {
                    var e;
                    return 8 >= tc && (e = a.currentStyle && a.currentStyle[b], "" === e && (e = "auto")), e = e || a.style[b], 8 >= tc && (e = "" === e ? c : e), e
                }
                a.style[b] = d
            },
            attr: function(a, b, d) {
                var e = oc(b);
                if (Wc[e]) {
                    if (!s(d)) return a[b] || (a.attributes.getNamedItem(b) || o).specified ? e : c;
                    d ? (a[b] = !0, a.setAttribute(b, e)) : (a[b] = !1, a.removeAttribute(e))
                } else if (s(d)) a.setAttribute(b, d);
                else if (a.getAttribute) {
                    var f = a.getAttribute(b, 2);
                    return null === f ? c : f
                }
            },
            prop: function(a, b, c) {
                return s(c) ? void(a[b] = c) : a[b]
            },
            text: function() {
                function a(a, c) {
                    var d = b[a.nodeType];
                    return r(c) ? d ? a[d] : "" : void(a[d] = c)
                }
                var b = [];
                return 9 > tc ? (b[1] = "innerText", b[3] = "nodeValue") : b[1] = b[3] = "textContent", a.$dv = "", a
            }(),
            val: function(a, b) {
                if (r(b)) {
                    if ("SELECT" === xc(a) && a.multiple) {
                        var c = [];
                        return f(a.options, function(a) {
                            a.selected && c.push(a.value || a.text)
                        }), 0 === c.length ? null : c
                    }
                    return a.value
                }
                a.value = b
            },
            html: function(a, b) {
                if (r(b)) return a.innerHTML;
                for (var c = 0, d = a.childNodes; c < d.length; c++) qa(d[c]);
                a.innerHTML = b
            },
            empty: Ba
        }, function(a, b) {
            oa.prototype[b] = function(b, d) {
                var e, f, g = this.length;
                if (a !== Ba && (2 == a.length && a !== va && a !== za ? b : d) === c) {
                    if (t(b)) {
                        for (e = 0; g > e; e++)
                            if (a === ua) a(this[e], b);
                            else
                                for (f in b) a(this[e], f, b[f]);
                        return this
                    }
                    for (var h = a.$dv, i = h === c ? Math.min(g, 1) : g, j = 0; i > j; j++) {
                        var k = a(this[j], b, d);
                        h = h ? h + k : k
                    }
                    return h
                }
                for (e = 0; g > e; e++) a(this[e], b, d);
                return this
            }
        }), f({
            removeData: sa,
            dealoc: qa,
            on: function Ee(a, c, d, e) {
                if (s(e)) throw Pc("onargs", "jqLite#on() does not support the `selector` or `eventData` parameters");
                var g = ta(a, "events"),
                    h = ta(a, "handle");
                g || ta(a, "events", g = {}), h || ta(a, "handle", h = Da(a, g)), f(c.split(" "), function(c) {
                    var e = g[c];
                    if (!e) {
                        if ("mouseenter" == c || "mouseleave" == c) {
                            var f = b.body.contains || b.body.compareDocumentPosition ? function(a, b) {
                                var c = 9 === a.nodeType ? a.documentElement : a,
                                    d = b && b.parentNode;
                                return a === d || !(!d || 1 !== d.nodeType || !(c.contains ? c.contains(d) : a.compareDocumentPosition && 16 & a.compareDocumentPosition(d)))
                            } : function(a, b) {
                                if (b)
                                    for (; b = b.parentNode;)
                                        if (b === a) return !0;
                                return !1
                            };
                            g[c] = [];
                            var i = {
                                mouseleave: "mouseout",
                                mouseenter: "mouseover"
                            };
                            Ee(a, i[c], function(a) {
                                var b = this,
                                    d = a.relatedTarget;
                                (!d || d !== b && !f(b, d)) && h(a, c)
                            })
                        } else Lc(a, c, h), g[c] = [];
                        e = g[c]
                    }
                    e.push(d)
                })
            },
            off: ra,
            one: function(a, b, c) {
                a = uc(a), a.on(b, function d() {
                    a.off(b, c), a.off(b, d)
                }), a.on(b, c)
            },
            replaceWith: function(a, b) {
                var c, d = a.parentNode;
                qa(a), f(new oa(b), function(b) {
                    c ? d.insertBefore(b, c.nextSibling) : d.replaceChild(b, a), c = b
                })
            },
            children: function(a) {
                var b = [];
                return f(a.childNodes, function(a) {
                    1 === a.nodeType && b.push(a)
                }), b
            },
            contents: function(a) {
                return a.contentDocument || a.childNodes || []
            },
            append: function(a, b) {
                f(new oa(b), function(b) {
                    (1 === a.nodeType || 11 === a.nodeType) && a.appendChild(b)
                })
            },
            prepend: function(a, b) {
                if (1 === a.nodeType) {
                    var c = a.firstChild;
                    f(new oa(b), function(b) {
                        a.insertBefore(b, c)
                    })
                }
            },
            wrap: function(a, b) {
                b = uc(b)[0];
                var c = a.parentNode;
                c && c.replaceChild(b, a), b.appendChild(a)
            },
            remove: function(a) {
                qa(a);
                var b = a.parentNode;
                b && b.removeChild(a)
            },
            after: function(a, b) {
                var c = a,
                    d = a.parentNode;
                f(new oa(b), function(a) {
                    d.insertBefore(a, c.nextSibling), c = a
                })
            },
            addClass: xa,
            removeClass: wa,
            toggleClass: function(a, b, c) {
                b && f(b.split(" "), function(b) {
                    var d = c;
                    r(d) && (d = !va(a, b)), (d ? xa : wa)(a, b)
                })
            },
            parent: function(a) {
                var b = a.parentNode;
                return b && 11 !== b.nodeType ? b : null
            },
            next: function(a) {
                if (a.nextElementSibling) return a.nextElementSibling;
                for (var b = a.nextSibling; null != b && 1 !== b.nodeType;) b = b.nextSibling;
                return b
            },
            find: function(a, b) {
                return a.getElementsByTagName ? a.getElementsByTagName(b) : []
            },
            clone: pa,
            triggerHandler: function(a, b, c) {
                var d, e, g, h = b.type || b,
                    i = (ta(a, "events") || {})[h];
                i && (d = {
                    preventDefault: function() {
                        this.defaultPrevented = !0
                    },
                    isDefaultPrevented: function() {
                        return this.defaultPrevented === !0
                    },
                    stopPropagation: o,
                    type: h,
                    target: a
                }, b.type && (d = l(d, b)), e = K(i), g = c ? [d].concat(c) : [d], f(e, function(b) {
                    b.apply(a, g)
                }))
            }
        }, function(a, b) {
            oa.prototype[b] = function(b, c, d) {
                for (var e, f = 0; f < this.length; f++) r(e) ? (e = a(this[f], b, c, d), s(e) && (e = uc(e))) : ya(e, a(this[f], b, c, d));
                return s(e) ? e : this
            }, oa.prototype.bind = oa.prototype.on, oa.prototype.unbind = oa.prototype.off
        }), Fa.prototype = {
            put: function(a, b) {
                this[Ea(a, this.nextUid)] = b
            },
            get: function(a) {
                return this[Ea(a, this.nextUid)]
            },
            remove: function(a) {
                var b = this[a = Ea(a, this.nextUid)];
                return delete this[a], b
            }
        };
        var Yc = /^function\s*[^\(]*\(\s*([^\)]*)\)/m,
            Zc = /,/,
            $c = /^\s*(_?)(\S+?)\1\s*$/,
            _c = /((\/\/.*$)|(\/\*[\s\S]*?\*\/))/gm,
            ad = d("$injector"),
            bd = d("$animate"),
            cd = ["$provide", function(a) {
                this.$$selectors = {}, this.register = function(b, c) {
                    var d = b + "-animation";
                    if (b && "." != b.charAt(0)) throw bd("notcsel", "Expecting class selector starting with '.' got '{0}'.", b);
                    this.$$selectors[b.substr(1)] = d, a.factory(d, c)
                }, this.classNameFilter = function(a) {
                    return 1 === arguments.length && (this.$$classNameFilter = a instanceof RegExp ? a : null), this.$$classNameFilter
                }, this.$get = ["$timeout", "$$asyncCallback", function(a, b) {
                    function c(a) {
                        a && b(a)
                    }
                    return {
                        enter: function(a, b, d, e) {
                            d ? d.after(a) : (b && b[0] || (b = d.parent()), b.append(a)), c(e)
                        },
                        leave: function(a, b) {
                            a.remove(), c(b)
                        },
                        move: function(a, b, c, d) {
                            this.enter(a, b, c, d)
                        },
                        addClass: function(a, b, d) {
                            b = u(b) ? b : Ec(b) ? b.join(" ") : "", f(a, function(a) {
                                xa(a, b)
                            }), c(d)
                        },
                        removeClass: function(a, b, d) {
                            b = u(b) ? b : Ec(b) ? b.join(" ") : "", f(a, function(a) {
                                wa(a, b)
                            }), c(d)
                        },
                        setClass: function(a, b, d, e) {
                            f(a, function(a) {
                                xa(a, b), wa(a, d)
                            }), c(e)
                        },
                        enabled: o
                    }
                }]
            }],
            dd = d("$compile");
        Oa.$inject = ["$provide", "$$sanitizeUriProvider"];
        var ed = /^(x[\:\-_]|data[\:\-_])/i,
            fd = d("$interpolate"),
            gd = /^([^\?#]*)(\?([^#]*))?(#(.*))?$/,
            hd = {
                http: 80,
                https: 443,
                ftp: 21
            },
            id = d("$location");
        mb.prototype = lb.prototype = kb.prototype = {
            $$html5: !1,
            $$replace: !1,
            absUrl: nb("$$absUrl"),
            url: function(a) {
                if (r(a)) return this.$$url;
                var b = gd.exec(a);
                return b[1] && this.path(decodeURIComponent(b[1])), (b[2] || b[1]) && this.search(b[3] || ""), this.hash(b[5] || ""), this
            },
            protocol: nb("$$protocol"),
            host: nb("$$host"),
            port: nb("$$port"),
            path: ob("$$path", function(a) {
                return a = null !== a ? a.toString() : "", "/" == a.charAt(0) ? a : "/" + a
            }),
            search: function(a, b) {
                switch (arguments.length) {
                    case 0:
                        return this.$$search;
                    case 1:
                        if (u(a) || v(a)) a = a.toString(), this.$$search = V(a);
                        else {
                            if (!t(a)) throw id("isrcharg", "The first argument of the `$location#search()` call must be a string or an object.");
                            f(a, function(b, c) {
                                null == b && delete a[c]
                            }), this.$$search = a
                        }
                        break;
                    default:
                        r(b) || null === b ? delete this.$$search[a] : this.$$search[a] = b
                }
                return this.$$compose(), this
            },
            hash: ob("$$hash", function(a) {
                return null !== a ? a.toString() : ""
            }),
            replace: function() {
                return this.$$replace = !0, this
            }
        };
        var jd, kd = d("$parse"),
            ld = {},
            md = Function.prototype.call,
            nd = Function.prototype.apply,
            od = Function.prototype.bind,
            pd = {
                "null": function() {
                    return null
                },
                "true": function() {
                    return !0
                },
                "false": function() {
                    return !1
                },
                undefined: o,
                "+": function(a, b, d, e) {
                    return d = d(a, b), e = e(a, b), s(d) ? s(e) ? d + e : d : s(e) ? e : c
                },
                "-": function(a, b, c, d) {
                    return c = c(a, b), d = d(a, b), (s(c) ? c : 0) - (s(d) ? d : 0)
                },
                "*": function(a, b, c, d) {
                    return c(a, b) * d(a, b)
                },
                "/": function(a, b, c, d) {
                    return c(a, b) / d(a, b)
                },
                "%": function(a, b, c, d) {
                    return c(a, b) % d(a, b)
                },
                "^": function(a, b, c, d) {
                    return c(a, b) ^ d(a, b)
                },
                "=": o,
                "===": function(a, b, c, d) {
                    return c(a, b) === d(a, b)
                },
                "!==": function(a, b, c, d) {
                    return c(a, b) !== d(a, b)
                },
                "==": function(a, b, c, d) {
                    return c(a, b) == d(a, b)
                },
                "!=": function(a, b, c, d) {
                    return c(a, b) != d(a, b)
                },
                "<": function(a, b, c, d) {
                    return c(a, b) < d(a, b)
                },
                ">": function(a, b, c, d) {
                    return c(a, b) > d(a, b)
                },
                "<=": function(a, b, c, d) {
                    return c(a, b) <= d(a, b)
                },
                ">=": function(a, b, c, d) {
                    return c(a, b) >= d(a, b)
                },
                "&&": function(a, b, c, d) {
                    return c(a, b) && d(a, b)
                },
                "||": function(a, b, c, d) {
                    return c(a, b) || d(a, b)
                },
                "&": function(a, b, c, d) {
                    return c(a, b) & d(a, b)
                },
                "|": function(a, b, c, d) {
                    return d(a, b)(a, b, c(a, b))
                },
                "!": function(a, b, c) {
                    return !c(a, b)
                }
            },
            qd = {
                n: "\n",
                f: "\f",
                r: "\r",
                t: "	",
                v: "",
                "'": "'",
                '"': '"'
            },
            rd = function(a) {
                this.options = a
            };
        rd.prototype = {
            constructor: rd,
            lex: function(a) {
                for (this.text = a, this.index = 0, this.ch = c, this.lastCh = ":", this.tokens = []; this.index < this.text.length;) {
                    if (this.ch = this.text.charAt(this.index), this.is("\"'")) this.readString(this.ch);
                    else if (this.isNumber(this.ch) || this.is(".") && this.isNumber(this.peek())) this.readNumber();
                    else if (this.isIdent(this.ch)) this.readIdent();
                    else if (this.is("(){}[].,;:?")) this.tokens.push({
                        index: this.index,
                        text: this.ch
                    }), this.index++;
                    else {
                        if (this.isWhitespace(this.ch)) {
                            this.index++;
                            continue
                        }
                        var b = this.ch + this.peek(),
                            d = b + this.peek(2),
                            e = pd[this.ch],
                            f = pd[b],
                            g = pd[d];
                        g ? (this.tokens.push({
                            index: this.index,
                            text: d,
                            fn: g
                        }), this.index += 3) : f ? (this.tokens.push({
                            index: this.index,
                            text: b,
                            fn: f
                        }), this.index += 2) : e ? (this.tokens.push({
                            index: this.index,
                            text: this.ch,
                            fn: e
                        }), this.index += 1) : this.throwError("Unexpected next character ", this.index, this.index + 1)
                    }
                    this.lastCh = this.ch
                }
                return this.tokens
            },
            is: function(a) {
                return -1 !== a.indexOf(this.ch)
            },
            was: function(a) {
                return -1 !== a.indexOf(this.lastCh)
            },
            peek: function(a) {
                var b = a || 1;
                return this.index + b < this.text.length ? this.text.charAt(this.index + b) : !1
            },
            isNumber: function(a) {
                return a >= "0" && "9" >= a
            },
            isWhitespace: function(a) {
                return " " === a || "\r" === a || "	" === a || "\n" === a || "" === a || " " === a
            },
            isIdent: function(a) {
                return a >= "a" && "z" >= a || a >= "A" && "Z" >= a || "_" === a || "$" === a
            },
            isExpOperator: function(a) {
                return "-" === a || "+" === a || this.isNumber(a)
            },
            throwError: function(a, b, c) {
                c = c || this.index;
                var d = s(b) ? "s " + b + "-" + this.index + " [" + this.text.substring(b, c) + "]" : " " + c;
                throw kd("lexerr", "Lexer Error: {0} at column{1} in expression [{2}].", a, d, this.text)
            },
            readNumber: function() {
                for (var a = "", b = this.index; this.index < this.text.length;) {
                    var c = oc(this.text.charAt(this.index));
                    if ("." == c || this.isNumber(c)) a += c;
                    else {
                        var d = this.peek();
                        if ("e" == c && this.isExpOperator(d)) a += c;
                        else if (this.isExpOperator(c) && d && this.isNumber(d) && "e" == a.charAt(a.length - 1)) a += c;
                        else {
                            if (!this.isExpOperator(c) || d && this.isNumber(d) || "e" != a.charAt(a.length - 1)) break;
                            this.throwError("Invalid exponent")
                        }
                    }
                    this.index++
                }
                a = 1 * a, this.tokens.push({
                    index: b,
                    text: a,
                    literal: !0,
                    constant: !0,
                    fn: function() {
                        return a
                    }
                })
            },
            readIdent: function() {
                for (var a, b, c, d, e = this, f = "", g = this.index; this.index < this.text.length && (d = this.text.charAt(this.index), "." === d || this.isIdent(d) || this.isNumber(d));) "." === d && (a = this.index), f += d, this.index++;
                if (a)
                    for (b = this.index; b < this.text.length;) {
                        if (d = this.text.charAt(b), "(" === d) {
                            c = f.substr(a - g + 1), f = f.substr(0, a - g), this.index = b;
                            break
                        }
                        if (!this.isWhitespace(d)) break;
                        b++
                    }
                var h = {
                    index: g,
                    text: f
                };
                if (pd.hasOwnProperty(f)) h.fn = pd[f], h.literal = !0, h.constant = !0;
                else {
                    var i = yb(f, this.options, this.text);
                    h.fn = l(function(a, b) {
                        return i(a, b)
                    }, {
                        assign: function(a, b) {
                            return ub(a, f, b, e.text, e.options)
                        }
                    })
                }
                this.tokens.push(h), c && (this.tokens.push({
                    index: a,
                    text: "."
                }), this.tokens.push({
                    index: a + 1,
                    text: c
                }))
            },
            readString: function(a) {
                var b = this.index;
                this.index++;
                for (var c = "", d = a, e = !1; this.index < this.text.length;) {
                    var f = this.text.charAt(this.index);
                    if (d += f, e) {
                        if ("u" === f) {
                            var g = this.text.substring(this.index + 1, this.index + 5);
                            g.match(/[\da-f]{4}/i) || this.throwError("Invalid unicode escape [\\u" + g + "]"), this.index += 4, c += String.fromCharCode(parseInt(g, 16))
                        } else {
                            var h = qd[f];
                            c += h || f
                        }
                        e = !1
                    } else if ("\\" === f) e = !0;
                    else {
                        if (f === a) return this.index++, void this.tokens.push({
                            index: b,
                            text: d,
                            string: c,
                            literal: !0,
                            constant: !0,
                            fn: function() {
                                return c
                            }
                        });
                        c += f
                    }
                    this.index++
                }
                this.throwError("Unterminated quote", b)
            }
        };
        var sd = function(a, b, c) {
            this.lexer = a, this.$filter = b, this.options = c
        };
        sd.ZERO = l(function() {
            return 0
        }, {
            constant: !0
        }), sd.prototype = {
            constructor: sd,
            parse: function(a) {
                this.text = a, this.tokens = this.lexer.lex(a);
                var b = this.statements();
                return 0 !== this.tokens.length && this.throwError("is an unexpected token", this.tokens[0]), b.literal = !!b.literal, b.constant = !!b.constant, b
            },
            primary: function() {
                var a;
                if (this.expect("(")) a = this.filterChain(), this.consume(")");
                else if (this.expect("[")) a = this.arrayDeclaration();
                else if (this.expect("{")) a = this.object();
                else {
                    var b = this.expect();
                    a = b.fn, a || this.throwError("not a primary expression", b), a.literal = !!b.literal, a.constant = !!b.constant
                }
                for (var c, d; c = this.expect("(", "[", ".");) "(" === c.text ? (a = this.functionCall(a, d), d = null) : "[" === c.text ? (d = a, a = this.objectIndex(a)) : "." === c.text ? (d = a, a = this.fieldAccess(a)) : this.throwError("IMPOSSIBLE");
                return a
            },
            throwError: function(a, b) {
                throw kd("syntax", "Syntax Error: Token '{0}' {1} at column {2} of the expression [{3}] starting at [{4}].", b.text, a, b.index + 1, this.text, this.text.substring(b.index))
            },
            peekToken: function() {
                if (0 === this.tokens.length) throw kd("ueoe", "Unexpected end of expression: {0}", this.text);
                return this.tokens[0]
            },
            peek: function(a, b, c, d) {
                if (this.tokens.length > 0) {
                    var e = this.tokens[0],
                        f = e.text;
                    if (f === a || f === b || f === c || f === d || !a && !b && !c && !d) return e
                }
                return !1
            },
            expect: function(a, b, c, d) {
                var e = this.peek(a, b, c, d);
                return e ? (this.tokens.shift(), e) : !1
            },
            consume: function(a) {
                this.expect(a) || this.throwError("is unexpected, expecting [" + a + "]", this.peek())
            },
            unaryFn: function(a, b) {
                return l(function(c, d) {
                    return a(c, d, b)
                }, {
                    constant: b.constant
                })
            },
            ternaryFn: function(a, b, c) {
                return l(function(d, e) {
                    return a(d, e) ? b(d, e) : c(d, e)
                }, {
                    constant: a.constant && b.constant && c.constant
                })
            },
            binaryFn: function(a, b, c) {
                return l(function(d, e) {
                    return b(d, e, a, c)
                }, {
                    constant: a.constant && c.constant
                })
            },
            statements: function() {
                for (var a = [];;)
                    if (this.tokens.length > 0 && !this.peek("}", ")", ";", "]") && a.push(this.filterChain()), !this.expect(";")) return 1 === a.length ? a[0] : function(b, c) {
                        for (var d, e = 0; e < a.length; e++) {
                            var f = a[e];
                            f && (d = f(b, c))
                        }
                        return d
                    }
            },
            filterChain: function() {
                for (var a, b = this.expression();;) {
                    if (!(a = this.expect("|"))) return b;
                    b = this.binaryFn(b, a.fn, this.filter())
                }
            },
            filter: function() {
                for (var a = this.expect(), b = this.$filter(a.text), c = [];;) {
                    if (!(a = this.expect(":"))) {
                        var d = function(a, d, e) {
                            for (var f = [e], g = 0; g < c.length; g++) f.push(c[g](a, d));
                            return b.apply(a, f)
                        };
                        return function() {
                            return d
                        }
                    }
                    c.push(this.expression())
                }
            },
            expression: function() {
                return this.assignment()
            },
            assignment: function() {
                var a, b, c = this.ternary();
                return (b = this.expect("=")) ? (c.assign || this.throwError("implies assignment but [" + this.text.substring(0, b.index) + "] can not be assigned to", b), a = this.ternary(), function(b, d) {
                    return c.assign(b, a(b, d), d)
                }) : c
            },
            ternary: function() {
                var a, b, c = this.logicalOR();
                return (b = this.expect("?")) ? (a = this.assignment(), (b = this.expect(":")) ? this.ternaryFn(c, a, this.assignment()) : void this.throwError("expected :", b)) : c
            },
            logicalOR: function() {
                for (var a, b = this.logicalAND();;) {
                    if (!(a = this.expect("||"))) return b;
                    b = this.binaryFn(b, a.fn, this.logicalAND())
                }
            },
            logicalAND: function() {
                var a, b = this.equality();
                return (a = this.expect("&&")) && (b = this.binaryFn(b, a.fn, this.logicalAND())), b
            },
            equality: function() {
                var a, b = this.relational();
                return (a = this.expect("==", "!=", "===", "!==")) && (b = this.binaryFn(b, a.fn, this.equality())), b
            },
            relational: function() {
                var a, b = this.additive();
                return (a = this.expect("<", ">", "<=", ">=")) && (b = this.binaryFn(b, a.fn, this.relational())), b
            },
            additive: function() {
                for (var a, b = this.multiplicative(); a = this.expect("+", "-");) b = this.binaryFn(b, a.fn, this.multiplicative());
                return b
            },
            multiplicative: function() {
                for (var a, b = this.unary(); a = this.expect("*", "/", "%");) b = this.binaryFn(b, a.fn, this.unary());
                return b
            },
            unary: function() {
                var a;
                return this.expect("+") ? this.primary() : (a = this.expect("-")) ? this.binaryFn(sd.ZERO, a.fn, this.unary()) : (a = this.expect("!")) ? this.unaryFn(a.fn, this.unary()) : this.primary()
            },
            fieldAccess: function(a) {
                var b = this,
                    c = this.expect().text,
                    d = yb(c, this.options, this.text);
                return l(function(b, c, e) {
                    return d(e || a(b, c))
                }, {
                    assign: function(d, e, f) {
                        var g = a(d, f);
                        return g || a.assign(d, g = {}), ub(g, c, e, b.text, b.options)
                    }
                })
            },
            objectIndex: function(a) {
                var b = this,
                    d = this.expression();
                return this.consume("]"), l(function(e, f) {
                    var g, h, i = a(e, f),
                        j = d(e, f);
                    return rb(j, b.text), i ? (g = sb(i[j], b.text), g && g.then && b.options.unwrapPromises && (h = g, "$$v" in g || (h.$$v = c, h.then(function(a) {
                        h.$$v = a
                    })), g = g.$$v), g) : c
                }, {
                    assign: function(c, e, f) {
                        var g = rb(d(c, f), b.text),
                            h = sb(a(c, f), b.text);
                        return h || a.assign(c, h = {}), h[g] = e
                    }
                })
            },
            functionCall: function(a, b) {
                var c = [];
                if (")" !== this.peekToken().text)
                    do c.push(this.expression()); while (this.expect(","));
                this.consume(")");
                var d = this;
                return function(e, f) {
                    for (var g = [], h = b ? b(e, f) : e, i = 0; i < c.length; i++) g.push(sb(c[i](e, f), d.text));
                    var j = a(e, f, h) || o;
                    sb(h, d.text), tb(j, d.text);
                    var k = j.apply ? j.apply(h, g) : j(g[0], g[1], g[2], g[3], g[4]);
                    return sb(k, d.text)
                }
            },
            arrayDeclaration: function() {
                var a = [],
                    b = !0;
                if ("]" !== this.peekToken().text)
                    do {
                        if (this.peek("]")) break;
                        var c = this.expression();
                        a.push(c), c.constant || (b = !1)
                    } while (this.expect(","));
                return this.consume("]"), l(function(b, c) {
                    for (var d = [], e = 0; e < a.length; e++) d.push(a[e](b, c));
                    return d
                }, {
                    literal: !0,
                    constant: b
                })
            },
            object: function() {
                var a = [],
                    b = !0;
                if ("}" !== this.peekToken().text)
                    do {
                        if (this.peek("}")) break;
                        var c = this.expect(),
                            d = c.string || c.text;
                        this.consume(":");
                        var e = this.expression();
                        a.push({
                            key: d,
                            value: e
                        }), e.constant || (b = !1)
                    } while (this.expect(","));
                return this.consume("}"), l(function(b, c) {
                    for (var d = {}, e = 0; e < a.length; e++) {
                        var f = a[e];
                        d[f.key] = f.value(b, c)
                    }
                    return d
                }, {
                    literal: !0,
                    constant: b
                })
            }
        };
        var td = {},
            ud = {},
            vd = d("$sce"),
            wd = {
                HTML: "html",
                CSS: "css",
                URL: "url",
                RESOURCE_URL: "resourceUrl",
                JS: "js"
            },
            xd = b.createElement("a"),
            yd = Mb(a.location.href, !0);
        Pb.$inject = ["$provide"], Rb.$inject = ["$locale"], Sb.$inject = ["$locale"];
        var zd = ".",
            Ad = {
                yyyy: Vb("FullYear", 4),
                yy: Vb("FullYear", 2, 0, !0),
                y: Vb("FullYear", 1),
                MMMM: Wb("Month"),
                MMM: Wb("Month", !0),
                MM: Vb("Month", 2, 1),
                M: Vb("Month", 1, 1),
                dd: Vb("Date", 2),
                d: Vb("Date", 1),
                HH: Vb("Hours", 2),
                H: Vb("Hours", 1),
                hh: Vb("Hours", 2, -12),
                h: Vb("Hours", 1, -12),
                mm: Vb("Minutes", 2),
                m: Vb("Minutes", 1),
                ss: Vb("Seconds", 2),
                s: Vb("Seconds", 1),
                sss: Vb("Milliseconds", 3),
                EEEE: Wb("Day"),
                EEE: Wb("Day", !0),
                a: Yb,
                Z: Xb
            },
            Bd = /((?:[^yMdHhmsaZE']+)|(?:'(?:[^']|'')*')|(?:E+|y+|M+|d+|H+|h+|m+|s+|a|Z))(.*)/,
            Cd = /^\-?\d+$/;
        Zb.$inject = ["$locale"];
        var Dd = q(oc),
            Ed = q(qc);
        ac.$inject = ["$parse"];
        var Fd = q({
                restrict: "E",
                compile: function(a, c) {
                    return 8 >= tc && (c.href || c.name || c.$set("href", ""), a.append(b.createComment("IE fix"))), c.href || c.xlinkHref || c.name ? void 0 : function(a, b) {
                        var c = "[object SVGAnimatedString]" === Ac.call(b.prop("href")) ? "xlink:href" : "href";
                        b.on("click", function(a) {
                            b.attr(c) || a.preventDefault()
                        })
                    }
                }
            }),
            Gd = {};
        f(Wc, function(a, b) {
            if ("multiple" != a) {
                var c = Pa("ng-" + b);
                Gd[c] = function() {
                    return {
                        priority: 100,
                        link: function(a, d, e) {
                            a.$watch(e[c], function(a) {
                                e.$set(b, !!a)
                            })
                        }
                    }
                }
            }
        }), f(["src", "srcset", "href"], function(a) {
            var b = Pa("ng-" + a);
            Gd[b] = function() {
                return {
                    priority: 99,
                    link: function(c, d, e) {
                        var f = a,
                            g = a;
                        "href" === a && "[object SVGAnimatedString]" === Ac.call(d.prop("href")) && (g = "xlinkHref", e.$attr[g] = "xlink:href", f = null), e.$observe(b, function(b) {
                            return b ? (e.$set(g, b), void(tc && f && d.prop(f, e[g]))) : void("href" === a && e.$set(g, null))
                        })
                    }
                }
            }
        });
        var Hd = {
            $addControl: o,
            $removeControl: o,
            $setValidity: o,
            $setDirty: o,
            $setPristine: o
        };
        cc.$inject = ["$element", "$attrs", "$scope", "$animate"];
        var Id = function(a) {
                return ["$timeout", function(b) {
                    var d = {
                        name: "form",
                        restrict: a ? "EAC" : "E",
                        controller: cc,
                        compile: function() {
                            return {
                                pre: function(a, d, e, f) {
                                    if (!e.action) {
                                        var g = function(a) {
                                            a.preventDefault ? a.preventDefault() : a.returnValue = !1
                                        };
                                        Lc(d[0], "submit", g), d.on("$destroy", function() {
                                            b(function() {
                                                Mc(d[0], "submit", g)
                                            }, 0, !1)
                                        })
                                    }
                                    var h = d.parent().controller("form"),
                                        i = e.name || e.ngForm;
                                    i && ub(a, i, f, i), h && d.on("$destroy", function() {
                                        h.$removeControl(f), i && ub(a, i, c, i), l(f, Hd)
                                    })
                                }
                            }
                        }
                    };
                    return d
                }]
            },
            Jd = Id(),
            Kd = Id(!0),
            Ld = /^(ftp|http|https):\/\/(\w+:{0,1}\w*@)?(\S+)(:[0-9]+)?(\/|\/([\w#!:.?+=&%@!\-\/]))?$/,
            Md = /^[a-z0-9!#$%&'*+\/=?^_`{|}~.-]+@[a-z0-9]([a-z0-9-]*[a-z0-9])?(\.[a-z0-9]([a-z0-9-]*[a-z0-9])?)*$/i,
            Nd = /^\s*(\-|\+)?(\d+|(\d*(\.\d*)))\s*$/,
            Od = {
                text: gc,
                number: hc,
                url: ic,
                email: jc,
                radio: kc,
                checkbox: lc,
                hidden: o,
                button: o,
                submit: o,
                reset: o,
                file: o
            },
            Pd = ["badInput"],
            Qd = ["$browser", "$sniffer", function(a, b) {
                return {
                    restrict: "E",
                    require: "?ngModel",
                    link: function(c, d, e, f) {
                        f && (Od[oc(e.type)] || Od.text)(c, d, e, f, b, a)
                    }
                }
            }],
            Rd = "ng-valid",
            Sd = "ng-invalid",
            Td = "ng-pristine",
            Ud = "ng-dirty",
            Vd = ["$scope", "$exceptionHandler", "$attrs", "$element", "$parse", "$animate", function(a, b, c, e, g, h) {
                function i(a, b) {
                    b = b ? "-" + _(b, "-") : "", h.removeClass(e, (a ? Sd : Rd) + b), h.addClass(e, (a ? Rd : Sd) + b)
                }
                this.$viewValue = Number.NaN, this.$modelValue = Number.NaN, this.$parsers = [], this.$formatters = [], this.$viewChangeListeners = [], this.$pristine = !0, this.$dirty = !1, this.$valid = !0, this.$invalid = !1, this.$name = c.name;
                var j = g(c.ngModel),
                    k = j.assign;
                if (!k) throw d("ngModel")("nonassign", "Expression '{0}' is non-assignable. Element: {1}", c.ngModel, T(e));
                this.$render = o, this.$isEmpty = function(a) {
                    return r(a) || "" === a || null === a || a !== a
                };
                var l = e.inheritedData("$formController") || Hd,
                    m = 0,
                    n = this.$error = {};
                e.addClass(Td), i(!0), this.$setValidity = function(a, b) {
                    n[a] !== !b && (b ? (n[a] && m--, m || (i(!0), this.$valid = !0, this.$invalid = !1)) : (i(!1), this.$invalid = !0, this.$valid = !1, m++), n[a] = !b, i(b, a), l.$setValidity(a, b, this))
                }, this.$setPristine = function() {
                    this.$dirty = !1, this.$pristine = !0, h.removeClass(e, Ud), h.addClass(e, Td)
                }, this.$setViewValue = function(c) {
                    this.$viewValue = c, this.$pristine && (this.$dirty = !0, this.$pristine = !1, h.removeClass(e, Td), h.addClass(e, Ud), l.$setDirty()), f(this.$parsers, function(a) {
                        c = a(c)
                    }), this.$modelValue !== c && (this.$modelValue = c, k(a, c), f(this.$viewChangeListeners, function(a) {
                        try {
                            a()
                        } catch (c) {
                            b(c)
                        }
                    }))
                };
                var p = this;
                a.$watch(function() {
                    var b = j(a);
                    if (p.$modelValue !== b) {
                        var c = p.$formatters,
                            d = c.length;
                        for (p.$modelValue = b; d--;) b = c[d](b);
                        p.$viewValue !== b && (p.$viewValue = b, p.$render())
                    }
                    return b
                })
            }],
            Wd = function() {
                return {
                    require: ["ngModel", "^?form"],
                    controller: Vd,
                    link: function(a, b, c, d) {
                        var e = d[0],
                            f = d[1] || Hd;
                        f.$addControl(e), a.$on("$destroy", function() {
                            f.$removeControl(e)
                        })
                    }
                }
            },
            Xd = q({
                require: "ngModel",
                link: function(a, b, c, d) {
                    d.$viewChangeListeners.push(function() {
                        a.$eval(c.ngChange)
                    })
                }
            }),
            Yd = function() {
                return {
                    require: "?ngModel",
                    link: function(a, b, c, d) {
                        if (d) {
                            c.required = !0;
                            var e = function(a) {
                                return c.required && d.$isEmpty(a) ? void d.$setValidity("required", !1) : (d.$setValidity("required", !0), a)
                            };
                            d.$formatters.push(e), d.$parsers.unshift(e), c.$observe("required", function() {
                                e(d.$viewValue)
                            })
                        }
                    }
                }
            },
            Zd = function() {
                return {
                    require: "ngModel",
                    link: function(a, b, d, e) {
                        var g = /\/(.*)\//.exec(d.ngList),
                            h = g && new RegExp(g[1]) || d.ngList || ",",
                            i = function(a) {
                                if (!r(a)) {
                                    var b = [];
                                    return a && f(a.split(h), function(a) {
                                        a && b.push(Fc(a))
                                    }), b
                                }
                            };
                        e.$parsers.push(i), e.$formatters.push(function(a) {
                            return Ec(a) ? a.join(", ") : c
                        }), e.$isEmpty = function(a) {
                            return !a || !a.length
                        }
                    }
                }
            },
            $d = /^(true|false|\d+)$/,
            _d = function() {
                return {
                    priority: 100,
                    compile: function(a, b) {
                        return $d.test(b.ngValue) ? function(a, b, c) {
                            c.$set("value", a.$eval(c.ngValue))
                        } : function(a, b, c) {
                            a.$watch(c.ngValue, function(a) {
                                c.$set("value", a)
                            })
                        }
                    }
                }
            },
            ae = bc({
                compile: function(a) {
                    return a.addClass("ng-binding"),
                        function(a, b, d) {
                            b.data("$binding", d.ngBind), a.$watch(d.ngBind, function(a) {
                                b.text(a == c ? "" : a)
                            })
                        }
                }
            }),
            be = ["$interpolate", function(a) {
                return function(b, c, d) {
                    var e = a(c.attr(d.$attr.ngBindTemplate));
                    c.addClass("ng-binding").data("$binding", e), d.$observe("ngBindTemplate", function(a) {
                        c.text(a)
                    })
                }
            }],
            ce = ["$sce", "$parse", function(a, b) {
                return {
                    compile: function(c) {
                        return c.addClass("ng-binding"),
                            function(c, d, e) {
                                function f() {
                                    return (g(c) || "").toString()
                                }
                                d.data("$binding", e.ngBindHtml);
                                var g = b(e.ngBindHtml);
                                c.$watch(f, function(b) {
                                    d.html(a.getTrustedHtml(g(c)) || "")
                                })
                            }
                    }
                }
            }],
            de = mc("", !0),
            ee = mc("Odd", 0),
            fe = mc("Even", 1),
            ge = bc({
                compile: function(a, b) {
                    b.$set("ngCloak", c), a.removeClass("ng-cloak")
                }
            }),
            he = [function() {
                return {
                    scope: !0,
                    controller: "@",
                    priority: 500
                }
            }],
            ie = {},
            je = {
                blur: !0,
                focus: !0
            };
        f("click dblclick mousedown mouseup mouseover mouseout mousemove mouseenter mouseleave keydown keyup keypress submit focus blur copy cut paste".split(" "), function(a) {
            var b = Pa("ng-" + a);
            ie[b] = ["$parse", "$rootScope", function(c, d) {
                return {
                    compile: function(e, f) {
                        var g = c(f[b], !0);
                        return function(b, c) {
                            c.on(a, function(c) {
                                var e = function() {
                                    g(b, {
                                        $event: c
                                    })
                                };
                                je[a] && d.$$phase ? b.$evalAsync(e) : b.$apply(e)
                            })
                        }
                    }
                }
            }]
        });
        var ke = ["$animate", function(a) {
                return {
                    transclude: "element",
                    priority: 600,
                    terminal: !0,
                    restrict: "A",
                    $$tlb: !0,
                    link: function(c, d, e, f, g) {
                        var h, i, j;
                        c.$watch(e.ngIf, function(f) {
                            S(f) ? i || (i = c.$new(), g(i, function(c) {
                                c[c.length++] = b.createComment(" end ngIf: " + e.ngIf + " "), h = {
                                    clone: c
                                }, a.enter(c, d.parent(), d)
                            })) : (j && (j.remove(), j = null), i && (i.$destroy(), i = null), h && (j = fa(h.clone), a.leave(j, function() {
                                j = null
                            }), h = null))
                        })
                    }
                }
            }],
            le = ["$http", "$templateCache", "$anchorScroll", "$animate", "$sce", function(a, b, c, d, e) {
                return {
                    restrict: "ECA",
                    priority: 400,
                    terminal: !0,
                    transclude: "element",
                    controller: Cc.noop,
                    compile: function(f, g) {
                        var h = g.ngInclude || g.src,
                            i = g.onload || "",
                            j = g.autoscroll;
                        return function(f, g, k, l, m) {
                            var n, o, p, q = 0,
                                r = function() {
                                    o && (o.remove(), o = null), n && (n.$destroy(), n = null), p && (d.leave(p, function() {
                                        o = null
                                    }), o = p, p = null)
                                };
                            f.$watch(e.parseAsResourceUrl(h), function(e) {
                                var h = function() {
                                        !s(j) || j && !f.$eval(j) || c()
                                    },
                                    k = ++q;
                                e ? (a.get(e, {
                                    cache: b
                                }).success(function(a) {
                                    if (k === q) {
                                        var b = f.$new();
                                        l.template = a;
                                        var c = m(b, function(a) {
                                            r(), d.enter(a, null, g, h)
                                        });
                                        n = b, p = c, n.$emit("$includeContentLoaded"), f.$eval(i)
                                    }
                                }).error(function() {
                                    k === q && r()
                                }), f.$emit("$includeContentRequested")) : (r(), l.template = null)
                            })
                        }
                    }
                }
            }],
            me = ["$compile", function(a) {
                return {
                    restrict: "ECA",
                    priority: -400,
                    require: "ngInclude",
                    link: function(b, c, d, e) {
                        c.html(e.template), a(c.contents())(b)
                    }
                }
            }],
            ne = bc({
                priority: 450,
                compile: function() {
                    return {
                        pre: function(a, b, c) {
                            a.$eval(c.ngInit)
                        }
                    }
                }
            }),
            oe = bc({
                terminal: !0,
                priority: 1e3
            }),
            pe = ["$locale", "$interpolate", function(a, b) {
                var c = /{}/g;
                return {
                    restrict: "EA",
                    link: function(d, e, g) {
                        var h = g.count,
                            i = g.$attr.when && e.attr(g.$attr.when),
                            j = g.offset || 0,
                            k = d.$eval(i) || {},
                            l = {},
                            m = b.startSymbol(),
                            n = b.endSymbol(),
                            o = /^when(Minus)?(.+)$/;
                        f(g, function(a, b) {
                            o.test(b) && (k[oc(b.replace("when", "").replace("Minus", "-"))] = e.attr(g.$attr[b]))
                        }), f(k, function(a, d) {
                            l[d] = b(a.replace(c, m + h + "-" + j + n))
                        }), d.$watch(function() {
                            var b = parseFloat(d.$eval(h));
                            return isNaN(b) ? "" : (b in k || (b = a.pluralCat(b - j)), l[b](d, e, !0))
                        }, function(a) {
                            e.text(a)
                        })
                    }
                }
            }],
            qe = ["$parse", "$animate", function(a, c) {
                function g(a) {
                    return a.clone[0]
                }

                function h(a) {
                    return a.clone[a.clone.length - 1]
                }
                var i = "$$NG_REMOVED",
                    j = d("ngRepeat");
                return {
                    transclude: "element",
                    priority: 1e3,
                    terminal: !0,
                    $$tlb: !0,
                    link: function(d, k, l, m, n) {
                        var o, p, q, r, s, t, u, v, w, x = l.ngRepeat,
                            y = x.match(/^\s*([\s\S]+?)\s+in\s+([\s\S]+?)(?:\s+track\s+by\s+([\s\S]+?))?\s*$/),
                            z = {
                                $id: Ea
                            };
                        if (!y) throw j("iexp", "Expected expression in form of '_item_ in _collection_[ track by _id_]' but got '{0}'.", x);
                        if (t = y[1], u = y[2], o = y[3], o ? (p = a(o), q = function(a, b, c) {
                                return w && (z[w] = a), z[v] = b, z.$index = c, p(d, z)
                            }) : (r = function(a, b) {
                                return Ea(b)
                            }, s = function(a) {
                                return a
                            }), y = t.match(/^(?:([\$\w]+)|\(([\$\w]+)\s*,\s*([\$\w]+)\))$/), !y) throw j("iidexp", "'_item_' in '_item_ in _collection_' should be an identifier or '(_key_, _value_)' expression, but got '{0}'.", t);
                        v = y[3] || y[1], w = y[2];
                        var A = {};
                        d.$watchCollection(u, function(a) {
                            var l, m, o, p, t, u, y, z, B, C, D, E, F = k[0],
                                G = {},
                                H = [];
                            if (e(a)) C = a, B = q || r;
                            else {
                                B = q || s, C = [];
                                for (u in a) a.hasOwnProperty(u) && "$" != u.charAt(0) && C.push(u);
                                C.sort()
                            }
                            for (p = C.length, m = H.length = C.length, l = 0; m > l; l++)
                                if (u = a === C ? l : C[l], y = a[u], z = B(u, y, l), da(z, "`track by` id"), A.hasOwnProperty(z)) D = A[z], delete A[z], G[z] = D, H[l] = D;
                                else {
                                    if (G.hasOwnProperty(z)) throw f(H, function(a) {
                                        a && a.scope && (A[a.id] = a)
                                    }), j("dupes", "Duplicates in a repeater are not allowed. Use 'track by' expression to specify unique keys. Repeater: {0}, Duplicate key: {1}, Duplicate value: {2}", x, z, Q(y));
                                    H[l] = {
                                        id: z
                                    }, G[z] = !1
                                }
                            for (u in A) A.hasOwnProperty(u) && (D = A[u], E = fa(D.clone), c.leave(E), f(E, function(a) {
                                a[i] = !0
                            }), D.scope.$destroy());
                            for (l = 0, m = C.length; m > l; l++) {
                                if (u = a === C ? l : C[l], y = a[u], D = H[l], H[l - 1] && (F = h(H[l - 1])), D.scope) {
                                    t = D.scope, o = F;
                                    do o = o.nextSibling; while (o && o[i]);
                                    g(D) != o && c.move(fa(D.clone), null, uc(F)), F = h(D)
                                } else t = d.$new();
                                t[v] = y, w && (t[w] = u), t.$index = l, t.$first = 0 === l, t.$last = l === p - 1, t.$middle = !(t.$first || t.$last), t.$odd = !(t.$even = 0 === (1 & l)), D.scope || n(t, function(a) {
                                    a[a.length++] = b.createComment(" end ngRepeat: " + x + " "), c.enter(a, null, uc(F)), F = a, D.scope = t, D.clone = a, G[D.id] = D
                                })
                            }
                            A = G
                        })
                    }
                }
            }],
            re = ["$animate", function(a) {
                return function(b, c, d) {
                    b.$watch(d.ngShow, function(b) {
                        a[S(b) ? "removeClass" : "addClass"](c, "ng-hide")
                    })
                }
            }],
            se = ["$animate", function(a) {
```
