## Supplemental Network for "Granger-Causal Testing for Irregularly Sampled Time Series with Application to Nitrogen Signaling in Arabidopsis": d3d07913.main.js

```
/*global _, angular */

angular.module('cyViewerApp')
    .controller('MainCtrl', function($scope, $http, $location, $routeParams, $window, Network, VisualStyles) {

        'use strict';

        var FILE_LIST_NAME = 'filelist.json';

        // Name of network tag in the DOM
        var NETWORK_SECTION_ID = '#network';

        // Default Visual Style name
        var DEFAULT_VISUAL_STYLE_NAME = 'default';

        var visualStyleFile;
        var networkData;

        $scope.LAYOUTS = [
            'preset', 'cola', 'random', 'grid', 'circle', 'concentric', 'breadthfirst', 'cose'
        ];


        // Application global objects
        $scope.networks = networks;
        $scope.currentVS = null;
        $scope.visualStyles = styles;
        $scope.visualStyleNames = Object.styles(styles);
        $scope.networkNames = Object.styles(networks);
        $scope.currentNetworkData = Network.get(
                {filename: $scope.networks[defaultNetworkName]},
                function () {
                    angular.element(NETWORK_SECTION_ID).cytoscape(options);
                    $scope.currentNetworkData = networkData;
                    $scope.currentNetwork = defaultNetworkName;
                });;


        // Show / Hide Table browser
        $scope.browserState = {
            show: false
        };

        // Show / Hide style selector UI
        $scope.overlayState = {
            show: true
        };

        // Show / Hide toolbar
        $scope.toolbarState = {
            show: true
        };

        // Background color
        $scope.bg = {
            color: '#FAFAFA'
        };

        $scope.columnNames = [];
        $scope.edgeColumnNames = [];
        $scope.networkColumnNames = [];

        // Basic settings for the Cytoscape window
        var options = {
            showOverlay: false,
            minZoom: 0.01,
            maxZoom: 200,
            boxSelectionEnabled: true,

            layout: {
                name: 'preset'
            },

            ready: function() {
                $scope.cy = this;
                $scope.cy.load(networkData.elements);

                VisualStyles.query(
                    {filename: visualStyleFile}, function (vs) {
                        init(vs);
                        dropSupport();
                        setEventListeners();
                        $scope.currentVS = DEFAULT_VISUAL_STYLE_NAME;
                        $scope.currentLayout = 'preset';
                        $scope.cy.style().fromJson($scope.visualStyles[DEFAULT_VISUAL_STYLE_NAME].style).update();
                        angular.element('.loading').remove();
                    });
            }
        };


        function dropSupport() {
            var dropZone = angular.element(NETWORK_SECTION_ID);
            dropZone.on('dragenter', function(e) {
                e.stopPropagation();
                e.preventDefault();
            });

            dropZone.on('dragover', function(e) {
                e.stopPropagation();
                e.preventDefault();
            });
            dropZone.on('drop', function(e) {
                e.preventDefault();
                var files = e.originalEvent.dataTransfer.files;
                var networkFile = files[0];
                var reader = new FileReader();
                reader.onload = function(evt) {
                    var network = JSON.parse(evt.target.result);
                    var networkName = 'Unknown';
                    // Check data section is available or not.
                    networkData = network.data;
                    if (networkData !== undefined) {
                        if (networkData.name !== undefined) {
                            networkName = networkData.name;
                            $scope.currentNetworkData = networkData;
                        }
                    }

                    while (_.contains($scope.networkNames, networkName)) {
                        networkName = networkName + '*';
                    }

                    $scope.$apply(function() {
                        $scope.networks[networkName] = network;
                        $scope.networkNames.push(networkName);
                        $scope.currentNetwork = networkName;
                    });
                    $scope.cy.load(network.elements);
                    reset();
                };
                reader.readAsText(networkFile);
            });
        }

        function init(vs) {
            $scope.nodes = networkData.elements.nodes;
            $scope.edges = networkData.elements.edges;
            initVisualStyleCombobox(vs);

            // Set network name
            var networkName = networkData.data.name;
            if (!$scope.networks[networkName]) {
                $scope.networks[networkName] = networkData;
                $scope.networkNames.push(networkName);
                $scope.currentNetwork = networkData.data.name;
            }
            // Get column names
            setColumnNames();

            if ($routeParams.bgcolor) {
                $scope.bg.color = $routeParams.bgcolor;
            }
        }

        function setColumnNames() {
            $scope.columnNames = [];
            $scope.edgeColumnNames = [];
            $scope.networkColumnNames = [];

            var oneNode = $scope.nodes[0];
            for (var colName in oneNode.data) {
                $scope.columnNames.push(colName);
            }
            var oneEdge = $scope.edges[0];
            for (var edgeColName in oneEdge.data) {
                $scope.edgeColumnNames.push(edgeColName);
            }
            for (var netColName in networkData.data) {
                $scope.networkColumnNames.push(netColName);
            }
        }

        function reset() {
            $scope.selectedNodes = {};
            $scope.selectedEdges = {};
        }

        /*
         Event listener setup for Cytoscape.js
         */
        function setEventListeners() {
            $scope.selectedNodes = {};
            $scope.selectedEdges = {};

            var updateFlag = false;

            // Node selection
            $scope.cy.on('select', 'node', function(event) {
                var id = event.cyTarget.id();
                $scope.selectedNodes[id] = event.cyTarget;
                updateFlag = true;
            });

            $scope.cy.on('select', 'edge', function(event) {
                var id = event.cyTarget.id();
                $scope.selectedEdges[id] = event.cyTarget;
                updateFlag = true;
            });

            // Reset selection
            $scope.cy.on('unselect', 'node', function(event) {
                var id = event.cyTarget.id();
                delete $scope.selectedNodes[id];
                updateFlag = true;
            });
            $scope.cy.on('unselect', 'edge', function(event) {
                var id = event.cyTarget.id();
                delete $scope.selectedEdges[id];
                updateFlag = true;
            });

            setInterval(function() {
                if (updateFlag && $scope.browserState.show) {
                    $scope.$apply();
                    updateFlag = false;
                }
            }, 300);

        }

        function initVisualStyleCombobox(vs) {
            _.each(vs, function(visualStyle) {
                $scope.visualStyles[visualStyle.title] = visualStyle;
                $scope.visualStyleNames.push(visualStyle.title);
            });

            $scope.currentVS = DEFAULT_VISUAL_STYLE_NAME;
        }


        $scope.toggleTableBrowser = function() {
            $scope.browserState.show = !$scope.browserState.show;
        };

        $scope.toggleOverlay = function() {
            $scope.overlayState.show = !$scope.overlayState.show;
        };

        $scope.toggleToolbar = function() {
            $scope.toolbarState.show = !$scope.toolbarState.show;
        };

        $scope.fit = function() {
            $scope.cy.fit();
        };


        // Apply Visual Style
        $scope.switchVS = function() {
            var vsName = $scope.currentVS.trim();
            var vs = $scope.visualStyles[vsName].style;
            // Apply Visual Style
            $scope.cy.style().fromJson(vs).update();
        };


        $scope.switchNetwork = function() {
            var networkFile = $scope.networks[$scope.currentNetwork];

            networkData = Network.get(
                {filename: networkFile},
                function (network) {
                    $scope.cy.load(network.elements);
                    $scope.currentNetworkData = networkData;
                    reset();
                    $scope.nodes = network.elements.nodes;
                    $scope.edges = network.elements.edges;
                    setColumnNames();
                });


        };

        $scope.switchLayout = function() {
            var layoutOptions = {
                name: $scope.currentLayout
            };
            $scope.cy.layout(layoutOptions);
        };

        ///////////////////// Start the loading process ////////////////


        /*$http.get(FILE_LIST_NAME).success(function(fileList) {
            visualStyleFile = styles;

            var defaultNetworkName = null;
            $scope.networks = networks;
            _.each(_.keys(networks), function(key) {
                $scope.networkNames.push(key)
            }

            _.each(_.keys(fileList), function(key) {
                if(key !== 'style') {
                    if(defaultNetworkName === null) {
                        defaultNetworkName = key;
                    }
                    $scope.networks[key] = fileList[key];
                    $scope.networkNames.push(key);
                }
            });
            
            networkData = Network.get(
                {filename: $scope.networks[defaultNetworkName]},
                function () {
                    angular.element(NETWORK_SECTION_ID).cytoscape(options);
                    $scope.currentNetworkData = networkData;
                    $scope.currentNetwork = defaultNetworkName;
                });
        });
    });*/
```
