## Supplemental Network for "Granger-Causal Testing for Irregularly Sampled Time Series with Application to Nitrogen Signaling in Arabidopsis": custom.js

```
$( document ).ready(function(){
  // Custom Cytoscape.JS code goes here.
  
  // Example: add linkouts to nodes that opens the "href" node attribute on click
  // cy.on('tap', 'node', function(){
  //   try { // your browser may block popups
  //     window.open( this.data('href') );
  //   } catch(e){ // fall back on url change
  //     window.location.href = this.data('href');
  //   }
  // });
  
  // For more options, check out http://js.cytoscape.org/
  
});
```
