## Supplemental Network for "Granger-Causal Testing for Irregularly Sampled Time Series with Application to Nitrogen Signaling in Arabidopsis": 22f5c459.network.js

```
/**
 * Created by kono on 2014/01/24.
 */

'use strict';

/**
 * Services that persists and retrieves TODOs from localStorage
 */
angular.module('cyViewerApp')
    .factory('Network', ['$resource', function ($resource) {
        return $resource('data/:filename', {filename: '@filename'});
    }]
);
```
