## Supplemental Network for "Granger-Causal Testing for Irregularly Sampled Time Series with Application to Nitrogen Signaling in Arabidopsis": ac1aeeaa.visual-styles.js

```
/**
 * Created by kono on 2014/01/24.
 */
/*
'use strict';

angular.module('cyViewerApp')
    .factory('VisualStyles', ['$resource', function ($resource) {
        return $resource('data/:filename', {filename: '@filename'});
    }]
);*/
```
