## Supplemental Network for "Granger-Causal Testing for Irregularly Sampled Time Series with Application to Nitrogen Signaling in Arabidopsis": main.html

### Visual Style:

### Layout:

|  |  |  |  |
| --- | --- | --- | --- |
| SUID {{columnName}}|  |  | | --- | --- | | {{ node.id() }} {{node.data(colName)}} |

|  |  |  |  |
| --- | --- | --- | --- |
| SUID {{columnName}}|  |  | | --- | --- | | {{ edge.id() }} {{edge.data(colName)}} |
