## Supplemental Methods for "Granger-Causal Testing for Irregularly Sampled Time Series with Application to Nitrogen Signaling in Arabidopsis"

#### 1 Model Conditions

Knowing that the parameters of interest $\boldsymbol{\theta}$ are included in the terms $\tilde{\mu}_i$ and $\boldsymbol{\Sigma}_i$, a series of conditions and assumptions need to be considered to estimate these parameters in a statistically sound manner. Aside from the assumption of normality mentioned above, the other assumptions are as follows:

**(A1)** The functions $f_j(\cdot)$, $h_j(\cdot)$ and $g_j(\cdot)$ have known structures for $j = 1, 2$.

**(A2)** The mapping $\boldsymbol{\theta} \to F_{\boldsymbol{\theta}}$ is injective.

**(A3)** The process $(t_i, \boldsymbol{Z}_i)$ is such that:

(i) $\mathbb{E}[\boldsymbol{Z}_{t_i}] = 0, \forall\, i \in \mathbb{N}^+$.

(ii) $cov(\boldsymbol{Z}_{t_i}, \boldsymbol{Z}_{t_k}) = \boldsymbol{\Sigma}(|t_i - t_k|)$.

where $\boldsymbol{\Sigma}(|t_i - t_k|)$ is a nonsingular covariance matrix for all $i, j \in \mathbb{N}^+$.

**(A4)** The functions $f_j(\cdot)$ and $h_j(\cdot)$, $j = 1, 2$, jointly denoted as $m_j(\cdot)$, are twice continuously differentiable and such that:

(i) $m_j(0) = 1$ and $\lim\limits_{\delta \to \infty} m_j(\delta) = 0$.

(ii) $0 \leq |m_j(\delta)| \leq 1, \forall\, \delta \in \mathbb{R}^+$.

**(A5)** The functions $g_j(\cdot)$, $j = 1, 2$, are twice continuously differentiable and such that:

(i) $g_j(0) = 0$ and $\lim\limits_{\delta \to \infty} g_j(\delta) = \sigma_j^2$, where $0 < \sigma_j^2 < \infty$.

(ii) $g_j(\delta) < g_j(\delta + h), \forall\ \delta, h \in \mathbb{R}^+$.

The above assumptions, along with other standard regularity conditions, are necessary in order to correctly estimate the model parameters and to test whether these parameters are significant.

### 2 Supplemental Methods

#### 2.1 Plant growth conditions and sampling

The time-evolved transcriptome of Arabidopsis roots and shoots was obtained as described in detail by Varala *et al.* (2018). Briefly, *Arabidopsis thaliana* (Col-0) was grown hydroponically on 1 mM $KNO_3^-$ for two weeks then transiently treated with nitrogen (N) (20 mM $KNO_3^-$ plus 20 mM $NH_4NO_3$) or control (20 mM KCl) for two hours. Samples were harvested at times 0, 5, 10, 15, 20, 30, 45, 60, 90, and 120 minutes, in which three replicates of roots and shoots were separated at harvest and immediately frozen in liquid nitrogen (see Fig. 1).

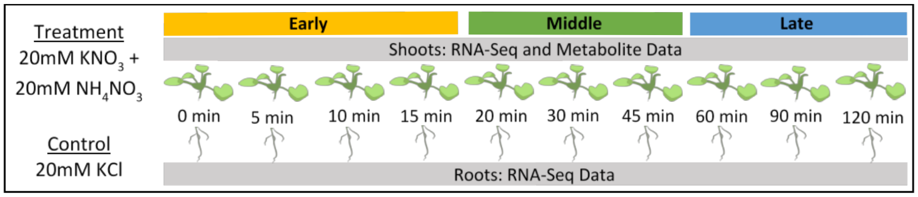


Figure 1: Tissue sampling scheme.

#### 2.2 Transcriptome analysis

As described in Varala *et al.* (2018), total RNA was extracted from approximately 100 mg of tissue using the Qiagen RNeasy Kit. RNA was then processed for paired-end Illumina Sequencing using standard protocols (Zhong *et al.*, 2011). The upper quartile normalization method from the `EDASeq` package in `R` was used to normalize the gene expression counts for both shoots and roots. The set of Differentially Expressed Genes (DEGs) (in response to N-treatment vs control over time) was derived using the spline fitting model in the `limma` package in `R`. The shoot gene set was determined at an FDR adjusted $p$-value $< 10^{-5}$, resulting in 2173 shoot DEGs, while the root gene set was determined at an FDR adjusted $p$-value $< 10^{-4}$, resulting in 568 root DEGs (see Varala *et al.*, 2018).

#### 2.3 Data pre-processing

The replicates were first combined by taking the average gene expression for each gene at each time point. This gave one single time series expression for each gene. Root genes and shoot genes were clustered separately using Multiple Experiment Viewer (MEV) (Saeed *et al.*, 2003). The gene time series were imported and normalized using the "Normalize Genes/Rows" function in MEV. This transforms the gene expression values using the mean and standard deviation of each time series. The genes were then clustered using the QT clustering algorithm, setting the maximum threshold to 0.25, and the minimum cluster population to 5. The list of genes in each cluster was then exported. In each cluster, the average gene expression at each time point was subtracted from the gene expression value at that time point for each time series. This method resulted in a de-trending of the root and shoot time series (thereby removing the mean $\mu_i^{(x)}$).

#### 2.4 Bioinformatic validation of the proposed method

In this section we provide an overview of the validation procedures followed in support of the results discussed in Section 3 in the main manuscript. Indeed, the proposed method merely suggests that a transcript level in one tissue is a result of a transcript in another tissue but the predicted relationships are not necessarily direct. Functional validation of predicted relationships focused on those that are likely caused by a direct, or potentially physical, interaction, such as protein:protein; protein:DNA; protein:metabolite, etc. A bioinformatics pipeline was developed to provide support for predicted relationships using network analysis, gene ontology, and text-mining to narrow down a manageable list of candidate genes for experimental testing.

##### 2.4.1 Gene Ontology (GO) Term Analysis

This analysis was performed using the GO enrichment analysis tool from the Gene Ontology Consortium (Botstein *et al.*, 2000; Consortium, 2019; Mi *et al.*, 2019). This tool returns a $p$-value from a Fisher's exact test in which the null hypothesis is that the biological function of the genes are distributed evenly throughout the subset of genes as compared to the the whole genome. A significant $p$-value indicates that the corresponding GO term appears more frequently than expected in the gene list compared to the the overall genome. In a GO analysis there are many class-subclass relationships, i.e. the GO term "nitrogen fixation" is a subclass of the "nitrogen cycle metabolic process". A Benjamini-Hochberg False Discovery Rate (FDR) correction is used to correct for the multiple testing and a cut-off of 0.05 is suggested by default for significant results (Mi *et al.*, 2019). GO terms were filtered using a FDR cut-off of 0.05 except in cases where there were too few GO terms (cutoff = 0.1) or no GO terms at the cutoff (no cutoff value).

##### 2.4.2 Network Analysis

Directed networks were generated where genes are represented as nodes, and the directional dependence, as determined by the model, is represented as edges between nodes. Biological networks have been shown to exhibit scale-free behavior such as the distribution in the network following a power-law (Albert, 2005). To provide some support towards the hypothesis that the proposed model-based network respects this feature, the predicted network was compared to random networks to determine how well it followed the power law for scale-free biological networks. $10^3$ random networks with the same number of nodes and edges were generated in `R` using the `sample_gnm` function as part of the `iGraph` package. For each generated random network, the $R^2$ was calculated for the degree, in-degree and out-degree.

###### 2.4.3 Promoter Analysis

The 2KB upstream region was obtained using Elefinder (Hudson, 2005). These regions were then used in Elefinder to determine over-represented transcription factor binding motifs. The results returned an E-value which is the likelihood of the result being returned by chance based on a binomial distribution. To search the 2KB upstream region for the significant occurrence, the FIMO tool from the MEME Suite (Bailey and Machanick, 2012) was used. Transcription factor binding motifs were first retrieved from the Plant Cistrome and EpiCistrome database (O'Malley *et al.*, 2016). Using FIMO, promoter regions obtained from Elefinder were then searched for the specific motif using the default settings.

###### 2.4.4 Nitrogen Response

For purposes of validation, particular attention was given to those genes previously implicated in the nitrogen response such as peptides (Araya *et al.*, 2014) and those involved in cytokinin biosynthesis Takei *et al.* (2004), carbon/nitrogen balance (Palenchar *et al.*, 2004), primary nitrogen metabolism and nitrogen transport (Perchlik and Tegeder, 2017; Krapp *et al.*, 2014).

###### 2.4.5 Microarray Data Analysis

GEO was searched for datasets with mutants of candidate causal genes. GSE8975, a DVL4 overexpression experiment, was analysed using GEO2R using the default settings. The results were scanned to see if any target genes were differentially expressed between the wild type and mutant plants ($p$-value $< 0.05$).

###### 2.4.6 Mobile Causal Gene Testing

In order to understand how well the proposed method detects known mobile causal genes, we performed a bootstrap procedure in which we considered all possible expression pairs (among all tested root and shoot expressions) and, from these pairs, we randomly selected the same amount of Granger-Causal pairs detected by our method. Among these we then randomly selected the causal gene in each pair and, once the list of causal genes was completed in this manner, this was compared to the list of known mobile genes. The latter list was obtained from the PlaMoM (Plant Mobile Macromolecules) database

(Guan *et al.*, 2016) and is made up of genes that produce a mobile product that has been previously experimentally shown to move from either root to shoot, shoot to root, or in both directions. Following this approach, we then counted the number of causal genes in the randomly selected list that also appeared in the list of known mobile genes. This procedure was repeated $10^3$ times and this distribution of counts was compared to the number of causal genes detected by our approach. This showed that the number of known mobile causal genes detected by the proposed method is always significantly higher than a method that simply randomly samples the same number of causal genes thereby supporting the validity of the proposed analysis.

#### 3 Supplemental Tables

The supplemental tables are made available in a supplemental Excel file called "Supplemental_Tables.xlsx". Below is a table that collects the name of each sheet in the file and what it contains.

| Sheet Name | Description |
|---|---|
| Supplemental Table 1 | Table showing predicted 3078 root-shoot interactions |
| Supplemental Table 2 | GO Terms for causal root genes at early time points. |
| Supplemental Table 3 | GO Terms for causal root genes at middle time points. |
| Supplemental Table 4 | GO Terms for causal root genes at late time points. |
| Supplemental Table 5 | GO Terms for causal shoot genes at early time points. |
| Supplemental Table 6 | GO Terms for causal shoot genes at middle time points. |
| Supplemental Table 7 | GO Terms for causal shoot genes at late time points. |
| Supplemental Table 8 | GO Terms for target root genes at early time points. |
| Supplemental Table 9 | GO Terms for target root genes at middle time points. |
| Supplemental Table 10 | GO Terms for target root genes at late time points. |
| Supplemental Table 11 | GO Terms for target shoot genes at early time points. |
| Supplemental Table 12 | GO Terms for target shoot genes at middle time points. |
| Supplemental Table 13 | GO Terms for target shoot genes at late time points. |
| Supplemental Table 14 | Table showing top 10 hubbiest genes by out-degree, and their network node properties. |
| Supplemental Table 15 | Nitrogen signal responsive gene families and their members appearing in the predicted model interactions. |
| Supplemental Table 16 | List of interactions in which the causal gene is a known signaling peptide. |
| Supplemental Table 17 | List of RNA-binding proteins and their known mobility according to the PLAMOM database. |

Table 1: Summary of tables contained in supplemental Excel file.
